# Systematic assessment of long-read RNA-seq methods for transcript identification and quantification

**DOI:** 10.1101/2023.07.25.550582

**Authors:** Francisco J. Pardo-Palacios, Dingjie Wang, Fairlie Reese, Mark Diekhans, Sílvia Carbonell-Sala, Brian Williams, Jane E. Loveland, Maite De María, Matthew S. Adams, Gabriela Balderrama-Gutierrez, Amit K. Behera, Jose M. Gonzalez, Toby Hunt, Julien Lagarde, Cindy E. Liang, Haoran Li, Marcus Jerryd Meade, David A. Moraga Amador, Andrey D. Prjibelski, Inanc Birol, Hamed Bostan, Ashley M. Brooks, Muhammed Hasan Çelik, Ying Chen, Mei R.M. Du, Colette Felton, Jonathan Göke, Saber Hafezqorani, Ralf Herwig, Hideya Kawaji, Joseph Lee, Jian-Liang Li, Matthias Lienhard, Alla Mikheenko, Dennis Mulligan, Ka Ming Nip, Mihaela Pertea, Matthew E. Ritchie, Andre D. Sim, Alison D. Tang, Yuk Kei Wan, Changqing Wang, Brandon Y. Wong, Chen Yang, If Barnes, Andrew Berry, Salvador Capella, Namrita Dhillon, Jose M. Fernandez-Gonzalez, Luis Ferrández-Peral, Natàlia Garcia-Reyero, Stefan Goetz, Carles Hernández-Ferrer, Liudmyla Kondratova, Tianyuan Liu, Alessandra Martinez-Martin, Carlos Menor, Jorge Mestre-Tomás, Jonathan M. Mudge, Nedka G. Panayotova, Alejandro Paniagua, Dmitry Repchevsky, Eric Rouchka, Brandon Saint-John, Enrique Sapena, Leon Sheynkman, Melissa Laird Smith, Marie-Marthe Suner, Hazuki Takahashi, Ingrid Ashley Youngworth, Piero Carninci, Nancy D. Denslow, Roderic Guigó, Margaret E. Hunter, Hagen U. Tilgner, Barbara J. Wold, Christopher Vollmers, Adam Frankish, Kin Fai Au, Gloria M. Sheynkman, Ali Mortazavi, Ana Conesa, Angela N. Brooks

**Affiliations:** Institute for Integrative Systems Biology, Spanish National Research Council (CSIC), Paterna, Spain; Department of Biomedical Informatics, The Ohio State University, Columbus, USA; Department of Computational Medicine and Bioinformatics, University of Michigan, Ann Arbor, USA; Developmental and Cell Biology; Center for Complex Biological Systems, University of California, Irvine, Irvine, USA; UC Santa Cruz Genomics Institute, University of California, Santa Cruz, Santa Cruz, USA; Centre for Genomic Regulation (CRG), The Barcelona Institute of Science and Technology, Dr. Aiguader 88, Barcelona 08003, Catalonia, Spain; Division of Biology and Biological Engineering, California Institute of Technology, Pasadena, USA; European Molecular Biology Laboratory, European Bioinformatics Institute, Wellcome Genome Campus, Hinxton, Cambridge CB10 1SD, UK; Department of Physiological Sciences, College of Veterinary Medicine; Center for Environmental and Human Toxicology, University of Florida, Gainesville, USA; Molecular Cell and Developmental Biology; Department of Biomolecular Engineering, University of California, Santa Cruz, Santa Cruz, USA; Flomics Biotech, Dr Aiguader 88, Barcelona 08003, Spain; Department of Molecular Physiology and Biological Physics, University of Virginia, Charlottesville, USA; Interdisciplinary Center for Biotechnology Research, University of Florida, Gainesville, USA; Department of Computer Science, University of Helsinki, Helsinki, Finland; Center for Bioinformatics and Algorithmic Biotechnology, Institute of Translational Biomedicine, St. Petersburg State University, St. Petersburg, Russia; Canada’s Michael Smith Genome Sciences Centre, BC Cancer, Vancouver, Canada; Biostatistics and Computational Biology Branch, National Institute of Environmental Health Sciences, Durham, USA; Genome Institute of Singapore (GIS), Agency for Science, Technology and Research (A*STAR), Singapore, Singapore; Walter and Eliza Hall Institute of Medical Research, Parkville, Australia; Department of Statistics and Data Science, National University of Singapore, Singapore, Singapore; Department Computational Molecular Biology, Max-Planck-Institute for Molecular Genetics, Berlin, Germany; Research Center for Genome & Medical Sciences, Tokyo Metropolitan Institute of Medical Science, Tokyo, Japan; Department of Neuromuscular Diseases, UCL Queen Square Institute of Neurology, London, UK; Department of Biomedical Engineering; Center for Computational Biology, Johns Hopkins University, Baltimore, USA; Department of Medical Biology, The University of Melbourne, Parkville, Australia; Yong Loo Lin School of Medicine, National University of Singapore, Singapore, Singapore; Barcelona Supercomputing Cente, Barcelona, Spain; Environmental Laboratory, US Army Engineer Research & Development Center, Vicksburg, USA; Biobam Bioinformatics SL, Valencia, Spain; Genetics Institute, University of Florida, Gainesville, USA; Cardiff University, Cardiff, UK; Department of Biochemistry & Molecular Genetics, University of Louisville, Louisville, USA; European Bioinformatics Institute, Wellcome Genome Campus, Hinxton, Cambridge CB10 1SD, UK, UK; Center for Integrative Medical Sciences, Laboratory for Transcriptome Technology, RIKEN, Yokohama, Japan; Department of Genetics, Stanford University, Palo Alto, USA; Human Technopole, Milano, Italy; Center for Environmental and Human Toxicology, Department of Physiological Sciences,, University of Florida, Gainesville, USA; Universitat Pompeu Fabra (UPF), Barcelona, Catalonia, Spain; U.S. Geological Survey, Wetland and Aquatic Research Center, Gainesville, USA; Brain and Mind Research Institute and Center for Neurogenetics, Weill Cornell Medicine, New York City, USA; Center for Public Health Genomics; UVA Cancer Center, University of Virginia, Charlottesville, USA; Microbiology and Cell Science Department, Institute for Food and Agricultural Sciences, University of Florida, Gainesville, USA

**Author notes:** Correspondence. These authors jointly supervised this work. These authors contributed equally to this work (Francisco J. Pardo-Palacios, Dingjie Wang, Fairlie Reese, Mark Diekhans, Sílvia Carbonell-Sala, Brian Williams, Jane E. Loveland, Maite De María). These authors contributed equally to this work (Matthew S. Adams, Gabriela Balderrama-Gutierrez, Amit K. Behera, Jose M. Gonzalez, Toby Hunt, Julien Lagarde, Cindy E. Liang, Haoran Li, Marcus Jerryd Meade, David A. Moraga Amador, Andrey D. Prjibelski).

## Abstract

The Long-read RNA-Seq Genome Annotation Assessment Project (LRGASP) Consortium was formed to evaluate the effectiveness of long-read approaches for transcriptome analysis. The consortium generated over 427 million long-read sequences from cDNA and direct RNA datasets, encompassing human, mouse, and manatee species, using different protocols and sequencing platforms. These data were utilized by developers to address challenges in transcript isoform detection and quantification, as well as *de novo* transcript isoform identification. The study revealed that libraries with longer, more accurate sequences produce more accurate transcripts than those with increased read depth, whereas greater read depth improved quantification accuracy. In well-annotated genomes, tools based on reference sequences demonstrated the best performance. When aiming to detect rare and novel transcripts or when using reference-free approaches, incorporating additional orthogonal data and replicate samples are advised. This collaborative study offers a benchmark for current practices and provides direction for future method development in transcriptome analysis.

## Introduction

There is a growing trend of using long-read RNA-seq (lrRNA-seq) data for transcript identification and quantification, primarily with Oxford Nanopore Technologies (ONT) and Pacific Biosciences (PacBio) platforms^1–4^. Consequently, there is a need to evaluate these approaches for transcriptome analysis to compare the impact of different sequencing platforms, multiple sequencing library preparation methods, and computational analysis methods^5–8^.

A previous effort by the RNA-Seq Genome Annotation Assessment Project (RGASP) Consortium^9,10^ involved evaluating short-read Illumina RNA-seq for transcript identification. It revealed limitations in recalling full-length transcript products due to the complexity of eukaryotic transcriptomes. To evaluate long-read approaches for transcriptome analysis, we formed the Long-read RNA-Seq Genome Annotation Assessment Project (LRGASP) Consortium modeled after the previous GASP^11^, EGASP^12^, and RGASP^9,10^ efforts. For this project, we aimed for an open community effort to be as transparent and inclusive as possible in evaluating technologies and computational methods (**Fig. 1**).

**Fig. 1.**
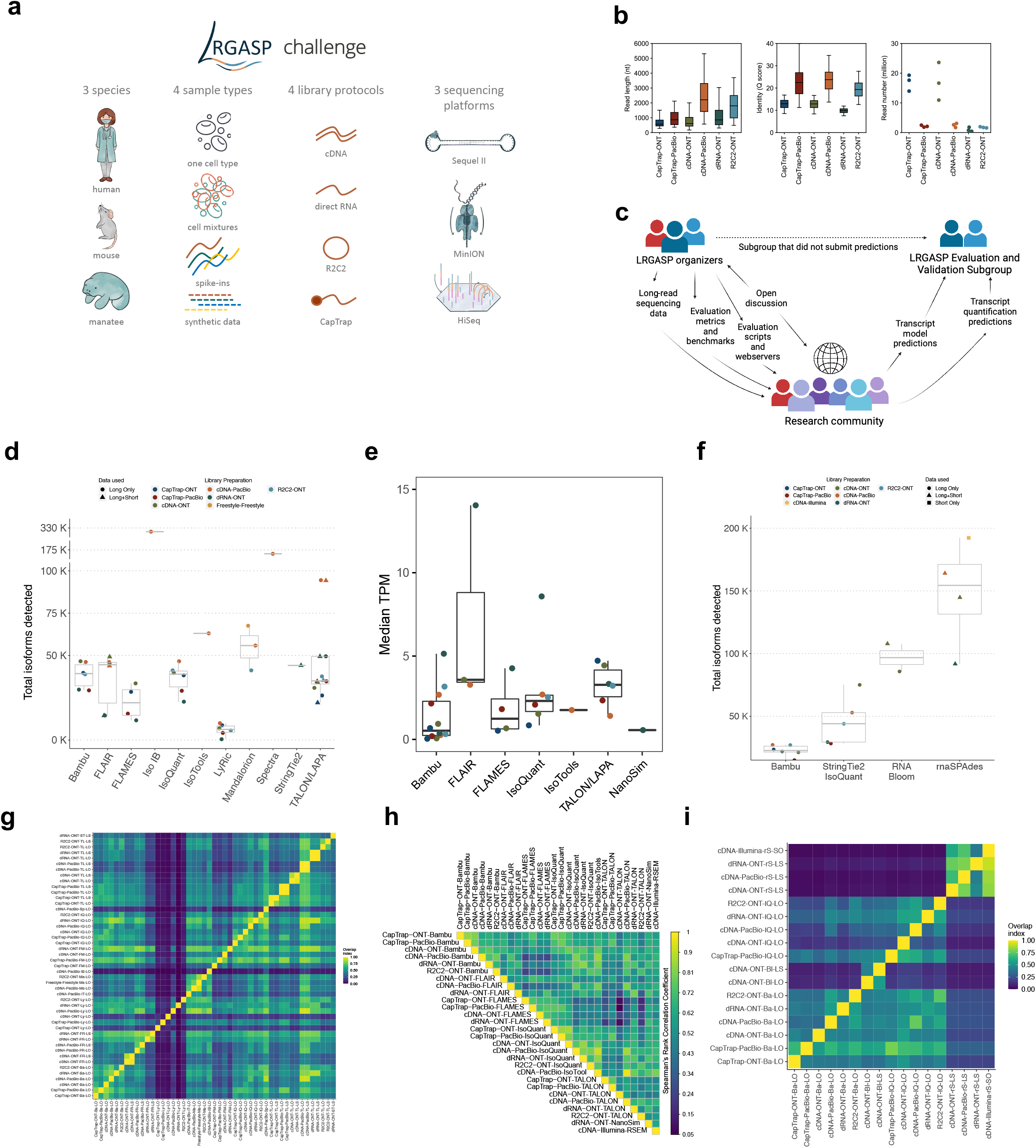
Overview of the Long-read RNA-seq Genome Annotation Assessment Project (LRGASP). **a,** Data produced for LRGASP consists of multiple species, multiple sample types, multiple library protocols, and multiple sequencing platforms for comparison. **b,** Distribution of read lengths, identify Q score, and sequencing depth (per biological replicate) for the WTC11 sample. **c**, LRGASP as an open research community effort for benchmarking and evaluating long-read RNA-seq approaches. **d,** Number of isoforms reported by each tool on different data types for the human WTC11 sample for Challenge 1. **e**, Median TPM value reported by each tool on different data types for the human WTC11 sample for Challenge 2. **f,** Number of isoforms reported by each tool on different data types for the mouse ES data for Challenge 3. **g,** Pairwise relative overlap of unique junction chains (UJCs) reported by each submission. The UJCs reported by a submission is used as a reference set for each row. The fraction of overlap of UJCs from the column submission is shown as a heatmap. For example, a submission that has a small, subset of many other UJCs from other submissions will have a high fraction shown in the rows, but low fraction by column for that submission. Data only shown for WTC11 submissions. **h,** Spearman correlation of TPM values between submissions to Challenge 2. **i,** Pairwise relative overlap of UJCs reported by each submission. The UJCs reported by a submission is used as a reference set for each row. The fraction of overlap of UJCs from the column submission is shown as a heatmap.

The LRGASP Consortium evaluated three fundamental aspects of transcriptome analysis. First, we assessed the reconstruction of full-length transcripts expressed in a given sample from a well-curated eukaryotic genome, such as from human or mouse. Second, we evaluated the quantification of the abundance of each transcript. Finally, we assessed the *de novo* reconstruction of full-length transcripts from samples without a high-quality genome, which would be beneficial for annotating genes in non-model organisms. These evaluations became the basis of the three challenges that comprised the LRGASP effort (**Box 1**).

#### BOX1: Overview of the LRGASP Challenges

##### Challenge 1: Transcript isoform detection with a high-quality genome

<u>Goal:</u> Identify which sequencing platform, library prep, and computational tool(s) combination gives the highest sensitivity and precision for transcript detection.

##### Challenge 2: Transcript isoform quantification

<u>Goal:</u> Identify which sequencing platform, library prep, and computational tool(s) combination gives the most accurate expression estimates.

##### Challenge 3: *De novo* transcript isoform identification

<u>Goal:</u> Identify which sequencing platform, library prep, and computational tool(s) combination gives the highest sensitivity and precision for transcript detection without a high-quality annotated genome.

Our benchmarking effort demonstrated the potential of long-read sequencing technologies in obtaining full-length transcripts and discovering new transcripts, even in well-annotated genomes. However, the agreement among bioinformatics tools was moderate, reflecting different goals in data analysis. Accurately detecting novel transcripts was more challenging than identifying transcript models already present in reference annotations. This indicates the need to improve *de novo* annotation of genomes using long-read data alone. Furthermore, transcript quantification accuracy varies significantly among bioinformatics tools depending on data scenarios, with long-read-based tools typically having lower quantitative accuracy than short-read-based tools due to low throughput and higher error rates. Improvements in long-read-based tools and increased throughput are expected to enhance their accuracy further. Evaluation analysis also highlights challenges in quantifying complex and lowly expressed transcripts. We experimentally validated many lowly expressed, single-sample transcripts, prompting further discussions on using long-read data for creating reference transcriptomes.

## Results

### LRGASP Data and study design

The LRGASP Consortium Organizers produced long-read and short-read RNA-seq data from aliquots of the same RNA samples using a variety of library protocols and sequencing platforms (**Fig. 1a**, **Table 1, Extended Data Table 1**). The Challenges 1 and 2 samples consist of human and mouse ENCODE biosamples with extensive chromatin-level functional data generated separately by the ENCODE Consortium. These include the human WTC11 iPSC cell line and a mouse 129/Casteneus ES cell line for Challenge 1 and a mix (H1-mix) of H1 human Embryonic Stem Cell (H1-hESC) and Definitive Endoderm derived from H1 (H1-DE) for Challenge 2. In addition, individual H1-hESC and H1-DE samples were sequenced on all platforms; however, those reads were only released after initial H1-mix predictions were submitted. All samples were grown as biological triplicates with the RNA extracted at one site, spiked with 5’-capped Spike-In RNA Variants (Lexogen SIRV-Set 4), and distributed to all production groups. A single pooled sample of manatee whole blood transcriptome was generated for Challenge 3. After sequencing, reads for human, mouse, and manatee samples were deposited at the ENCODE Data Coordination Center (DCC) for community access, including but not limited to usage for the challenges. We performed different cDNA preparation methods for each sample, including an early-access ONT cDNA kit (PCS110), ENCODE PacBio cDNA, and R2C2 for increased sequence accuracy ONT data, and CapTrap^13^ to enrich for 5’-capped RNAs (see Methods). CapTrap is derived from the CAGE technique^14^ and was adapted for lrRNA-seq. We also performed direct RNA sequencing (dRNA) with ONT.

**Table 1:** Overview of LRGASP sequencing data.

| Sample | # of Reps | cDNA-PacBio | cDNA-ONT | dRNA-ONT | R2C2-ONT | CapTrap-PacBio | CapTrap-ONT | cDNA-Illumina |
| --- | --- | --- | --- | --- | --- | --- | --- | --- |
| Mouse ES | 3 | Yes | Yes | Yes | Yes | Yes | Yes | Yes |
| Human WTC11 | 3 | Yes | Yes | Yes | Yes | Yes | Yes | Yes |
| Human H1-mix | 3 | Yes | Yes | Yes | Yes | Yes | Yes | Yes |
| Human H1-hESC | 3 | Yes | Yes | Yes | Yes | Yes | Yes | Yes |
| Human H1-DE | 3 | Yes | Yes | Yes | Yes | Yes | Yes | Yes |
| Manatee PBMCs | 1 | Yes | Yes | No | No | No | No | Yes |

The quality of the LRGASP datasets was extensively assessed (**Supplementary information: LRGASP Data QC**). cDNA-PacBio and R2C2-ONT datasets contained the longest read-length distributions, while sequence quality (assessed as % identity after genome mapping) was higher for CapTrap-PacBio, cDNA-PacBio, and R2C2-ONT than other experimental approaches. We obtained approximately ten times more reads from CapTrap-ONT and cDNA-ONT than with other methods (**Fig. 1b**).

The overall design of the LRGASP Challenge aimed for a fair and transparent process of evaluating long-read methods. The LRGASP effort was announced to the broader research community via social media and the GENCODE main website to recruit tool developers to submit transcript detection and quantification predictions based on the LRGASP data. Having tool developers provide results, rather than a third party, was likely to result in the best-performing parameters for that tool, making for a more fair comparison. In addition to data, the tool developers were provided detailed descriptions of the evaluation metrics and benchmarks, including scripts for challenge evaluations (**Fig. 1c**, detailed in **Supplementary information: Challenge submissions and timeline, Data and code availability**). Benchmarks included ground truth datasets known and unknown to participants before submission. Additionally, several community video conferences were held to answer questions about the data and evaluation and receive feedback on improving the study. Tool developers submitted challenge predictions, which were evaluated and validated by a subgroup of the LRGASP organizers that did not submit predictions (**Fig. 1c**). The separation of evaluators from prediction submitters was made to reduce any bias in the evaluation.

The evaluation of the challenges consisted of both bioinformatic and experimental approaches. SQANTI3^15^ was used to obtain transcript features and performance metrics (**Table 2)** computed based on SIRV-Set 4 spike-ins, simulated data, and a set of undisclosed, manually curated transcript models defined by GENCODE^16^. Human models were further compared to CAGE and Quant-seq data from the same samples. In addition, multiple evaluation metrics were designed for transcript quantification performance assessment both with and without known ground truth in different data scenarios. Experimental validation was performed on a select number of loci with either high agreement or disagreement between sequencing platforms or analysis pipelines, as well as several GENCODE annotated transcript models.

**Table 2:** Transcript Classifications and Definitions used by the LRGASP computational evaluation.

| Classification | Description |
| --- | --- |
| Full Splice Match (FSM) | Transcripts matching a reference transcript at all splice junctions |
| Incomplete Splice Match (ISM) | Transcripts matching consecutive, but not all, splice junctions of the reference transcripts |
| Novel in Catalog (NIC) | Transcripts containing new combinations of 1) already annotated splice junctions, 2) novel splice junctions formed from already annotated donors |
|  | and acceptors, or 3) unannotated intron retention |
| Novel Not in Catalog (NNC) | Transcripts using novel donors and/or acceptors |
| Reference Match (RM) | FSM transcript with 5' and 3' ends within 50 nts of the transcription start site (TSS)/transcription termination site (TTS) annotation |
| 3' poly(A) supported | Transcript with poly(A) signal sequence support or short-read 3' end sequencing (e.g. QuantSeq) support at the 3' end |
| 5' CAGE supported | Transcript with CAGE support at the 5' end |
| 3' reference supported | Transcript with 3' end within 50 nts from a reference transcript TTS |
| 5' reference supported | Transcript with 5' end within 50 nts from a reference transcript TSS |
| Supported Reference Transcript Model (SRTM) | FSM/ISM transcript with 5' end within 50 nts of the TSS or has CAGE support AND 3' end within 50 nts of the TTS or has poly(A) signal sequence support or short-read 3' end sequencing support |
| Supported Novel Transcript Model (SNTM) | NIC/NNC transcript with 5' end within 50 nts of the TSS or CAGE support AND 3' end within 50 nts of the TTS or has poly(A) signal sequence support or short-read 3' end sequencing support AND Illumina read support at novel junctions |
| % Long Read Coverage (%LRC) | Fraction of the transcript model sequence length mapped by one or more long reads |
| Read multiplicity | Number of assigned transcripts per read |
| Redundancy | # LR transcript models / reference model |
| Longest Junction Chain<br>ISM<br>NIC / NNC | # junctions in ISM / # junctions reference<br># reference junctions / # junctions in NIC/NNC |
| Intron retention (IR) level | Number of IR within the NIC category |
| Illumina Splice Junction (SJ) Support | % SJ in transcript model with Illumina support |
| Full Illumina Splice Junction Support | % transcripts in category with all SJ supported |
| % Novel Junctions | # of new junctions / total # junctions |
| % Non-canonical junctions | # of non-canonical junctions / total # junctions |
| % Non-canonical transcripts | % transcripts with at least one non-canonical junction |
| Intra-priming | Evidence of intra-priming (described in <sup>15</sup>) |
| RT-switching | Evidence of RT-switching (described in <sup>15</sup>) |

### LRGASP Challenge submissions

A total of 14 tools and labs participated in LRGASP, submitting predictions for each of the three challenges. (**Extended Data Table 2**) While submitters could choose the type of experimental procedure (a combination of library preparation and sequencing platform) they wished to participate in, predictions were required to be provided for all biological samples in the chosen experimental procedure to assess pipeline consistency. We received 141, 143, and 25 submissions for Challenges 1, 2, and 3, respectively.

We observed a large variability in the number and quantification of transcript models predicted by each submission, with differences of up to 10-fold in each challenge (**Figs. 1d-f**). Moreover, there was little overlap in the transcripts identified by any two pipelines in Challenges 1 and 3, and low pairwise correlations were detected for Challenge 2 quantification results (**Figs. 1g-i, Extended Data Table 3**). These results highlight the importance of the comprehensive benchmarking presented here.

### Challenge 1 Results and Evaluation: Transcript isoform detection with a high-quality genome

The evaluation of transcript model predictions in Challenge 1 used a variety of datasets, each of them addressing different performance aspects. When looking at predictions using datasets of real samples, we noticed that the fraction of the long reads that were used to build the transcript model (Percentage of Reads Used, PRU) greatly varied among and within methods, with tools such as LyRic^13^ only utilizing less than 20% of the available reads and others, e.g., Iso_IB^17^, FLAMES^18^, and IsoQuant^19^ showing PRU greater than 100%, indicating that the same read supported more than one transcript model (**Extended Data Fig. 1**), and revealing very different strategies in the reconstruction of the transcriptomes. We evaluated the overall capacity of the experimental methods and tools for transcript and gene detection based on SQANTI categories and various orthogonal datasets, as described in **Table 3**. Results were consistent for all three samples (**Extended Data Tables 4-6**). **Fig. 2a** shows data for WTC11, while **Extended Data Figs. 2,3** contain the results for H1-mix and mouse ES samples, respectively. We found a large variation in the number of known genes (399-23,647) and transcripts (524-329,131) detected by the LyRic-only (**Fig. 2a**), although there was a consistent relationship between both figures, with most pipelines reporting on average 3-4 transcripts per gene, except for Spectra^20^ and Iso_IB that reported a huge number of transcripts (∼170K and 330K, respectively). Due to pipeline variation, we did not find a clear relationship between the number of reads, the read length, and read quality with the number of detected transcripts (**Extended Data Figs. 4-6**). Roughly the number of detected genes and transcripts were determined to a great extent by the analysis tool (**Fig. 2a** and **Extended Data Figs. 7,8**). In general, pipelines detected more genes using cDNA library preparations and the PacBio sequencing platform. At the same time, there was not a clear best library preparation method for ONT in terms of the number of detected genes **(Extended Data Fig. 9**) or transcripts (**Extended Data Fig. 10**). The same trend was observed when comparing results within each analysis tool, (**Extended Data Figs. 11-14**).

**Figure 2:**
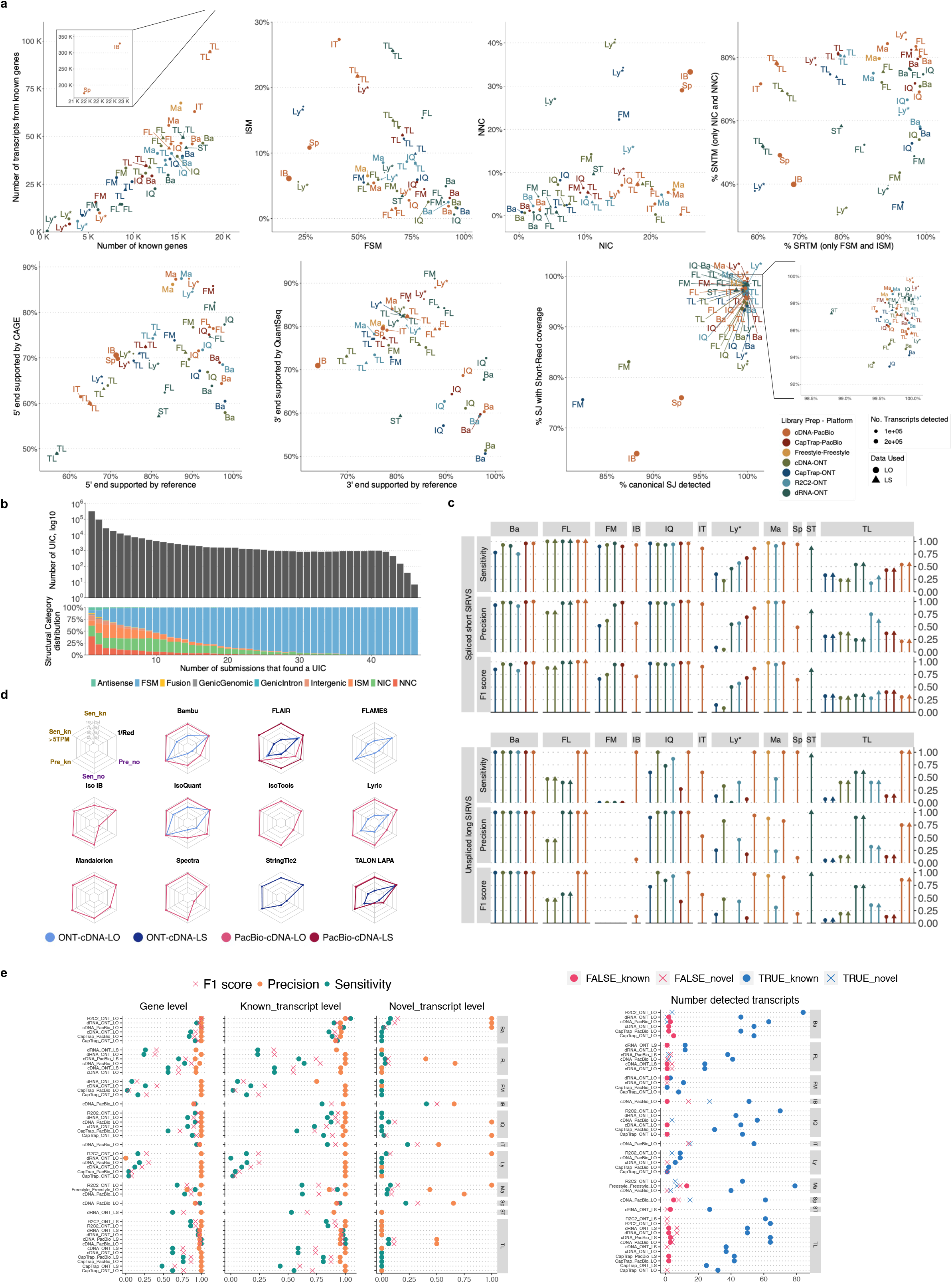
Overview of evaluation for Challenge 1: transcript identification with a reference annotation. **a)** Number of genes and transcripts per submission. Abundance of the main structural categories and support by external data. b) Agreement in transcript detection as a function the number of detecting pipelines. c) Performance based on for spliced-short and unspliced-long SIRVs. d) Performance based on simulated data. e) Performance for known and novel transcripts based on 50 manually-annotated genes by GENCODE. Ba: Bambu, FM: Flames, FR: FLAIR, IQ: IsoQuant, IT: IsoTools, IB: Iso_IB, Ly: LyRic, Ma: Mandalorion, TL: TALON-LAPA, Sp: Spectra, ST: StringTie2.

**Table 3:**
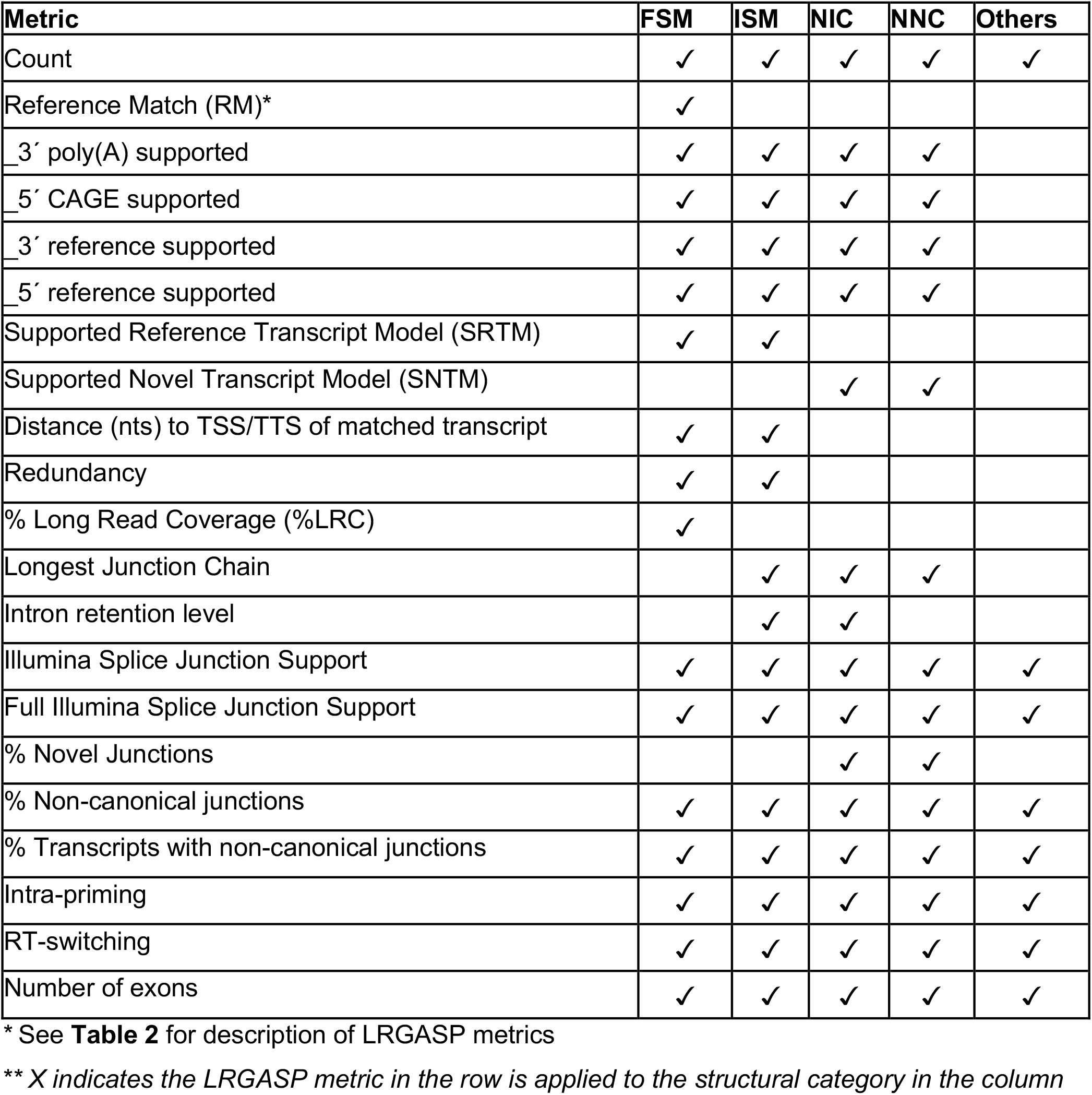
Metrics for evaluation against GENCODE annotation.

Pipelines also greatly varied in detecting transcripts annotated in GENCODE (Full-Splice Match, FSM), containing annotated splice junctions but unannotated transcript start or ends (Incomplete Splice Match, ISM), containing novel junctions between annotated GENCODE splice sites (Novel In Catalog, NIC), or containing novel splice sites with respect to GENCODE (Novel Not in Catalog, NNC). Bambu^21^, FLAIR^22^, FLAMES, and IsoQuant consistently detected a high percentage of FSM and a low proportion of ISM transcripts. In contrast, TALON^23,24^, IsoTools^25^, and LyRic detected a relatively high number of ISM (**Fig. 2a**). It was noted by the Lyric submission group that they did not use existing annotations to guide their analysis, which can explain their results. As for novel transcripts, Bambu was the tool that reported the lowest values for NNC and NIC. Also, IsoQuant and TALON had a low discovery of novel transcripts. FLAIR and Mandalorion^26^ pipelines typically returned around 20% NIC and low NNC percentages. LyRic and FLAMES were among the pipelines with the highest percentages of novel transcript detections, and so were Iso_IB and Spectra, which generally returned many isoforms, with only a small fraction of them being FSM. When these values were further analyzed stratified by library preparation and sequencing platform, a similar pattern was observed for the total number of transcripts: cDNA library preps, especially in combination with PacBio, more frequently had the highest number of FSM, NIC, and NNC while there was not a clear pattern for the detection of ISM (**Extended Data Figs. 15-20**). Within each analysis tool, SQANTI categories Antisense and Genic Genomic were more frequently detected using ONT sequencing data combined with cDNA or CapTrap. Intergenic transcripts were often found in cDNA library preparations (**Extended Data Figs. 21,22**).

We then evaluated the support of transcript models by reference annotation and orthogonal data derived from short-read sequencing (i.e., cDNA sequencing, CAGE, and Quant-seq). We observed that many pipelines had a high percentage of known transcripts with full support at TTS, TSS, and junctions (SRTM, see Methods), which were not mirrored by full support in novel transcript models (SNTM) (**Fig. 2a**). In general, tools analyzing cDNA-PacBio and cDNA-ONT data had high values of full support both for novel and known transcripts. A significant number of TALON pipelines had only moderate full support of known transcripts, possibly due to their high number of ISM. However, TALON showed consistent full support of novel transcripts in most cases. In contrast, methods such as LyRic, IsoQuant, FLAMES, and Bambu, which had high full support values for novel transcripts when analyzing cDNA-PacBio data, returned novel transcripts models with lower support when processing ONT library preparations. When looking at the specific evidence for 5’ and 3’ ends, we found that, in general, pipelines were more successful in reporting experimentally supported 3’ ends than 5’ ends. At the same time, transcript models matched the reference TTS and TSS annotations relatively well. There were, though, significant differences between pipelines. Tools such as Bambu and IsoQuant reported a high percentage of transcripts matching reference TTSs and TSSs. However, their support by matched CAGE and Quant-seq experimental data was relatively low. On the contrary, some of the LyRic and FLAMES submissions contained transcript models well supported by both types of experimental data, and Mandalorion had the highest CAGE support rates. TALON, Iso_IB, Spectra, and StringTie2^27^ had intermediate support at both transcript ends, considering the reference annotation or the orthogonal experimental data. This result suggested that a number of lrRNA-seq analysis pipelines that rely on reference annotations may use this information to adjust or complete transcript model predictions. To assess this hypothesis, we evaluated to which extent the actual long read data supported the predicted transcript sequences provided by each analysis pipeline by calculating the Long Read Coverage (LRC), i.e., the percentage of transcript model nucleotides mapped by at least one long read, based on our independent alignments. We found that FLAMES, Iso_IB, IsoTools, LyRic, and Mandalorion had nearly 100% of their transcript models fully supported (>98% coverage). While for FLAIR, Spectra, TALON, IsoQuant, Bambu, and StringTie2, these percentages dropped to approximately 90, 90, 85, 75, 60, and 45%, respectively (**Extended Data Fig. 23**). This suggests that this second set of tools employs a distinct strategy of the read alignments from the ones used for the assessment, or partially complete their transcript sequences using additional information such as the reference annotation or short-read data. Finally, we looked at the percentage of junctions with Illumina reads support and canonical splice sites. We found these values were generally very high for all pipelines except Spectra, Iso_IB, and FLAMES using cDNA-ONT and CapTrap-ONT data, with LyRic_PacBio showing the highest percentage of splice junctions supported by Illumina reads (**Fig. 2a**). The distribution of detected gene biotypes was, however, very similar for all methods, with the largest majority of detected genes being protein-coding (84.5%±6.5%), followed by lncRNAs (9.8%±4.1%) and pseudogenes (3.9%±2.2%) (**Extended Data Fig. 24**). Only Bambu reported an unusual number of non-coding genes when analyzing several library preparations of the H1-mix sample (**Extended Data Fig. 25**).

We then evaluated the agreement in the detection of known and novel Unique Intron Chains (UICs) across sequencing platforms, library preparations, and analysis pipelines (**Extended Data Tables 3-7**). When considering all 47 WTC11 submissions, detection by only one pipeline was the most frequent transcript class. The number of transcripts consistently detected by an increasing number of pipelines steadily decreased (**Fig. 2b**, **Extended Data Fig. 26**). Moreover, there was a clear relationship between the transcript’s SQANTI structural category and the number of pipelines it was detected by, with novel transcripts being more frequently identified by few pipelines and FSMs being nearly the only type of transcripts that were found by more than 40 pipelines. Overall, the overlap in detection between any two pipelines was higher for genes and junctions than for UICs, even when we only considered dominant UICs accounting for over 50% of the gene expression (**Extended Data Figs. 27-29**), highlighting the disparity in the identification of transcript models across methodologies. We then re-evaluated the agreement in UIC detection by looking at the overlap of each analysis method with the rest and discarding those tools with a large number of detections (Spectra and Iso_IB). Results were highly reproducible for the three samples evaluated in Challenge 1. Overall, a significant number of UICs were detected by all analysis pipelines when considering each library preparation/sequencing platform combination separately. However, important differences were observed for different SQANTI categories and experimental methods (**Extended Data Figs. 30-35**). FSMs were the most consistently detected transcript type across tools, in tens of thousands of different UICs in cDNA-PacBio, and R2C2-ONT datasets, while ISM, NIC, and NNC consistent detections were in the hundreds range for these experimental protocols. Other types of combinations (CapTrap-PacBio, CapTrap-ONT, and dRNA-ONT) returned agreement rates that were one order of magnitude lower: in the range of thousands for FSM and tens for ISM and NNC. Although detected by all individual tools, other structural categories, such as Fusion, Antisense, Genic Genomic, and Genic Intron, disappeared when imposing replicability across analysis methods. We also investigated the consistency in UIC detections for the same tool across different experimental datasets with similar conclusions: FSM transcripts were the most repeatedly detected throughout experimental protocols, while ISM, NIC, and NNC counts dropped quickly as more protocols were considered, with a few hundred models being present in over three datasets. One exception was Mandalorion, which reported consistent numbers of UICs regardless of the experimental data used, even in the ISM, NIC, and NNC categories. Multiple experimental protocols did not consistently detect UICs of other SQANTI categories (**Extended Data Fig. 36**). LyRic was the tool with the lowest number of consistently detected transcript models. Finally, we defined a set of “frequently detected transcripts” (FDT) as UICs present in at least three experimental datasets and detected by at least three analysis tools. This set contained around 45,000 UICs in each biological sample, representing ∼8% of the total number of UICs found by any method. Of these, the great majority (∼80%) were FSM, followed by NIC transcripts that accounted for ∼10% of the FDT (**Extended Data Fig. 37**). This implies that compared to all detections, FSM was enriched by 4-5 fold in the FDT set, while the other categories were significantly depleted. Finally, the set of FDT was more frequently found by IsoQuant, Mandalorion, Bambu, TALON-LAPA, and FLAMES than by other tools, even when correcting by the number of analysis pipelines submitted by each lab (**Extended Data Fig. 37**).

To understand which characteristics were driving differences in the detection of transcripts by the long read technologies, we compared expression level, transcript length and the number of exons of transcripts exclusively detected by each experimental method. We found that transcripts found only by pipelines processing PacBio reads were longer and had more exons and lower expression values than transcripts detected by Nanopore only or by both technologies (**Extended Data Fig. 38**). This pattern was also seen in the transcripts exclusively identified when tools analyzed cDNA data –and to a lesser extent R2C2-ONT, compared to those only detected by other library preparations methods (**Extended Data Fig. 39**). The combination cDNA-PacBio was the experimental procedure where their exclusively detected transcripts were the longest and had significantly lower expression (**Extended Data Fig. 40**). These results are in keeping with the read quality data and reveal that global capacity for transcript detection is associated with quality parameters of the sequencing reads. Interestingly, although these differences in transcript length were broadly recapitulated by analysis method, large transcripts (>10,000 nts in length) were exclusively reported by Bambu, IsoQuant, StringTie2, and TALON-LAPA (**Extended Data Fig. 41**).

The evaluation based on cell-line data lacks a ground truth that can be used to estimate the actual accuracy of the methods. We used a combination of spike-ins (SIRV), simulated data, and GENCODE manual annotation to estimate sensitivity, precision, and other performance metrics. Using the spliced SIRVs Lexogen dataset, most tools showed high sensitivity, except for TALON, while LyRic and Bambu sensitivity varied as a function of the library preparation method (**Fig. 2c, Extended Data Tables 4-6**). LyRic only had a sensitivity above 0.8 for the cDNA-PacBio sample, while Bambu showed lower sensitivity with R2C2-ONT and CapTrap-ONT. Precision was generally high for Bambu, IsoQuant, IsoTools, and Mandalorion methods and low for TALON, Iso_IB, and Spectra, with FLAMES, FLAIR, and LyRic showing variable results. F1 score figures were, in general, similar to the precision values, with IsoQuant, Mandalorion, FLAIR, Bambu, and IsoTools being the highest. However, It should be noted that conclusions on performance consistency are biased by the actual samples analyzed by the different tools. For example, the IsoTools team only analyzed the cDNA-PacBio data and performed well, comparable to FLAIR and Bambu. However, these methods also submitted predictions for ONT samples and yielded more variable results.

The SIRVs dataset also contained long, non-spliced transcripts that were evaluated separately. Interestingly, performance was very different with this subset (**Fig. 2c**, **Extended Data Tables 3-5)**. FLAMES did not report any due to the lack of support for single-exon transcripts. While Bambu had 100% sensitivity and precision in all library preparation protocols, possibly due to their support by reference annotation data (**Extended Data Fig. 42**). In general, sensitivity was lower across all analysis methods, except when using the cDNA-PacBio library preparation, which was 100% for most tools, including TALON and LyRic. Interestingly, when tools used CapTrap data, sensitivity was generally low, even in combination with PacBio sequencing, suggesting that the CapTrap library preparation was limited for capturing long molecules. Precision values were more variable and dependent on the tool and library preparation protocol. Since these SIRVs do not contain splice sites, low precision values indicate false variability at TTS and TSS. As results obtained with the long non-spliced dataset may be the combination of the ability of the analysis tool to process non-spliced data and accurately define TSS and TTS, but also of the capacity of the experimental protocol to capture long molecules, we looked specifically at the long read coverage for long SIRVs (**Extended Data Fig. 42**). We found that cDNA-PacBio was the combination where long reads more uniformly covered long SIRVs. However, coverage strongly dropped towards the 5’ end for the longest molecules. CapTrap library preparations -sequenced with PacBio and Nanopore-showed a strong coverage drop, frequently down to 0, at the middle of the SIRV sequence. R2C2 preparations lacked complete coverage for the extremely long SIRVs, while dRNA data showed more uniform coverage, albeit with striped patterning denoting sequencing errors.

SIRVs-based evaluation, while allowing assessment of experimental protocols, represents a reduced dataset that might be biased by the knowledge of the ground truth and cannot evaluate the capacity of tools to accurately discover new transcripts, which is one of the major claims of long-read sequencing platforms. We used simulated data to provide a more extensive assessment scenario, including novel transcripts in the synthetic dataset but not made available to the participants (**Extended Data Tables 7,8**). Using PacBio simulated data, we found that Mandalorion generally had good prediction and sensitivity for annotated and novel transcripts (**Fig. 2d** and **Extended Data Fig. 43**). IsoTools and LyRic also generally performed well except for the precision of novel transcripts. Bambu and FLAIR showed high sensitivity and precision when considering known transcripts but considerably lower figures when looking for novel isoforms. This was particularly noticeable for FLAIR, when not using short-read data, could not discover new transcripts accurately. Spectra and Iso_IB conversely had high sensitivity for novel isoforms at the cost of lower precision. For all tools, sensitivity increased on highly expressed transcripts, and redundancy values were close to 1, except for Iso IB and Spectra, which returned a higher number of redundant predictions. Simulation of Nanopore data resulted in datasets where most tools returned very low-performance figures, possibly due to lower coverage of the NanoSim^28^ reads of the simulated transcript models (**Extended Data Fig. 44**). Exceptions were Bambu and IsoQuant, which had good precision for ONT-known simulated transcripts, and StringTie at metrics other than those related to novel transcript discovery. Overall, results using simulated data indicated lower sensitivity and prediction for novel than known transcripts, with some tools being these differences considerable (**Fig. 2d**).

Synthetic data, allowing performance assessment on large datasets and novel transcripts, is limited by the properties of the simulation algorithms that are designed to reproduce sequencing errors but ignore other sources of artifacts such as library preparation and biological noise. To evaluate performance considering the complexity of real data, a set of 50 genes were selected for which the ground truth was set by exhaustive manual annotation by GENCODE annotators, and loci were unknown to submitters. Manually annotated loci were chosen for having mapped reads in all six library preparation/sequencing platform combinations and average to moderately high expression levels (**Extended Data Fig. 45**). GENCODE annotators evaluated the long read data for each experimental procedure independently and called transcript models in each case (See Supplementary Methods). Globally, 271 models were accepted as true transcripts in the WTC11 sample. cDNA-PacBio and cDNA-ONT were the experimental methods with the highest number of annotated transcripts (129 and 125, respectively), followed by R2C2-ONT and CapTrap-ONT (118 and 110, respectively). CapTrap-PacBio and dRNA-ONT had the lowest number (63 and 67, respectively), due to overall fewer aligned reads to the selected regions. Interestingly, the majority of transcripts in this subset were novel, with NNC being the most predominant SQANTI category. However, while FSM accounted only for 30% of these transcript models, they were the only category present within the transcripts detected in five or more experimental conditions. Most novel transcripts were found only in one dataset (**Extended Data Fig. 45**). Similar results were obtained on 50 loci annotated from the ES_mouse sample (**Extended Data Fig. 46**).

When pipelines were assessed based on this set of loci, we found that the analysis pipeline drove performance differences. All methods showed high precision at the gene level, but sensitivity was lower than with previous evaluating datasets. FLAMES and LyRic showed overall low sensitivity, as did FLAIR on dRNA data and TALON on CapTrap and cDNA-ONT datasets. Bambu, IsoTools, IsoQuant, and Spectra showed the highest sensitivity at the gene detection level (**Fig. 2e**), followed by TALON and Mandalorion but were more affected by the data type. A similar pattern of sensitivity and prediction was observed when considering transcripts already present in the reference annotation (**Fig. 2e**). ONT-known sensitivity dramatically dropped for the transcripts classified as novel by the GENCODE annotators, being close to 0 in most cases, and only IB had a sensitivity higher than 0.4. On the contrary, precision values greatly varied, ranging from 1 to 0 to non-computable even within the same tool, due to a generally low number of novel discoveries (<4) by most pipelines (**Fig. 2f**). Results were similar for the mouse ES annotated dataset (**Extended Data Figs. 47,48**).

Since our analysis of the manually annotated transcripts indicated that many new isoforms were found in only one dataset, we also calculated performance metrics considering only those transcripts detected in at least two experimental samples (114 transcripts) or with more than two reads (94 transcripts). These filters did not significantly affect the sensitivity of the methods and worsened the precision for novel transcript detection (**Extended Data Figs. 49,50**). When reevaluating performance based on library preparation method and sequencing platform, no clear best experimental method was evident; however, there was slightly higher precision for novel transcripts and higher sensitivity for known transcripts for tools working on PacBio, rather than Nanopore datasets (**Extended Data Figs. 51,52**).

In summary, the library preparation method, sequencing platform, and transcript analysis tool influenced many differences in the reported transcriptomes. Interestingly, the number of transcripts detected was not associated with the number of reads. Some tools are more reliant on annotation than others. For example, some tools correct their transcript models based on the annotation (Bambu, FLAMES, FLAIR, and IsoQuant), while other methods allow more novelty based on the actual data (Iso_IB, IsoTools, Mandalorion, Spades, TALON, LyRic).

We observe discrepancies among performance results depending on the benchmark. For example, LyRic performed poorly on SIRVS and GENCODE manual annotation but good on simulated data. In general, sensitivity for novel transcripts when using simulated data resulted in better performance than when using the GENCODE manual annotation, indicating challenges in benchmarking approaches. It could be argued that the GENCODE manual annotation is based on real data that presents a more realistic annotation challenge; however, the manual review resulted in a bias in locus selection that had relatively fewer reads to review with fewer replicates in different libraries.

### Challenge 2 Results and Evaluation: Transcript isoform quantification

To evaluate the performance of transcript quantification, both with and without a ground truth, we generated 84 real sequencing datasets (including SIRV-set 4) from four human cell lines (H1-hESC, H1-DE, H1-mix, and WTC11) and six simulation datasets for Nanopore (NanoSim), PacBio (IsoSeqSim^29^) and Illumina (RSEM^30^) reads (**Fig. 1a**). We received a total of 143 submitted datasets from seven quantification tools (IsoQuant, Bambu, TALON, FLAIR, FLAMES, NanoSim and IsoTools) on six combinations of protocols-platforms (cDNA-PacBio, cDNA-ONT, dRNA-ONT, CapTrap-PacBio, CapTrap-ONT, and R2C2-ONT). As a control, short-read datasets (cDNA-Illumina) were also quantified using the RSEM tool with GENCODE reference annotation. We designed nine metrics for performance assessment under different data scenarios (**Fig. 3a** and **Table 4**). We further provided a benchmarking web application (https://lrrna-seq-quantification.org/), allowing users to upload their transcript quantification results and produce interactive evaluation reports in HTML and PDF format.

**Fig. 3.**
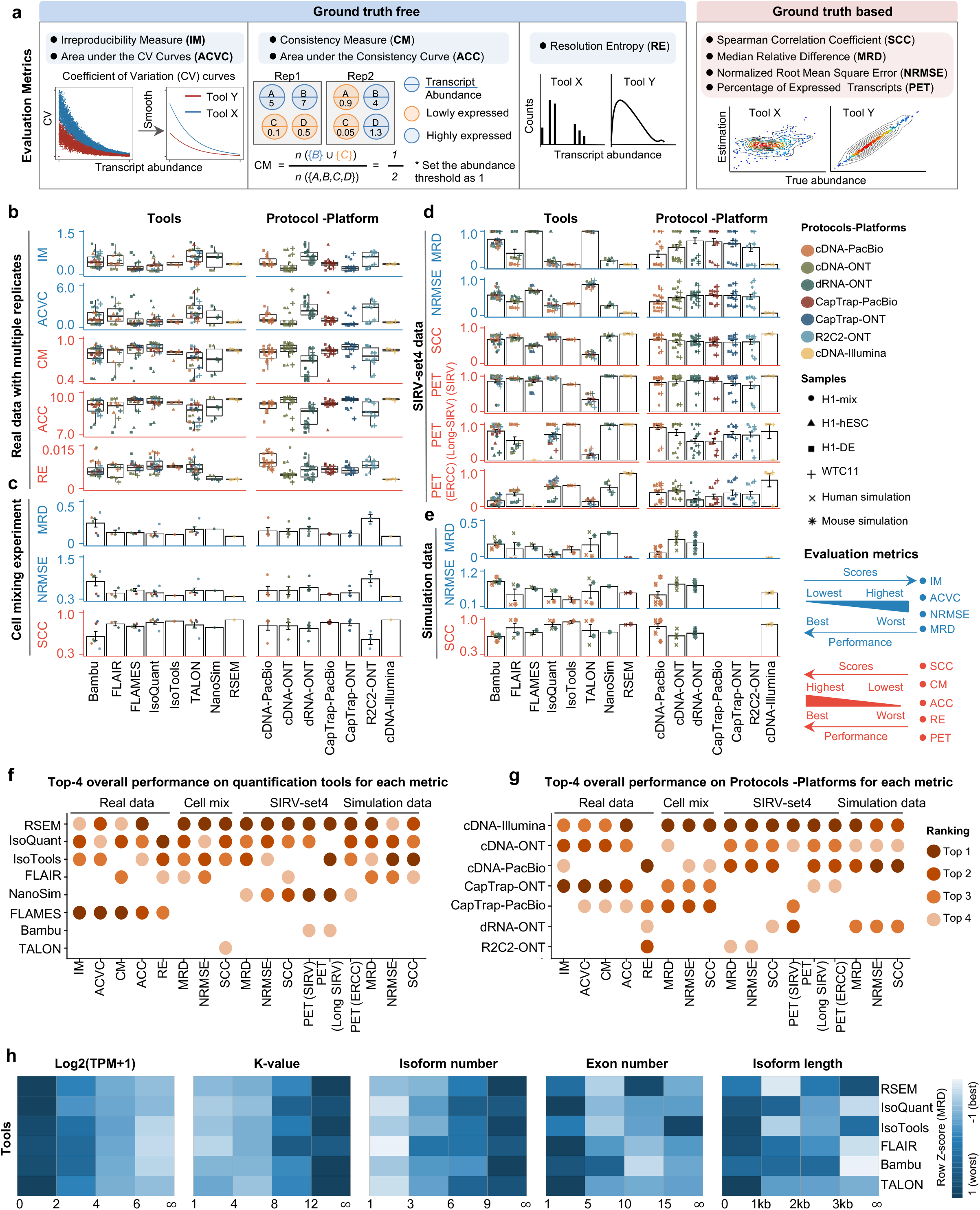
Overview of performance evaluation for Challenge 2: transcript isoform quantification. **(a)** Cartoon diagrams are used to explain 9 evaluation metrics under the ground truth given or not given. **(b) - (e)** Overall evaluation results of 8 quantification tools and 7 protocols-platforms on real data with multiple replicates, cell mixing experiment, SIRV-set4 data and simulation data. **(f) - (g)** Top-4 overall performance on quantification tools and protocols-platforms for each metric. **(h)** Evaluation of quantification tools with respect to multiple transcript features, including the number of isoforms, number of exons, isoform length and a customized statistic K-value representing the complexity of exon-isoform structures. Here, we use the normalized MRD metric to evaluate performance on human cDNA-PacBio simulation data. Additionally, we show RSEM evaluation results with respect to transcript features based on human short-read simulation data as a control.

**Table 4:** Metrics for Challenge 2 evaluation.

| Metrics | Description |
| --- | --- |
| Irreproducibility Measure (IM) and Area under the Coefficient of Variation Curve (ACVC) | IM and ACVC characterize the coefficient of variation of abundance estimates among multiple replicates. |
| Consistency Measure (CM) and Area under the Consistency Curve (ACC) | CM and ACC characterize the similarity of abundance profiles between pairs of replicates. |
| Resolution Entropy (RE) | RE characterizes the resolution of abundance estimation. |
| Spearman Correlation Coefficient (SCC) | SCC evaluates the monotonic relationship between the estimation and the ground truth. |
| Median Relative Difference (MRD) | MRD is the median of the relative difference of abundance estimates among all transcripts. |
| Normalized Root Mean Square Error (NRMSE) | NRMSE measures the normalized root mean square error between the estimation and the ground truth, which characterizes the variability of the quantification accuracy. |
| Percentage of Expressed Transcripts (PET) | PET characterizes the percentage of truly expressed transcripts in SIRV-set4 data |

First, we assessed the overall performance of the eight quantification tools with different protocols-platforms under four different data scenarios by considering the metrics evaluated above (**Figs. 3b-f**, and **Extended Data Fig. 53**).

For real data with multiple replicates, four metrics (Irreproducibility <u>M</u>easure/IM, <u>A</u>rea under the <u>C</u>oefficient of <u>V</u>ariation <u>C</u>urve/ACVC, <u>C</u>onsistency <u>M</u>easure/CM, and <u>A</u>rea under the <u>C</u>onsistency <u>C</u>urve/ACC) were designed to evaluate the reproducibility and consistency of transcript abundance estimates among multiple replicates (**Fig. 3b**, **Extended Data Figs. 54-55 and Extended Data Table 5**). Overall, FLAMES and IsoQuant, together with IsoTools, had similar overall performance: low IM and ACVC and high CM and ACC. They achieved comparable performance with the control RSEM tool (e.g., 0.18-0.33 vs. 0.35 and 0.65-0.86 vs. 0.77 for IM and ACVC, 0.81-0.88 vs. 0.84 and 9.31-9.46 vs. 9.51 for CM and ACC). FLAIR and Bambu followed closely behind, performing slightly worse than RSEM but outperforming the other quantification tools (**Fig. 3b**). Specifically, IsoQuant on cDNA-ONT and CapTrap-ONT, as well as FLAMES on CapTrap-ONT, demonstrated the top 3 performances across all datasets (<0.15 and <0.53 for IM and ACVC, >0.89 and >9.53 for CM and ACC). Of note, all tools showed poor performance on dRNA-ONT (mean IM=0.66, ACVC=2.62, CM=0.64, and ACC=8.31), which may be attributed to the low throughput of dRNA-ONT (<1 Million for each replicate).

In addition, the <u>R</u>esolution <u>E</u>ntropy (RE) metric was also used to characterize the resolution of transcript abundance estimates among multiple replicates in real data (**Fig. 3b**, **Extended Data Fig. 56**, and **Extended Data Table 5**). Among the REs, the top 6 tools (IsoQuant, IsoTools, FLAMES, FLAIR, TALON, and Bambu) achieved comparable resolution of transcript abundance, which were at least 2.7 -fold higher RE compared to the NanoSim and RSEM. This may be that NanoSim and RSEM use the GENCODE reference annotation, which contains a large number of transcripts not expressed in a specific sample, resulting in many transcripts with low expression in the quantification results (79.02% and 58.04% of transcripts with TPM<=1 in H1-hESC sample).

Due to the challenges of transcript-level quantification and the lack of a gold standard in real data, we designed an evaluation strategy using a cell mixing experiment (see details in **Extended Data Fig. 57a**), in which an undisclosed ratio of two samples (H1-hESC and H1-DE) is mixed before sequencing and provided to the participants at an initial phase. After predictions of transcript abundance in the mixed sample, sequencing data from the individual H1-hESC and H1-DE samples were released, and participants submitted quantification separately on these two datasets. The quantification of the mix samples should be equivalent to the expected ratios from the quantification of the individual cell lines. Three metrics (<u>S</u>pearman <u>C</u>orrelation <u>C</u>oefficient/SCC, <u>M</u>edian <u>R</u>elative <u>D</u>ifference/MRD, and <u>N</u>ormalized <u>R</u>oot <u>M</u>ean <u>S</u>quare <u>E</u>rror/NRMSE) were used to evaluate quantification accuracy by comparing expected and observed abundance (**Fig. 3c** and **Extended Data Table 10**. Most quantification tools generally had a good correlation (0.74-0.87 for mean SCC) between expected abundance and observed abundance, except for Bambu (0.53). In particular, RSEM had the best performance in cell mixing experiments (**Fig. 3c** and **Extended Data Fig. 57b**), which had the highest SCC (0.87) and lowest MRD (0.13) and NRMSE (0.38) values. Compared with the long-read-based tools, IsoQuant on cDNA-ONT had the best performance in MRD (0.14) and SCC (0.85), and FLAIR on cDNA-ONT had the lowest NRMSE (0.43).

Furthermore, SIRV-set 4 and the simulation data were used to evaluate how close the estimations and the ground truth values are by four metrics: <u>P</u>ercentage of <u>E</u>xpressed <u>T</u>ranscripts (PET), SCC, MRD, and NRMSE (**Figs. 3d,e**, **Extended Data Figs. 58,59** and **Extended Data Tables 11,12**). Generally, for SIRV-set 4 data, there is a large variation in the number of SIRV transcripts with TPM>0 quantified by tools (28-136). RSEM outperformed other long-read-based tools in average SCC (0.84 vs. 0.29-0.78), MRD (0.12 vs. 0.13-1.00), and NRMSE (0.45 vs. 0.89-2.19) scores. Especially in long-read-based tools, NanoSim (SCC=0.78, MRD=0.23, and NRMSE=0.89) and IsoQuant (0.76, 0.19, and 0.89) had the best overall performance, and IsoTools (0.69, 0.13 and 1.02), FLAIR (0.73, 0.42 and 1.13) and Bambu (0.68, 0.79 and 1.55) followed behind. All tools except TALON and FLAMES could better quantify the regular and long SIRV transcripts with TPM>0 (PET>80%). Conversely, most methods had poor performance for quantifying ERCC transcripts with TPM>0 (PET<50%), which may be attributed to the low expression levels of many ERCC transcripts ^31,32^.

For simulation data, all tools performed significantly better on PacBio data than ONT data (0.96 vs. 0.69 for mean SCC, 0.07 vs. 0.23 for mean MRD, and 0.25 vs. 0.78 for mean NRMSE). In particular, FLAIR, IsoQuant, IsoTools, and TALON on cDNA-PacBio had the highest correlation (SCC>0.97) between the estimation and the ground truth compared with other pipelines, which was slightly better than RSEM (SCC=0.90) and outperformed other long-read pipelines (SCC<0.83). Moreover, we observed that the transcript annotation accuracy significantly impacted quantitative accuracy. Specifically, when using an inaccurate annotation, RSEM yielded mean NRMSE values 2.74 and 3.27 times higher than those of LR-based tools and RSEM with accurate annotation, respectively (**Extended Data Fig. 60**, mean NRMSE: 1.70 vs. 0.62 vs. 0.52).

This emphasizes the critical importance of identifying a sample-specific accurate annotation for accurate transcript quantification.

Next, we evaluated the overall performance of the seven protocols-platforms across all quantification tools (**Figs. 3b-e**, **Fig. 3g**, and **Extended Data Fig. 53**).

Among the reproducibility and consistency in real data (**Fig. 3b**), CapTrap-ONT, CapTrap-PacBio, cDNA-PacBio, and cDNA-ONT had similar overall performance: low IM and ACVC, and high CM and ACC, which was significantly better than dRNA-ONT and R2C2-ONT (0.20-37 vs. 0.61-0.63 for IM, 0.50-1.05 vs. 2.39-2.87 for ACVC, 0.82-0.89 vs. 0.69-0.76 for CM and 9.19-9.51 vs. 8.61-8.68 for ACC). This is probably due to the relatively lower sequencing depth of dRNA-ONT and R2C2-ONT compared with other protocols-platforms (**Fig. 1b**). In particular, CapTrap-ONT and cDNA-ONT have the lowest irreproducibility (mean IM= 0.19 and 0.20, and ACVC=0.50 and 0.51) and highest consistency (mean CM= 0.89 and 0.86, and ACC=9.49 and 9.51). In terms of the abundance resolution, cDNA-PacBio, and R2C2-ONT dominated the other protocols-platforms, which is at least 2-fold higher RE compared to the cDNA-ONT (**Fig. 3b**). Notably, there are bimodal distributions of read length for some of the protocols-platforms (cDNA-PacBio, CapTrap-PacBio, dRNA-ONT definitely for R2C2-ONT, **Extended Data Fig. 61**) in real data. The sequencing error rate varies across the different sequencing platforms (**Fig. 1b**). Some tools may have a distinct advantage when dealing with particular data types if it takes into account the characteristics of these data (**Fig. 3f**).

For the cell mixing experiments (**Fig. 3c** and **Extended Data Fig. 57b**), CapTrap-PacBio, CapTrap-ONT, cDNA-PacBio, cDNA-ONT, and dRNA-ONT have similar performances with mean SCC scores between 0.73 and 0.83, while the remaining R2C2-ONT did not get to mean SCC scores above 0.60. MRD and NRMSE showed consistent evaluation results with SCC (e.g., 0.16-0.21 vs. 0.32 for MRD and 0.45-0.63 vs. 0.88 for NRMSE). In particular, CapTrap-PacBio exhibited the best overall quantification accuracy (SCC=0.83, MRD=0.16, and NRMSE=0.45) between expected abundance and observed abundance compared with other long-read-based protocols-platforms, which can achieve comparable performance with cDNA-Illumina.

For the SIRV-set 4 data, cDNA-PacBio had the best overall performance in long-read-based protocols-platforms, e.g., highest SCC (0.70 vs. 0.60-0.66) and lowest MRD (0.40 vs. 0.58-0.75) and NRMSE values (1.14 vs. 1.38-1.52). cDNA-ONT followed behind and outperformed the other four protocols-platforms (**Fig. 3d**). Notably, the ability of all protocols-platforms to quantify ERCC transcripts with TPM>0 (Mean PET=33.01%) was significantly worse than that of regular SIRV (Mean PET=82.17%) and long SIRV transcripts (Mean PET=69.75%). In particular, cDNA-ONT, cDNA-PacBio, CapTrap-ONT, and R2C2-ONT achieved similar performance with PET between 34.39% and 43.27% in quantifying ERCC transcripts, which was higher than dRNA-ONT (18.35%) and CapTrap-PacBio (27.99%). With respect to the long SIRV transcripts, all protocols-platforms, except for CapTrap-PacBio and dRNA-ONT, can quantify more than 70% of transcripts with TPM>0. All protocols-platforms performed well for the regular SIRV transcripts, among which dRNA-ONT, CapTrap-PacBio, cDNA-PacBio, and cDNA-ONT were the most prominent (PET>82.00%). Similar to SIRV-set 4 data, the simulation study showed that the cDNA-PacBio exhibited better performance in SCC, MRD, and NRMSE than cDNA-ONT and dRNA-ONT (**Fig. 3e** and **Extended Data Fig. 59b**).

Finally, we evaluated the performance of quantification tools for different sets of genes/transcripts grouped by transcript features, including transcript abundance, isoform number, exon number, isoform length, and a customized statistic K-value representing the complexity of exon-isoform structures (**Fig. 3h** and **Extended Data Figs. 54b,c and 62**).

For real data with multiple replicates, all quantification tools revealed a decreased <u>C</u>oefficient of <u>V</u>ariation (CV) and increased CM on six protocols-platforms with increasing transcript abundances (**Extended Data Figs. 54b,c**). Furthermore, we explored the changes of normalized MRD metric with different gene/transcript features based on the human cDNA-PacBio simulation data (**Fig. 3h)**. The MRD scores on all quantification tools increased dramatically when the transcript abundance was TPM≤2. Therefore, there existed greater variabilities and errors of abundance estimation for lowly expressed transcripts on all quantification tools. In addition, we found that all quantification tools had poor performance at the high K-value and isoform number, which revealed that more complex gene structures tend to be harder to quantify accurately. Notably, most tools had difficulty accurately quantifying transcripts with less than five exons but performed well with isoform counts between 5 and 15 exons, except for RSEM and Bambu. Interestingly, transcripts shorter than 1 kb had larger quantification errors, but the performance of different tools for transcripts longer than 1 kb varied significantly.

In summary, our evaluation results demonstrated that the performance of quantification tools varies depending on the features of genes and transcripts, and it remains challenging to accurately quantify transcripts with low expression and complex structure. Notably, there are significant performance differences among quantification tools under different data scenarios (**Fig. 3f**). Overall, RSEM outperformed long-read-based tools across different protocols-platforms and evaluation metrics. IsoQuant, IsoTools, and FLAIR exhibited superior overall performance among long-read-based tools across different data scenarios. Additionally, FLAMES demonstrated good reproducibility and consistency across multiple replicates in real data, while NanoSim achieved superior performance in quantifying SIRV transcripts. Generally, in different protocols-platforms, cDNA-Illumina showed the best overall performance, ranking among the top 3 in all evaluation metrics except for RE (**Fig. 3g**). Meanwhile, cDNA-PacBio, cDNA-ONT, CapTrap-ONT, and CapTrap-PacBio exhibited overall good performance across various data scenarios, surpassing dRNA-ONT and R2C2-ONT. It is worth noting that some protocols-platforms, such as cDNA-PacBio, CapTrap-PacBio, dRNA-ONT definitely for R2C2-ONT, exhibited bimodal distributions of read length in real data, and the error rate in sequencing varies among different sequencing platforms. Therefore, considering the characteristics of the data, some tools may have a distinct advantage in dealing with specific data types.

Currently, the throughput of long reads is relatively low, and most of the long-read-based tools mainly focus on transcript identification. However, with the improvement of long-read-based tools and the increase in throughput, the quantification accuracy of long-read-based tools is likely to be further improved.

### Challenge 3 Results and Evaluation: *De novo* transcript isoform identification

We evaluated the potential of long-read methods for transcriptome identification without a reference annotation using two types of samples. The mouse ES sample analyzed in Challenge 1 represents a case of high-quality genome assembly and data. The manatee white blood cell data depicts a typical scenario for a non-model species where limited genomics resources are available, and samples are directly taken from a field experiment. Additionally, the manatee sample had an excess of SIRV spike-ins, representing a challenging dataset. A draft genome of the manatee was assembled using Nanopore and Illumina genomic reads (**Extended Data Fig. 63**) and provided to submitters to support analysis. Still, no genome annotation was allowed in either the manatee or mouse analyses of Challenge 3. Matched short-read RNA-seq data was available to all submitters.

Four different tools were evaluated in Challenge 3: Bambu, StringTie2+IsoQuant, RNA-Bloom^33^, and rnaSPAdes^34^, which submitted a total of 17 and 8 transcriptome prediction sets for the mouse ES and manatee samples, respectively (**Fig. 1f**). While all long-read methods predicted high transcript mapping rates to the genome, both for mouse and manatee data (**Extended Data Fig. 64**), the number of detected transcripts varied among analysis tools, ranging from ∼20K to ∼150K in mouse ES and from ∼2K to ∼500K in the manatee sample (**Figs. 4a,b**, **Extended Data Tables 12-13**). rnaSPADes predicted the largest number of transcripts and the largest fraction of non-coding sequences, followed by RNA-Bloom (**Fig. 4a**). Bambu has the least number of predicted transcripts in all analyzed samples. Since the actual genome annotation of the mouse is available, a SQANTI structural classification analysis of this sample is possible. Remarkably, most transcripts detected by all methods under the reference annotation-free scenario for the mouse data were novel transcripts. rnaSPAdes predicted a large fraction of intergenic and antisense transcripts, RNA-Bloom was enriched in NNC, StringTie2+IsoQuant reported a high number of NIC and NNC, while for Bambu the most abundant novel class was ISM in most scenarios (**Fig. 4a**). This strongly contrasts with results obtained by these last two tools in Challenge 1 when guided by the reference annotation, where a large majority of transcript detections were FSM (**Fig. 2a** and **Extended Data Fig. 65**) and shows that, at least for Bambu and IsoQuant, the utilization of a reference annotation has a strong impact in their predicted transcript models. Since no reference annotation is available for manatee, a structural category analysis is not possible. However, we evaluated the number of transcripts per locus detected by each method. Bambu, rnaSPAdes, and RNA-Bloom predicted a single transcript for most loci. In contrast, StringTie2+IsoQuant, especially when using cDNA-ONT data, predicted two or more transcripts for nearly half of the loci, especially when using cDNA-ONT data.

**Figure 4.**
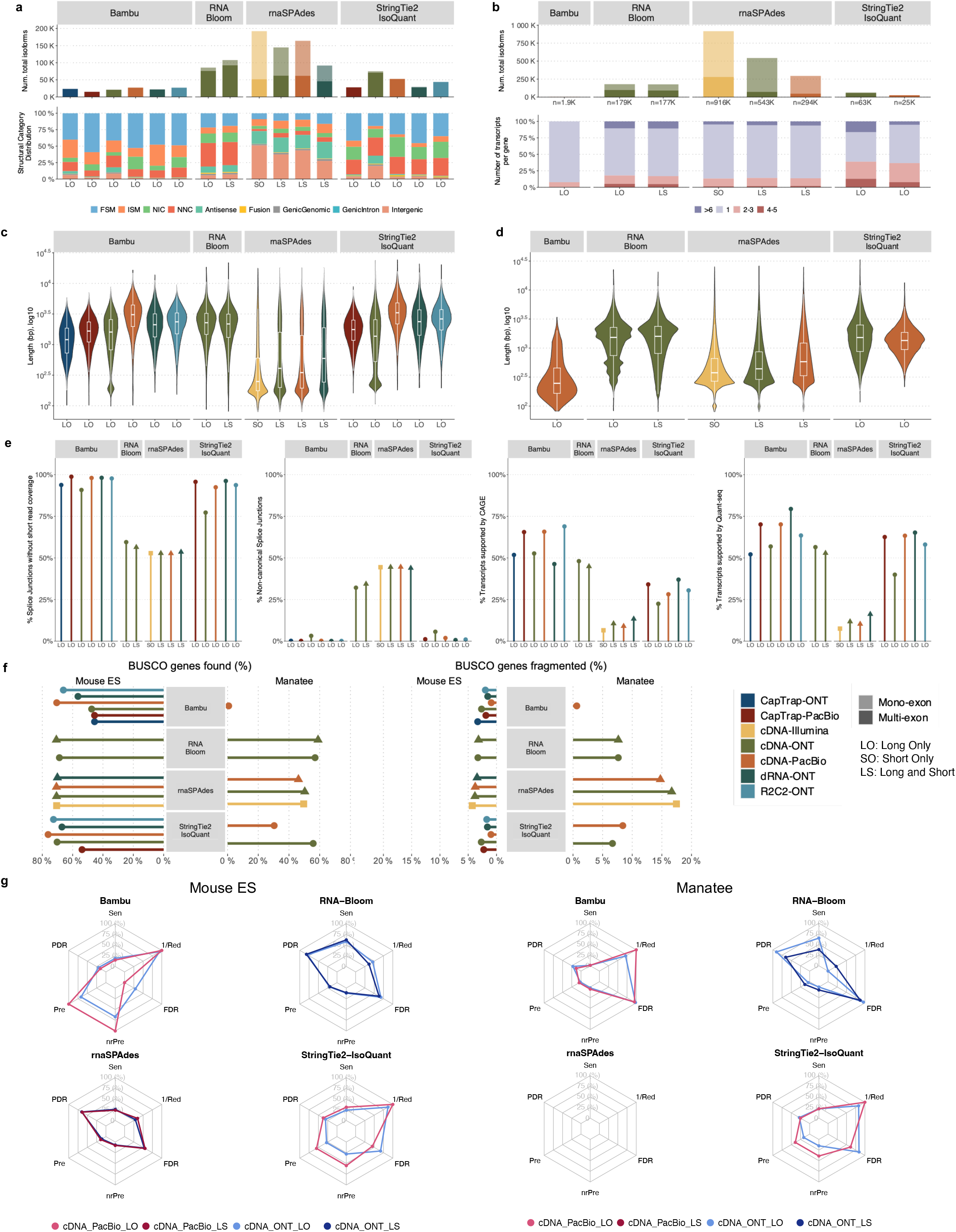
Evaluation of Challenge 3: transcript identification without a reference annotation. **a)** Number of detected transcripts and distribution of SQANTI structural categories, Mouse ES sample. **b)** Number of detected transcripts and distribution of transcripts per loci, Manatee sample. **c)** Length distribution of Mouse ES transcripts predictions. **d)** Length distribution of Manatee transcripts predictions. **e)** Support by orthogonal data. **f)** BUSCO metrics. **g)** Performance metrics based on SIRVs. Sen: Sensitivity, PDR: Positive Detection Rate, Pre: Precision, nrPred: non-redundant Precision, FDR: False Discovery Rate, 1/Red: Inverse of Redundancy.

Without a reference annotation, Bambu, StringTie2+IsoQuant, and RNA-Bloom could predict most transcripts models with lengths ranging between 1 kb and 3 kb. However, both Bambu and StringTie2+IsoQuant reported a significant number of short transcripts from the mouse ES cDNA-ONT dataset (**Fig. 4c**), possibly due to the large fraction of shorter reads in this sample (**Fig. 1c**) and Bambu had a lower transcript length distribution on the manatee cDNA-PacBio dataset (**Fig. 4d**). For all mouse ES and manatee samples rnaSPAdes returned a large number of short transcripts that strongly pulled down the length distributions (**Figs. 4c,d**).

When looking at supporting orthogonal data for the mouse ES transcript models, we found a clear relationship between the total number of predicted transcripts and the orthogonal support: rnaSPAdes, which predicted a large number of transcripts, had the least support for Illumina, CAGE, and Quantseq junctions, 5’ end and 3’ end respectively, and the largest (∼40%) number of non-canonical splice junctions. Bambu, on the contrary, with the least number of predicted transcripts, generally had the highest orthogonal support. RNA-Bloom had transcripts with only 50% of junctions supported by Illumina and a considerable fraction of non-canonical junctions, while support at 3’ and 5’ ends was moderate. StringTie2+IsoQuant produced transcripts with quality junctions but low CAGE support (**Fig. 4e**).

A majority of transcripts identified by Bambu and StringTie2+IsoQuant were predicted to be protein-coding both in the manatee and mouse ES samples, except for CapTrap-ONT and cDNA-ONT datasets, with ∼25% of transcript models predicted to be non-coding, which could be related to the higher number of reads in this dataset. RNA-Bloom predicted a similar percentage of protein-coding genes on the cDNA-ONT sample but was significantly lower in rnaSPAdes transcript models. Moreover, all tools predicted a lower fraction of transcripts with coding potential in the manatee sample, ranging from ∼70% for IsoQuant and Bambu and less than 20% for all rnaSPAdes predictions (**Extended Data Fig. 66**).

We used the BUSCO^35^ database of highly conserved genes to assess the relative completeness of the predicted transcriptomes. Interestingly, despite the observed differences in protein-coding transcript rates, BUSCO analysis indicated a relatively good performance in most cases. On the mouse ES sample, rnaSPAdes, RNA-Bloom detected above 60% of complete BUSCO genes, while Bambu only reached this threshold when analyzing cDNA-PacBio and R2C2-ONT data. In the case of the manatee sample, the highest BUSCO completeness (∼50%) was achieved by IsoQuant and RNA-Bloom on the Nanopore datasets, rnaSPAdes returned ∼30% of complete BUSCO genes, and performance for Bambu was poor. Following these results, the fraction of incomplete BUSCO genes was generally lower in the mouse ES than in the manatee sample, and the highest ratio of incomplete BUSCO genes was provided by rnaSPAdes in the manatee sample (**Fig. 4f**).

Performance analysis using SIRV spike-ins revealed significant differences between tools and samples. On the mouse ES sample SIRVs were detected with relatively high sensitivity (∼70%) by RNA-Bloom. However, the precision was low, and the False Discovery Rate (FDR) was high.

A similar pattern was observed by rnaSPAdes, although this tool had low sensitivity and a high Positive Detection Rate, indicating the detection of SIRVs by incomplete transcript models. RNA-Bloom and rnaSPAdes had low values for 1/Red, implying that multiple transcripts models were predicted for the same SIRV. On the contrary, StringTie2+IsoQuant and Bambu had low sensitivity (∼25%) but medium to high precision and better control of the FDR. For both tools, precision figures were better when analyzing cDNA-PacBio data (**Fig. 4g**). Analysis of the manatee sample, which contained an excess of SIRVs, resulted in generally poorer performance results for all pipelines, especially for Bambu, which was unable to recover the spiked RNAs. On the manatee sample, the best sensitivity for SIRVs was obtained by RNA-Bloom (∼70%) and the best precision by StringTie2+IsoQuant on cDNA-PacBio (∼40%) (**Fig. 4g**). To understand whether the limited sensitivity of the pipelines was a data or algorithm issue we evaluated mapped SIRVs reads with SQANTI. Interestingly, we found that over 90% and 46% of the SIRV reads were FSMs and that 90% and 64% of the SIRVs had at least one reference-match read in the manatee cDNA-PacBio and cDNA-ONT datasets (**Extended Data Fig. 67**), suggesting that sufficient quality data was present and that sensitivity may be challenged at the correct processing of the noisy reads.

As expected, transcript detection without a reference annotation is a challenging problem. Interestingly, transcripts with higher coverage, as in our SIRV spike-ins in the manatee sample, led to poorer performance for all tools, suggesting that accurate detection from highly expressed genes may be problematic. Overall, Bambu and IsoQuant had moderate to good precision but low sensitivity. RNA-Bloom had high sensitivity but low precision, and rnaSPAdes produced many fragmented and short transcripts with a high FDR.

### Experimental validation of transcript isoform predictions

To experimentally validate isoforms, we targeted isoform-specific regions for PCR amplification followed by gel electrophoresis and sequencing of pooled amplicons via ONT and PacBio sequencing (**Fig. 5a**). To support isoform-specific PCR design, we developed Primers-JuJu, a program employing Primer3^36^ for semi-automated isoform-specific primer design (see **Supplementary Information**). Due to practical limitations of the scale of experimental validation, we prioritized targets of interest in three comparison groups: GENCODE known and novel, consistently versus rarely identified novel isoforms, and ONT and PacBio preferential isoforms. The first comparison group included isoforms that GENCODE Annotators identified from LRGASP data and were classified as occurring in the GENCODE annotation (known) or not (novel). The second group compared isoforms identified consistently by more than half the pipelines or rarely by 1 or 2 pipelines. The final group was isoforms preferentially identified in ONT libraries or only in PacBio libraries. From these comparison groups, we designed primers for 178 target regions, in which the length of the amplified region ranged from 120 to 4406 bp long, with a median of 488 bp and 25-75 interquartile range of 305 to 795 bps.

**Figure 5.**
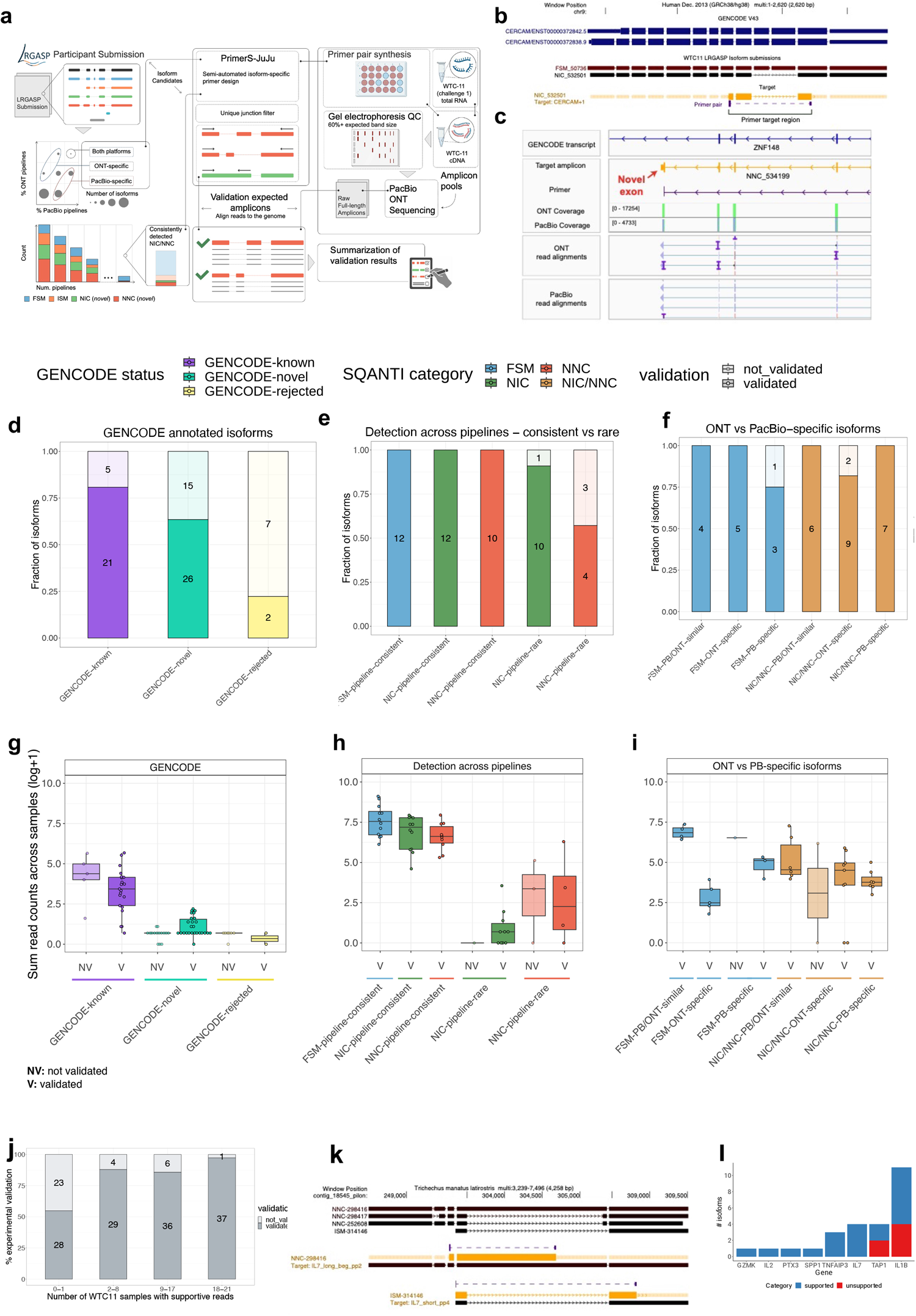
Experimental validation of known and novel isoforms. **a)** Schematic for the experimental validation pipeline. **b)** Example of a consistently detected NIC isoform (detected in over half of all LRGASP pipeline submissions) which was successfully validated by targeted PCR. The primer set amplifies a novel event of exon skipping (NIC). Only transcripts above ∼5 CPM and and part of the GENCODE Basic annotation are shown. **c)** Example of a successfully validated novel terminal exon, with ONT amplicon reads shown in the IGV track (PacBio produce similar results). **d)** Recovery rates for GENCODE annotated isoforms that are reference-matched (known), novel, and rejected. **e)** Recovery rates for consistently versus rarely detected isoforms, for known and novel isoforms. **f)** Recovery rates between isoforms that are more frequently identified in ONT versus PacBio pipelines. **g-i)** Relationship between estimated transcript abundances (calculated as the sum of reads across all WTC11 sequencing samples) and validation success for GENCODE (**g)**, consistent versus rare **(h**), and platform-preferential **(i)** isoforms. **j)** Fraction of validated transcripts as a function of the number of WTC11 samples in which supportive reads were observed. **k)** Example of two *de novo* isoforms in Manatee validated through isoform-specific PCR amplification, blue corresponds to supported transcripts and red to unsupported transcripts. **l)** PCR validation results for manatee isoforms for seven target genes.

Target validation was confirmed upon sequencing of the PCR-based amplicons on both ONT and PacBio platforms, and reads were aligned to determine support for the isoform (**Fig. 5a**, see **Methods**). Examples of a validated exon skipping event (NIC) and a novel terminal exon (NNC) are shown in **Figs. 5b,c**.

To evaluate GENCODE annotated isoforms, we compared groups of randomly selected isoforms that were 1) annotated (GENCODE-known, n=26), 2) novel and confirmed through manual annotation (GENCODE-novel, n=41), and 3) unsupported isoforms that were investigated but did not pass rigorous manual curation (GENCODE-rejected n=9). As expected, we found a high validation rate for GENCODE-known, 81% (**Fig. 5d**). Of the GENCODE-known isoforms, we found that 5 of the 28 targets failed to validate. Re-checking these isoforms confirmed a high degree of support for annotated introns from short-read RNA-seq datasets in the recount3 database, and we speculate that the failed validation reflects suboptimal primer or PCR conditions. GENCODE-novel isoforms validated at a slightly lower but still high validation rate (63%) compared to GENCODE-known. Manual review again confirmed GENCODE-novel isoforms that failed to validate as correctly annotated but showed that they tended to be lower in abundance compared to their successfully validated counterparts (**Fig. 5g**). In some cases, GENCODE annotators had confidently annotated a full-length isoform from a single long read sequenced in one experiment, and such examples were more difficult to validate. As expected, only two out of nine GENCODE-rejected isoforms were amplified (22% validation rate). Notably, upon close examination, we found that validated examples were mismapping cases due to tandem repeats, where the validated transcript sequence was correct but not the original junction model, which was correctly rejected by GENCODE as not supported by the initial aligned reads.

A large number of novel isoforms were detected in this study (e.g., 279,791 novel isoforms in WTC11, **Fig. 1b**). We found that 743 novel isoforms were detected consistently, but a vast majority, or 242,125 isoforms, were rarely detected, and found in only 1 or 2 of the pipelines. We obtained a 100% validation rate for consistently novel isoforms, underscoring their experimental amenability to PCR and sequencing-based processing (**Fig. 5e**). Strikingly, for isoforms with exceedingly low reproducibility across pipelines, we found a surprisingly high validation rate of 91% and 57% for NIC and NNC isoforms, respectively. Though the validation rate was lower for rarely detected isoforms, as they comprise a vast majority of novel isoforms reported in this study, a simple linear extrapolation would imply that many isoforms may be expressed in WTC11. Abundance correlated with a validation rate (**Fig. 5h**), as found for the GENCODE validation set.

Lastly, we determined the validation rates of known and novel isoforms in common or preferentially detected in the ONT and PacBio platforms for their respective cDNA preparations. For example, an isoform detected in more than 50% of ONT pipelines but less than 50% of PacBio pipelines would be considered ONT-preferential and vice versa. We found that all known and novel isoforms found frequently (more than 50% of all pipelines) across both platforms were validated (**Fig. 5f-i)**. We note that most of all validated isoforms were identified by amplicon sequencing on both ONT and PacBio. While the consistency of the validation is remarkable, we acknowledge that this subset’s relatively small sample size limits drawing general conclusions regarding validation rates for platform-preferential isoforms.

The validation experiments underscore that the transcript models from long-read pipelines will likely capture biologically real isoforms. In other words, novel isoform predictions generally have high accuracy, even if such isoforms are not consistently predicted across pipelines and platforms. Notably, our results suggest that validation success was related to the frequency of detection of the isoform, measured either by the number of WTC11 datasets (resulting from different combinations of library preparation and sequencing technologies) in which the isoform was detected (**Fig. 5j**) or by the total number of read counts supporting the isoform across WTC11 datasets (**Extended Data Fig. 68**). Other genomic features, such as transcript lengths, did not show a clear relationship with validation rates (**Extended Data Fig. 69**). While long-read transcript alignments are often able to resolve regions that are problematic for more accurate short-read alignments, these results highlight “blind spots” in a long-read-based transcript annotation, including imperfectly aligned reads (e.g., tandem repeats, inversions, short exon overhangs) in which isoform alignment can be highly affected by assumptions of the aligner. While expert human intervention can resolve many difficult cases, others remain unsolvable.

To validate long-read-based isoform discovery without a reference annotation, we focused on the manatee dataset. Compared to Challenges 1 and 2, Challenge 3 had fewer submissions; therefore, we established a goal of not explicitly comparing pipelines but, rather, assessing the ability of the long-read RNA-seq datasets, collectively, to return accurate transcript isoform annotation. To select genes and isoforms of interest, we used a targeted approach, focusing on a small, pre-defined list of genes related to immune pathways (see **Methods**).

Genes and their respective isoform targets were manually selected based on visualization and evaluation of the isoform structures on a custom UCSC Genome Browser track. We designed 22 primers that could potentially amplify 26 transcript predictions, with some of the primers targeting multiple transcripts. The length of the amplified region ranged from 78 to 2633 bp long, with a median of 1038 bp and a 25-75 interquartile range of 379 to 1379 bps. Validation of targets was confirmed upon PacBio sequencing of the amplicons. In total, we assessed isoforms from seven manatee genes (example shown in **Fig. 5k**). For five of the genes, those with one or a few isoforms, all isoforms were validated. For the two genes for which many isoform models were predicted and for which there was more variability across participants, approximately half of the targets were validated (**Fig. 5l**).

Overall, we find a greater technical challenge and variability of “field collected” non-model organism LR data, as compared to the human or mouse datasets. Though our sample population was small, they tend to validate when many pipelines coincide in their predictions of the number and identity of isoforms. In addition, ONT platforms tended to predict more isoforms than PacBio platforms, although with a higher false positive rate (**Extended Data Fig. 70**).

## Discussion

The LRGASP project aimed to provide a thorough and impartial evaluation of long-read sequencing methods for characterizing the transcriptome. Replicated data was gathered for various sequencing platforms and library preparation methods, and 14 bioinformatics approaches took part in one or more of the three challenges. Predictions were submitted by the tool developers, allowing for optimal utilization of the methodologies, and evaluation scripts were provided along with the data to enable self-assessment and transparent review.

We found significant differences among sequencing platforms and library preparation methods in terms of the number and quality of reads obtained. ONT sequencing of cDNA and CapTrap libraries produced ten times more reads than other combinations, while cDNA-PacBio and R2C2-ONT provided the longest and more accurate reads. Interestingly, more reads did not consistently lead to more transcripts, indicating that read quality and length are important factors for transcript identification. We also found a large influence of the analysis tool on the results and identified fundamental differences in the strategy followed by each algorithm. Some tools used long reads as evidence to recall known transcripts, leading to predictions biased toward the reference annotation. Other methods inferred transcript models from the actual reads and were more receptive to novelty. Some of these approaches returned a broad set of transcript predictions based on the processed sequences neglecting possible RNA degradation and library preparation errors. In contrast, others targeted the identification of novel transcripts with high confidence. These different conceptions of the data analysis challenged our evaluation strategy, prompting us to examine different aspects of the transcript model support. Thanks to the numerous evaluation metrics and orthogonal data sources used in the benchmark, we accomplished this.

Remarkably, our three approaches for which ground truth was available yielded different conclusions about performance. Assessment using SIRVs, where the annotation was known, concluded high accuracy for the majority of methods. In contrast, the simulated data indicated poorer performance for novel transcripts, and the GENCODE manual annotation revealed that most tools failed to predict most of the transcript models inferred by human annotators, likely due to their low expression. However, our experimental validation results confirmed the presence of many novel transcripts, highlighting the relevance of long-read sequencing technologies in profiling the complexity of transcriptomes.

The novel transcript class with the highest overall consensus and validation was the NIC, implying that novel combinations of splice junctions, TTS, and TSS in the transcriptome are to be expected, at least for well-characterized organisms like mouse and human. Surprisingly, ∼50% of the tested NNCs were validated, indicating additional novelty and high false discovery in this category. These results point to the utility of having a deep reference annotation capturing the largest possible catalog of splice features. Interestingly, many of these novel transcripts were detected by just one or few reads, found in only one or few samples, yet still validated by PCR. This suggests that many rare RNA molecules are present in specific samples, raising questions about considering them when reporting transcriptome composition using long reads. Arguably, while the comprehensive profiling of the RNA molecular content of a particular sample may require the inclusion of any detected transcript when defining the transcriptional signature of cell types and cell states, including only consistently detected transcripts might be advisable. The LRGASP results show that the analysis strategies implemented by the different tools are differently suited for these two scenarios.

Long-read transcriptome sequencing is increasingly being used to support the annotation of reference genomes, and, likely, projects such as VGP and BioGenome will extensively use lrRNA-seq. While the LRGASP challenge was not designed to assess the full annotation of new genomes, our Challenge 3 results suggest that *de novo* annotation of genomes based solely on long-read sequencing is challenging. Results were highly variable, more tools should be developed and tested, and data shows room for improvement. Reassuringly, experimental validation of a few manatee loci with multiple predicted transcripts again demonstrated the potential of long-read methods in revealing the transcriptome complexity of non-model organisms.

Notably, our evaluation results in Challenge 2 highlight several important factors that may impact the performance of transcript quantification tools. Firstly, the accuracy of transcript annotations significantly influences quantitative accuracy, underscoring the pressing need to identify accurate sample-specific annotations using long-read sequencing technology. Additionally, the bimodal read length distributions observed for several protocols-platforms in real data, including cDNA-PacBio, R2C2-ONT, CapTrap-PacBio, and dRNA-ONT, demonstrate the need for transcript quantification tools that can accurately handle varying read lengths. Furthermore, the variation in sequencing error rates across different sequencing platforms emphasizes the importance of robust tools that can adapt to different data scenarios. Moreover, the relatively low throughput of long reads limits their use in quantification, with most long-read-based tools focused on transcript identification, leading to less impressive performance than short-read-based tools. However, we anticipate the continued development of long-read-based tools and increased throughput will improve quantification accuracy. Meanwhile, our results also reveal the persistent challenge of accurately quantifying structurally complex and lowly expressed transcripts. These transcripts may require different quantification strategies to overcome the technical limitations of existing tools. Thus, selecting appropriate quantification tools and optimizing them for specific data scenarios will be crucial for achieving accurate transcript quantification.

We acknowledge some limitations of our benchmark. LRGASP did not address the evaluation of long-read mapping methods, and we noticed that all tools, including SQANTI3, used minimap2^37^ as the aligner. However, the GENCODE manual curation identified mapping errors that led to incorrect junction models. Therefore, a separate evaluation of long-read mapping algorithms is advisable. Moreover, due to the multiple datasets and analysis modalities present in the Challenge, participants were allowed to submit only one set of predictions in each case, and it was up to them to decide how to configure their methods for the best performance. However, most of these tools are highly customizable and allow fine-tuning depending on the user’s preferences, which, as discussed, is relevant to adapt to specific analysis needs. Furthermore, to facilitate participation, submitters were allowed to choose which data modalities they contributed to, and many chose a subset, resulting in unbalanced data when evaluating methods, which might have influenced some of our conclusions. For example, tools returning the largest number of transcripts only used PacBio datasets and did not analyze Nanopore data. We tried to consider this in our analyses, but balanced participation would have been more useful. Additionally, despite our outreach efforts, several long-read analysis tools (e.g., TAMA^38^, IsoSeq3^39^, LIQA^40^, and ESPRESSO^41^) did not participate in LRGASP or were released after the start of the effort. To address this limitation, we offer the LRGASP datasets and evaluation strategy through OpenEBench^42^, an online platform that allows continuous benchmarking of tools and comparison to previous approaches. We encourage the community to use this resource to develop new lrRNA-seq analysis tools. Finally, as sequencing technologies continue to improve in throughput and accuracy, the dataset’s quality is expected to change in the future, and some of our conclusions may need revision.

### Final recommendations

Based on the results of LRGASP, the Consortium recommends the following suggestions to improve the analysis of transcriptomes using long reads:

1. For transcript identification, longer and more accurate sequences are preferable to having more reads. Therefore, the cDNA-PacBio and R2C2-ONT datasets are the best options. However, if the goal is quantification, especially if it is based on a reference annotation, cDNA-ONT is the best choice.
2. When choosing a bioinformatics tool, it is crucial to consider the study’s objective:

a. If the goal is to identify a sample-specific transcriptome in a well-annotated organism when only minimal novel transcripts are expected, Bambu, IsoQuant, and FLAIR are the most effective tools
b. If the aim is to detect lowly expressed or rare transcripts, use a tool that allows novelty and includes orthogonal data. Mandalorion and FLAIR, combined with short reads, are among the best performers for detecting novel transcripts. If feasible, experimental validation of the rare transcripts should also be included.
3. If quantification is essential, IsoQuant, IsoTools, and FLAIR are the best options and can perform comparably to short-read tools.
4. To create a reference for genome annotation, we recommend using high-quality data and including replication. It is also recommended to use extensive orthogonal data to validate predictions. The accuracy of the transcript calls can be improved by imposing a filter on transcript levels and combining more than one analysis tool.

## Methods

### GENCODE manual annotation

Expert GENCODE human annotators sought to establish the baseline for annotating genuine alternative isoforms using the long transcriptomic data generated by the LRGASP consortium at selected loci. To do this, an exhaustive and fully manual investigation of all aligned reads from the sequence data generated by the LRGASP consortium was undertaken at selected loci, and all isoforms passing GENCODE annotation criteria that were present were captured as annotated transcript models.

GENCODE expert human annotation has very high sensitivity and specificity as every read can be individually considered on its merits for use in supporting a transcript model that is subsequently included in the annotation set. Consequently, the fully human annotation process where every read is manually reviewed and every transcript model built manually is time-consuming. The speed of the process limits the number of loci that could be considered for the LRGASP project, with 50 human and 50 mouse loci being selected.

The selection of loci was random within constraints based on the properties of the locus, identification of aligned reads in all libraries sequenced (where possible), the number of aligned reads, and an indication of the presence of valid isoforms at the locus.

Properties required of the locus to be considered for annotation included the presence of multiple exons and compactness. Loci with very long introns were excluded. While criteria required that (where possible) all libraries had at least one aligned read at the locus, a maximum number of aligned reads also had to be added. This additional requirement was necessary for two reasons; firstly, to allow the manual consideration of every read at the locus to determine whether it could support an isoform, and secondly, because the large number of reads generated from the LRGASP sequencing experiments exposed bugs affecting the consistent display of transcripts when loaded into the Otter/Zmap^43^ tools used for manual annotation, raising the possibility of erroneous exclusion of reads and transcripts that failed to display properly.

Preliminary analysis was conducted to identify plausible alternative splicing in aligned reads using Tmerge^44^ at most permissive settings to create a set of putative transcripts from the LRGASP long transcriptomic data. The introns of this set of putative transcripts were assessed using Recount3^45^ data, and those putative transcripts where all introns were supported by at least two RNAseq reads from the GTEx^46^ dataset captured by Recount3 analysis were considered to be alternatively spliced for locus selection. The transcript models generated that this stage was not directly included in the GENCODE annotation.

A long list of human loci fitting the criteria for annotation was then compared to an equivalent list derived from a similar analysis in mouse, and candidate loci were defined. Human and mouse shortlists were manually reviewed to confirm selection criteria were met. Genuine alternative splicing events were maximized, and, where possible orthologous loci were selected for human and mouse.

Expert human annotation was carried out independently for each library prep method. Independence was defined as not using reads from one library prep to support reads in another. For example, a longer or higher quality read from one library prep method supported the interpretation of a truncated or low-quality read in another library.

Effectively this necessitated 12 independent sets of annotation, six in human and six in mouse, to support complete flexibility in downstream analysis.

While annotation was performed independently for each library, orthogonal data external to LRGASP was used to support the interpretation of the long transcriptomic data. Specifically, Recount3 intron data was used to support the interpretation of splice sites and Fantom CAGE^47^ data, the definition of transcription start sites (TSS), and thereby the 5’ completeness of a transcript. External l,ong transcriptomic datasets were not used to support this annotation in any way.

All transcript models passing standard Ensembl-GENCODE manual annotation criteria for splicing and supported by at least one long transcriptomic read were annotated as transcript models. GENCODE annotation criteria require that introns are canonical (GT-AG, GC-AG, or AT-AC with evidence of U12 splicing) or those non-canonical introns are supported by evidence of evolutionary conservation or constraint of the splice site. Read data was required to align such that a canonical intron could be unambiguously resolved, which generally requires that there is no equally plausible alignment of the read that could give a non-canonical ‘intron,’ for example, where sequence aligns equally well at the putative donor and acceptor splice site but can be forced into a canonical splice site by the initial alignment method, or where a read has an indel near a splice site that leads to an error in its initial alignment. Where necessary, annotators could realign the read to the genomic sequence using various methods, including the Exonerate pairwise alignment software and the Dotter^48^ dot plot tool. Introns identified by spanning RNAseq reads by the Recount3 project were used as orthogonal data to support the interpretation of splice sites.

Transcript models were extended to the full length of the homology between the read (or reads) supporting a transcript model and the genome sequence. 5’ transcript ends were not modified (clipped or extended) based on annotation already present at the locus before LRGASP (including the MANE Select^49^ transcript). While Fantom CAGE data was used to identify TSS at loci in both human and mouse, CAGE data was not used to modify the TSS. Similarly, at the 3’ end, transcripts were extended to the full length of the homology between the read and the genome. Where a poly(A) site was identified, the transcript model was not extended further unless another, longer read with the same intron chain was identified. In addition to the annotation of alternative isoforms, expert human annotators also defined sets of polyadenylation sites and signals at every locus. Again these were annotated independently per library prep based on the presence of a poly(A) tail on one or more reads aligned to the locus and a poly(A) site hexamer (AATAAA or ATTAAA) within 50 bases upstream of the poly(A) site. Multiple poly(A) sites and their corresponding poly(A) signals could be annotated at any locus. However, only one poly(A) site was annotated per poly(A) signal.

In total, 635 transcript models and 641 poly(A) sites were annotated in human. The mean number of isoforms annotated per locus per library ranged from 2.58 (cDNA-PacBio) to 1.28 (CapTrap-PacBio) in human and from 2.5 (cDNA-PacBio) to 1.38 (dRNA-ONT). The mean number poly(A) sites annotated per locus per library ranged from 2.42 (cDNA-PacBio) to 1.12 (dRNA-ONT) in human and 2.26 (CapTrap-ONT) to 1.2 (dRNA-ONT) in mouse. The loci with the most annotated alternative splicing were ANGPT1 (cDNA-PacBio library), RERG ZADH2 (CapTrap-ONT) with six isoforms. ZADH2 also had the most poly(A) sites, with 12 reported in three libraries (cDNA-PacBio, cDNA-ONT, and CapTrap-ONT). For mouse, nine isoforms were annotated at Rcan1 (cDNA-PacBio), while Gan displayed the most diversity in polyadenylation with ten poly(A) sites annotated (cDNA-PacBio).

Expert human annotators reviewed transcript models that were included in the annotation set but failed to validate by RT-PCR, and also, a set of reads were rejected as supporting valid transcript models by the first pass annotation. In both cases, the initial annotation of the transcript model was supported by review (**Extended Data Table 15)**.

## Computational evaluation of transcript isoform detection and quantification

### Challenge 1 Evaluation: Transcript isoform detection

Four sets of transcripts were used for the evaluation of transcript calls made on human and mouse lrRNA-seq data

1. Lexogen SIRV-Set 4^50^ (SIRV-Set 3 plus 15 new long SIRVs with sizes ranging from 4 to 12 kb)
2. Comprehensive GENCODE annotation: human v39, mouse vM28. GENCODE human v38 and vM27 were available at the time of the LRGASP data release, and new versions of GENCODE were released after the close of LRGASP submissions.
3. A set of transcripts from a subset of undisclosed genes which were manually annotated by GENCODE. These transcripts are considered high-quality models derived from LRGASP data
4. Simulated data for both Nanopore (NanoSim) and PacBio (IsoSeqSim) reads

The rationale for including these different types of transcript data is that each set creates a different evaluation opportunity but has its limitations. For example, SIRVs and simulated data provide a clear ground truth that allows the calculation of standard performance metrics such as sensitivity, precision, or false discovery rate. Evaluation of SIRVs can identify potential limitations of both library preparation as well as sequencing, but the SIRVs themselves represent a dataset of limited complexity, and the SIRV annotation is known to submitters. Higher complexity can be generated when simulating long reads based on actual sample data. However, read simulation algorithms only capture some potential biases of the sequencing technologies (e.g., error profiles) and not of the library preparation protocols. In any case, both types of data approximate but do not fully recapitulate real-world datasets. Evaluation against the GENCODE annotation^16^ represents this real dataset scenario, although in this case the ground truth is not entirely known. This limitation was partially mitigated by the identification of a subset of GENCODE transcript models that were manually annotated by GENCODE annotators, and by follow-up experimental validation for a small set of transcripts using semi-quantitative RT-PCR and quantitative PCR (qPCR) approaches. In this way, although an exhaustive validation of the real data is not possible, estimates of the methods’ performances can be inferred. By putting together evaluation results obtained with all these different benchmarking datasets, insights can be gained on the performance of the library preparation, sequencing, and analysis approaches both in absolute and in relative terms.

The evaluation of the transcript models was guided by the use of SQANTI categories (**Fig. 2a**), implemented in the SQANTI3 software. It incorporated additional definitions and performance metrics to provide a comprehensive framework for transcript model assessment (**Table 2**). The evaluation considers the accuracy of the transcript models both at splice junctions and at 3’/ 5’ transcript ends. It took into account external sources of evidence such as CAGE and Quant-seq data, poly(A) annotation, and support by Illumina reads (**Fig. 2b**).. The evaluation script was provided to participants at the time of data release (**Data and code availability**).

Given the LRGASP definitions, evaluation metrics were specified for each type of data.

#### SIRVs

To evaluate SIRVs, we extracted from each submission all transcript models that associate to SIRV sequences after SQANTI3 analysis. This includes FSM and ISM isoforms of SIRVs and NIC, NNC, antisense, and fusion transcripts mapping to SIRV loci. The metrics for SIRV evaluation are shown in **Table 5**.

**Table 5:** Metrics and definitions for evaluation against SIRVs.

|  |  |
| --- | --- |
| Reference SIRV (rSIRV) | Ground truth SIRV model |
| SIRV_transcripts | Transcripts mapping to a SIRV chromosome |
| SIRV_RM | SIRV_transcripts associated to at True Positive |
| True Positive detections (TP) | rSIRVs identified as RM |
| Partial True Positive detections (PTP) | rSIRVs identified as ISM or FSM_non_RM |
| False Negative (FN) | rSIRVs without FSM or ISM |
| False Positive (FP) | NIC + NNC + antisense + fusion SIRV_transcripts |
| Sensitivity | TP/rSIRVs |
| Precision | RM/SIRV_transcripts |
| Non_redundant Precision | TP/SIRV_transcripts |
| Positive Detection Rate | unique(TP+PTP)/rSIRVs |
| False Discovery Rate | (SIRV_transcripts - SIRV_RM)/SIRV_transcripts |
| False Detection Rate | FP/SIRV_transcripts |
| Redundancy | (FSM + ISM)/unique(TP+PTP) |

#### Simulated Data

The simulated data contained both transcript models based on the current GENCODE annotation and several simulated novel transcripts that will result in valid NIC and NNC annotations. Transcript models generated from simulated data were analyzed by SQANTI3, providing a GTF file that includes all simulated transcripts (GENCODE and novel) and excludes all transcripts for which reads were not simulated. The evaluation metrics for simulated data are shown in **Table 6**.

**Table 6:** Metrics and definitions for evaluation against simulated data.

|  |  |
| --- | --- |
| Metric | All simulated transcripts |
| True Positive (TP)<br>TP_ref | RM<br>RM to GENCODE mdels |
| TP_novel | RM to simulated novel transcript models |
| Partial True Positive (PTP)<br>PTP_ref<br>PTP_novel | ISM or FSM_non_RM<br>ISM or FSM_non_RM of GENCODE models<br>ISM or FSM_non_RM of simulated novel models |
| False Negative (FN)<br>FN_ref<br>FN_novel | Simulated transcripts without RM or PTP calls<br>Simulated GENCODE models without RM or PTP calls<br>Simulated novel models without RM or PTP calls |
| False Positive (FP) | NIC + NNC + antisense + fusion |
| Sensitivity<br>Sens_ref<br>Sens_novel | TP_ref/P(GENCODE)<br>TP_novel/P(Simulated novel) |
| Precision | $TP/(TP+PTP+FP)$ |
| Positive Detection Rate | $(TP+PTP)/P$ |
| False Discovery Rate | $(FP+PTP)/(TP+PTP+FP)$ |
| False Detection Rate | $FP/(TP+PTP+FP)$ |
| Redundancy | # FSM and ISM per simulated transcript model |

#### Comprehensive GENCODE annotation

Submitted transcript models were analyzed with SQANTI3 using the newly released GENCODE annotation (version 39 for human and M28 for mouse) and different metrics were obtained for FSM, ISM, NIC, NNC and Other models according to the scheme depicted below. Transcripts from new genes included in the latest annotation release were cataloged as “Intergenic” initially but considered FSM, ISM, NIC, or NNC with an updated GENCODE annotation. This will allow the evaluation of gene and transcript discovery on unannotated regions.

*High-confidence transcripts derived from LRGASP data* (Positives P are the set of all high-confidence transcripts)

Finally, a set of manually curated transcript models were used to estimate sensitivity and precision on real data. Metrics that will be applied in this transcript set are TP, PTP, and FN, Sensitivity, Positive Detection Rate, Redundancy, and %LRC (**Table 7**)

**Table 7:** Metrics for evaluation of manually annotated transcript models.

|  |  |
| --- | --- |
| TP | RM |
| PTP | ISM or FSM_not_RM |
| FN | Curated GENCODE transcripts without FSM or ISM |
| Sensitivity | TP_ref/Curated GENCODE transcripts |
| Positive Detection Rate | (TP+PTP)/Curated GENCODE transcripts |
| Redundancy | (FSM + ISM)/unique(TP+PTP) |

\* See **Table 2** for description of LRGASP metrics

#### Analysis of transcript model identification across pipelines

We will evaluate the characteristics of the transcripts detected as a function of the experimental factors of the LRGASP study, e.g., sequencing platform or library protocol. To do that, we will compare detected transcripts across pipelines at the level of Unique Intron Chain (UIC), allowing for variability in the 3’ and 5’ definition, and annotate the pipelines that detected each UIC. A barcode for each UIC indicates the type and number of pipelines where it was detected, together with general transcript properties. The fields of the UIC barcode are described in **Extended Data Table 16 UIC Barcode,** and the location of BED files of the UICs with barcodes can be found in **Supplementary Information, Data and code availability.**

The barcode enables interrogation of transcript characteristics associated with consistent detection by pipelines using specific types of data. For example, we can ask which transcript properties are associated with transcripts that tend to be pipeline-specific versus detected by most pipelines, or length differences between transcripts detected by most PacBio pipelines and not by Nanopore, or by dRNA and not by other library preparation methods. We will systematically screen transcript properties associated with the LRGASP experimental factors to identify biases.

Transcript models were visualized in the UCSC Genome Browser^51^ using the Track Hub facility^52^. The track hub displayed consolidated transcript models from the submissions with metadata, color-coding, and filtering by attributes. This allowed us to efficiently explore the significant quality of LRGASP results in the genomic context.

### Challenge 2 Evaluation: Transcript isoform quantification

We evaluated transcript isoform quantification performance with four data scenarios (real data with multiple replicates, cell mixing experiment, SIRV-set 4 data, and simulation data). We designed nine metrics for performance assessment both with and without known ground truth (**Table 4 and Fig. 3a**). The participants of the Challenge 2 were able to run these evaluations by submitting their quantification results at the website https://lrrna-seq-quantification.org/ that generates an interactive report in the HTML and PDF formats (See **Data and code availability**).

#### Ground truth is available

We evaluated how close the estimations and the ground truth values are by three metrics as follows.

Denote Θ̂ = (θ̂_1_,…,θ̂_*I*_)^*T*^ and Θ = (θ,…,θ_*I*_)^*T*^ as the estimation and ground truth of the abundance of transcript isoforms in a sample, respectively. Here, we use the **T**ranscripts **P**er **M**illion (**TPM**) as the unit of transcript abundance. Then, four metrics can be calculated by the following formulas.

##### Spearman Correlation Coefficient (*SCC*)

*SCC* evaluates the monotonic relationship between the estimation and the ground truth, which is based on the rank for transcript isoform abundance. It is calculated by

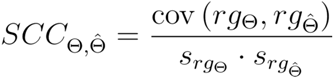

where *rg*Θ and *rg*Θ̂ are the ranks of Θ and Θ̂, respectively, and *cov* (*rg*Θ, *rg*Θ̂) is the covariance of the corresponding ranks, *s*_*rg*_Θ and *s*_*rg*_Θ̂ are the sample standard deviations of *rg*Θ and *rg*Θ̂, respectively.

##### Median Relative Difference (*MRD*)

*MRD* is the median of the relative difference of abundance estimates among all transcript isoforms within a sample, is calculated by

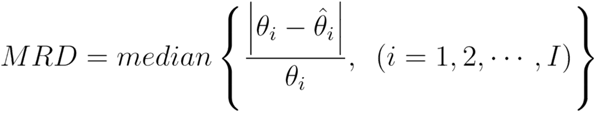

A small *MRD* value indicates the good performance of abundance estimation.

##### Normalized Root Mean Square Error (*NRMSE*)

*NRMSE* provides a measure of the extent to which the one-to-one relationship deviates from a linear pattern. It can be calculated by

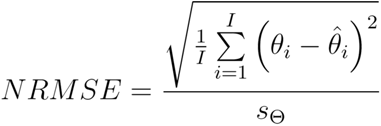

where *s*_Θ_ is the sample standard deviation of Θ.

A good performance of abundance estimation should have a small value of *NRMSE*.

In the case of LRGASP, the above metrics were calculated with the cell mixing experiment, simulation data and SIRVs.

##### Ground truth is unavailable

For multiple replicates under different conditions without the ground truth, we evaluated a quantification method by the “goodness” of its statistical properties, including irreproducibility, consistency, and resolution entropy that is also calculated for single sample data.

##### Irreproducibility

The irreproducibility statistic characterizes the average coefficient of variation of abundance estimates among different replicates (**Extended Data Fig. 55**), which is calculated by

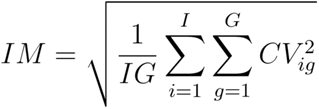

Here, *CV_ig_* is the coefficient of variation of *log* (θ̂_*igr*_ + 1) (*r* = 1,2,…,*R*), which is calculated by

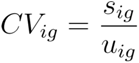

where *s_ig_* and *u_ig_* are the sample standard deviation and mean of abundance estimates, which are calculated by

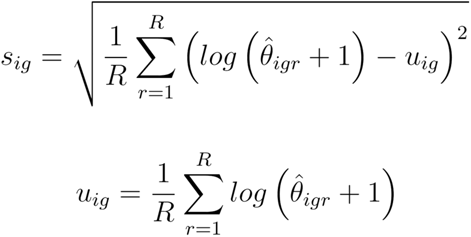

By plotting *CV_ig_* versus average abundance *u_ig_*, we examined how the coefficient of variation changes with respect to the abundance and the <u>a</u>rea under the <u>CV</u> <u>c</u>urve (ACVC) was calculated as a secondary statistic. With a small value of irreproducibility and ACVC scores, the method has high reproducibility.

##### Consistency

A good quantification method consistently characterizes abundance patterns in different replicates. Here, we propose a consistency measure *C*(α) to examine the similarity of abundance profiles between mutual pairs of replicates (**Extended Data Fig. 55**), which is defined as:

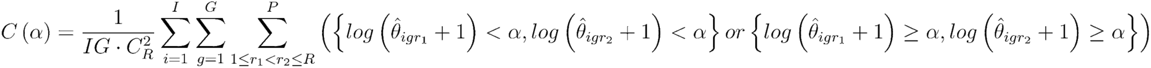

Where α is a customized threshold defining whether a transcript is expressed or not.

We plotted the abundance threshold α versus consistency measure *C*(α) to evaluate how *C*(α) changes with respect to the abundance threshold. The <u>a</u>rea under the *<u>C</u>*(α) <u>c</u>urve (ACC) can be used as the second metric to characterize the degree of similarity of transcript expression. With a large value of consistency and ACC scores, the method has a higher similarity of abundance estimates among multiple replicates.

##### Resolution Entropy (*RE*)

A good quantification method should have a high resolution of abundance values. For a given sample, a Resolution Entropy (*RE*) statistic characterizes the resolution of abundance estimation (**Extended Data Fig. 56**):

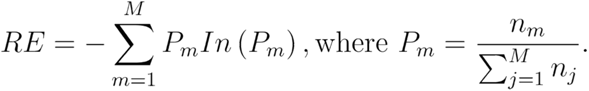

Here, the abundance estimates are binned into *M* groups, where *n_m_* represents the number of transcript isoforms with the abundance estimate Θ̂ ∈ [*m* · α, (*m* + 1) · α), and α = *max* (Θ̂)/*M*. *RE* = 0 if all transcript isoforms have the same estimated abundance values, while it obtains a large value when the estimates are uniformly distributed among *M* groups.

##### Evaluation with respect to multiple transcript features

Different transcript features, such as exon-isoform structure and the true abundance level could influence quantification performance. Thus, we also evaluated the quantification performance for different sets of genes/transcripts grouped by transcript features, including a number of isoforms, number of exons, ground truth abundance values, and a customized statistic K-value representing the complexity of exon-isoform structures.

##### K-value

Most methods for transcript isoform quantification assign sequencing coverage to isoforms; therefore, the exon-isoform structure of a gene is a key factor influencing quantification accuracy. Here, we used a statistic K-value (manuscript in preparation, **Extended Data Fig. 62**) to measure the complexity of exon-isoform structures for each gene. Suppose a gene of interest has *I* transcript isoforms and *E* exons and define *A* = (*a_ie_*, (*i* = 1,2,…, *I*; *e* = 1,2,…, *E*) as the exon-isoform binary matrix, where

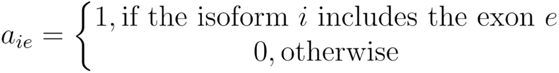

The K-value is the condition number of the exon-isoform binary matrix *A*, which is calculated by

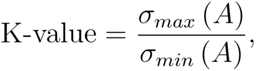

where σ_*max*_ (*A* and σ_*min*_ (*A* are maximum and minimum singular values of the matrix, respectively.

### Challenge 3 Evaluation: De novo transcript isoform detection without a high-quality genome

Challenge 3 evaluated the applicability of lrRNA-seq for *de novo* delineation of transcriptomes in non-model organisms to assess the capacity of technologies and analysis pipelines for both defining accurate transcript models and for correctly identifying the complexity of expressed transcripts at genomic loci when genome information is limited.

The challenge includes three types of datasets. The mouse ES transcriptome data (**Table 1**) was used to request the reconstruction of mouse transcripts without making use of the available genome or transcriptome resources for this species. Models were compared to the true annotations set with the same parameters as in Challenge 1. As FASTA rather than GTF files are provided in Challenge 3, we used the same mapper, minimap2, for all the submissions to transform sequence information into a genome annotation file. While this dataset allows for a quantitative evaluation of transcript predictions in Challenge 3, it might deliver unrealistic results if analysis pipelines were somehow biased by information derived from prior knowledge of the mouse genome. To avoid this problem, a second dataset was used corresponding to the Floridian manatee’s whole blood transcriptome (*Trichechus matatus*). An Illumina draft genome of this organism exists (https://www.ncbi.nlm.nih.gov/assembly/GCF_000243295.1/), and the LRGASP consortium has generated a long-read genome assembly to support transcript predictions for this species. Additionally, Illumina data has been generated for this challenge. Again, we will evaluate pipelines that obtain transcript models without genome annotation but with these draft genome sequences and without genome assembly data. Since no curated gene models exist for the manatee, Challenge 1 metrics cannot be applied. Instead, the evaluation of this dataset will involve a comparative assessment of the reconstructed transcriptomes and experimental validation. For comparative assessment, the following parameters will be calculated.

a. Total number of transcripts
b. Mapping rate of transcripts to the draft genomes (for pipelines not using genome data)
c. Length of the transcript models
d. Number of mono- and multi-exon transcripts
e. % of junctions with Illumina coverage
f. % of transcripts with Illumina coverage at all junctions
g. % of chained junctions supported by at least one 454 read.
h. % junctions and transcripts with non-canonical splicing
i. % of transcripts with predicted coding potential
j. Predicted RT switching incidence
k. Predicted intra-priming
l. Number of transcripts/loci
m. Benchmarking sets of universal single-copy orthologs (BUSCO)^35^ analysis:

i. Number of complete BUSCO genes detected by a single transcript
ii. Number of complete BUSCO genes detected by multiple transcripts
iii. Number of fragmented BUSCO genes detected
iv. Redundancy level for complete and fragmented BUSCO genes
v. Number of transcript models with a BUSCO hit

The BUSCO analysis will include the percentage of eutherian BUSCO genes (lineage *eutheria_odb10*) that were fully detected by a single transcript (complete single-copy) or by multiple transcript models (complete duplicated) and that was partially detected (fragmented). We do not expect a BUSCO-complete transcriptome recovery since only one tissue or cell type per organism was sequenced. We expect that good-performing pipelines will obtain longer transcripts, well supported by Illumina data, with a high mapping rate to the draft genomes, most of them coding, and with a higher number of complete BUSCO genes and Blast2GO annotation potential.

Finally, the manatee long reads data also contain spiked-in SIRVs, which will be used to compute performance metrics for Challenge 3 analysis settings, using the same type of metrics as described for Challenge 1.

## Experimental validation of transcript models

### Gene selection process

#### Challenge 1 - WTC11

In order to semi-systematically select isoforms for comparison in the validation experiments, we binned isoforms based on the frequency by which they were detected in certain pipeline parameters. Isoform test groups were defined based on their presence across various pipelines and library preparations. In general, during the isoform selection stage, we prioritized isoforms with expression higher than 10 TPM, wherever possible, and isoforms that contained distinguishable sequence regions. During the primer design process, we considered as “present” all isoforms in the GENCODE annotation and all isoform models submitted by all participants.

For the assessment of abundance measures between isoforms annotated by GENCODE as known versus novel (e.g., GENCODE-known, GENCODE-novel, GENCODE-rejected, the sum of all read counts across all WTC-11 datasets (e.g., cDNA-PacBio, cDNA-ONT, etc.) were summed.

We chose groups of novel isoforms (NIC or NNC) that were preferentially detected by pipelines using ONT versus PacBio platform, using the cDNA preparations, and a control group of isoforms frequently detected by all pipelines across both platforms. Additionally, we chose known (FSM) isoforms for all three of these groups.

### Semi-automated primer selection process with Primers-Juju

The aggregate of all transcript models from all pipelines underwent visualization in a UCSC Browser track hub ^52^ to design primers that target specific transcript features. The process identified uniquely mapping sub-segments of isoforms and selected flanking 5’ end and 3’ end regions for primer design via the “Highlight” function within the UCSC Browser ^51^. The system then recorded the genomic coordinates of the regions and transcript identifiers.

Primers-JuJu (**Supplementary Information**) processed the primer region specifications, obtained the DNA sequence for the predicted RNA, and employed Primer3 ^36^ for primer design. The primer pairs are evaluated for off-target genome and transcriptome hits using In-Silico PCR^53^. The resulting primer pairs then add to the track hub for visualization (see **Extended Data Table 17 f**or the list of primers).

### cDNA synthesis

Replicate 2 and 3 of the same WTC11 total RNA aliquots that were used as input for the sequencing runs were used for cDNA syToproximately 1.3 ug of total RNA from each replicate was converted to cDNA using the NEBNext Single Cell/Low Input cDNA Synthesis & Amplification Module kit, with 14 cycles of PCR being performed.

### PCR of targets, QC, and amplicon pooling

Aliquots of 2uL cDNA (∼0.45 ug) was used as template for PCR reactions in which isoform-specific or isoform-partially-specific primers were used for amplification. One round of PCRs was done using the Replicate 2 cDNA, and a second round was done using the Replicate 3 cDNA. A touch-down PCR method was employed with a total of 35 cycles.

Each PCR reaction was run in a 1% agarose e-gel with SYBR Safe, and the band sizes were recorded. We found excellent agreement between the predicted and experimentally measured product length, with more than 60% of the bands matching closely. The products were pooled in equal volumes, 3uL of each product, to create a pool of all amplicons derived from PCR of both Replicate 2 and 3, in batches (**Table 8, Extended Data Table 18: <u>Challenge 1 WTC11 Validation Batches</u>**). After quantification of the amplicon pool via Qubit, they were subjected to ONT and PacBio sequencing.

**Table 8:** WTC-11 Validation Batches.

| **Amplicon Pool** | **Pooled Amplicon Concentration (ng/uL)** | **Total Volume Extracted From Amplicons in Pool (uL)** | **Total Amount of DNA (ng) in Pools** |
| --- | --- | --- | --- |
| Batch_1 WTC 'POOL' | 71 | 51 | 3,621 |
| Batch_2 WTC_Rep_2 | 79.4 | 144 | 11,433.60 |
| Batch_2 WTC_Rep_3 | 70.2 | 144 | 10,108.80 |
| Batch_3 WTC_Rep_2 | 47.2 | 132 | 6,230.40 |
| Batch_3 WTC_Rep_3 | 73.8 | 132 | 9,741.60 |
| Batch_4 WTC_Rep_2 | 20.8 | 243 | 5,054.40 |
| Batch_4 WTC_Rep_3 | 64.4 | 243 | 15,649.20 |

Libraries for ONT sequencing were prepared using an SQK-LSK114 kit and 300ng of pooled cDNA library was loaded on an R10.4.1 flow cell and sequenced at 260bps/sec. The run was stopped after 21h with ∼5.6e6 reads with an N50 of 1.2Kbp. Nanopore data was basecalled with guppy version 6.2.11 with the high-accuracy configuration (dna_r10.4.1_e8.2_260bps_hac.cfg). The reads, aligned to the human genome assembly, were deposited in the Sequence Read Archive (SRA) under the accession number SRR23881262.

For PacBio sequencing, 492 ng of the cDNA was combined with 123 ng of the manatee cDNA (pooled in a 1:5 ratio manatee: WTC-11 sample, see description in the section below) and subsequently converted into a SMRTBell library. The library was sequenced on a PacBio Sequel II, generating HiFi reads. The reads, which were aligned to the human and manatee genome assemblies, have been deposited in the SRA under the accession numbers SRR24680098 and SRR24680099, respectively.

### Challenge 1: Targeted PCR validation of isoforms in WTC11

cDNA was synthesized from Replicate 1 and Replicate 3 of the WTC11 total RNA, and used as a template for two sets of PCR reactions, respectively. After targeted PCR, amplicons were analyzed via agarose electrophoresis, and sizes were estimated. We found that at least 60% of all targets produced a single band corresponding to the expected amplicon size, indicating a moderately high success rate. All amplicons were pooled and sequenced with ONT minIon and PacBio Sequel II. Amplicon reads were aligned to the genome as well as expected target sequences (a subset of the test transcript), and all targets with at least one gapless (<2 bp), high identity long-read alignment was considered validated.

### Analysis of the amplicon reads to determine the support of targets

Using minimap2 (version 2.24-r1122), we aligned the RT-PCR sequences to the human genome assembly, with the targeted transcripts serving as junction specifications. We aligned the expected amplicon sequences extracted from the genome to control for difficult-to-align cases. We aligned the ONT sequences with the ‘splice’ minimap2 preset while using the ‘splice:hq’ preset for the PacBio and control sequences.

We aligned the WTC11 RT-PCR sequences to the GRC3h38 assembly, which includes SIRVs and an EBV sequence provided to LRGASP participants (syn25683364). In addition to genomic alignments, we aligned the RT-PCR sequences to a reference composed of the predicted amplicon sequences. Furthermore, we aligned the amplicon alignments to the amplicon reference. This approach facilitated the identification of difficult-to-align amplicons to the genome and cases where alignments to different isoforms might not be detected. For ONT sequences, we used the minimap2 ‘map-ont’ preset for amplicon reference alignments and the ‘map-pb’ preset for the PacBio and control alignments.

We evaluated each data class (ONT, PacBio, or control) by examining the counts of supporting reads for each amplicon on the genomic and amplicon reference alignments. To validate intron chains in the genomic alignments, we ensured the read alignment had the same intron chain as the targeted amplicon. We used minimap2 to identify introns in reads, rejecting those with adjacent indels.

For the amplicon reference alignments, we evaluated two metrics: Indel similarity and the maximum number of indels. The similarity metric is length-independent, while the absolute difference can distinguish subtle differences such as NAGNAG split sites. We filtered the intron chain results for only those reads with no more than two bases difference in indels.

We gathered these statistics in a table (**Extended Data Table 19**) and manually classified them as ‘supported,’ ‘likely,’ or ‘unsupported.’ In cases where there were low read counts or conflicted data (generally less than 50), we examined them using IGV^54^.

One confounding issue when using RT-PCR for validation is that the primers crossing introns may result in the ends of some amplicons not aligning across the introns. In such cases, the ends would sometimes align into the intron with a similar sequence while the remainder of the amplicon was soft-clipped. The control alignments of the amplicons are a good indication of this issue, and targets exhibiting the unaligned end regions could be classified as supported based on other evidence.

An interesting case was the ALG6 WTC11 target. Here minimap2 forced a rare but annotated GT-AT U12 intron into a GT-AG intron and genomic deletion, leading to none of the pipelines correctly identifying the isoforms of this gene containing this intron (**Extended Data Fig. 71: ALG6 alignment**).

To estimate the counts across WTC11 datasets for each experimentally tested transcript, long reads derived from different combinations of library preparation and sequencing methods were first mapped to hg38 using minimap2 (PacBio: minimap2 -ax splice:hq -uf --MD -t 40; ONT: minimap2 -ax splice --MD -t 30; ONT direct RNA sequencing: minimap2 -ax splice -uf -k14 –MD -t 30). Primary read alignments were converted from BAM to GTF format to extract the unique intron chains (UIC) for each read. Then, read counts matching the UIC of a given transcript were summed across all WTC11 datasets. The number of WTC11 datasets where at least one read supported the corresponding UIC was computed, with R2C2 samples with and without size selection being treated independently.

#### Challenge 3: Targeted PCR validation of isoforms in manatee

Since the manatee transcriptome is not annotated, we employed *ab initio* gene finding and transcript annotation. Genes were predicted from the manatee genome assembled as part of this LRGASP project and the program GeneMark ^55^. BUSCO^35^ analysis was used to evaluate the completeness of the transcript assembly and for annotation of the proteins represented from translations of the transcript sequences, thus quantifying the coverage and completeness of the ORFs.

We targeted a small set of relevant genes for the immune system. To validate our approach, we selected two genes for which a clear single isoform was consistently identified across pipelines: Secreted phosphoprotein 1 (*SPP1*) and Granzyme K (*GZMK*). Secreted phosphoprotein 1 is involved in immune regulation and tumor progression^56,57^, and *GZMK* modulates the proinflammatory immune cytokine response ^58^.

Next, we selected six genes that were present in the BUSCO database, showed variability in the number of isoforms predicted by each pipeline, and had a role in the immune response: interleukin 2 (*IL-2*), interleukin 7 (*IL-7*), interleukin 1 beta (*IL-1B*), Pentraxin 3 (*PTX3*), transporter 1, ATP binding cassette subfamily B member (*TAP1*) and TNF alpha-induced protein 3 (*TNFAIP3*). IL2 cytokine that regulates survival, proliferation and differentiation of T-cells ^58,59^. *IL7* is important for B and T cell development. *IL1B* is an inflammatory cytokine, an amplifier of an immune response, involved in cell proliferation, differentiation, and apoptosis ^60^. *PTX3* regulates complement activation of the immune system in the innate immune system and is in the same family of proteins as C-reactive protein ^61^. *TNFAIP3* negative regulation of inflammation and immunity, targeted for drug development ^56^. *IL2, PTX3*, *TNFAIP3* had 1 or 3 isoforms predicted by pipelines using only ONT data. *IL7* had three isoforms predicted by ONT and 3 by PacBio pipelines. While *TAP1* had four isoforms predicted only by ONT pipelines. *IL1B* had 11 isoforms predicted by ONT data and one by PacBio.

For manual target selection, a similar protocol employed for Challenge 1 targets, using Primers-Juju, was used to select regions of isoforms with unique junction chains that could be confirmed by generation of a PCR amplicon product. Whenever possible, the full span of the isoform, up to ∼2 kb, was selected. In some regions, multiple primer sets were designed.

Aliquots of the original nine individual manatee RNA samples used in Challenge 3 were stored at −80°C until the validation stage (**Extended Data Table 20**). RNA quality was re-verified using BioAnalyzer PicoChip for mRNA (Agilent, Santa Clara, CA). Approximately 400 ng of RNA from each manatee sample was pooled to prepare cDNA. cDNA was synthesized using Maxima H minus First Strand cDNA Synthesis kit (Thermo Scientific, Waltman, MA). Following the manufacturer’s instructions, we used a combination of oligo dT and random hexamer primers for the cDNA synthesis. Controls lacking RT enzyme and controls lacking template were prepared in tandem with test samples.

#### Primer selection and RT-PCR

In the case of the manatee, 2 or 4 primer sets were designed for each gene of interest. The process used for primer design was similar to that used for Challenge 1, using a semi-automated Primers-Juju approach (see **Extended Data Table 21** for the list of primers).

Manatee PCR was performed using KAPA HiFi HotStart Ready Mix (Roche, Switzerland) due to its high sensitivity and low error rate. Approximately 0.01 ng of cDNA was used as a template for individual PCR reactions. PCR protocol was also a touchdown approach with an initial annealing temperature of 70°C for 15 seconds, with a reduction of this temperature 1°C per cycle during 12 cycles. The second amplification phase was carried out for 21 cycles and 2 minutes of extension. When a PCR fragment larger than 1500 pb was expected, another PCR was run for that primer set, including 25 cycles and 5 min of extension. PCR products were quantified and sized using Agilent Bioanalyzer7000 DNA chip (Agilent, Santa Clara, CA).

The obtained PCR products were cleaned using QIAquick PCR purification kit (QIAgen, Germany). All PCR products were pooled as an equimolar pool for PacBio sequencing. We prepared the equimolar solution based on the BioAnalyzer molarity quantification for the band corresponding to the intended PCR product. Additionally, 25 µl of each PCR product that did not show a quantifiable band on the BioAnalyzer were added to the final sample equimolar pool for sequencing.

Analysis of the long-read amplicon reads was done in the same manner as for Challenge 1. The manatee RT_PCR sequences were aligned to the pre-submission manatee genome assembly used in LRGASP (GenBank accession JARVKP000000000.1). The resulting statistics were gathered (**Extended Data Table 22)** for manual analysis.

## Supporting information

Supplementary Data Report

## Data and code availability

An overview and documentation about the LRGASP Consortium can be found at https://www.gencodegenes.org/pages/LRGASP/, All data and code is available and how to access it is described in the **Supplementary Information** document.

## Acknowledgments

We thank Lexogen, Oxford Nanopore Technologies (ONT), and Pacific Biosciences for helpful discussions. ONT provided partial support of flow cells and reagents. We thank Xingjie Ren and Yin Shen for providing WTC11 cells, Takayo Sasaki and Dave Gilbert for providing the F121-9 hybrid mouse ES cells, and Alyssa Cousineau, Krishna Mohan Parsi, and Rene Maehr for providing human H1-hESC and H1-DE cells. We also thank Mark Akeson and Miten Jain for providing resources and technical advice for Nanopore sequencing. We thank Julia Visser for contributing artwork that gives an overview of the LRGASP Consortium. The project is supported by the following grants: Pew Charitable Trust (A.N.B.), NIGMS R35GM138122(A.N.B.), NIGMS R35GM142647 (G.M.S.), NHGRI U41HG007234 (J.L, M.D., R.G., S.C-S, J.E.L., J.M.G, T.H., I.B., A.E.B., J.M.M., and A.F.), NHGRI F31HG010999 (A.D.T.), and UM1 HG009443 (A.M. and B.W.), NHGRI R01HG008759, R01HG011469, and R01GM136886 (K.F.A., D.W., and H.L.); an institutional fund from the Department of Biomedical Informatics, The Ohio State University (K.F.A., D.W., and H.L.), an institutional fund from the Department of Computational Medicine and Bioinformatics, University of Michigan (K.F.A., D.W., and H.L.), SPBU 73023672 (A.P). Wellcome Trust [WT108749/Z/15/Z], and European Molecular Biology Laboratory (A.F.). We acknowledge Michael T. Walsh (University of Florida and Ellie Schiller (Homosassa Springs Park) for providing archive Lorelei blood samples. We acknowledge the support of the Spanish Ministry of Science and Innovation to the EMBL partnership, Centro de Excelencia Severo Ochoa, and CERCA Programme / Generalitat de Catalunya and the support of the German Federal Ministry of Education and Research with the grant 161L0242A The content of this manuscript is solely the responsibility of the authors and does not necessarily represent the official views of the National Institutes of Health. Any use of trade, firm, or product names is for descriptive purposes only and does not imply endorsement by the United States Government.

---

**Extended Data Fig.1.**
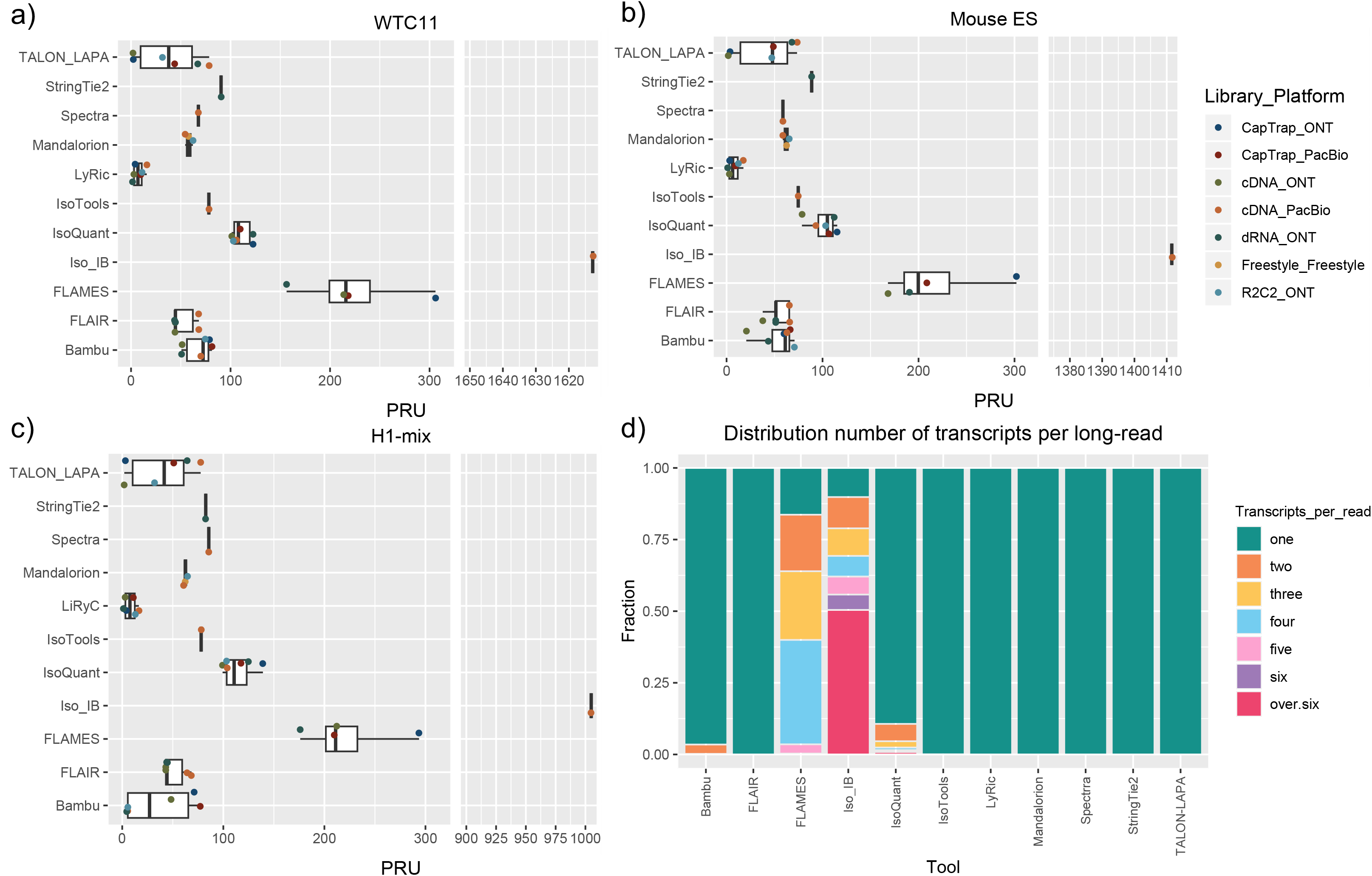
Read usage by analysis tool. a-c) The Percentage of Reads Used (PRU) is calculated as the fraction between the number of reads in transcript models provided in the submission of each pipelines and the number of available reads in the dataset. Values > 100 indicate the same read is assigned to more than one transcript model. Values < 100 indicate that not all available reads were used to predict transcript models. d) Distribution of the number of transcripts assigned to each long-read in the submitted reads2transcripts files. Values are aggregated for all submissions of the same tool.

**Extended Data Fig. 2.**
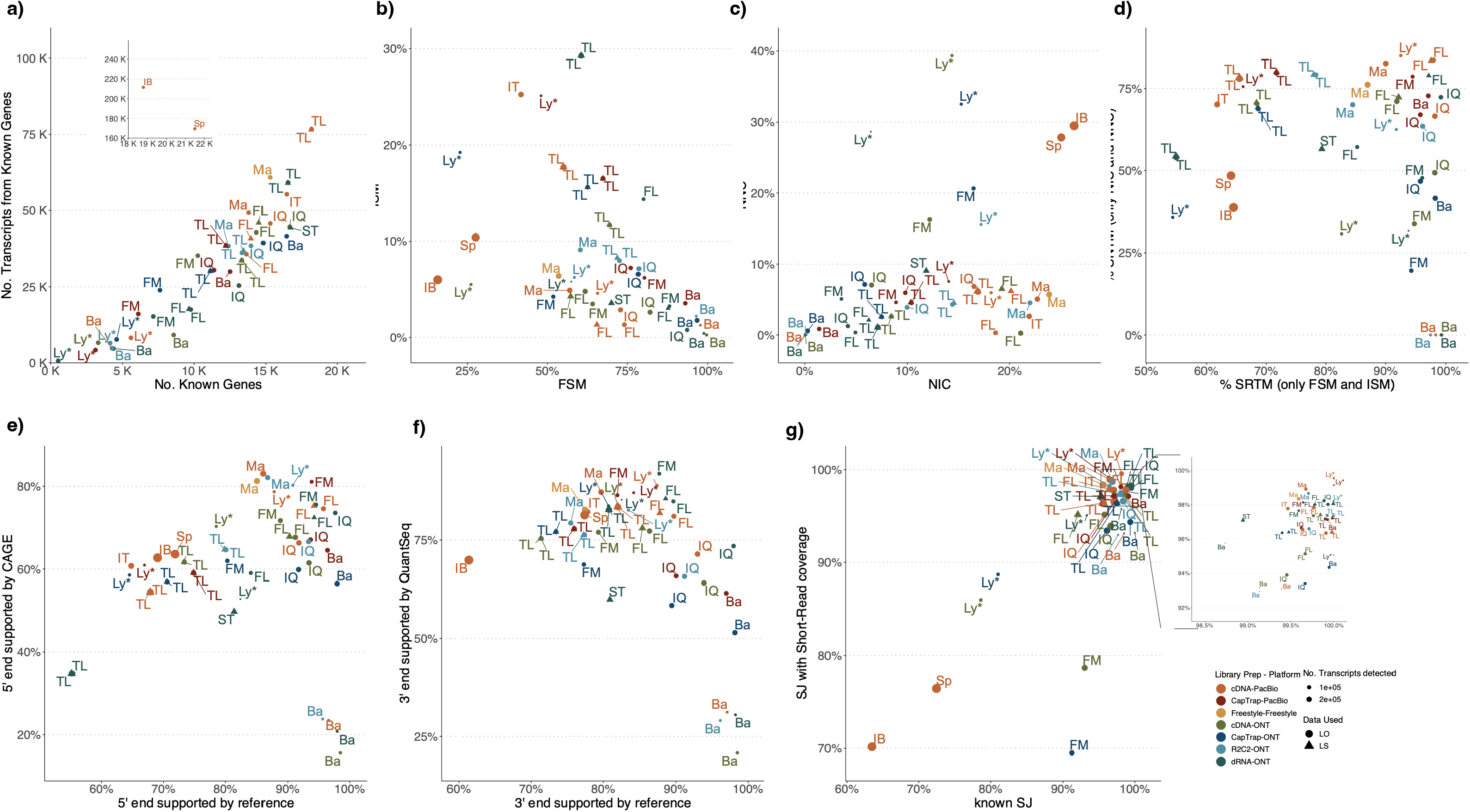
SQANTI3 evaluation of LRGASP submissions of the H1-mix dataset. Labels correspond to analysis tools and the color code indicates the combination of library preparation and sequencing platform. a) Number of gene and transcript detections. b) Number of Full Splice Match and Incomplete Splice Match transcripts. c) Number of Novel in Catalogue and Novel Not in Catalogue transcripts. d) Number of known and novel transcripts with full support at

**Extended Data Fig. 3.**
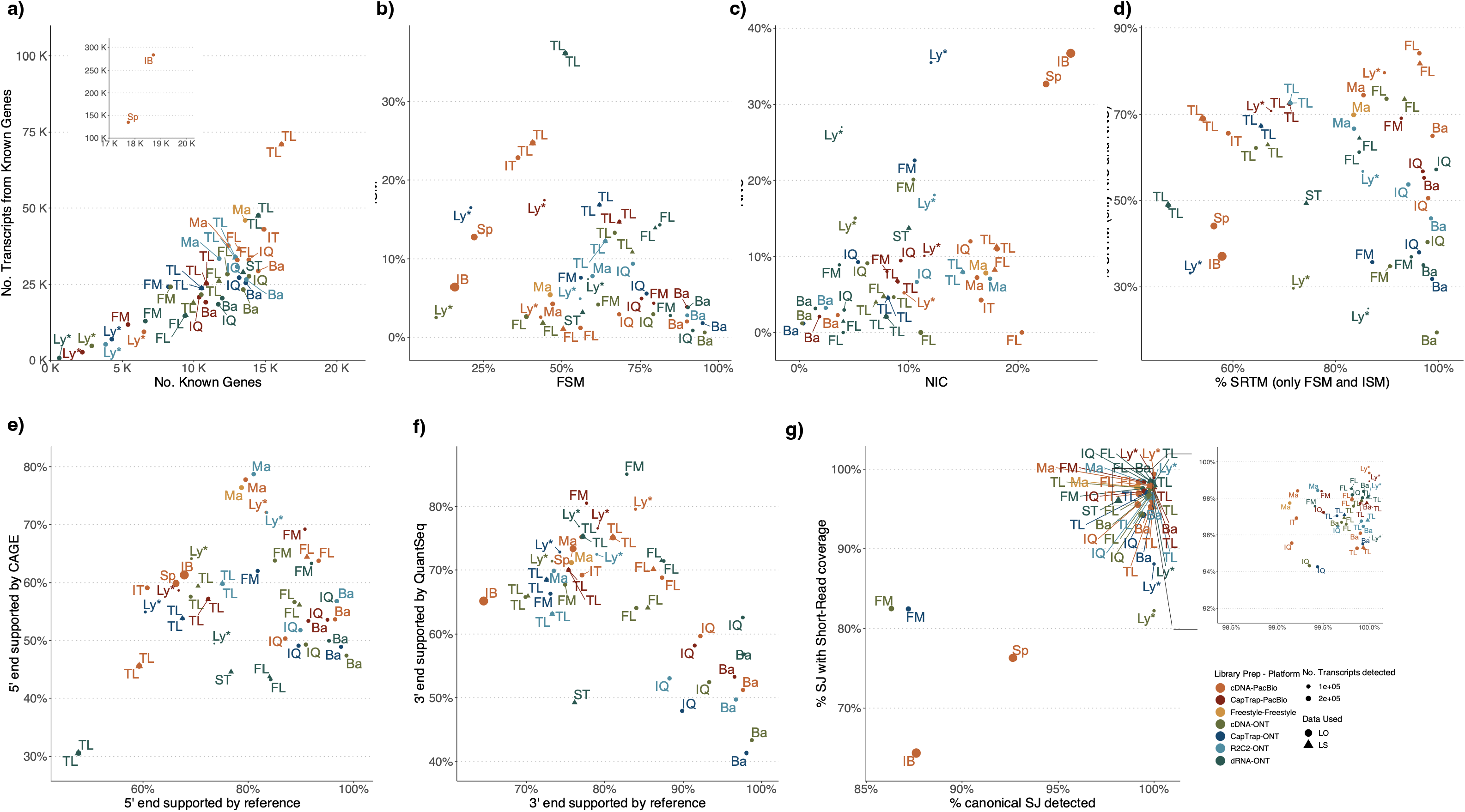
SQANTI3 evaluation of LRGASP submissions of the mouse ES dataset. Labels correspond to analysis tools and the color code indicates the combination of library preparation and sequencing platform. a) Number of gene and transcript detections. b) Number of Full Splice Match and Incomplete Splice Match transcripts. c) Number of Novel in Catalogue and Novel Not in Catalogue transcripts. d) Number of known and novel transcripts with full support at

**Extended Data Fig. 4.**
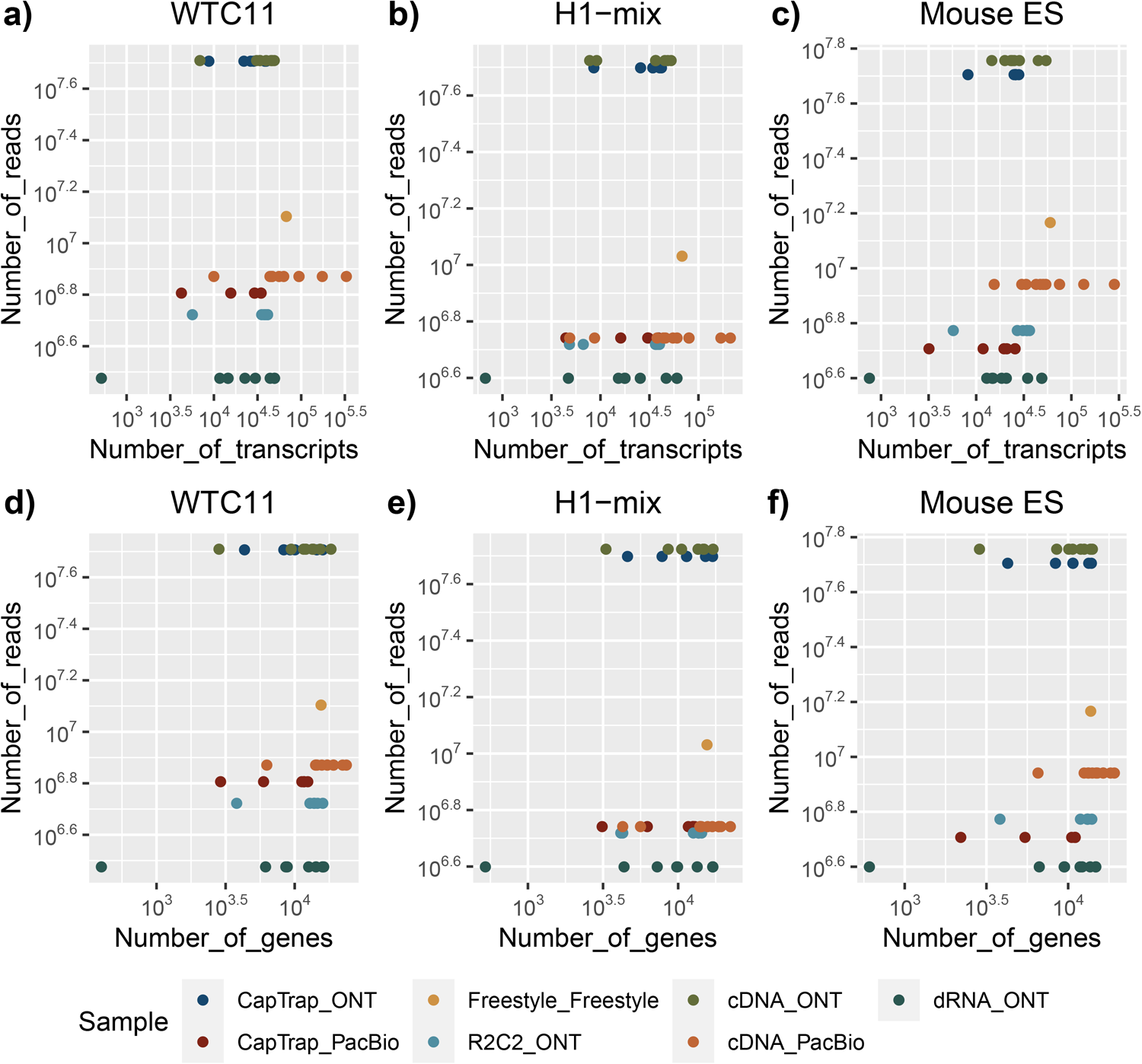
Relationship between sequencing depth and number of detected features. a−c) Transcripts, d−f) Genes.

**Extended Data Fig. 5.**
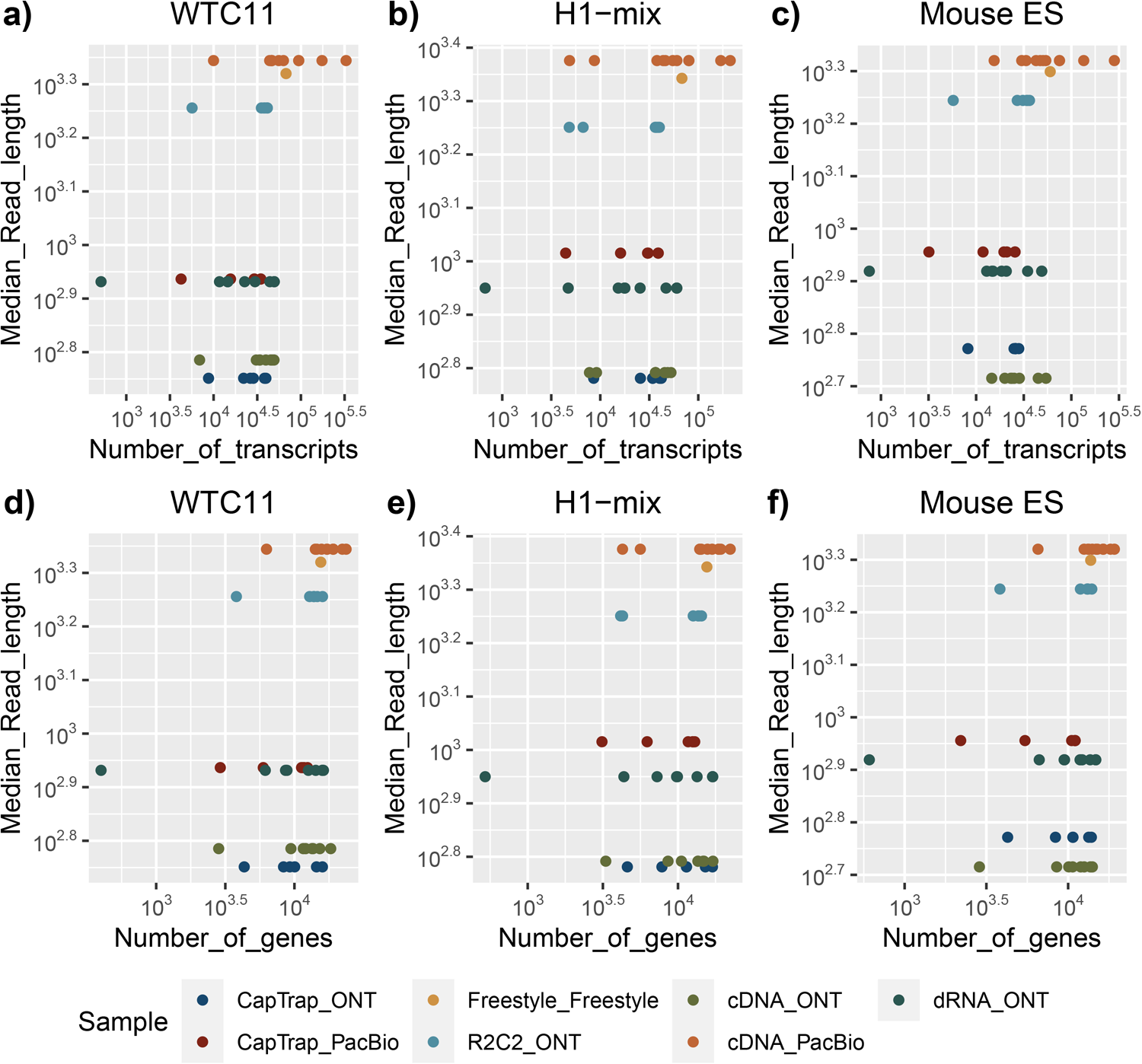
Relationship between read length and number of detected features. a−c) Transcripts, d−f) Genes.

**Extended Data Fig. 6.**
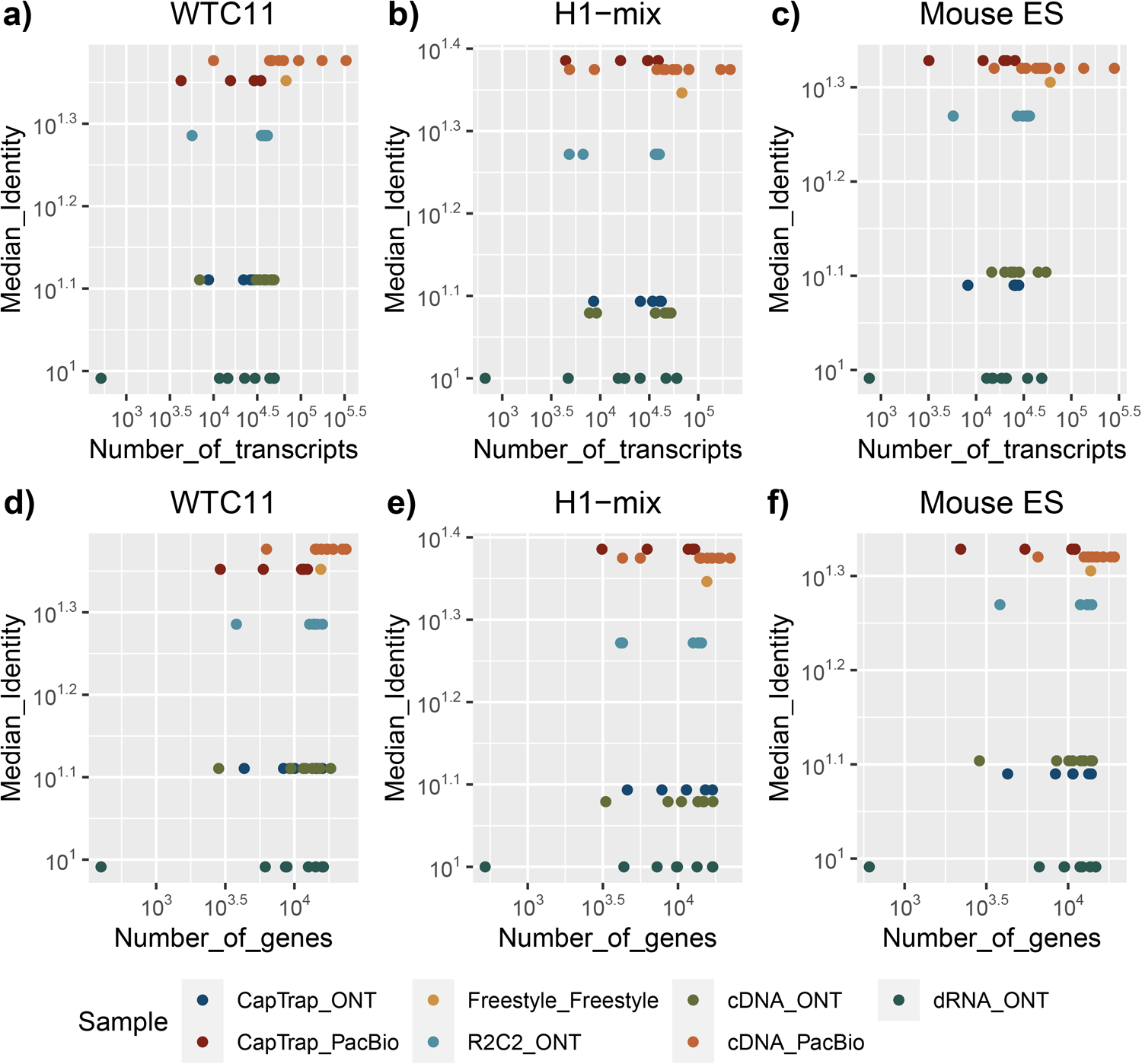
Relationship between read quality and number of detected features. a−c) Transcripts, d−f) Genes.

**Extended Data Fig. 7.**
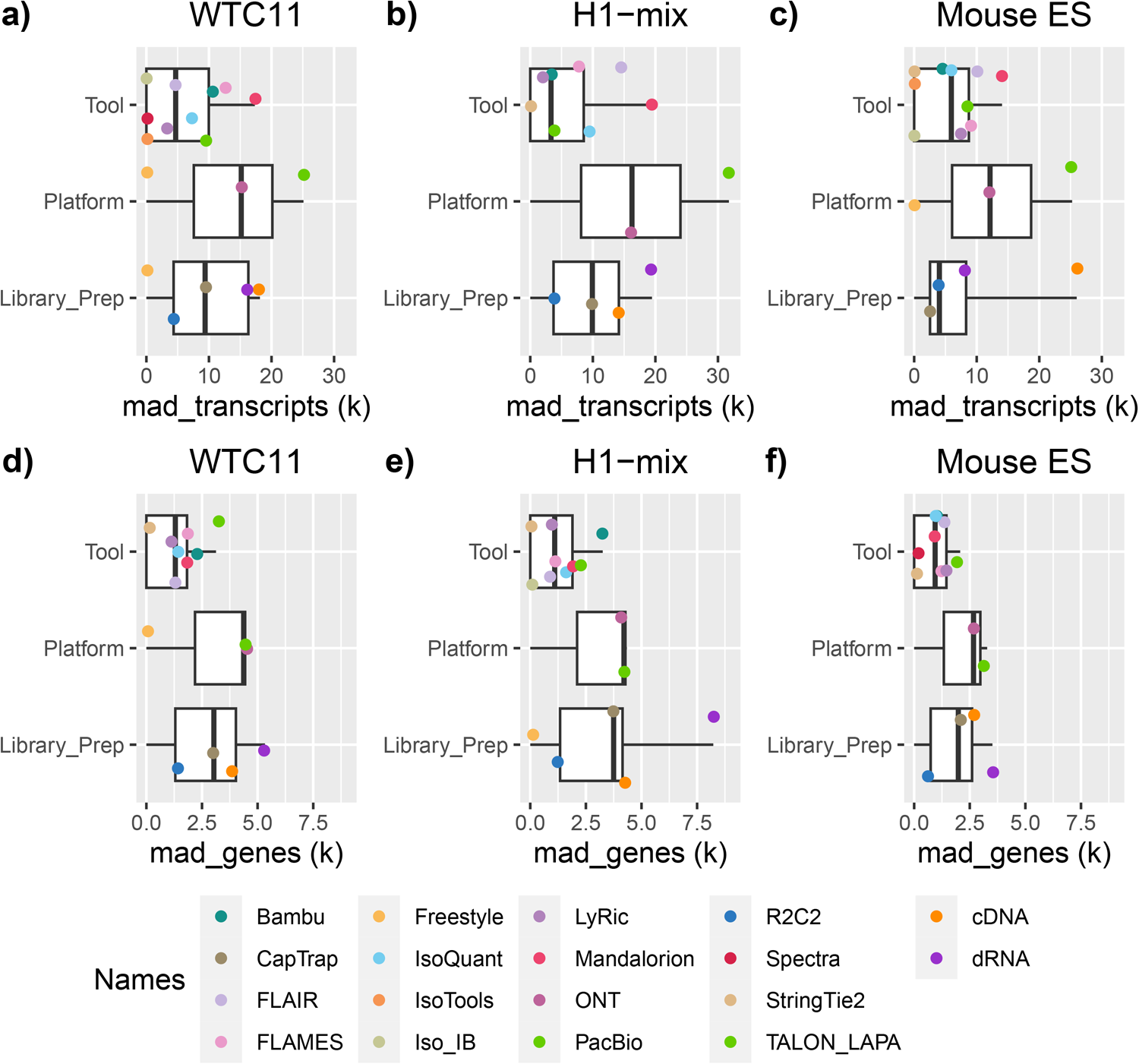
Median Absolute Deviance of detected features by experimental factor. a−c) Transcripts, d−f) Genes.

**Extended Data Fig. 8.**
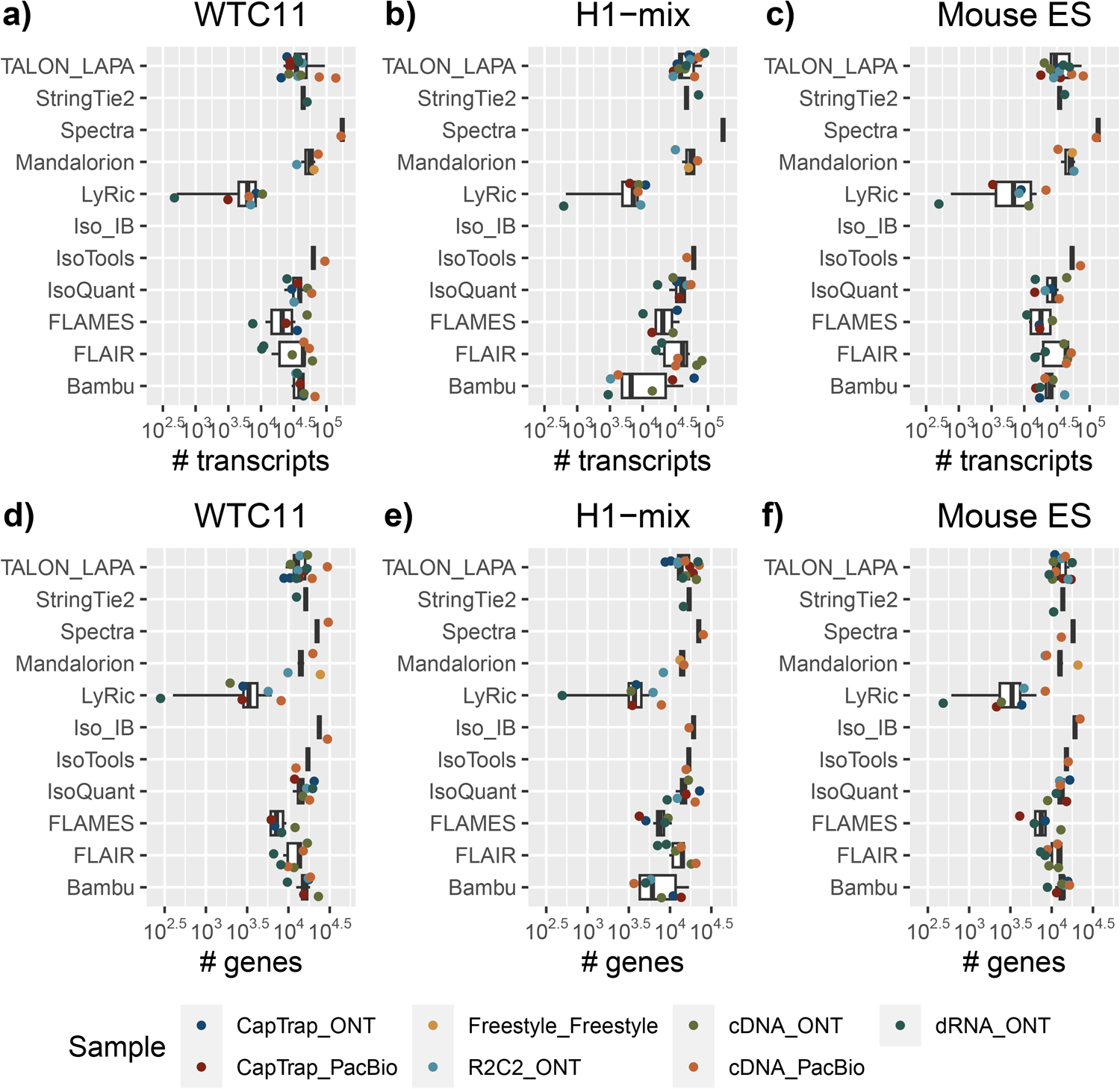
Number of detected transcripts and genes per analysis tool. a−c) Transcripts, d−f) Genes.

**Extended Data Fig. 9.**
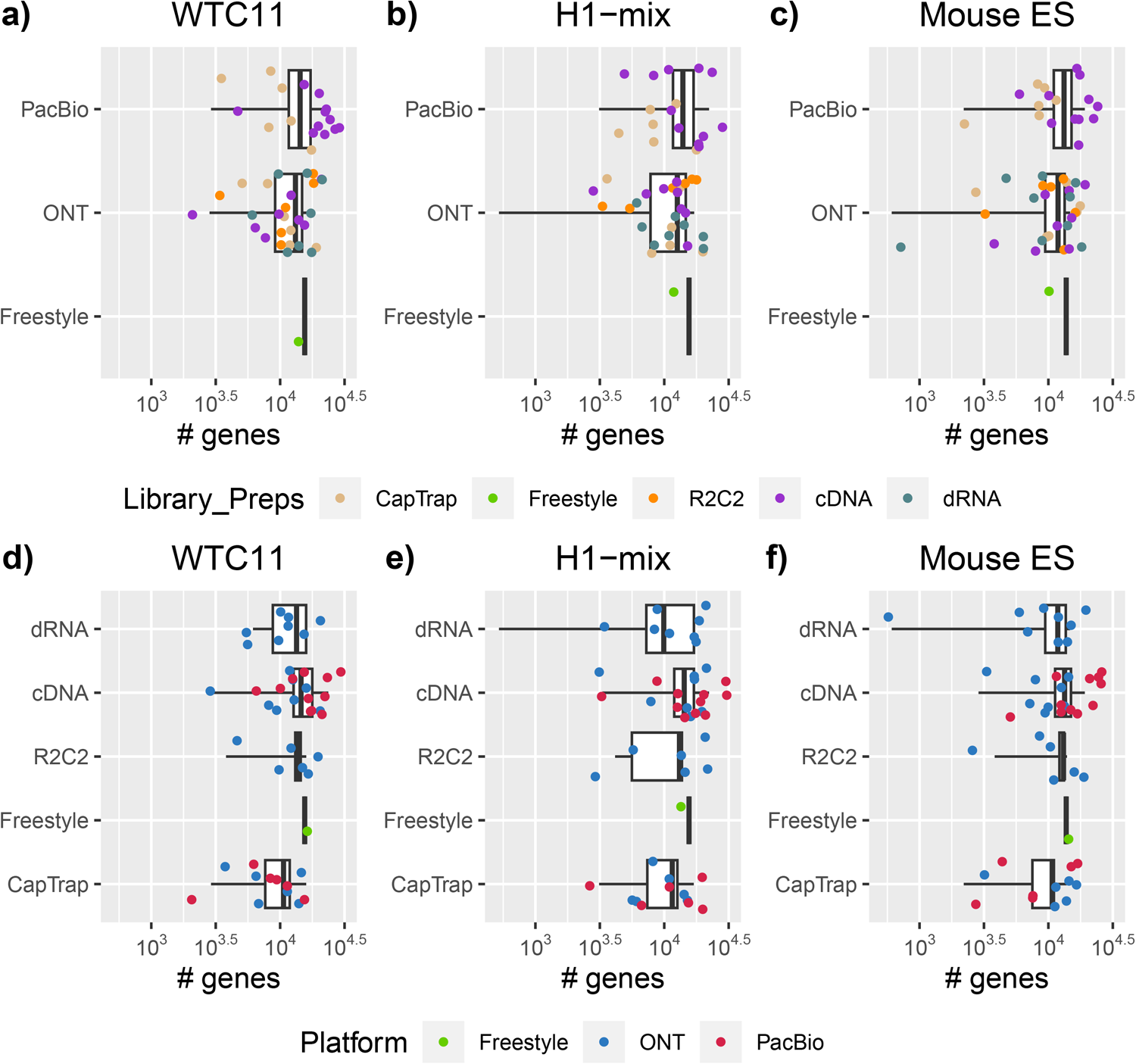
Number of detected genes per Platform and Library Preparation. a−c) Platform, d−f) Library Preparation.

**Extended Data Fig. 10.**
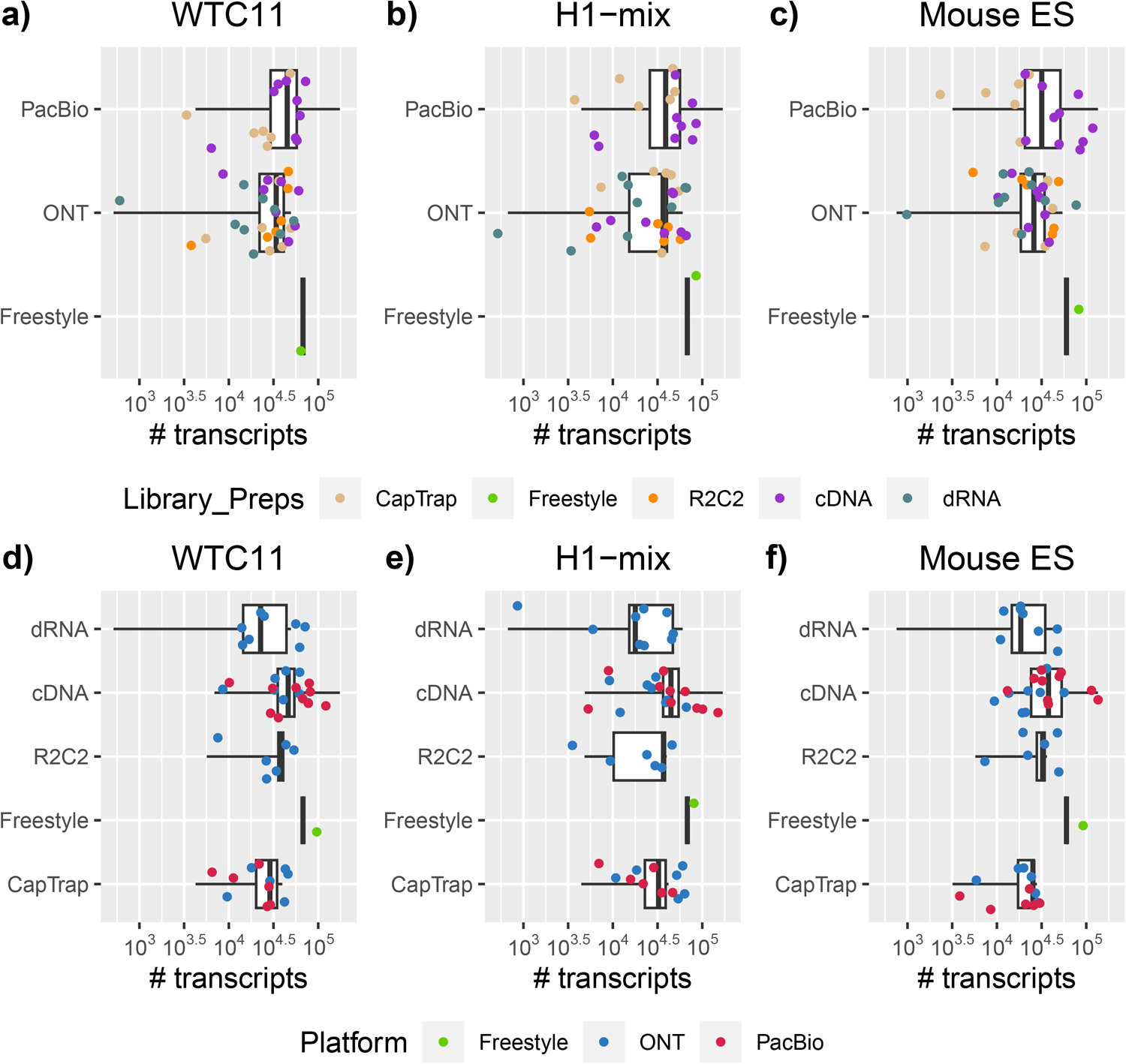
Number of detected transcripts per Platform and Library Preparation. a−c) Platform, d−f) Library Preparation.

**Extended Data Fig. 11.**
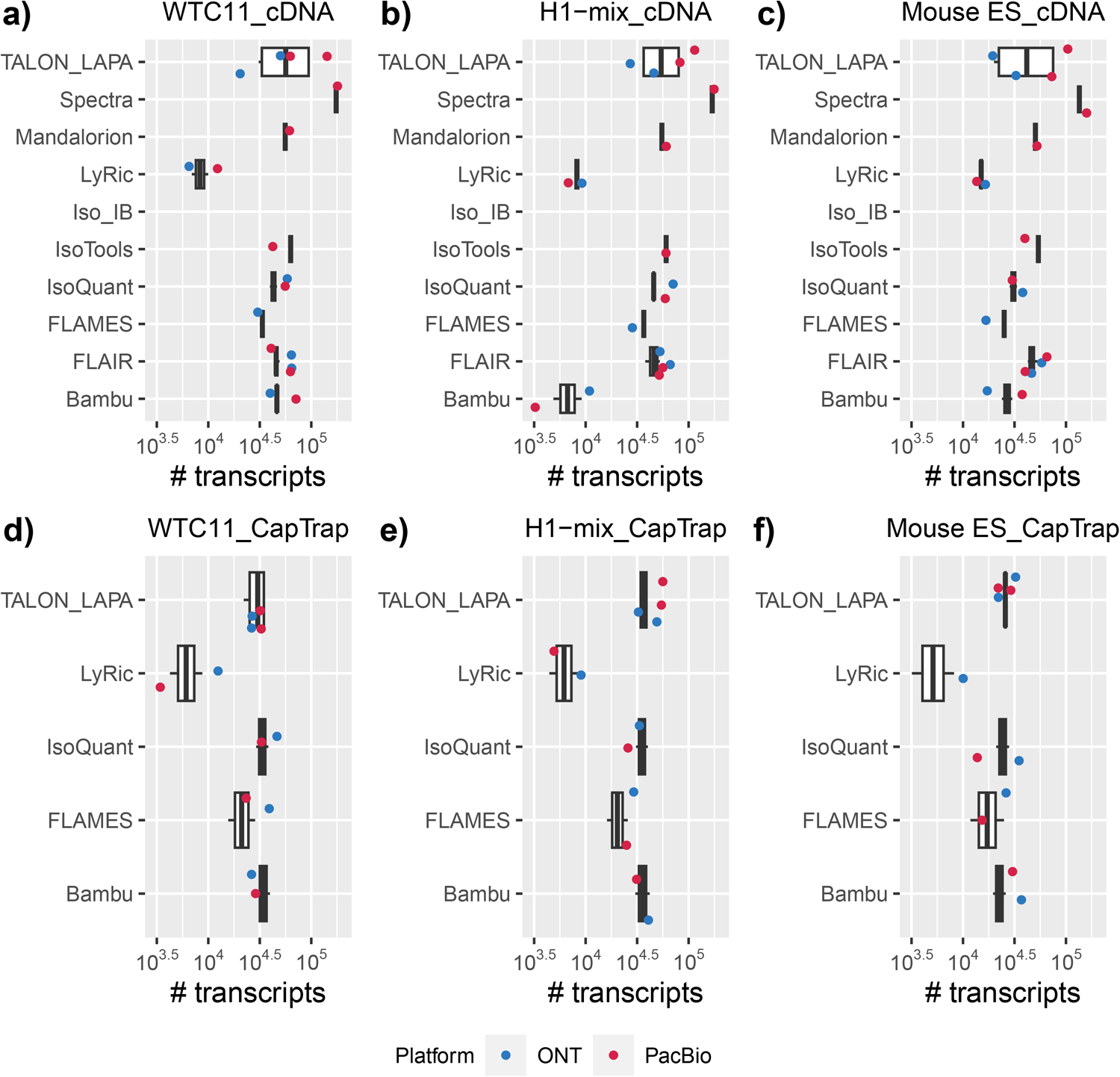
Number of detected transcripts in cDNA and CapTrap libraries. a−c) cDNA, d−f) CapTrap.

**Extended Data Fig. 12.**
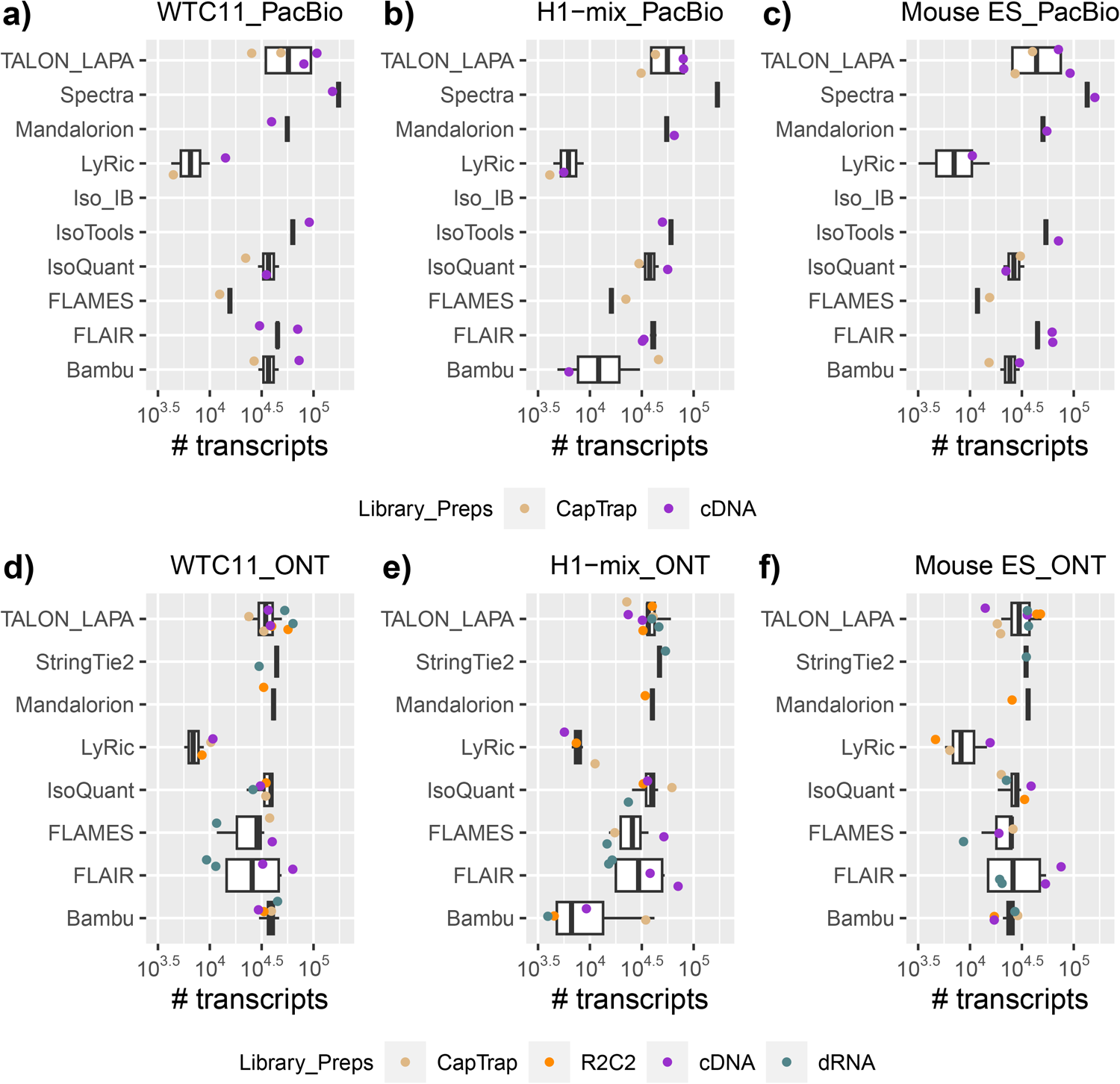
Number of detected transcripts in PacBio and Nanopore platforms. a−c) PacBio, d−f) Nanopore.

**Extended Data Fig. 13.**
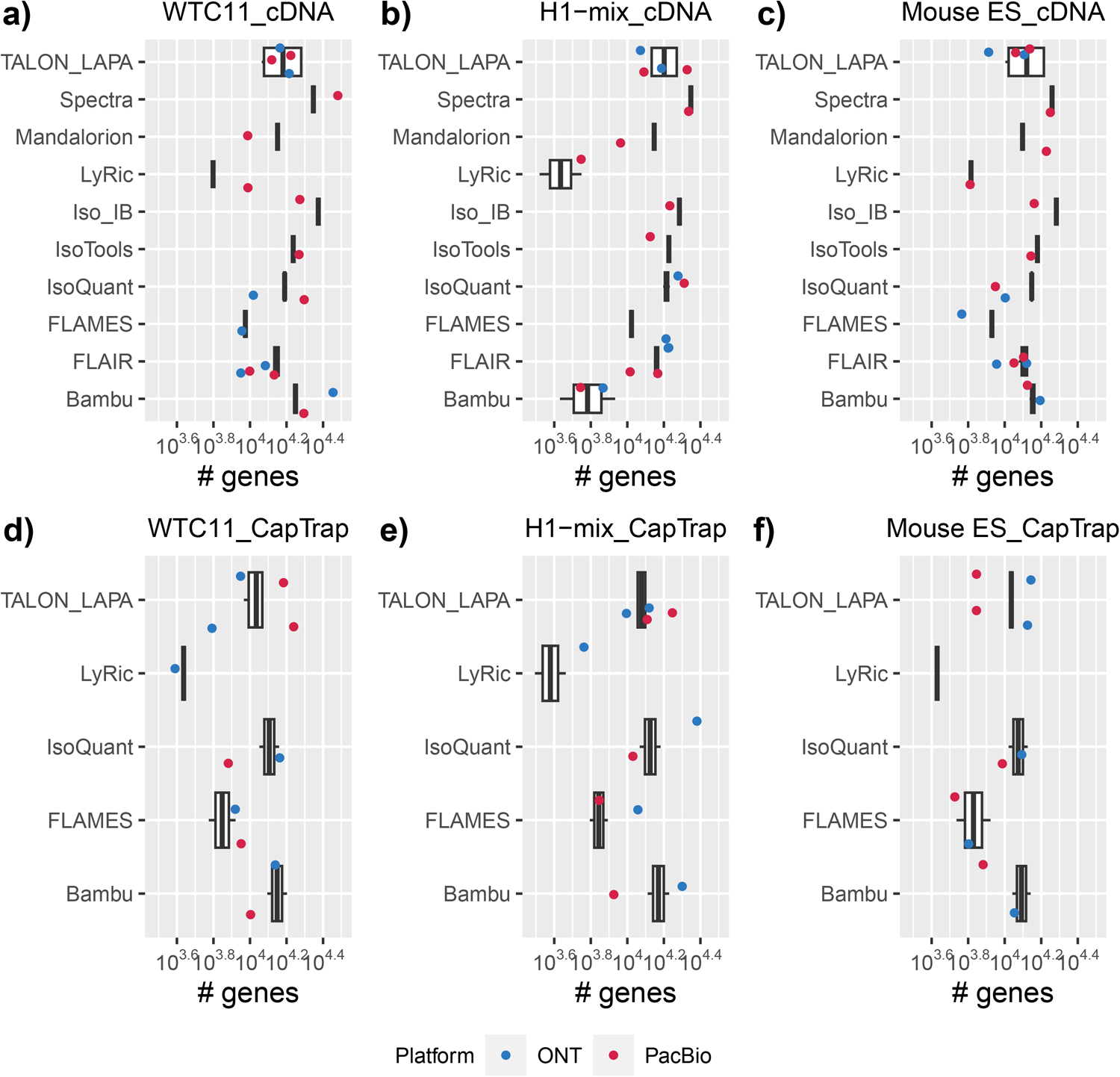
Number of detected genes in cDNA and CapTrap libraries. a−c) cDNA, d−f) CapTrap.

**Extended Data Fig. 14.**
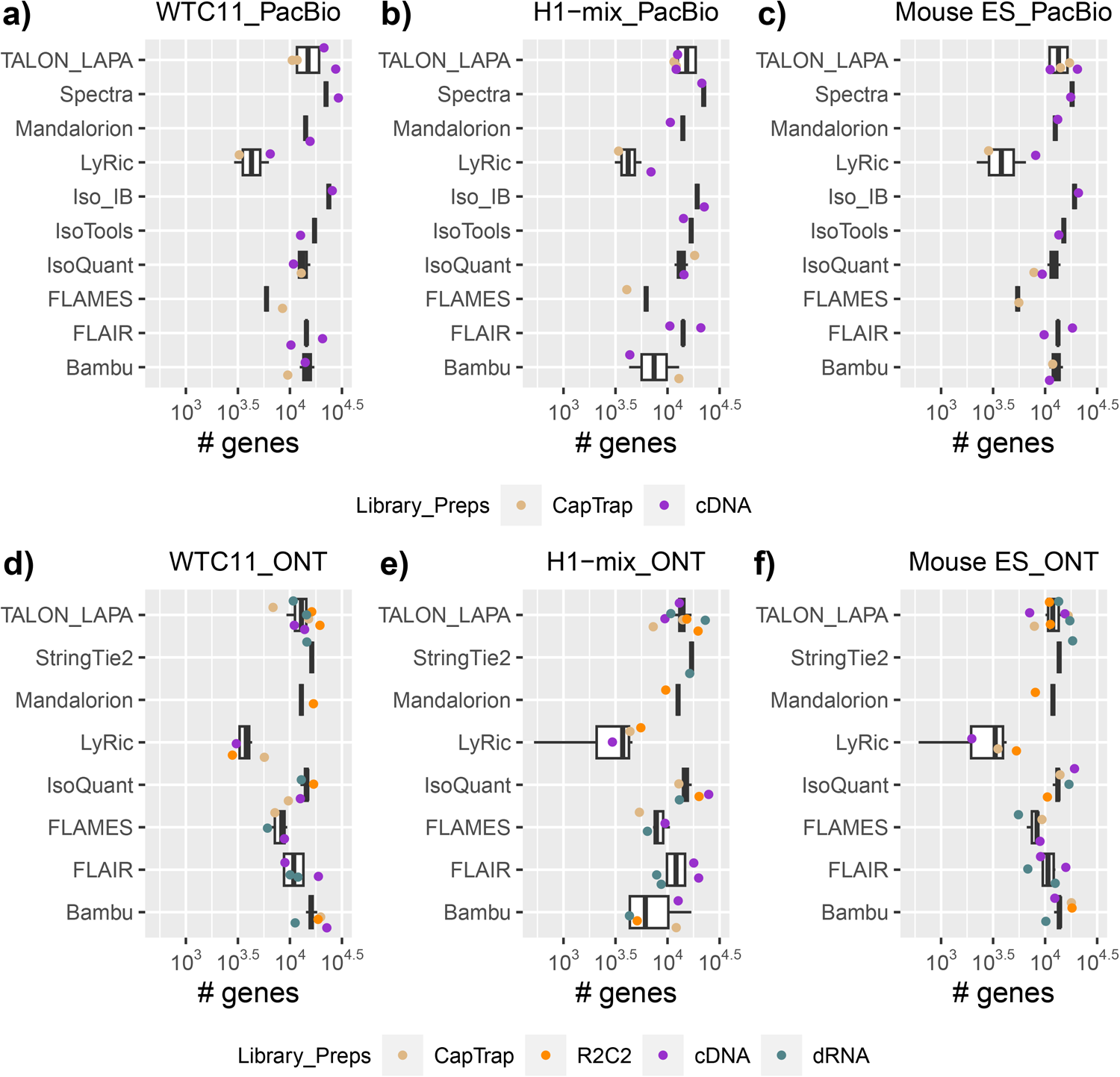
Number of detected genes in PacBio and Nanopore platforms. a−c) PacBio, d−f) Nanopore.

**Extended Data Fig. 15.**
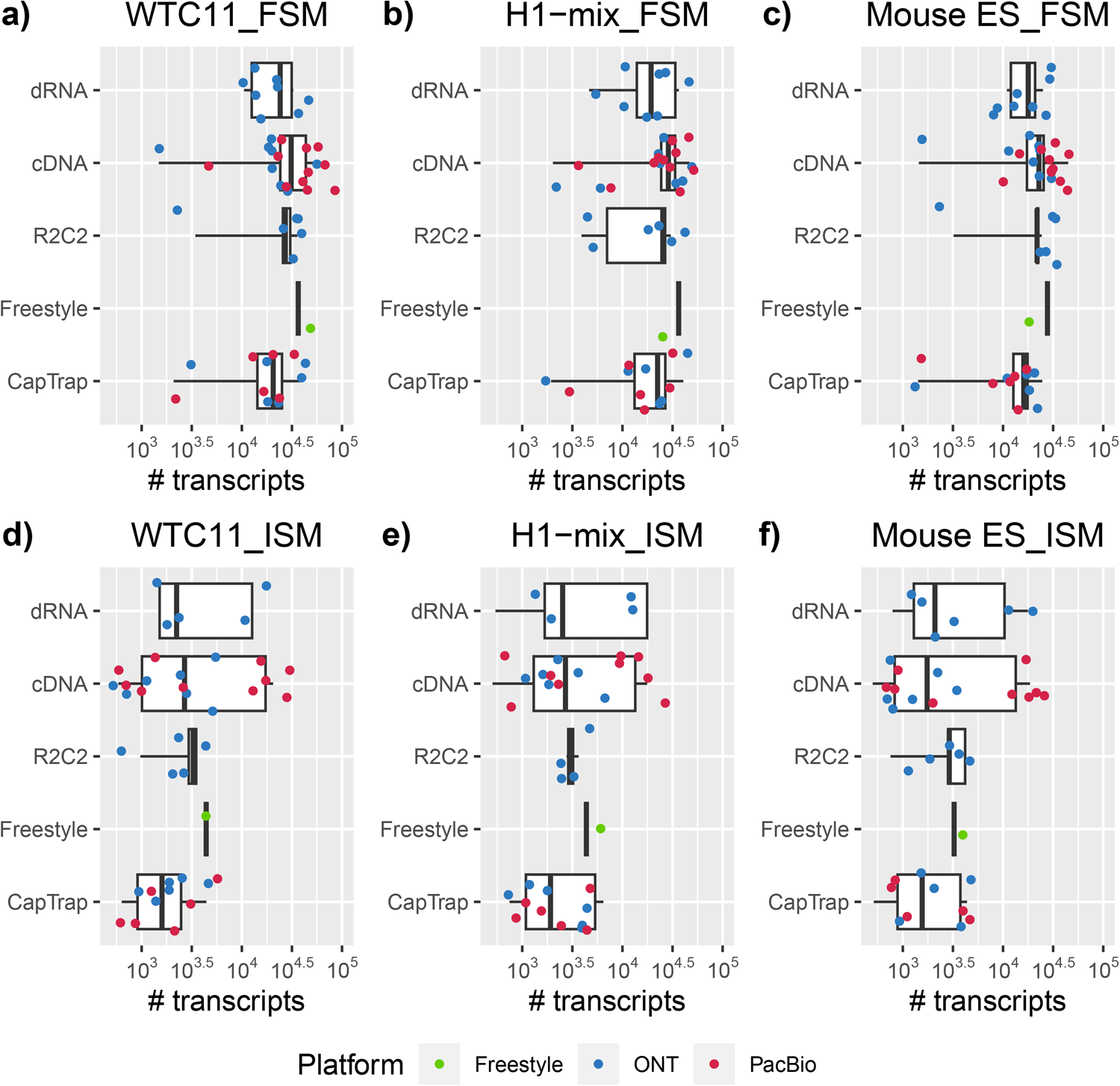
Number of FSM and ISM by sequencing platform and library preparation. a−c) FSM, d−f) ISM.

**Extended Data Fig. 16.**
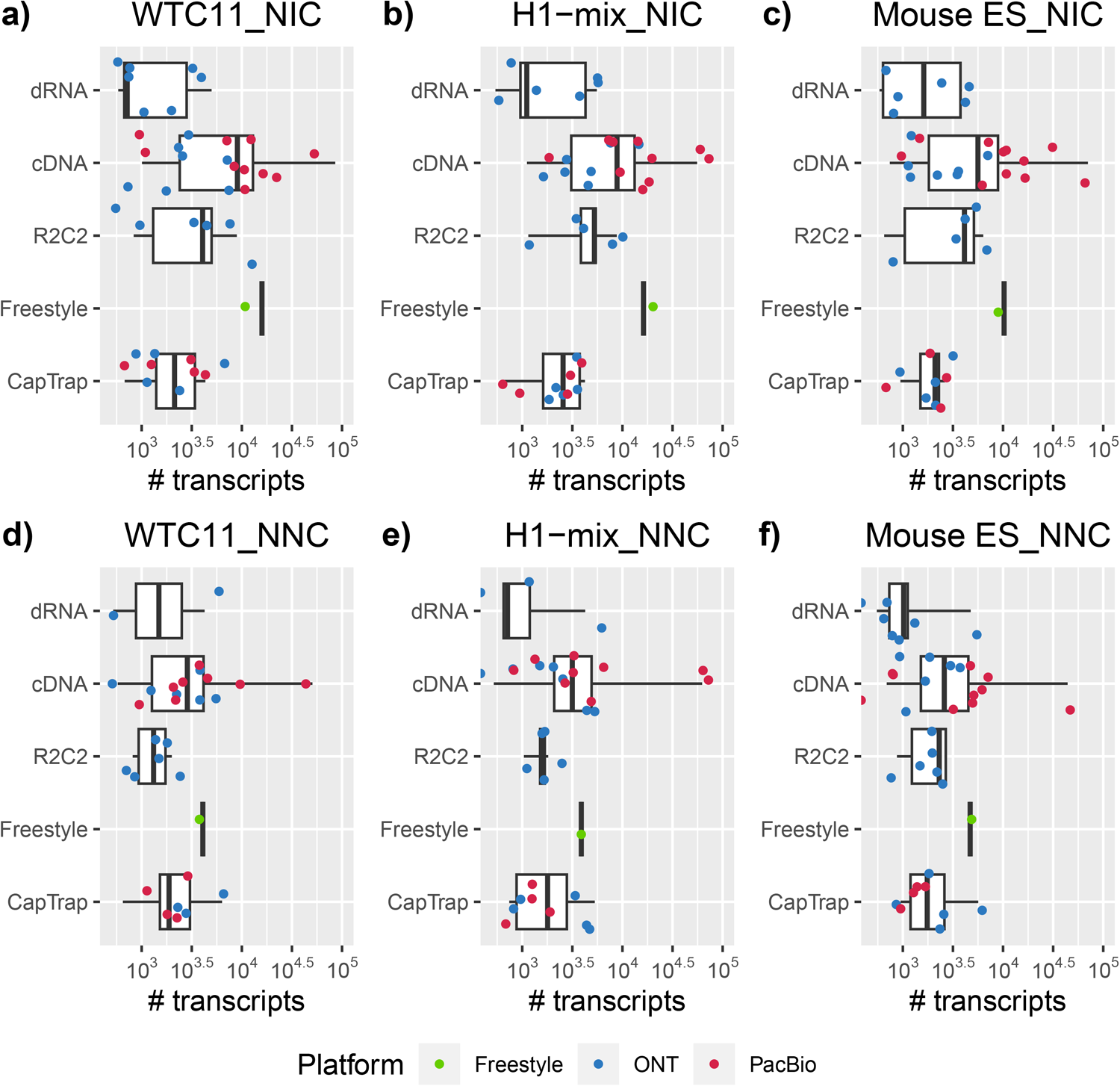
Number of NIC and NNC by sequencing platform and library preparation. a−c) NIC, d−f) NNC.

**Extended Data Fig. 17.**
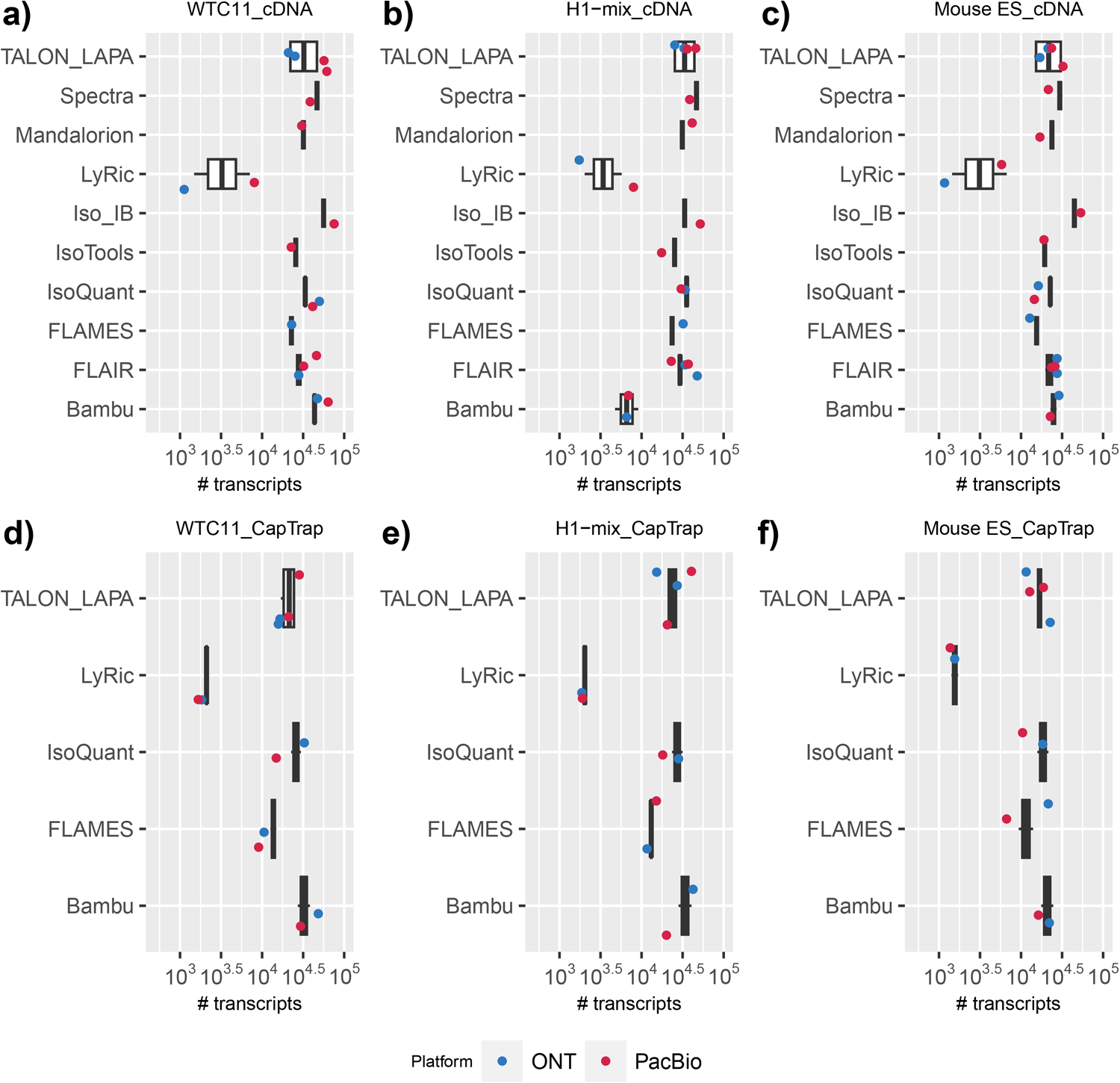
Number of FSM transcripts by library preparation and analysis tool. a−c) cDNA. d−f) CapTrap.

**Extended Data Fig. 18.**
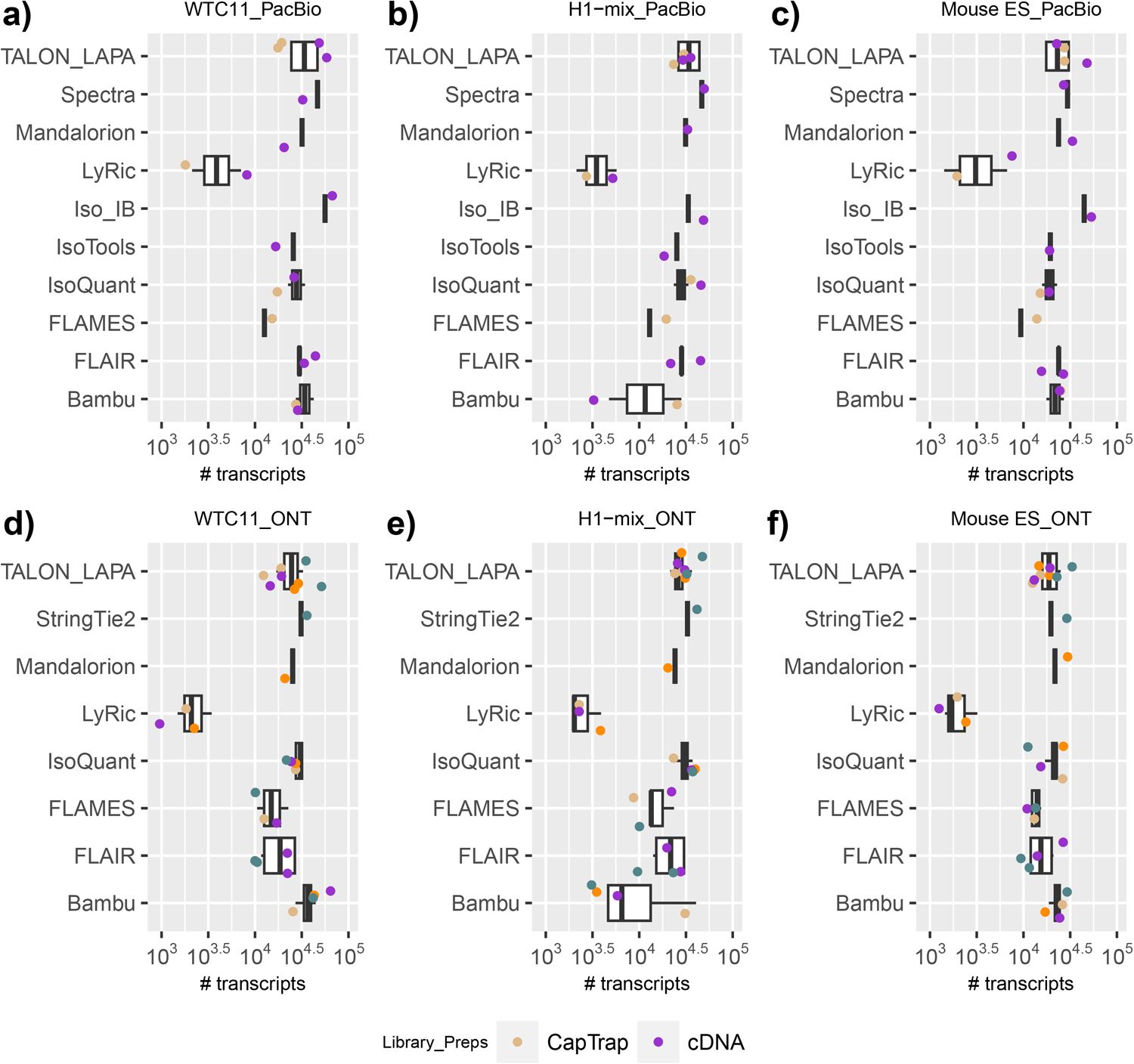
Number of FSM transcripts by sequencing platform and analysis tool. a−c) PacBio, d−f) Nanopore.

**Extended Data Fig. 19.**
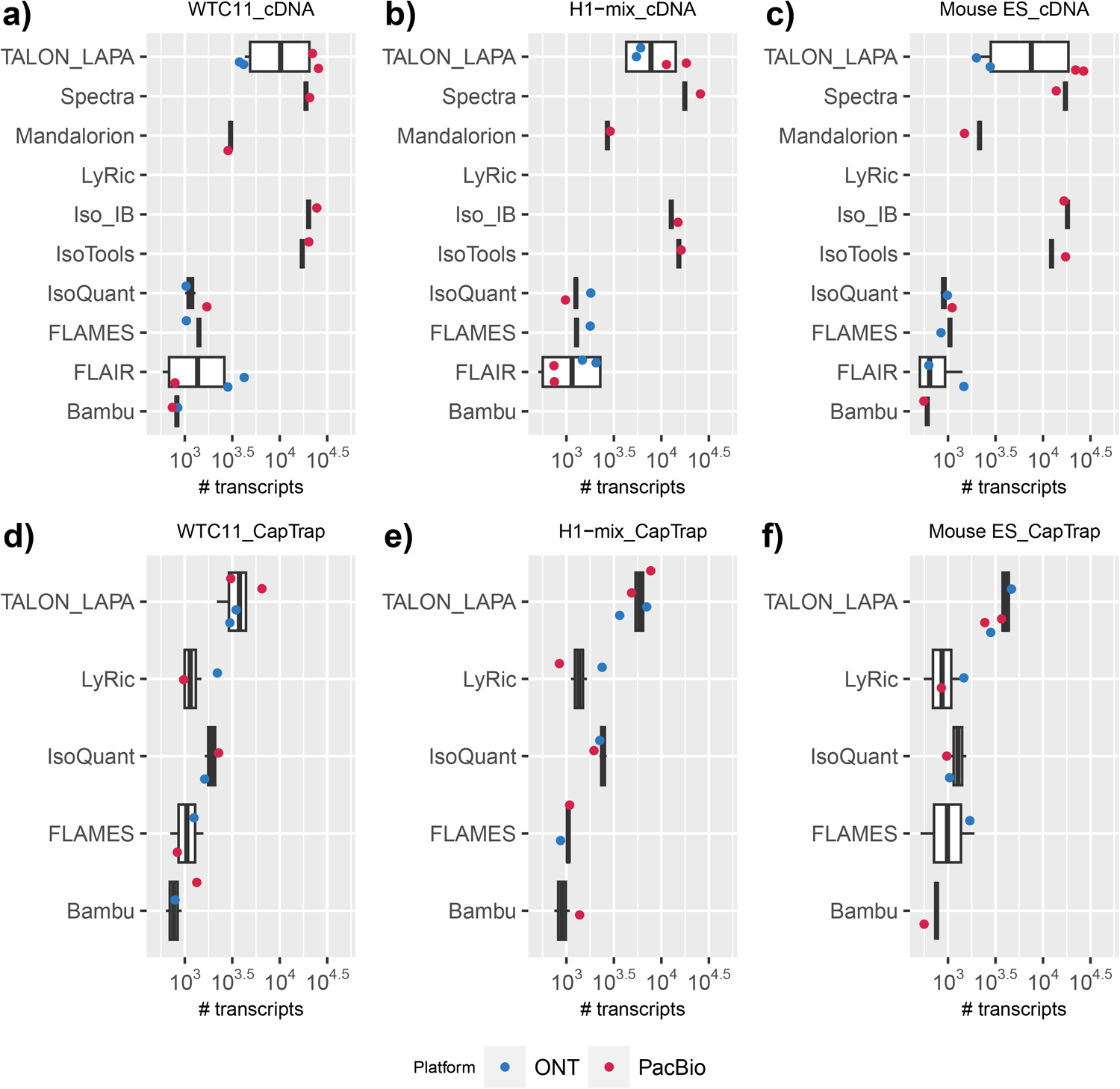
Number of ISM transcripts by library preparation and analysis tool. a−c) cDNA. d−f) CapTrap.

**Extended Data Fig. 20.**
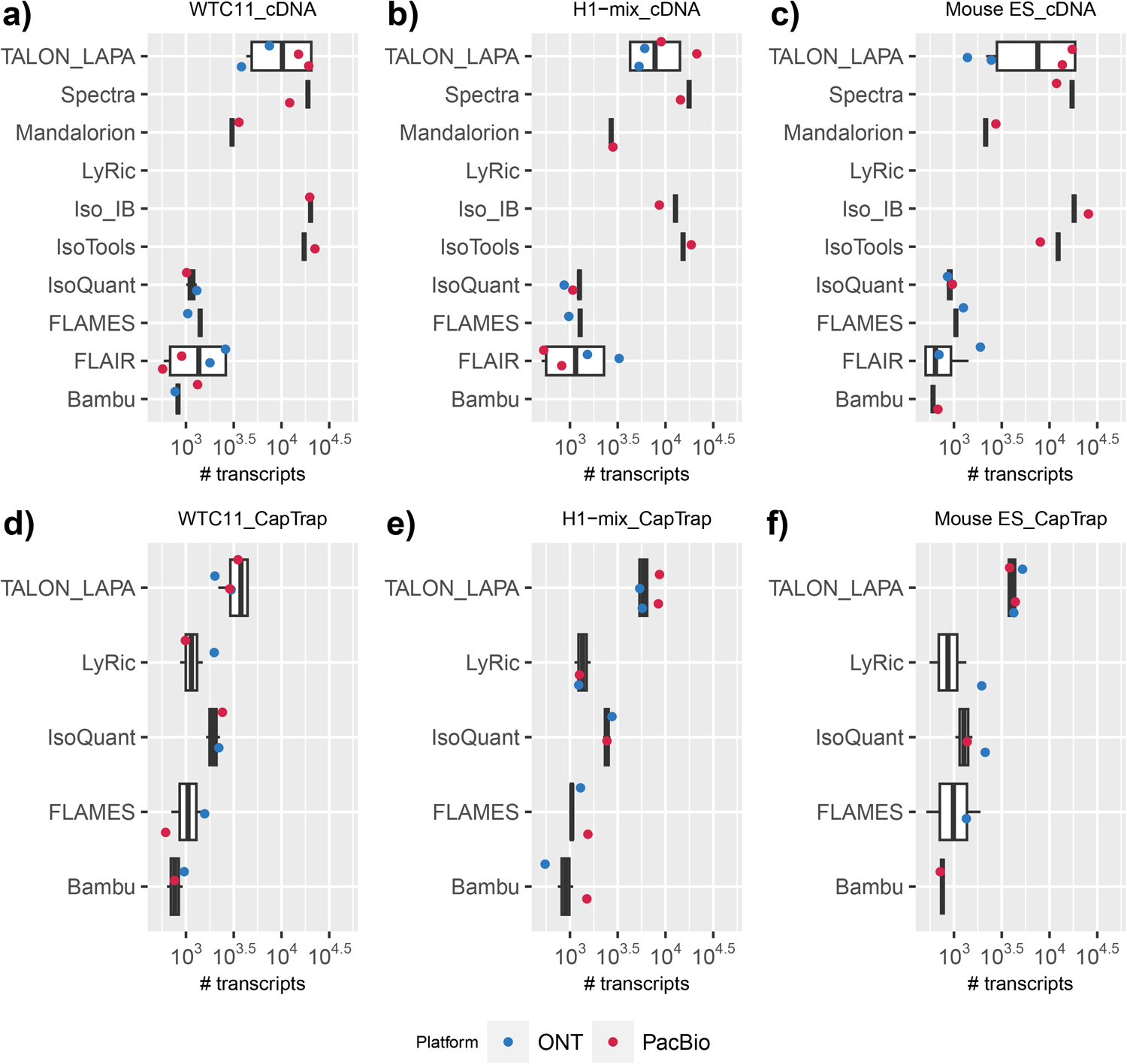
Number of ISM transcripts by sequencing platform and analysis tool. a−c) Intergenic. d−f) GenicGenomic.

**Extended Data Fig. 21.**
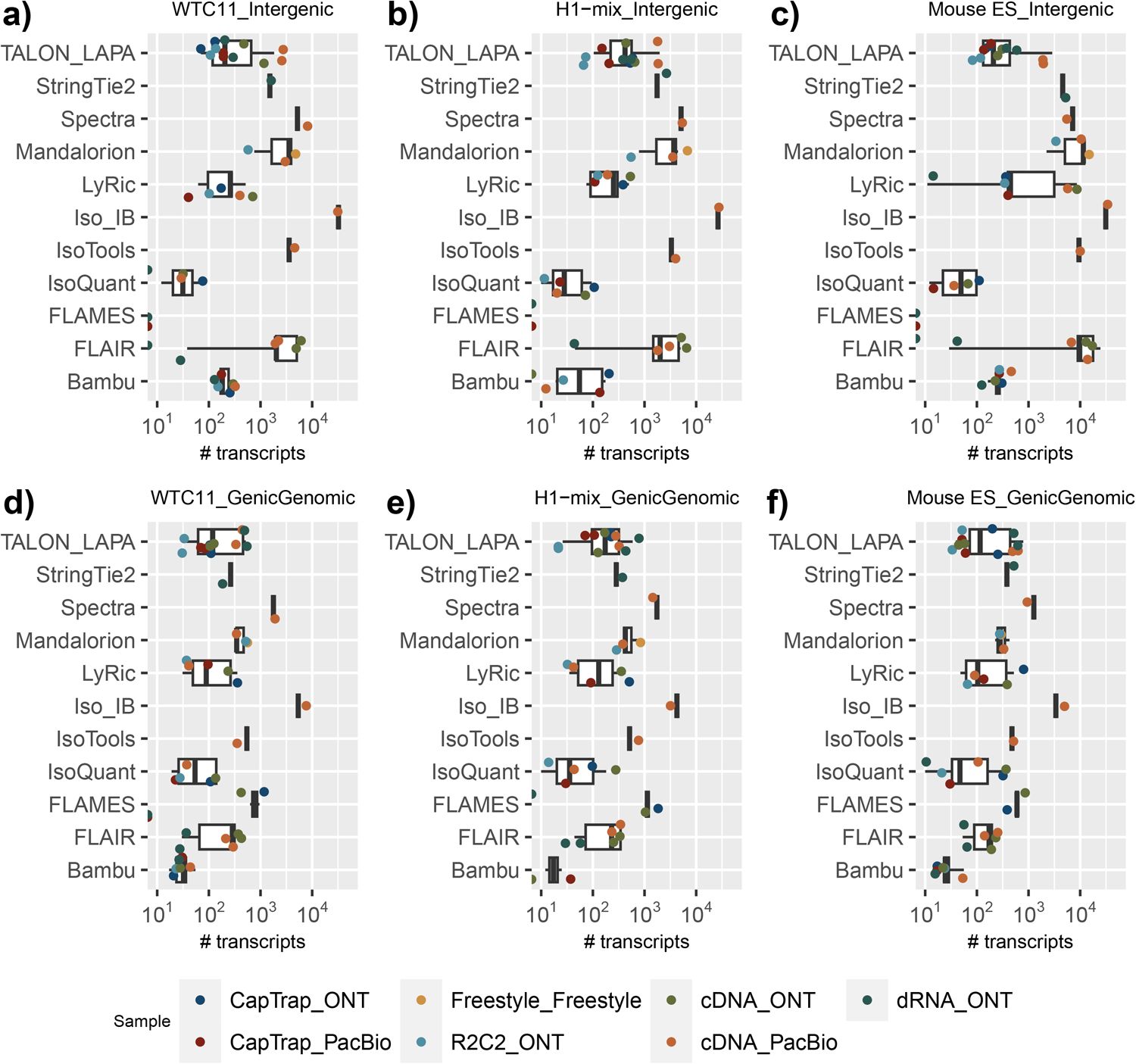
Number of Intergenic and GenicGenomic by sequencing platform and library preparation. a−c) Intergenic, d−f) GenicGenomic.

**Extended Data Fig. 22.**
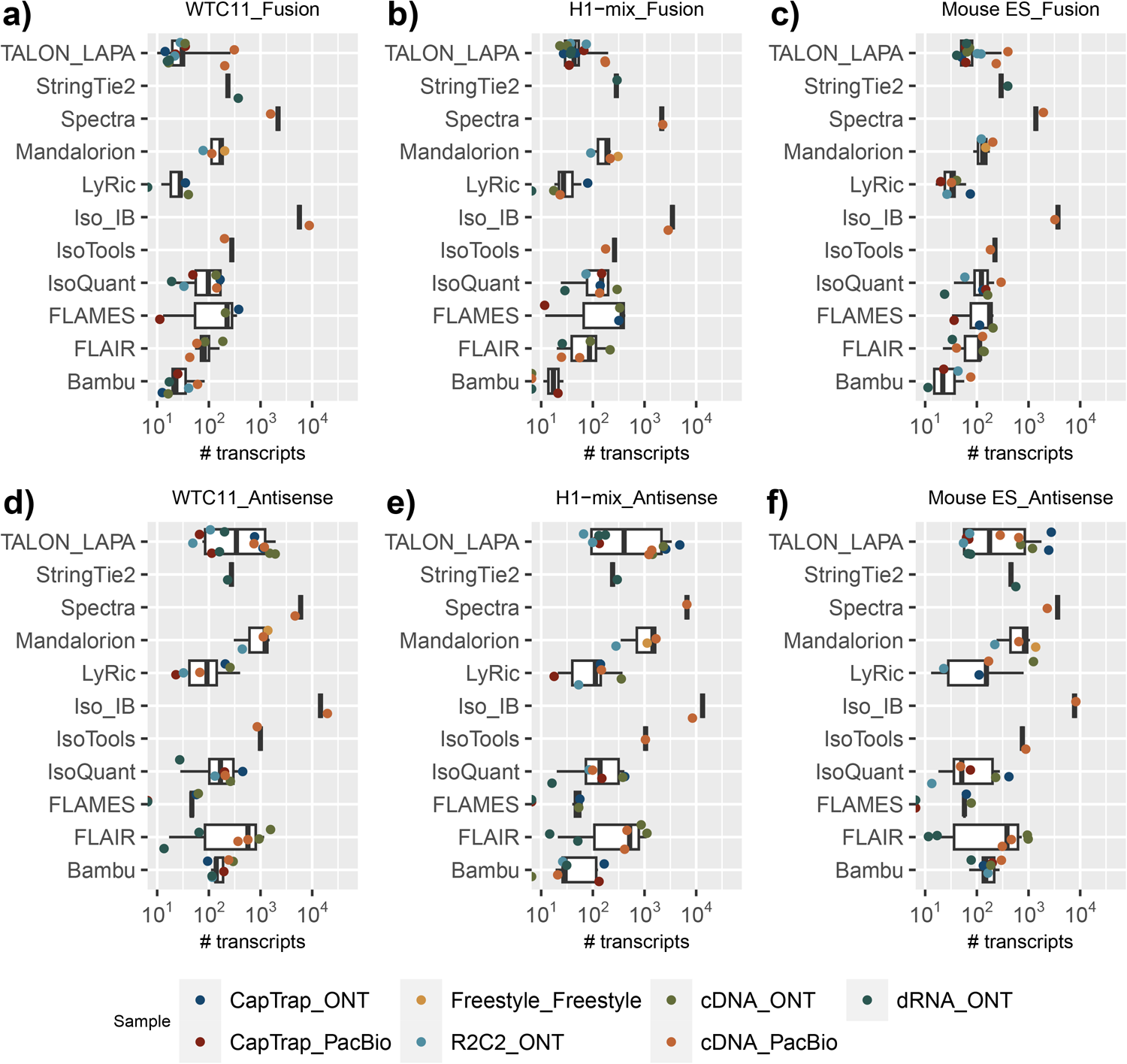
Number of Fusion and Antisense by sequencing platform and library preparation. a−c) Fusion. d−f) Antisense.

**Extended Data Fig. 23.**
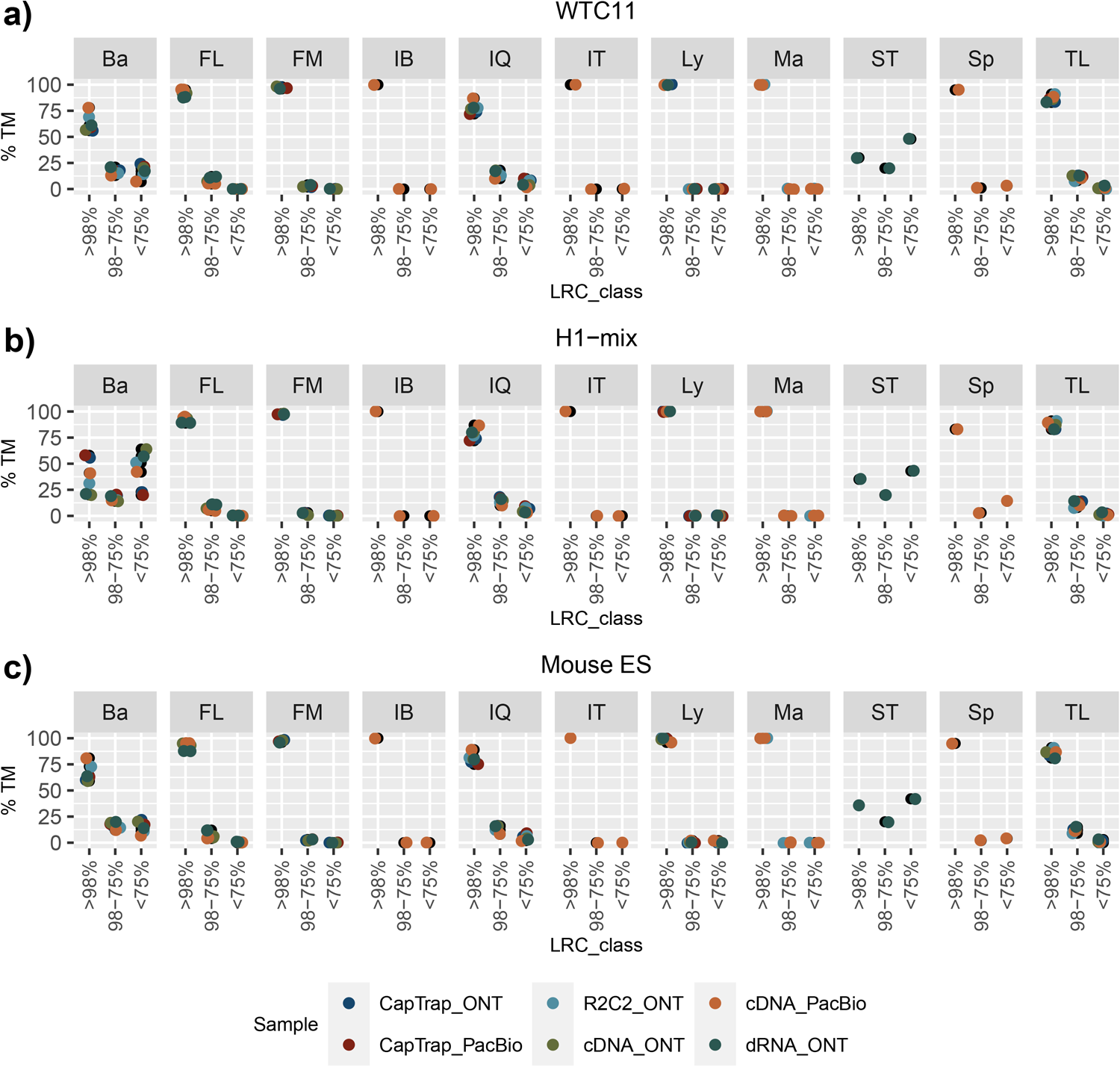
Percentage of transcript models (TM) with different ranges of sequence coverage by long reads. a) WTC11. c) H1−mix. c) Mouse ES. Ba: Bambu, FM: FLAMES, FL: FLAIR, IQ: IsoQuant, IT: IsoTools, IB: Iso_IB, Ly: LyRic, Ma: Mandalorion, TL: TALON−LAPA, Sp: Spectra, ST: StringTie2.

**Extended Data Fig. 24.**
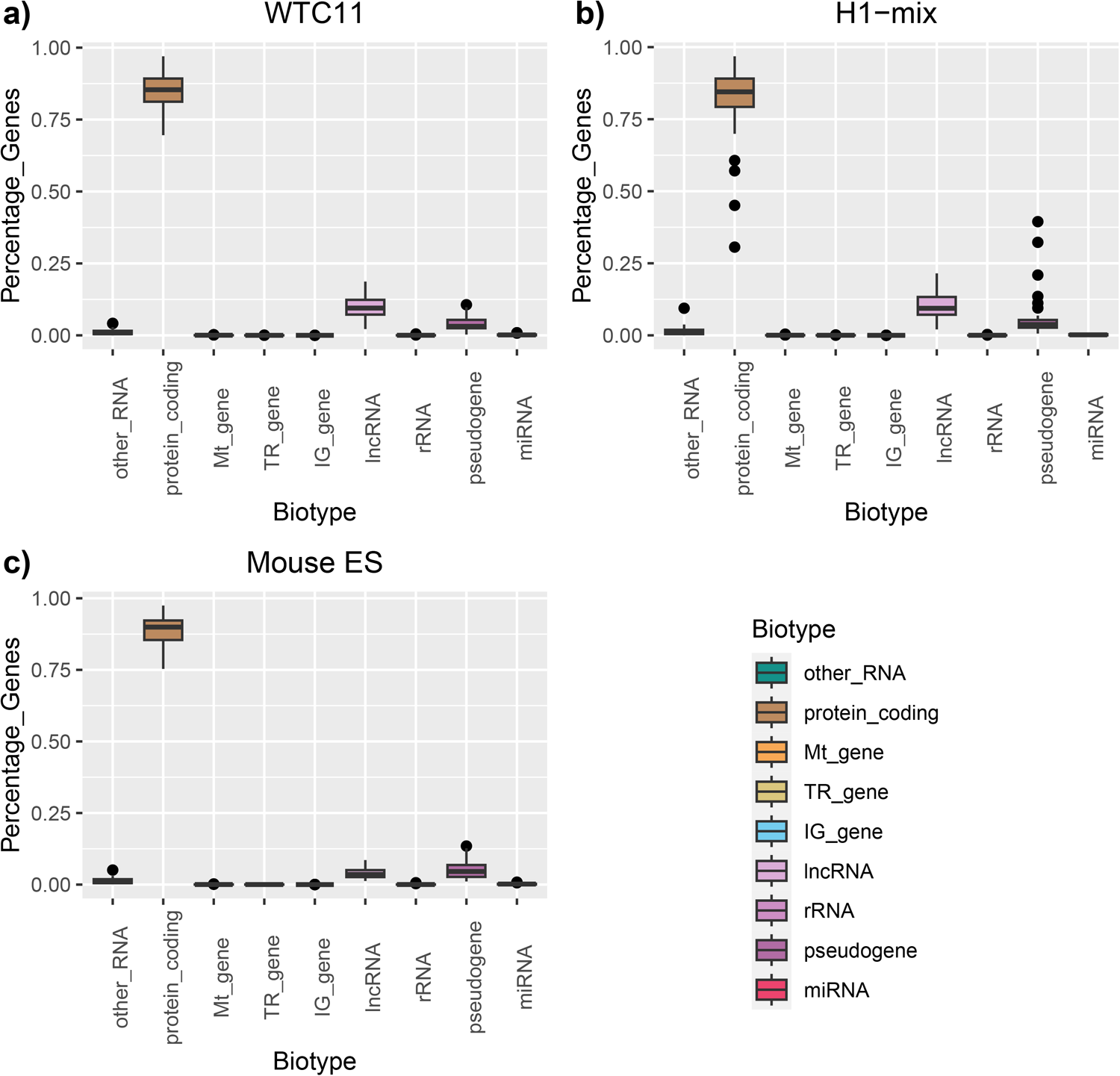
Distribution of Biotypes across pipelines. a) WTC11, c) H1−mix, c) Mouse ES.

**Extended Data Fig. 25.**
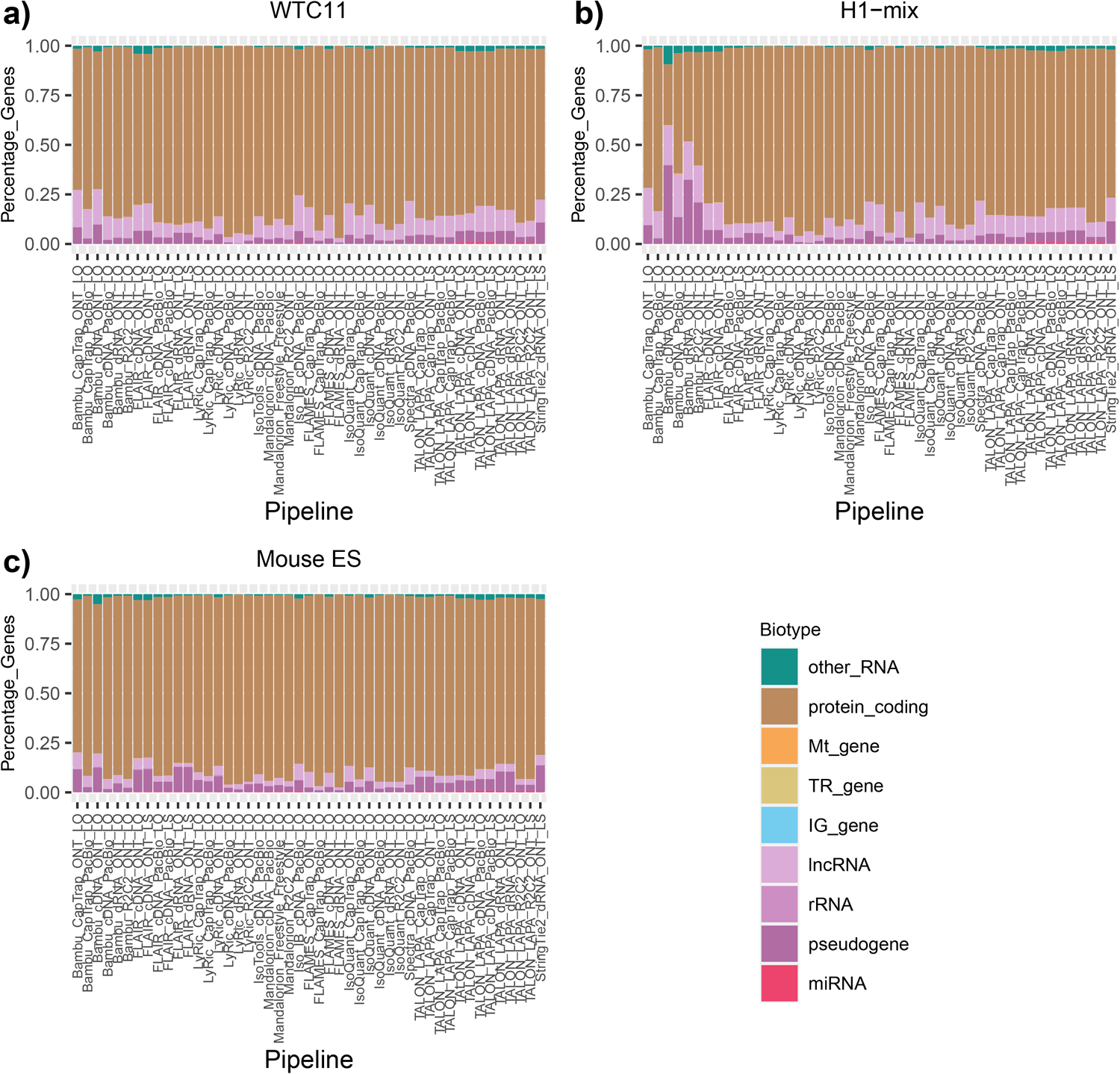
Biotypes per pipeline. a) WTC11, c) H1−mix, c) Mouse ES.

**Extended Data Fig. 26.**
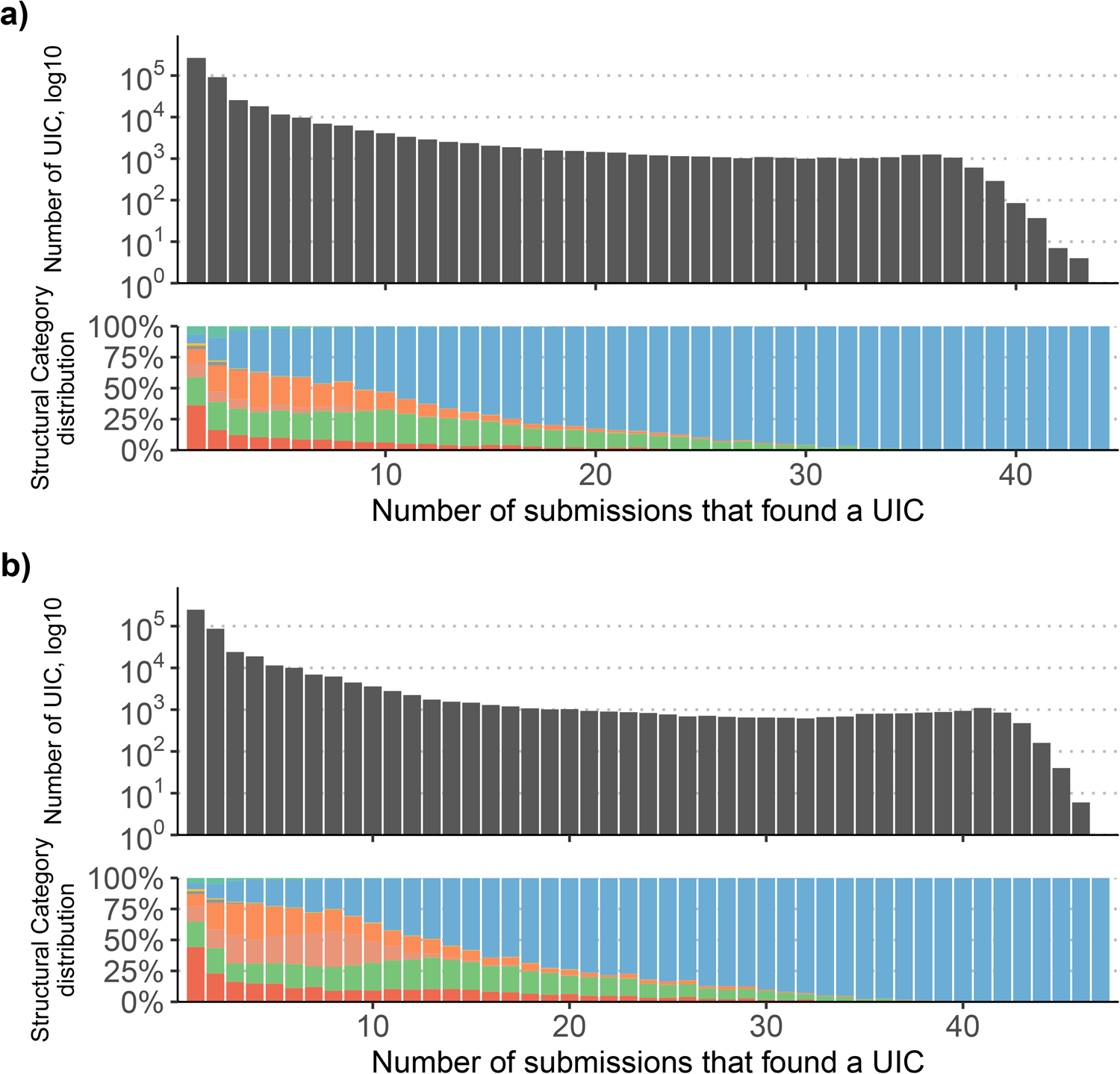
Number and SQANTI category distribution of Unique Intron Chain (UIC) consistently detected by an increasing number of submissions. a) H1−mix sample, b) Mouse ES sample.

**Extended Data Fig. 27.**
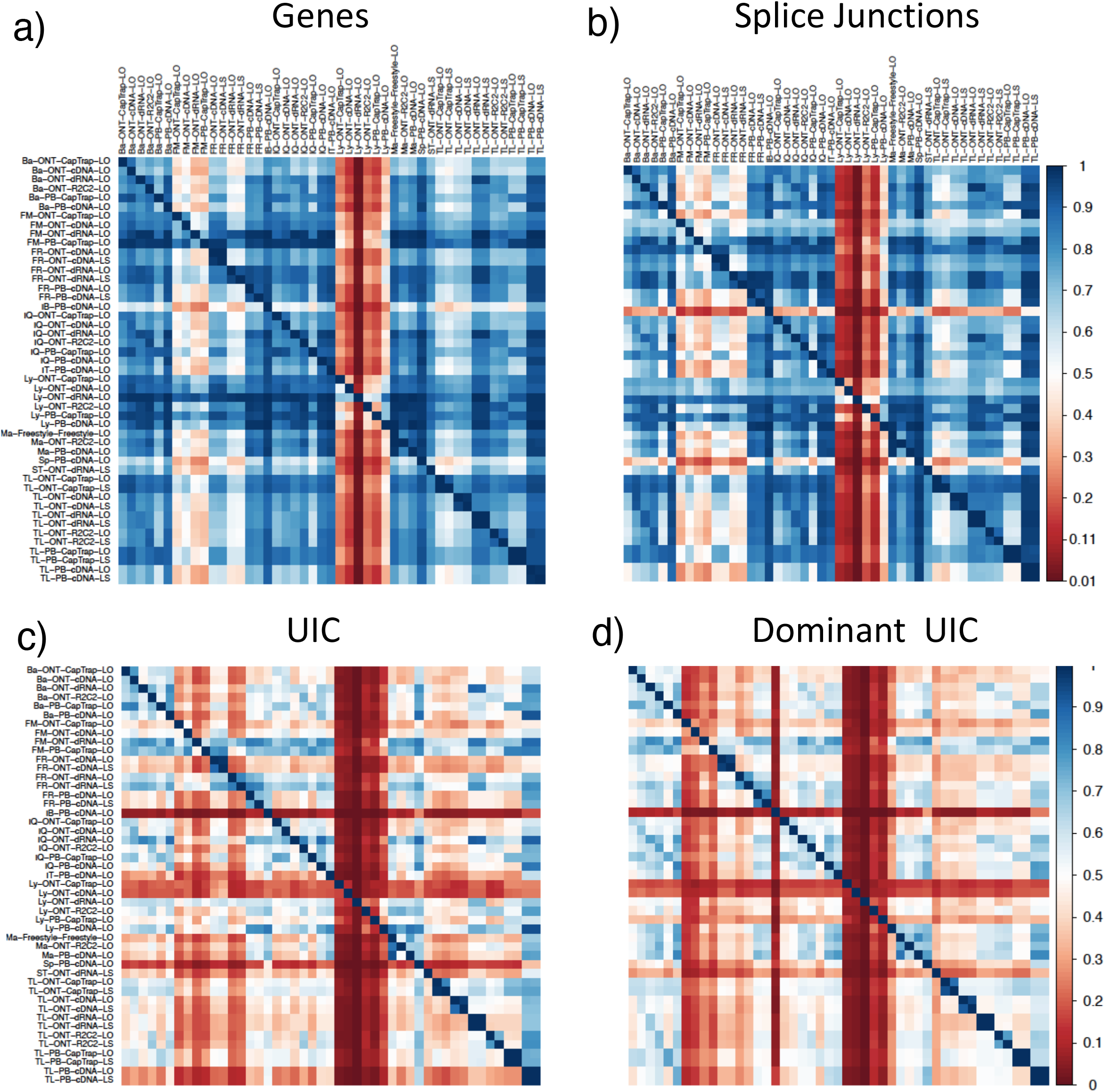
Pair-wise overlap in the detection of features between pipelines; WTC11 sample. Each value represents the feature intersection between column and row pipelines divided by the number of detections in the row pipeline. a) Genes, b) Splice junctions, c) Unique Intron Chains (UIC), c) Top UIC accounting for at least 50% of the gene expression.

**Extended Data Fig. 28.**
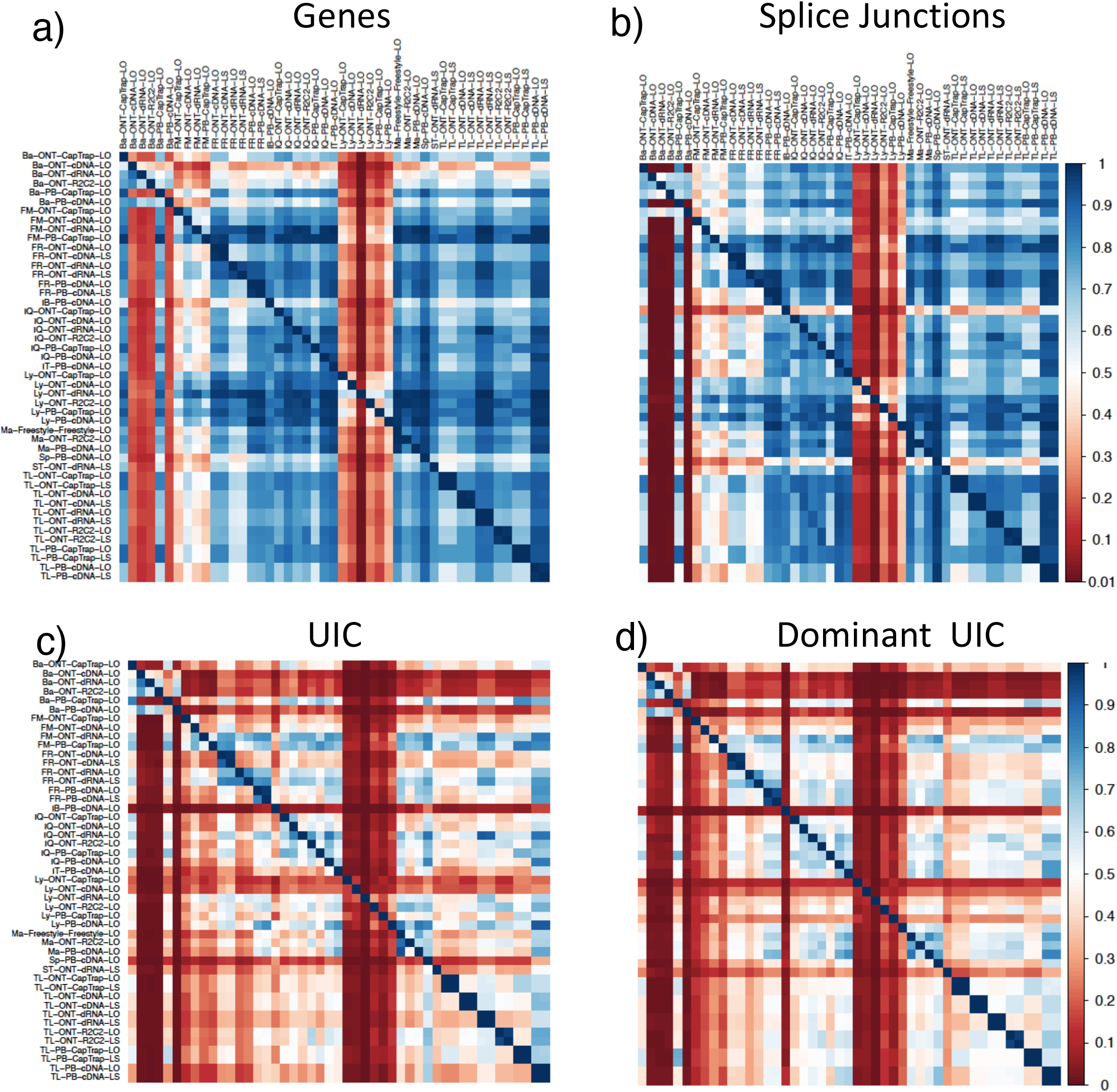
Pair-wise overlap in the detection of features between pipelines; H1-mix sample. Each value represents the feature intersection between column and row pipelines divided by the number of detections in the row pipeline. a) Genes, b) Splice junctions, c) Unique Intron Chains (UIC), c) Top UIC accounting for at least 50% of the gene expression.

**Extended Data Fig. 29.**
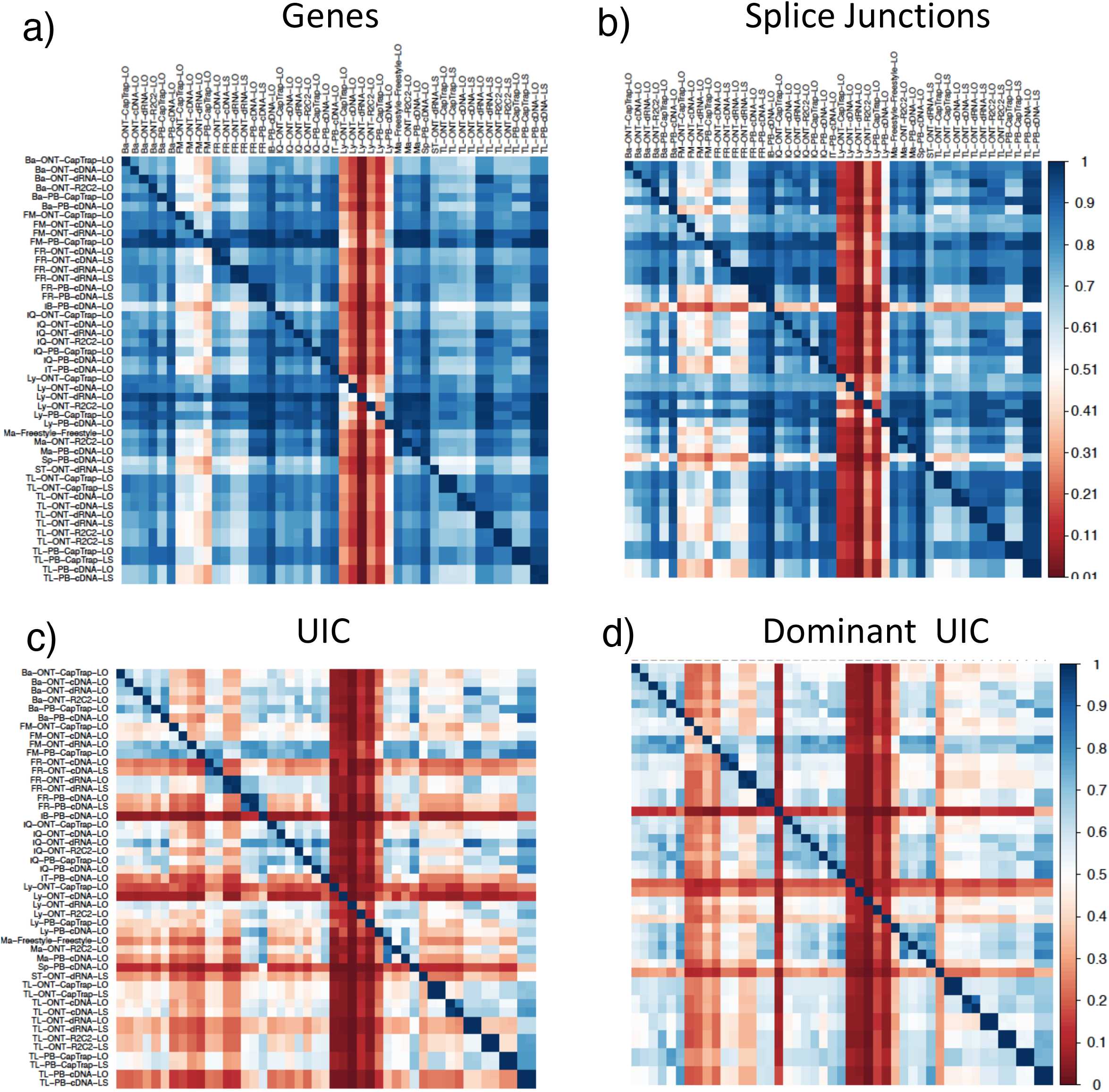
Pair-wise overlap in the detection of features between pipelines; ES mouse sample. Each value represents the feature intersection between column and row pipelines divided by the number of detections in the row pipeline. a) Genes, b) Splice junctions, c) Unique Intron Chains (UIC), c) Top UIC accounting for at least 50% of the gene expression.

**Extended Data Fig. 30.**
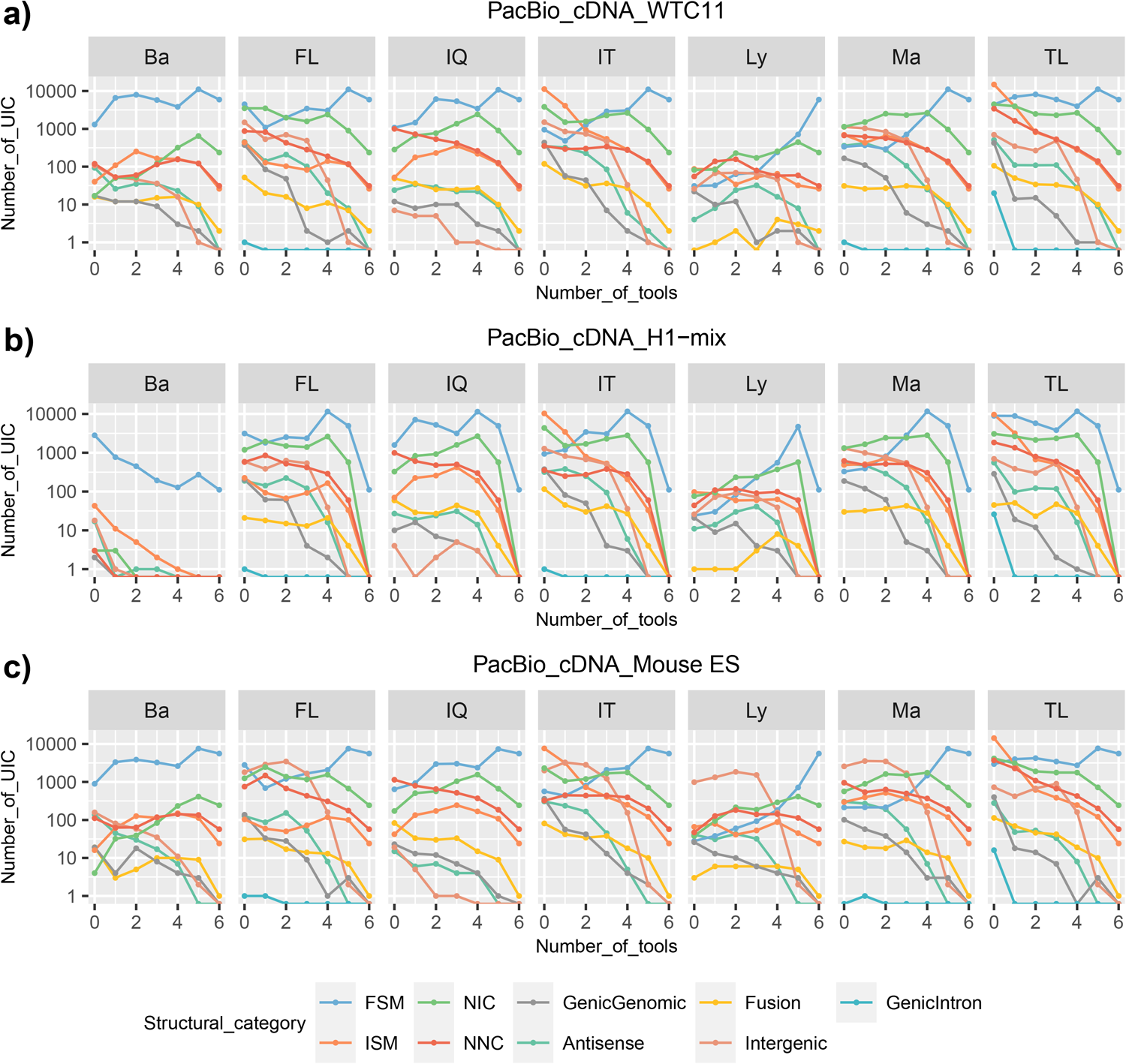
Number of UIC detected by a tool and shared with an increasing number of other tools, processing PacBio_cDNA data. a) WTC11, c) H1−mix, c) Mouse ES.

**Extended Data Fig. 31.**
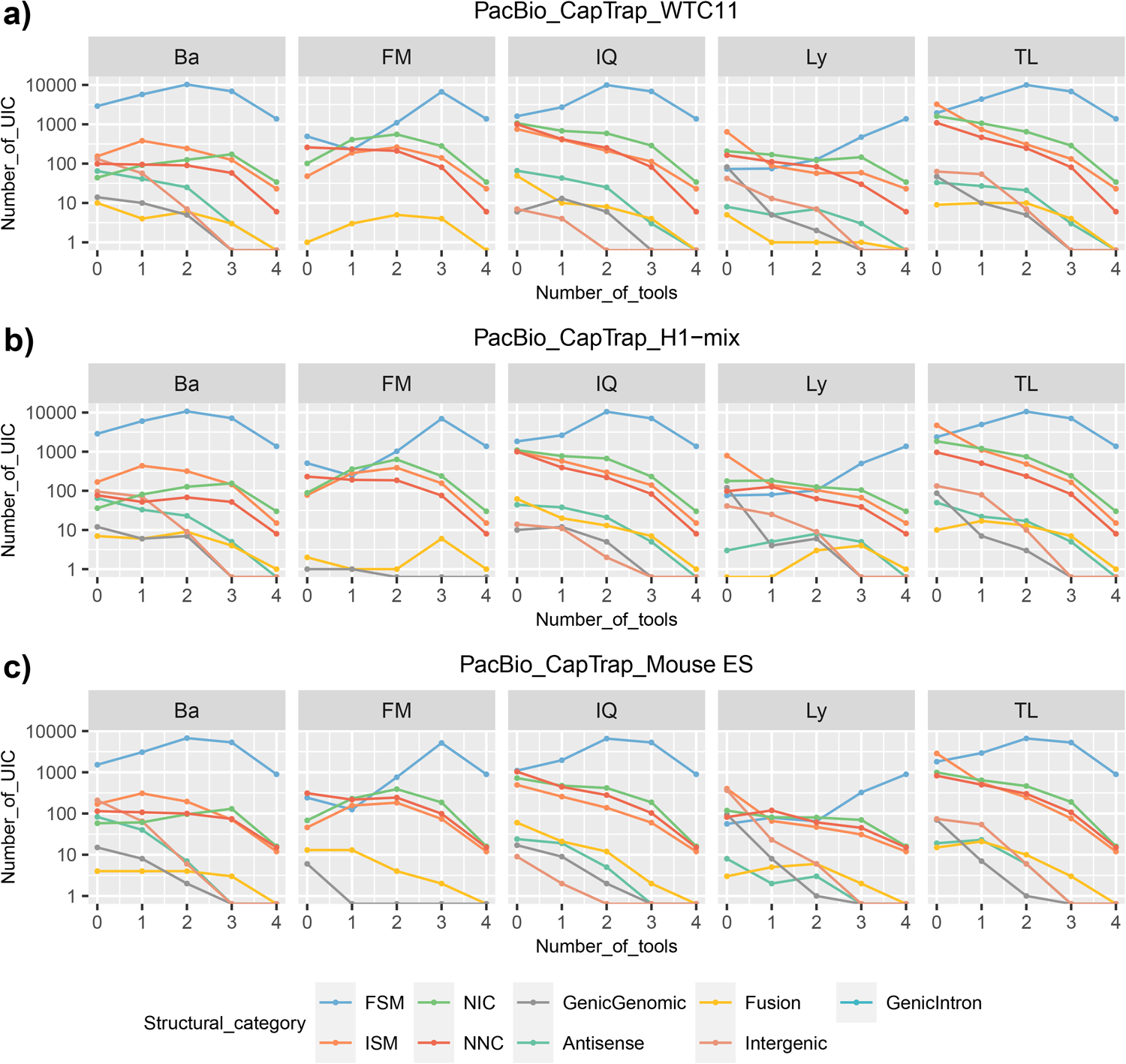
Number of UIC detected by a tool and shared with an increasing number of other tools, processing PacBio_CapTrap data. a) WTC11, c) H1−mix, c) Mouse ES.

**Extended Data Fig. 32.**
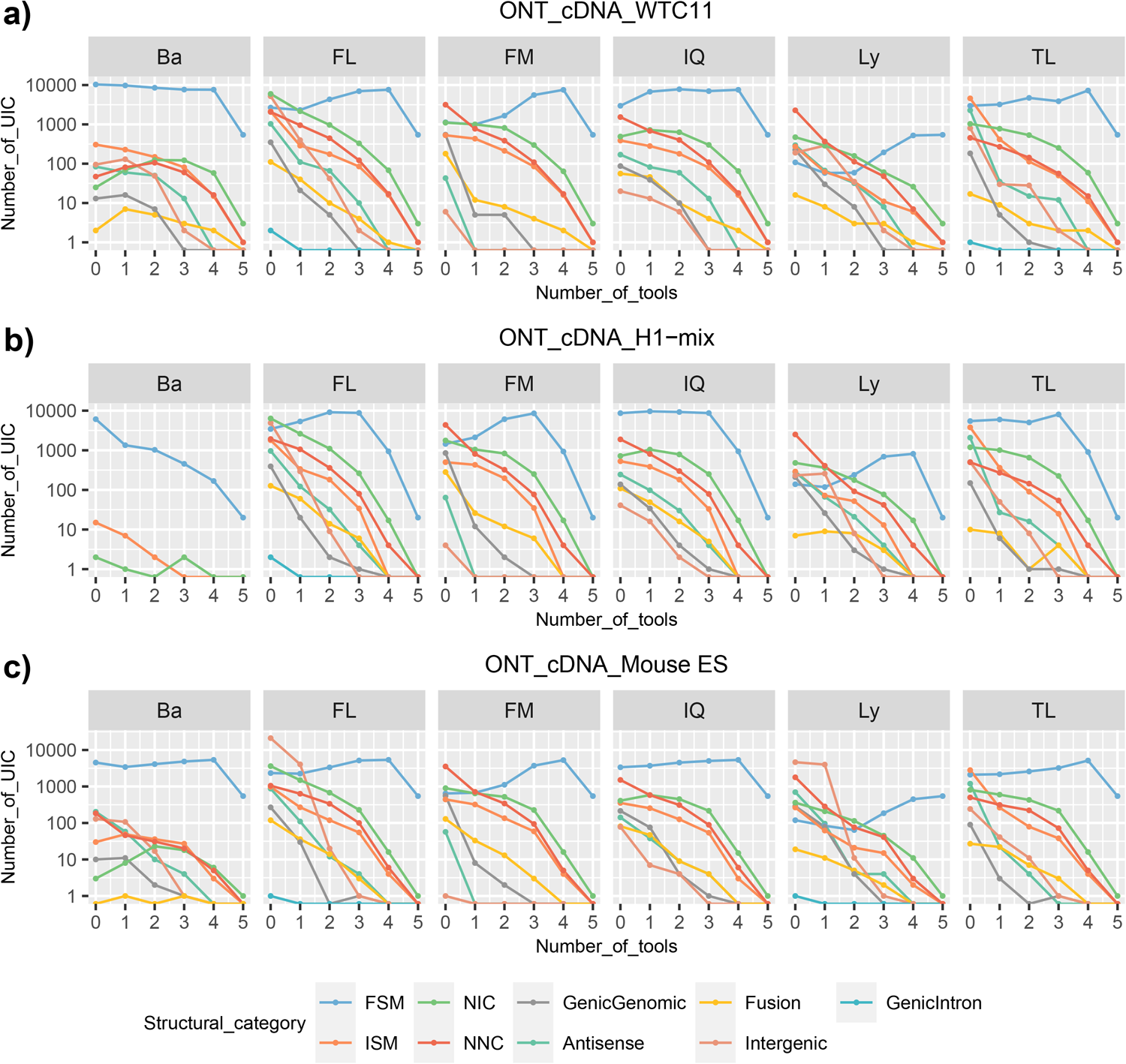
Number of UIC detected by a tool and shared with an increasing number of other tools, processing PacBio_CapTrap data. a) WTC11, c) H1−mix, c) Mouse ES.

**Extended Data Fig. 33.**
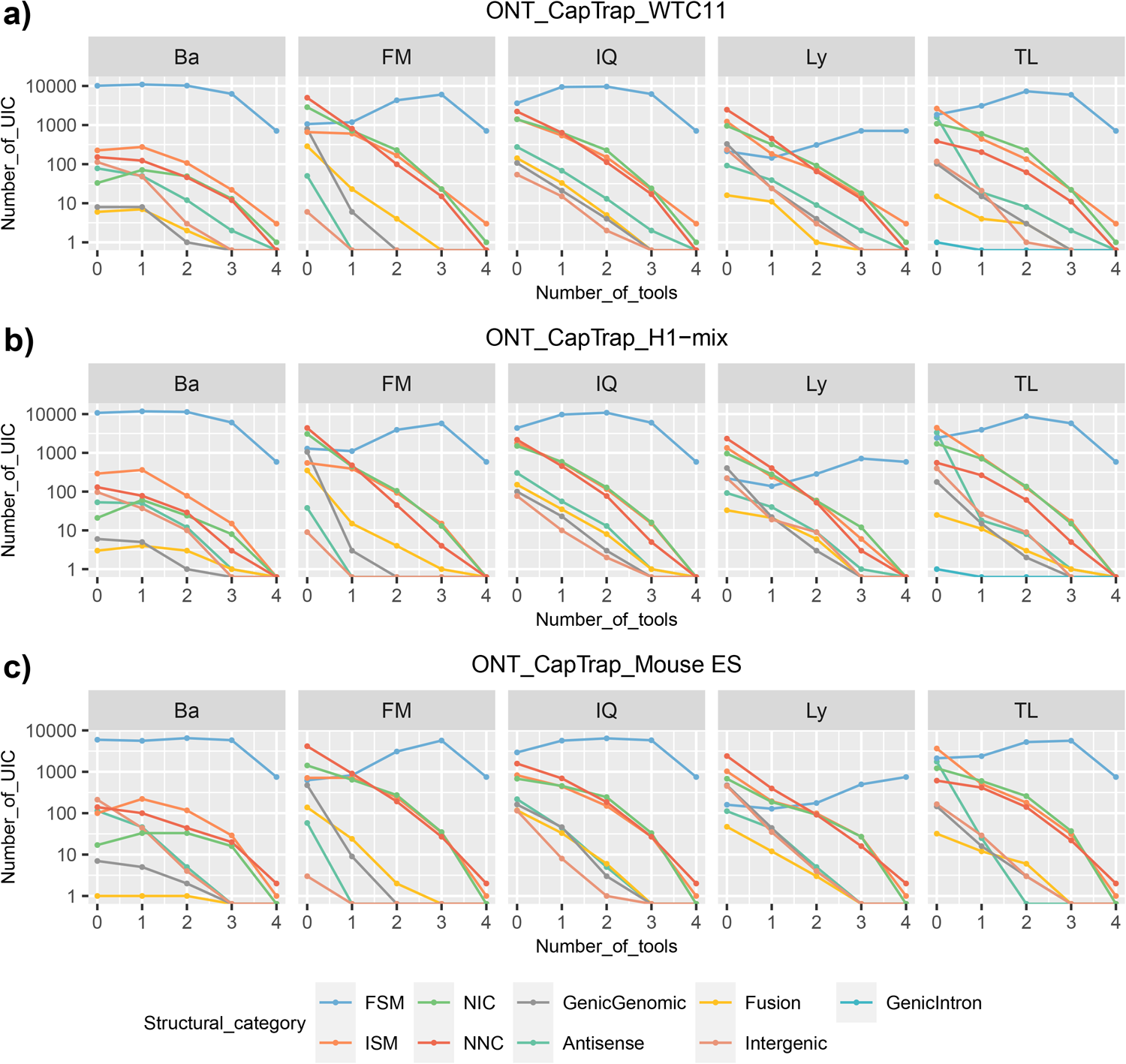
Number of UIC detected by a tool and shared with an increasing number of other tools, processing ONT_CapTrap data. a) WTC11, c) H1−mix, c) Mouse ES.

**Extended Data Fig. 34.**
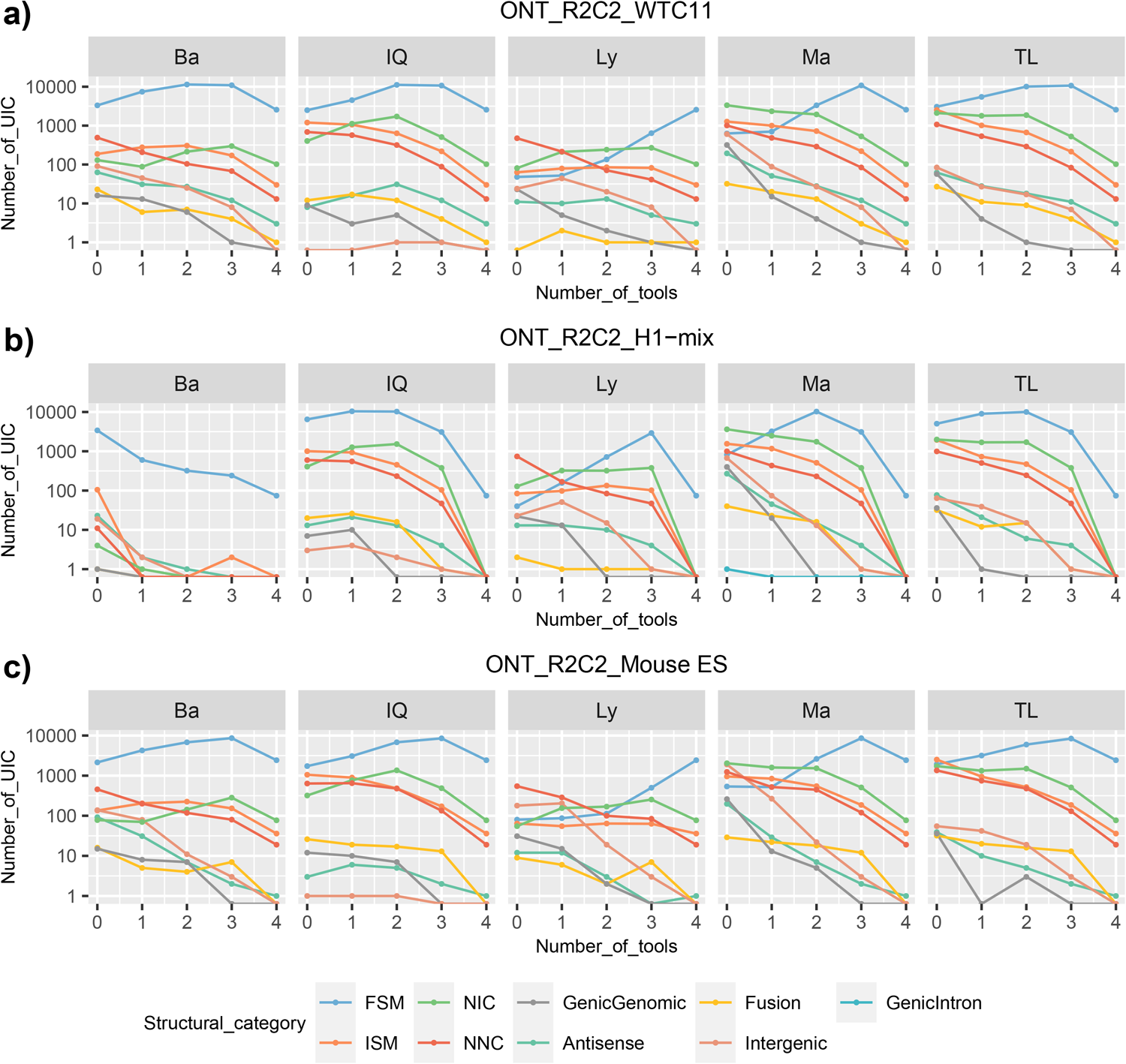
Number of UIC detected by a tool and shared with an increasing number of other tools, processing ONT_R2C2 data. a) WTC11, c) H1−mix, c) Mouse ES

**Extended Data Fig. 35.**
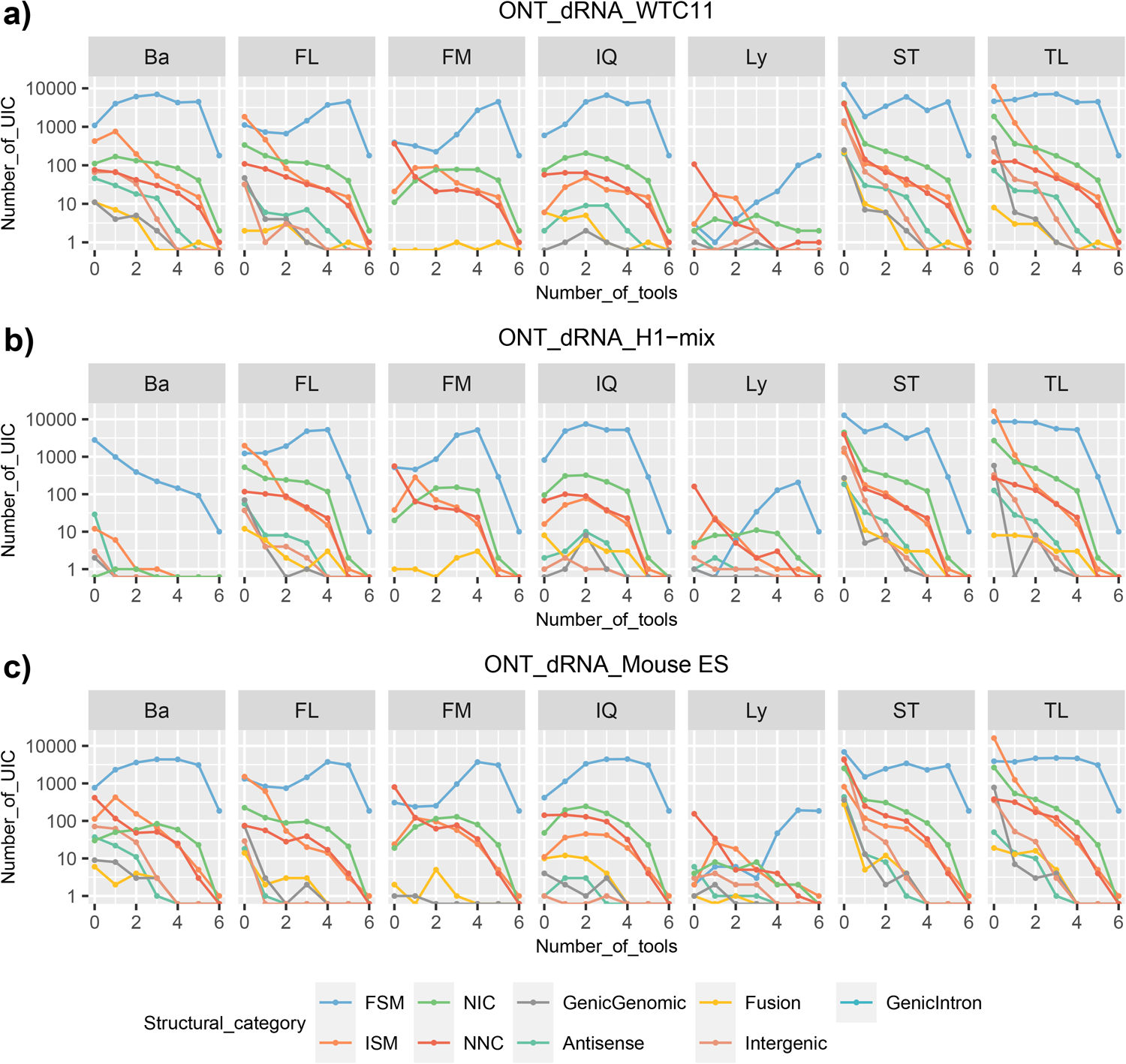
Number of UIC detected by a tool and shared with an increasing number of other tools, processing ONT_dRNA data. a) WTC11, c) H1−mix, c) Mouse ES

**Extended Data Fig. 36.**
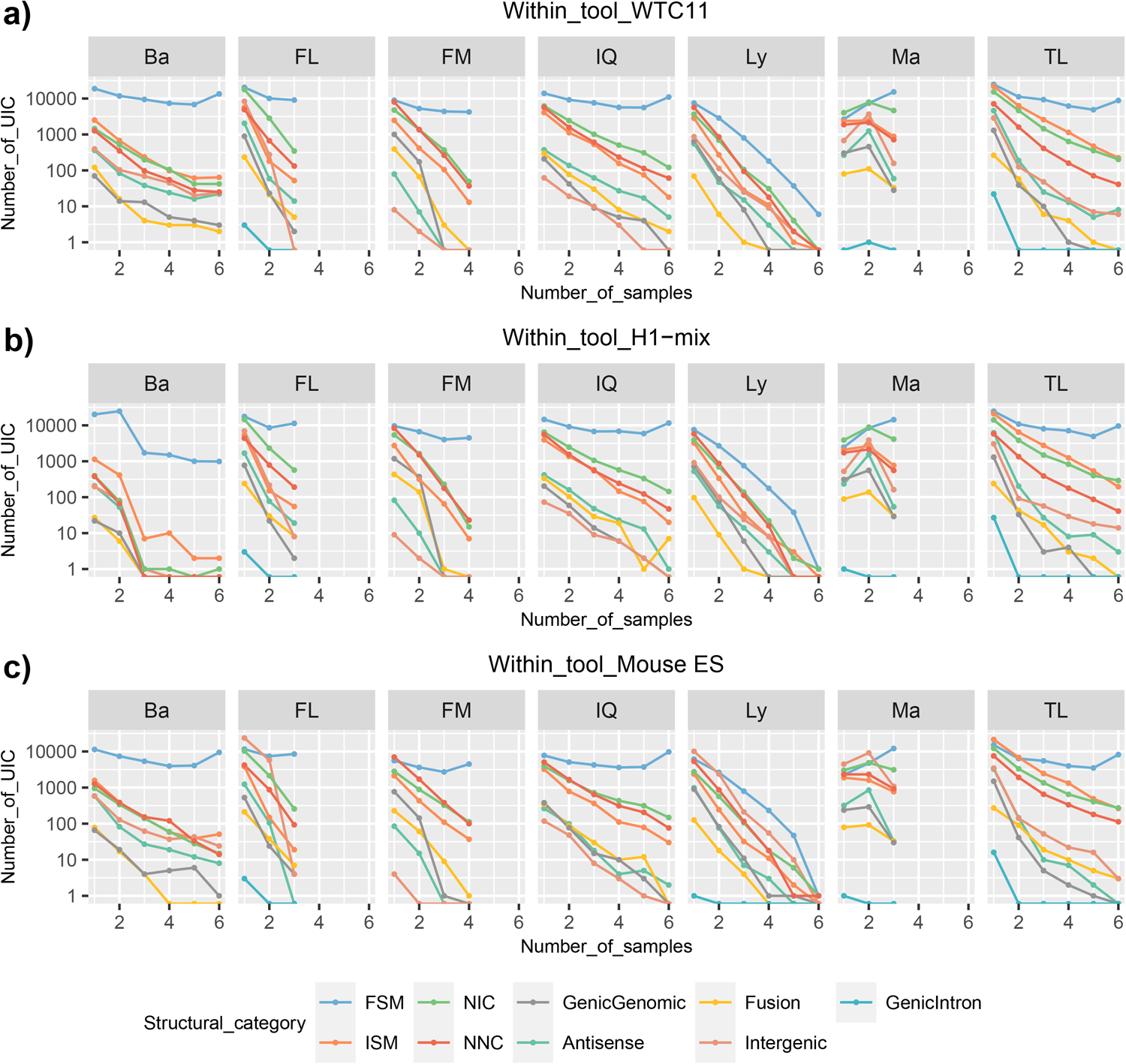
Number of UIC consistently detected by a tool across samples. a) WTC11, c) H1−mix, c) Mouse ES

**Extended Data Fig. 37.**
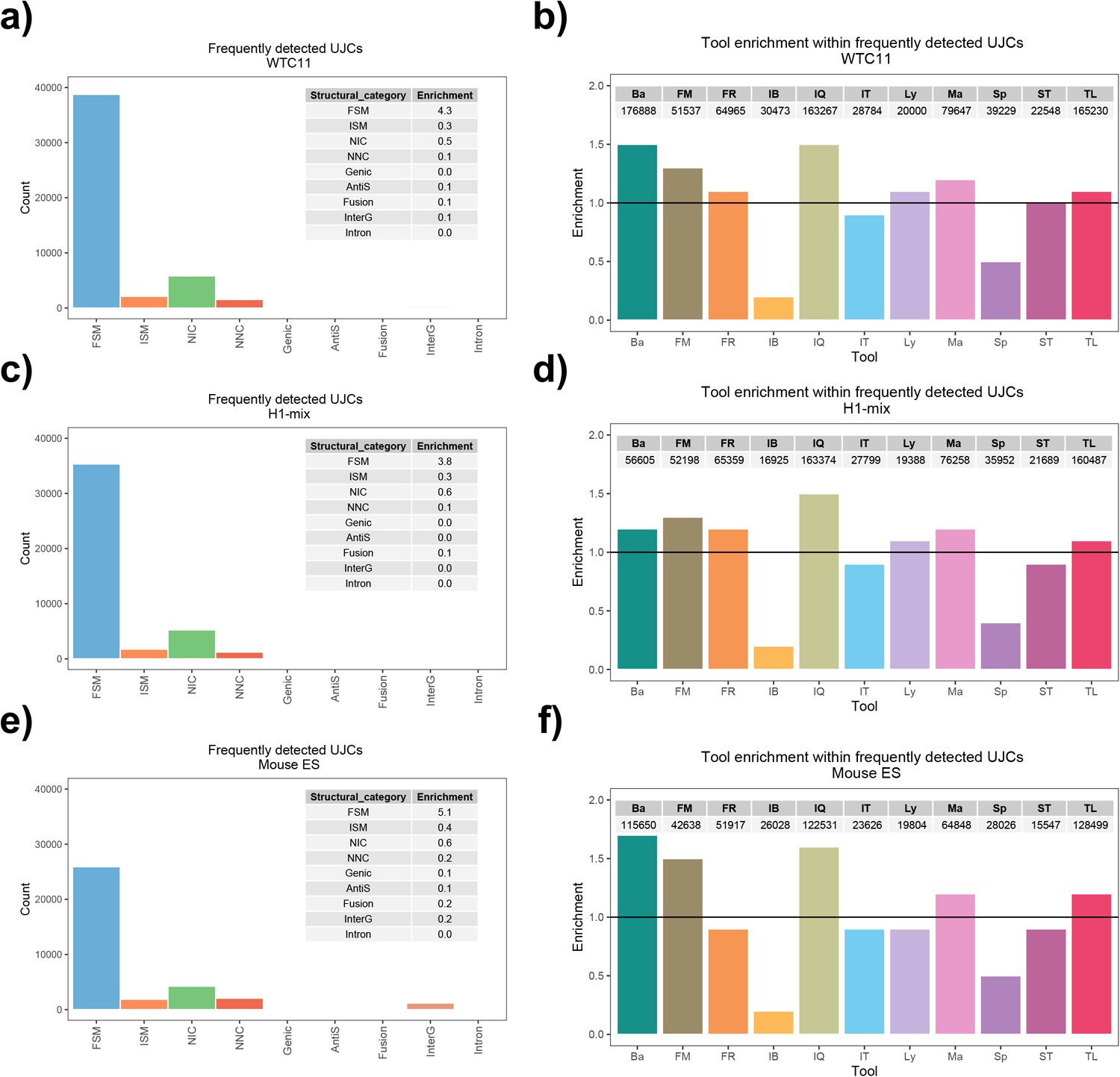
Characterization of frequently detected UICs (FDU). a,c,e) Structural category distribution of FDU. The table indicates the fold enrichment of each structural category within the frequently detected transcripts respect to their global count. b,d,f) Tools identifying FDU. The graph shows the enrichment in the number FDU found by a tool with respect to their global number of reported transcripts. The table reports the total number of FDU detected by the tool.

**Extended Data Fig. 38.**
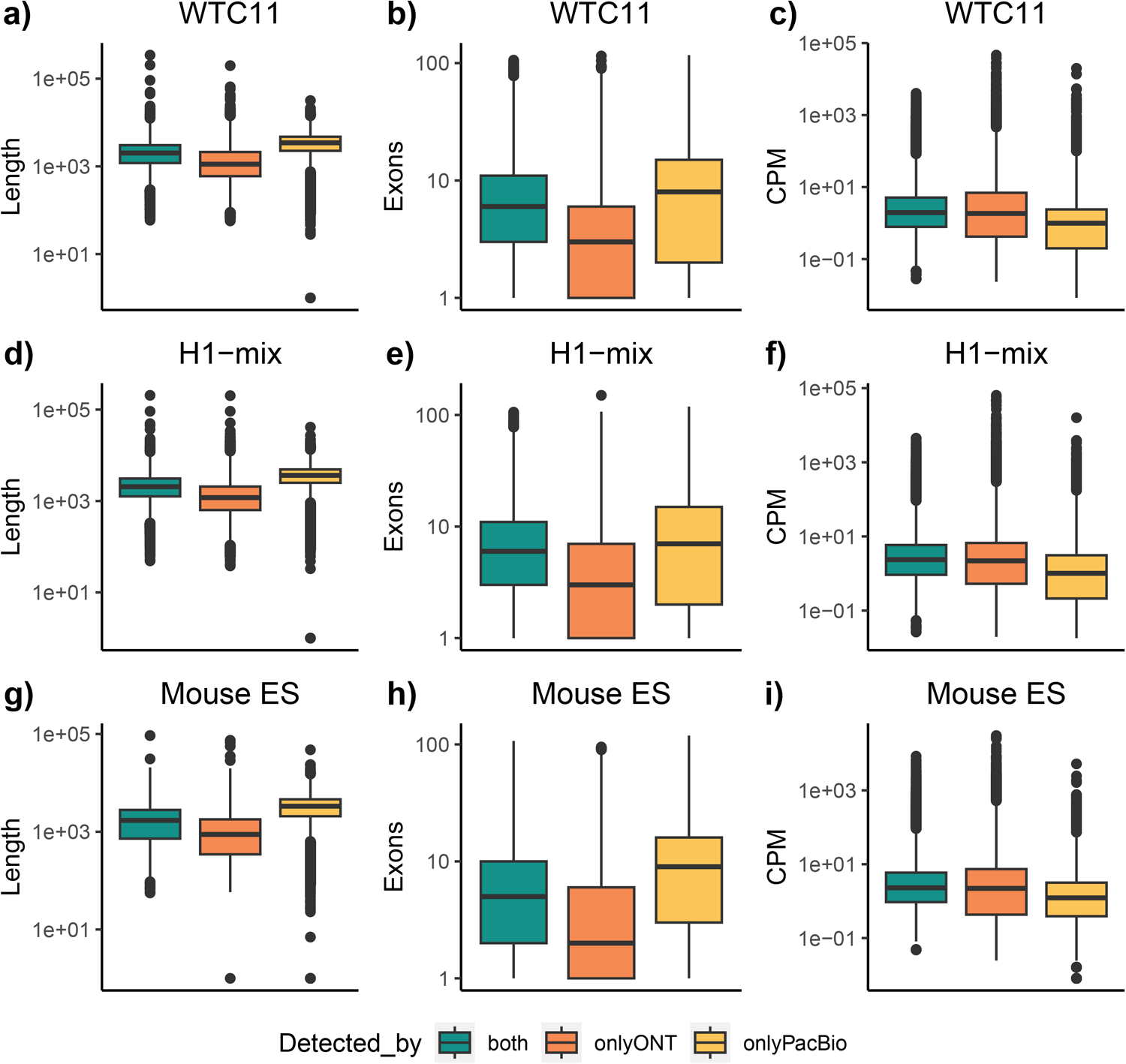
Properties of detected transcripts by library preparation. a,d,g) Length distribution. b,e,h) Exon number distribution. c,f,i) Counts per million

**Extended Data Fig. 39.**
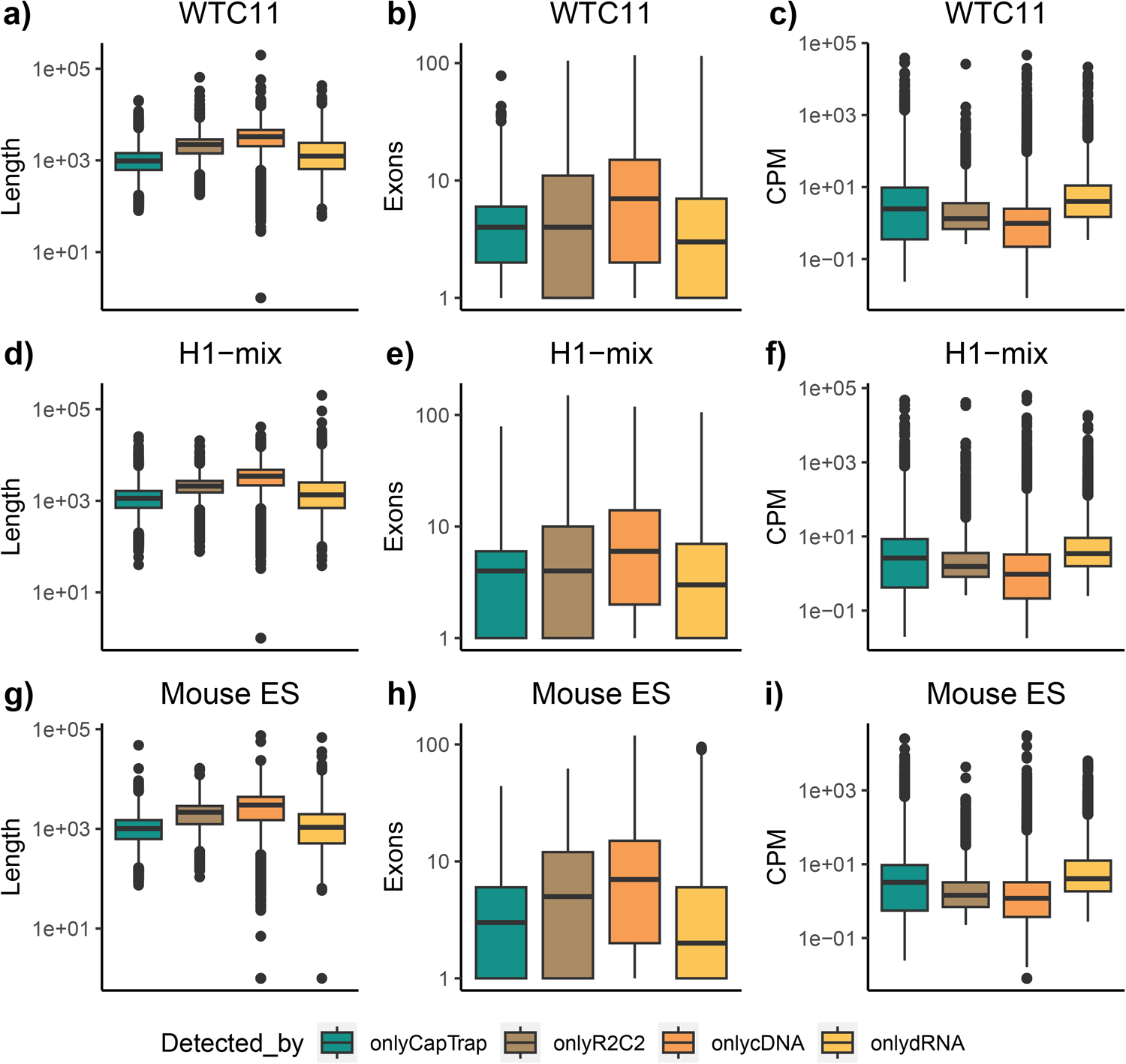
Properties of detected transcripts by library preparation. a,d,g) Length distribution. b,e,h) Exon number distribution. c,f,i) Counts per million

**Extended Data Fig. 40.**
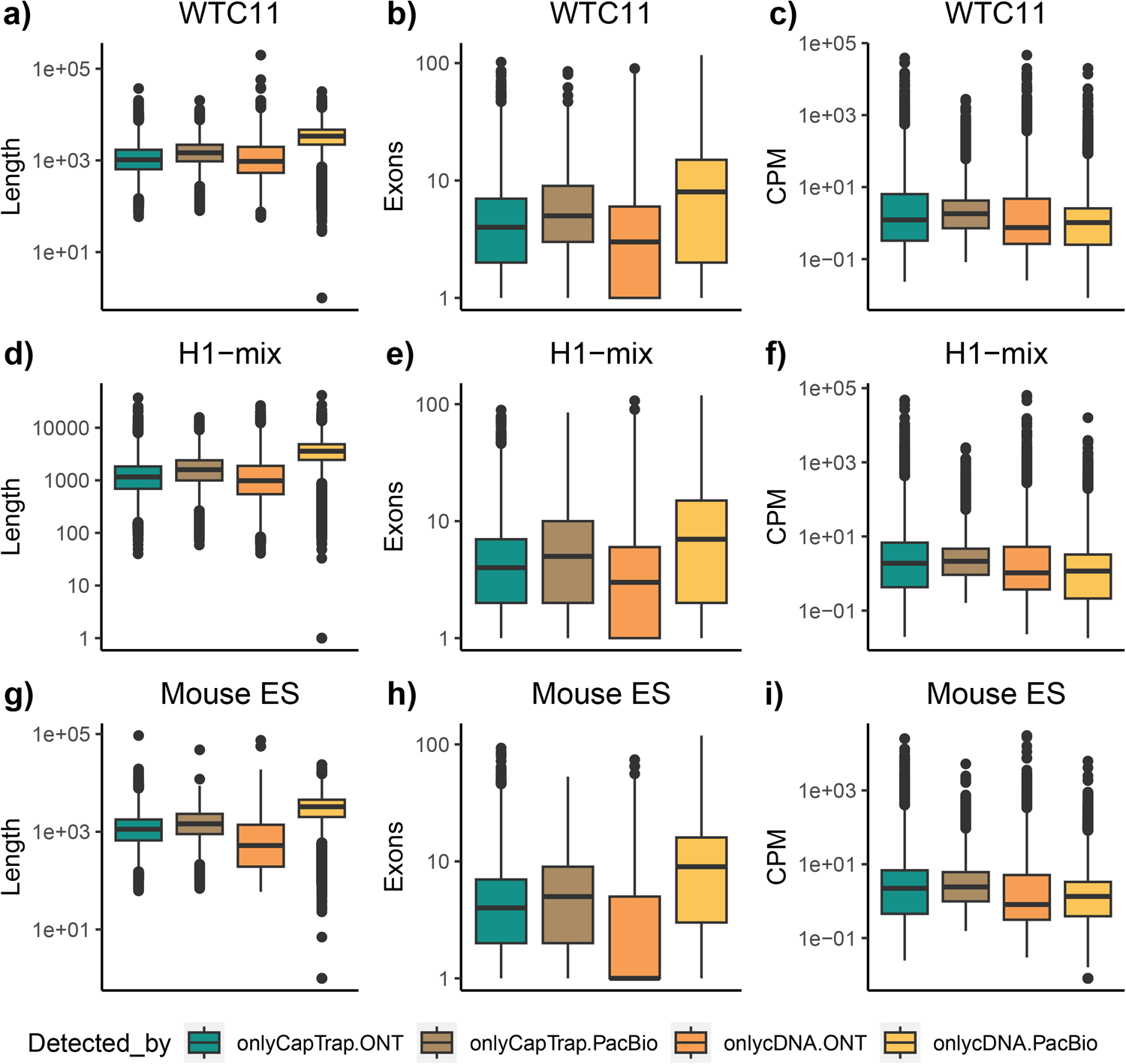
Properties of detected transcripts by experimental protocol. a,d,g) Length distribution. b,e,h) Exon number distribution. c,f,i) Counts per million

**Extended Data Fig. 41.**
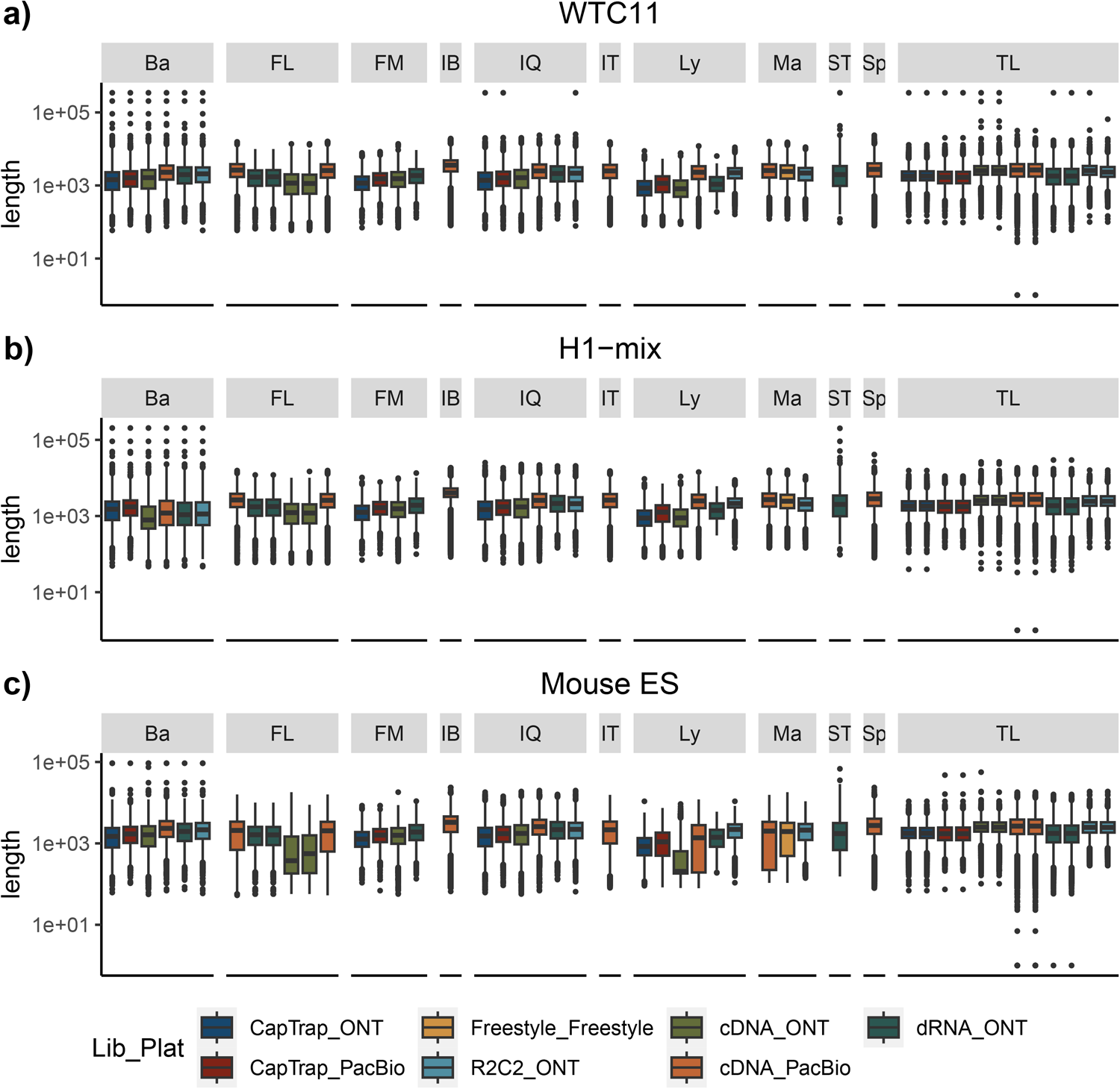
Distribution of transcript length by analysis tool.

**Extended Data Fig. 42a.**
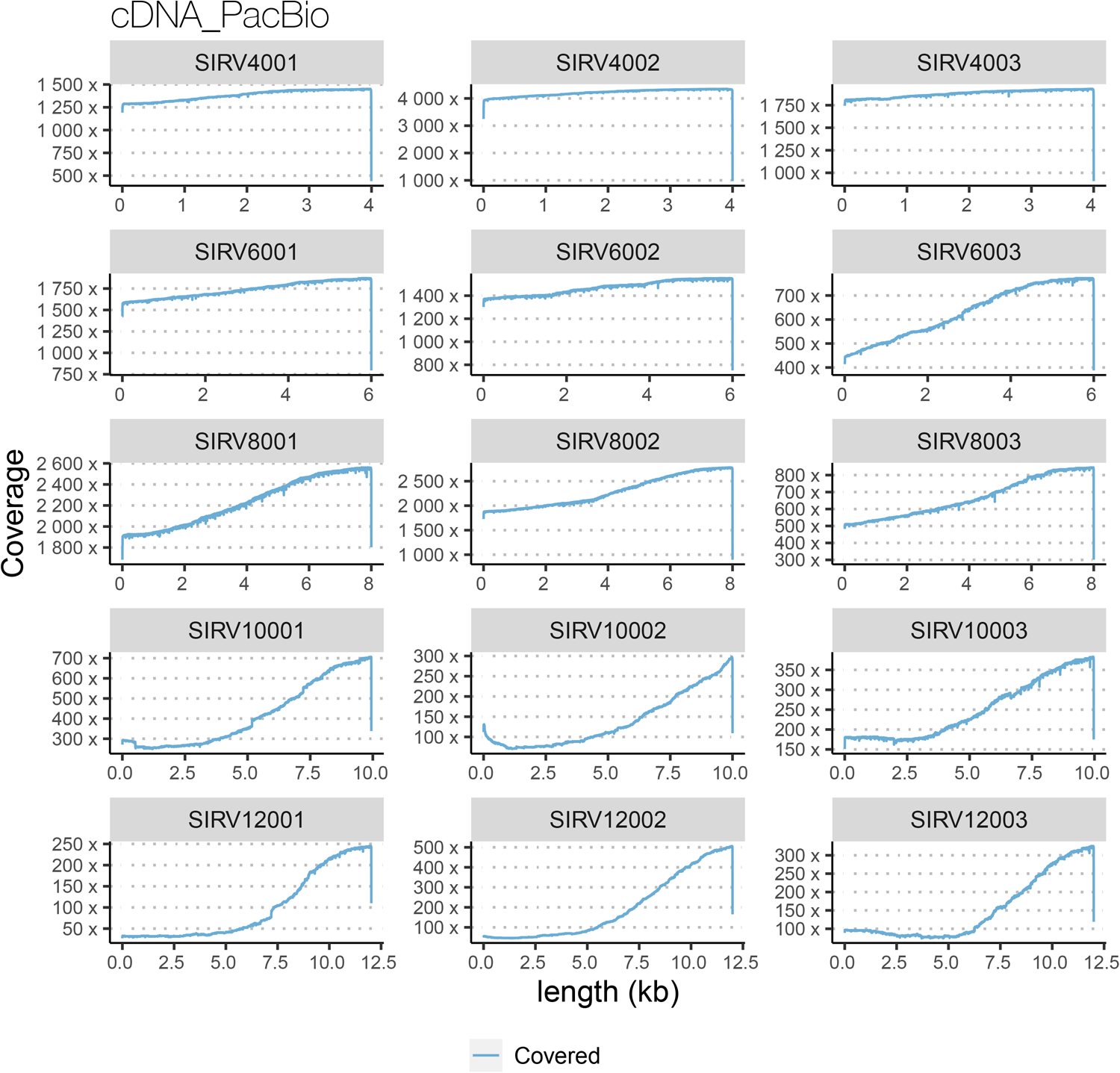
Positional coverage of SIRV transcript sequences by long reads in the cDNA_PacBio sample.

**Extended Data Fig. 42b.**
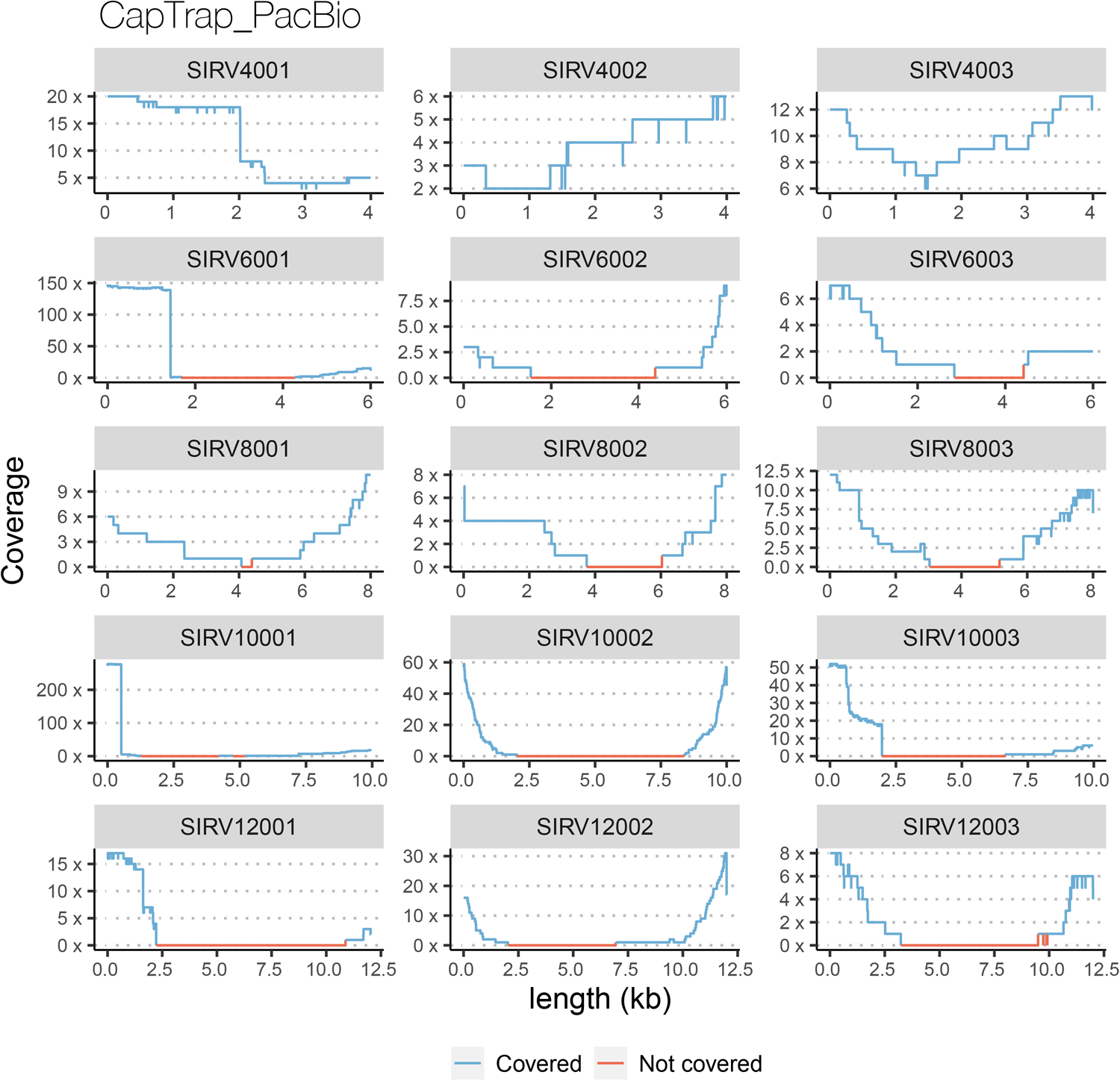
Positional coverage of SIRV transcript sequences by long reads in the CapTrap_PacBio

**Extended Data Fig. 42c.**
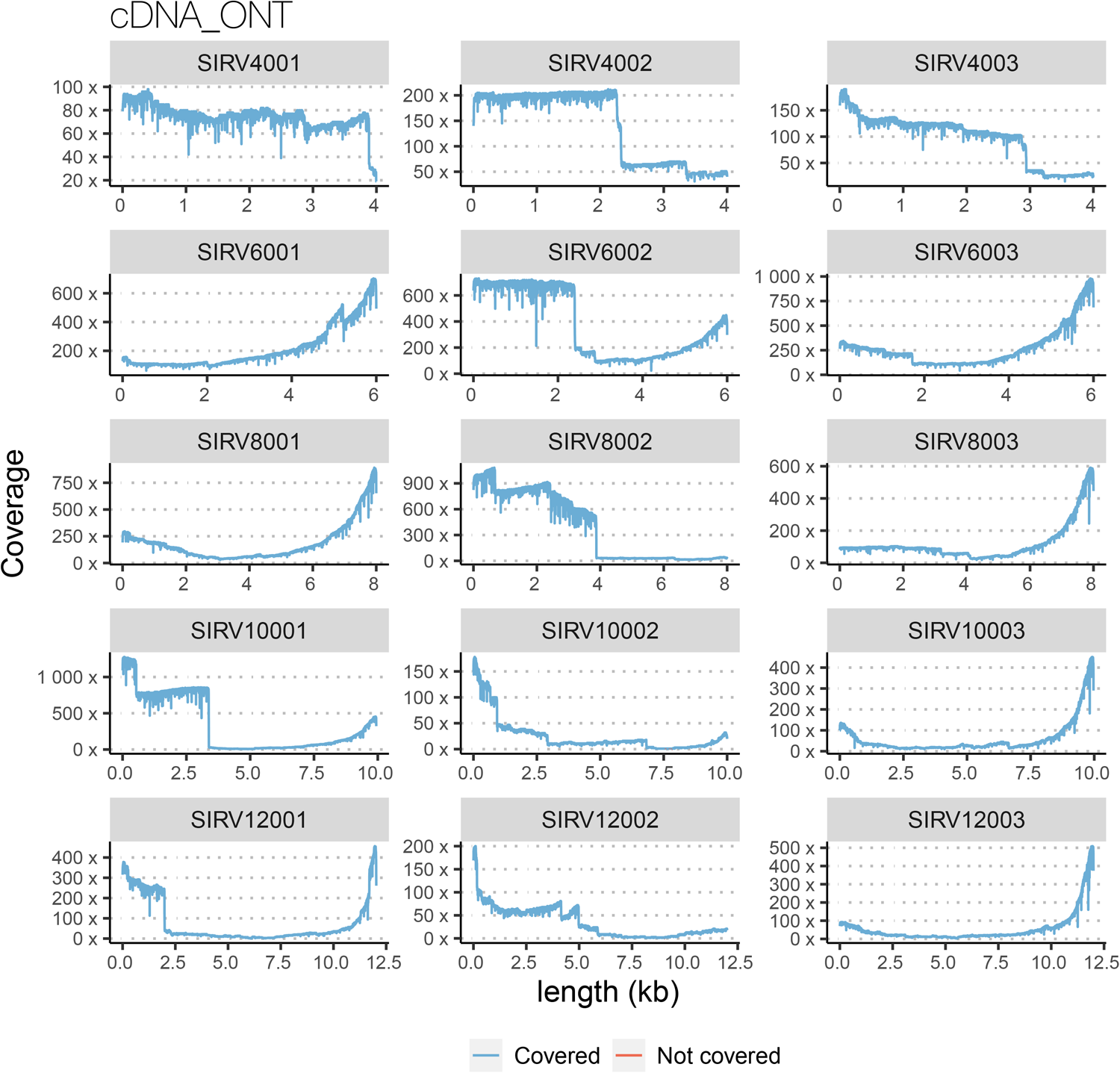
Positional coverage of SIRV transcript sequences by long reads in the cDNA_ONT sample.

**Extended Data Fig. 42d.**
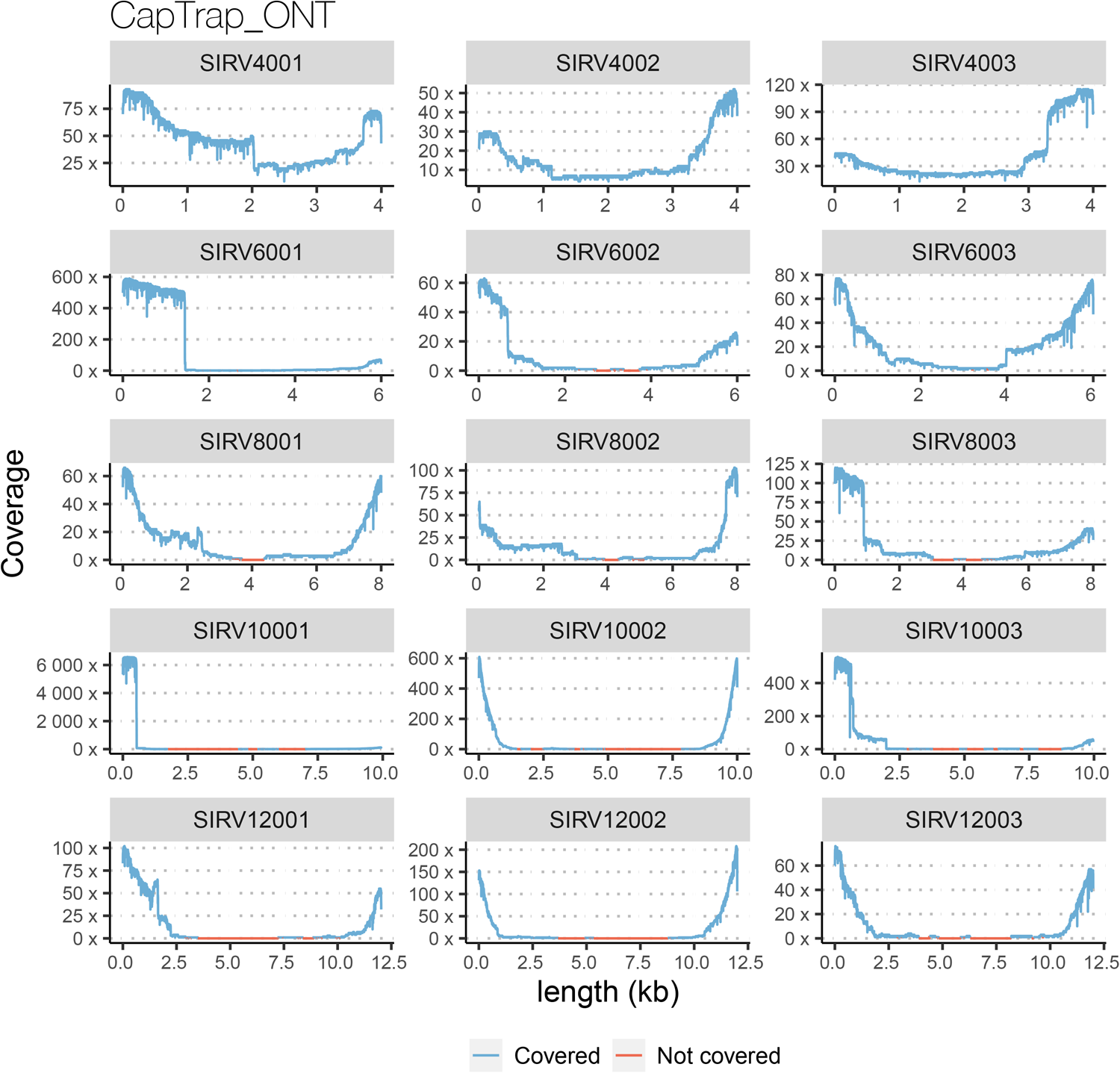
Positional coverage of SIRV transcript sequences by long reads in the CapTrap_ONT

**Extended Data Fig. 42e.**
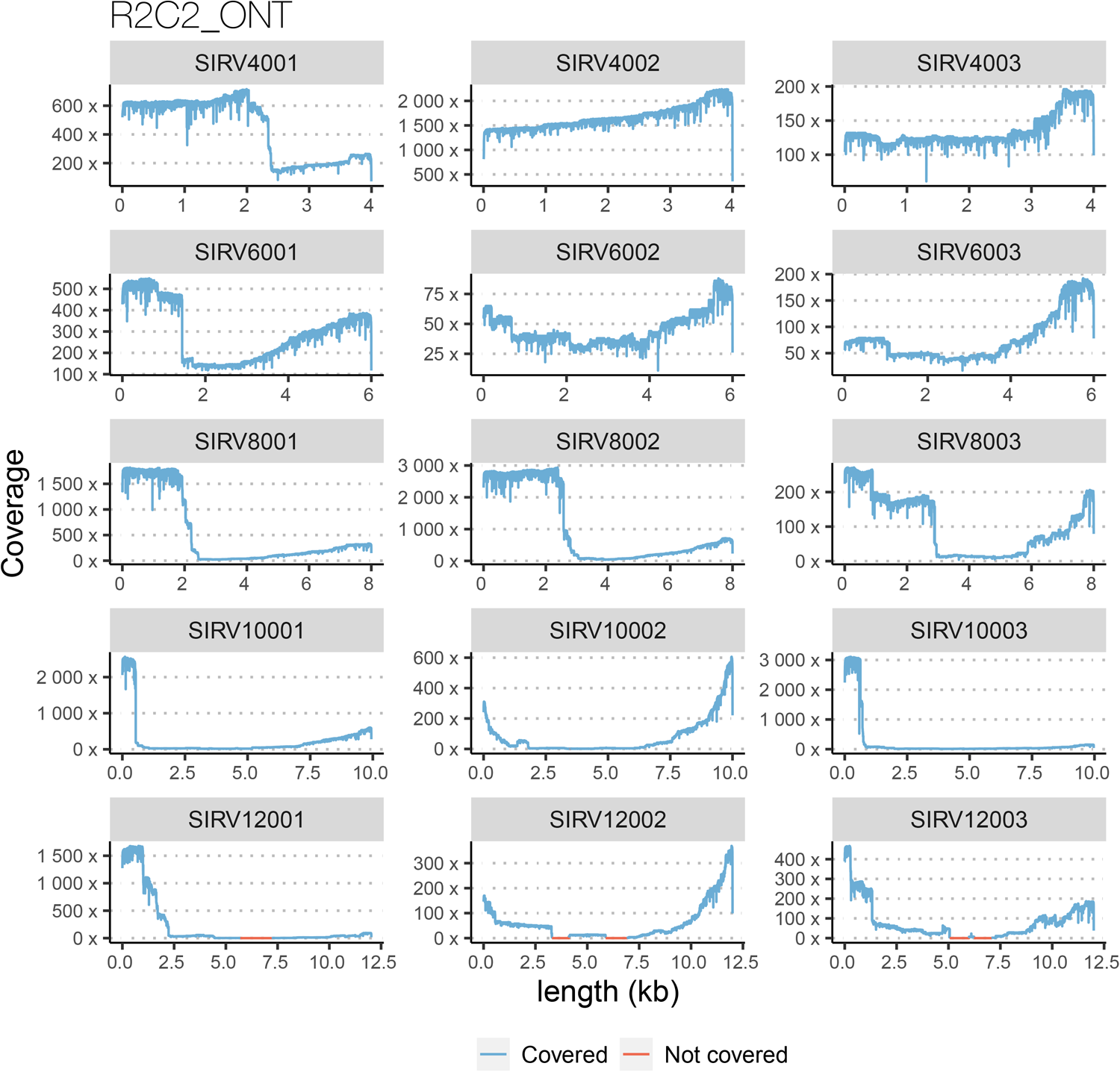
Positional coverage of SIRV transcript sequences by long reads in the R2C2_ONT sample.

**Extended Data Fig. 42f.**
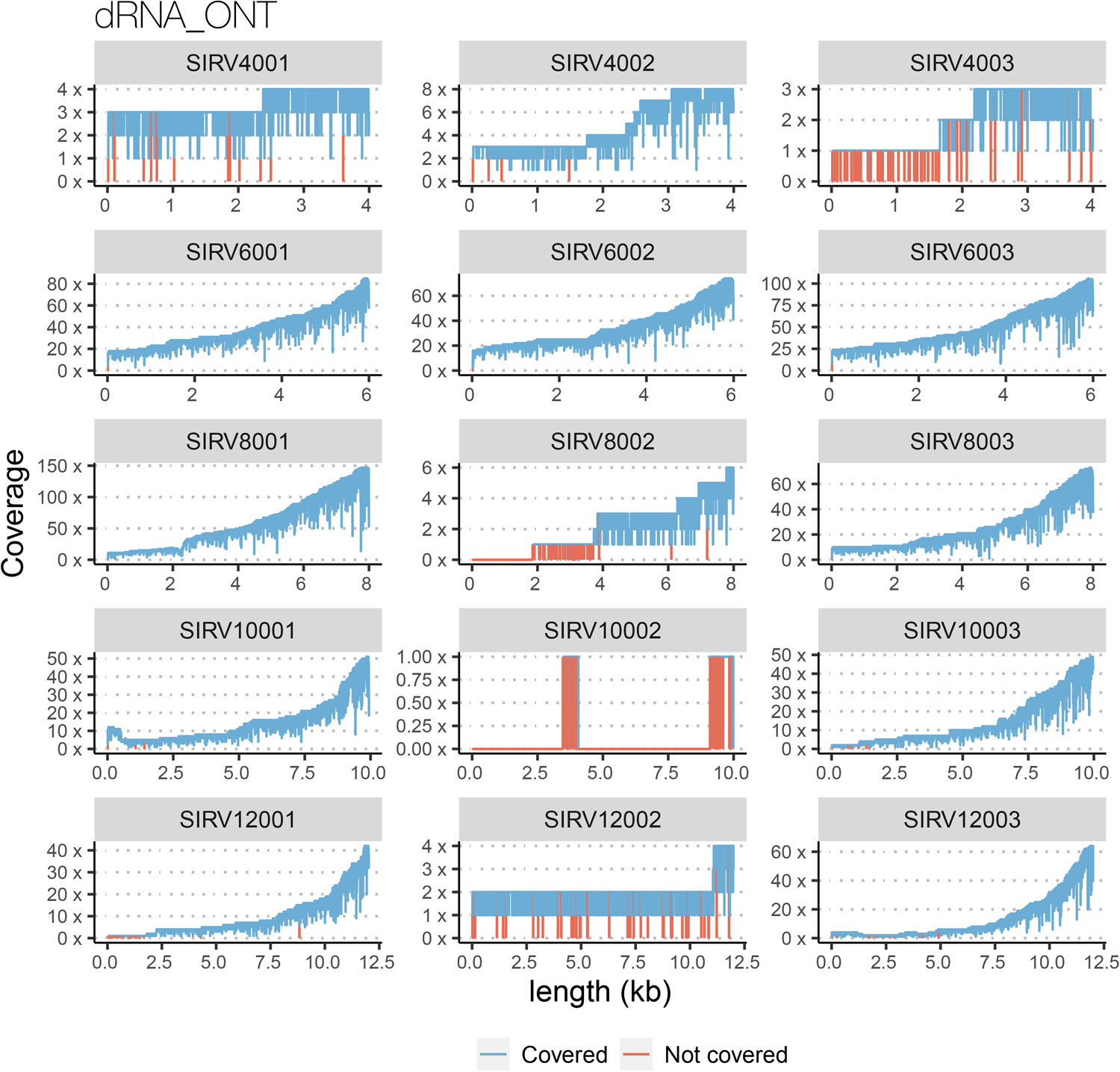
Positional coverage of SIRV transcript sequences by long reads in the dRNA_ONT sample.

**Extended Data Fig.43.**
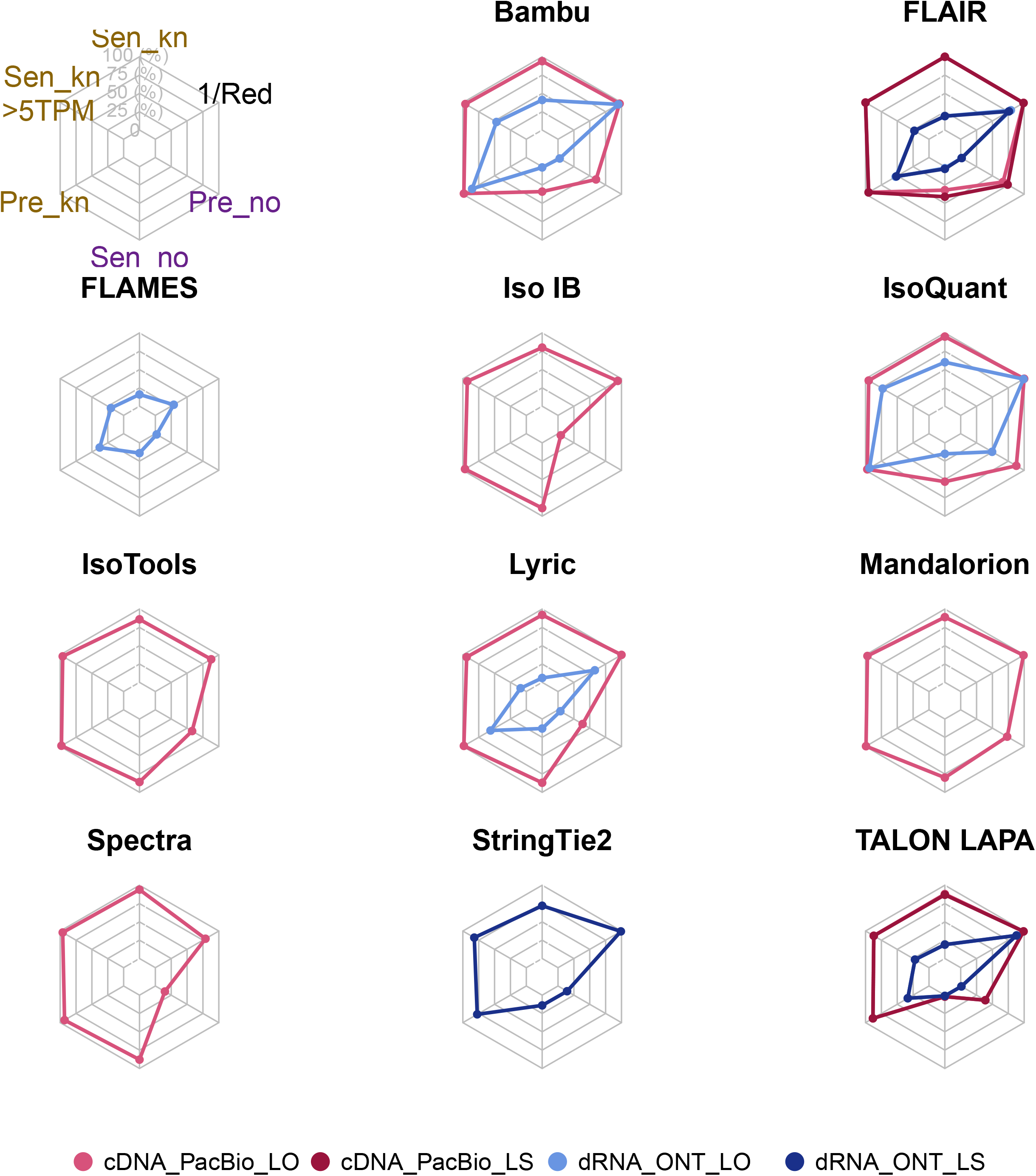
Performance metrics on mouse simulated data. Sen_kn: sensitivity known transcripts, Sen_kn > 5TMP: sensitivity known transcripts with expression > 5 TPM, Pre_kn: precision known transcripts, Sen_no: sensitivity novel transcripts, Pre_no: precision novel transcripts, 1/Red: inverse of redundancy.

**Extended Data Fig. 44.**
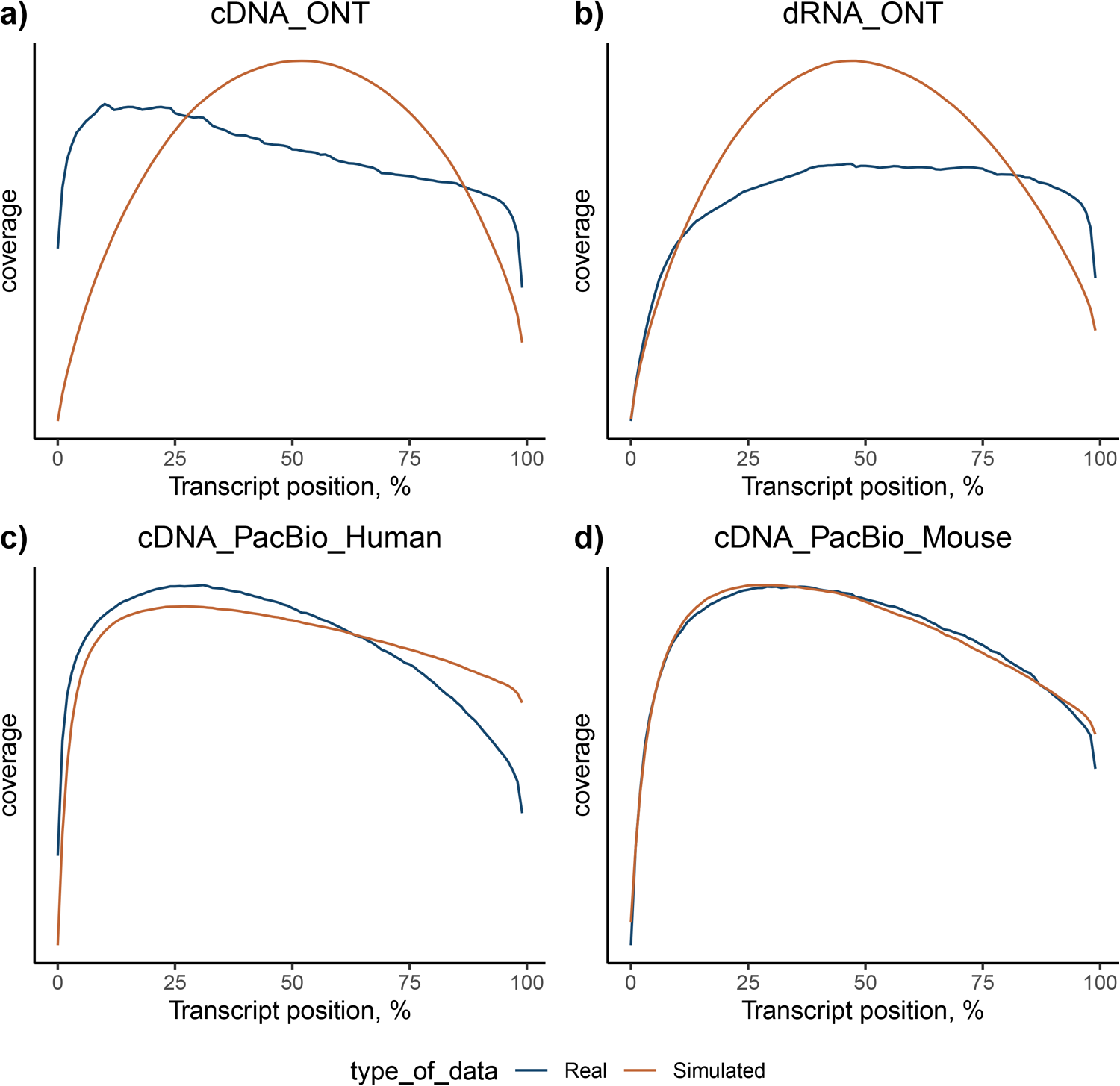
Comparison of long−read transcript coverage between real and simulated datasets.

**Extended Data Fig. 45.**
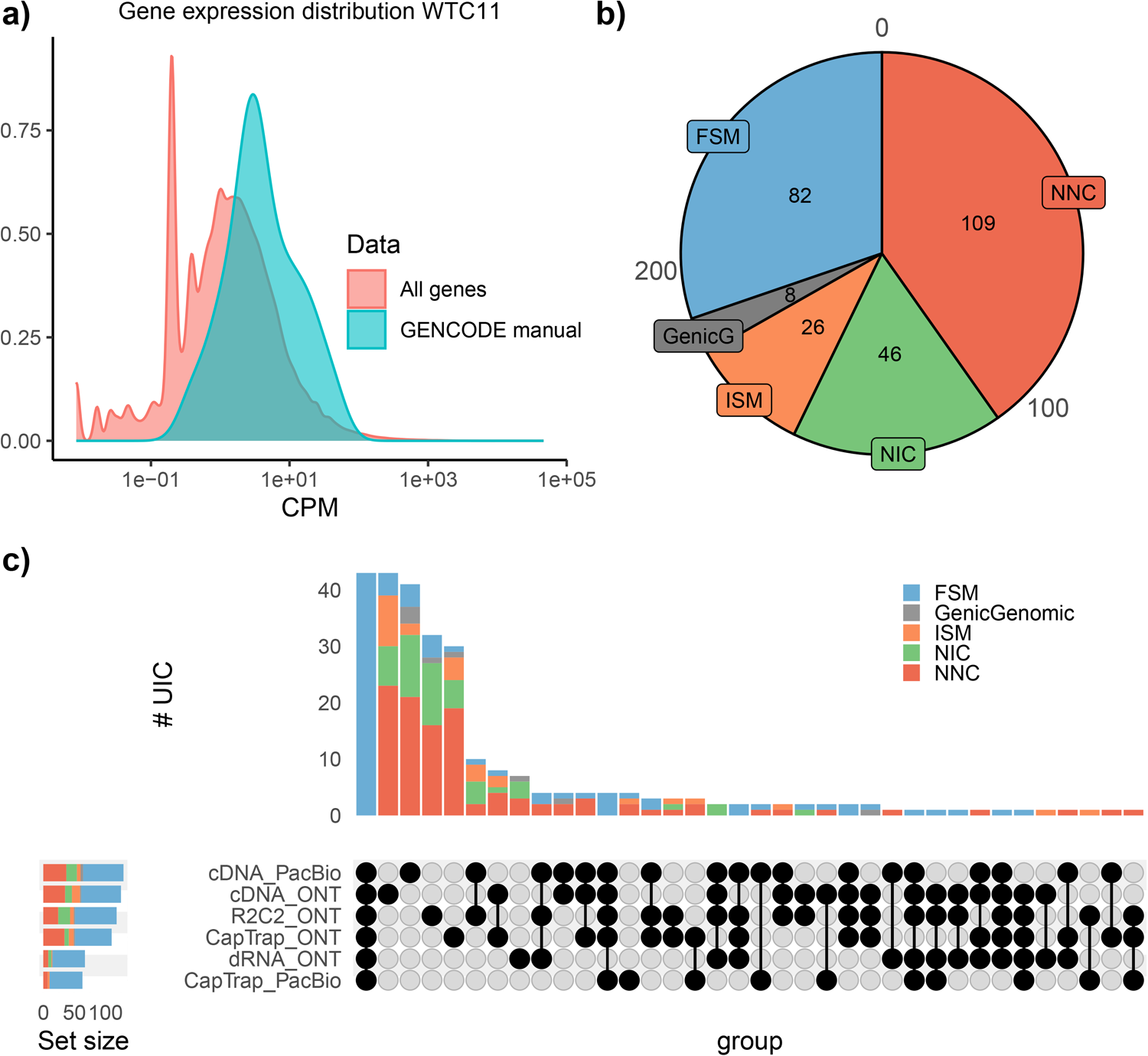
Properties of GENCODE manually annotated loci for WTC11 sample.a) Distributon of gene expression. b) Distribution of SQANTI categories. c) Intersection of Unique Intron Chains (UIC) among experimental protocols.

**Extended Data Fig. 46.**
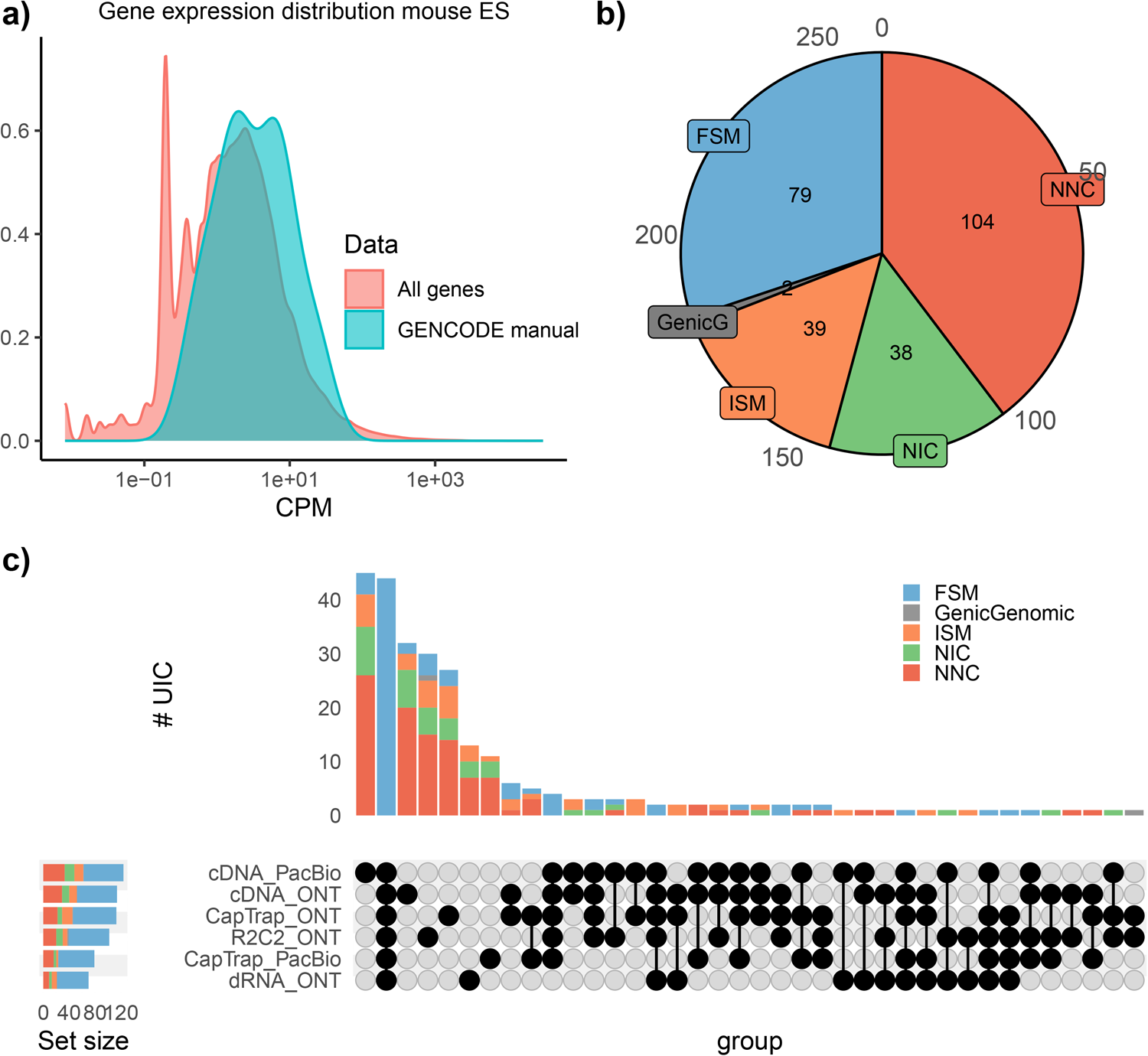
Properties of GENCODE manually annotated loci for mouse ES sample.a) Distributon of gene expression. b) Distribution of SQANTI categories. c) Intersection of Unique Intron Chains (UIC) among experimental protocols.

**Extended Data Fig. 47.**
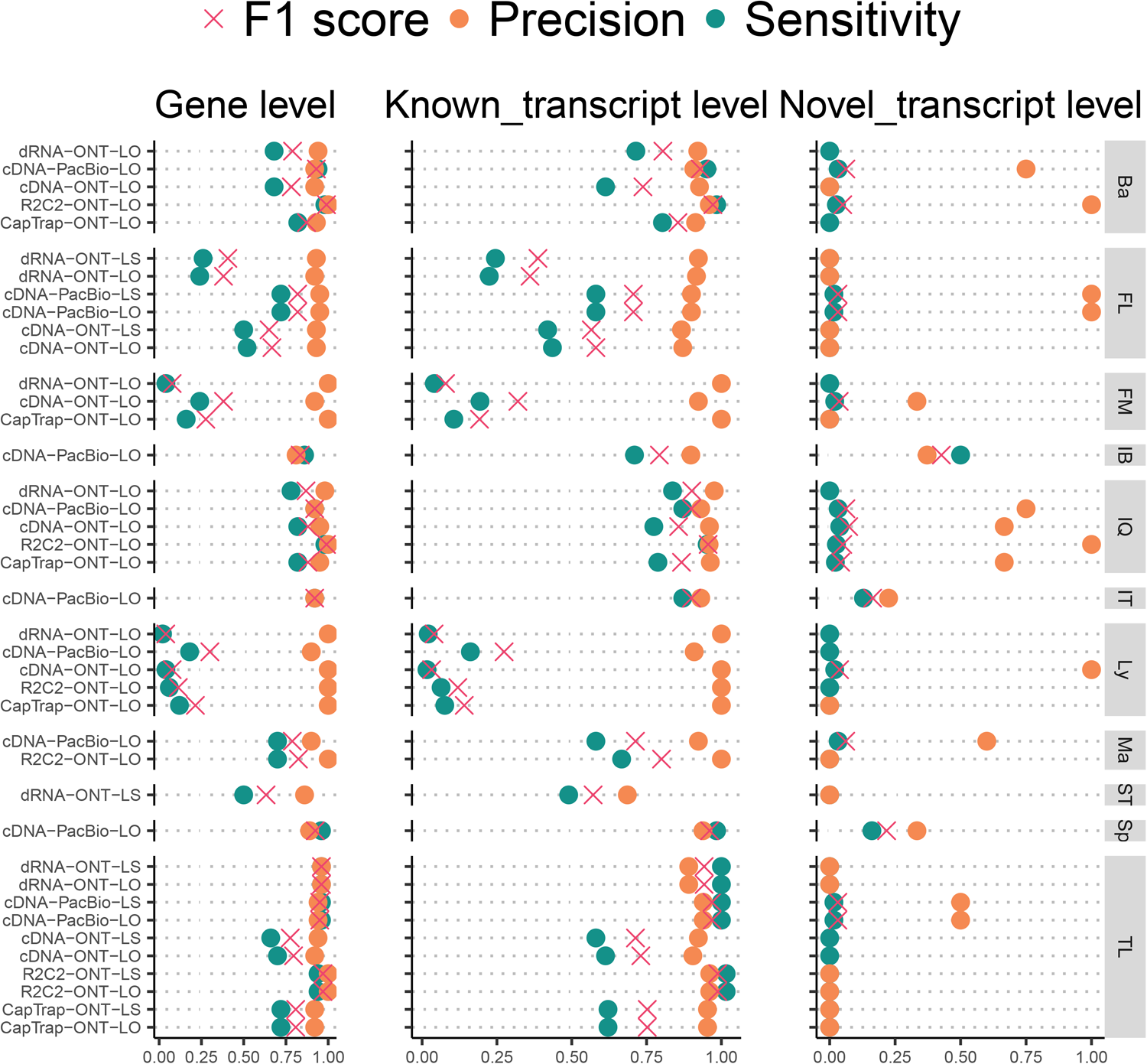
Performance metrics of LRGASP pipelines evaluate against GENCODE manual annotation of mouse ES sample. Ba: Bambu, FM: Flames, FR: FLAIR, IQ: IsoQuant, IT: IsoTools, IB: Iso_IB, Ly: LyRic, Ma: Mandalorion, TL: TALON−LAPA, Sp: Spectra, ST: StringTie2.

**Extended Data Fig. 48.**
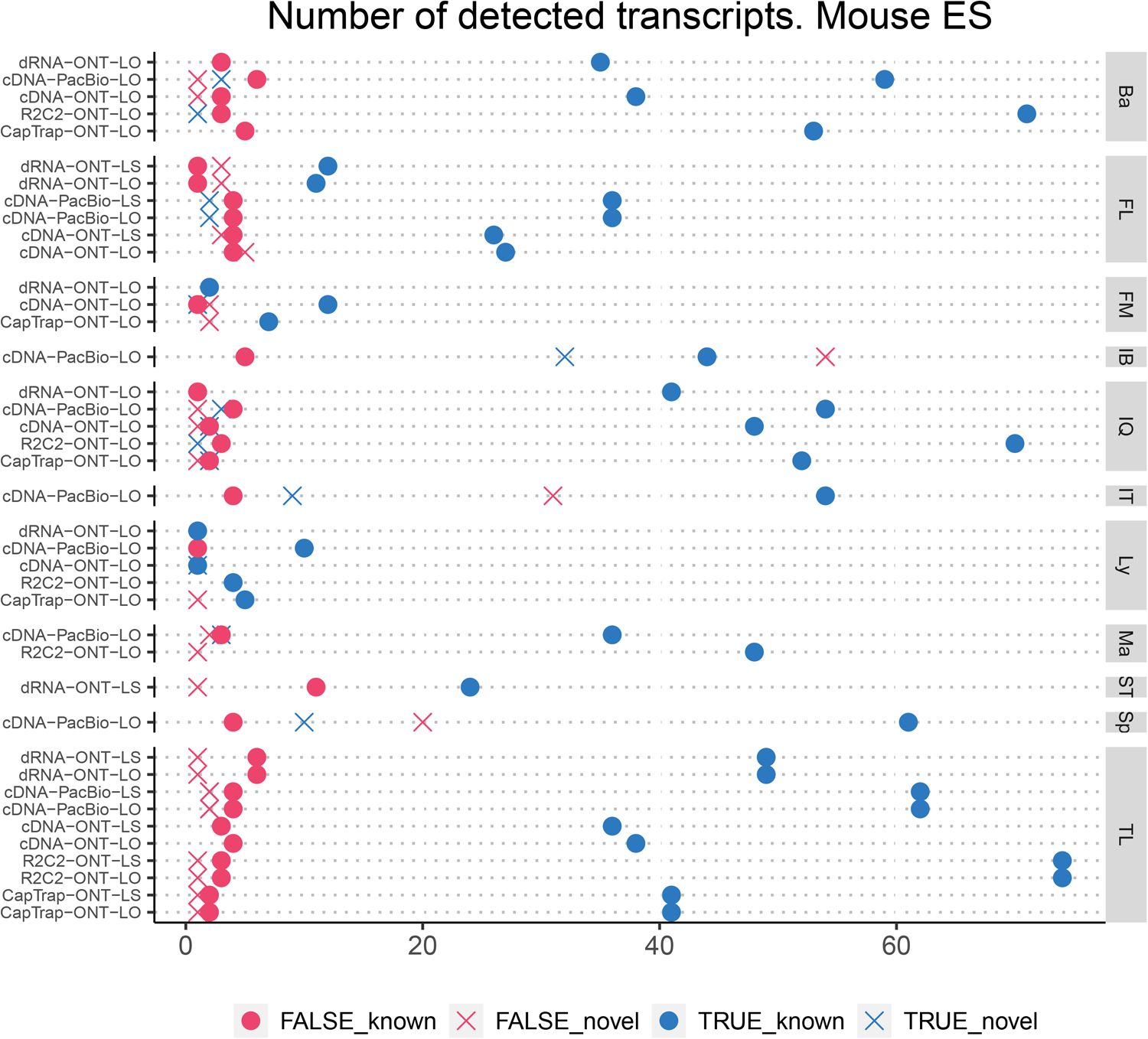
Detection of Unique Intron Chains (UIC) at GENCODE manual annotation loci. Ba: Bambu, FM: Flames, FL: FLAIR, IQ: IsoQuant, IT: IsoTools, IB: Iso_IB, Ly: LyRic, Ma: Mandalorion, TL: TALON−LAPA, Sp: Spectra, ST: StringTie2.

**Extended Data Fig. 49.**
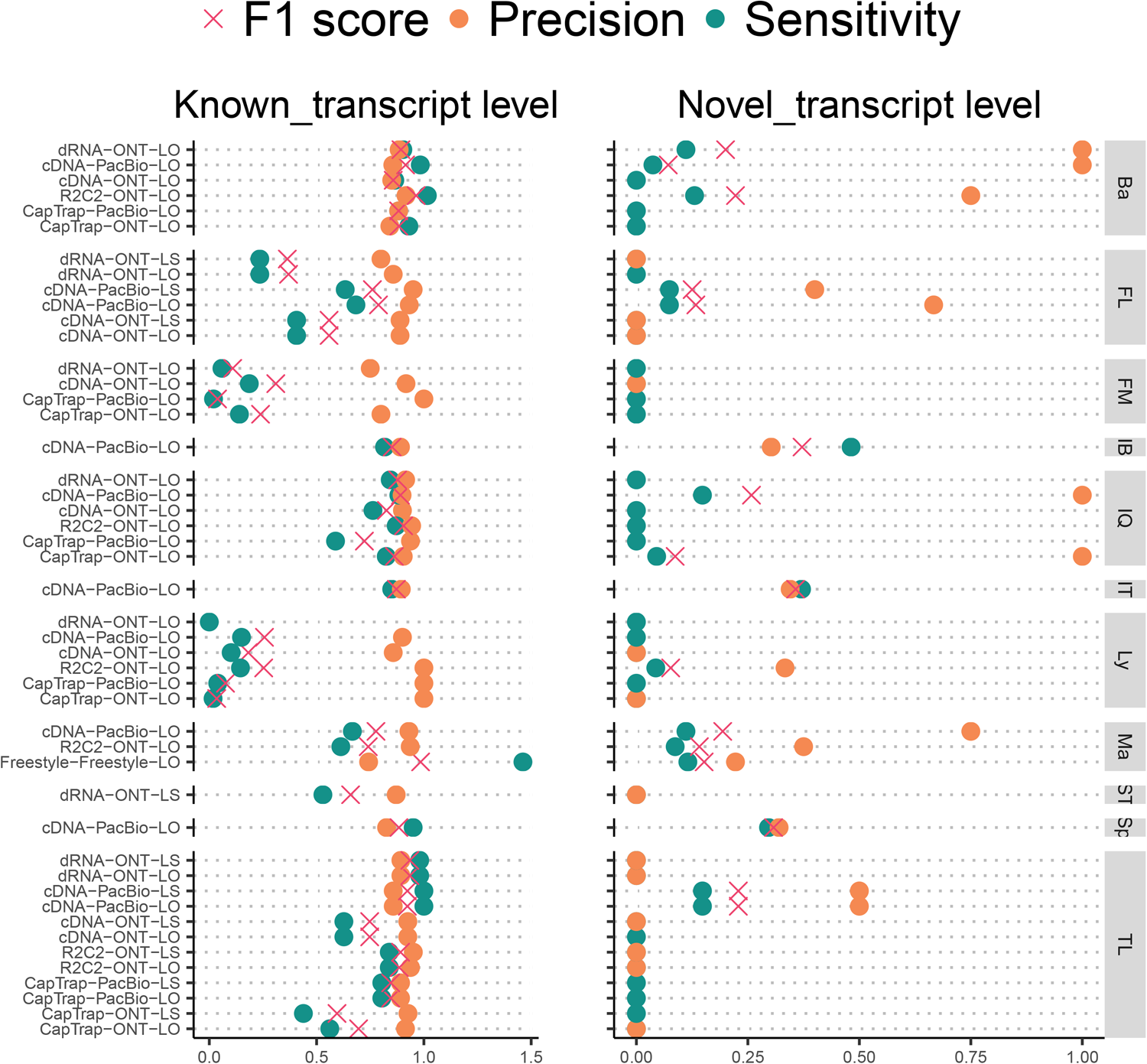
Performance on GENCODE manually curated data. Curated transcripts selected to be present in at least two experimental datasets. Ba: Bambu, FM: Flames, FL: FLAIR, IQ: IsoQuant, IT: IsoTools, IB: Iso_IB,

**Extended Data Fig. 50.**
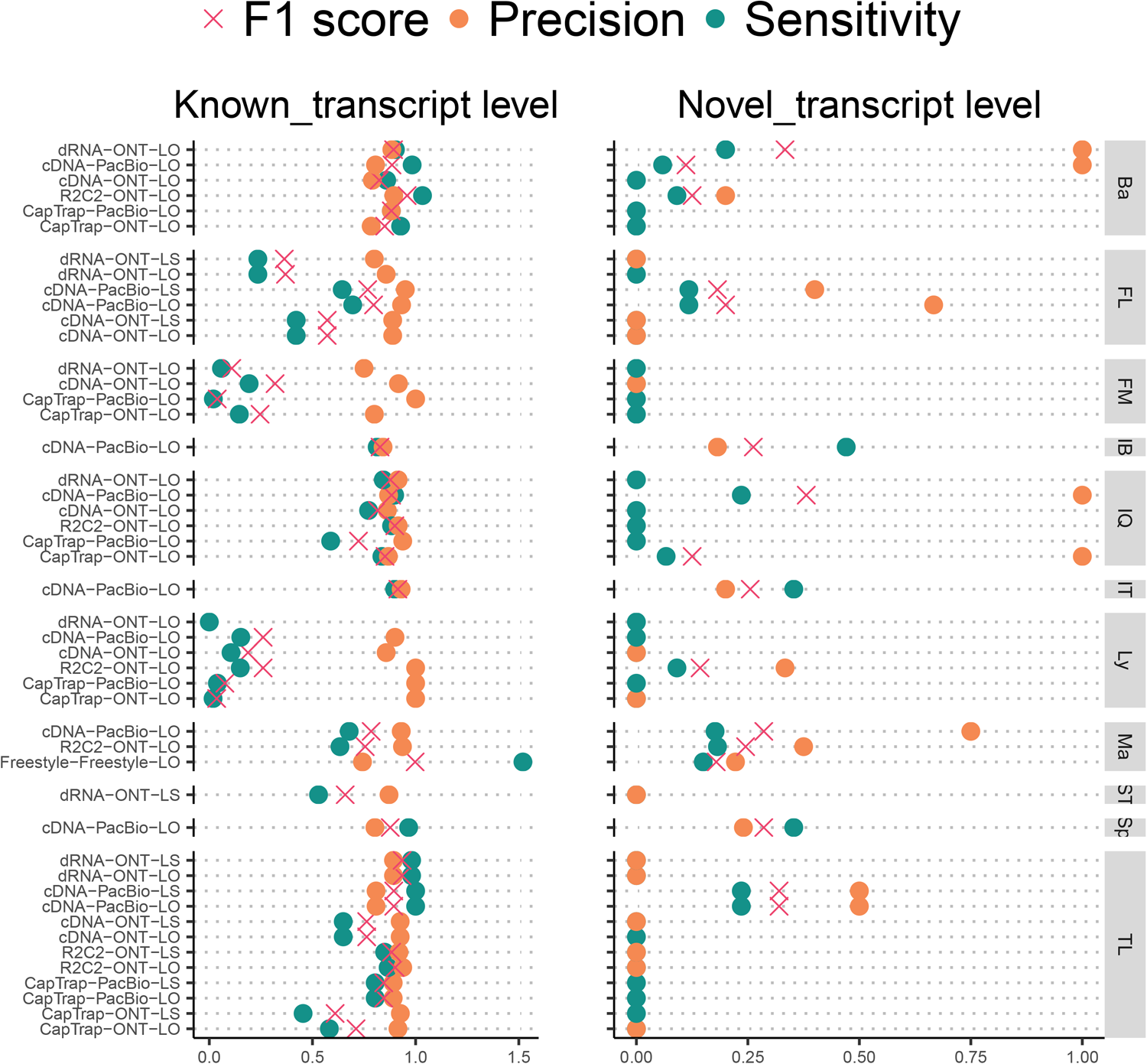
Performance on GENCODE manually curated data. The ground truth is the set of manually annotated transcripts with more than two reads. Ba: Bambu, FM: Flames, FL: FLAIR, IQ: IsoQuant, IT: IsoTools,

**Extended Data Fig. 51.**
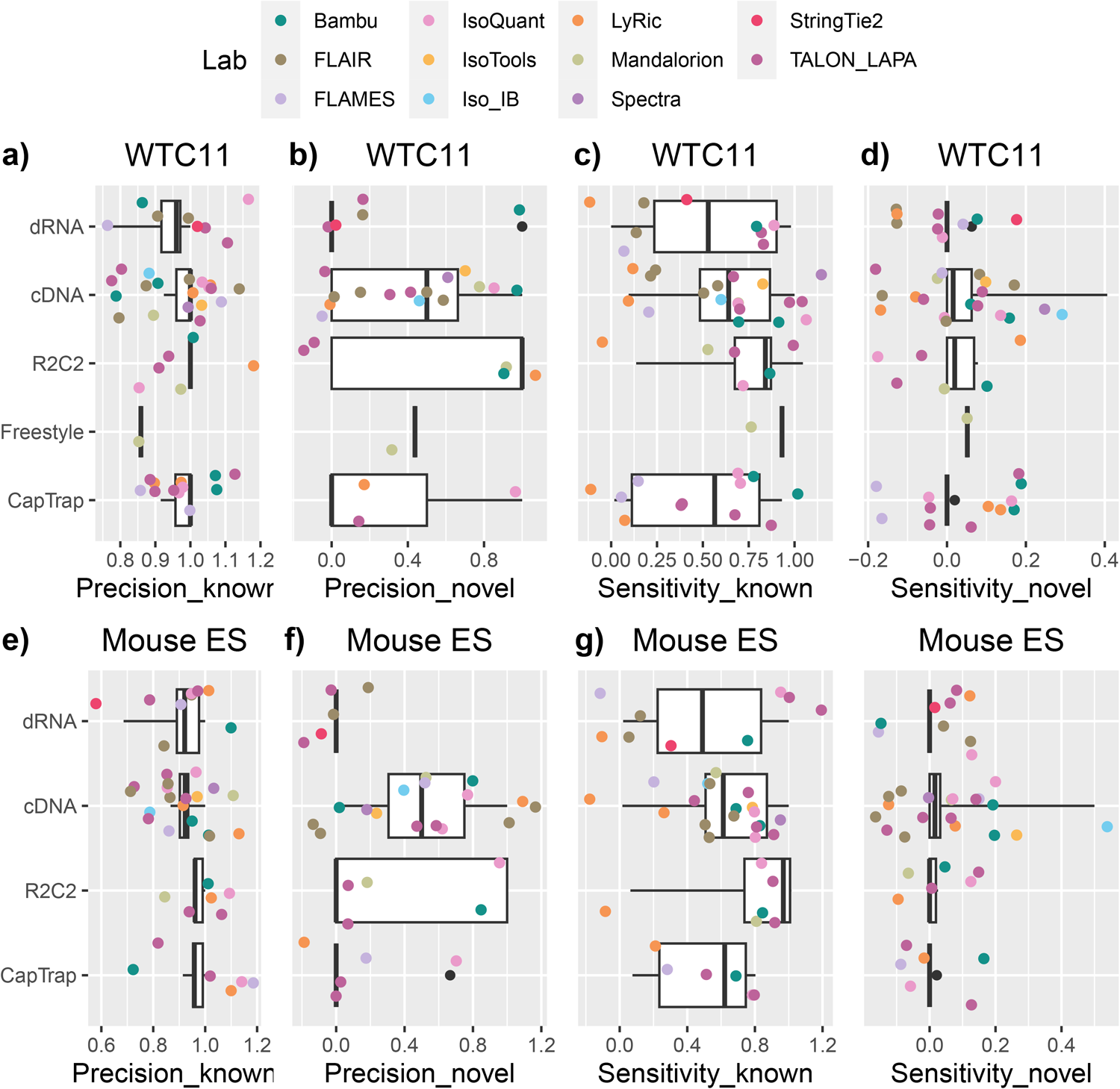
Performance on GENCODE manually curated data by Library Preparation.

**Extended Data Fig. 52.**
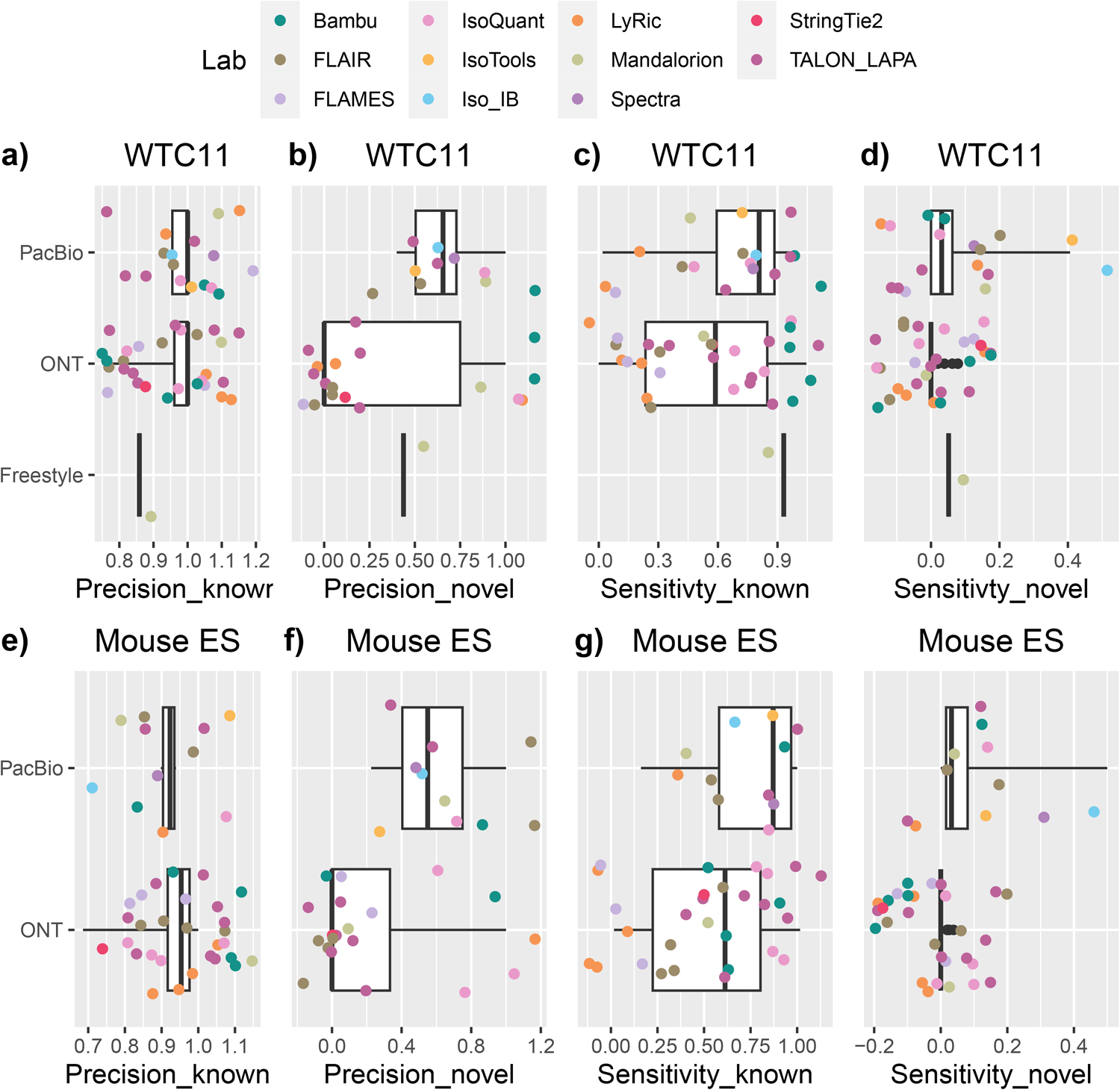
Performance on GENCODE manually curated data by Platform.

**Extended Data Fig. 53.**
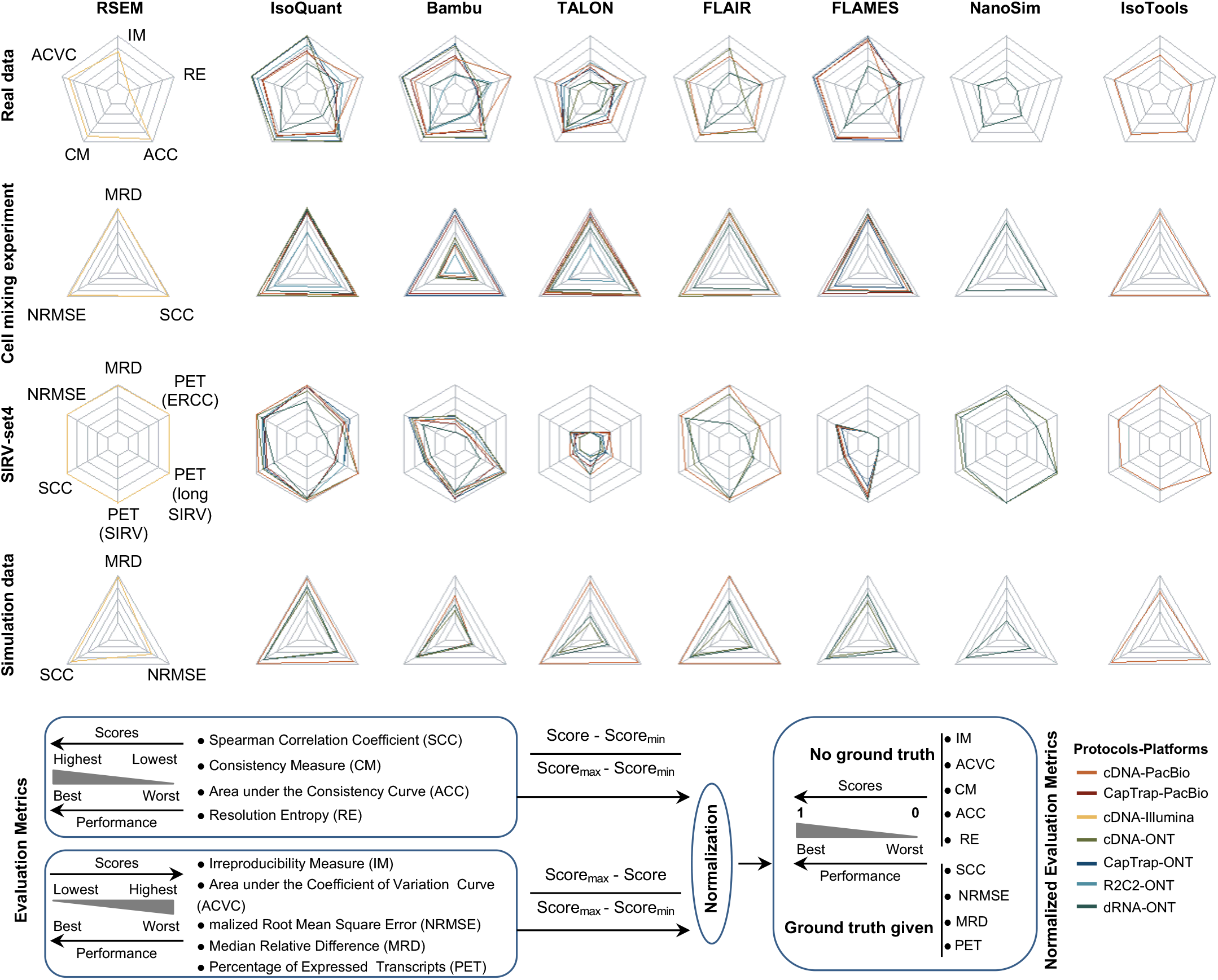
Radar plot of overall evaluation results of 8 quantification tools with 7 protocols-platforms on 4 data scenarios: real data with multiple replicates, cell mixing experiment, SIRV-set4 data and simulation data. To display the evaluation results more effectively, we normalized all metrics to 0-1 range: 0 corresponds to the worst performance and 1 corresponds to the best performance.

**Extended Data Fig. 54.**
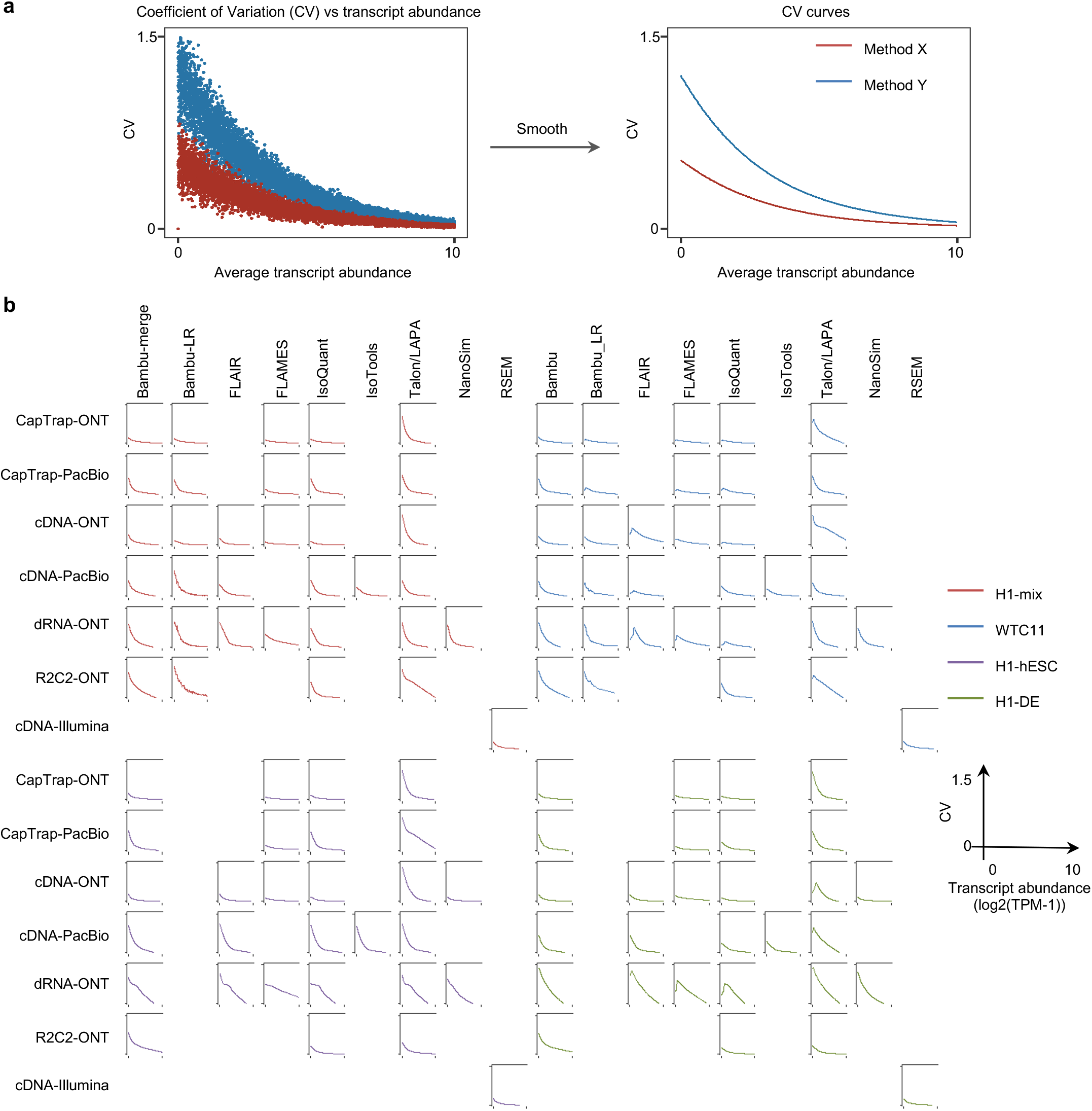
Overall evaluation results of irreproducibility on real data with multiple replicates. (a) The diagram illustrates the calculation of irreproducibility. By fitting the <u>c</u>oefficient of <u>v</u>ariation (CV) versus average transcript abundance into a smooth curve, it can be shown that Method X has lower coefficient of variation and higher reproducibility. (b) The overall results of CV curves with different transcript abundances on four samples (H1-mix, WTC11, H1-hESC and H1-DE) with different protocols and platforms. Here, Bambu-merge represents the transcript quantification using Bambu with GENCODE plus LR-specific annotation. And Bambu-LR represents the transcript quantification using only LR-specific annotation.

**Extended Data Fig. 55.**
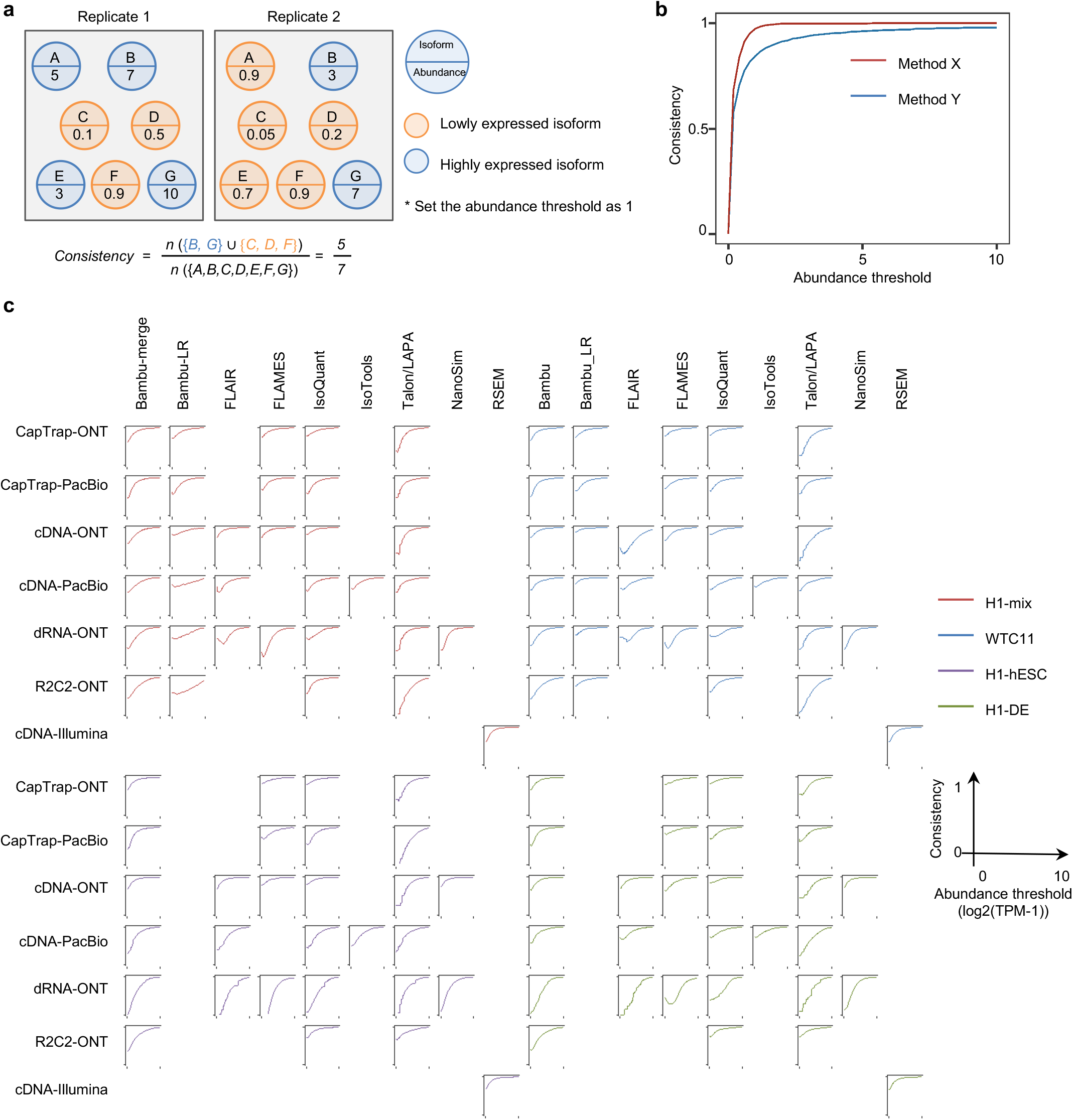
Overall evaluation results of consistency on real data with multiple replicates. (a) The diagram illustrates the calculation of consistency. By setting an expression threshold (i.e. 1 in this toy example), we can define which set of transcripts express (in blue) or not (in orange). This statistic is to measure the consistency of the expressed transcripts sets between replicates. (b) A toy example to show the consistency curves with different abundance threshold. Here, method X performs the better consistency of transcript abundance estimation across multiple replicates than method Y. (c) The detailed evaluation results of consistency curves with different abundance thresholds on four samples (H1-mix, WTC11, H1-hESC and H1-DE) with different protocols and platforms.

**Extended Data Fig. 56.**
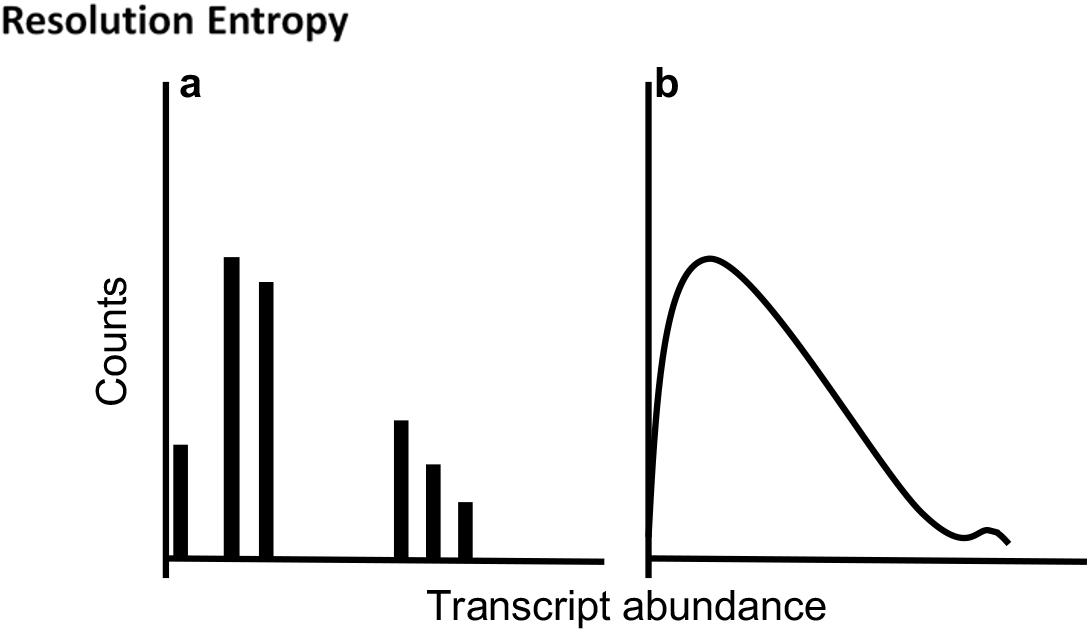
Resolution Entropy. (a) The software output only a few certain discrete values has lower resolution entropy as it cannot capture the continuous and subtle difference of gene expressions. (b) The software with continuous output values has higher resolution entropy

**Extended Data Fig. 57.**
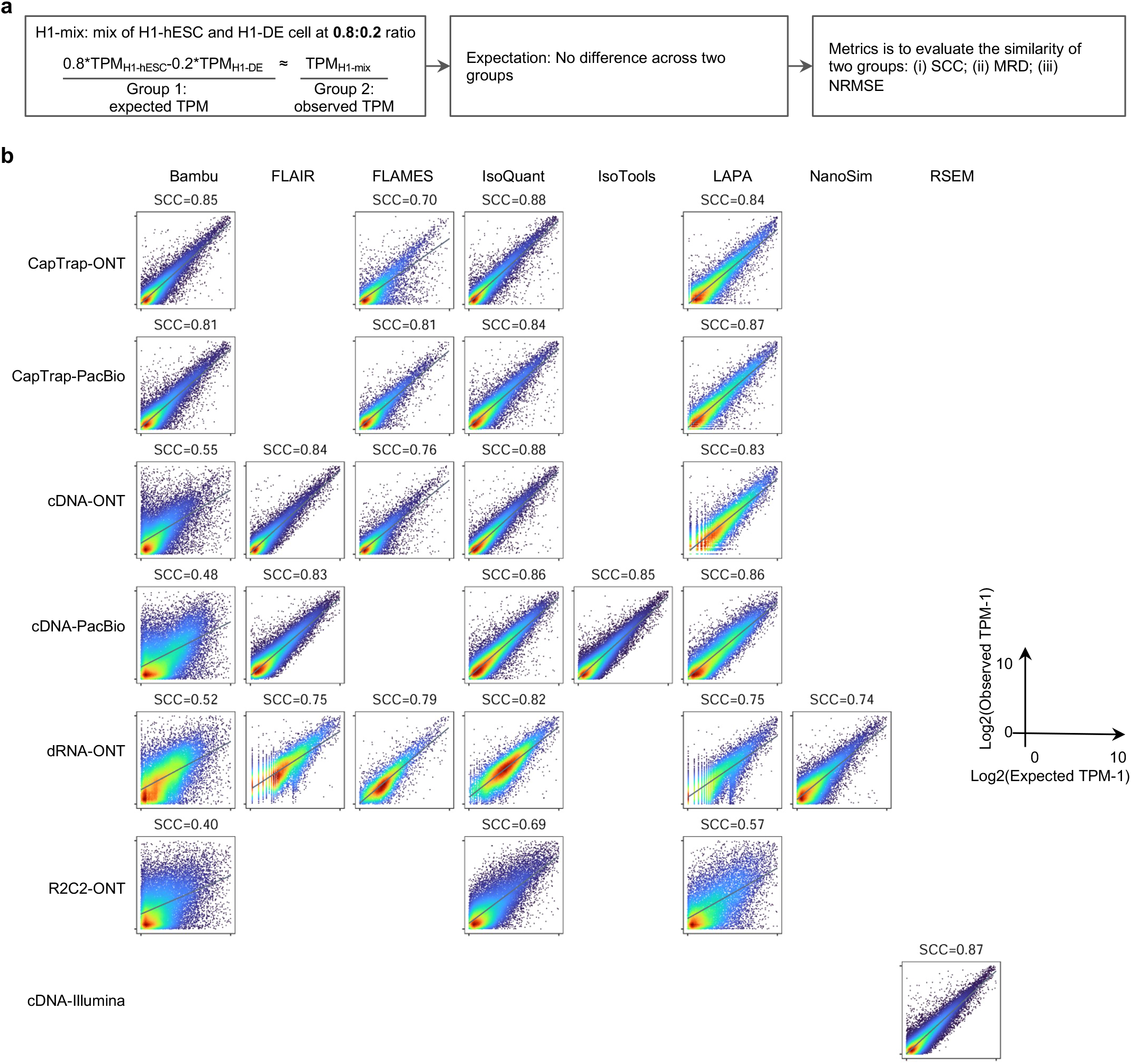
Performance evaluation on cell mixing experiment. (a) Schematic diagram of evaluation strategy using the cell mixing experiment. Here, H1-mix was initially provided for quantification which was a mix of H1-hESC cells and H1-DE cells at an undisclosed ratio. After the initial submission, the individual H1-hESC and H1-DE samples were released and participants submitted quantifications for each. (b) Scatter plot of expected abundance and observed abundance for 7 participant’s tools with different protocols and platforms.

**Extended Data Fig. 58.**
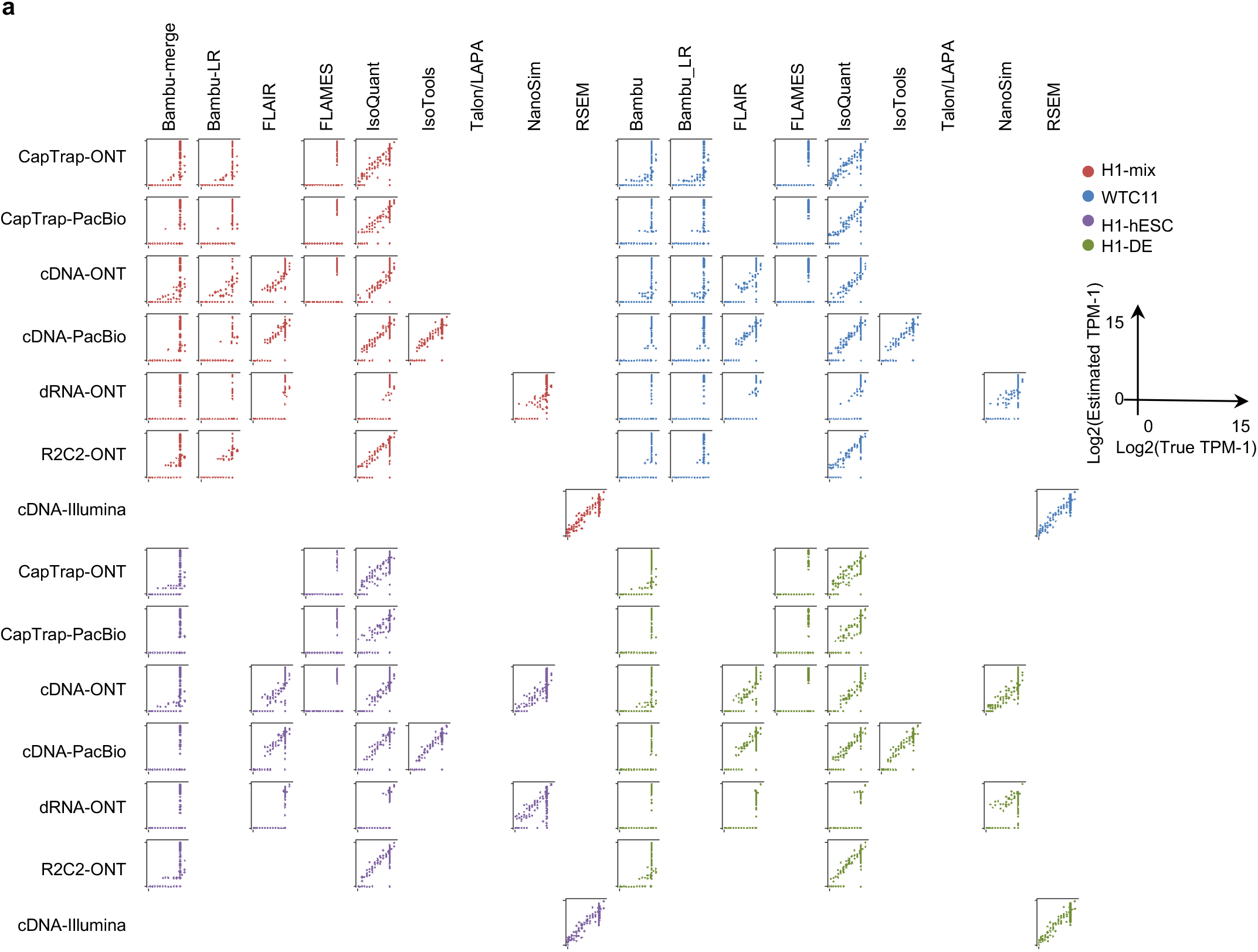
Performance evaluation on SIRV-set4 data. (a) Scatter plot of true abundance and estimated abundance on SIRV-set4 data with different protocols and platforms.

**Extended Data Fig. 59.**
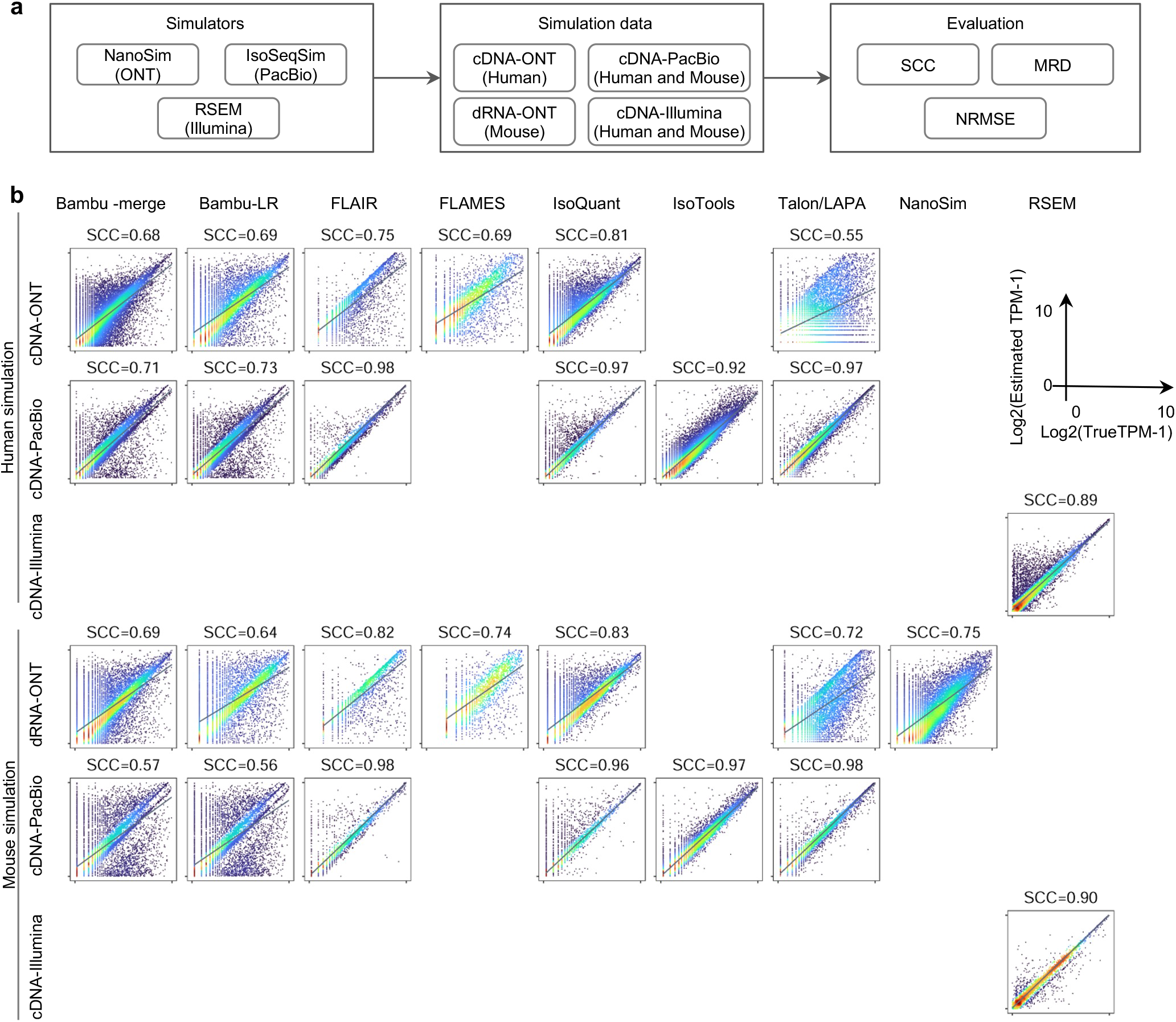
Performance evaluation on simulation data. (a) The flow chart of simulation study. (b) Scatter plot of true abundance and estimated abundance on simulation data.

**Extended Data Fig. 60.**
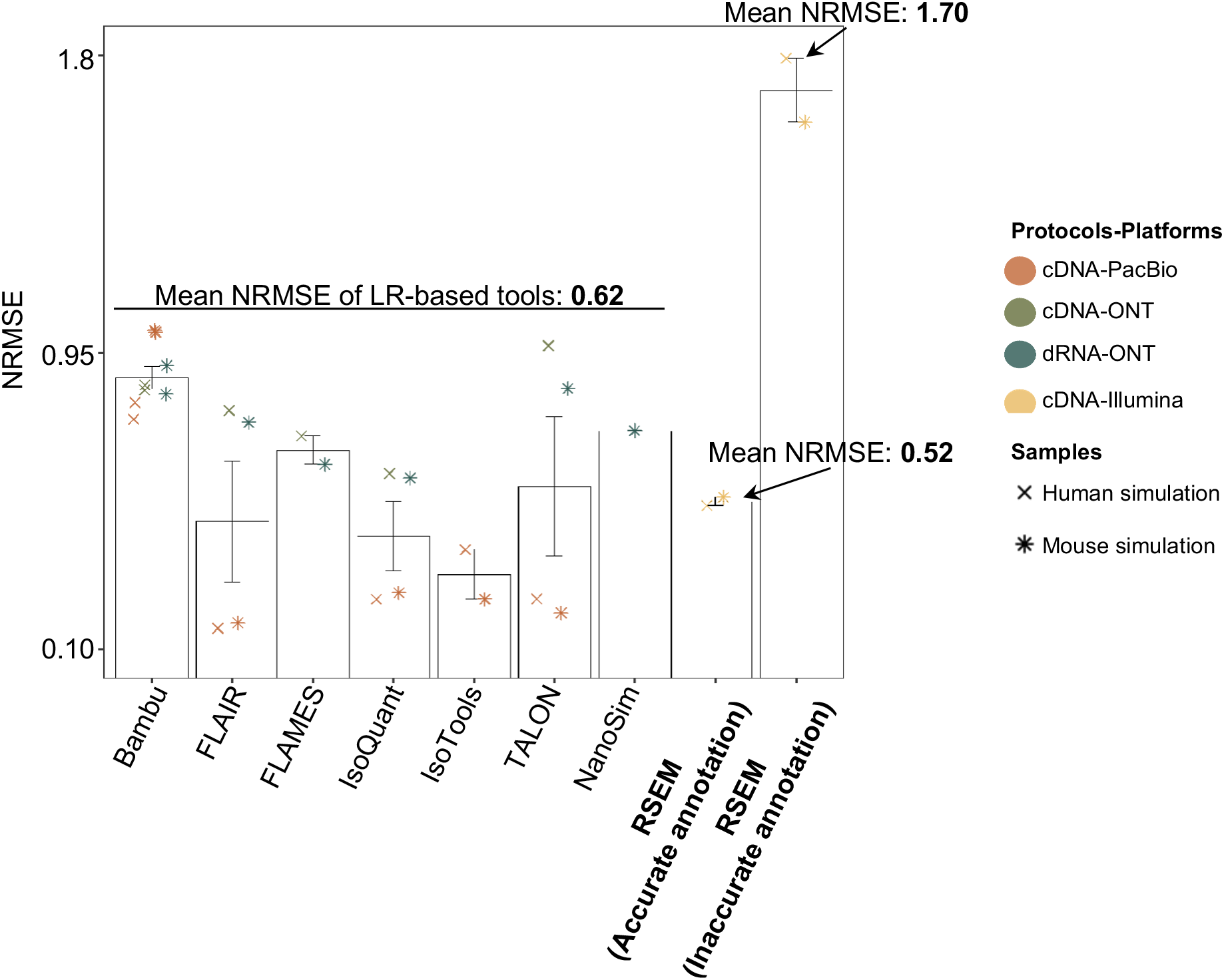
Impact of annotation accuracy on transcript quantification. We assessed the performance of RSEM and LR-based tools (Bambu, FLAIR, FLAMES, IsoQuant, IsoTools, TALON, and NanoSim) with different annotations. The NRMSE metric was used to evaluate their performance on simulated data for human and mouse. For LR-based tools, the transcript quantification annotations were derived from sample-specific annotations identified by the participant using long-read RNA-seq data. As for RSEM, we present quantification results based on two annotations: a completely accurate annotation (i.e., the ground truth transcripts generated by the simulation data) and an inaccurate annotation (i.e., the common GENCODE reference annotation, which contains numerous false negative and false positive transcripts specific to the sample).

**Extended Data Fig. 61.**
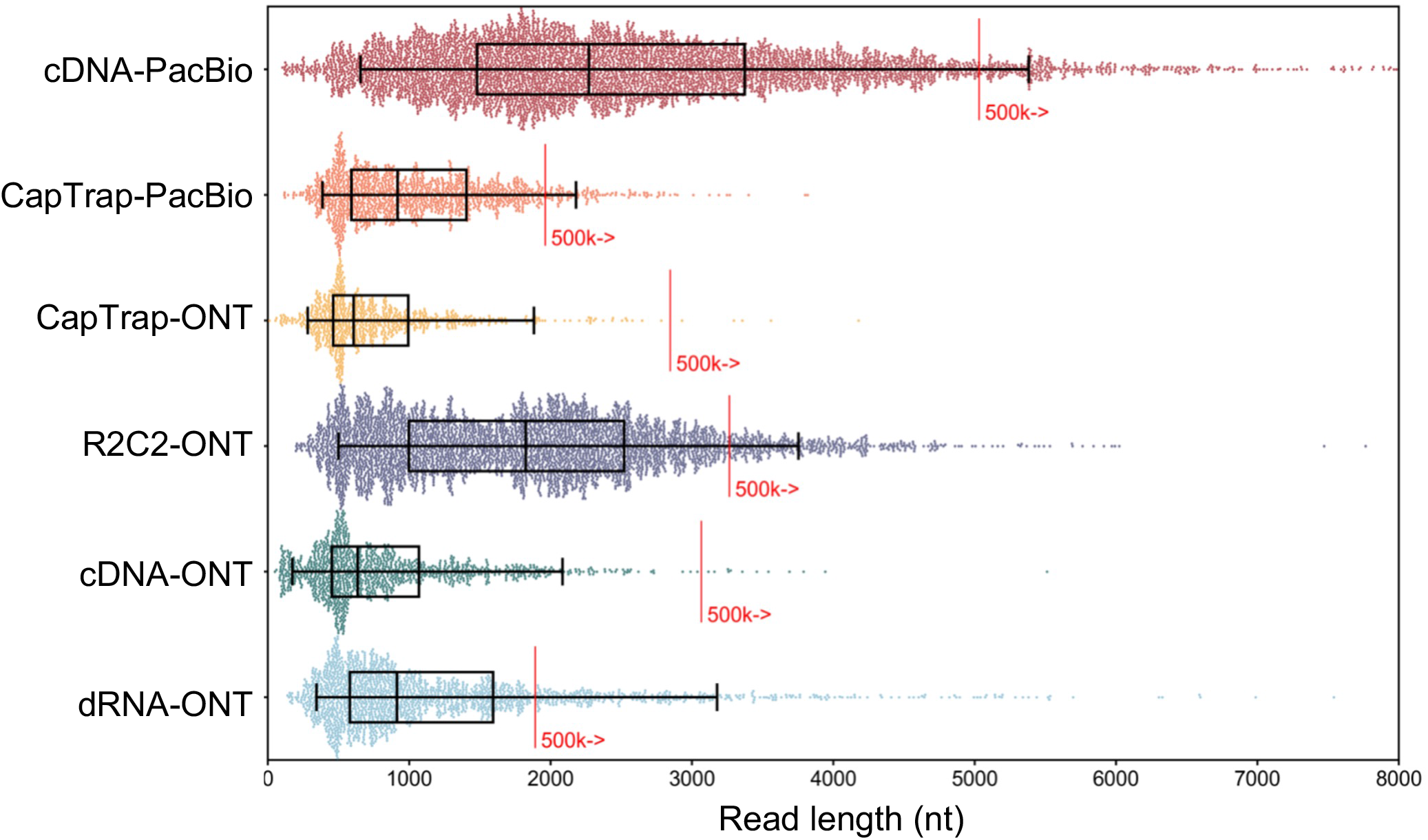
Read length distributions in six protocols-platforms.

**Extended Data Fig. 62.**
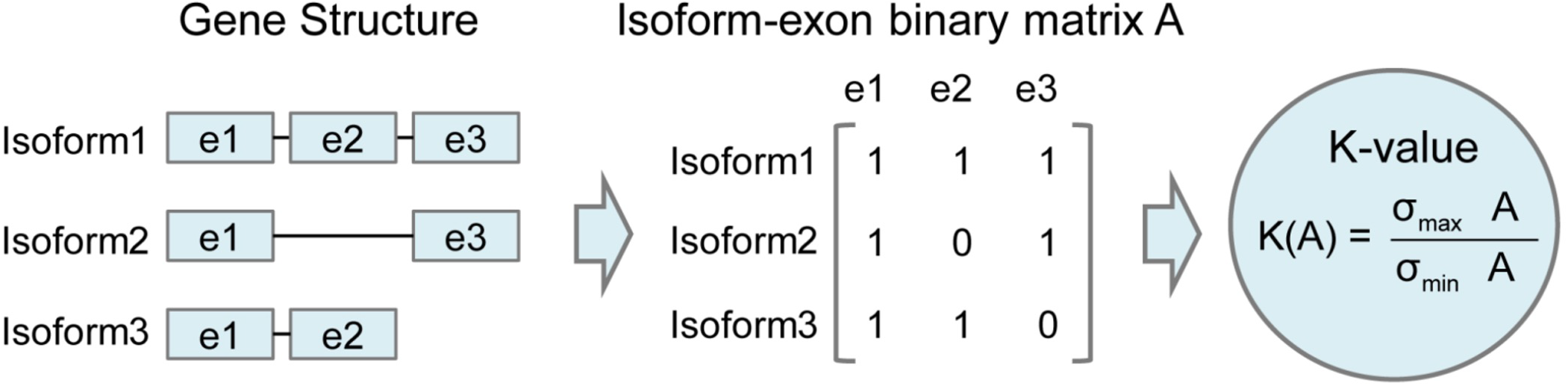
Description of K-value. A measure of the complexity of exon-isoform structures for each gene.

**Extended Data Fig. 63.**
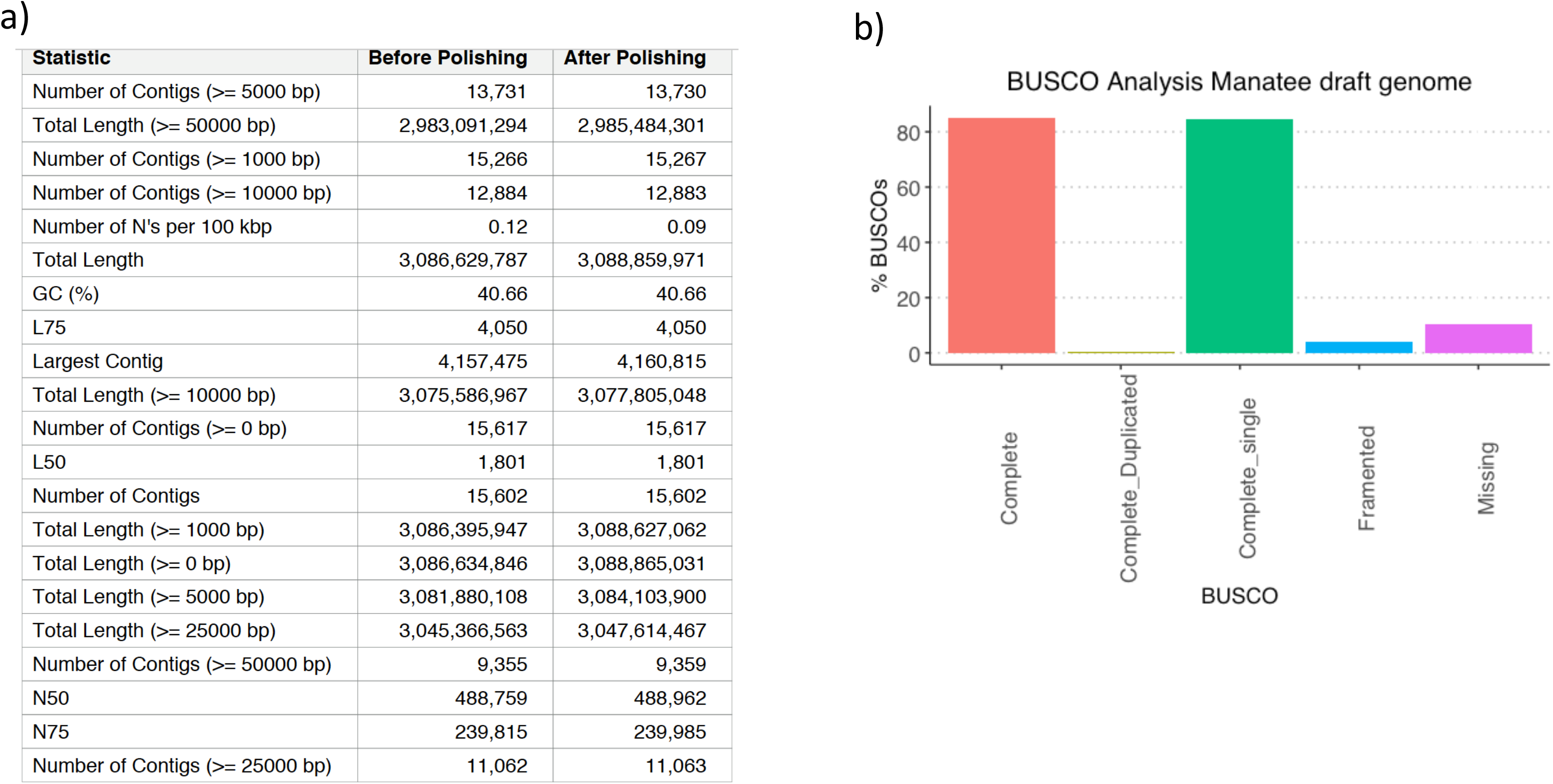
Manatee genome assembly statistics. **a** Nanopore reads were used to obtain a draft genome of the Floridian manatee with Flye. The resulting assembly was polished with existing Illumina reads using Pilon. **b** BUSCO completeness.

**Extended Data Fig.64.**
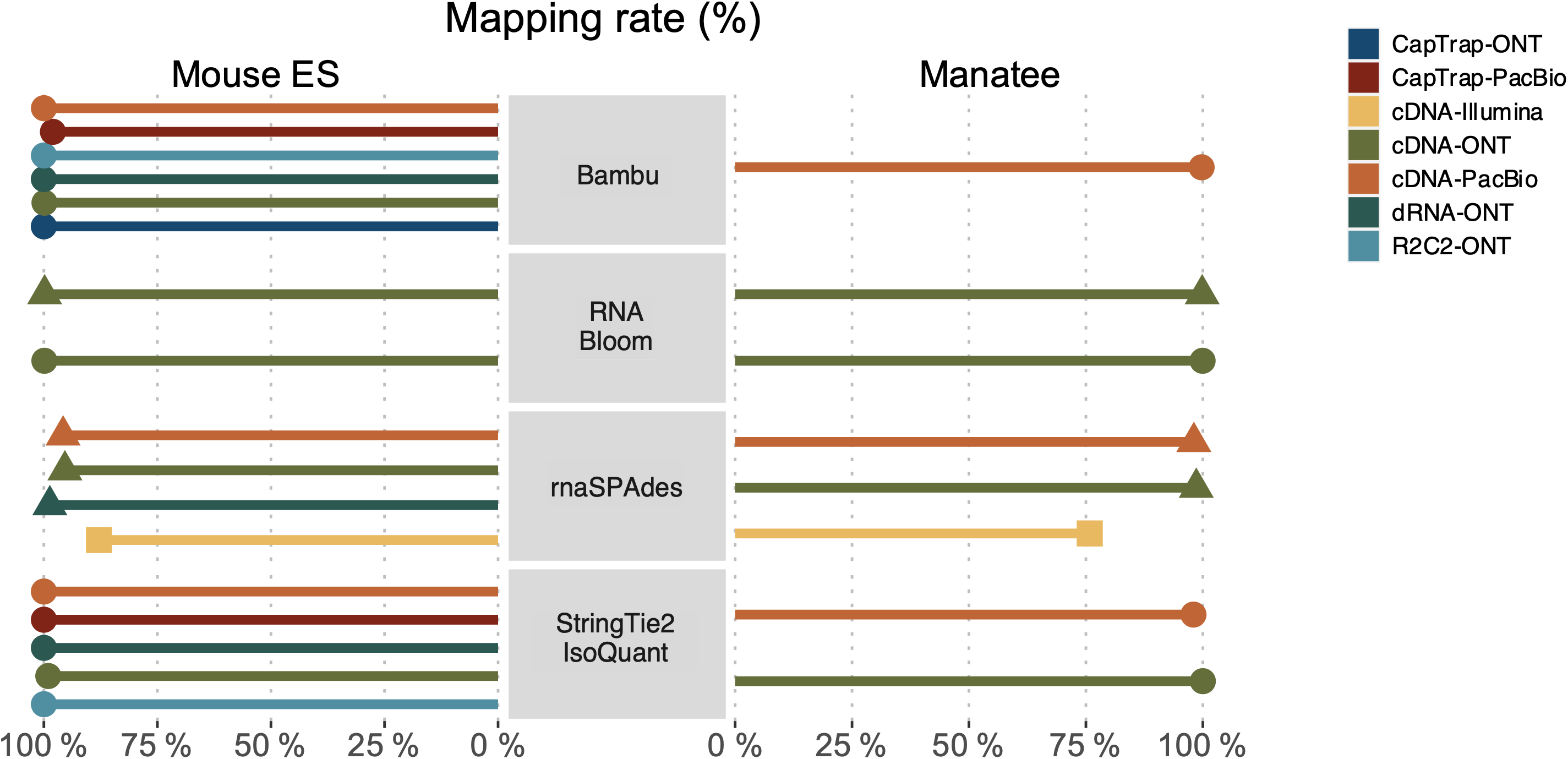
Mapping rate of transcript detected by Challenge 3 submissions.

**Extended Data Fig. 65.**
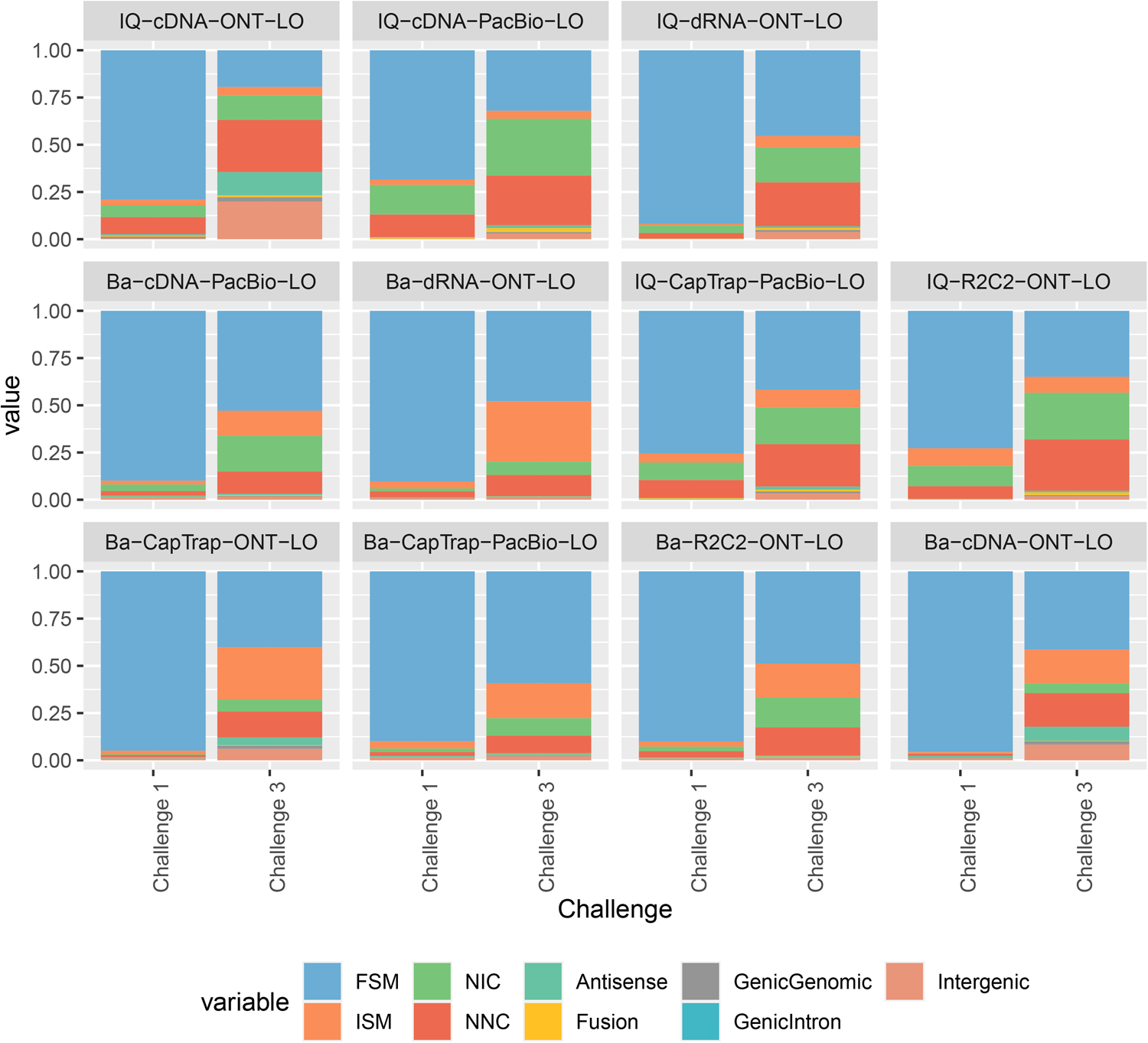
SQANTI category classification of transcript models detected by the same tools in Challenge 1 and Challenge 1 predictions used the reference annotation and Challenge 3 predictions did not. Ba = Bambu, IQ = StringTie2/IsoQuant.

**Extended Data Fig. 66.**
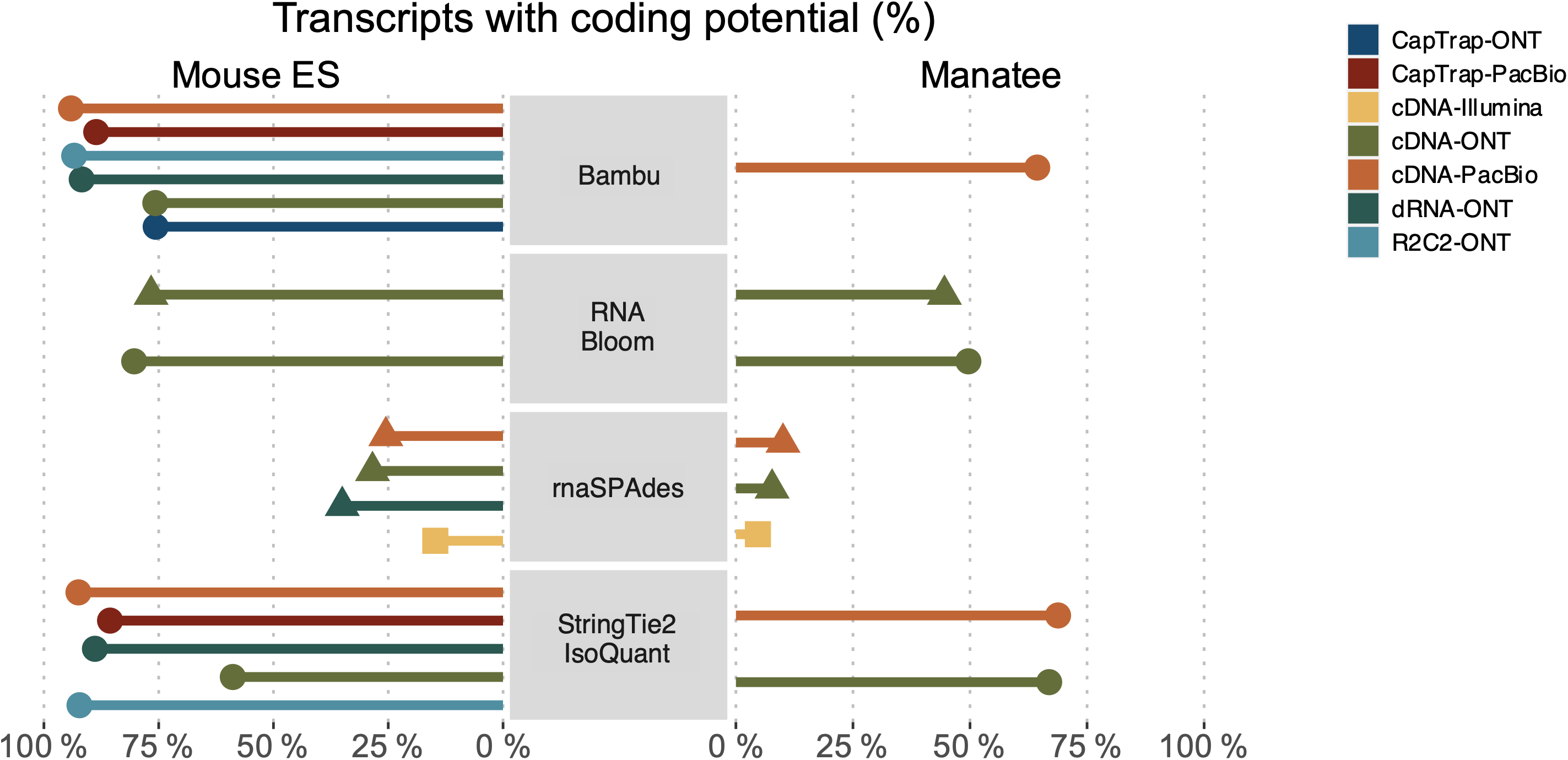
Coding potential of transcripts detected by Challenge 3 submissions.

**Extended Fig. 67.**
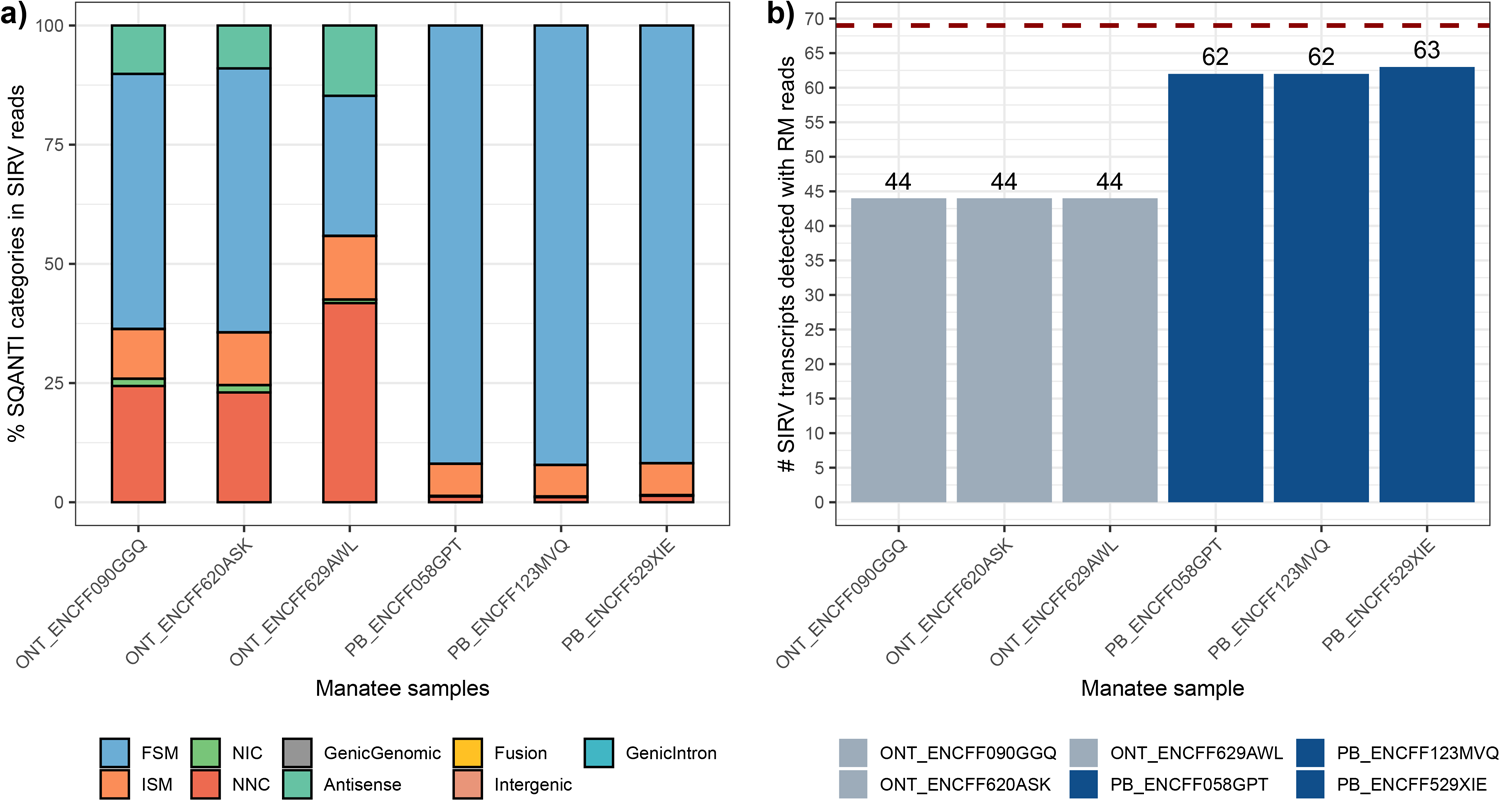
SQANTI3 analysis of SIRV reads in manatee samples. a) SQANTI3 categories for reads mapping to SIRVs in cDNA−PacBio and cDNA−ONT replicates.

**Extended Data Fig. 68.**
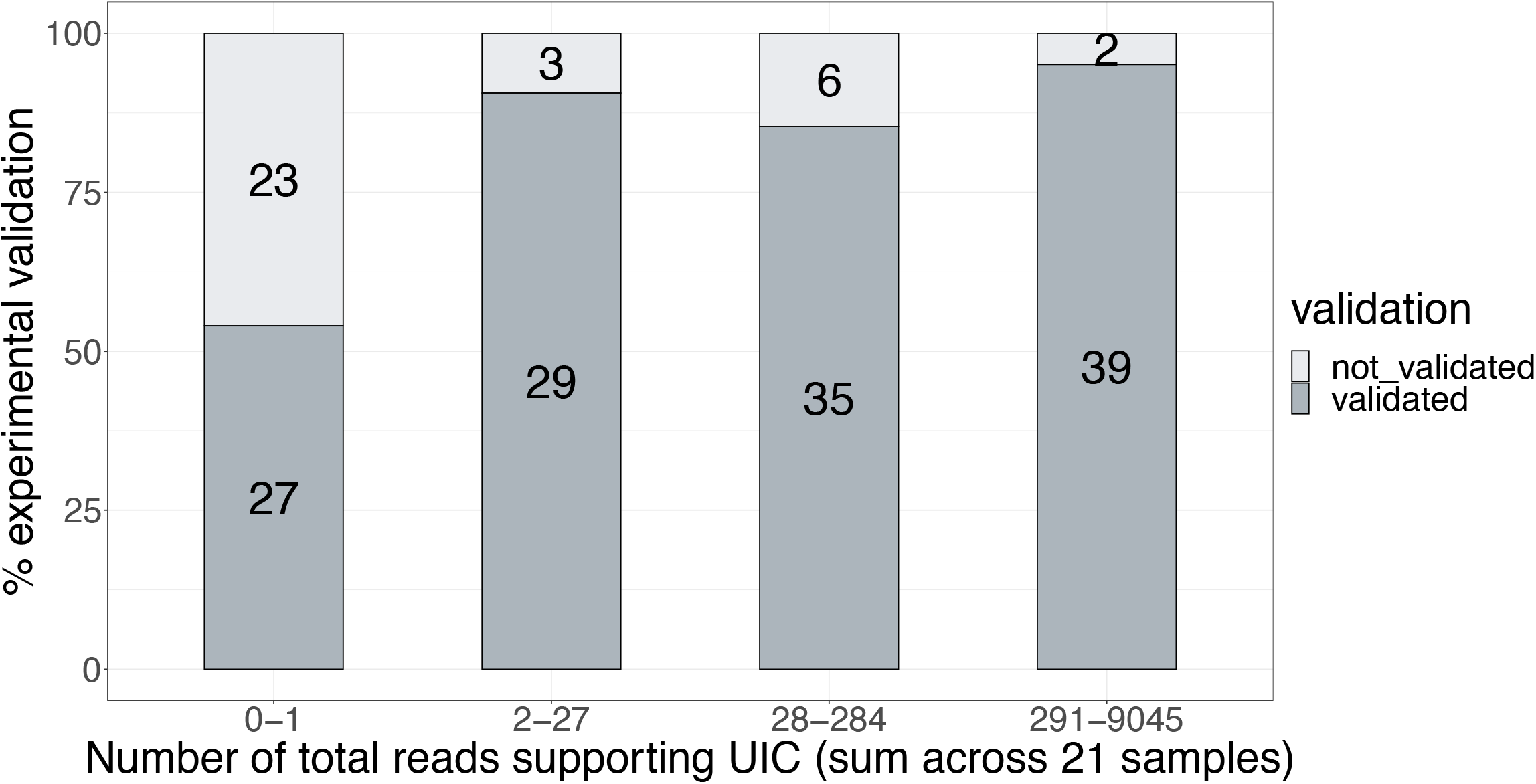
Fraction of validated transcripts as a function of the total numbers of long reads that were observed across the 21 library preparations (e.g., PacBio cDNA, ONT cDNA, PacBio CapTrap).

**Extended Data Fig. 69.**
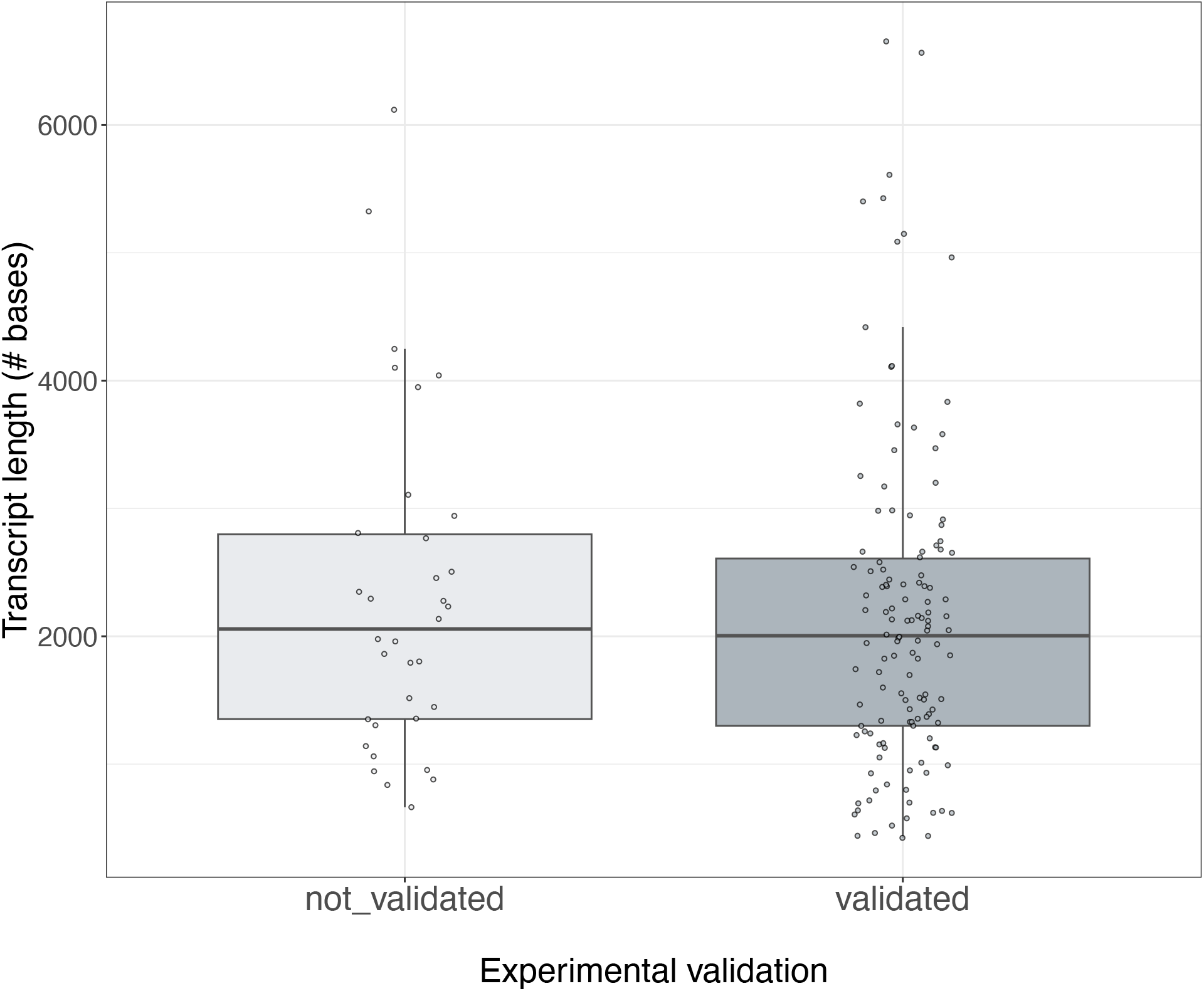
The distribution of lengths corresponding to the target transcript isoform across the entire validation experiment (including GENCODE, Platform, and Consistency groups), broken down by their validation status.

**Extended Data Fig. 70.**
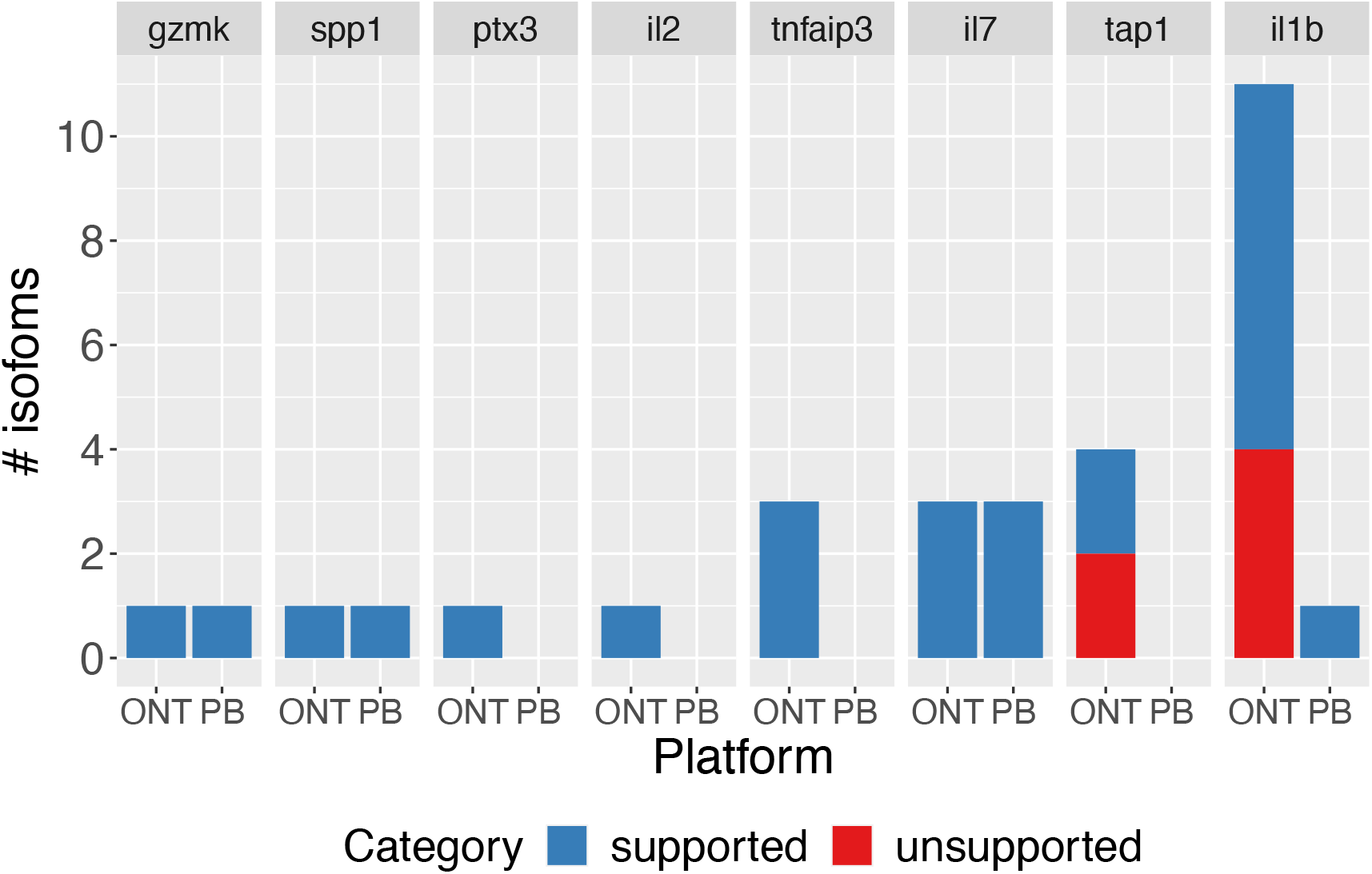
PCR validation results for manatee isoforms for seven target genes (data shown in **Figure 5l**) broken down by the platform (ONT or PacBio) underlying the pipelines that led to the identification of the isoform.

**Extended Data Fig. 71.**
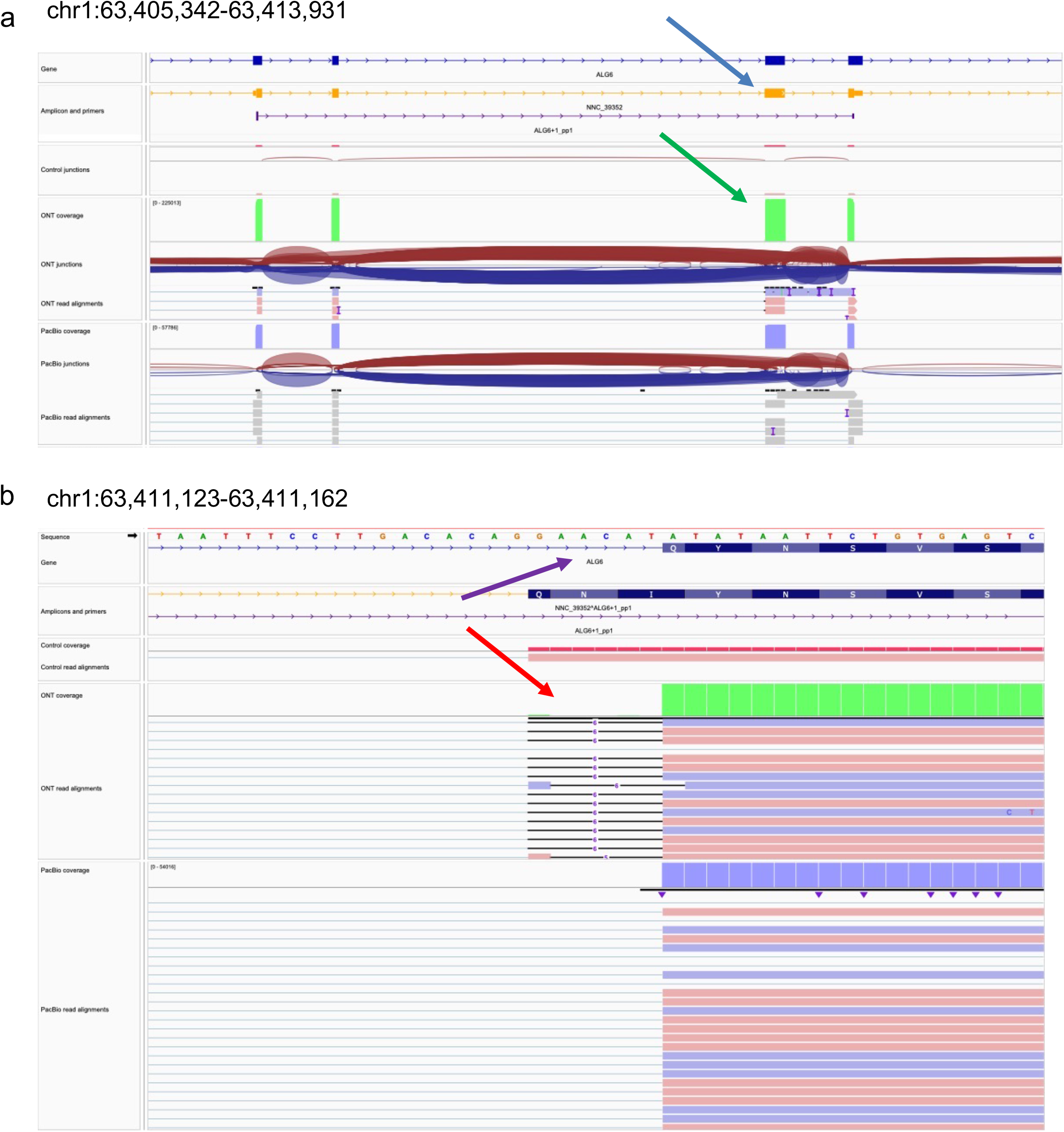
Validation of ALG6 U12 Intron with WTC11 Reads. In panel (a), a novel transcript model, NCC_39352 (blue arrow), appears to corroborate the exon within the ALG6 GENCODE annotation. The mapped amplicon in the control junction tracks provides evidence of the preceding intron. The green arrow indicates the ONT and PacBio read alignment coverage over the exon, but the junction tracks shows a lack of support for the splice junction at the exon’s 5’ end. In panel (b), GENCODE’s annotation of a rare U12 GT-AT intron (purple arrow), which is unsupported by minimap2. Instead, minimap2 forces a GT-AG intron by reporting a six-base deletion in the reference genome (red arrow). As all pipelines relied on minimap2, correct annotation of this transcript was unattainable, illustrating the challenges difficult-to-align regions can pose to annotation with long-read transcripts.

