## Supplementary material for "Systematic assessment of long-read RNA-seq methods for transcript identification and quantification": Supplemenatry Information

### LRGASP Supplementary Information

This document contains supplementary information, figures, and tables supporting *Systematic assessment of long-read RNA-seq methods for transcript identification and quantification*.


**LRGASP Pilot** 2
**LRGASP Challenge** 7
**Submissions and timeline** 7
**Capping SIRVs** 11
**Mouse and human RNA sample preparation** 12
**Manatee RNA sample preparation** 12
**Manatee genome sample preparation** 13
**cDNA preparation for Illumina and PacBio sequencing of human and mouse** 13
**PacBio library preparation of human and mouse libraries** 14
**CapTrap preparation for PacBio and ONT sequencing of human and mouse** 14
**R2C2 preparation for ONT sequencing of human and mouse** 16
**cDNA preparation for ONT sequencing of human and mouse** 16
**Direct RNA (dRNA) preparation for ONT sequencing of human and mouse** 17
**Manatee ONT genome sequencing** 17
**Manatee cDNA PacBio library preparation and sequencing** 18
**Manatee cDNA Nanopore library preparation and sequencing** 19
**Long-read data processing** 20
**Reference genome and annotations** 22
**Simulated data** 22
**CAGE data of WTC11 samples for validation of transcript 5' ends** 24
**QuantSeq of human and mouse samples for validation of transcript 3' ends** 25
LRGASP Data QC 25
GENCODE benchmarks and computational evaluation 26
**Computational pipeline description from submitters** 27
**Challenge 1** 27
**Challenge 2** 33
**Challenge 3** 38
**Method changes from registered report phase 1** 39
**Primers-JuJu bulk PT_PCR primer design tool** 41
**Data and code availability** 41
**Bibliography** 43

#### LRGASP Pilot

To demonstrate and test our evaluation metrics, we implemented our approaches on several PacBio and Nanopore transcriptomics datasets analyzed with different pipelines to verify that the proposed metrics were able to reveal differences among experimental and bioinformatics lrRNA-seq methods.

##### Challenge 1 mock evaluation

The Challenge 1 mock-run dataset consisted of available PacBio Sequel II (cDNA) (ENCODE ENCSR838WFC)[1,2] and Nanopore directRNA[3] lrRNA-seq experiments from the GM12878 cell line. Data was analyzed with 4 different algorithms (A, B, C, and E), most of them applying two different sets of parameters (permissive and restrictive), resulting in a total of 11 analysis pipelines. We next discuss the results of the mock run analysis to illustrate how metrics were to be interpreted and to anticipate differences among submissions. Note that pipeline optimization was not attempted at the mock run, therefore no conclusions can be extracted at this point on the performance of any of the methods included in this test. Hence, the mock run analysis serves for the only purpose of assessing metrics, and not long-read methodologies.

The evaluation of Challenge 1 on mock data indicated that a great variability in transcript detection is to be expected among long reads sequencing platforms, library preparation protocols and analysis pipelines. **Supplementary Data Report** provides an exhaustive comparative evaluation of Challenge 1 predictions according to the LRGASP evaluation metrics, while Error! Reference source not found. highlights some r epresentative results. First, the total number of detected isoforms compared as Unique Intron Chains (UIC) varied from ~2000 in the ONT_E2 to over 8000 in the ONT_E1 pipelines. FSM was the most abundant SQANTI category in most cases, except for two ONT pipelines that detected a similar number of ISM, NIC and NNC, revealing a very different detection rate for known and novel transcripts by the different methods (**Supplementary Fig. 2a**). While the number of isoforms per genes was roughly similar (**Pilot Data Report, page 5**), the distribution in SQANTI structural categories was very different (**Pilot Data Report, page 7**). Remarkably, the overlap in detected transcripts among pipelines was low (Jaccard Index < 0.5) except for algorithm A applied to the same data with different parameters (**Supplementary Fig. 2b**). This indicated that we might find a strong algorithm-bias in the LRGASP results that would require attention. The great majority of the detected transcripts were found by only one pipeline (**Pilot Data Report, page 17**), although this single pipeline detected transcripts were mostly novel isoforms (ISM, NIC, NNC and antisense) and an enrichment in

FSMs was observed for those transcripts detected by more pipelines, indicating that agreement is more frequent for known transcript models than for novel isoforms. Similarly, transcripts detected by many pipelines showed higher expression values, regardless of the SQANTI category (**Supplementary Fig. 2c** and **Supplementary Data Report, pages 18 to 24**), indicating that high expression value was a signature of consistent detection. No association was found between consistent detection and transcript length or exon number (**Pilot Data Report, pages 25 to 38**).

Using LRGASP metrics we evaluated general characteristics of transcript models, including how well they are supported by the GENCODE annotation and by orthogonal data (CAGE, poly(A) motifs and Illumina reads). We found significant differences in the definition of 5’ and 3’ ends when pipelines were compared (**Supplementary Fig. 2d**). While methods A and B used to provide FSM transcript models with 5’ and 3’ ends closely matching the reference TSS and TTS, respectively, pipelines C and E showed greater variability. These differences were maintained regardless the sequencing platform (PacBio or ONT), suggesting an algorithm rather than a data property. Interestingly, similar distribution of distances to closest CAGE peaks were found both for FSM (**Pilot Data Report, page 13**) and ISM (**Pilot Data Report, page 14**) when pipelines are compared. NIC are novel transcripts that contain novel combinations of annotated donor or acceptor sites. One type of NIC is intron retention. Pipelines also varied greatly in the number (**Pilot Data Report, page 100**) and percentage (**Supplementary Fig. 2e**) of transcripts showing an intron retention event, with pipelines based on ONT data having a general higher incidence. Similarly, the percentage of NNC transcripts having at least one non-canonical splice junction varied from 0% for B and E algorithms, to values above 20% for other computational methods (**Supplementary Fig. 2f** and **Supplementary Data Report, page 115**), indicating significant differences among algorithms in their control of canonical junctions. Finally, there were large differences in the number of novel transcripts (NIC: **Supplementary Data Report, pages 99 and 106**, and **Supplementary Fig. 2g** and **Supplementary Data Report, page 123**) with complete orthogonal support (3’, 5’ ends and splice junctions) among pipelines, especially for permissive versions of method E, regardless of the sequencing platform. These evaluations provide evidence of the importance of algorithmic choices in calling transcript models.

The utilization of LRGASP evaluation metrics on the full transcript model dataset allowed us to qualitatively compare pipelines, reveal their specific biases for transcript detection, and provided a means to select candidates for experimental validation. However, as no ground truth exists in this case, formal performance metrics cannot be calculated with these data. The incorporation of spike-ins (SIRVs), simulated data and a set of highly curated GENCODE genes in the LRGASP challenge allowed for evaluation against different types of ground-truth datasets. Since our mock data included the Lexogen SIRV-Set3 with 69 spiked-in isoforms, LRGASP performance metrics can be illustrated with these data (Supplementary Fig. 2 **h-l**).

Out of the 11 pipelines in our mock run, 8 provided predictions for SIRVs. Pipelines predicted between 26 and 75 SRIV transcripts (**Supplementary Fig. 2h**), indicating a great diversity in the isoform calls returned by different methods. Also, some pipelines detected many partial transcripts while others did not show this problem at all (**Pilot Data Report, page 46**). Since analysis pipelines could include multiple transcript models matching the same SIRV transcript, for example having the same set of junctions but different 3' or 5' ends, we introduced the redundancy and non-redundant prediction metrics to evaluate these cases. While some pipelines had consistent redundancy levels of one, indicating each SIRV was detected by one single transcript model, others had mean redundancy values greater than one (**Pilot Data Report, page 56**), which suggests that analysis methods follow different strategies for using reference annotation to consolidate their transcript models. Finally, metrics such as False Negatives (**Supplementary Fig. 2i**), Precision (**Supplementary Fig. 2j**), Sensitivity (**Supplementary Fig. 2k**), and False Detection Rates (**Supplementary Fig. 2l**) reported very different values across pipelines and were generally more influenced by the algorithm than by the sequencing platform.

We identified UJCs to compare transcripts across pipelines in our pilot experiment and computed barcodes as described in the methods section **Supplementary Fig. 3** gives examples of analyses performed based on barcode information. Figures 5a-f show transcript model characteristics as a function of increasing number of Nanopore or PacBio pipelines where the UCJ was detected. We found a number of FSMs detected by Nanopore but not by PacBio, but also a concentration of this category in the detection by both sequencing platforms. Interestingly, NIC were frequently found by only one pipeline, although there were also examples of NICs found by all PacBio pipelines (**Supplementary Fig. 3b**). A similar pattern, though lower in number could be seen in Fusion transcripts (**Supplementary Fig. 3c**). This suggests that Nanopore might have a higher capacity for identifying transcripts present in the reference while novel transcripts are strongly pipeline and sequencing platform-specific with a slightly higher percentage of PacBio novel transcripts being robust to pipeline choices. When looking at transcript properties, the analysis shows that PacBio recovers more transcripts with a higher predicted number of exons (**Supplementary Fig. 3d**) and length (**Supplementary Fig. 3c**), and that, in general, highly expressed transcripts are those identified by all pipelines regardless the long reads sequencing platform (**Supplementary Fig. 3f**). This conclusion is corroborated when we looked at aggregated values per sequencing platform by setting our barcode selection to filter by > 1 for dRNA-ONT (position 2) to 0 for all others (position 1 and 3). Transcript models predicted exclusively by dRNA-ONT were more highly expressed (**Supplementary**

**Fig. 3g**), with fewer exons (**Supplementary Fig. 3h**) and shorter (**Supplementary Fig. 3i**) than all other transcripts.

In summary, our mock-run analysis demonstrated the ability of LRGASP metrics to highlight important differences between both experimental and computational lrRNA-seq methods

#### Challenge 2 mock evaluation

To test the validity of the proposed metrics in LRGASP Challenge 2, we conducted the performance evaluation of two different pipelines (referred as "Pipelines 1 and 2") on two types of lrRNA-seq data (cDNA-PacBio and cDNA-ONT) from GM12878 cells. We evaluated the accuracy of transcript abundance estimation by examining the variation and similarity of estimates among multiple replicates by the metrics irreproducibility, ACVC, consistency and ACC scores (**Supplementary Fig**. 4**a-b**). In addition, SIRV set-3 data was used to evaluate how close the estimations and the ground truth values are by four metrics: SCC, NRMSE, MRD and ARR (**Supplementary Table 1**).

**Supplementary Table 1: Metrics for Challenge 2 evaluation**

| **Metrics** | **Description** |
|---|---|
| Spearman Correlation Coefficient (SCC) | SCC evaluates the monotonic relationship between the estimation and the ground truth. |
| Abundance Recovery Rate (ARR) | ARR is the percentage of the estimation over the ground truth. |
| Median Relative Difference (MRD) | MRD is the median of the relative difference of abundance estimates among all transcripts. |
| Normalized Root Mean Square Error (NRMSE) | NRMSE provides a measure of the extent to which the one-to-one relationship deviates from a linear pattern |
| Irreproducibility and ACVC (Area under the Coefficient of Variation Curve) | Irreproducibility and ACVC characterize the coefficient of variation of abundance estimates among different replicates. |
| Consistency and ACC (Area under the Consistency Curve) | Consistency and ACC characterize the similarity of abundance profiles between mutual pairs of replicates. |

| | |
|---|---|
| Resolution Entropy (RE) | RE characterizes the resolution of abundance estimation. |

Pipeline 2 has the lowest coefficient of variation (ACVC=0.92) and the highest reproducibility (irreproducibility=0.07) among multiple replicates in cDNA-ONT data (**Supplementary Fig. 4c**), which is different from in cDNA-PacBio (ACVC=1.28, irreproducibility=0.11). It indicates the performance variability of the same pipeline in different data. In addition, both pipelines on the mock data showed decreasing coefficient of variation with transcript abundances (**Supplementary Fig. 4d**), so quantification of lowly expressed transcripts remains a challenge.

Similarly, Pipeline 2 has the highest ACC value (14.57) as well as the best consistency (0.98) of transcript abundance estimation across multiple replicates on cDNA-ONT data (**Supplementary Fig. 4e**). The consistency scores of both pipelines decreased dramatically when the abundance threshold was 5 or smaller (**Supplementary Fig. 4f**). Therefore, there existed greater variabilities and errors of abundance estimation by both pipelines for lowly expressed transcripts.

Finally, SIRV data also demonstrated the highest correlation between the ground truth and the estimations by Pipeline 2 (SCC=0.69, **Supplementary Fig. 4g**) and showed its best performance on cDNA-ONT data (Supplementary Fig. 4**h**, NRMSE=0.91, MRD=0.37) compared to the other test combinations of pipeline plus data. However, both pipelines overestimated transcript abundances, because 63.33%, 45.08% and 30.83% of transcripts had ARR larger than 100% for the three scenarios **(Supplementary Fig. 4**).

#### Challenge 3 mock evaluation

For the Challenge 3 mock evaluation, we used the different long-read libraries - Sequel I, Sequel II and three MinIon (**Supplementary Table 2**) - generated for the manatee sample, and processed them independently with the Isoseq3 algorithm[4], resulting in five different "pipelines" providing transcript model predictions for the manatee. FASTA sequences were mapped to the manatee draft genome using minimap2[5]. Note that Challenge 3 instructions for manatee data indicate that all reads from each sequencing platform must be combined to predict transcript models, therefore our mock pipelines use data subsets of the actual competition. Supplementary Fig. 5shows the results of this analysis. While the number of predicted transcript models was very different for each pipeline (**Supplementary Fig. 5a)**, in all cases the mapping rate was high (**Supplementary Fig. 5b**) with PacBio transcript models reaching 100% mapping rate and a majority of detected transcripts were multiexon **(Supplementary Fig. 5c**). The distribution of transcript length showed that ONT3 and both PacBio pipelines

had higher values than ONT1 and ONT2 between and median values varied between 1094 and 1394 nts (**Supplementary Fig. 5d**). Clear differences were observed between PacBio and Nanopore pipelines on the number of transcripts with complete short-read junction support (**Supplementary Fig. 5e**), the total number of non-canonical junctions (**Supplementary Fig**. **5f**) and the number of transcripts containing at least one non-canonical junction (**Supplementary Fig. 5**), with PacBio pipelines showing, in general, higher support and less incidence of junctions non-canonical. Also, the predicted coding potential for PacBio pipelines was higher than for Nanopore (**Supplementary Fig. 5h**). As for BUSCO analysis, ONT pipelines returned a higher number of either BUSCO complete, BUSCO incomplete and BUSCO duplicated sequences than PacBio pipelines, although the relative numbers were roughly similar, except for the PB2 pipeline that had a lower fraction of BUSCO complete genes (**Supplementary Fig. 5i**).
This analysis indicated that our LRGASP metric were able to capture differences between analysis pipelines and revealed that although in all cases transcript models can be mapped to the genome, their number and sequence and splice site accuracy is very different, with ONT pipelines returning more complete transcriptomes, and PacBio pipelines returning better supported and accurate splice sites.

In these mock evaluations for all three challenges on published GM12878 data, we highlighted the variability between sequencing platforms and computational methods. This further motivates the need for the LRGASP effort to highlight these differences in a real study and to use our benchmarks for evaluation.

#### LRGASP Challenge

##### Submissions and timeline

Participants submitted challenge predictions on Synapse (https://www.synapse.org/#!Synapse:syn25007472).

The following is an overview of the data used for each challenge and the result files that were submitted (**Supplementary Fig. 5-Supplementary** Fig. 7).

- Challenge 1: transcript isoform detection with a high-quality genome (iso_detect_ref)
    - Samples
        - WTC11 (human iPSC cell line)
        - H1-mix (human H1 ES cell line mixed with human Definitive Endoderm derived from H1)
        - ES (mouse ES cell line)

        - human simulation - simulated human reads (Illumina,cDNA-ONT, and cDNA-PacBio)
        - mouse simulation - simulated mouse reads (Illumina and cDNA-PacBio, dRNA-ONT)
    - Result files:
        - models.gtf.gz
        - read_model_map.tsv.gz
- Challenge 2: transcript isoform quantification (iso_quant)
    - Samples
        - WTC11 (human iPSC cell line)
        - H1-mix (human H1 ES cell line mixed with human Definitive Endoderm derived from H1)
        - Human simulation - simulated human reads (Illumina, cDNA-ONT, and cDNA-PacBio)
        - Mouse simulation - simulated mouse reads (Illumina and cDNA-PacBio, dRNA-ONT)
    - Result files:
        - expression.tsv.gz
        - models.gtf.gz
- Challenge 3: de novo transcript isoform detection (iso_detect_de_novo)
    - Samples
        - Manatee (manatee whole blood)
        - ES (mouse ES cell line)
    - Result files:
        - rna.fasta.gz
        - read_model_map.tsv.gz

Computational methods may have been developed and tuned to a specific sequencing platform, library prep approach (e.g., dRNA-ONT), or use of additional orthogonal data; therefore, entries were organized such that a comparison can be made across different tools using the same type of data. Additionally, it was important to evaluate how robust computational tools are to transcript analysis in different species or biological samples. Thus, for each entry to a challenge, a team selected a data category, library prep, and sequencing platform and submitted experiments for all samples that are available for the challenge + library prep + sequencing platform combination (**Supplementary Fig. 5)**. The samples that are available for a challenge + library prep + sequencing platform combination can be found in **Supplementary Table 2**. Note that there are also simulated samples that were also be available for Challenges 1 and 2.

Each entry must have met the following requirements:

*Requirements for Challenge 1 and 2*

At least one experiment must have been supplied for each sample available for a given challenge, library prep, and sequencing platform combination that is selected. Human and mouse samples have biological replicates that must have been used for the entry.

A major goal of LRGASP is to assess the capabilities of long-read sequencing for transcriptome analysis and how much improvement there is over short-read methods. Additionally, long-read computational pipelines vary in their use of only long-read data or if they incorporate additional data for transcript analysis. To facilitate comparisons between long-read and short-read methods and variation in tool parameters, we broke down submissions into different categories:

- long-only - Use only LGRASP-provided long-read RNA-seq data from a single sample, library preparation method and sequencing platform.
- short-only - Use only LGRASP-provided short-read Illumina RNA-seq data from a single sample. This is to compare with long-read approaches.
- long and short - Use only LGRASP-provided long-read and short-read RNA-Seq data from a single long-read library preparation method and the Illumina platform. Additional accessioned data in public genomics data repositories can also be used.
- freestyle - Any combination of at least one LRGASP data set as well as any other accessioned data in public genomics data repositories. For example, multiple library methods can be combined (e.g. cDNA-PacBio + CapTrap-PacBio , cDNA-ONT + CapTrap-ONT + R2C2-ONT + dRNA-ONT, all data, etc.).

In all the above categories, the genome and transcriptome references specified by LRGASP were used. For the long and short and freestyle category, additional transcriptome references could be used.

All replicates must have been used in each experiment. Challenge 2 must have reported replicates separately in the expression matrix. Each team could submit multiple entries for each challenge; however, they can only submit one entry per challenge + data type + library prep + sequencing platform combination. This was to encourage tool development that is robust to different library preps and sequencing platforms, but prevent multiple entries that are subtle parameter changes.

For Challenge 1, the submitted GTF file only contained transcripts that had been assigned a read. For Challenge 2, submitters had the option of quantifying against the reference transcriptome or a transcriptome derived from the data (i.e., results from Challenge 1). The GTF used for quantification was included as part of the Challenge 2 submission.

The type of platform and library preparation method used in a given experiment, except for freestyle experiments, was limited to data from a single library preparation method plus sequencing technology (long-only). LRGASP Illumina short-read data of the same sample could optionally be used in an experiment with the LRGASP long-read data (long and short):

- cDNA-Illumina - short-only
- cDNA-PacBio - long-only or long and short
- CapTrap-PacBio - long-only or long and short
- cDNA-ONT - long-only or long and short
- CapTrap-ONT - long-only or long and short
- R2C2-ONT - long-only or long and short
- dRNA-ONT - long-only or long and short

*Requirements for Challenge 3*

At least one experiment had to be supplied for each sample available for a given library prep and sequencing platform combination that was selected. Mouse samples had biological replicates that were used for the entry. Manatee samples only had cDNA library preparation type and sequencing data from Illumina, ONT, and PacBio.

For similar reasons as described above, the data used for a given experiment had to fit into one of the following categories:

- long-only - Use only LGRASP-provided long-read RNA-Seq data from a single sample, library preparation method and sequencing platform. No genome reference can be used.
- short-only - Use only LGRASP-provided short-read Illumina RNA-Seq data from a single sample. This is to compare with long-read approaches. No genome reference can be used.
- long and short - Use only LGRASP-provided long-read and short-read RNA-Seq data from a single long-read library preparation method and the Illumina platform. No genome reference can be used.
- long and genome - Use only LGRASP-provided long-read RNA-Seq data from a single long-read library preparation method. A genome reference sequence can be used.

- freestyle - Any combination of at least one LRGASP data set as well as any other accessioned data in public genomics data repositories. For example, multiple library methods can be combined (e.g. cDNA-PacBio + CapTrap-PacBio, cDNA-ONT + CapTrap-ONT + R2C2-ONT + dRNA-ONT, all data, etc.).

In all the above categories, except for freestyle, a transcriptome reference could not be used. The submitted FASTA file only contained transcripts that had been assigned a read. Each team could submit multiple entries for each challenge; however, they could only submit one entry per challenge + data type + library prep + sequencing platform combination.

LRGASP biological data was available at the ENCODE DCC (https://www.encodeproject.org/search/?type=Experiment&internal_tags=LRGASP). The simulated data was available from Synapse (https://www.synapse.org/#!Synapse:syn25683370). The competition was launched on May 1, 2021, and challenge submissions were closed on October 8, 2021. Figures giving a summarized overview of the challenges, including specific samples used and expected entry files (**Supplementary Fig. 8**), challenge evaluations (**Supplementary Fig. 11-**Supplementary Fig. 12), and experimental validation (**Supplementary Fig. 14**), are provided in the **Supplementary Figures** section below.

#### Supplementary Methods

Additional details of all protocols for library preparation and sequencing can be found at the ENCODE DCC and is linked to each dataset produced by LRGASP (**Extended Data Table 1**).

##### Capping SIRVs

Exogenous synthetic RNA references (spike-ins) are widely used to calibrate measurements in RNA assays, but they lack the 7-Methylguanosine ($m^7G$) cap structure that most natural eukaryotic RNA transcripts bear at their 5’ end. This characteristic makes commercial spike-in mixes unsuitable for library preparation protocols involving 5’ cap enrichment steps. Therefore, we enzymatically added the appropriate $m^7G$ structure to the SIRV standards used in this challenge. Specifically, the pp5'N structure present at the 5’ end of spike-in sequence was used as a template for the Vaccinia capping enzyme (catalog num M2080S, New England BioLabs) to add the $m^7G$ structure to SIRV-Set 4 (Iso Mix E0 / ERCC / Long SIRVs, catalog num 141.03, Lexogen). A total of ten vials of SIRV-Set 4 (100 µl) were employed to perform the capping reaction (final total mass of 535 ng). The reaction was performed following the recommendations of the manufacturer’s capping protocol with two minor changes: 3.5 µl of RNAse inhibitors (RNasin Plus RNase Inhibitor, catalog num N2611, Promega) were added to the capping reaction to avoid RNAse degradation, and the incubation time was extended from 30 minutes to two hours, following a recommendation from New England BioLabs technical support scientists. The final capping reaction was purified by using 1.8x AMPure RNA Clean XP beads (catalog num. A63987, Beckman Coulter) and resuspended in 100 µl of nuclease-free water.

**Mouse and human RNA sample preparation**

Prior to distribution of the biosample total RNA aliquots to each of the participating labs, 110 µg of each biosample total RNA was spiked with Lexogen Long SIRV Set-4 quantification standards (catalog # 141.03) at approximately 3% of the estimated mRNA mass present (~1% of total RNA). The mass of capped SIRVs used was 29.5 ng and the mass of uncapped SIRVS used was 28.9 ng. In the case of direct RNA sequencing of one replicate of WTC11 (ENCODE library accession ENCLB926JPE) and one replicate of mouse ES cells (ENCODE library accession ENCLB386NNT), only uncapped SIRV 4.0 were spiked in at approximately 3% of the estimated mass. Appropriate volumes of the spiked total RNA mixture to meet the input mass requirements for each library preparation method were then aliquoted separately, stored at -80 C, and shipped on dry ice to participating labs.

**Manatee RNA sample preparation**

Blood samples from Florida manatees were collected during health assessments by the U.S Geological Survey (USGS) Sirenia Project, the Florida Fish and Wildlife Conservation Commission (FWC), and the University of Florida under U.S. Fish and Wildlife Service (USFWS) permit # MA791721-5 in Crystal River (Citrus County, Florida, USA) and in Satellite Beach (Brevard County, Florida, USA) in December and January of 2018 and 2019 respectively. Samples were processed under the University of Florida USFWS permit #MA067116-2 following a protocol approved by the ethics committee (IACUC # 201609674 & IACUC # 201909674). Whole blood from minimally restrained Florida manatees were collected from the medial interosseous space between the ulna and radio from the pectoral flippers. Samples were drawn using Sodium Heparin 10-mL BD vacutainers (BD BioScience, New Jersey, U.S.A). Blood samples were spun on-site, and the plasma was aliquoted, stored in liquid nitrogen or ice, and transferred to -80 ºC once in the lab. The buffy coat (white blood cells) was flash-frozen in liquid nitrogen on-site and total RNA was extracted subsequently in the lab using STAT 60 (Tel-test Friendswood, TX) reagent. Approximately 350 µL of the frozen buffy coat was added to 1 ml of STAT 60 and vortexed for 30 seconds, 250 µL of chloroform was added and the tube was centrifuged 20,800 x g for 15 minutes at 4 ºC, to extract the RNA. This step was repeated and then RNA was precipitated from the supernatants overnight at -20ºC by the addition of 700 µL isopropanol with 1.5 µL of GlycoBlueTM (15 mg/mL) (Ambion, Invitrogen, Austin, TX) as a coprecipitant. Following centrifugation at 20,800 x g for 45 minutes, the pellet was washed with ethanol 70%, air-dried, and resuspended in 20 mL of RNA secure

(Ambion, Austin, TX). A DNAse treatment was performed using Turbo DNA-freeTM kit (Ambion, Austin, TX). A total of nine good-quality RNA samples were selected to create an RNA pool. These samples included 6 females, one calf, one lactating female and one male and had RIN values from 8.0 to 8.8.

**Manatee genome sample preparation**

The genome of the Florida manatee Lorelei was sequenced using Nanopore and PacBio. Lorelei is the same individual manatee for which an Illumina-based genome assembly was released by the Broad Institute in 2012 [6]. An EDTA, -80ºC whole blood sample aliquot was used. gDNA was extracted from 1400 µl of blood using the DNeasy kit (QIAGEN, MD, USA) following the companies' specifications for 100 µl aliquots of blood. Thawed blood was diluted 1:1 with RNA free Phosphate buffered saline 1x (Gibco, UK), 20 µl of proteinase K (QIAGEN, MD, USA), and 200 ul of AL lysis buffer (QIAGEN, MD, USA) and vortexed immediately. It was incubated at 56 °C for 10 minutes. Then, we added 200 µl of ethanol 96% and mixed it thoroughly. The mixture was added to the DNeasy mini spin-column and centrifuged at 6,000 x g for 1 minute. The column was washed with 500 µl of AW1 solution (QIAGEN, MD, USA) and centrifuged at 6,000 x g for 1 minute and followed with a wash with 500 µl AW2 (QIAGEN, MD, USA ) and centrifuged 20,000 x g for 3 minutes. gDNA was eluted twice with 100 µl of AE buffer added to the center of the column, incubated for 1 minute, and centrifuged 6,000 x g for 1 minute. The first and second elution from the DNeasy mini spin-column were pooled and concentrated using a speed vacuum for 20 minutes in which each preparation was reduced from 200 to 50 µl. All gDNA tubes were pooled and the DNA was cleaned with AM Pure magnetic beads (Beckman Coulter-Life Sciences, IN, USA) at a ratio of 0.5:1, beads volume to gDNA volume (50 µl of beads to 100 µl of gDNA). gDNA bound to the beads was washed twice with 1 ml of 70% ethanol. Ethanol traces were removed by quick spin to the bottom of the tube and removed with a pipette. Then, the beads were dried for 2 minutes and gDNA was eluded in 55 µl of EB buffer (QIAGEN, MD, USA) at 37 °C with 10 minutes of incubation. This process was repeated twice. Quantification of gDNA was performed with a Qubit™ fluorometer (Thermo Fisher Scientific) and the quality of the gDNA was assessed using a Genomic Tape on the Agilent TapeStation (Santa Clara, CA, USA). The final DNA quantity was 28.8 µg of DNA at a concentration of 267 ng/µl. The DNA Integrity Number (DIN) was 8.8 and the peak size was 54.5 kb.

**cDNA preparation for Illumina and PacBio sequencing of human and mouse**

PacBio cDNA synthesis was performed using a modified version of the Picelli protocol[7] substituting the Maxima H- reverse transcriptase. Total RNA (400 ngs) spiked with SIRV standards was combined in a priming reaction with RNAse inhibitor, oligo dT, dNTP's and water, incubated at 72ºC for 3 minutes, then ramped down to 50ºC for an additional 3 minutes. We then added a first strand synthesis buffer (5x RT buffer, TSO oligo, Maxima H(-

) RT and water) that had previously been equilibrated to 50ºC, to the priming reaction. First strand synthesis was carried out as follows: (Extension at 50ºC for 90 min, 85ºC for 5 min and held at 4ºC). To the first strand reaction, we then added 2x SeqAmp (Takara) reaction buffer, IS primers, water and SeqAmp polymerase). First strand cDNA was amplified for 11 cycles as follows: (95ºC 1 min, 98ºC 15 sec, 65ºC 30 sec and 68ºC 13 min), and finished off by incubation at 72ºC for 10 min and holding at 4ºC. The amplified products were purified using SPRI beads, quantified on Qubit, and checked for length distribution on the Agilent Bioanalyzer. The short-read protocol is described in the *Nextera DNA Flex Library Prep Reference Guide*[8] and the long-read protocol in *Long read cDNA prep with Maxima H(-) (no exonuclease version)*[9]. 50 ng sub-aliquots of the full-length cDNA libraries were tagmented for Illumina short-read sequencing using the Illumina Nextera DNA Flex Library prep kit, according to the manufacturer's protocol.

**PacBio library preparation of human and mouse libraries**

To build PacBio libraries, we followed the SMRTbellTM Express Template Prep Kit 2.0 protocol. We started from 500 ng of poly(A) selected cDNA. The ends of the cDNA were repaired first in order for the cDNA molecule to be suitable for ligation of SMRTbell adapters. We added a damage repair reaction (DNA prep buffer, NAD and DNA damage repair) and then incubated at 37ºC for 30 min. Then End prep mix was added and incubated at 20ºC for 30 min and 65ºC 20 min. Ligation of the adapter at the ends of the cDNA was done by adding a ligation mix (PacBio adapters, ligation mix, ligation enhancer and ligation additive), followed by incubation at 20ºC for 60 min. Final libraries were cleaned up using SPRI beads and we recorded the size and concentration of samples. Once the ligation step was done and the libraries passed the QC, a sequencing primer was annealed to the adapters in the UCI GHTF sequencing facility to allow for the binding of the polymerase during sequencing.

**CapTrap preparation for PacBio and ONT sequencing of human and mouse**

CapTrap is a technique developed by the Guigó laboratory (CRG, Barcelona, Spain) in collaboration with the group of Piero Carninci in RIKEN, Japan. The method enriches for full-length transcripts by selection of the 7-Methylguanosine (m7G) cap structure present at the 5' ends of RNA transcripts, followed by specific cap- and poly(A)- dependent linker ligations. The cDNA libraries generated using this method are compatible with long-read sequencing platforms (ONT or PacBio). The protocol starts with first strand synthesis (PrimeScript II Reverse Transcriptase, catalog num. 2690A, Takara) where 5 µg of total RNA poly(A) + RNAs are fully reverse transcribed using a 16-mer anchored dT oligonucleotide. First strand synthesis was performed at 42 ºC for 60 minutes. Resulting products were purified with 1.8x AMPure RNA Clean XP beads (catalog num. A63987, Beckman Coulter). After the first-strand generation, the m7G cap structure at the 5' end of the transcripts is selectively captured using the CAP-trapper technique [10,11], which leads to the removal of uncapped RNAs. The diol group on the $m^7G$ cap is oxidized with 1M NaOAc (pH 4.5) and NaIO4 (250 mM). Tris HCl (1M, pH 8.5) was added to stop the reaction and the whole reaction was purified with 1.8x AMPure RNA Clean XP beads. Aldehyde groups were biotinylated using a mixture containing NaOAc (1M, pH 6.0) and Biotin (Long Arm) Hydrazide (100 mM, catalog num. SP-1100, Vector Laboratories). The resulting mixture was then incubated for 30 minutes at 40ºC and purified with 1.8x AMPure RNA Clean XP beads. Single strand RNA was degraded by RNase ONE Ribonuclease (catalog num. M4261, Promega) for 30 minutes at 37ºC and purified with 1.8x AMPure RNA Clean XP beads. The m7G cap structure bound to biotin is then selected using M-270 streptavidin magnetic beads (catalog num. 65305, Thermo Fisher Scientific). M-270 streptavidin magnetic beads were equilibrated with CapTrap Lithium chloride/Tween 20 based binding buffer. Sample recovered after RNase ONE purification was bound to equilibrated M-270 streptavidin magnetic beads (incubation at 37ºC for 15 minutes), washed 3 times with CapTrap Tween20 based washing buffer and released by heat shock for 5 minutes at 95ºC and quickly cooled on ice. A second release was performed, and the supernatant was also collected and mixed with the eluate from the previous release. The released sample was treated with RNase H (60 U/μl, Ribonuclease H <RNase H>, catalog num. 2150, Takara), RNase ONE (10 U/μl) and CapTrap release buffer (incubated at 37ºC for 30 minutes), purified with 1.8x AMPure XP beads (catalog num. A63881, Beckman Coulter) and concentrated by using a speed vac. After this cap specific selection, two double-stranded linkers, carrying a unique molecular identifier (UMI), are specifically ligated to the first strand cDNA [12]. Linker ligation (DNA Ligation Kit <Mighty Mix>, catalog num. 6023, Takara) was performed in two separate steps. First the 5' linker was ligated, purified twice, to completely eliminate the non-incorporated linkers, with 1.8x AMPure XP beads and concentrated by using a speed vac. Then the 3' linker was ligated, purified once with 1.8x AMPure XP beads and finally concentrated by using a speed vac. The double stranded linkers are converted into single strand by Shrimp Alkaline Phosphatase (1 U/μl SAP, catalog num. 78390, Affymetrix) and Uracil-Specific Excision Reagent (1 U/μl USER, catalog num. M5505L, NEB) treatment. This reaction was incubated for 30 minutes at 37ºC, 5 minutes at 95ºC and finally placed on ice. The sample was then purified with 1.8x AMPure XP beads. After this treatment, the two linkers which serve as priming sites for the polymerase (2x HiFi KAPA mix, catalog num. 7958927001-KK2601, Kapa), enable the synthesis of the full-length second strand. The mixture was incubated for 5 minutes at 95ºC, 5 minutes at 55ºC, 30 minutes at 72ºC and finally held at 4ºC until 1 μl Exonuclease I (20U/μl, catalog num. M0293S, NEB) was added to each sample. The sample was then incubated for 30 minutes at 37ºC and afterwards, purified twice with 1.8x and 1.4x (respectively) AMPure XP beads and finally concentrated in a speed vac. The resulting cDNA is amplified (TaKaRa LA Taq, catalog num. RR002M, Takara) via long and accurate PCR (LA PCR) protocol. In order to minimize PCR duplicates, each sample was split in two PCR independent reactions and amplified 16 cycles with 15 seconds at 55ºC for annealing, and 8 minutes at 65ºC for extension. The 2 PCR replicates were

merged and purified with 1x AMPure XP beads. Samples were quantified with Qubit (Qubit 4 Fluorometer, Thermo Fisher Scientific) and quality checked with BioAnalyzer (Agilent 2100 Bioanalyzer, Agilent Technologies).
CapTrap MinION cDNA sequencing was performed with 500 ng of cDNA sample coming from CapTrap cDNA protocol and strictly following the SQK-LSK109 adapter ligation protocol (ONT). The cDNA sequencing on MinION platform was performed using ONT R9.4 flow cells and the standard MiniKNOW protocol.
PacBio Sequel II sequencing was performed using 500 ng of CapTrap samples following the SMRTbell™ Express Template Prep Kit 2.0 protocol.

**R2C2 preparation for ONT sequencing of human and mouse**

For each biological replicate, two libraries were created, a regular (non-size selected), and a size selected library of cDNA over 2 kb in length to achieve higher coverage of longer transcripts. For each RNA sample, 400 ng was used to generate full-length single stranded cDNA using an indexed oligo(dT) primer and a template switching oligo (TSO). PCR was used to generate the second strand and amplify the library. The cDNA was then isolated by SPRI bead clean up. For the size selected libraries, cDNA was run on a 1% low melt agarose gel. A smear in the range of 2–10 kb was excised from the gel and digested with beta-agarase followed by SPRI bead clean up. At this point, indexed cDNA from each biological replicate was pooled together equally. cDNA was circularized using a short DNA splint with sequence complementary to the cDNA ends by Gibson Assembly (NEBuilder, NEB) with a 1:1 cDNA:splint ratio (100 ng each). After Gibson assembly, a linear digestion (ExoI, ExoIII, and Lambda Exonuclease) was performed to eliminate non-circularized DNA. The circular Gibson assembly product was cleaned up using SPRI beads. The circularized library was used as template for rolling circle amplification (RCA) using Phi29 polymerase and random hexamer primers. Following the RCA reaction, T7 endonuclease was used to debranch the DNA product. A DNA clean and concentrator column was used to purify the DNA. Purified RCA product was size-selected using a 1% low melt agarose gel. The main band just over the 10 kb marker was excised from the gel and digested with beta-agarase followed by SPRI bead clean up. The cleaned and size selected RCA product was sequenced using the ONT 1D Genomic DNA by Ligation sample prep kit (SQK-LSK109) and MinION flow cells (R9.4.1) following the manufacturer's protocol. Flow cells were nuclease flushed and reloaded with additional library according to the ONT Nuclease Flush protocol.

**cDNA preparation for ONT sequencing of human and mouse**

Library preparation was done from total RNA (200ng) using SQK-PCS110 kit from ONT for PCR-cDNA sequencing. Briefly, cDNA RT adapters were annealed and ligated to full length RNAs using NEBNext® Quick

Ligation Reaction Buffer (NEB B6058) and T4 DNA Ligase (NEB M0202). Bead clean up was done using Agencourt RNAClean XP beads. Purified RNA with CRTA top strand, RT primers, and dNTPs (NEB N0447) were incubated at RT for 15 mins to generate primer-annealed RNA. Reverse transcription and strand-switching was performed with Maxima H Minus RT enzyme in presence of strand-switching primers at $42^0$C for 90 mins followed by heat inactivation at $85^0$C for 5 mins. Reverse transcribed samples were PCR amplified using cDNA primers and LongAmp Hot Start Master Mix (NEB, M0533S). Samples were treated with NEB exonuclease I (NEB, M0293) for 15 mins at $37^0$C to degrade linear single-stranded DNA, followed by enzyme inactivation at $80^0$C for 15 mins. Samples were purified with Agencourt AMPure XP beads. Elution was done with 12 ul of elution buffer. 1ul of libraries was electrophoresed on TapeStation screentapes to assess size distribution, quantity and quality of library. FLO-MIN106D flow cells were primed with EXP-FLP002 kit reagents followed by loading of PCR-cDNA library mixed with rapid adapter F (along with sequencing buffer and loading beads). Sequencing of the library was performed without any size selection using MinION Mk1B devices and MinKNOW software interface.

**Direct RNA (dRNA) preparation for ONT sequencing of human and mouse**

Direct RNA libraries were prepared from 75ug total RNA. RNA samples were poly-A selected using the NEXTFLEX poly-A kit. Purified mRNA was eluted in 12uL nuclease-free $H_2O$. Library preparation was performed on purified mRNA using the SQK-RNA002 kit. Direct RNA RT adapters were annealed and ligated to full-length mRNA using T4 DNA Ligase, NEBNext Quick Ligation Reaction Buffer, and Nanopore's RNA CS. Adapter-ligated mRNA was incubated with dNTPs, 5x first-strand buffer, nuclease-free water, SuperScript IV, and 0.1M DTT to create a cDNA-RNA hybrid. This reverse-transcription (RT) step is recommended by Nanopore to reduce secondary structure formation of the mRNA as it is being sequenced. RTed RNA was purified using RNAClean XP beads. Nanopore adapters were ligated onto the RTed RNA using NEBNext Quick Ligation Reaction Buffer and T4 DNA Ligase. Following RNAClean XP bead cleanup, the libraries were eluted in 21uL of Nanopore's Elution Buffer. 1 uL of each library was quantified on the TapeStation to ensure nucleic acid concentration was at minimum ~200ng. Libraries were loaded into MinION flow cells using the EXP-FLP002 Flow Cell Priming Kit. Libraries were sequenced for 72 hour runs.

**Manatee ONT genome sequencing**

Two µg of genomic DNA in a total volume of 100 µl was fragmented by the g-Tube fragmentation method (Covaris, Woburn, MA, USA) by centrifuging at 6,000x g for 1 min. The large DNA fragments were enriched by using 0.85x volume of Agencourt AMPure XP beads (Beckman Coulter, Brea, CA, USA) in the purification procedure. The enriched DNA fragments were subjected to library preparation with Nanopore Genomic DNA

Ligation Sequencing Kit (Oxford Nanopore Technologies, Oxford, UK) following the manufacturer's protocol. A total of 700 ng of final library product was loaded on a flow cell and sequenced with a Nanopore GridION sequencer (Oxford Nanopore Technologies, Oxford, UK) for a 72-hr run. A total of 5 flow-cell runs were conducted for this project.

**Manatee cDNA PacBio library preparation and sequencing**

Approximately 280 ng of total pooled RNA were processed according to a modified IsoSeq protocol. The sample was spiked-in with the uncapped E2 RNA variant control mix (SIRVs, Lexogen, Cat # 025.03) at a 2.83% mass proportion relative to the total RNA. The resulting mixture was subjected to a globin removal step using the QIAseq FastSelect™- HRM Globin removal reagent (cat # 334376). This kit was designed for globin removal from human, mouse, and rat tissues and was found to perform with various degrees of efficiency on blood from a wide variety of samples of mammalian origin. Globin removal was performed as recommended in the QIAseq FastSelect™- -rRNA HRM -Globin Handbook (Oct 2019) in the NEBNext Ultra II section, except that the high-temperature fragmentation step was omitted. The globin removal reaction (9 µl) contained: 280 ng sample (RNA plus 2.83% SIRVs), QIAseq FastSelect globin removal reagent, 2 µl NEBNext Single Cell RT Primer Mix (NEB #6421), and 2.25 µl of NEBNext Single Cell RT buffer (4x). This mixture was prepared in a 0.2 ml PCR tube and subjected to a stepwise series of 2 min incubations each of 75°C, 70°C, 65°C, 60°C, 55°C, 37°C and 25°C. At this point, the sample was snap-cooled by transferring to a pre-chilled freezer block until ready for the RT and amplification steps. From this point on, cDNA synthesis was done as described in the "Protocol for Low Input RNA: cDNA Synthesis and Amplification" (NEB #E6421) starting on section 2.3. More specifically, the template "RT and Template Switching" reaction consisted of 9 µl of globin-removed RNA, 2.75 µl NEBNext Single Cell RT Buffer (4x), 1 µl of NEBNext Template Switching Oligo, 2 µl of NEBNext Single Cell RT Enzyme Mix and enough water to bring the total to 20 µl. The reaction was incubated in a thermocycler for 90 min at 42 °C and 10 min at 72 °C. The cDNA products were split into four aliquots for PCR amplification (100 µl) reactions containing 2 µl NEBNext Single Cell cDNA PCR Primer, 0.5 µl 10X NEBNext Cell Lysis Buffer, 50 µl NEBNext Single Cell cDNA PCR Master Mix, 5 µl RT and Template Switching reaction and water. Amplified cDNA was purified by AMPure, one round at 0.8 to 1.0 beads to sample ratio and one round at 0.65:1.0 ratio. The yield of amplified cDNA by this modified protocol (300-400 ng) was about 10-fold lower than the standard protocol (i.e., without globin-removal). The average cDNA size was ~1400 bp. When increased amounts of cDNA were desired the cDNA was amplified by 5 additional PCR cycles.

Two preps obtained with the above-described protocol were pooled together and 500 ng were loaded on an electrophoretic lateral fractionation system (ELF, SageScience). Fragments above 2.5 kb were collected, re-amplified (10 cycles), and re-pooled equimolarly with non-size-selected cDNA fragments. This re-pooled cDNA prep is referred to as "enriched cDNA_>2.5kb". Both non_enriched cDNA and enriched cDNA_>2.5kb cDNA were used for SMRT bell library construction starting with 1 µg of cDNA as described in the PacBio IsoSeq protocol 101-070-200 Version 06, September 2018. Briefly, SMRTbell adaptors (Iso-SeqTM) were added using reagents from the PacBio SMRTbell Template Prep Kit 1.0-SPv3 starting with either 200 ng (for enriched cDNA >2.5kb) or 700 ng (for non enriched cDNA). The main steps included: DNA Damage Repair, End Repair, Blunt-end ligation of SMRT bell adaptors, and ExoIII/ExoVII treatment. This procedure resulted in ~25-30% yield. Finally, libraries were eluted in 15 ul of 10 nM Tris HCl, pH 8.0. Library fragment size was estimated by the Agilent TapeStation (genomic DNA tapes), and this data was used for calculating molar concentrations.

The enriched cDNA >2.5 kb library was diffusion-loaded on a single SEQUEL SMRT cell (University of Florida, Interdisciplinary Center for Biotechnology Research (ICBR)-NGS core lab) using a loading concentration of 10 pM, 4-hr pre-extension, 20 hr movies and v3 chemistry reagents (for binding and sequencing). All other steps for sequencing were done according to the recommended protocol by the PacBio SMRT Link Sample Setup and Run Design modules (SMRT Link 6.0).

The non enriched cDNA library was loaded on three Sequel II SMRT cells at University of California, Irvine.

**Manatee cDNA Nanopore library preparation and sequencing**

One hundred and fifty nanograms of total pooled RNA were processed according to a modified ONT cDNA-PCR Sequencing protocol (cDNA-PCR-PCS109, version PCS_9085 v109 revJ Aug 14, 2019). Spike-in and globin depletion treatment was conducted as described for PacBio library preparation. In this case, the globin removal reaction (11 ul) contained: sample (RNA plus SIRVs), globin removal reagent, 1 mM dNTP, 0.2 µM VPN primer from the Nanopore cDNA synthesis protocol (i.e., in place of random primers), and 1X RT buffer (ThermoFisher). This mixture was prepared in a 0.2 ml PCR tube and submitted to a stepwise series of 2 min incubation for each of 75 °C, 70 °C, 65 °C, 60 °C, 55 °C, 37 °C and 25 °C. At this point, the sample was snap-cooled by transferring to a pre-chilled freezer block until ready for the RT and amplification steps. From this point on, cDNA synthesis was done as described in the cDNA-PCR Sequencing (SQK-PCS109) Oxford Nanopore manual starting on page 9 (Version: PCS_90985_v109_revJ_14Aug2019). A single globin removal and cDNA synthesis reaction was split into four PCR reactions for amplification. This process resulted in approximately 2 micrograms of "full-length" cDNA with an average size of ~1800 bp. One size-selected library was constructed by loading 1500 ng of this cDNA on an electrophoretic lateral fractionation system (ELF, SageScience), collecting >2.5 kb fragments, re-ampliying (6 cycles) and re-pooling with non-size-selected cDNA fragments. Adaptor ligation and sequencing were performed according to the cDNA-PCR Sequencing (SQK-PCS109) Nanopore manual. Between 120-140 fmol of cDNA was loaded on a FLO-MIN106D (R9.4 SpotON) flow cell for sequencing on the minION device. Two runs were done on non-size-selected manatee cDNA, while only one run was done on the cDNA that had been enriched with >2.5 kb fragments. Sequencing runs were allowed to proceed for 48 hours.

**Long-read data processing**

Base calling of ONT data from human, mouse and manatee was performed with Guppy 4.2.2 and hac 9.4.1 config file, with default parameters, except: --qscore_filtering --min_qscore 7 (these non-default parameters were used in all cDNA-ONT runs except for R2C2 datasets). Direct RNA basecalling was also performed with Guppy 4.4.2 with the following configurations: --qscore_filtering yes --min_qscore 7 --reverse_sequence yes --u_substitution yes

PacBio full-length non-chimeric (FLNC) reads were generated with CCS 4.2.0 (parameters: --noPolish --minLength=10 --minPasses=3 --min-rq=0.9 --min-snr=2.5), Lima 1.11.0 (parameters: FASTA with the appropriate adapters --isoseq --min-score 0 --min-end-score 0 --min-signal-increase 10 --min-score-lead 0), and Refine 3.3.0 (parameters: --min-polya-length 20 --require-polya).

Consensus R2C2 reads were generated with C3POa v1.0.0 (https://github.com/rvolden/C3POa/tree/gonk) with default options.

Sequence data are provided in FASTQ format. For PacBio data, subreads are provided in unaligned BAM format and for R2C2 data, subreads are provided in FASTQ (**Supplementary Table 2**).

**Supplementary Table 2: Summary statistics for LRGASP data**

| Sample | ES | | | | | |
|---|---|---|---|---|---|---|
| Method | dRNA | cDNA | R2C2 | CapTrap | CapTrap | cDNA |
| Tech | ONT | ONT | ONT | ONT | PacBio | PacBio |
| Platform | MinION | MinION | MinION | MinION | SequelII | SequelII |
| # of Flowcells/SMRT cells | 3 | 3 | 6 | 3 | 3 | 9 |
| # of raw reads | 4,325,200 | 59,746,818 | 7,862,883[1] | 56,684,765 | 9,689,619 | 23,487,808 |
| # of supplied reads | **3,975,725** | **57,055,583** | **5,930,487** | **50,697,997** | **5,090,848** | **8,733,814** |
| # of aligned reads | 3,836,020 | 44,873,564 | 5,914,779 | 49,741,194 | 5,028,403 | 8,199,908 |
| # of aligned reads with adapters | N/A | 40,190,805 | 5,914,779 | 32,206,495 | 5,028,403 | 8,199,908 |
| Median Read length | 830 | 519 | 1,755 | 591 | 903 | 2,090 |

| | | | | | | |
|---|---|---|---|---|---|---|
| Median Identity (Q score) | 9.8 | 12.7 | 18.6 | 12.3 | 21.3 | 20.9 |
| % Directionality | 99.54 | 98.59 | 99.74 | 94.66 | 99.88 | 99.55 |
| % of spike-in reads | 0.71 | 1.02 | 2.03 | 2.41 | 1.77 | 1.85 |
| Pearson r2 (gene level) | 0.99 | 0.99 | 0.98 | 0.99 | 0.98 | 0.97 |

For each sample, replicates were combined when reporting statistics.
[1]R2C2 libraries for ES and WTC11 libraries were multiplexed, and raw reads cannot be demultiplexed directly. Raw read numbers for these libraries are therefore calculated based on the ES/WTC11 ratio of demultiplexed supplied consensus reads and total number of subreads.

| Sample | WTC11 | | | | | |
|---|---|---|---|---|---|---|
| Method | dRNA | cDNA | R2C2 | CapTrap | CapTrap | cDNA |
| Tech | ONT | ONT | ONT | ONT | PacBio | PacBio |
| Platform | MinION | MinION | MinION | MinION | SequelII | SequelII |
| # of Flowcells/SMRT cells | 3 | 3 | 6 | 3 | 3 | 9 |
| # of raw reads | 3,229,571 | 53,463,774 | 6,994,789[1] | 56,730,485 | 13,463,712 | 28,567,150 |
| # of supplied reads | **2,988,430** | **51,194,535** | **5,275,737** | **50,902,303** | **6,399,632** | **7,424,923** |
| # of aligned reads | 2,931,482 | 43,085,527 | 5,271,334 | 49,930,350 | 6,304,610 | 7,373,147 |
| # of aligned reads with adapters | N/A | 37,275,068 | 5,271,334 | 31,348,191 | 6,304,610 | 7,373,147 |
| Median Read length | 854 | 610 | 1,802 | 564 | 864 | 2,209 |
| Median Identity (Q score) | 9.8 | 12.9 | 19.3 | 12.9 | 22.5 | 23.8 |
| % Directionality | 99.76 | 99.11 | 99.92 | 96.28 | 99.92 | 99.67 |
| % of spike-in reads | 0.6 | 1.45 | 2.27 | 2.79 | 2.26 | 2.25 |
| Pearson r2 (gene level) | 0.92 | 0.96 | 0.94 | 0.99 | 0.96 | 0.90 |

For each sample, replicates were combined when reporting statistics.
[1]R2C2 libraries for ES and WTC11 libraries were multiplexed, and raw reads cannot be demultiplexed directly. Raw read numbers for these libraries are therefore calculated based on the ES/WTC11 ratio of demultiplexed supplied consensus reads and total number of subreads.

| Sample | H1-mix | | | | | |
|---|---|---|---|---|---|---|
| Method | dRNA | cDNA | R2C2 | CapTrap | CapTrap | cDNA |
| Tech | ONT | ONT | ONT | ONT | PacBio | PacBio |
| Platform | MinION | MinION | MinION | MinION | SequelII | SequelII |
| # of Flowcells/SMRT cells | 3 | 3 | 6 | 3 | 3 | 6 |
| # raw reads | 4,223,164 | 55,927,828 | 7,093,671 | 54,055,468 | 10,534,880 | 24,290,762 |
| # of supplied reads | **3,969,603** | **52,927,595** | **5,231,255** | **49,883,469** | **5,511,853** | **5,511,357** |
| # of aligned reads | 3,905,742 | 43,026,016 | 5,229,686 | 48,424,901 | 5,436,170 | 5,480,635 |
| # of aligned reads with adapters | N/A | 36,653,422 | 5,229,686 | 28,099,080 | 5,436,170 | 5,480,635 |
| Median Read length | 891 | 619 | 1,782 | 604 | 1,036 | 2,376 |
| Median Identity (Q score) | 10.0 | 12 | 18.7 | 12.4 | 24.3 | 23.7 |
| % Directionality | 99.8 | 99.19 | 99.74 | **76.15**[1] | 99.91 | 99.63 |
| % of spike-in reads | 0.77 | 1.5 | 1.69 | 1.59 | 1.33 | 1.97 |
| Pearson r2 (gene-level) | 0.99 | 0.997 | 0.98 | 0.96 | 0.98 | 0.98 |

[1]Replicate 3 of the H1_mix sample appears to be an outlier among the CapTrap ONT library type. Replicates 1 and 2 show % directionality ~95% similar to what is observed in the other samples for this library type.

| Sample | Manatee | Manatee |
|---|---|---|
| Method | cDNA | cDNA |
| Tech | ONT | PacBio |
| Platform | MinION | Sequel I + Sequel II |
| # of Flowcells/SMRT cells | 3 | 1+3 |
| # of supplied reads | **40,948,571** | **6,883,684** |
| # of aligned reads | 32,833,840 | 6,877,181 |
| # of aligned reads with adapters | 27,381,394 | 6,877,181 |
| Median Read length | 540 | 894 |
| Median Accuracy (Q score) | 12.5 | 25.2 |
| % Directionality | 97.2 | 99.76 |
| % of spike-in reads | **14.05*** | **33.78*** |
| *spike-in percentage is higher than expected | | |

**Reference genome and annotations**

For submissions of transcript models and quantification, transcript annotations and genome models corresponding to GENCODE human v38 and mouse M27 were used. Submissions of challenge predictions were expected to end in Fall 2021, prior to the release of GENCODE human v39 and mouse M28. The newly released GENCODE annotations would, therefore, be used for the evaluations. GRCh38 was the reference genome sequence for human, and GRCm39 was used for mouse. GENCODE annotations were based on these genomes. Please note that GENCODE M25 and earlier annotation releases were based on GRCm38.

**Simulated data**

Simulating RNA reads simply from the reference transcriptome would only allow the assessment reconstruction of known transcript models. Thus, we extended both human and mouse annotations with artificial novel transcripts. To obtain those, we mapped reference transcripts of an undisclosed mammalian organism to the human and mouse genomes and converted the alignments into transcript models using SQANTI[13]. We then arbitrarily selected isoforms of known genes that have only canonical splice sites (GT-AG, GC-AG and AT-AC) and merged them into human and mouse GENCODE Basic annotations.

To generate realistic isoform expression profiles, we selected undisclosed human and mouse long read datasets and quantified them simply by mapping to the reference transcripts with minimap2 v2.17 [34]. Artificial novel isoforms were assigned arbitrary expression values. The generated expression profile was then used for simulating short and long reads. Finally, poly(A) tails were attached to the 3' end of reference transcript sequences prior to running the simulation.

To simulate reads produced by different sequencing platforms we used existing simulation methods. Illumina 2x150bp read pairs were generated with the RSEM simulator[14] using an error model obtained from real RNA-Seq data[15] (accession number ERR1474891).

ONT reads were simulated with NanoSim[16] using pre-trained cDNA and dRNA models available in the package with average error rate of 15.9% (4.8% substitutions, 6.0% deletions, 5.1% insertions) and 11.2% (2.8% substitutions, 5.9% deletions, 2.5% insertions) respectively. NanoSim exploits models trained on real data to produce realistic sequencing error patterns, read length distribution and unaligned sequences at reads ends typical for ONT sequencing. The complete list of Nanopore data characteristics is described in the Trans-NanoSim manuscript[16]. Manual inspection revealed that as the transcript truncation is done randomly in Trans-NanoSim, no 3'/5' bias is introduced. Thus, simulated ONT data may have slightly different coverage profiles compared to the real cDNA-ONT and dRNA-ONT data.

PacBio CCS reads were obtained with IsoSeqSim (https://github.com/yunhaowang/IsoSeqSim), which truncates input reference transcript sequences and uniformly inserts errors according to the given probabilities. Uniform error distribution appears to be a reasonable choice according to the previously developed tool for simulating genomic PacBio reads[17]. Error rate was estimated using real cDNA-PacBio CCS reads obtained in this work as 1.6% (0.4% substitutions, 0.6% deletions, 0.6% insertions). To create a realistic coverage profile, for read truncation in IsoSeqSim we used pre-computed Sequel II truncation probabilities provided along with the package.

To verify generated data we mapped real and simulated reads to the respective genomes with minimap2[5] in spliced mode and computed empirical error rates (**Supplementary Table 3**). As the table shows, with the exception of cDNA-ONT data, error rates appear to be similar. For cDNA-ONT, however, real data sequenced within this work is more accurate compared to NanoSim-generated reads.

**Supplementary Table 3: Error rates in percentage for real and simulated data of different types obtained via read alignment.**

| Data type | Error type | Real data | Simulated |
|---|---|---|---|
| | Mismatches | 0.25 | 0.46 |

| | | | |
|---|---|---|---|
| cDNA-PacBio | Insertions | 0.57 | 0.57 |
| | Deletions | 0.45 | 0.64 |
| | Total | 1.27 | 1.67 |
| cDNA-ONT | Mismatches | 2.5 | 4.2 |
| | Insertions | 3.3 | 5.1 |
| | Deletions | 1.6 | 4.1 |
| | Total | 7.4 | 13.4 |
| dRNA-ONT | Mismatches | 7.0 | 6.0 |
| | Insertions | 5.2 | 5.4 |
| | Deletions | 2.9 | 2.1 |
| | Total | 15.1 | 13.5 |

We simulated two datasets containing reads from all 3 platforms listed above but with slightly different properties. Human datasets were simulated with 100 million Illumina read pairs, 30 million cDNA-ONT and 10 million PacBio reads. Mouse datasets also contained 100 million Illumina read pairs, but equal amounts of PacBio CCS and dRNA-ONT reads were generated (20 million sequences each).

To allow users to simulate their own data, the methods described above are implemented as simple command-line scripts which are available at https://github.com/LRGASP/lrgasp-simulation/.

**CAGE data of WTC11 samples for validation of transcript 5' ends**

To validate novel 5' ends, we used a recently generated deep coverage CAGE data on the WTC11 line.

The 15 µg of WTC11 RNAs from each biological replicate, ENCODE BioSample Accession ENCBS944CBA and ENCBS474NOC, were used for the single strand (ss)CAGE library preparation followed in the *Low Quantity Single Strand CAGE* protocol[18]. Briefly, the 15 µg RNAs were aliquoted to 5 µg in three tubes and reverse transcribed to cDNAs with random primers, and the capped RNA-cDNA hybrids were trapped by streptavidin beads. The single strand cDNAs were released from the beads and ligated to the Illumina adaptors with Index, and 1080 amols of the cap-trapped single strand cDNAs from each biological replicate were sequenced by Illumina HiSeq Rapid SBS Kits v2 (SR, 150 cycles, 1 lane for each).

CAGE data from WTC11 samples was produced for the validation of transcript 5' ends and was not released until after the close of the challenge submissions. CAGE data was obtained from two RNA biological replicates of WTC11, using the exact same RNA that was used for long-read sequencing.

The 15 µg of WTC11 RNAs from each biological replicate, ENCODE BioSample Accession ENCBS944CBA and ENCBS474NOC, were used for the single strand (ss)CAGE library preparation described in the published protocol[18]. Briefly, the 15 µg RNAs were aliquoted to 5 µg in three tubes and reverse transcribed to cDNAs with random primers, and the RNA-cDNA hybrids were cap-trapped by the streptavidin beads. The single strand cDNAs were released from the beads and ligated to the Illumina adaptors with an index. 1080 amols of the cap-trapped single strand cDNAs from each biological replicate were sequenced by Illumina HiSeq Rapid SBS Kits v2 (SR, 150 cycles, 1 lane for each), producing approximately 40 million reads per sample.

**QuantSeq of human and mouse samples for validation of transcript 3' ends**

QuantSeq data (3' end sequencing) from challenge 1 and 2 samples was produced for validation of 3' ends and was not released until the close of the challenge submissions. Data was be obtained from two RNA biological replicates of WTC11, from the same exact RNA used for long-read sequencing.
To validate novel polyadenylation sites, we collected poly(A)-seq data using the Quant-Seq method from Lexogen, which can map poly(A) sites *de novo*.

#### LRGASP Data QC

Initial quality control (QC) metrics were determined for the LRGASP data (**Supplementary Fig. 1)**. Reads (ONT cDNA, dRNA, CapTrap) or consensus reads (PacBio cDNA and CapTrap and ONT R2C2) were aligned to the human or mouse genome as appropriate using minimap2 with the following parameters: -ax splice --secondary=no -G 400k. For each data type, the reads and their resulting alignments in sam format were parsed for the following parameters:

1) Number of aligned reads
2) Number of aligned reads with adapters on both ends

    For ONT dRNA this is not applicable as this workflow does not attach an adapter to the 5' end of molecules. For ONT cDNA and CapTrap this percentage was determined by pyChopper. For all other data types, all provided reads are assumed to have adapters on both ends as the pre-processing pipelines (lima and C3POa) discard reads otherwise.

3) median read length

   measured by the number of aligned bases (matches or mismatches)

4) median accuracy

   measured by matches/(matches+mismatches+indels)),

5) Percent of aligned reads where the orientation of the reads as determined by 5’ and 3’ adapter sequences agree with the direction of the read alignment

   determined by minimap2 through splice site context (calculated only for the subset of reads with splice alignments with the ts:A: flag in their sam entry),

6) Percent of reads originating from spike-in molecules

   determined by alignment to the SIRVomeERCC FASTA entry in the genome sequence files

7) Pearson correlation between replicates

   determined by quantifying gene expression for each replicate and calculating the pearson r value based on those expression values.

#### GENCODE benchmarks and computational evaluation

Full manual annotation was undertaken on 50 selected loci on both the human and mouse reference genomes. Transcript models were only annotated during this exercise based on their support from long transcriptomic datasets generated by the consortium specifically for LRGASP. No transcript annotation was based on transcriptomic data from externally produced datasets, although annotators used any publicly available orthogonal data to aid in the interpretation of aligned consortium data. For example, Fantom 5 CAGE datasets were used to help identify transcription start sites and transcript 5' ends, and RNA-seq-supported introns derived from high-throughput reanalysis pipelines such as Recount were used to support putative introns identified in the alignments of long transcriptomic data.

Manual annotation was performed according to the guidelines of the HAVANA (Human And Vertebrate Analysis aNd Annotation) group[19,20]. Transcriptomic data was aligned to the human and mouse reference genome using appropriate methods. The benefits of aligning the transcriptomic data using multiple methods were tested to reduce the impact of alignment errors and artifacts.

Annotators also tppl advantage of local alignment tools integrated into annotation software to give further alternative views of alignments and improve annotation accuracy. Transcript models where manually extrapolated from the alignments by annotators using the otter annotation interface [21]. Alignments were navigated using the Blixem alignment viewer [22,23] and, where required, visual inspection of the dot-plot output from the Dotter tool[24] was used to resolve any alignment with the genomic sequence that was unclear or absent from Blixem. Short alignments (<15 bases) that cannot be visualized using Dotter were detected using Zmap DNA Search[24] (essentially a pattern matching tool). The construction of exon-intron boundaries required the presence of canonical splice sites (defined as GT-AG, GC-AG and AT-AC) and any deviations from this rule were given clear explanatory tags (for example non-canonical splice site supported by evolutionary conservation). All non-redundant splicing transcripts at an individual locus were used to build transcript models, and all alternatively spliced transcripts were assigned an individual biotype based on their putative functional potential. Once the correct transcript structure was ascertained, the protein-coding potential of the transcript was determined based on its context within the locus, similarity to known protein sequences, the sequences of orthologous and paralogous proteins, candidate coding regions (CCRs) identified by PhyloCSF, evidence of translation from mass spectrometry and Ribo-seq data, the presence of Pfam functional domains, the presence of possible alternative ORFs, the presence of retained intronic sequence, and the likely susceptibility of the transcript to nonsense-mediated mRNA decay (NMD).
Although the annotation of transcript functional biotype and CDS is not required of submitters, they were added to transcripts as a matter of routine manual annotation and may be used to investigate the detection or non-detection of groups of transcripts by submitters. When necessary, annotations were checked by a second annotator to ensure the completeness and consistency of annotation between the genes annotated for LRGASP and the remainder of the Ensembl/GENCODE gene set.

#### Computational pipeline description from submitters

##### Challenge 1

**Name: Bambu**
**Description:** Bambu trains a transcript discovery model on each sample using the known reference annotation to predict if novel aligned reads are likely to represent full-length transcripts. This optimizes several parameters relevant to transcript discovery and reduces this down to a single tunable parameter which is customized to the specific sample transcriptome, the novel discovery rate (NDR). By ranking novel transcripts with the NDR Bambu can extend the annotations across a large range of sensitivity and precision.
**Version:** The development version of Bambu 0.9.1 was used during LRGASP.
**Team:** Göke, Genome Institute of Singapore
**URL:** https://github.com/GoekeLab/bambu and https://bioconductor.org/packages/bambu/
**Citations:** Chen, Y., Sim, A., Wan, Y.K. *et al.* Context-aware transcript quantification from long-read RNA-seq data with Bambu. *Nat Methods* (2023). https://doi.org/10.1038/s41592-023-01908-w
**Config:** Bambu was run with the following parameters (NOTE: many of these parameters are deprecated in the updated version of Bambu) min.txScore.multiEx
on = 0, min.txScore.singleExon = 1, max.txNDR = 0.2, min.geneScore = 0, min.samp
leNumber = 1, remove.subsetTx = FALSE, min.readFractionByGene = 0). Please see the documentation for best practices when using Bambu's latest version.
**Notes:** Bambu uses an NDR threshold which allows the user to influence the sensitivity and precision of novel transcripts. By default Bambu is calibrated to report a precision selection of novel transcripts, with a threshold

of 0.2 being used in the LRGASP challenge. However a higher (and less stringent) NDR value can be used to greatly increase the number of transcripts reported by Bambu.
**Funding:** A.S, Y.C, J.J.X.L, Y.K.W, and J.G are supported by funding from the Agency for Science, Technology and Research (A*STAR)and the National Medical Research Council (NMRC).

**Name: FLAIR**
**Description:** FLAIR is a tool for RNA isoform exploration with long reads with the optional pairing of short reads. FLAIR contains modules for correcting noisy reads, isoform definition, isoform quantification, and analysis of alternative splicing in long read data.
**Version:** 2
**Team:** Brooks Lab, University of California, Santa Cruz
**URL:** https://github.com/BrooksLabUCSC/flair/
**Citations:** Detecting haplotype-specific transcript variation in long reads with FLAIR2
Alison D Tang, Eva Hrabeta-Robinson, Roger Volden, Christopher Vollmers, Angela N Brooks
bioRxiv 2023.06.09.544396; https://doi.org/10.1101/2023.06.09.544396
**Config:** We ran the FLAIR2 isoform discovery pipeline using the flair-collapse module with the --annotation_reliant, --check_splice, and --stringent parameters. The short-and-long submissions used short-read data to identify confident splice junctions.
**Notes:** FLAIR2 run with the --annotation_reliant argument invokes an alignment of the reads to an annotated transcriptome first, followed by novel isoform detection. When including --check_splice, this enforces higher quality matching specifically around each splice site for read-to-isoform assignment steps.
**Funding:** A.D.T. is supported by NIH NHGRI F31 HG010999. This work was also supported by NIH NIGMS R35GM138122 (A.N.B.).

**Name: FLAMES**
**Description:** A tool developed for full-length transcript quantification, mutation and splicing analysis of long-read RNA-seq data.
**Version:** 0.1.0
**Team:** Ritchie Lab, Walter and Eliza Hall Institute of Medical Research
**URL:** https://github.com/LuyiTian/FLAMES
**Citations:** Tian, L., Jabbari, J.S., Thijssen, R. *et al.* Comprehensive characterization of single-cell full-length isoforms in human and mouse with long-read sequencing. *Genome Biol* **22**, 310 (2021). https://doi.org/10.1186/s13059-021-02525-6
**Config:** The following parameters were adjusted in the configuration file for LRGASP: has_UMI:false, Min_sup_cnt:10, Min_cnt_pct:0.01, strand_specific:1, remove_incomp_reads:5, no_flank:true, min_tr_coverage:0.75, and min_read_coverage:0.75.
**Notes:** FLAMES provides a default set of parameters, which can be changed in the configuration JSON file. The 'pipeline_parameters' section specifies the steps to be executed in the pipeline (all by default). The 'isoform_parameters' section determines the results of isoform detection.
**Funding:** FLAMES development was supported by funding from the Chan Zuckerberg Initiative DAF, an advised fund of Silicon Valley Community Foundation (Grant No. 2019-002443 to M.E.R.) and Australian National Health and Medical Research Council (NHMRC) Investigator Grant (2017257 to M.E.R.).

**Name: Iso_IB**
**Description:** An IsoSeq evidence-based approach to predict gene models, alternative splices and isoforms using a custom path from the cDNA-cupcake workflow

**Version:** CD-HIT: 4.8.1; minimap2: 2.17-r974; collapse_isoforms_by_sam.py: 22.0.0; gffread: v0.12.7;
**Team:** Integrative Bioinformatics, National Institute of Environmental Health Sciences
**URL:** https://github.com/weizhongli/cdhit
**Config:** The following parameters were used for each step of the workflow CD-HIT: cd-hit-est -M 0 -c 0.99 -G 0 -aL 0.90 -AL 100 -aS 0.99 -AS 30; minimap2: -ax splice -t 40 -uf --secondary=no -C5; collapse_isoforms_by_sam.py --fq -s; gffread -E -T -o-;
**Notes:** The reference workflow and repo for cDNA_cupcake can be found at https://github.com/Magdoll/cDNA_Cupcake. The conda instance can be also found at https://github.com/PacificBiosciences/IsoSeq. In this workflow, no reference gene model/isoforms/alternative splicing prediction/annotation was used to enhance or validate the models. The resulted set was run in de novo mode, solely on the basis of evidence sequences obtained from the IsoSeq sequencing data provided by the consortium.
**Funding:** This work was supported by the Intramural Research Program of the National Institute of Environmental Health Sciences ZIC ES103371

**Name: IsoQuant**
**Description:** IsoQuant is a reference-based approach for transcript discovery and quantification using long RNA reads. Since version 3.0 it also supports annotation-free transcript discovery.
**Version:** 2.0.0
**Team:** Center for Algorithmic Biotechnology, Saint Petersburg State University
**URL:** https://github.com/ablab/IsoQuant
**Citations:** Prjibelski, A.D., Mikheenko, A., Joglekar, A. *et al.* Accurate isoform discovery with IsoQuant using long reads. *Nat Biotechnol* (2023). https://doi.org/10.1038/s41587-022-01565-y
**Config:** --data_type nanopore for ONT data, --data_type pacbio_ccs for PacBio data
**Notes:** The tool can be installed via conda. Reads can be provided in BAM or in FASTQ format. In the later case they will be automatically mapped using minimap2.
**Funding:** St. Petersburg State University (grant ID: 94030965), European Research Council (ERC) under the European Union's Horizon 2020 research and innovation programme (grant agreement No. 851093, SAFEBIO)

**Name: IsoTools**
**Description:** IsoTools is a Python module for Long Read Transcriptome Sequencing (LRTS) analysis, providing transcriptome reconstruction, filtering, and quantification, along with explorative analysis, alternative splicing detection both on isoform as well as splice-site level and differential analysis. IsoTools version > 0.3.2 also includes functional annotation and interpretation including ORF prediction, NMD prediction, and domain annotation.
**Version:** 0.2.5
**Team:** Herwig Lab, Max Planck Institute for Molecular Genetics
**URL:** https://isotools.readthedocs.io
**Citations:** Matthias Lienhard and others, IsoTools: a flexible workflow for long-read transcriptome sequencing analysis, *Bioinformatics*, 2023;, btad364, https://doi.org/10.1093/bioinformatics/btad364
**Config:** We filtered transcripts based on splice-site matches and read-count thresholds. Transcripts with novel splice sites must be supported by at least 5 reads and at least 5% of the total gene read support, and are discarded if they contain more than 50% A content downstream or a direct repeat of length 6 or longer at a novel splice-site. Transcripts with all splice-sites matching reference splice-sites need support from at least 2 reads and at least 2% of the total gene read support. For more configuration details, see https://github.molgen.mpg.de/lienhard/LRGASP_IsoTools.

**Notes:** The filtering strategy used in the IsoTools pipeline has a significant impact on the number and nature of identified transcripts. While the strategy described above was developed specifically for the challenge, it may not be optimal for all datasets and research questions. Please consult the documentation for detailed instructions on how to use IsoTools for your specific dataset and research question. Note that the filtering syntax has been improved since pre-release version 0.2.5 used for the submission.
**Funding:** This work was supported by the German Research Foundation (DFG) with the grant HE4607/7-1 and the Federal Ministry of Education and Research with the grant SafetyNet (161L0242A).

**Name: LyRic**
**Description:** An end-to-end workflow for long-read based transcriptome annotation, visualization and analysis
**Version: v1.0.4**
**Team:** Guigó Lab, Centre de Regulació Genòmica
**URL**: https://github.com/guigolab/LyRic
**Citations**: CapTrap-Seq: A platform-agnostic and quantitative approach for high-fidelity full-length RNA transcript sequencing
Silvia Carbonell-Sala, Julien Lagarde, Hiromi Nishiyori, Emilio Palumbo, Carme Arnan, Hazuki Takahashi, Piero Carninci, Barbara Uszczynska-Ratajczak, Roderic Guigo
bioRxiv 2023.06.16.543444; https://doi.org/10.1101/2023.06.16.543444
**Config:** LyRic was run with default parameters except as specified in its LRGASP configuration file, available at https://github.com/guigolab/LyRic/blob/master/config_LRGASP.json. The LyRic working directory was staged for LRGASP data processing using the *setup_LRGASP.sh* script (https://github.com/guigolab/LyRic/blob/master/setup_LRGASP.sh) All PacBio sequencing data were re-processed from BAM files using the *pb_gen* pipeline (https://github.com/guigolab/pb_gen) with default parameters, except for adapter sequences ('*PB_ADAPT*' parameter), which were changed according to LRGASP organizers' adapter sequence specifications. PacBio and ONT FASTQ files were processed by LyRic using a *filter_SJ_Qscore* of 30 and 10, respectively. Briefly, *filter_SJ_Qscore* represents the minimum average Phred sequencing quality of read sequences +/- 3 nts around all their splice junctions for a spliced read to be considered for transcript model building. See LyRic documentation (https://guigolab.github.io/LyRic/documentation.html), and input LRGASP sample annotation file (https://github.com/guigolab/LyRic/blob/master/sample_annotations_LRGASP.tsv) for more details.
Reads were merged into transcript models with the *tmerge* utility (https://github.com/guigolab/tmerge) using the following two-step nested approach. First, reads were merged separately within replicates, requiring a minimum of two reads supporting each transcript model. The resulting transcript models were then merged again, this time across all three replicates of each LRGASP sample, requiring replicate-specific transcript models to be detected at least once in every replicate.
**Notes:** LyRic's output transcript models are completely agnostic to any pre-existing reference annotation. In other words, LyRic does not adjust the coordinates of the transcript models it produces based on a reference annotation.
**Funding:** National Human Genome Research Institute of the US National Institutes of Health (grant 2U24HG007234-09). We acknowledge support of the Spanish Ministry of Science and Innovation to the EMBL partnership, Centro de Excelencia Severo Ochoa and CERCA Programme / Generalitat de Catalunya.

**Name: Mandalorion**
**Description:** Mandalorion uses PacBio or ONT-based R2C2 consensus reads. It aligns those reads using minimap2, then parses those alignments to generate models of isoforms. Mandalorion then generates read-based consensus sequences for each isoform using pyabpoa and racon tools. Mandalorion then aligns these isoform

consensus sequences and filters the isoforms based on these alignments and their abundance. The final isoforms are reported as both FASTA and PSL/GTF files.
**Version: v3.6**
**Team**: Vollmers Lab, University of California, Santa Cruz
**URL:** https://github.com/christopher-vollmers/Mandalorion
**Citations:** Identifying and quantifying isoforms from accurate full-length transcriptome sequencing reads with Mandalorion, Roger Volden, Kayla Schimke, Ashley Byrne, Danilo Dubocanin, Matthew Adams, Christopher Vollmers bioRxiv 2022.06.29.498139 https://doi.org/10.1101/2022.06.29.498139
**Config:** all runs were performed using the "-R 3" and "-I 150" flags, setting minimum read number and minimum length of isoforms, respectively.
**Notes:** Mandalorion only uses individual splice sites in any provided GTF annotation file. It discards information on how splice sites are connected into splice junctions. It also ignores annotated transcription start sites and poly(A) sites when constructing isoforms. Because Mandalorion doesn't heavily rely on information in the annotation file, performance is very similar for novel or annotated isoforms and the ends of identified isoforms will agree with read alignments rather than annotated TSS and poly(A) sites. Another consequence of not relying heavily on an annotation file is that Mandalorion will not "assemble" isoforms that are longer than the provided reads, which is obvious for some of the "long SIRV" data.
**Funding:** NIH/NIGMS R35GM133569 to Christopher Vollmers

**Name: Spectra**
**Description:** Spectra is a tool to build gene models based on full-length cDNA reads, not fragmented or incomplete ones, through a guide of genome alignments. The resulting gene models are entirely (end-to-end) supported with one or more observations of reads.
**Version:** v0.1a
**Team:** Hideya Kawaji, Tokyo Metropolitan Institute of Medical Science
**URL:** https://github.com/hkawaji/spectra
**Config:** PacBio's consensus sequence is computed with a set of helper scripts bundled in the repository.
**Notes:** The development of this tool was motivated by the notion that the read counts of long RNA molecules are depleted in the contributed data sets, even with the best protocol. Discoveries supported by experimental evidence were maximized by selecting high-quality data and setting a minimum read count threshold.
**Funding:** AMED (Grant Number 21kk0305013h0002).

**Name: StringTie2**
**Description:**StringTie2 is a guided transcriptome assembler, able to assemble either short or long RNA-seq read data, even in the absence of a reference annotation. Since version 2.2.0 it is also capable of handling mixed transcriptomic data that includes both short and long RNA-seq reads sequenced from the same sample.
**Version**: 2.2.1
**Team:** Pertea Lab, Johns Hopkins University
**URL:** https://github.com/gpertea/stringtie
**Citation:** Kovaka, S., Zimin, A.V., Pertea, G.M. *et al.* Transcriptome assembly from long-read RNA-seq alignments with StringTie2. *Genome Biol* **20**, 278 (2019). https://doi.org/10.1186/s13059-019-1910-1
**Config**: -L option for long read alignments, or --mix for both short and long read alignments
**Notes:** An up-to-date documentation, and usage manual can be consulted at https://ccb.jhu.edu/software/stringtie/index.shtml**.**
**Funding:** NSF grant DBI-1759518

**Name: TALON_LAPA**

**Description:** Minimap2, STAR, TranscriptClean, TALON, LAPA

- TranscriptClean: TranscriptClean corrects common long-read sequencing artifacts such as microindels and mismatches.
  - Noncanonical splice junctions, if not provided in the input set of splice junctions, will be corrected to the nearest canonical splice sites in places where possible, otherwise they are discarded.

- TALON: TALON annotates long reads to their transcripts of origin, quantifies the expression of annotated transcripts, and filters novel transcript models based on reproducibility and evidence of internal priming
  - Any read meeting coverage and identity filters with new splice sites will be constitute a novel model
  - Any read with an intron chain that matches that of a reference model or a novel model that's already been cataloged in the database that also 5'/3' ends within a certain distance of the cataloged model will be assigned to that model. Otherwise, it will be used to create a new model with new 5'/3' ends.
  - Transcripts are quantified simply by counting the number of reads that belong to each cataloged transcript model.
  - Unannotated (novel) transcripts are filtered for reproducibility and for those that display evidence of internal priming (see settings used in the config section)

- LAPA: LAPA is used to refine the 3' end calls made by TALON.
  - In the analysis, we called poly(A)-sites with LAPA and updated the 3' ends of the transcripts based on those poly(A)-sites with the most likely poly(A)-site assigned to each intron chain (transcript) based on the number of reads ending in the poly(A)-cluster.
  - Transcripts with low expression may not map to any poly(A)-cluster. In this case, we chose the longest end as a 3' end of the transcript.

**Version:** LAPA 0.0.1
**Team:** Mortazavi Lab, University of California, Irvine
**URL:** https://github.com/lh3/minimap2, https://github.com/mortazavilab/TranscriptClean/, https://github.com/mortazavilab/TALON/, https://github.com/mortazavilab/lapa/, https://github.com/mortazavilab/lrgasp-talon/
**Citations:** Analysis of alternative polyadenylation from long-read or short-read RNA-seq with LAPA
Muhammed Hasan Çelik, Ali Mortazavi
bioRxiv 2022.11.08.515683; https://doi.org/10.1101/2022.11.08.515683
A technology-agnostic long-read analysis pipeline for transcriptome discovery and quantification
Dana Wyman, Gabriela Balderrama-Gutierrez, Fairlie Reese, Shan Jiang, Sorena Rahmanian, Stefania Forner, Dina Matheos, Weihua Zeng, Brian Williams, Diane Trout, Whitney England, Shu-Hui Chu, Robert C. Spitale, Andrea J. Tenner, Barbara J. Wold, Ali MortazavibioRxiv 672931; https://doi.org/10.1101/672931
**Config:** All arguments are default unless otherwise specified
**TranscriptClean:** --canonOnly --spliceJns {short read + GENCODE splice junctions for the short+long, otherwise just GENCODE splice junctions)
**talon_label_reads:** --ar 20
**talon_initialize_database:** --5p 500 --3p 300
**talon:** --cov 0.9 --identity 0.8
**talon_filter_transcripts:** --maxFracA 0.5 --minCount 2 minDatasets 2
**Notes:**

- Known problems with submission:

- For the ONT reads (dRNA, cDNA, CapTrap), the adapters were not removed in contrast to the PacBio reads. This affected our coverage and identity filters which require certain mapping rates to include each read. Thus, many ONT reads with the adapters still on had large unalignable regions and were discarded, leading to erroneous results.
- For the ONT cDNA and cDNA CapTrap, the reads are unstranded while our tools expect only 5'-3' oriented alignments as input. This gave us a lot of antisense transcripts that should have been real transcripts and were discarded by the filter that takes novelty into account.
- Spike-in models were not included in our reference transcriptome annotation. Therefore, we treated them like novel transcripts and were subject to the reproducibility filter; leading many to them being erroneously discarded. Our performance on the spike-ins represents our efforts to do reference-free annotation and should be interpreted as such. This likely explains the discrepancies between our performance on the spike-in and the simulated data.
- TALON labels transcript models using the SQANTI novelty categories. Users have the option of filtering out all ISMs, even those that pass the reproducibility and internal priming filters. Internally, we have found that our precision on spike-ins is much better when we do so (data available on request). However for this submission we did not, leading to the expected high levels of ISMs reported in our dataset.
- In general, we ran all our tools with default parameters regardless of the protocol which is similar to what the average user can do.

**Funding:** NHGRI UM1HG009443

#### Challenge 2

**Name: Bambu**
**Description:** Bambu performs quantification after performing transcript discovery.
**Version:** The development version of Bambu 0.9.1 was used during LRGASP.
**Team:** Göke, Genome Institute of Singapore
**URL:** https://github.com/GoekeLab/bambu and https://bioconductor.org/packages/bambu/
**Citations:** Chen, Y., Sim, A., Wan, Y.K. *et al.* Context-aware transcript quantification from long-read RNA-seq data with Bambu. *Nat Methods* (2023). https://doi.org/10.1038/s41592-023-01908-w
**Config:** Bambu was run with the following parameters (NOTE: many of these parameters are deprecated in the updated version of Bambu): NDR = 0.2, and opt.em=list(degradationBias = TRUE). Please see the documentation for best practices when using Bambu's latest version.
**Notes:** We use a development version of Bambu for quantification in LRGASP challenge. Now Bambu uses an improved quantification model. We recommend using the latest version of Bambu.
**Funding:** A.S, Y.C, J.J.X.L, Y.K.W, and J.G are supported by funding from the Agency for Science, Technology and Research (A*STAR) and the National Medical Research Council (NMRC).

**Name: FLAIR**
**Description:** FLAIR is a tool for RNA isoform exploration with long reads with the optional pairing of short reads. FLAIR contains modules for correcting noisy reads, isoform definition, isoform quantification, and analysis of alternative splicing in long read data.
**Version:** 2
**Team:** Brooks Lab, University of California, Santa Cruz

**Citations:** Detecting haplotype-specific transcript variation in long reads with FLAIR2
Alison D Tang, Eva Hrabeta-Robinson, Roger Volden, Christopher Vollmers, Angela N Brooks
bioRxiv 2023.06.09.544396; https://doi.org/10.1101/2023.06.09.544396
**URL:** https://github.com/BrooksLabUCSC/flair
**Config:** We ran the flair-quantify module with the --stringent and --tpm parameters.
**Notes:** None
**Funding:** A.D.T. is supported by NIH NHGRI F31 HG010999. This work was also supported by NIH NIGMS R35GM138122 (A.N.B.).

**Name: FLAMES**
**Description:** A tool developed for full-length transcript quantification, mutation and splicing analysis of long-read RNA-seq data.
**Version:** 0.1.0
**Team:** Ritchie Lab, Walter and Eliza Hall Institute of Medical Research
**URL:** https://github.com/LuyiTian/FLAMES
**Citations:** Tian, L., Jabbari, J.S., Thijssen, R. *et al.* Comprehensive characterization of single-cell full-length isoforms in human and mouse with long-read sequencing. *Genome Biol* **22**, 310 (2021). https://doi.org/10.1186/s13059-021-02525-6
**Config:** The following parameters were adjusted in the configuration file for LRGASP: has_UMI:false, Min_sup_cnt:10, Min_cnt_pct:0.01, strand_specific:1, remove_incomp_reads:5, no_flank:true, min_tr_coverage:0.75, and min_read_coverage:0.75.
**Notes:** FLAMES provides a default set of parameters, which can be changed in the configuration JSON file. The 'pipeline_parameters' section specifies the steps to be executed in the pipeline. The 'isoform_parameters' section determines the results of isoform detection.
**Funding:** FLAMES development was supported by funding from the Chan Zuckerberg Initiative DAF, an advised fund of Silicon Valley Community Foundation (Grant No. 2019-002443 to M.E.R.) and Australian National Health and Medical Research Council (NHMRC) Investigator Grant (2017257 to M.E.R.).

**Name: IsoQuant**
**Description:** IsoQuant is a reference-based approach for transcript discovery and quantification using long RNA reads. Since version 3.0 it also supports annotation-free transcript discovery.
**Version:** 2.0.0
**Team:** Center for Algorithmic Biotechnology, Saint Petersburg State University
**URL:** https://github.com/ablab/IsoQuant
**Citations:** Prjibelski, A.D., Mikheenko, A., Joglekar, A. *et al.* Accurate isoform discovery with IsoQuant using long reads. *Nat Biotechnol* (2023). https://doi.org/10.1038/s41587-022-01565-y
**Config:** --data_type nanopore for ONT data, --data_type pacbio_ccs for PacBio data
**Notes:** The tool can be installed via conda. Reads can be provided in BAM or in FASTQ format. In the latter case they will be automatically mapped using minimap2.
**Funding:** St. Petersburg State University (grant ID: 94030965), European Research Council (ERC) under the European Union's Horizon 2020 research and innovation programme (grant agreement No. 851093, SAFEBIO)

**Name: IsoTools**
**Description:** IsoTools is a Python module for Long Read Transcriptome Sequencing (LRTS) analysis, providing transcriptome reconstruction, filtering, and quantification, along with explorative analysis, alternative splicing detection both on isoform as well as splice-site level and differential analysis. IsoTools version > 0.3.2

also includes functional annotation and interpretation including ORF prediction, NMD prediction, and domain annotation.
**Version:** 0.2.5
**Team:** Herwig Lab, Max Planck Institute for Molecular Genetics
**URL:** https://isotools.readthedocs.io
**Citations:** Matthias Lienhard and others, IsoTools: a flexible workflow for long-read transcriptome sequencing analysis, *Bioinformatics*, 2023;, btad364, https://doi.org/10.1093/bioinformatics/btad364
**Config:** To account for transcript length distribution differences between PacBio Isoseq data and reference annotation, TPM values were normalized using a lognormal model fit to the reference and observed transcript length distributions.The quotient of the two distributions served as the normalization factor. Due to poor fit in the left tail, the factor was set constant for the first 1% percentile of the observed transcript length model.
For more configuration details, see https://github.molgen.mpg.de/lienhard/LRGASP_IsoTools.
**Funding:** This work was supported by the German Research Foundation (DFG) with the grant HE4607/7-1 and the Federal Ministry of Education and Research with the grant SafetyNet (161L0242A).

**Name: NanoSim**
**Description:** NanoSim is a fast and scalable read simulator that captures the technology-specific features of ONT data, and allows for adjustments upon improvement of nanopore sequencing technology.
**Version:** 3.0.0 (See "Notes" below)
**Team**: Birol Lab, University of British Columbia, Vancouver
**URL:** https://github.com/bcgsc/lrgasp_nanosim
**Citations:** Chen Yang and others, NanoSim: nanopore sequence read simulator based on statistical characterization, *GigaScience*, Volume 6, Issue 4, April 2017, gix010, https://doi.org/10.1093/gigascience/gix010
**Config:** python read_analysis.py quantify -t THREADS -e trans -rt REFERENCE_TRANSCRIPTS.fasta -i READS.fastq -o nanosim
**Notes:** Please note that this is a completely separate repository that was branched from the primary repository at https://github.com/bcgsc/NanoSim.
**Funding:** This work was supported by Genome Canada and Genome BC (281ANV); and by the National Human Genome Research Institute of the National Institutes of Health (R01HG007182). Scholarship funding was provided by the University of British Columbia, and the Natural Sciences and Engineering Research Council of Canada.

**Name: TALON_LAPA**
**Description:** Minimap2, STAR, TranscriptClean, TALON, LAPA

- TranscriptClean: TranscriptClean corrects common long-read sequencing artifacts such as microindels and mismatches.
    - Noncanonical splice junctions, if not provided in the input set of splice junctions, will be corrected to the nearest canonical splice sites in places where possible, otherwise they are discarded.

- TALON: TALON annotates long reads to their transcripts of origin, quantifies the expression of annotated transcripts, and filters novel transcript models based on reproducibility and evidence of internal priming
    - Any read meeting coverage and identity filters with new splice sites will be constitute a novel model

- Any read with an intron chain that matches that of a reference model or a novel model that's already been cataloged in the database that also 5'/3' ends within a certain distance of the cataloged model will be assigned to that model. Otherwise, it will be used to create a new model with new 5'/3' ends.
- Transcripts are quantified simply by counting the number of reads that belong to each cataloged transcript model.
- Unannotated (novel) transcripts are filtered for reproducibility and for those that display evidence of internal priming (see settings used in the config section)

- LAPA: LAPA is used to refine the 3' end calls made by TALON.
    - In the analysis, we called poly(A)-sites with LAPA and updated the 3' ends of the transcripts based on those poly(A)-sites with the most likely poly(A)-site assigned to each intron chain (transcript) based on the number of reads ending in the poly(A)-cluster.
    - Transcripts with low expression may not map to any poly(A)-cluster. In this case, we chose the longest end as a 3' end of the transcript.

**Version:** LAPA 0.0.1
**Team:** Mortazavi Lab, University of California, Irvine
**URL:** https://github.com/lh3/minimap2, https://github.com/mortazavilab/TranscriptClean/, https://github.com/mortazavilab/TALON/, https://github.com/mortazavilab/lapa/, https://github.com/mortazavilab/lrgasp-talon/
**Citations:** Analysis of alternative polyadenylation from long-read or short-read RNA-seq with LAPA
Muhammed Hasan Çelik, Ali Mortazavi
bioRxiv 2022.11.08.515683; https://doi.org/10.1101/2022.11.08.515683
A technology-agnostic long-read analysis pipeline for transcriptome discovery and quantification
Dana Wyman, Gabriela Balderrama-Gutierrez, Fairlie Reese, Shan Jiang, Sorena Rahmanian, Stefania Forner, Dina Matheos, Weihua Zeng, Brian Williams, Diane Trout, Whitney England, Shu-Hui Chu, Robert C. Spitale, Andrea J. Tenner, Barbara J. Wold, Ali Mortazavibio Rxiv 672931; https://doi.org/10.1101/672931
**Config:**
All arguments are default unless otherwise specified.
**TranscriptClean:** --canonOnly --spliceJns {short read + GENCODE splice junctions for the short+long, otherwise just GENCODE splice junctions)
**talon_label_reads:** --ar 20
**talon_initialize_database:** --5p 500 --3p 300
**talon:** --cov 0.9 --identity 0.8
**talon_filter_transcripts:** --maxFracA 0.5 --minCount 2 minDatasets 2
**Notes:**

- Known problems with submission:
    - For the ONT reads (dRNA, cDNA, CapTrap), the adapters were not removed in contrast to the PacBio reads. This affected our coverage and identity filters which require certain mapping rates to include each read. Thus, many ONT reads with the adapters still on had large unalignable regions and were discarded, leading to erroneous results.
    - For the ONT cDNA and cDNA CapTrap, the reads are unstranded while our tools expect only 5'-3' oriented alignments as input. This gave us a lot of antisense transcripts that should have been real transcripts and were discarded by the filter that takes novelty into account.
    - Spike-in models were not included in our reference transcriptome annotation. Therefore, we treated them like novel transcripts and were subject to the reproducibility filter; leading many to them being erroneously discarded. Our performance on the spike-ins represents our efforts to do

reference-free annotation and should be interpreted as such. This likely explains the discrepancies between our performance on the spike-in and the simulated data.
  - TALON labels transcript models using the SQANTI novelty categories. Users have the option of filtering out all ISMs, even those that pass the reproducibility and internal priming filters. Internally, we have found that our precision on spike-ins is much better when we do so (data available on request). However for this submission we did not, leading to the expected high levels of ISMs reported in our dataset.
  - In general, we ran all our tools with default parameters regardless of the protocol which is similar to what the average user can do.

**Funding:** NHGRI UM1HG009443

**Name: RSEM**
**Description:** For comparison against long-read quantification, the LRGASP organizers ran RSEM against the GENCODE reference transcripts.
**Version**: RSEM v1.3.3
**Team:** Dewey Lab
**URL:** https://deweylab.github.io/RSEM/
**Citations:** Li, B., Dewey, C.N. RSEM: accurate transcript quantification from RNA-Seq data with or without a reference genome. *BMC Bioinformatics* **12**, 323 (2011). https://doi.org/10.1186/1471-2105-12-323
**Config:**

- Preparing Reference Sequences
  **RSEM-1.3.3**/rsem-prepare-reference
  **-gtf** reference annotation(GTF file) \
  **--bowtie2** \
  reference genome(fasta file) \
  rsem_index(index path)

- Calculating Expression Values
  **RSEM-1.3.3**/rsem-calculate-expression \
  **-p** 20 \
  **--bowtie2** \
  **--sort-bam-by-coordinate** \
  **--sort-bam-memory-per-thread** 10G \
  **--paired-end** paired_end_1.fq paired_end_2.fq \
  rsem_index(index_path) \
  quantification_results(output_quantification_result_path)

**Notes:**

RSEM quantification consisted of two main steps:
(1) Preparing Reference Sequences;
(2) Calculating Expression Values.
The specific parameter design is described as above. The versions of RSEM and bowtie2 are:
Bowtie2: version 2.4.1
RSEM: v1.3.3

**Challenge 3**

**Name: Bambu**
**Description:** Bambu uses a reference annotation trained model to predict if novel aligned reads are likely to represent full-length transcripts. As Challenge 3 involved not using reference annotations, Bambu instead uses its internal pretrained model.
**Version:** The development version of Bambu 0.9.1 was used during LRGASP. Bambu's current version is 3.0.8.
**Team:** Göke, Genome Institute of Singapore
**URL:** https://github.com/GoekeLab/bambu and https://bioconductor.org/packages/bambu/
**Citations:** Chen, Y., Sim, A., Wan, Y.K. *et al.* Context-aware transcript quantification from long-read RNA-seq data with Bambu. *Nat Methods* (2023). https://doi.org/10.1038/s41592-023-01908-w
**Config:** Bambu was run with the following parameters (NOTE: many of these parameters are deprecated in the updated version of Bambu) min.txScore.multiExon = 0, min.txScore.singleExon = 1, max.txNDR = 0.7, min.geneScore = 0, min.sampleNumber = 1, remove.subsetTx = FALSE, min.readFractionByGene = 0. Please see the documentation for best practices when using Bambu's latest version.
**Notes:** To achieve more sensitive results, Bambu can be run with a less stringent NDR threshold. Without a reference annotation, Bambu cannot perform NDR calibration. Therefore, NDR threshold selection instead uses the undying prediction score and should be made much more sensitive than when reference annotations are provided to Bambu. When performing *de novo* transcript discovery, if there is a similar but well annotated dataset for a related organism, it is possible to retrain the model used so that it is more applicable. **Funding:** A.S, Y.C, J.J.X.L, Y.K.W, and J.G are supported by funding from the Agency for Science, Technology and Research (A*STAR) and the National Medical Research Council (NMRC).

**Name: IsoQuant**
**Description:** IsoQuant is a reference-based approach for transcript discovery and quantification using long RNA reads. Since version 3.0 it also supports annotation-free transcript discovery. As version 2.0 did not support annotation-free transcript discovery, IsoQuant was launched using GTF obtained with StringTie2 (2.15).
**Version:** 2.0.0
**Team:** Center for Algorithmic Biotechnology, Saint Petersburg State University
**URL:** https://github.com/ablab/IsoQuant
**Citations:** Prjibelski, A.D., Mikheenko, A., Joglekar, A. *et al.* Accurate isoform discovery with IsoQuant using long reads. *Nat Biotechnol* (2023). https://doi.org/10.1038/s41587-022-01565-y
**Config:** --data_type nanopore for ONT data, --data_type pacbio_ccs for PacBio data. StringTie2 was launched using -L option.
**Notes:** IsoQuant can be installed via conda. Reads can be provided in BAM or in FASTQ format. In the later case they will be automatically mapped using minimap2.
**Funding:** St. Petersburg State University (grant ID: 94030965), European Research Council (ERC) under the European Union's Horizon 2020 research and innovation programme (grant agreement No. 851093, SAFEBIO)

**Name: RNA-Bloom**
**Description:** RNA-Bloom uses a reference-free approach to assemble long transcriptomics reads.
**Version:** 1.4.3
**Team:** Birol Lab, University of British Columbia, Vancouver
**URL:** https://github.com/bcgsc/RNA-Bloom

**Citations:** Nip, K.M., Hafezqorani, S., Gagalova, K.K. *et al.* Reference-free assembly of long-read transcriptome sequencing data with RNA-Bloom2. *Nat Commun* **14**, 2940 (2023). https://doi.org/10.1038/s41467-023-38553-y
**Config:** For long + short reads (ONT + Illumina): `-t 48 -ntcard -artifact -long fulllength.fastq rescued.fastq unclassified.porechop.fastq -sef paired_1.fastq paired_2.fastq unpaired_1.fastq unpaired_2.fastq -fpr 0.005 -indel 20 -p 0.75 -Q 15 -overlap 100 -length 150`. For long reads only (ONT): `-t 48 -ntcard -artifact -long fulllength.fastq rescued.fastq unclassified.porechop.fastq -fpr 0.005 -indel 20 -p 0.75 -Q 15 -overlap 100 -length 150`. For short reads only (Illumina), `-t 48 -ntcard -stranded -left paired_1.fastq -right paired_2.fastq -rcr -sef unpaired_1.fastq -ser unpaired_2.fastq -fpr 0.005 -k 25 -indel 2 -q 15 -Q 15 -length 150`.
**Notes:** All tools and resources used to generate the assemblies are documented at https://github.com/bcgsc/lrgasp_birol
**Funding:** The development of RNA-Bloom was supported by Genome Canada and Genome British Columbia (243FOR); the National Institutes of Health (2R01HG007182-04A1); the Natural Sciences and Engineering Research Council of Canada (NSERC); and the Canadian Institutes of Health Research (CIHR).

**Name: rnaSPAdes**
**Description:** RnaSPAdes is a part of SPAdes package — a toolkit for various sequence assembly pipelines. RnaSPAdes is a transcriptome assembler preliminary designed for Illumina data, but can handle long-read data as a supplementary information.
**Version:** 3.15.3
**Team**: Pevzner Lab, St. Petersburg Academic University
**URL:** https://github.com/ablab/spades
**Citations**: Prjibelski, A., Antipov, D., Meleshko, D., Lapidus, A., & Korobeynikov, A. (2020). Using SPAdes de novo assembler. Current Protocols in Bioinformatics, 70, e102. https://doi.org/10.1002/cpbi.102
**Config:** (indicate if any special commands for specific library types)
**Notes:** Sequencing data was provided using the appropriate options from the user manual. Strand specificity was indicated using --ss rf flag.
**Funding:** St. Petersburg State University (grant ID: 94030965).

#### Method changes from registered report phase 1

While the great majority of the analysis indicated in the Registered Report are present in the final version of the manuscripts, some modifications were introduced. These are listed in the **Supplementary Table 4**.

**Supplementary Table 4. Method Changes**

| Modification | Section | Description |
|---|---|---|
| Orthogonal data | Introduction | Orthogonal data for human was Illumina, CAGE and Quant-seq. We did not generate or use ChIP-seq or ATAC-seq |
| Analysis of novel gene transcripts | Challenge 1 | Exhaustive analysis of transcripts in novel genes and SQANTI categories other than FSM, ISM, NIC, NNC was not performed, although these numbers are included in these supplementary tables. |

| | | |
|---|---|---|
| %LRC analysis | Challenge 1 | Fraction of the transcript model sequence length mapped by one or more long reads (%LRC) analysis for GENCODE annotated transcripts was not performed as GENCODE transcripts were called de novo by annotators, rather than validated from submissions, to avoid biases. |
| Percentage of Expressed Transcripts (PET) | Challenge 2 | A newly added metric used to characterize the percentage of truly expressed transcripts in SIRV-set4 data. |
| Abundance Recovery Rate (ARR) skipped | Challenge 2 | Considering the redundancy of multiple evaluation metrics, the ARR metric was skipped. |
| ROC-based metrics skipped | Challenge 2 | Current real data cannot obtain the truly differentially expressed transcripts (i.e., the ground truth) due to lack of qPCR validation. So, ROC-based metrics were skipped. |
| Assessment without a reference genome | Challenge 3 | This assessment was not launched and Challenge 3 restricted to transcript identification without a reference annotation but with a reference genome. This turned out to be quite challenging already. |
| Number of transcripts/loci | Challenge 3 | This information was not asked to submitters, as initially planned, but computed during the analysis |
| Blast2GO analysis skipped | Challenge 3 | Functional annotation of transcript predictions was skipped due to long computing times. BUSCO analysis is a proxy for this analysis |
| Validation of TM with manatee 454 data skipped | Challenge 3 | We found limitations when accessing the data. |
| Validation of quantitative levels of isoforms for Challenge 2 | Validation | We could not find a tractable and cost-manageable technique for providing reliable isoform quantity estimates. Isoform-specific qPCR was cost prohibitive and for the targets of interest, not amenable to analysis. |
| Target selection for Challenge 1. | Validation | We added additional test categories including novel and suspect transcripts from the GENCODE manual annotation. We report data for the human WTC11 sample. No mouse targets were validated due to insufficient material and resources. |

#### Primers-JuJu bulk PT_PCR primer design tool

To facilitate the design of a large number of RT-PCR primers for validating a subset of the predicted isoforms, we developed a tool called Primers-Juju. It provides a semi-automated interface between the visualization of the transcript models in the UCSC browser and the Primer3 primer design package.

The design process starts with a UCSC track hub containing the consolidated transcript models from all pipelines. Unique features for transcripts to validate are identified by visualization. A pair of genomic regions that could contain a primer pair that would amplify the targeted transcript are manually defined. The region may be within an exon or two exons spanning a splice junction. These regions are marked using the UCSC Browser region highlight facility. **Supplementary Fig. 15** shows an example of specifying the design region primers for a unique intron.

The genomic coordinates of the pair of regions are recorded in a spreadsheet, along with the transcript identifier. Additional transcripts that would also be amplified by the primers may be included for validation.

Primers-Juju provides a command line tool that takes the specification spreadsheet with multiple targets and transcript annotations and does primer design and validation. The input specifications are validated against the targeted transcripts, with minor adjustments for inexact bounds. The sequence for the transcripts is obtained from the genomic sequence of the exons and the regions converted to transcript coordinates.

The Primer3 programmatic library is given each transcript sequence and region pair and will attempt to design a stable primer pair to amplify this transcript, returning up to five possible primer pairs per region. The in-Silico PCR command line tool is used to check for potential off-target primer pairs. Queries are done against the genome sequences and transcriptome sequences, which consist of the known annotations as well as the LRGASP consolidated transcripts.

Primers-Juju generates additional tracks for the hub with primer pairs and amplicon sequences, as seen in **Supplementary Fig. 16.** It also produces reports and recommends the most stable primer with no off-target hits to order.

Primers-Juju source code is available at https://github.com/diekhans/PrimerS-JuJu/ and was developed by The University of California, Santa Cruz (UCSC) and El Centre de Regulació Genòmica (CRG).

### Data and code availability

Biological sequencing data is available from the ENCODE Portal (https://www.encodeproject.org/) and are described in the RNA-Seq data matrix (**Extended Data Table 1**).

Experimental data used in GENCODE manual evaluation:

- ssCAGE WTC11
  GEO: GSE185917 https://www.ncbi.nlm.nih.gov/geo/query/acc.cgi?acc=GSE185917
- WTC11 QuantSeq

ENCODE: ENCSR322MWL https://www.encodeproject.org/experiments/ENCSR322MWL/
GEO: GSE219685 https://www.ncbi.nlm.nih.gov/geo/query/acc.cgi?acc=GSE219685

- H1 QuantSeq
  ENCODE: https://www.encodeproject.org/experiments/ENCSR813AOB/
  GEO: GSE219788 https://www.ncbi.nlm.nih.gov/geo/query/acc.cgi?acc=GSE219788
- H1-DE QuantSeq
  ENCODE: ENCSR198UNH https://www.encodeproject.org/experiments/ENCSR198UNH/
  GEO: GSE219571 https://www.ncbi.nlm.nih.gov/geo/query/acc.cgi?acc=GSE219571

Reads generated for experimental validation are available in the Sequence Read Archive (SRA):

- SRR24680099: Manatee whole blood RT-PCR mixed with human WTC11
- SRR24680098: Human WTC11 mixed with manatee whole blood RT-PCR
- SRR23881262: LRGASP WTC-11 experimental validation RT-PCR/ONT

LRGASP-specific code is available in the GitHub LRGASP project (https://github.com/LRGASP/):

- LRGASP submission commands, which includes documentation on submission metadata and format: https://github.com/LRGASP/lrgasp-submissions/
- Read simulation pipeline:
  https://github.com/LRGASP/lrgasp-simulation/
- Challenge 1 evaluation code:
  https://github.com/LRGASP/lrgasp-challenge-1-evaluation/
- Challenge 2 evaluation code:
  https://github.com/LRGASP/lrgasp-challenge-2-evaluation/
- Challenge 3 evaluation code:
  https://github.com/LRGASP/lrgasp-challenge-3-evaluation/
- Code to generate Challenge 1 figures for the paper:
  https://github.com/LRGASP/Challenge1_Figures_Code/
- Code to generate Challenge 2 figures for the paper:
  https://github.com/LRGASP/Challenge2_Figures_Code/
- Code to generate Challenge 3 figures for the paper:
  https://github.com/LRGASP/Challenge3_Figures_Code/

Other data provide to participants, participant submissions, evaluation results, and data for generating the paper figures is available from the LRGASP project on Synapse (https://www.synapse.org/#!Synapse:syn25683370)

- LRGASP reference genomes and annotations:
  Files/data/references/ (syn25683363)
  https://www.synapse.org/#!Synapse:syn25683363
- LRGASP simulation data:
  Files/data/simulation/ (syn25007472)
  https://www.synapse.org/#!Synapse:syn25683370
- Participant submissions:
  Files/submissions/ (syn51603880)
  https://www.synapse.org/#!Synapse:syn51603880
- Evaluations results for all challenges:
  Files/results/ (syn26964358)

https://www.synapse.org/#!Synapse:syn26964358

- Data for generate Challenge 1 figures for the paper:
  https://www.synapse.org/#!Synapse:syn51602848
  Files/paper/Challenge1_Figures_Data.zip (syn51602848)
- Data for generate Challenge 2 figures for the paper:
  Files/paper/Challenge2_Figures_Data.zip (syn51401823)
  https://www.synapse.org/#!Synapse:syn51401823
- Data for generate Challenge 3 figures in paper:
  Files/paper/Challenge3_Figures_Data.zip (syn51603662)
  https://www.synapse.org/#!Synapse:syn51603662
- Spearman correlations of TPM for each Challenge 2 pipeline:
  Files/paper/Spearman_correlation_of_TPM_values.zip (syn51666778)
  https://www.synapse.org/#!Synapse:syn51666778
- Non-redundant genome annotations derived from the submitted annotations:
  Files/ annotations/ (syn25007472)
  https://www.synapse.org/#!Synapse:syn51728612

### Supplementary Figures

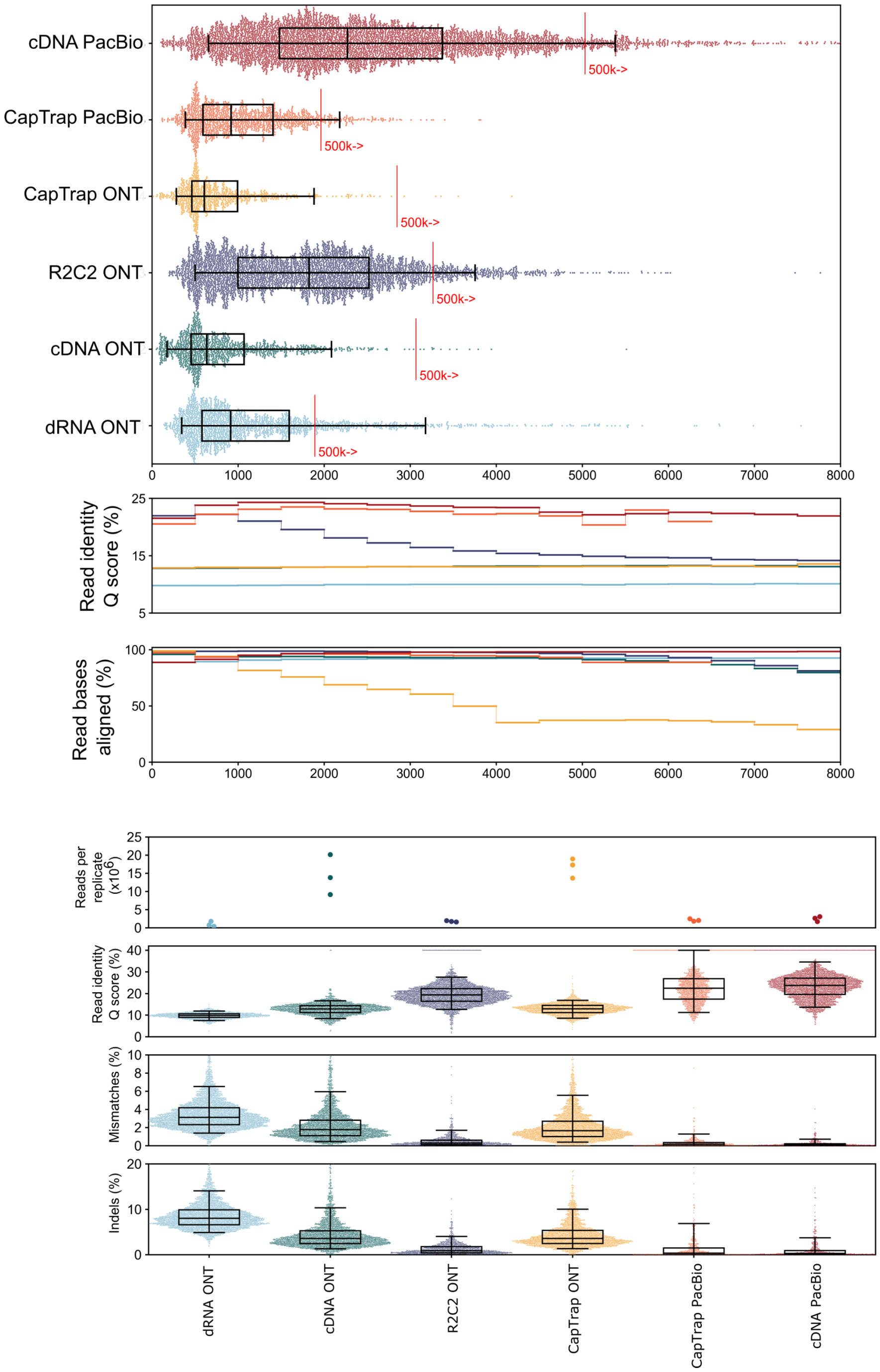


**Supplementary Fig. 1. Summary of LRGASP Data.**

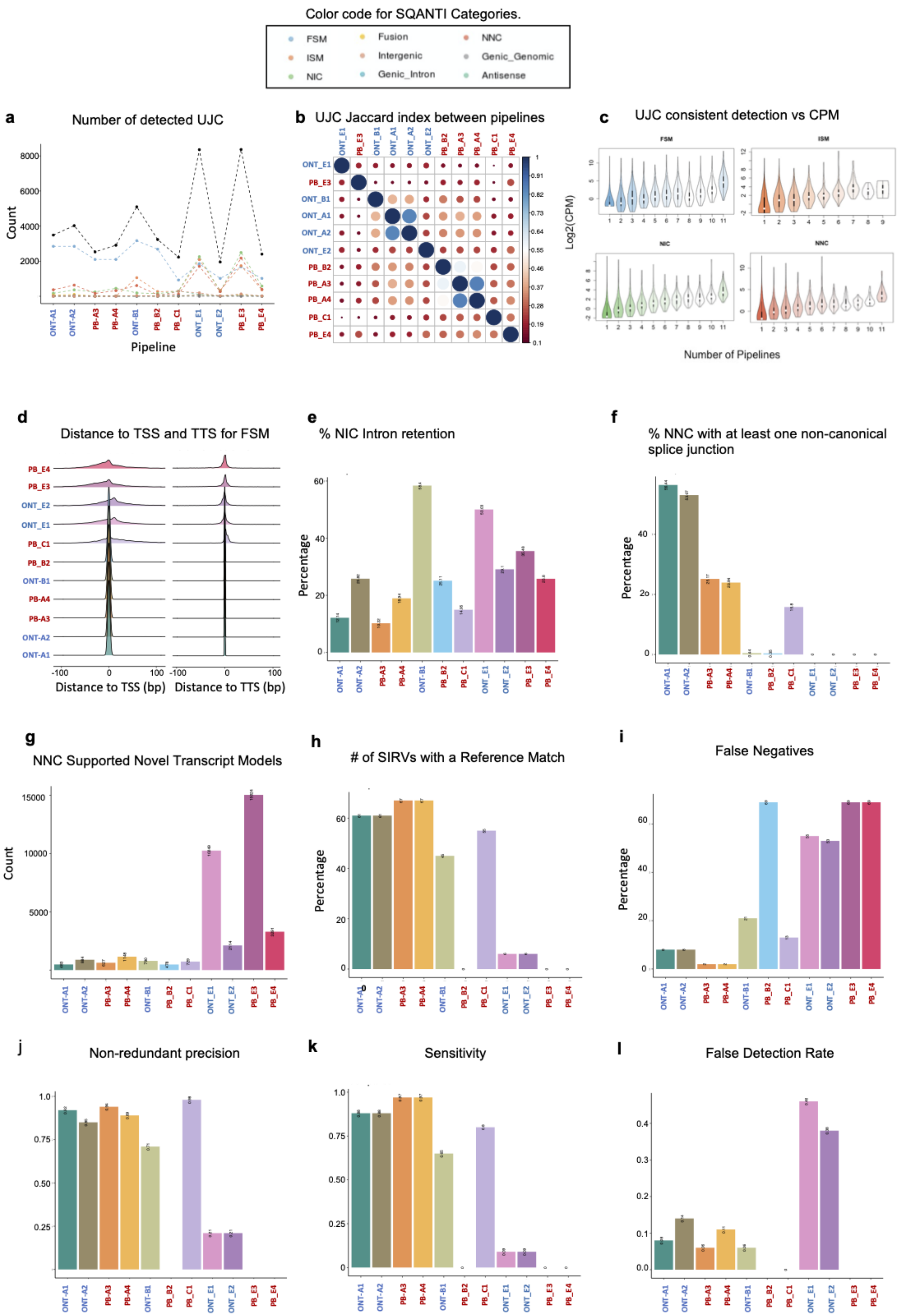


**Supplementary Fig. 2. Example of Challenge 1 evaluation metrics on Pilot Data.** Pipeline names indicate the sequencing platform (PB: PacBio, ONT: Oxford Nanopore), the undisclosed analysis software (A,B,C,E) and the undisclosed software parameters (1 to 4). a-g: Results on whole transcriptome data. a: Number of detected unique junction chains (UJC) per pipeline for each SQANTI category. Black line indicates the total number of UJCs. b: Jaccard index plot for similarity in UJC detection between pipelines. c: Violin plots of the expression values of UJC as a function of the number of pipelines where they were detected, broken down by SQANTI categories. d: Density plot of the distance from the transcript model 5´and 3´genome mapping positions to the Transcription Start Site (TSS) and Transcription Termination Site (TTS), respectively, of the corresponding reference transcript. Wider distributions indicate greater deviations from the reference. e: Percentage of Novel In Catalogue transcript models showing Intron Retention. f: Percentage of Novel Not in Catalogue transcript models containing at least one non-canonical splice junction. g: Percentage of Novel Not in Catalogue transcripts classified as Supported Novel Transcript Models. h-l: results on SIRV data. h Number of SIRVs with at least one Reference Match. i: False Negatives. j: Non-redundant precision. k: Sensitivity, i: False Detection Rate. See **Supplementary Table 1** for metrics definitions. log2(CPM) log2 of the median counts per million of the UJC in the pipelines where it was detected. FSM: Full Splice Match; ISM: Incomplete Splice Match; NIC: Novel In Catalogue; NNC: Novel Not in Catalogue.

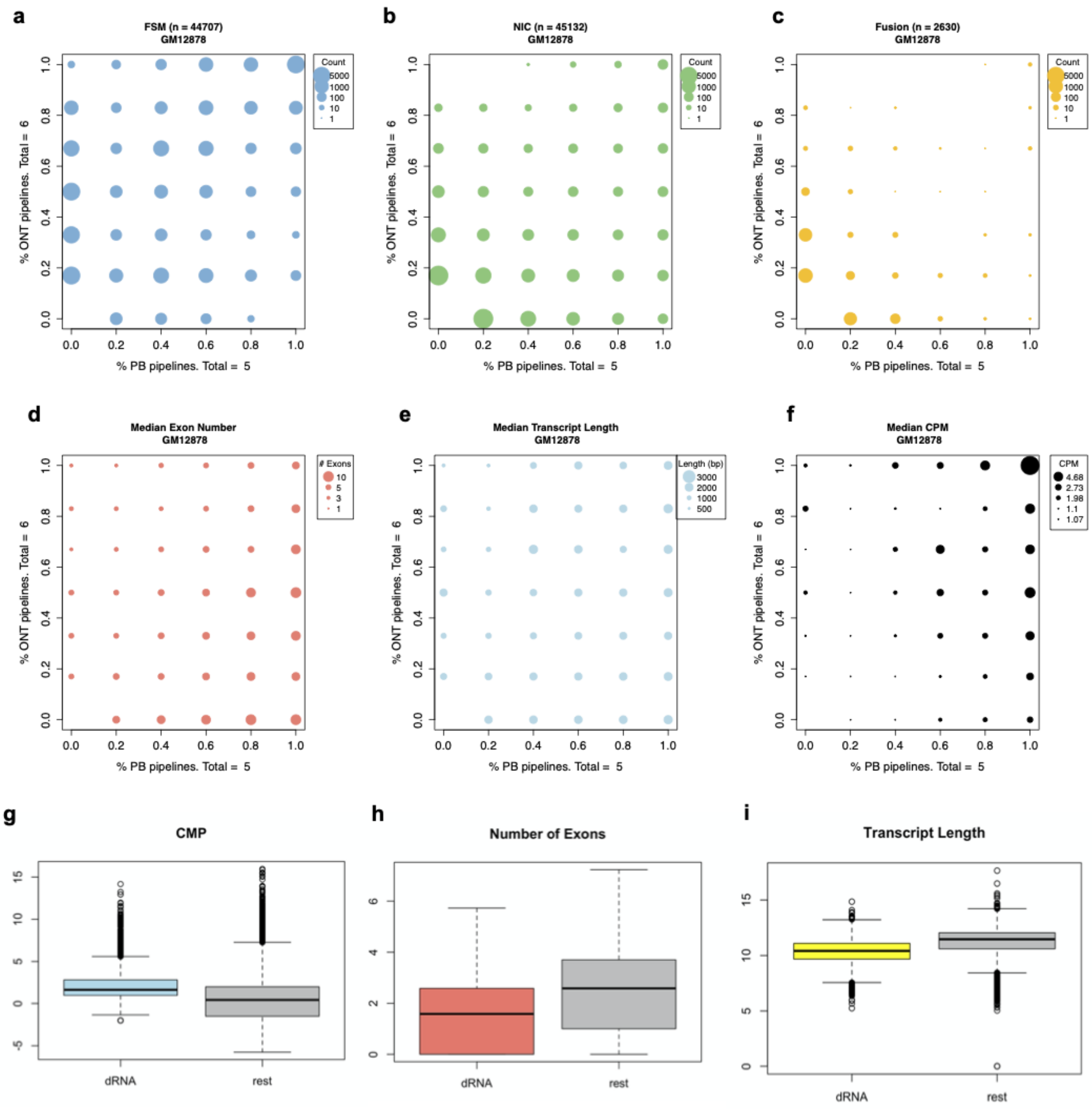


**Supplementary Fig. 3. Examples of UJC barcode-supported analyses for GM12878 data.** Number of detected FSM a:, NIC b: and Fusion transcript c: as a function of the percentage of PacBio and Nanopore pipelines detecting the UJC. Median Exon Number d: Median Transcript Length e: and Median Counts Per Millions f: of the UJC as a function of the percentage of PacBio and Nanopore pipelines detecting the UJC. Comparison of the distribution of Counts Per Million g:, Number of Exons h: and Transcript Length i: for transcript models detected exclusively by directRNA Nanopore sequencing with those detected by all other pipelines. FSM: Full Splice Match; Match, NIC: Novel In Catalogue.

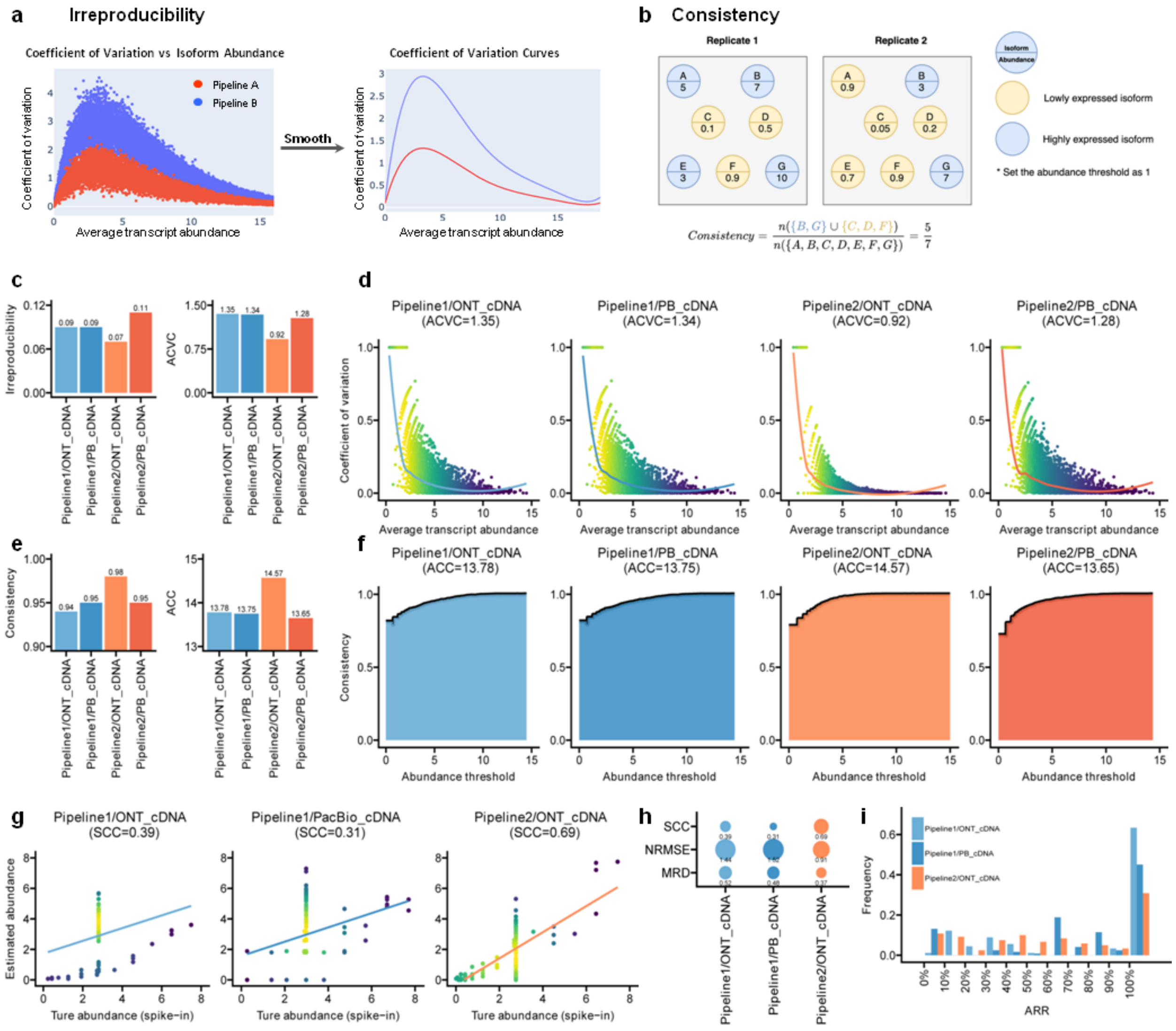


**Supplementary Fig. 4. Performance evaluation with the proposed metrics for Challenge 2 in the published lrRNA-seq GM12878 from PacBio and ONT sequencing.** a and b illustrate the calculation of irreproducibility and consistency. a: By fitting the coefficient of variation versus average isoform abundance into a smooth curve, it can be shown that Pipeline A has lower coefficient of variation and higher reproducibility. b: By setting an expression threshold (i.e. 1 in this toy example), we can define which set of genes express (in blue) or not (in yellow). This statistic is to measure the consistency of the expressed gene sets between replicates. c-i: perform the irreproducibility, consistency and SIRV transcript analysis of two pipelines in lrRNA-seq GM12878 data. The evaluation results reveal Pipeline 2 has the best performance on the GM12878 ONT cDNA samples. c-d: Irreproducibility and ACVC scores. e-f: Consistency and ACC scores. g-i:, SCC, NRMSE, MRD and ARR for SIRV data.

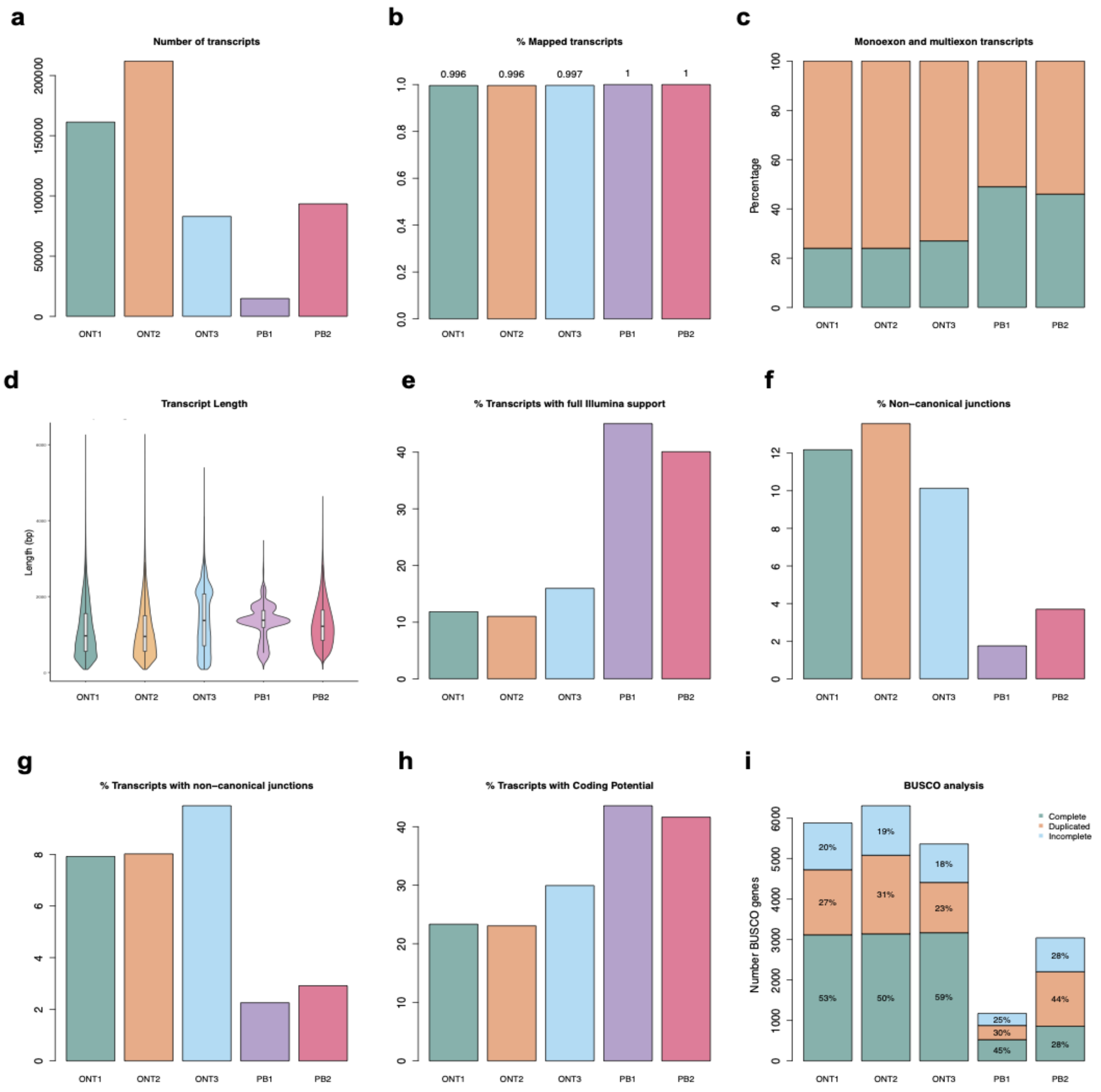


**Supplementary Fig. 5. Example of Challenge 3 evaluation metrics on Pilot Data**. Pipelines in bars represent Isoseq3 analysis on different subsets of the LRGASP manatee Nanopore (ONT1, ONT2, ONT3) and PacBio (PB1, PB2) data.a: Total number of detected transcripts. b: Percentage of successfully mapped transcript to the LRGASP manatee genome assembly. c: Distribution of mono and multi-exon transcript models. Multi-exon shown in orange. d: Distribution of transcript lengths. e: Percentage of transcript models with short reads support at all splice junctions. f: Percentage of non-canonical junctions, g: Percentage of Transcript models with at least one non canonical junction. h: Percentage of transcript models with coding potential. i BUSCO analysis results indicating the percentage of BUSCO genes identified as complete, duplicated or incomplete sequences.

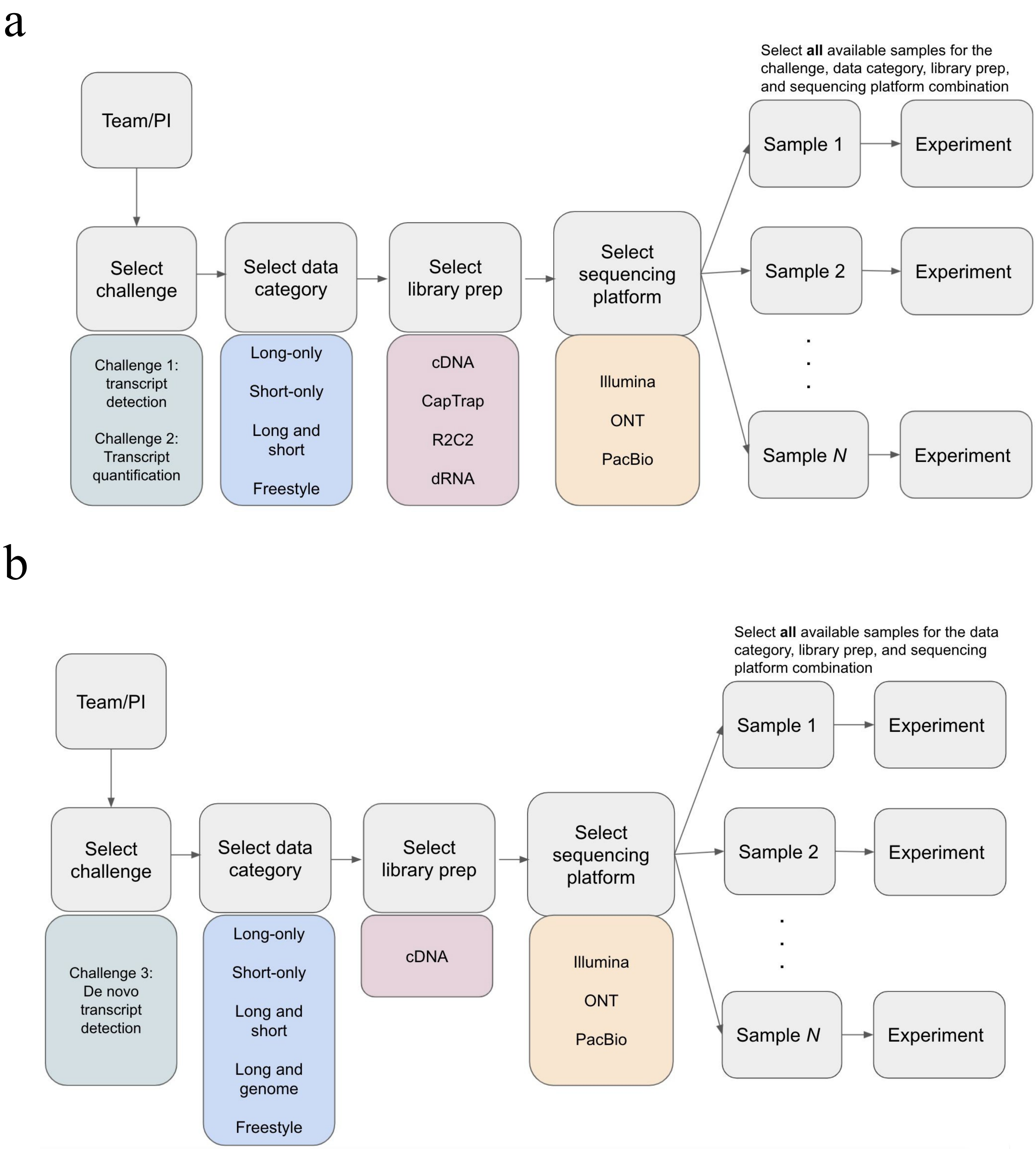


**Supplementary Fig. 6: Challenge submission.** a, Overview of submissions to Challenges 1 and 2. Each entry was derived from a specific data category, library prep, and sequencing platform combination. All available samples for the selected combination must be included in an entry b, Overview of submissions for Challenge 30.34.

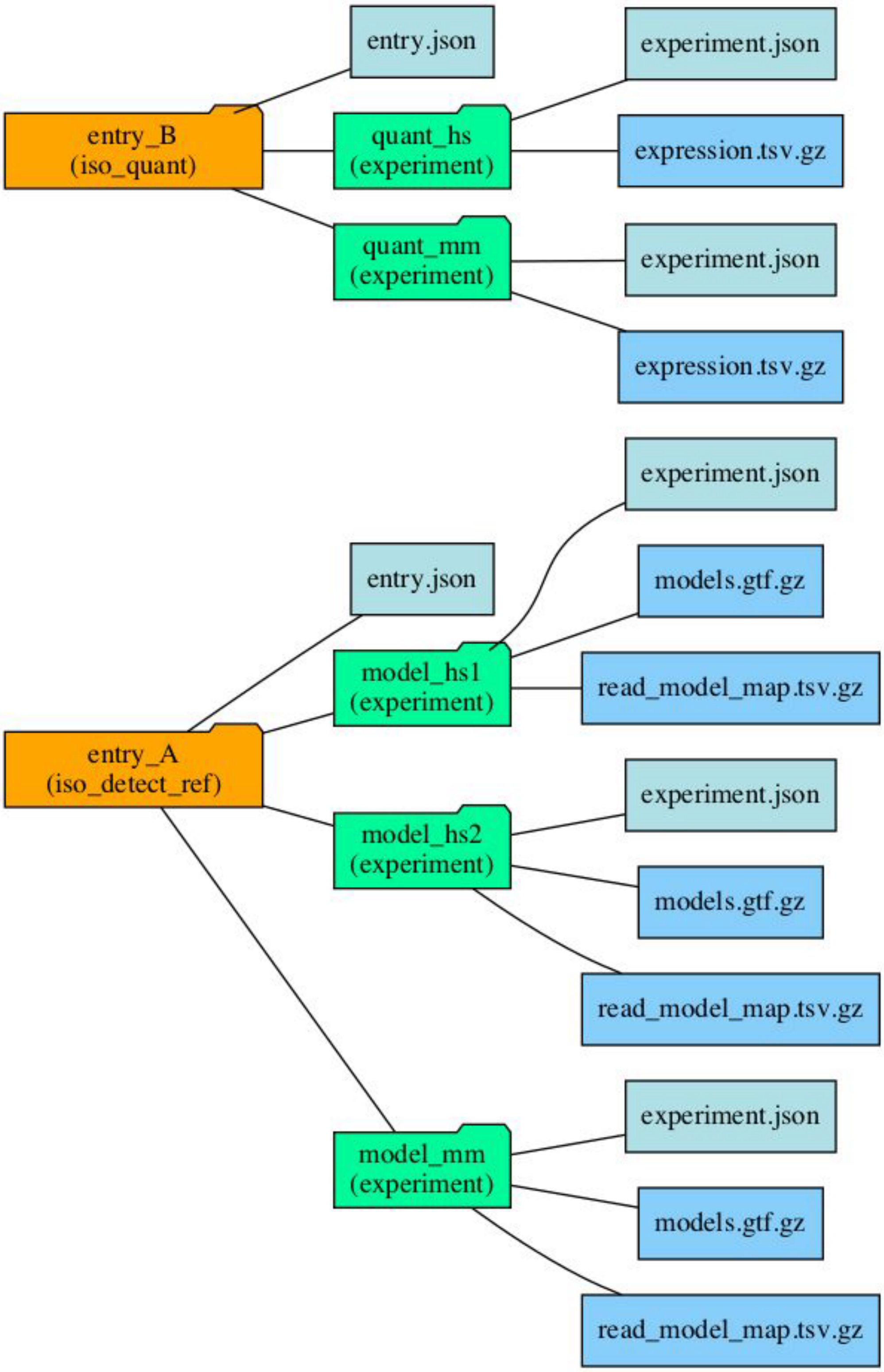


**Supplementary Fig. 7. Schematic of directory structure and files that would be included in each entry.**

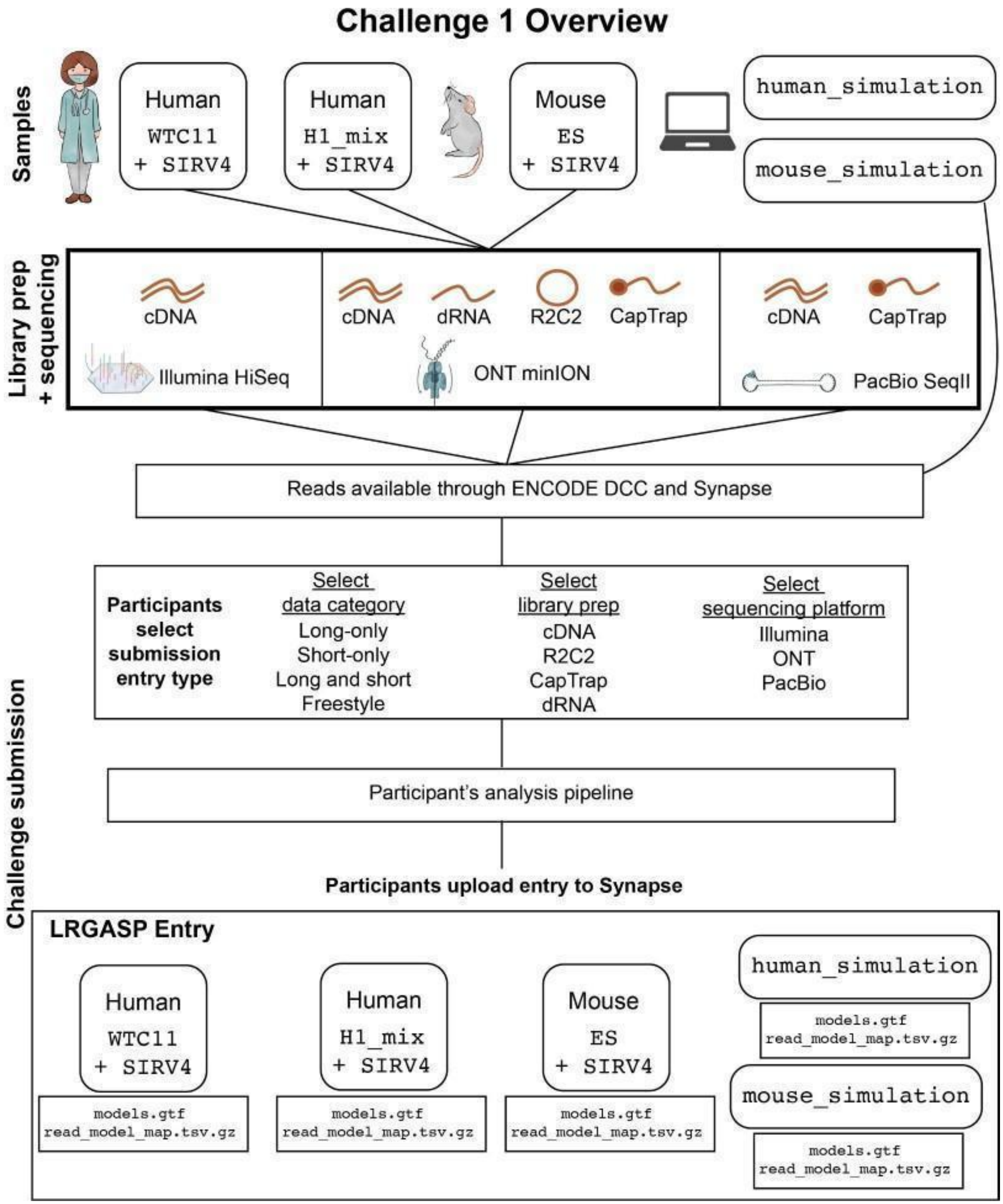


**Supplementary Fig. 8. Flow diagram of Challenge 1:** Transcript isoform detection with a high-quality genome. Samples, library prep methods, and sequencing platforms used in the challenge are indicated at the top. Participants select which data category, library prep, and sequencing platform to analyze, run their pipelines to generate transcript predictions, and submit an entry which includes predictions for all samples. The entries include a GTF file of the transcript models and a TSV file that assigns reads that supported each transcript model.

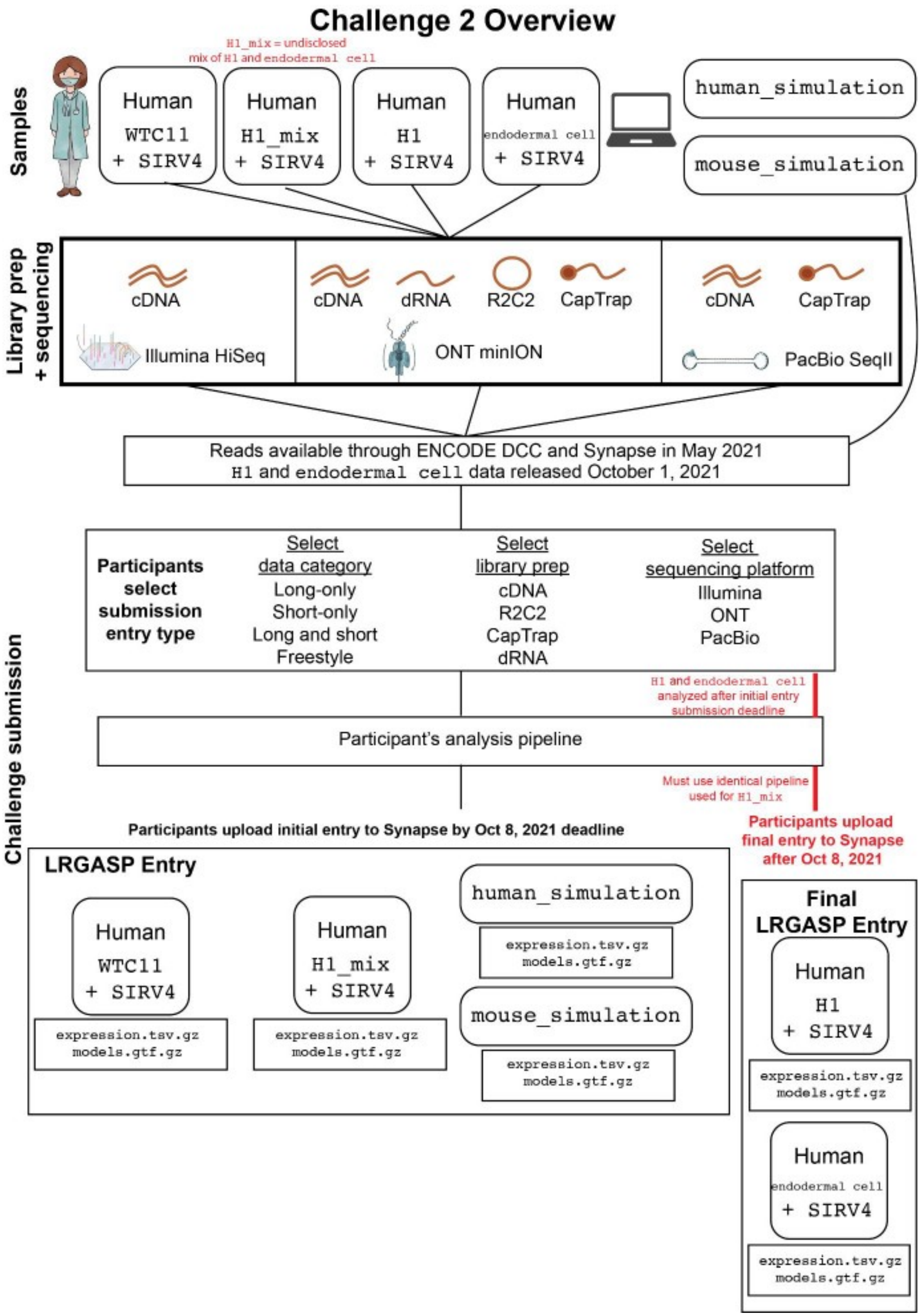
Challenge 2 Overview
Samples
H1_mix = undisclosed mix of H1 and endodermal cell
Human WTC11 + SIRV4
Human H1_mix + SIRV4
Human H1 + SIRV4
Human endodermal cell + SIRV4
human_simulation
mouse_simulation
Library prep + sequencing
cDNA
Illumina HiSeq
cDNA
dRNA
R2C2
CapTrap
ONT minION
cDNA
CapTrap
PacBio SeqII
Reads available through ENCODE DCC and Synapse in May 2021
H1 and endodermal cell data released October 1, 2021
Challenge submission
Participants select submission entry type
Select data category
Long-only
Short-only
Long and short
Freestyle
Select library prep
cDNA
R2C2
CapTrap
dRNA
Select sequencing platform
Illumina
ONT
PacBio
H1 and endodermal cell analyzed after initial entry submission deadline
Participant's analysis pipeline
Must use identical pipeline used for H1_mix
Participants upload initial entry to Synapse by Oct 8, 2021 deadline
Participants upload final entry to Synapse after Oct 8, 2021
LRGASP Entry
Human WTC11 + SIRV4
expression.tsv.gz
models.gtf.gz
Human H1_mix + SIRV4
expression.tsv.gz
models.gtf.gz
human_simulation
expression.tsv.gz
models.gtf.gz
mouse_simulation
expression.tsv.gz
models.gtf.gz
Final LRGASP Entry
Human H1 + SIRV4
expression.tsv.gz
models.gtf.gz
Human endodermal cell + SIRV4
expression.tsv.gz
models.gtf.gz

**Supplementary Fig. 9. Flow diagram of Challenge 2:** Transcript isoform quantification. Samples, library prep methods, and sequencing platforms used in the challenge are indicated at the top. Participants select which data category, library prep, and sequencing platform to analyze, run their pipelines to generate transcript predictions, and submit an entry which includes predictions for all samples. The entries include a GTF file of the transcript models that are quantified and a TSV file of the expression quantification. The H1 and endodermal cell samples were released after the initial submission deadline and participants were required to submit the quantification after the deadline Supplementary Fig. 9. Flow diagram of Challenge 2: Transcript isoform quantification. Samples, library prep methods, and sequencing platforms used in the challenge are indicated at the top. Participants select which data category, library prep, and sequencing platform to analyze, run their pipelines to generate transcript predictions, and submit an entry which includes predictions for all samples. The entries include a GTF file of the transcript models that are quantified and a TSV file of the expression quantification. The H1 and endodermal cell samples were released after the initial submission deadline and participants were required to submit the quantification after the deadline.

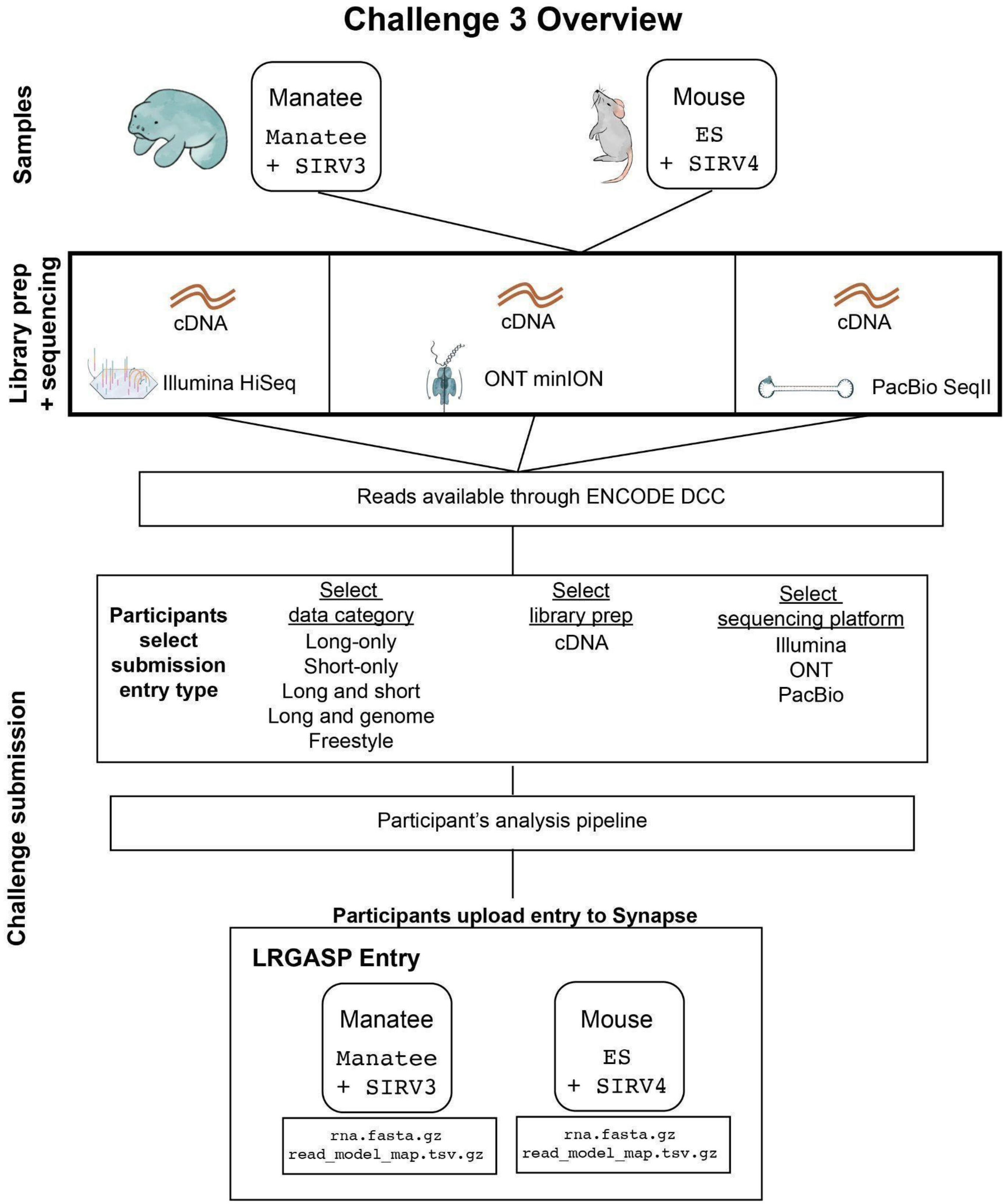


**Supplementary Fig. 10. Flow diagram of Challenge 3.** Samples, library prep methods, and sequencing platforms used in the challenge are indicated at the top. Participants select which data category and sequencing platform to analyze, run their pipelines to generate transcript predictions, and submit an entry which includes predictions for all samples. The entries include a FASTA file of the transcript models and a TSV file that assigns reads that supported each transcript model.

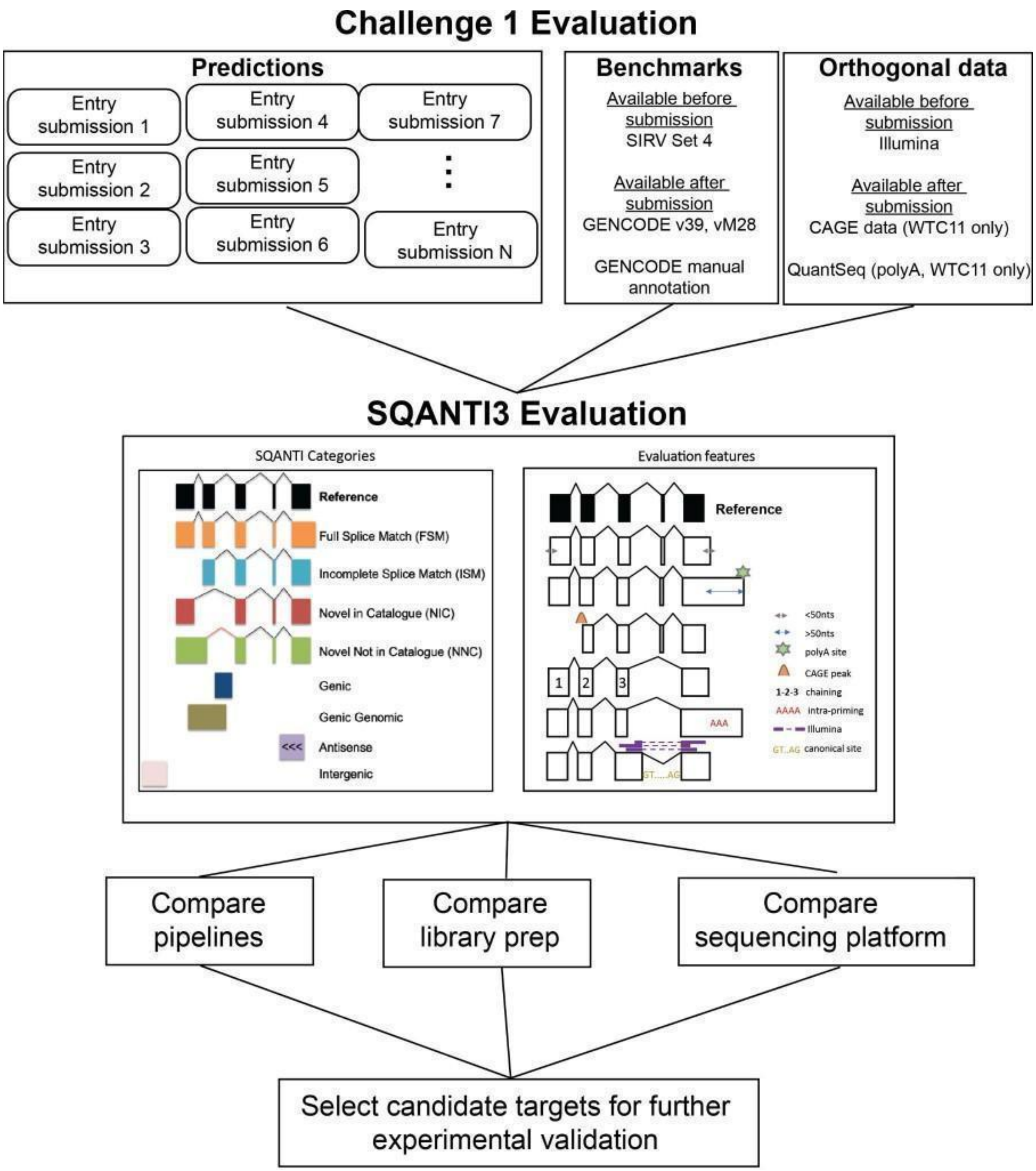


**Supplementary Fig. 11. Flow diagram of the evaluation for Challenge 1.** Benchmarks and additional orthogonal data that was used for the evaluation are indicated. For example, CAGE and QuantSeq data from WTC11 cells were generated and made available only after participant submissions; therefore, they represent "hidden" data. These was used to define 5' transcript starts and 3' ends.

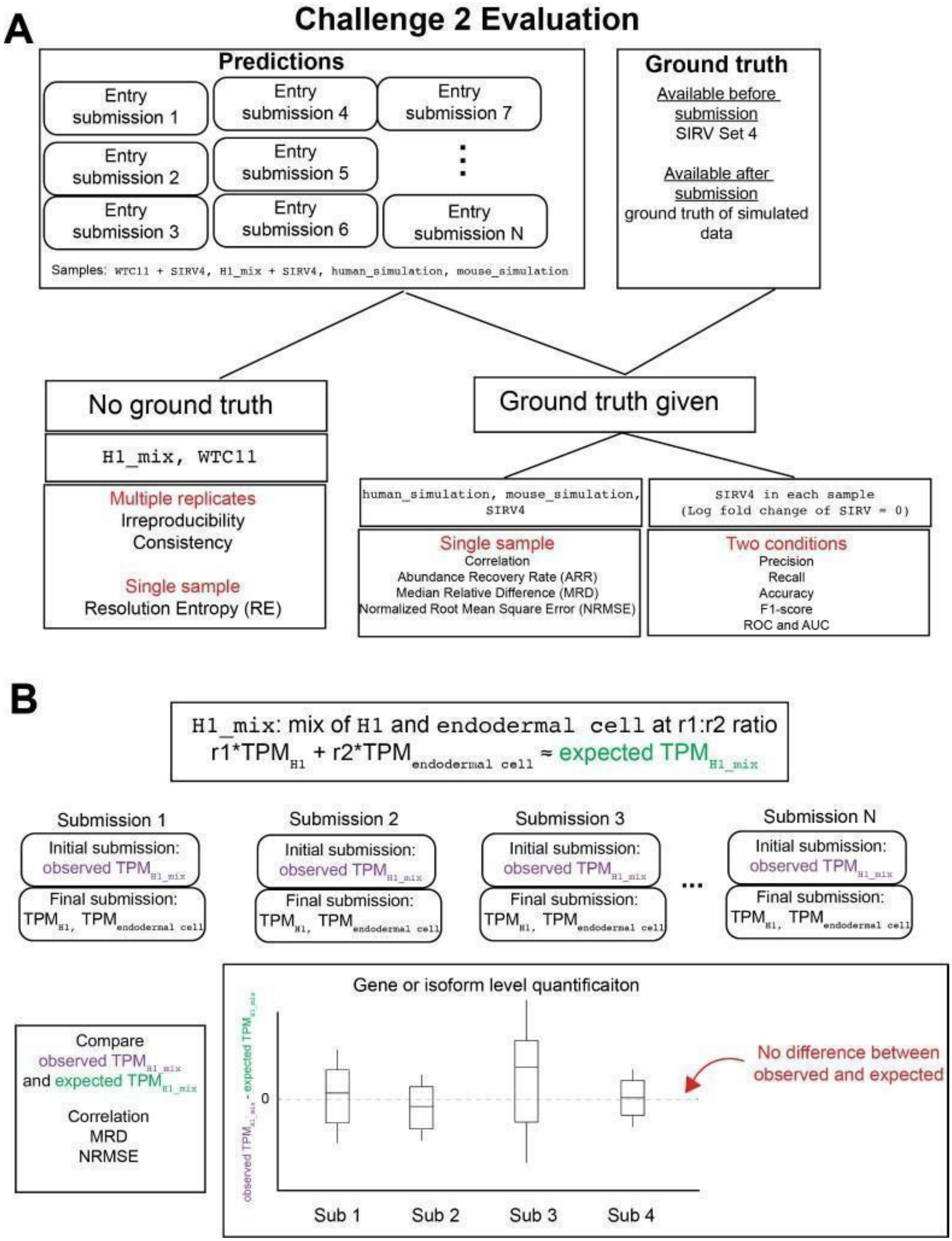


**Supplementary Fig. 12. Flow diagram of the evaluation for Challenge 2.** (A) Evaluation of Challenge 2 can be separated into metrics when a ground truth is known or a ground truth is unknown. (B) Example analyses to evaluate transcript expression using the cell mixing experiment. A sample, H1_mix, was initially provided for quantification which was a mix of H1 cells and endodermal cells at an undisclosed ratio. After the initial submission, the individual H1 and endodermal cell samples were released and participants submitted quantifications for each.

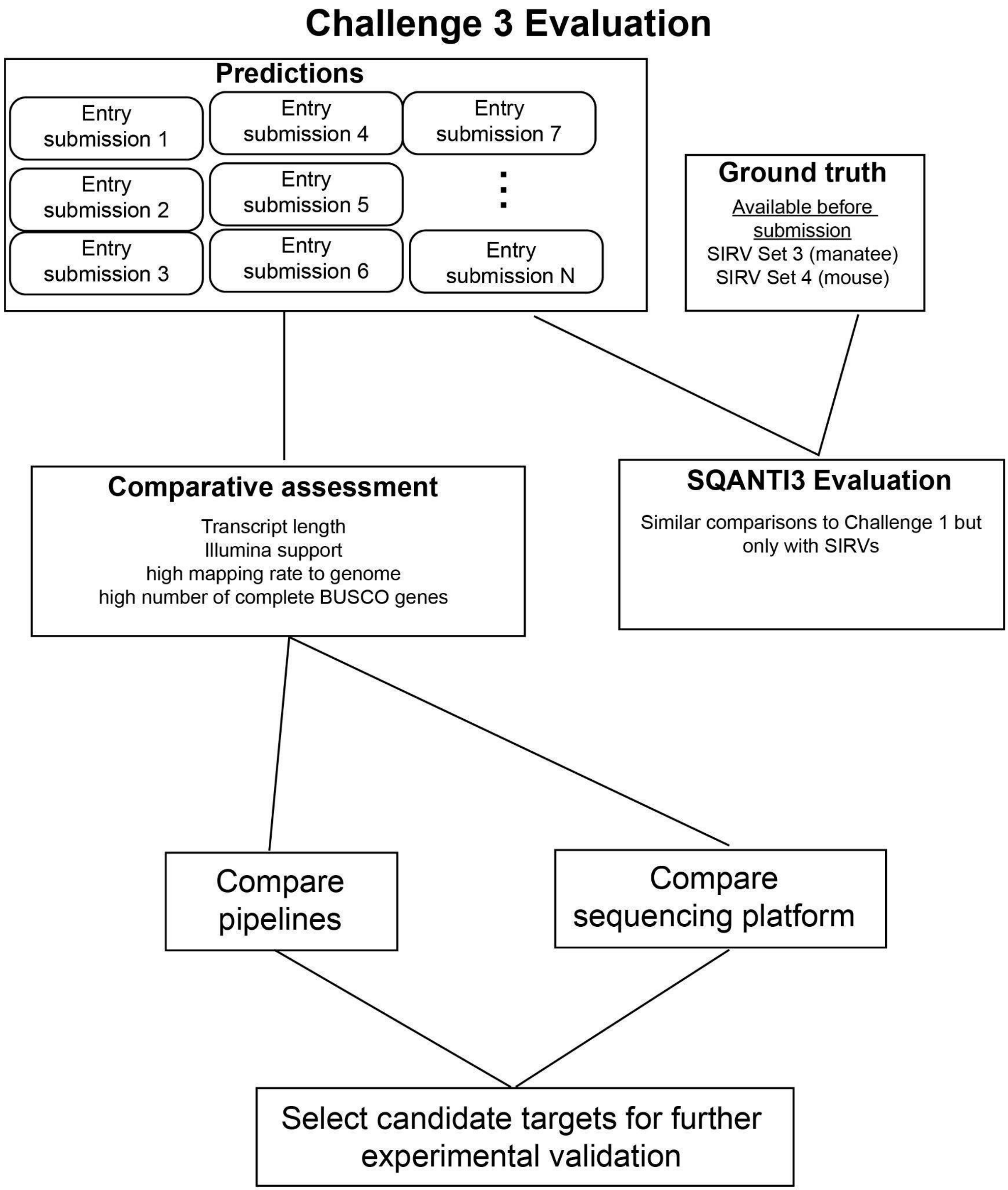


**Supplementary Fig. 13**. **Flow diagram of the evaluation for Challenge 3.** Only SIRVs are available for ground truth information. The evaluation was based on a comparative assessment of the predictions followed by targeting specific candidates for further validation.

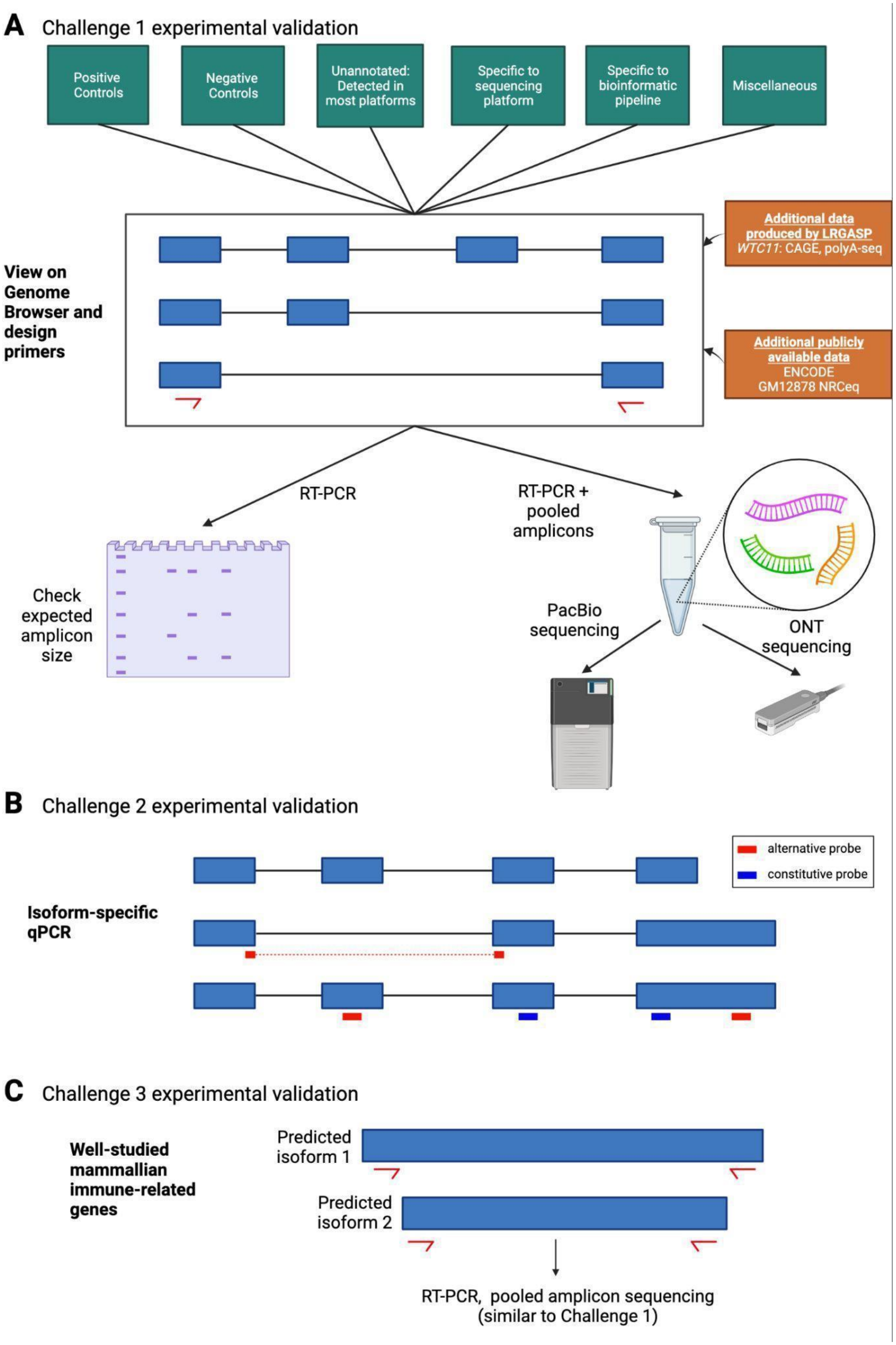
A
Challenge 1 experimental validation
Positive Controls
Negative Controls
Unannotated: Detected in most platforms
Specific to sequencing platform
Specific to bioinformatic pipeline
Miscellaneous
View on Genome Browser and design primers
Additional data produced by LRGASP
WTC11: CAGE, polyA-seq
Additional publicly available data
ENCODE
GM12878 NRCeq
RT-PCR
RT-PCR + pooled amplicons
Check expected amplicon size
PacBio sequencing
ONT sequencing
B
Challenge 2 experimental validation
alternative probe
constitutive probe
Isoform-specific qPCR
C
Challenge 3 experimental validation
Well-studied mammallian immune-related genes
Predicted isoform 1
Predicted isoform 2
RT-PCR, pooled amplicon sequencing
(similar to Challenge 1)

**Supplementary Fig. 14. Experimental validation approaches for the LRGASP challenges.** (A) Multiple categories of types of transcript were selected for validation (shown in green boxes). These loci will be viewed in the UCSC Genome Browser along with additional datasets to aid in the manual design of primers. Amplicons will be analyzed by fragment size and pooled to perform long-read sequencing with PacBio and ONT (B) A select number of genes were selected for transcript isoform-specific qPCR. A combination of probes detecting constitutive and alternative regions will be used. (C) RT-PCR validation will be performed similar to Challenge 1, except transcript were selected from well-studied mammalian immune-related genes.

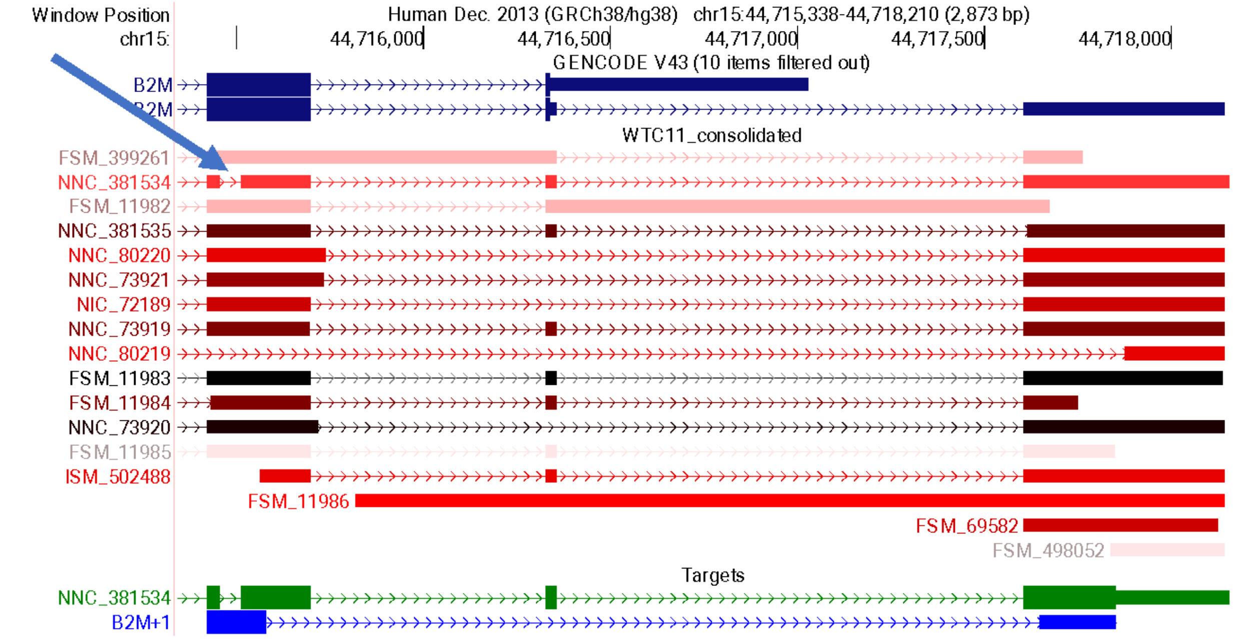


**Supplementary Fig. 15. An example of a unique intron in transcript NNC_381534 to validation.** The green and blue region vertical highlights indicate the manually selected primer pair regions. The 'Targets' track, produced by Primers-Juju, recapitulates the region as blue item B2M+1, and transcript with the maximal possible amplicon drawn in thick boxes.

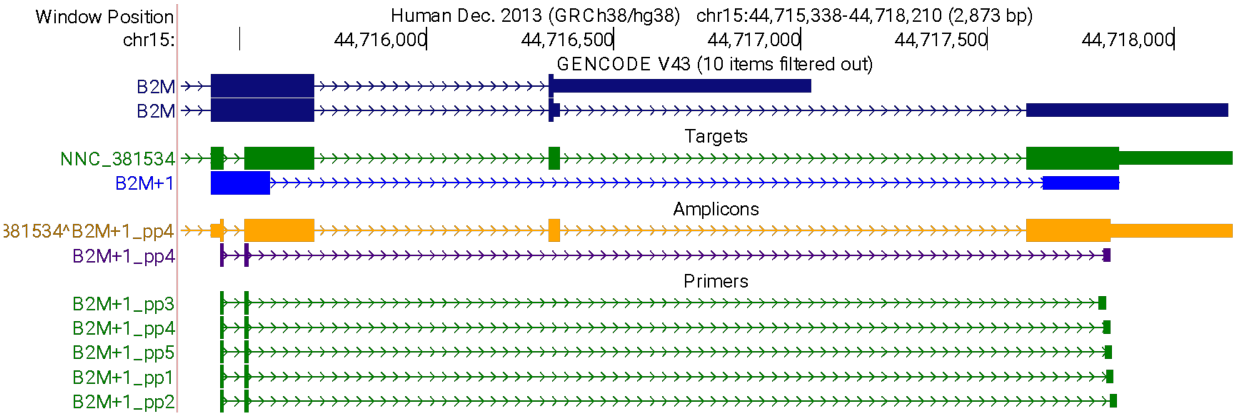


**Supplementary Fig. 16. The Primers-Juju track hub with the addition of the primer pairs design.** This adds Primer3 results (Primers track) and the most stable primer along with the amplicon sequence for the target transcript (Amplicons track).
