## Supplementary Data Report for "Systematic assessment of long-read RNA-seq methods for transcript identification and quantification"

### *LRGASP Challenge 1 comparison report*

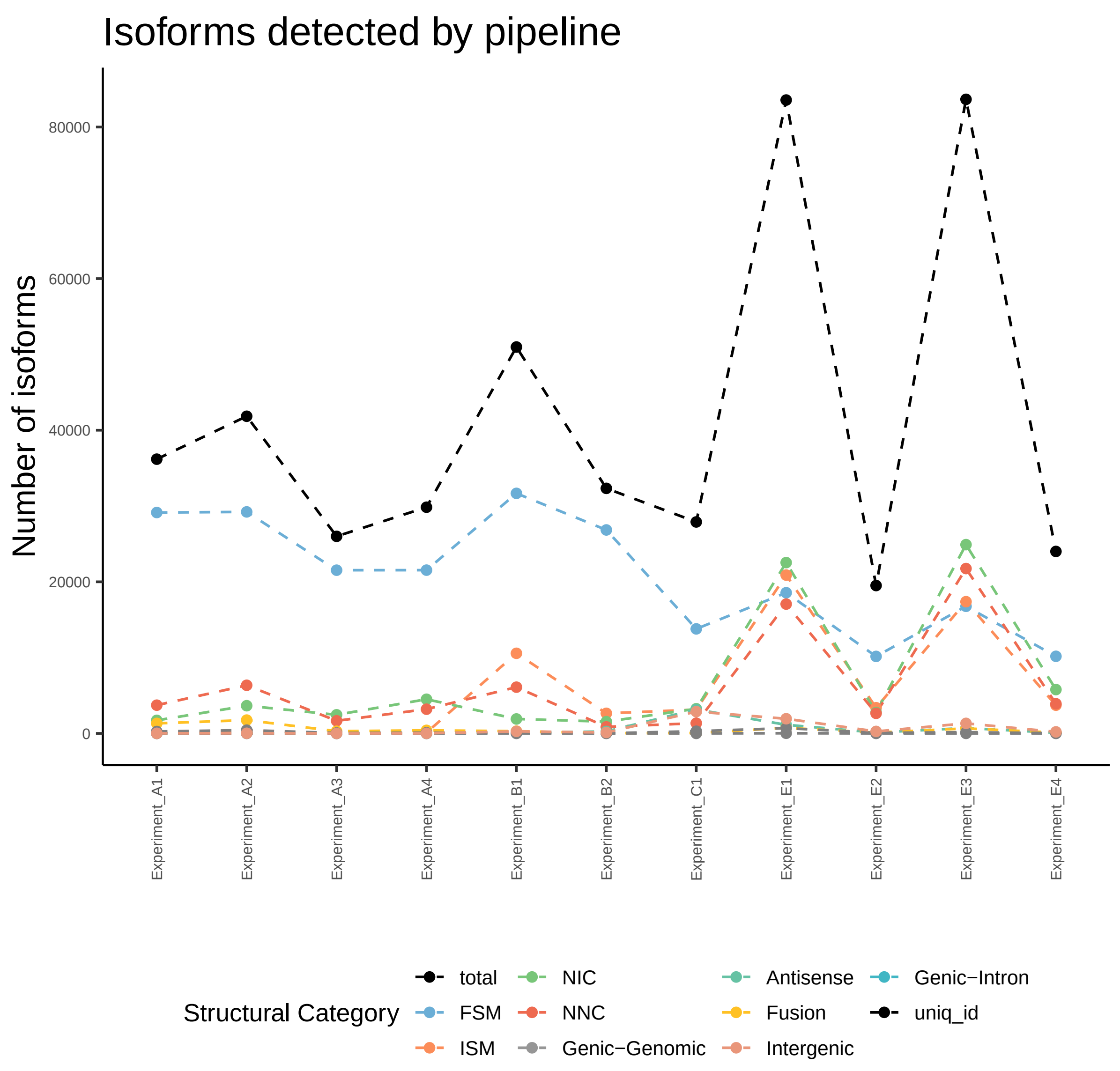
Isoforms detected by pipeline
Number of isoforms
80000
60000
40000
20000
0
Experiment_A1
Experiment_A2
Experiment_A3
Experiment_A4
Experiment_B1
Experiment_B2
Experiment_C1
Experiment_E1
Experiment_E2
Experiment_E3
Experiment_E4
Structural Category
total
FSM
ISM
NIC
NNC
Genic−Genomic
Antisense
Fusion
Intergenic
Genic−Intron
uniq_id

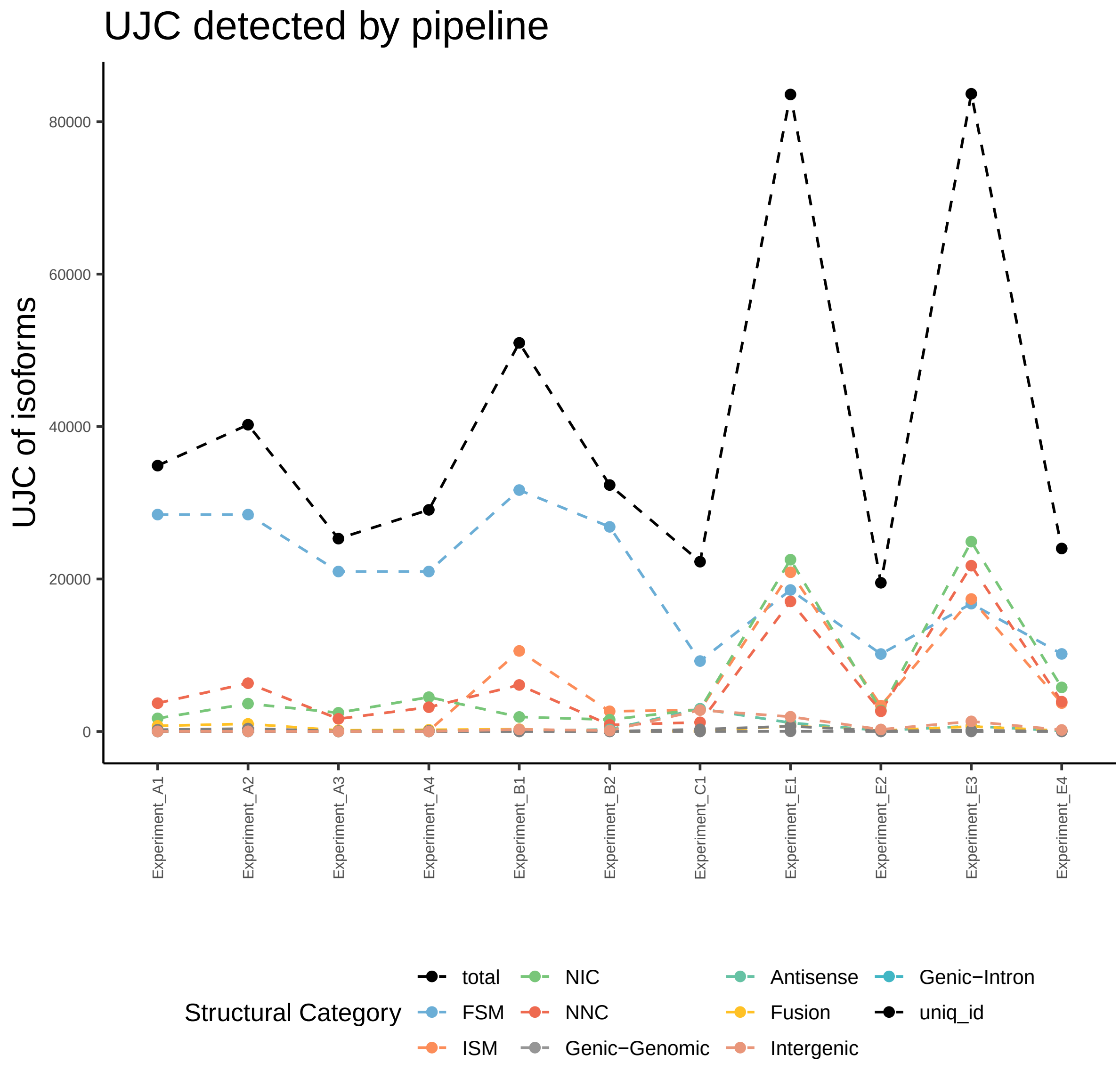

UJC detected by pipeline
UJC of isoforms
80000
60000
40000
20000
0
Experiment_A1
Experiment_A2
Experiment_A3
Experiment_A4
Experiment_B1
Experiment_B2
Experiment_C1
Experiment_E1
Experiment_E2
Experiment_E3
Experiment_E4
Structural Category
total
FSM
ISM
NIC
NNC
Genic−Genomic
Antisense
Fusion
Intergenic
Genic−Intron
uniq_id

*Gene Characterization*

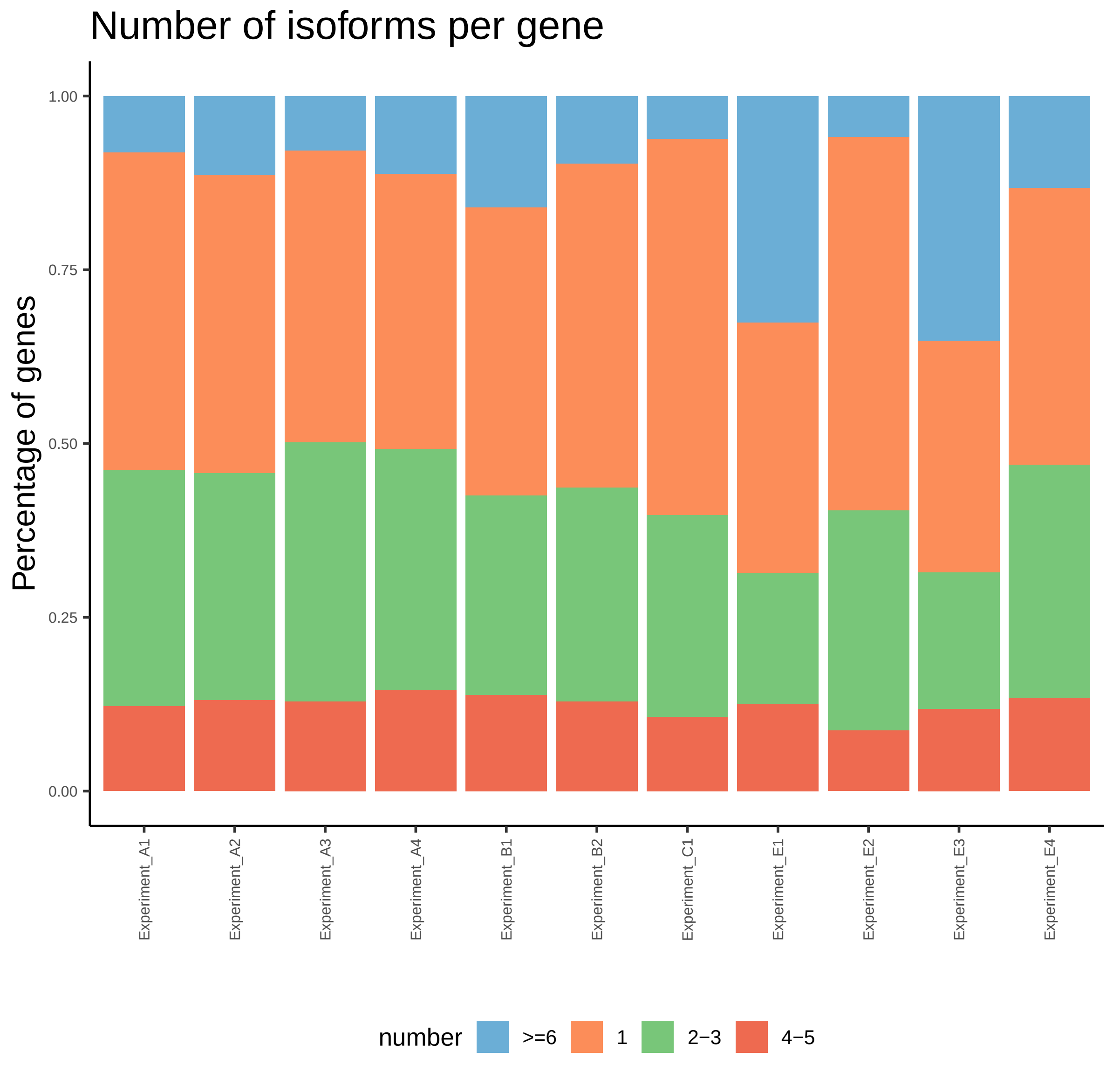

Number of isoforms per gene
Percentage of genes
1.00
0.75
0.50
0.25
0.00
Experiment_A1
Experiment_A2
Experiment_A3
Experiment_A4
Experiment_B1
Experiment_B2
Experiment_C1
Experiment_E1
Experiment_E2
Experiment_E3
Experiment_E4
number
>=6
1
2−3
4−5

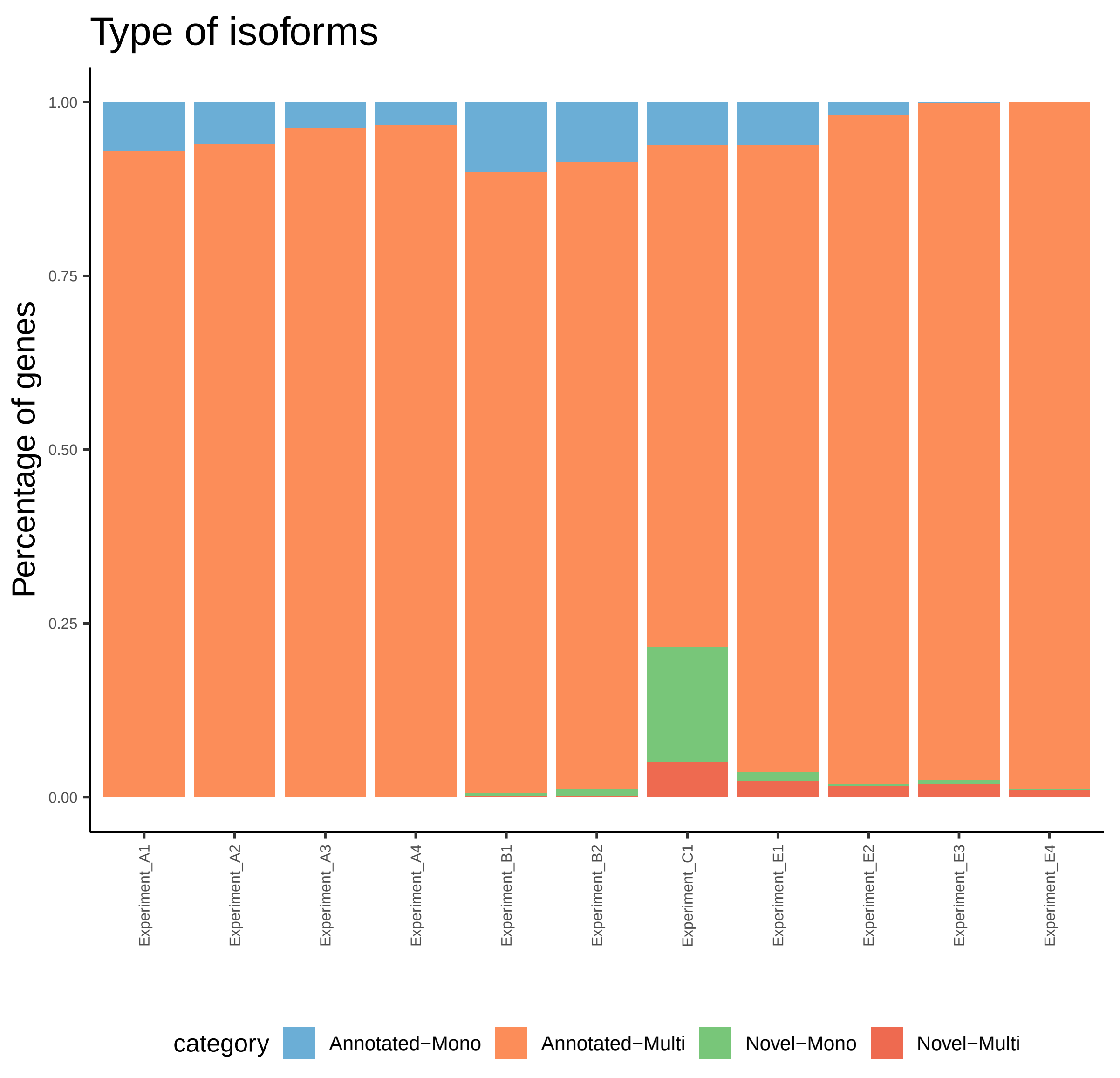

Type of isoforms
Percentage of genes
1.00
0.75
0.50
0.25
0.00
Experiment_A1
Experiment_A2
Experiment_A3
Experiment_A4
Experiment_B1
Experiment_B2
Experiment_C1
Experiment_E1
Experiment_E2
Experiment_E3
Experiment_E4
category
Annotated−Mono
Annotated−Multi
Novel−Mono
Novel−Multi

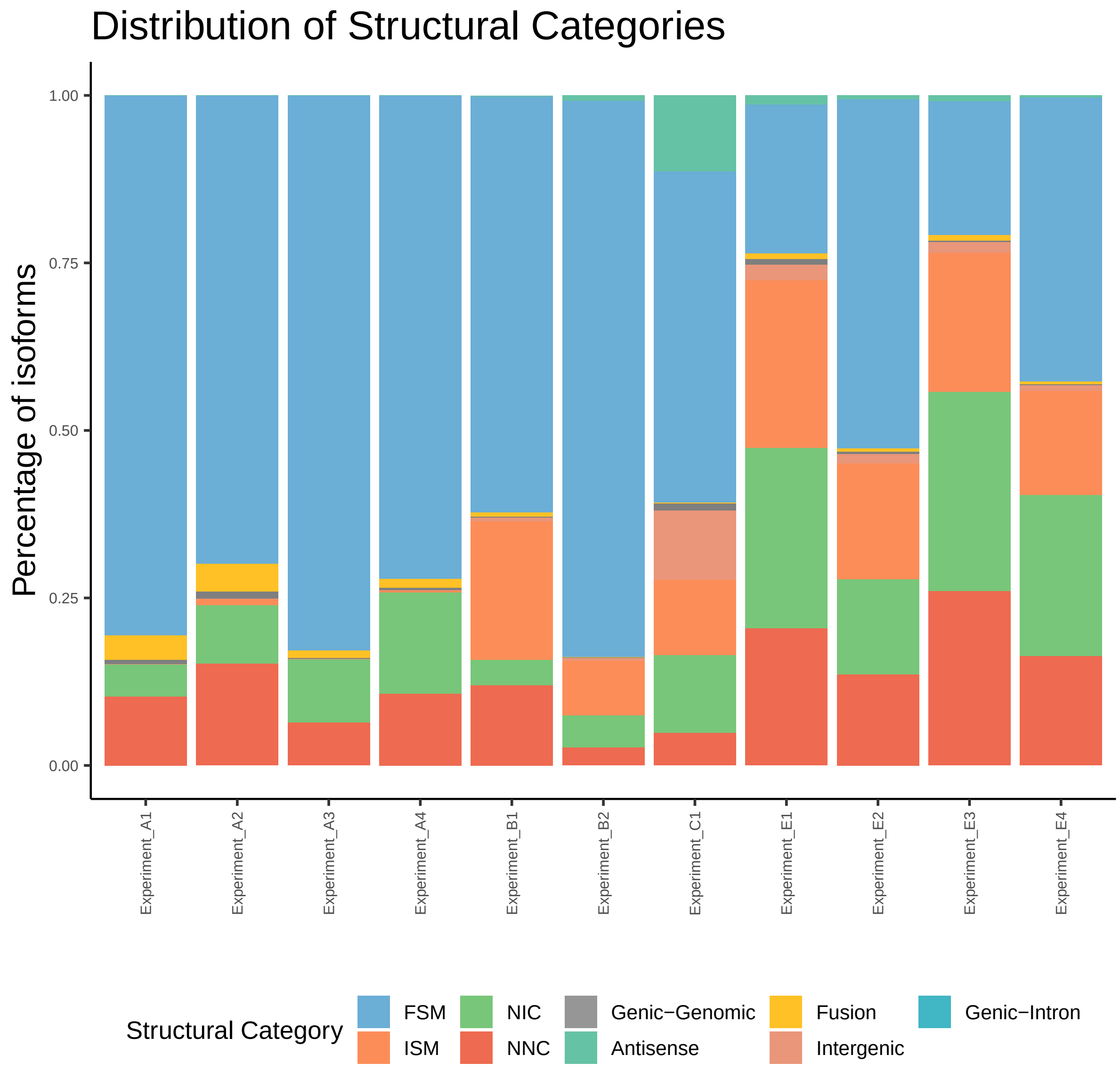

Distribution of Structural Categories
Percentage of isoforms
1.00
0.75
0.50
0.25
0.00
Experiment_A1
Experiment_A2
Experiment_A3
Experiment_A4
Experiment_B1
Experiment_B2
Experiment_C1
Experiment_E1
Experiment_E2
Experiment_E3
Experiment_E4
Structural Category
FSM
ISM
NIC
NNC
Genic−Genomic
Antisense
Fusion
Intergenic
Genic−Intron

*Distance to annotated TSS and TTS*

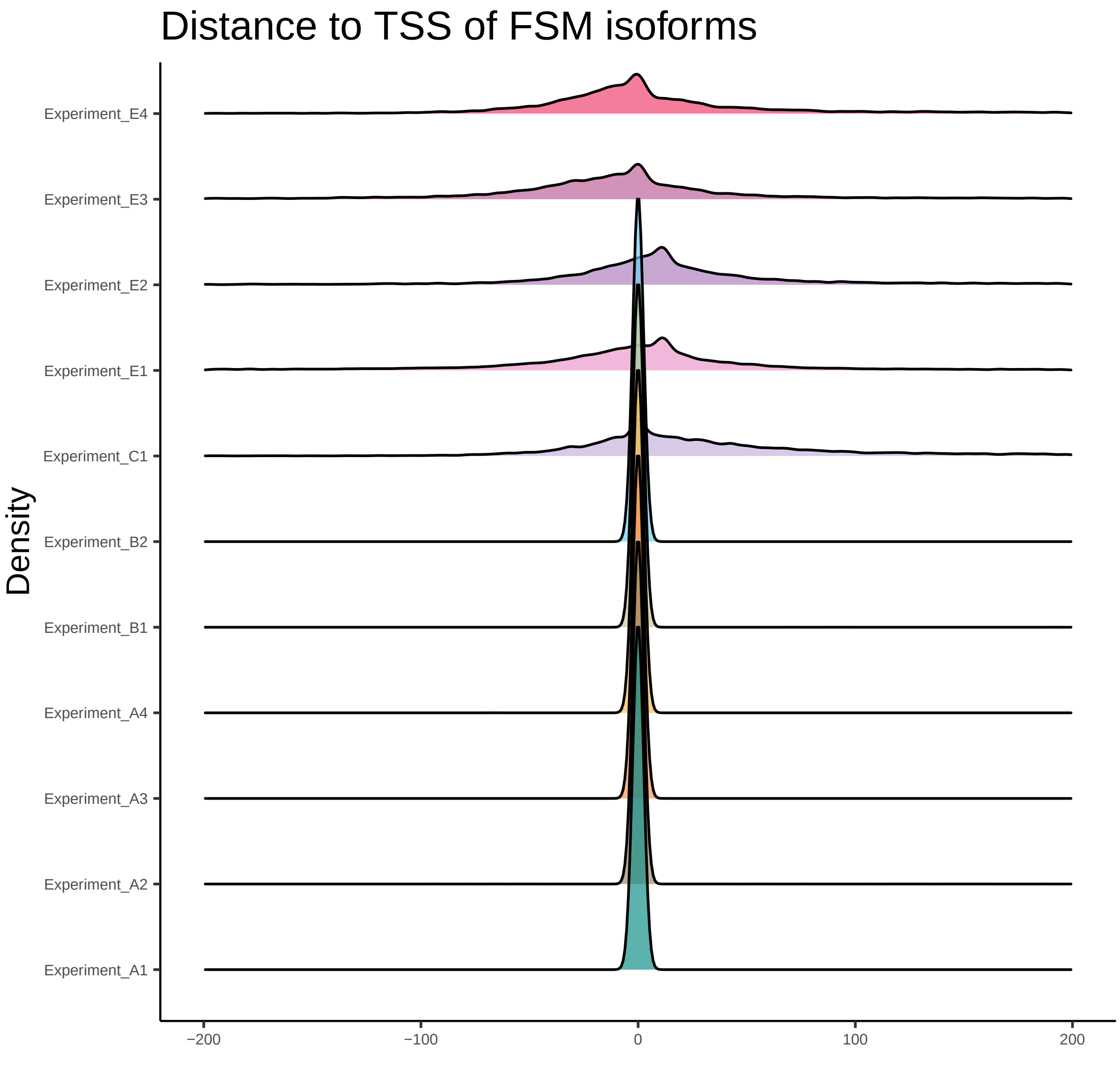

Distance to TSS of FSM isoforms
Density
Experiment_E4
Experiment_E3
Experiment_E2
Experiment_E1
Experiment_C1
Experiment_B2
Experiment_B1
Experiment_A4
Experiment_A3
Experiment_A2
Experiment_A1
−200
−100
0
100
200

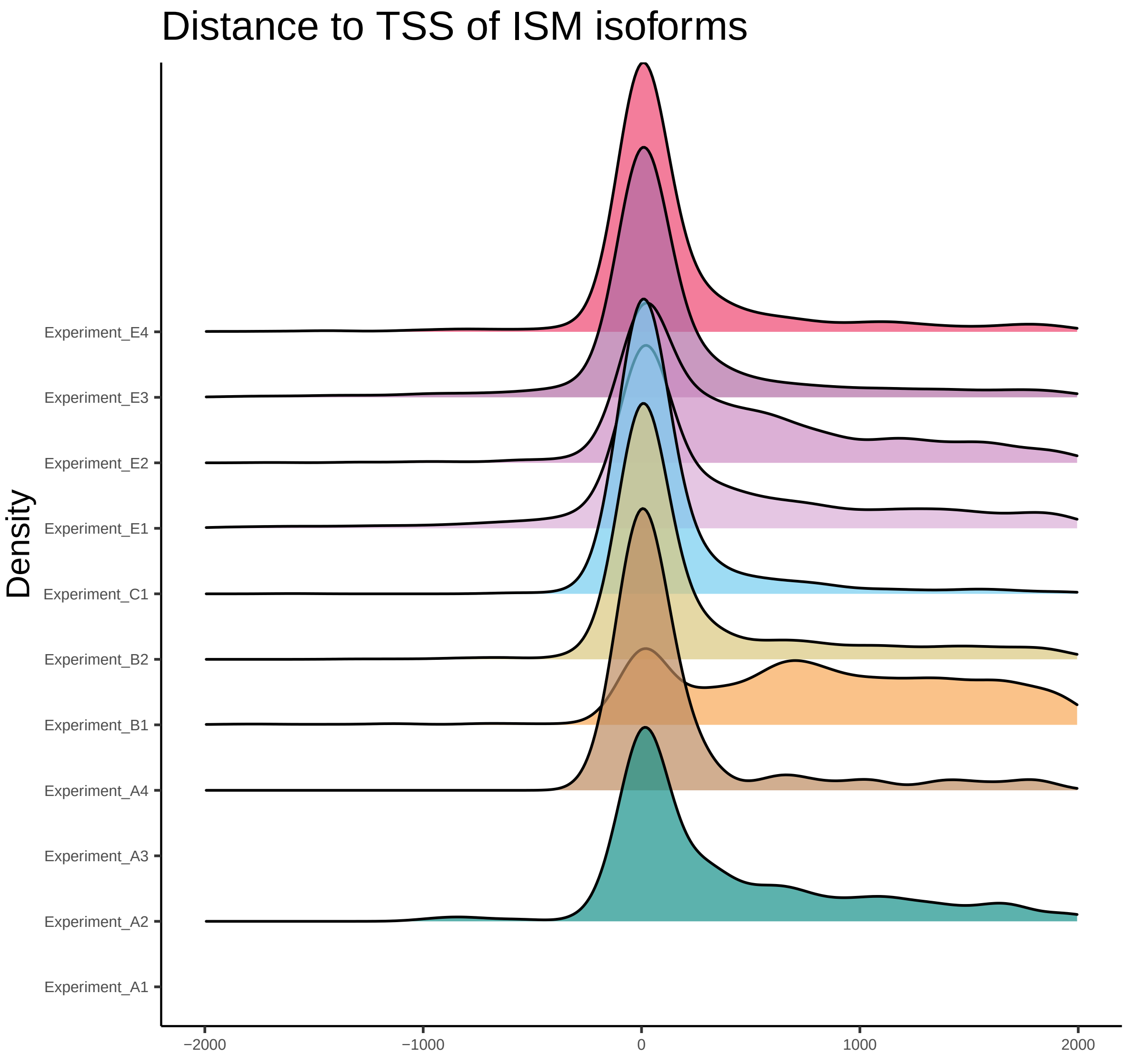

Distance to TSS of ISM isoforms
Density
Experiment_E4
Experiment_E3
Experiment_E2
Experiment_E1
Experiment_C1
Experiment_B2
Experiment_B1
Experiment_A4
Experiment_A3
Experiment_A2
Experiment_A1
−2000
−1000
0
1000
2000

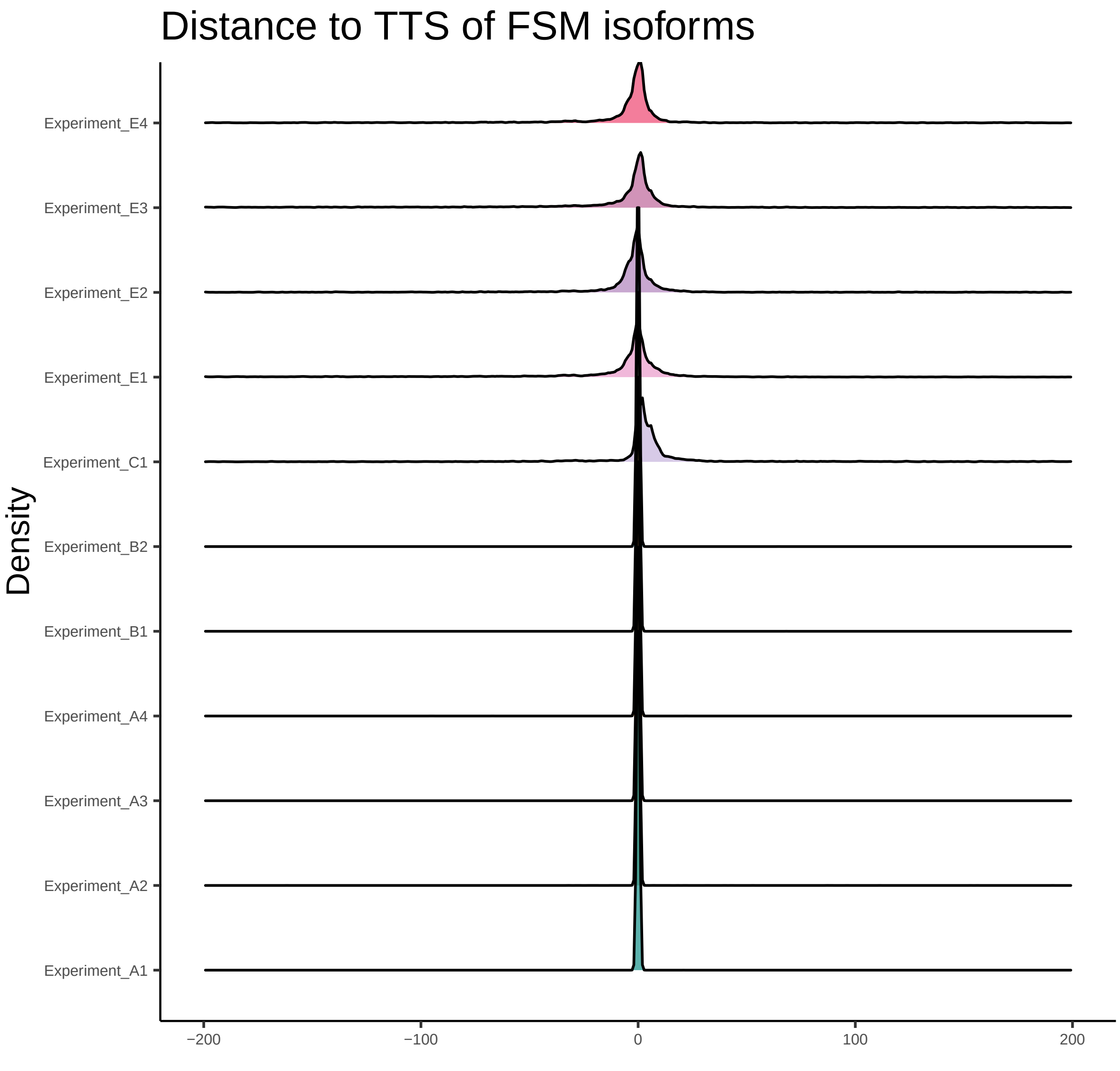
Distance to TTS of FSM isoforms
Density
Experiment_E4
Experiment_E3
Experiment_E2
Experiment_E1
Experiment_C1
Experiment_B2
Experiment_B1
Experiment_A4
Experiment_A3
Experiment_A2
Experiment_A1
−200
−100
0
100
200

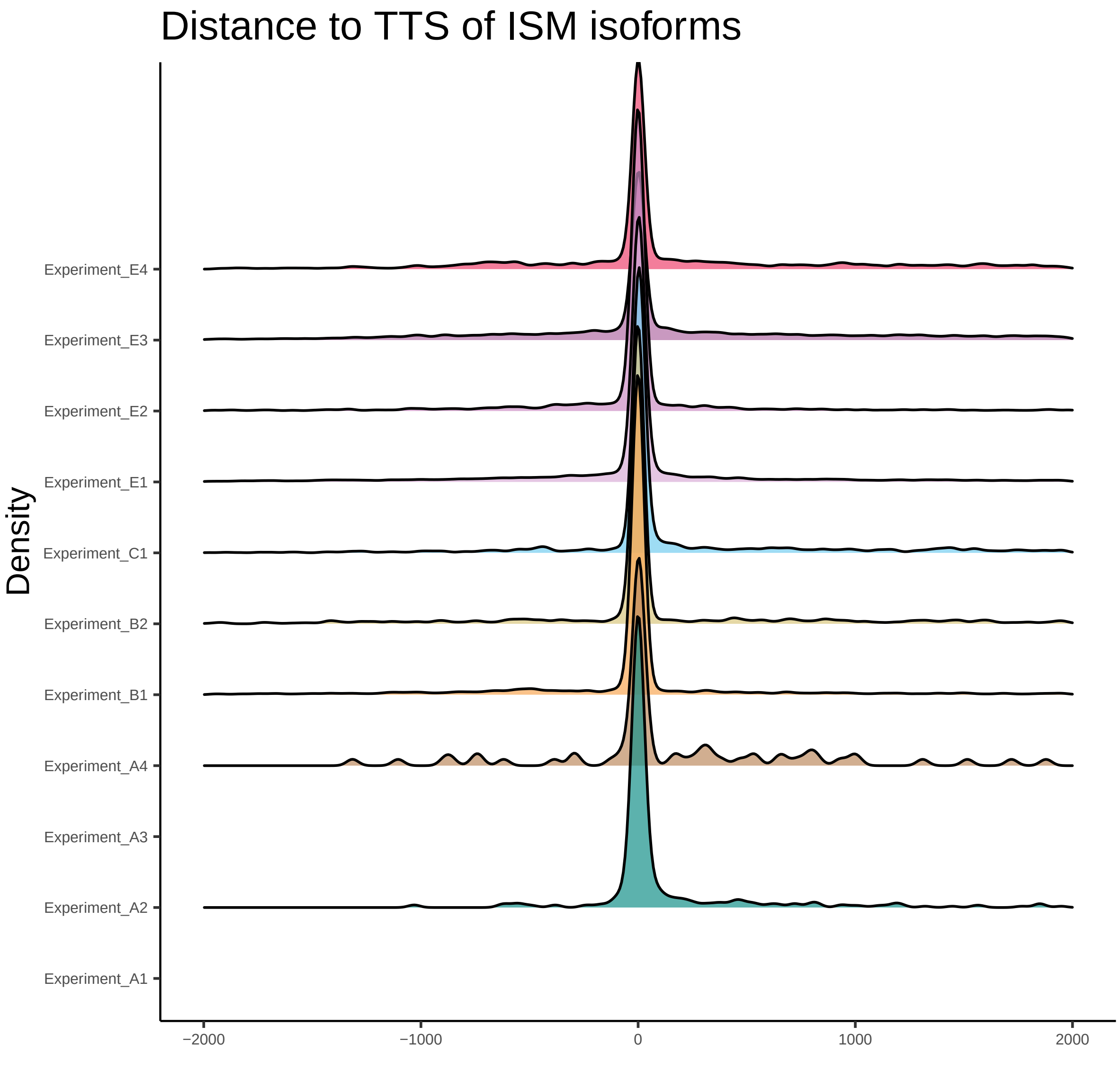

Distance to TTS of ISM isoforms
Density
Experiment_E4
Experiment_E3
Experiment_E2
Experiment_E1
Experiment_C1
Experiment_B2
Experiment_B1
Experiment_A4
Experiment_A3
Experiment_A2
Experiment_A1
−2000
−1000
0
1000
2000

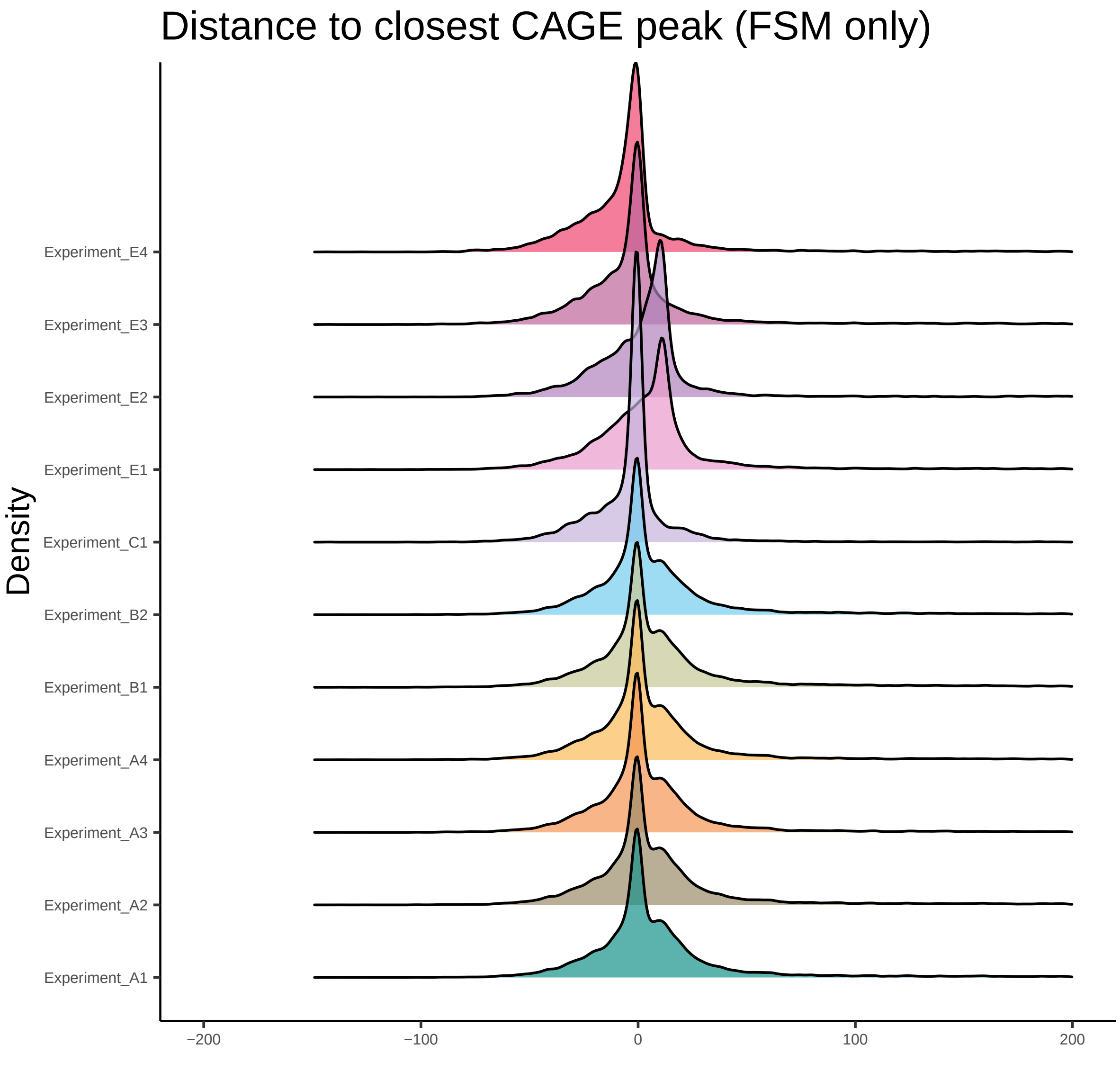
Distance to closest CAGE peak (FSM only)
Density
Experiment_E4
Experiment_E3
Experiment_E2
Experiment_E1
Experiment_C1
Experiment_B2
Experiment_B1
Experiment_A4
Experiment_A3
Experiment_A2
Experiment_A1
−200
−100
0
100
200

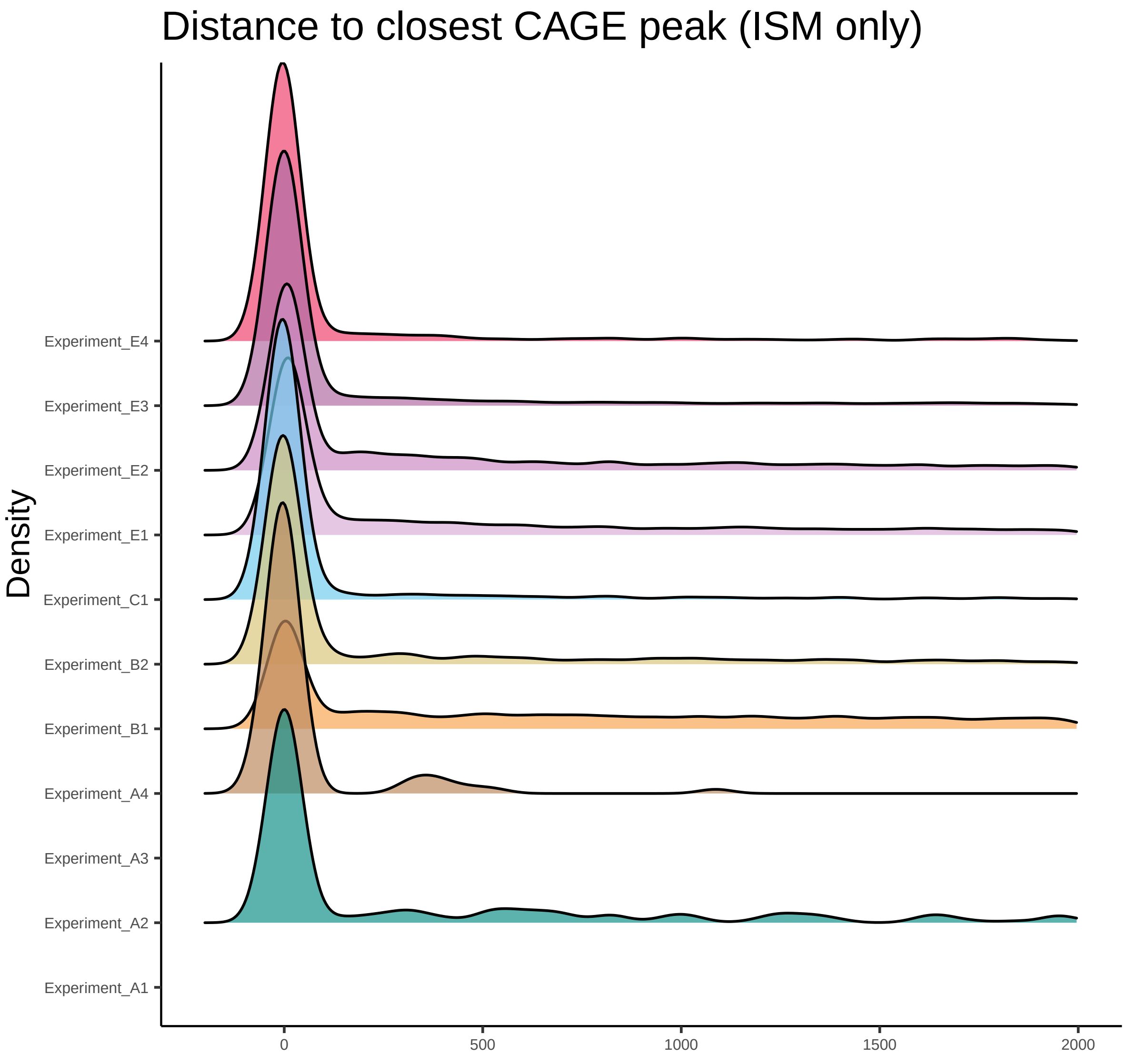
Distance to closest CAGE peak (ISM only)
Density
Experiment_E4
Experiment_E3
Experiment_E2
Experiment_E1
Experiment_C1
Experiment_B2
Experiment_B1
Experiment_A4
Experiment_A3
Experiment_A2
Experiment_A1
0
500
1000
1500
2000

*Presence/absence analysis of UJC across pipelines*

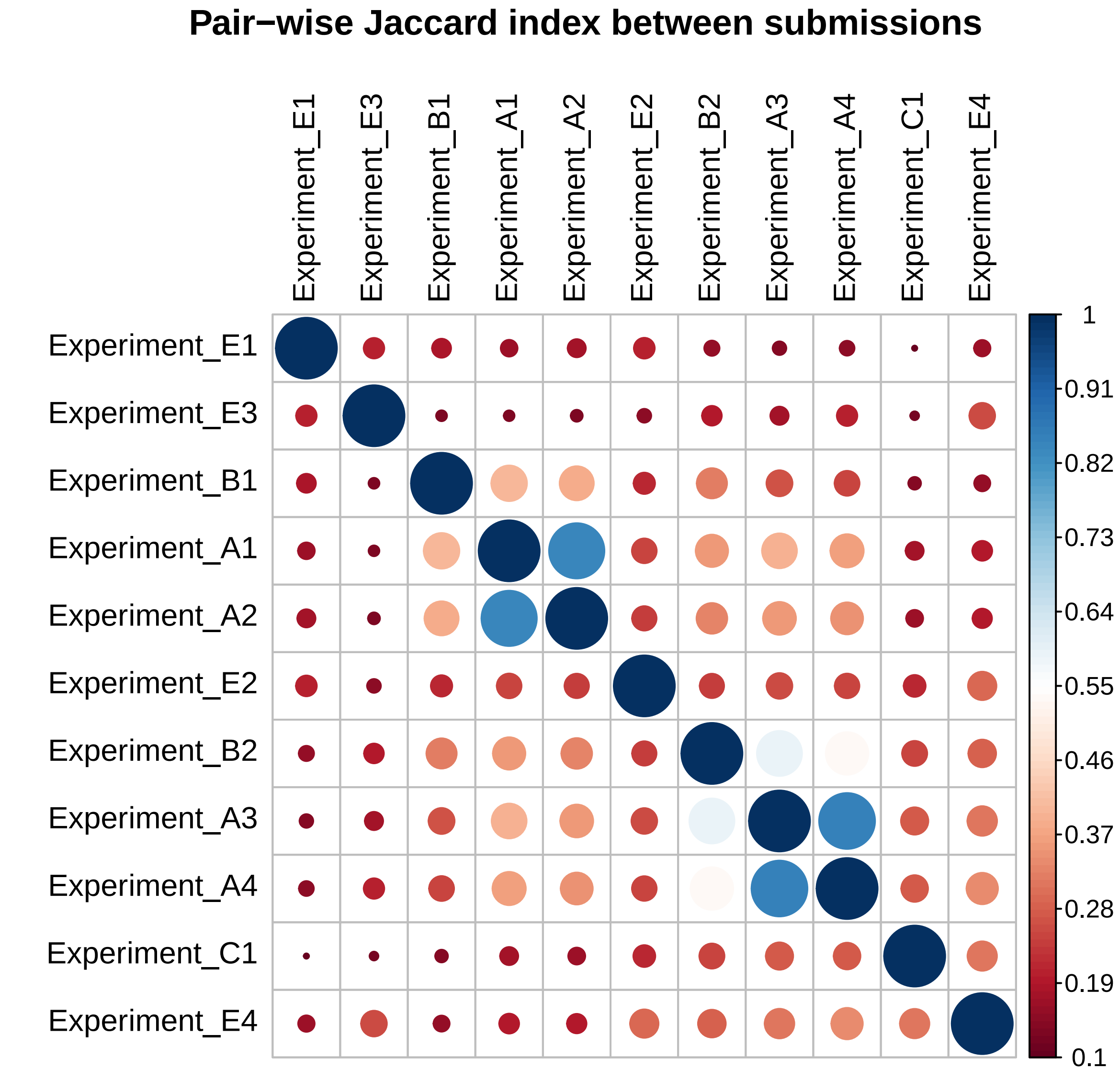
Pair−wise Jaccard index between submissions
Experiment_E1
Experiment_E3
Experiment_B1
Experiment_A1
Experiment_A2
Experiment_E2
Experiment_B2
Experiment_A3
Experiment_A4
Experiment_C1
Experiment_E4
1
0.91
0.82
0.73
0.64
0.55
0.46
0.37
0.28
0.19
0.1

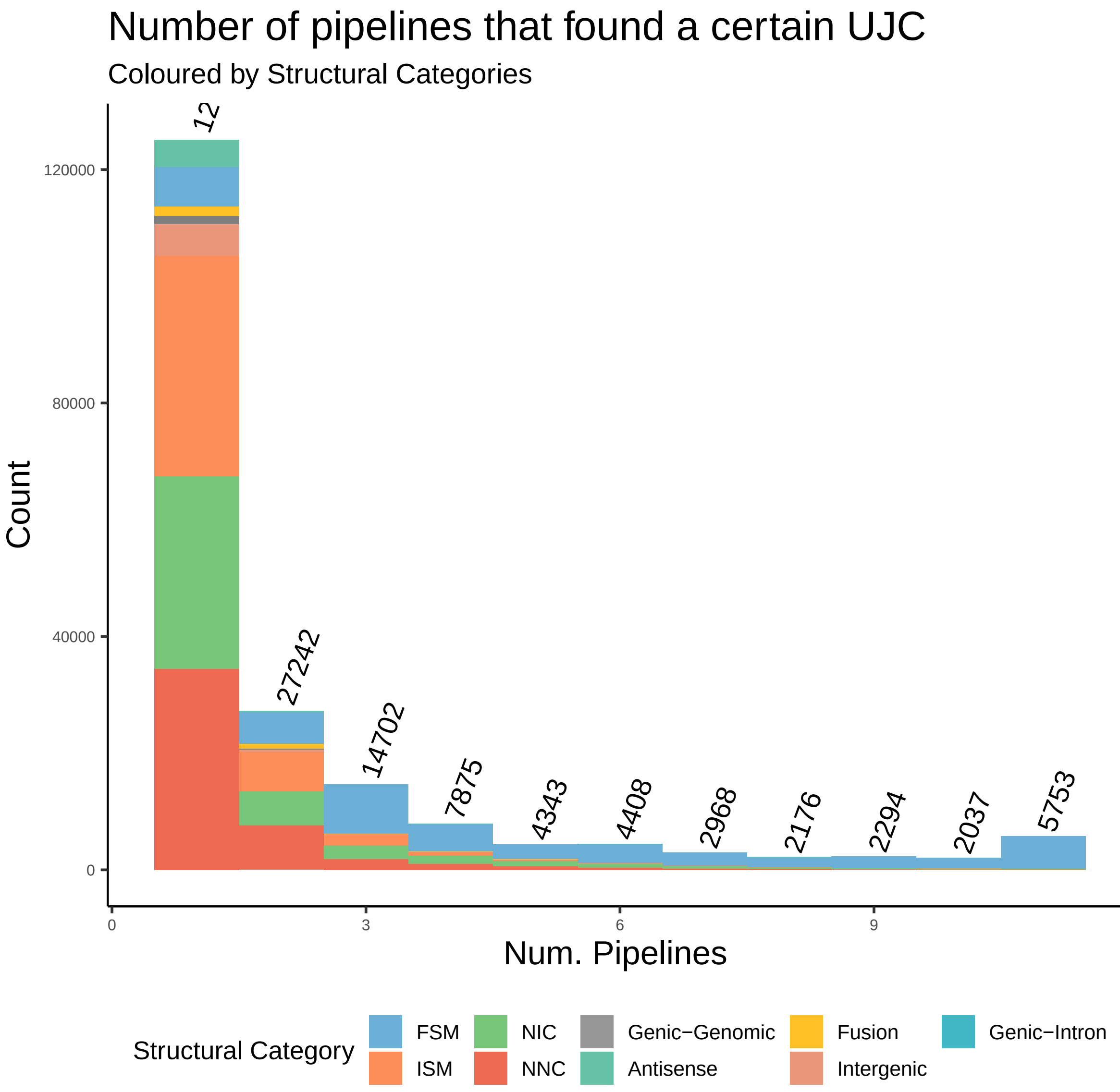

Number of pipelines that found a certain UJC
Coloured by Structural Categories
Count
Num. Pipelines
0
40000
80000
120000
0
3
6
9
12
12
27242
14702
7875
4343
4408
2968
2176
2294
2037
5753
Structural Category
FSM
ISM
NIC
NNC
Genic−Genomic
Antisense
Fusion
Intergenic
Genic−Intron

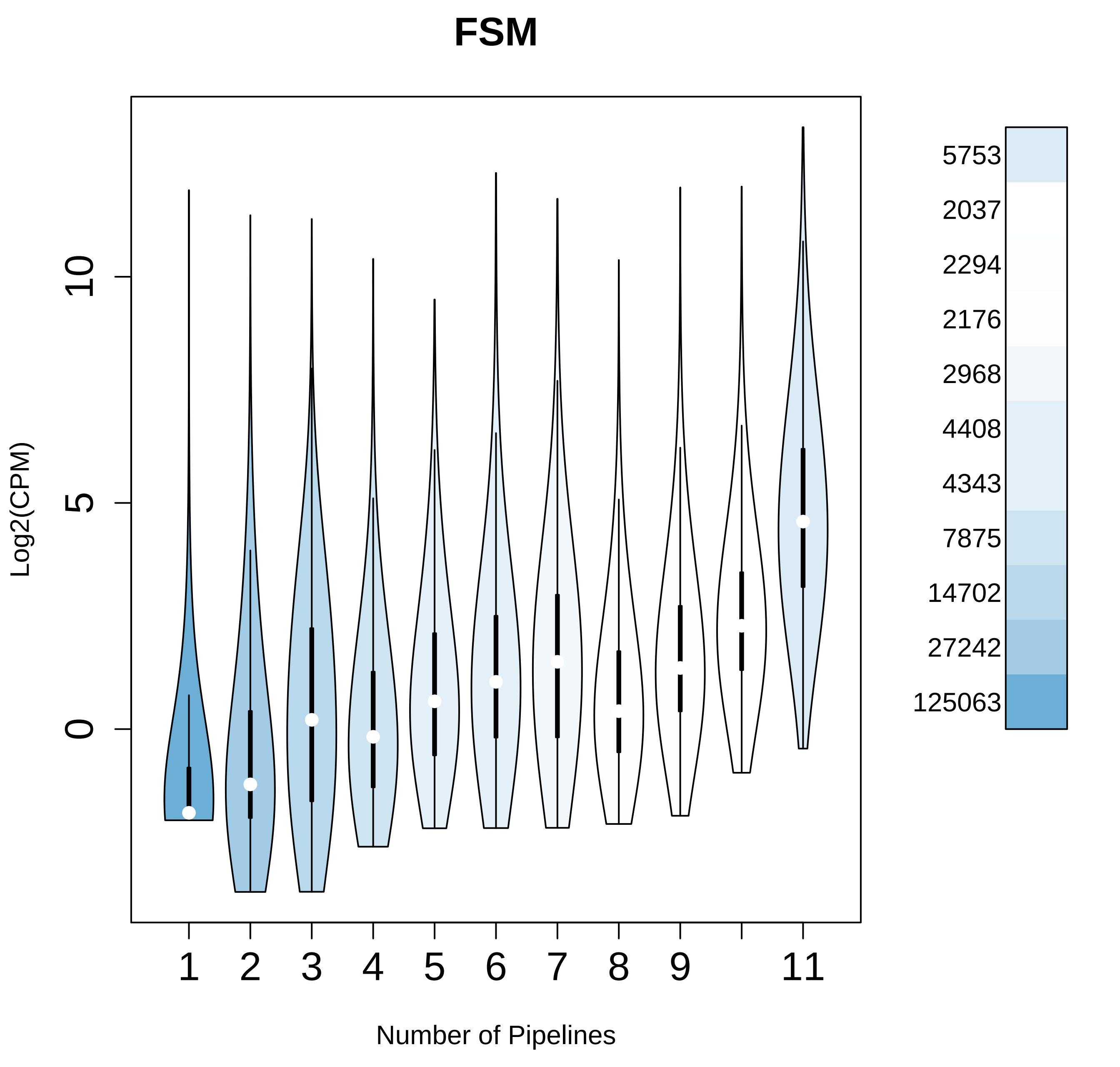

FSM
Log2(CPM)
0
5
10
1
2
3
4
5
6
7
8
9
11
Number of Pipelines
5753
2037
2294
2176
2968
4408
4343
7875
14702
27242
125063

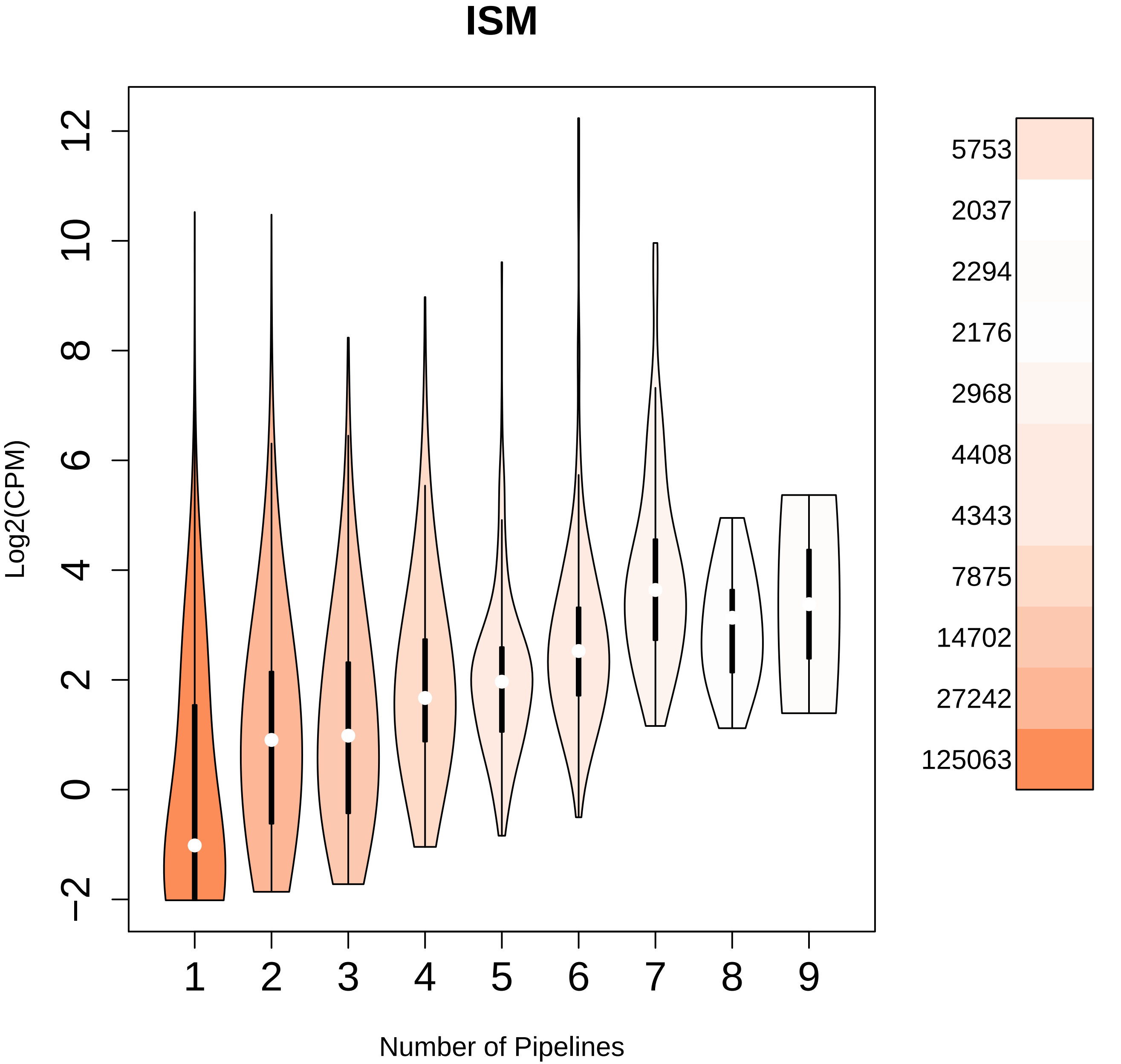
ISM
Log2(CPM)
12
10
8
6
4
2
0
−2
1
2
3
4
5
6
7
8
9
Number of Pipelines
5753
2037
2294
2176
2968
4408
4343
7875
14702
27242
125063

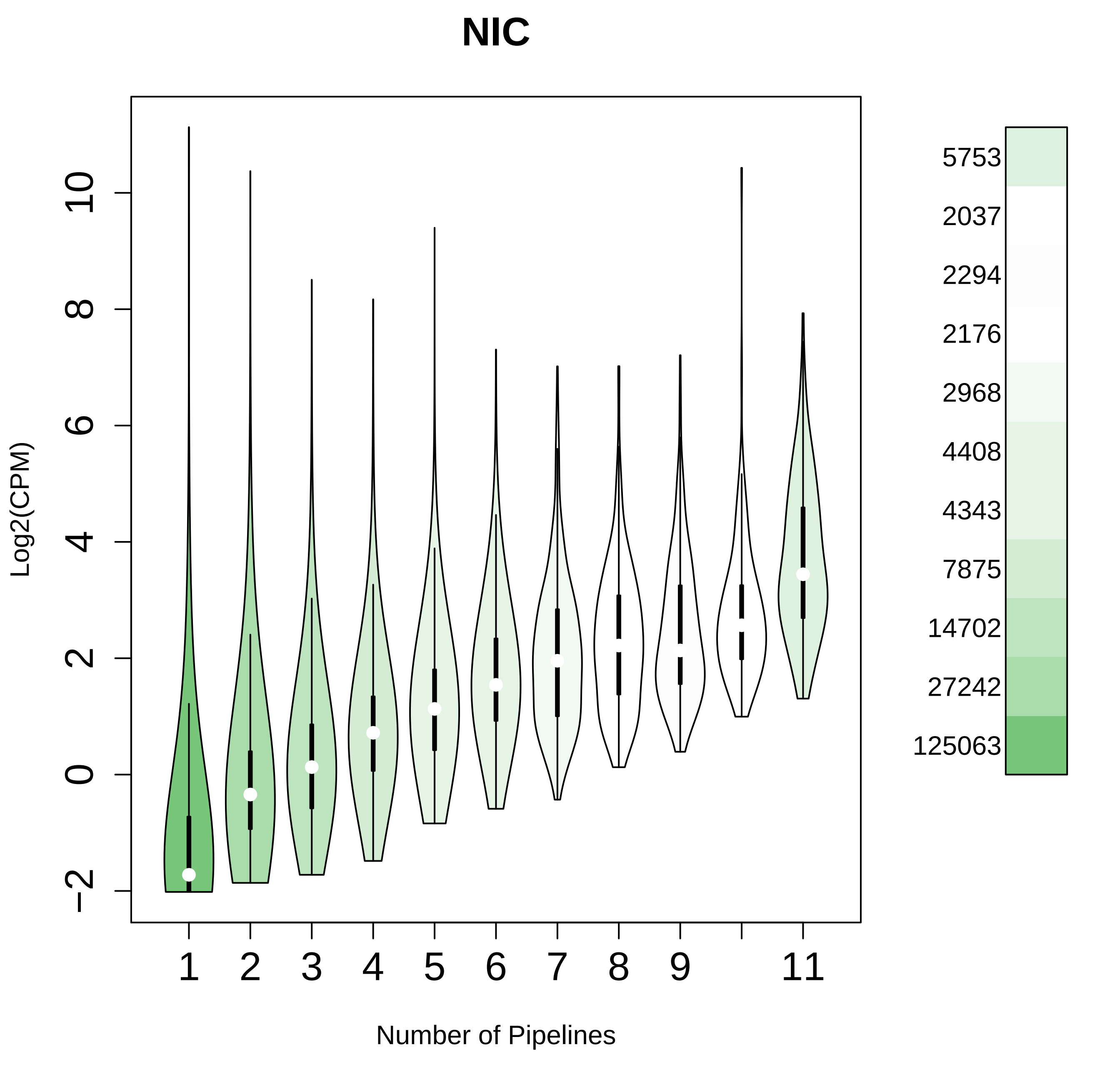
NIC
Log2(CPM)
−2
0
2
4
6
8
10
1
2
3
4
5
6
7
8
9
11
Number of Pipelines
5753
2037
2294
2176
2968
4408
4343
7875
14702
27242
125063

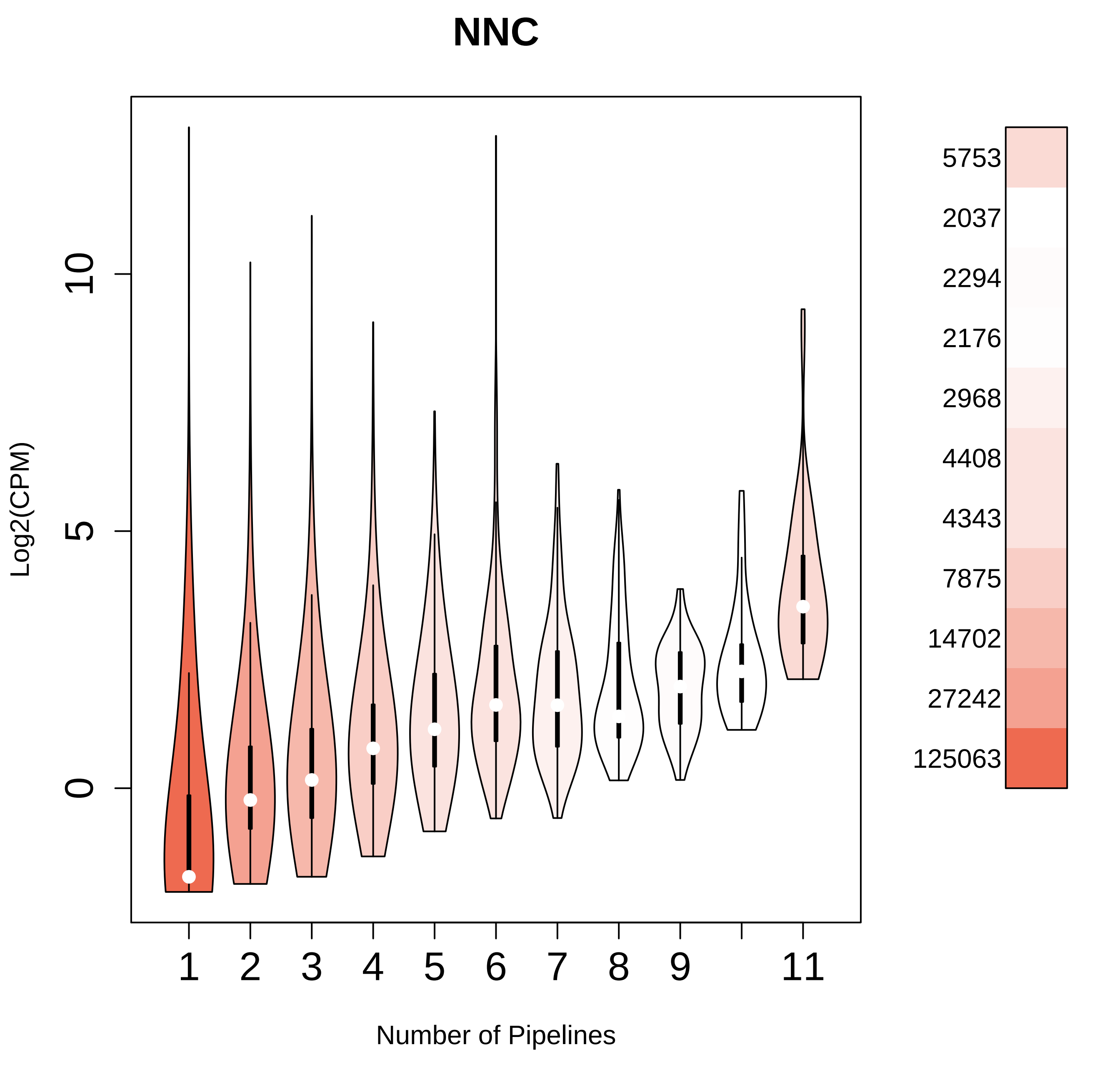
NNC
Log2(CPM)
0
5
10
1
2
3
4
5
6
7
8
9
11
Number of Pipelines
5753
2037
2294
2176
2968
4408
4343
7875
14702
27242
125063

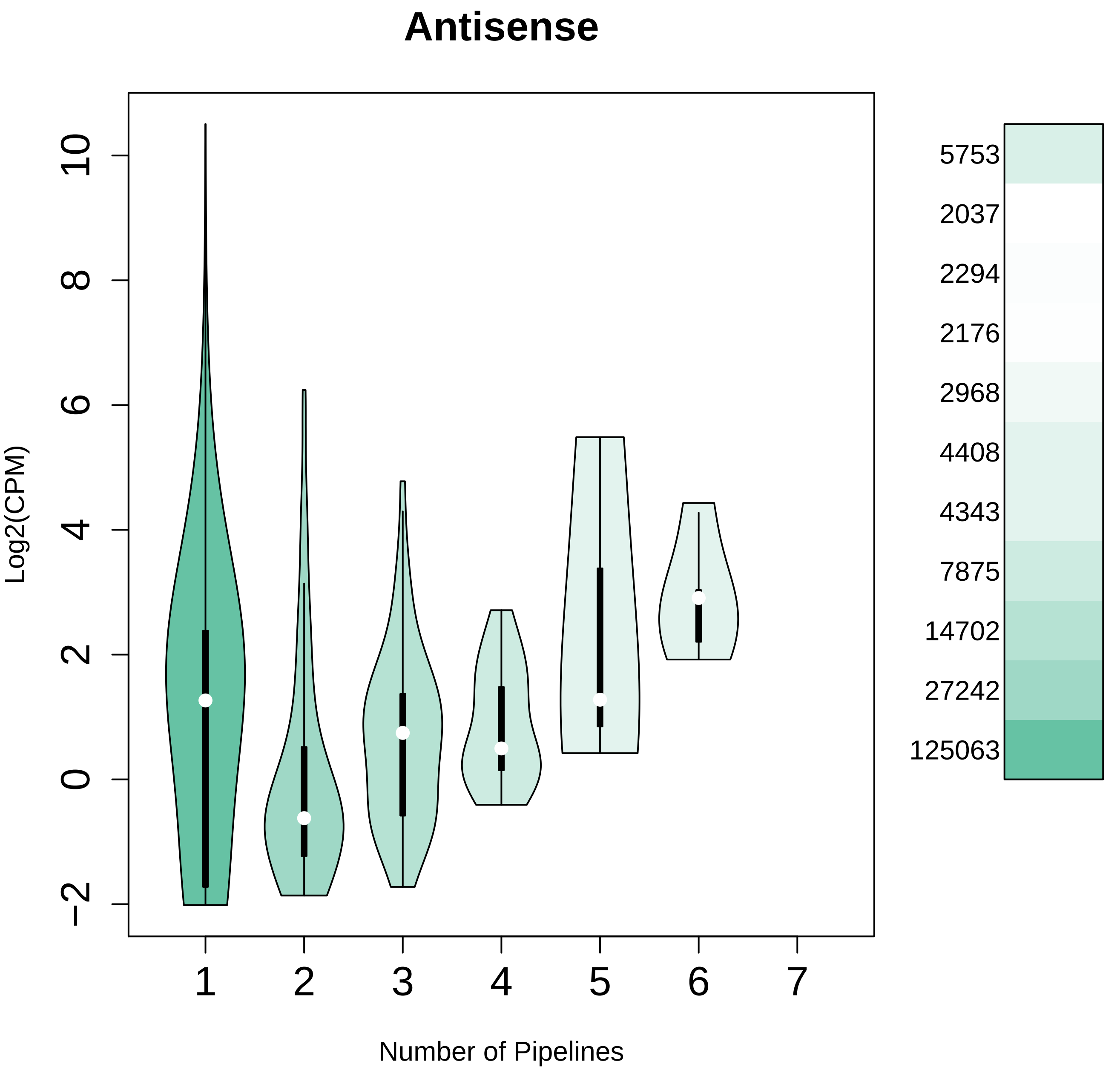

Antisense
Log2(CPM)
Number of Pipelines
−2
0
2
4
6
8
10
1
2
3
4
5
6
7
5753
2037
2294
2176
2968
4408
4343
7875
14702
27242
125063

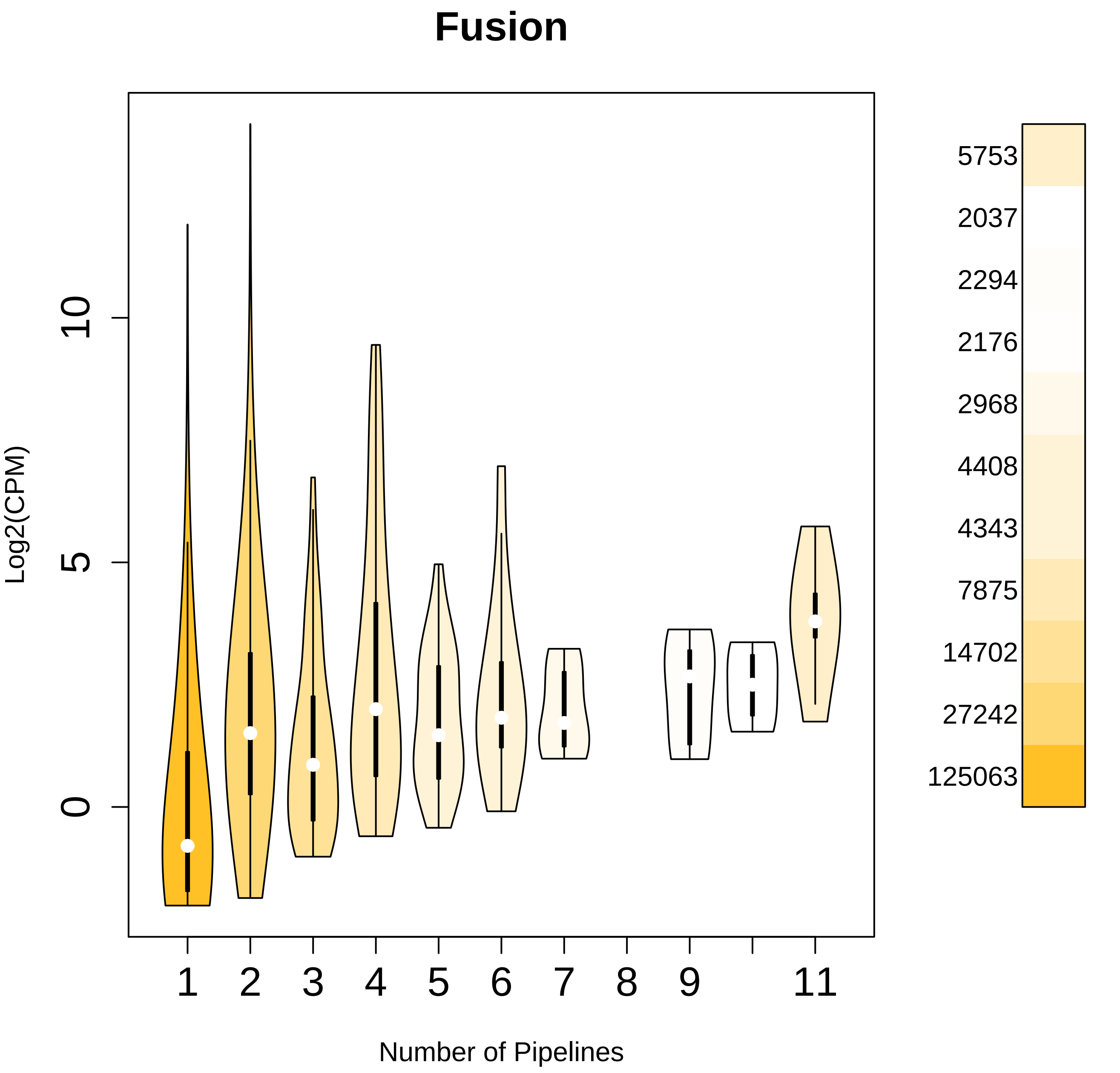

Fusion
Log2(CPM)
0
5
10
1
2
3
4
5
6
7
8
9
11
Number of Pipelines
5753
2037
2294
2176
2968
4408
4343
7875
14702
27242
125063

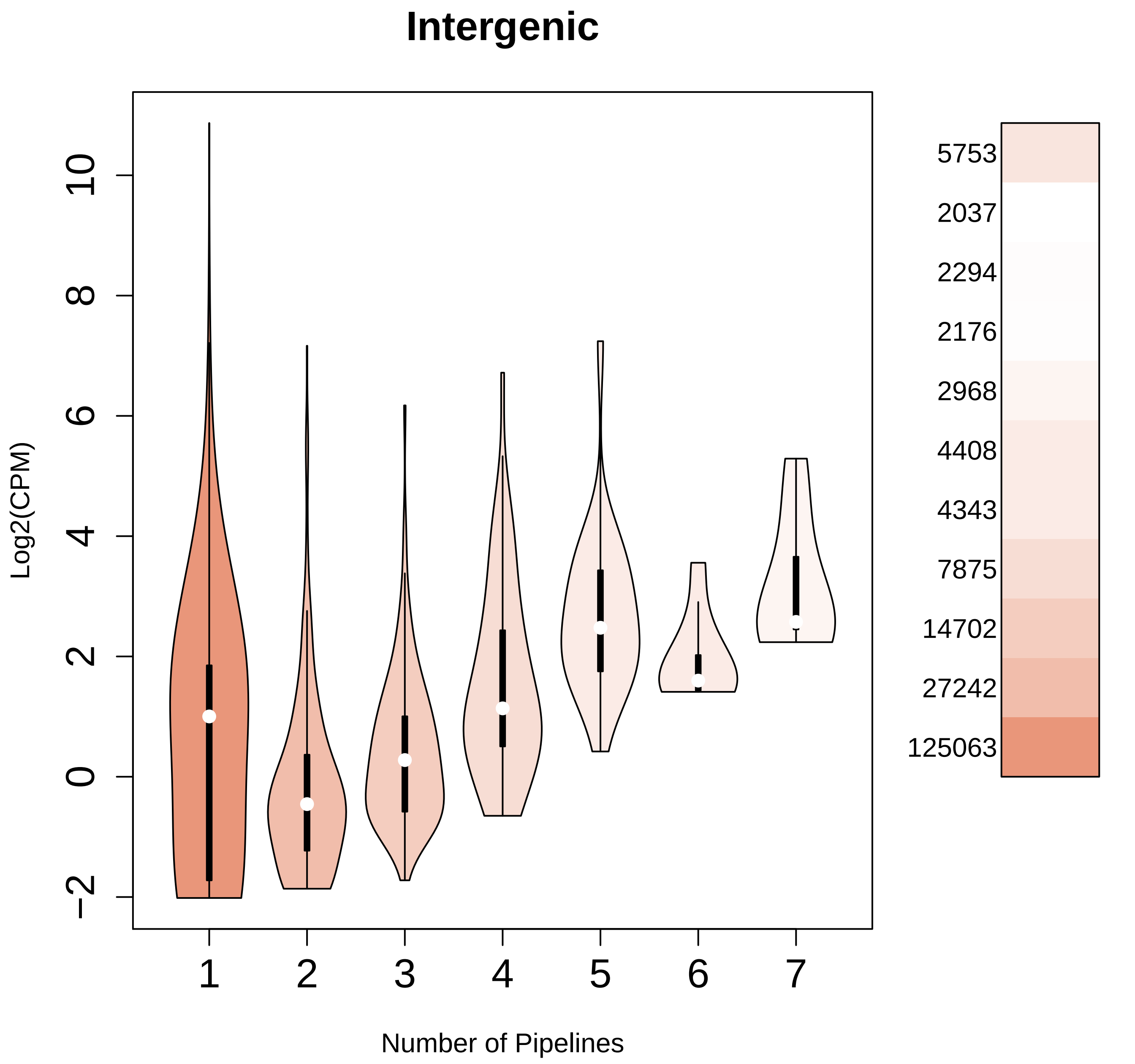

Intergenic
Log2(CPM)
10
8
6
4
2
0
−2
1
2
3
4
5
6
7
Number of Pipelines
5753
2037
2294
2176
2968
4408
4343
7875
14702
27242
125063

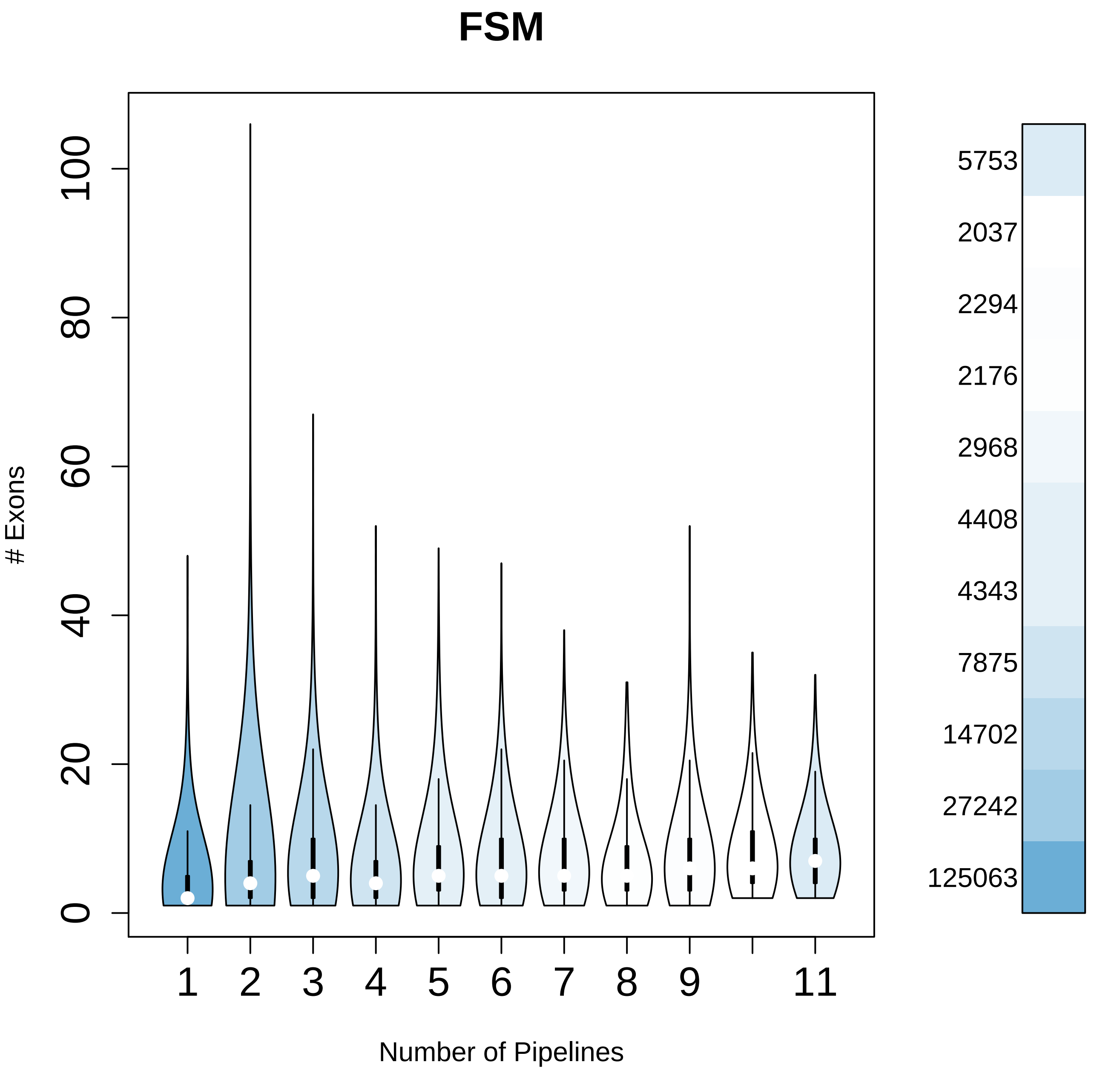

FSM
### Exons
0
20
40
60
80
100
1
2
3
4
5
6
7
8
9
11
Number of Pipelines
5753
2037
2294
2176
2968
4408
4343
7875
14702
27242
125063

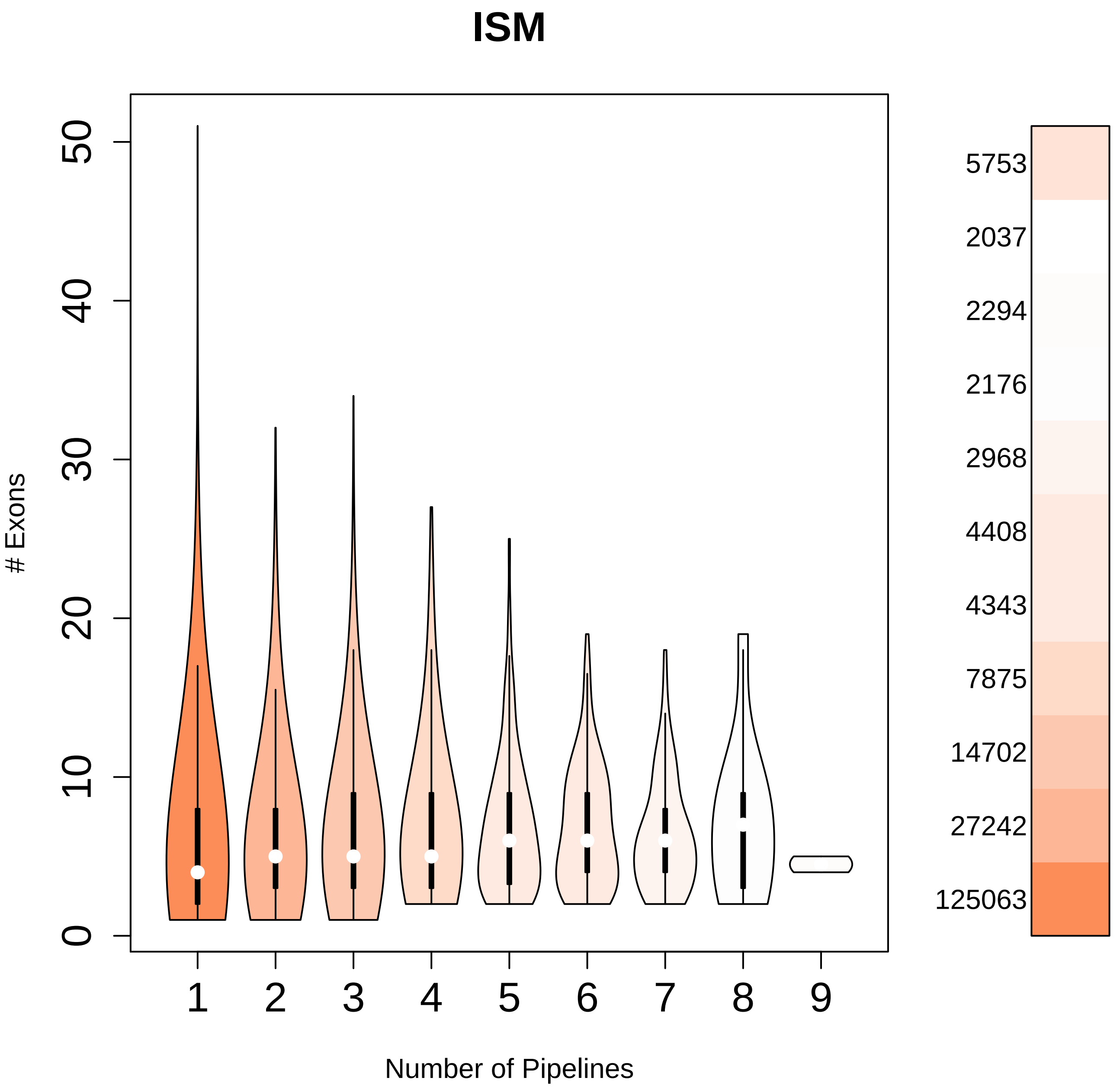

ISM
### Exons
0
10
20
30
40
50
1
2
3
4
5
6
7
8
9
Number of Pipelines
5753
2037
2294
2176
2968
4408
4343
7875
14702
27242
125063

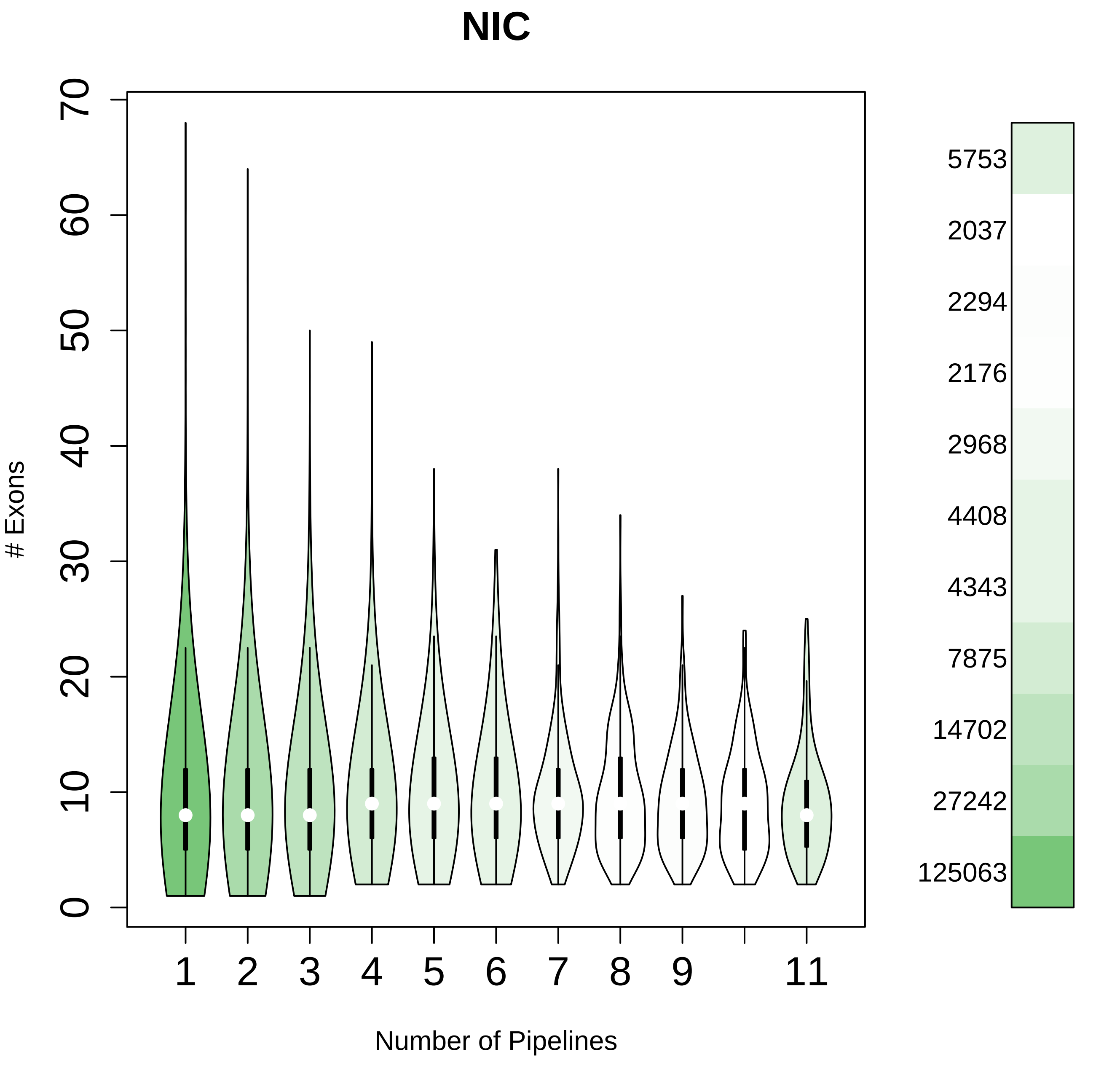

NIC
### Exons
0
10
20
30
40
50
60
70
1
2
3
4
5
6
7
8
9
11
Number of Pipelines
5753
2037
2294
2176
2968
4408
4343
7875
14702
27242
125063

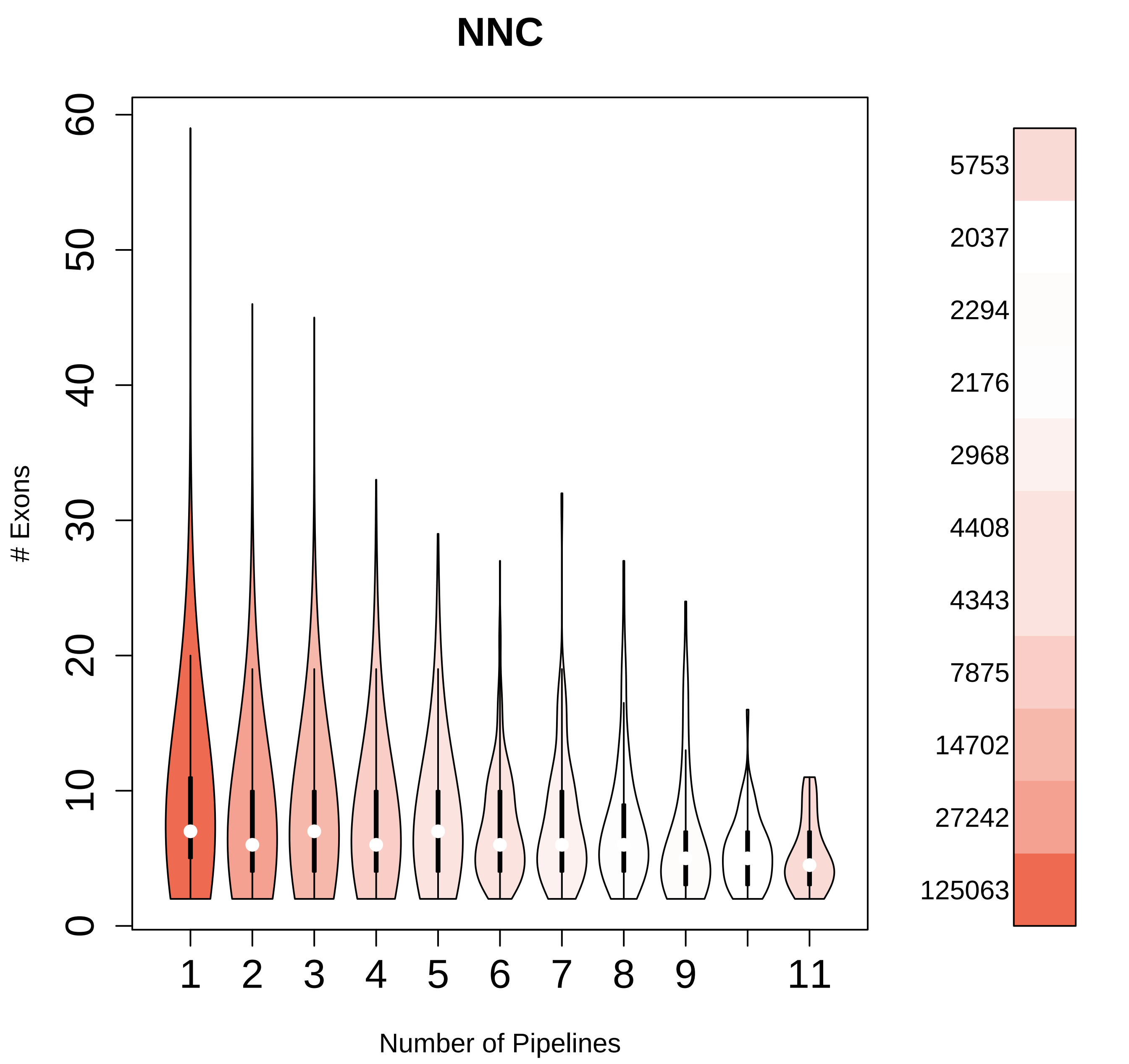

NNC
### Exons
Number of Pipelines
0
10
20
30
40
50
60
1
2
3
4
5
6
7
8
9
11
5753
2037
2294
2176
2968
4408
4343
7875
14702
27242
125063

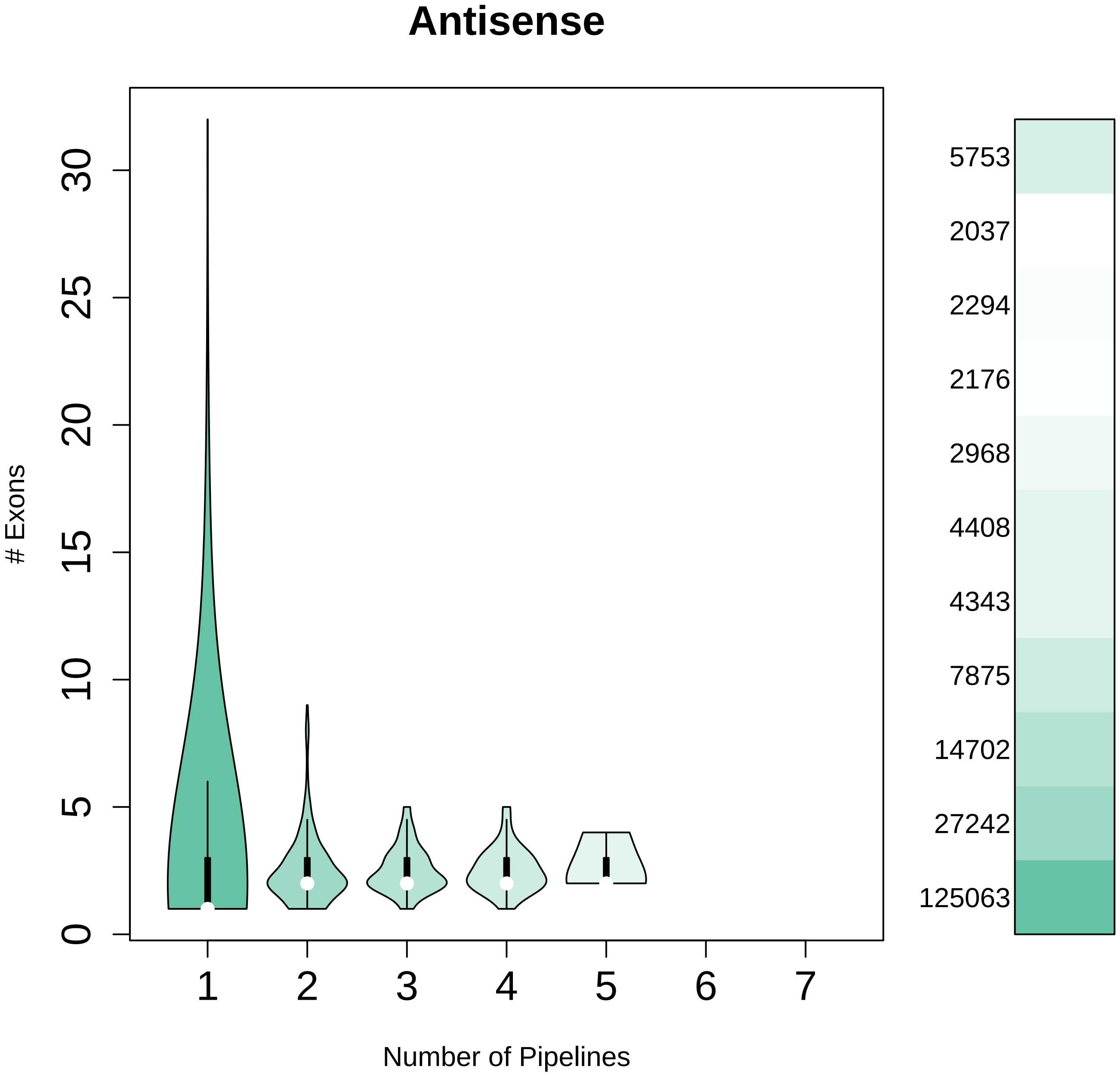
Antisense
### Exons
0
5
10
15
20
25
30
1
2
3
4
5
6
7
Number of Pipelines
5753
2037
2294
2176
2968
4408
4343
7875
14702
27242
125063

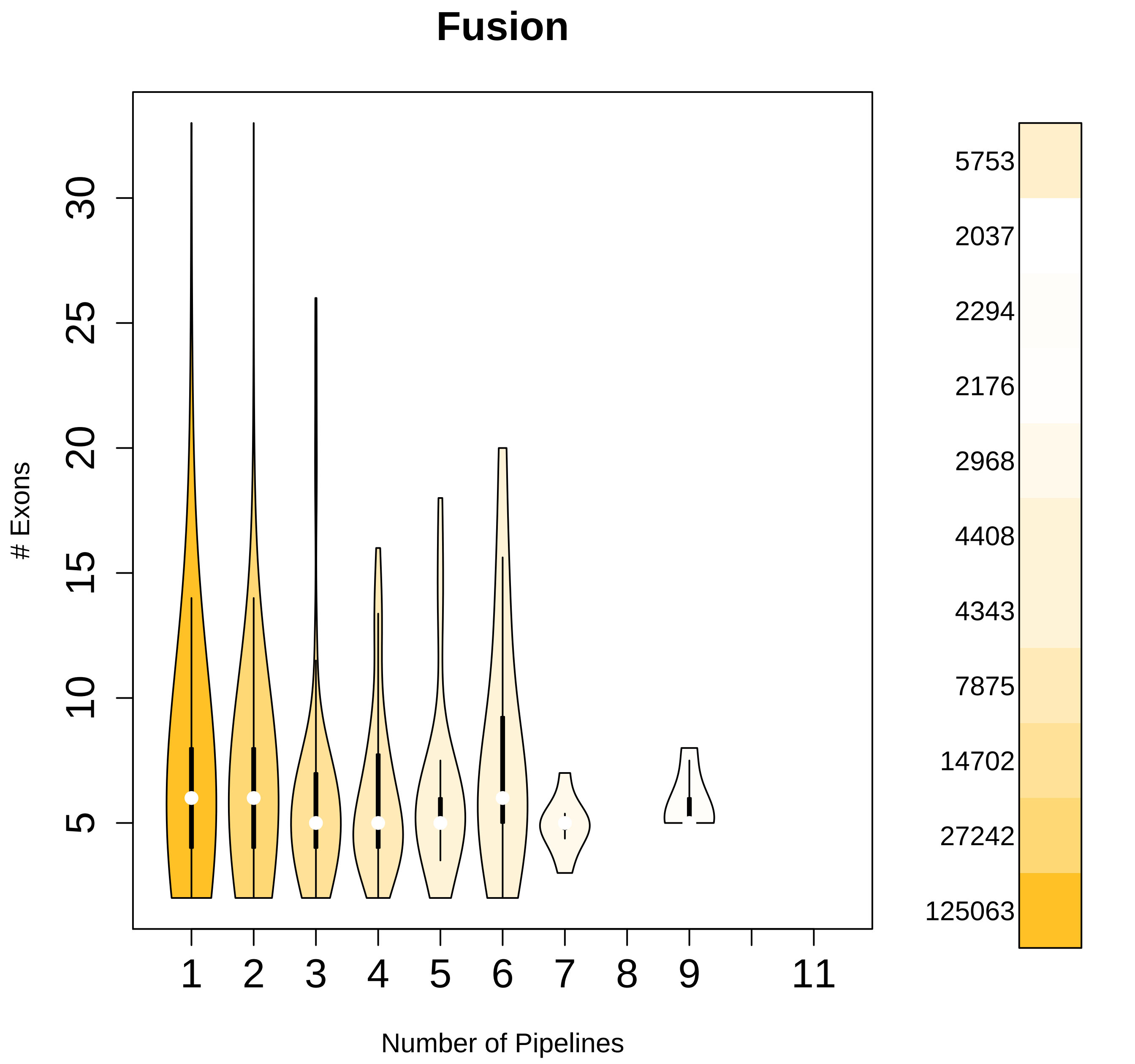

Fusion
30
25
20
15
10
5
### Exons
1
2
3
4
5
6
7
8
9
11
Number of Pipelines
5753
2037
2294
2176
2968
4408
4343
7875
14702
27242
125063

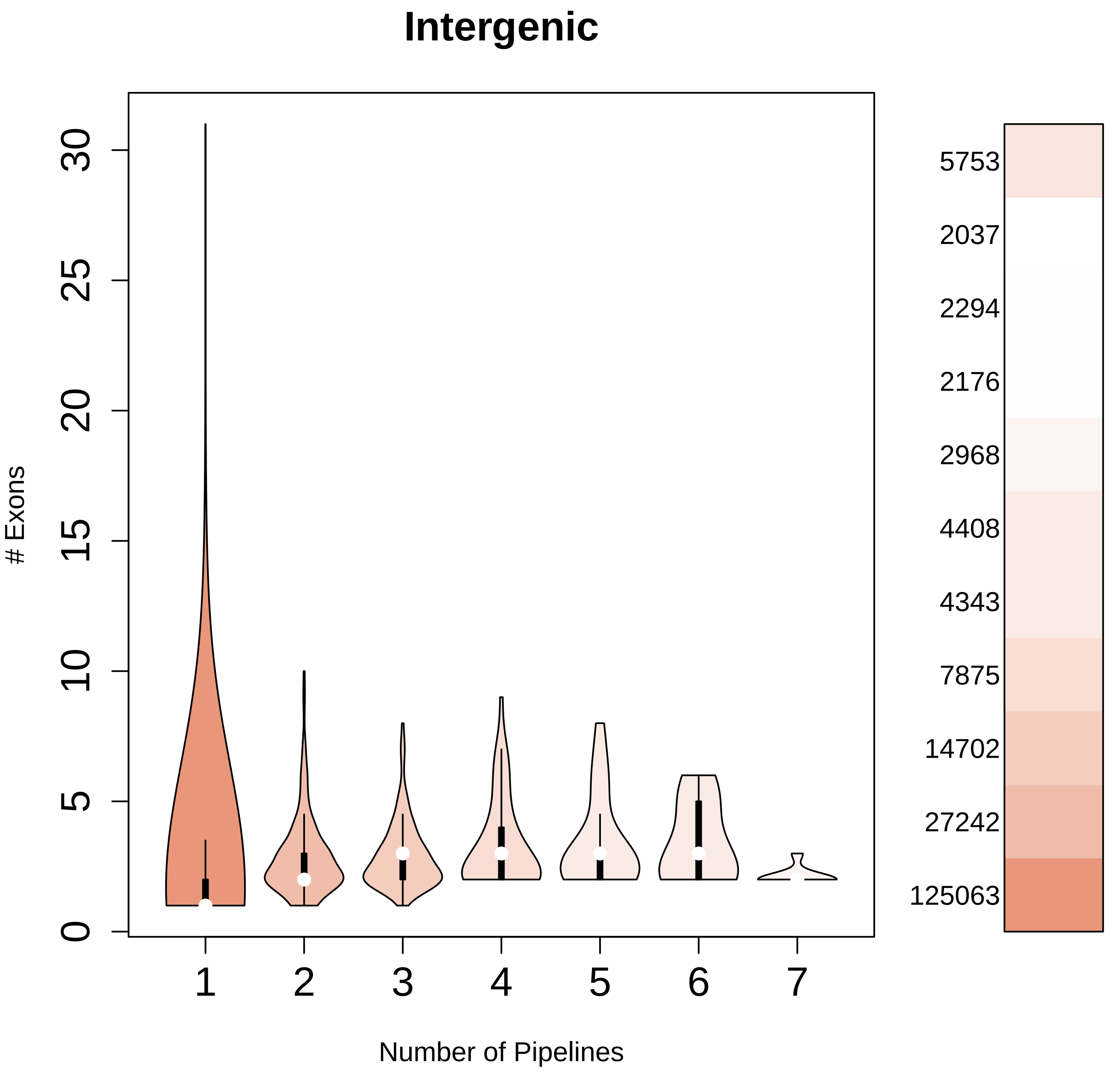

Intergenic
### Exons
0
5
10
15
20
25
30
1
2
3
4
5
6
7
Number of Pipelines
5753
2037
2294
2176
2968
4408
4343
7875
14702
27242
125063

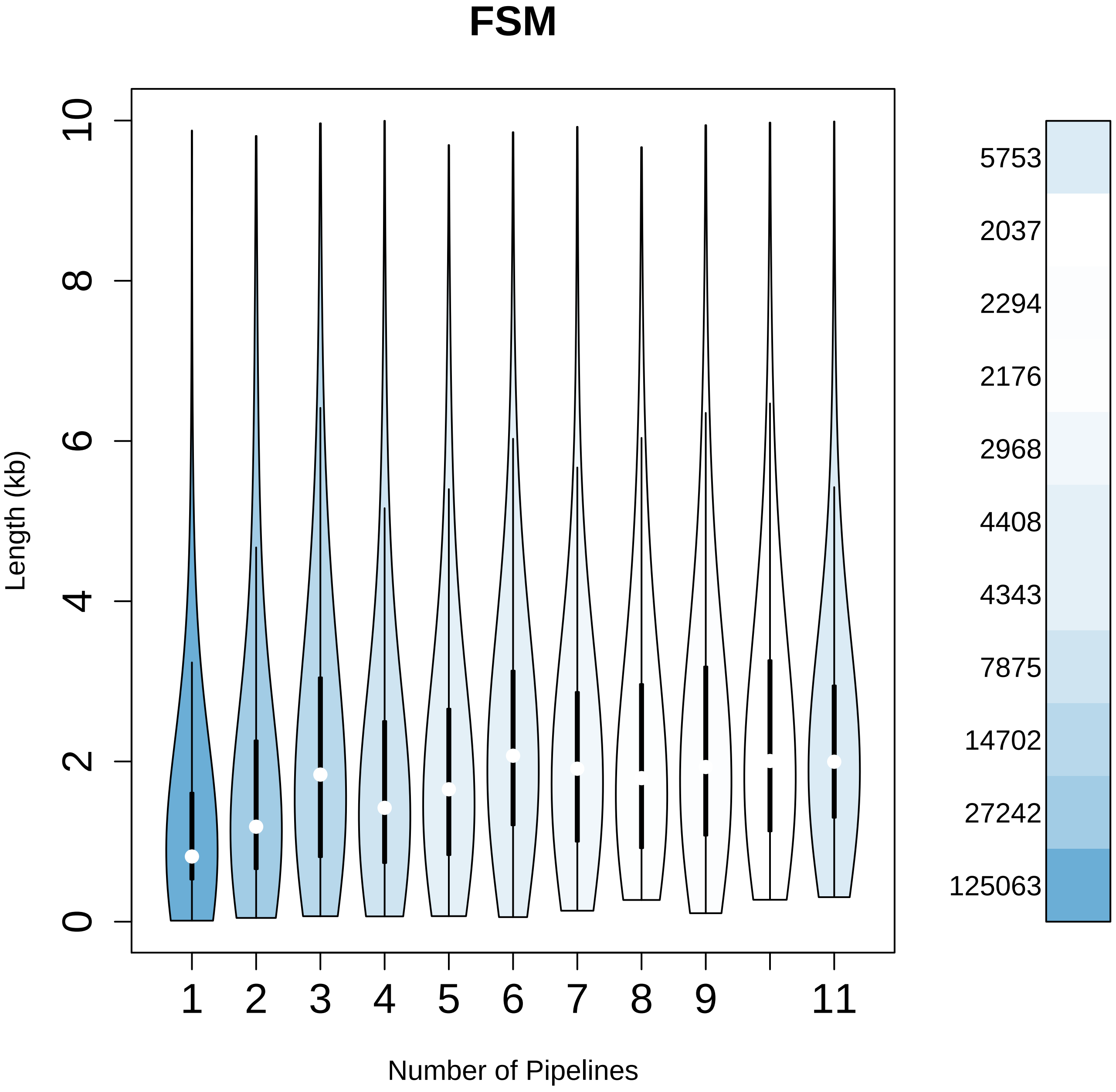
FSM
Length (kb)
0
2
4
6
8
10
1
2
3
4
5
6
7
8
9
11
Number of Pipelines
5753
2037
2294
2176
2968
4408
4343
7875
14702
27242
125063

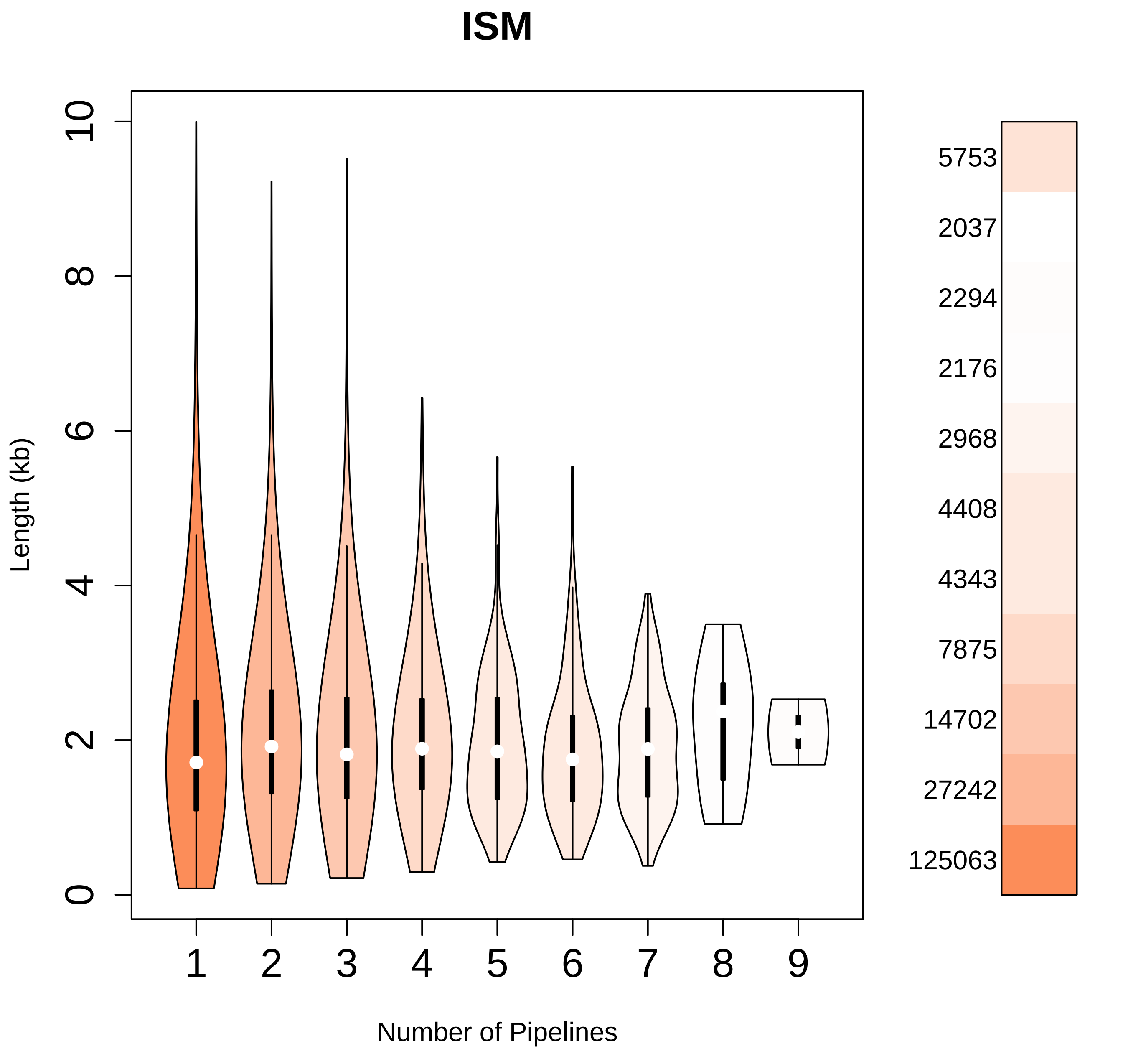

ISM
Length (kb)
0
2
4
6
8
10
1
2
3
4
5
6
7
8
9
Number of Pipelines
5753
2037
2294
2176
2968
4408
4343
7875
14702
27242
125063

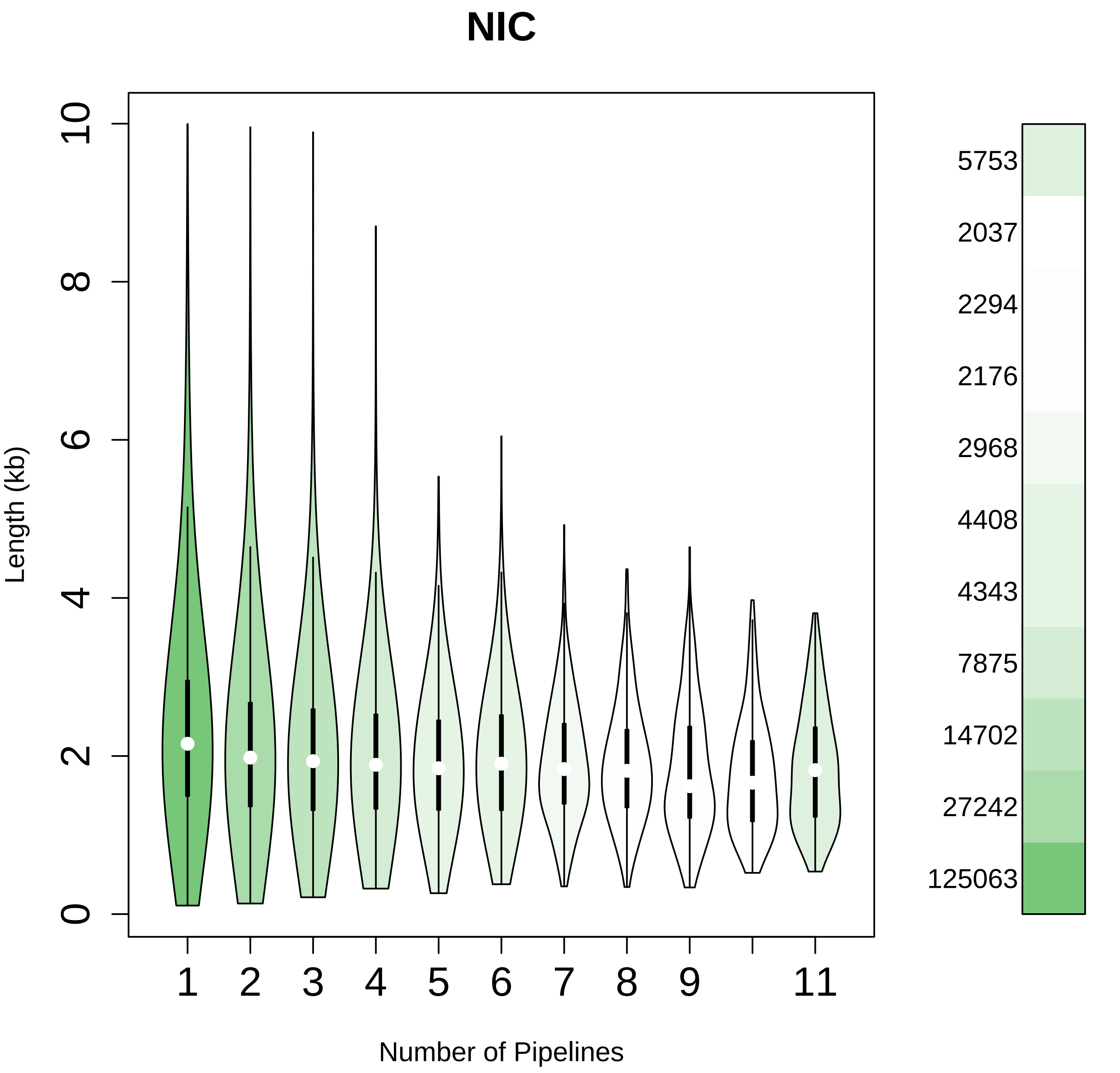

NIC
Length (kb)
0
2
4
6
8
10
1
2
3
4
5
6
7
8
9
11
Number of Pipelines
5753
2037
2294
2176
2968
4408
4343
7875
14702
27242
125063

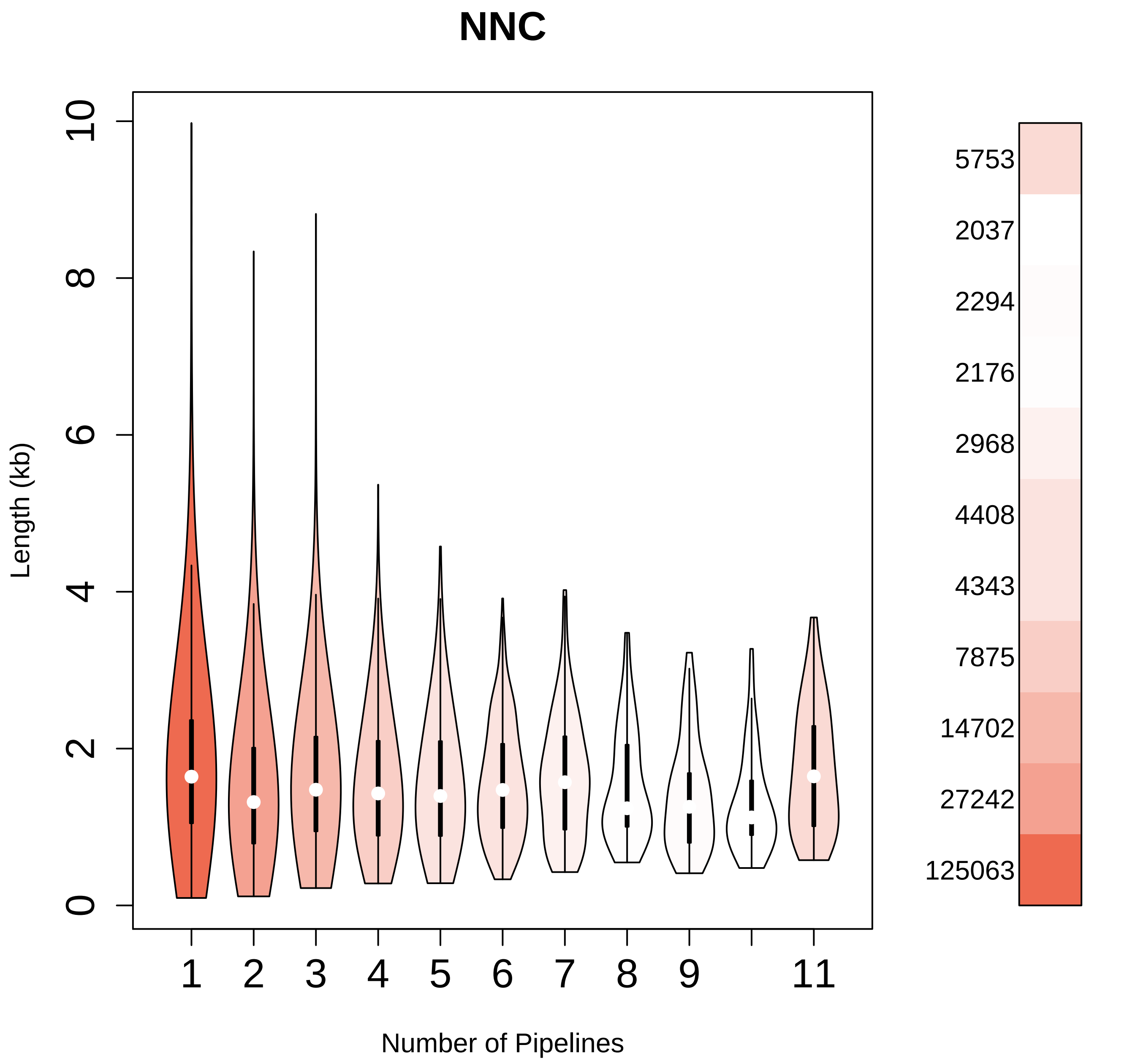

NNC
Length (kb)
0
2
4
6
8
10
1
2
3
4
5
6
7
8
9
11
Number of Pipelines
5753
2037
2294
2176
2968
4408
4343
7875
14702
27242
125063

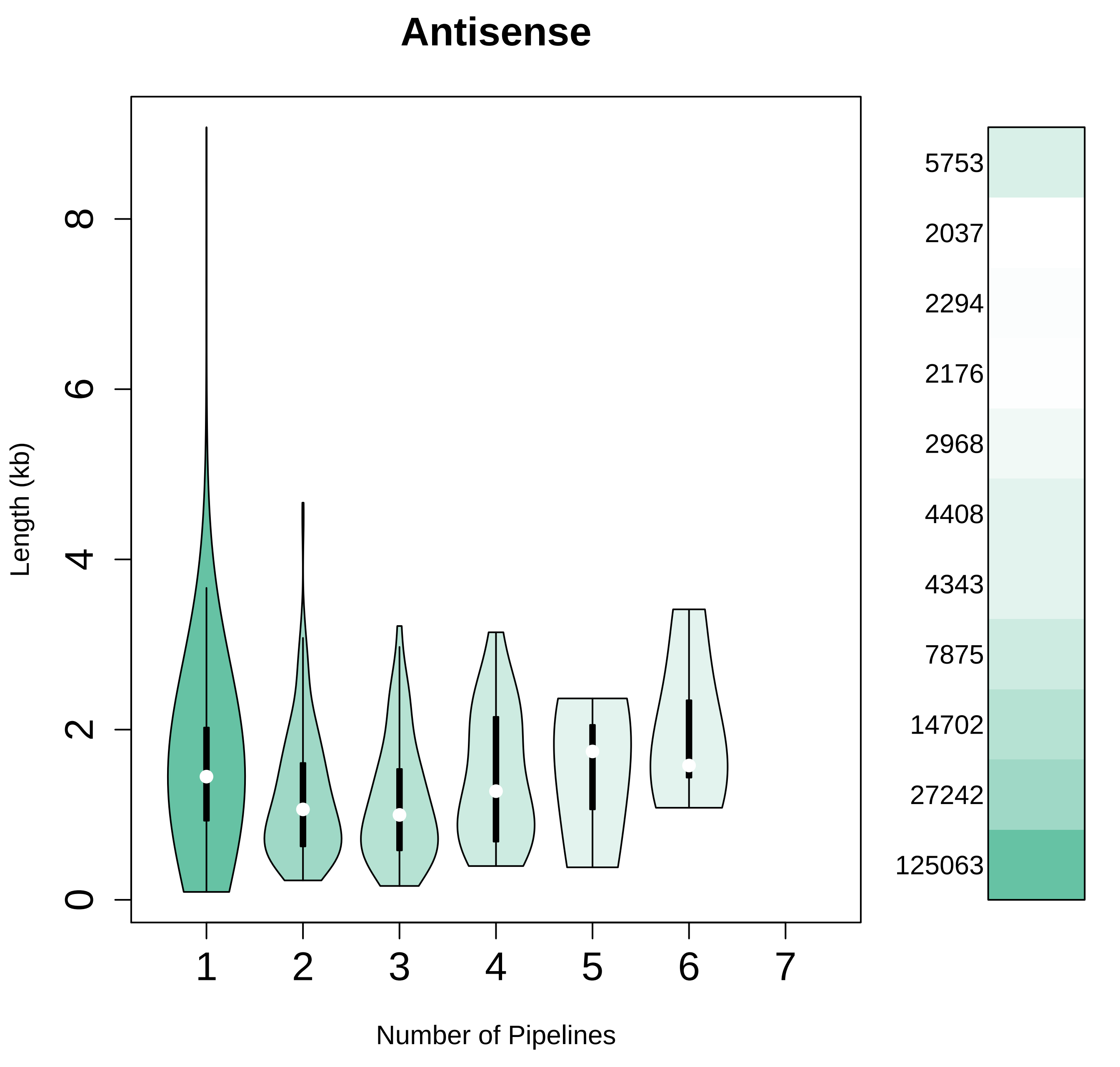
Antisense
Length (kb)
0
2
4
6
8
1
2
3
4
5
6
7
Number of Pipelines
5753
2037
2294
2176
2968
4408
4343
7875
14702
27242
125063

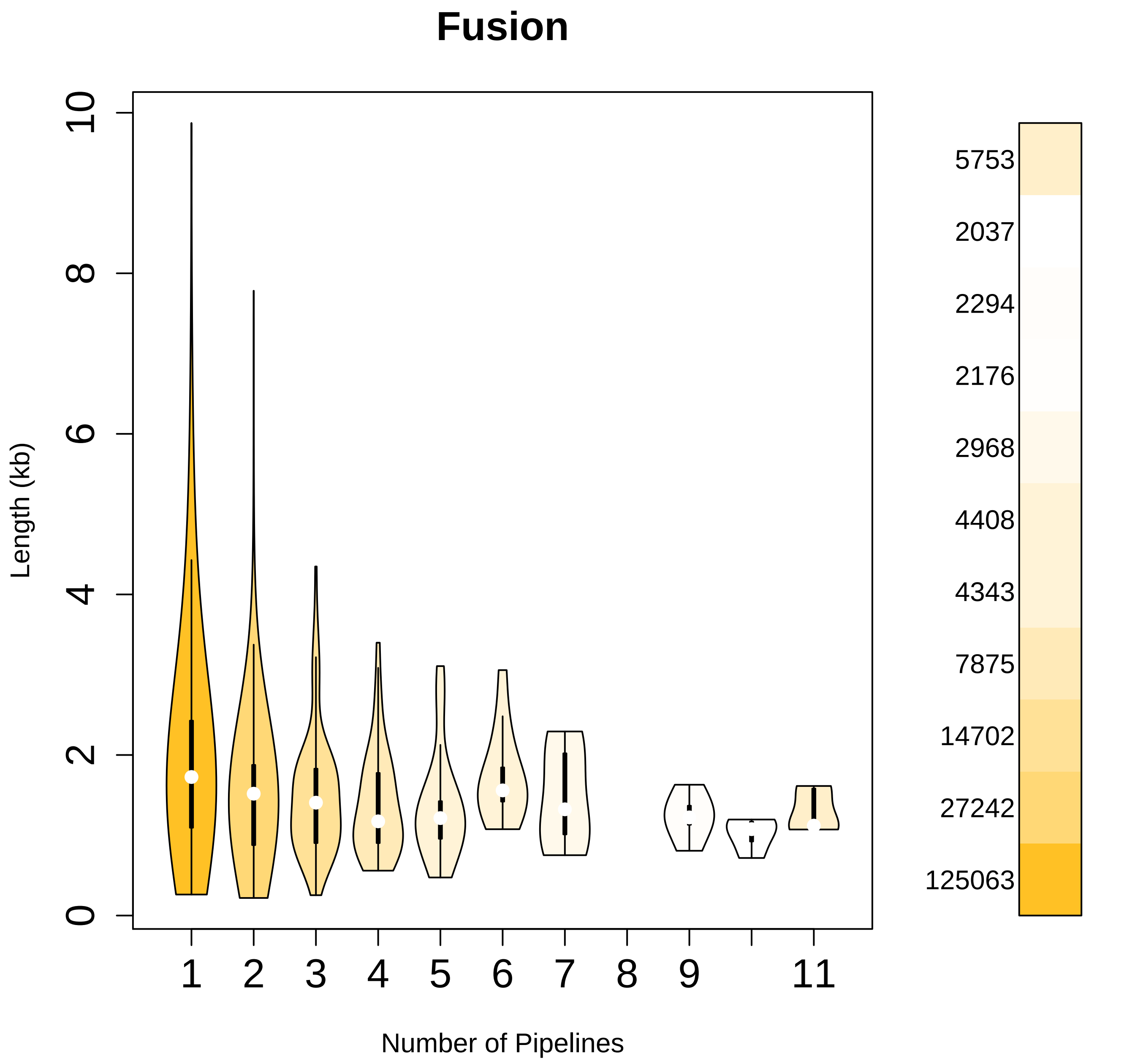
Fusion
Length (kb)
0
2
4
6
8
10
1
2
3
4
5
6
7
8
9
11
Number of Pipelines
5753
2037
2294
2176
2968
4408
4343
7875
14702
27242
125063

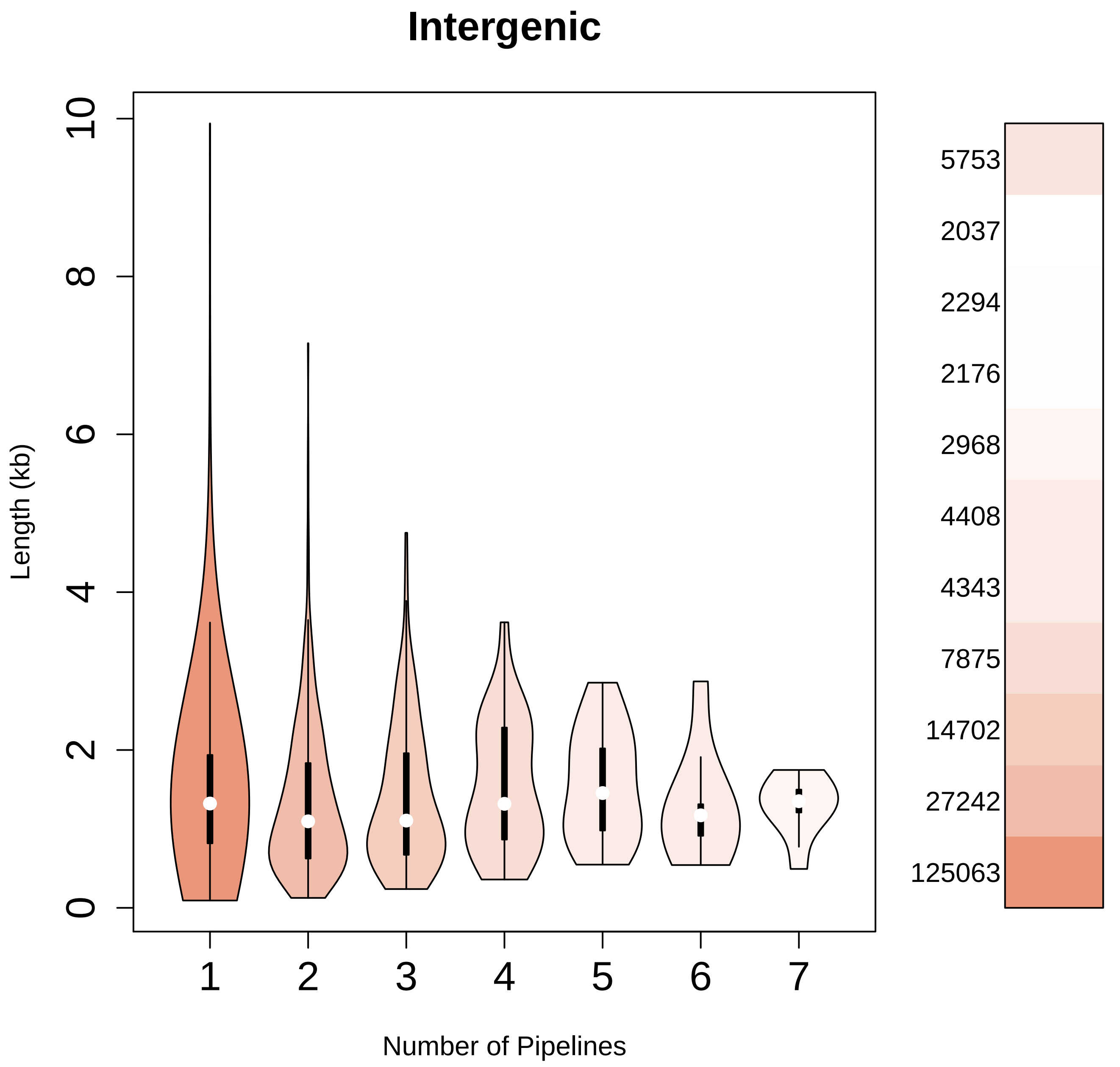

Intergenic
Length (kb)
0
2
4
6
8
10
1
2
3
4
5
6
7
Number of Pipelines
5753
2037
2294
2176
2968
4408
4343
7875
14702
27242
125063

*Standard Deviation of TSS and TTS*

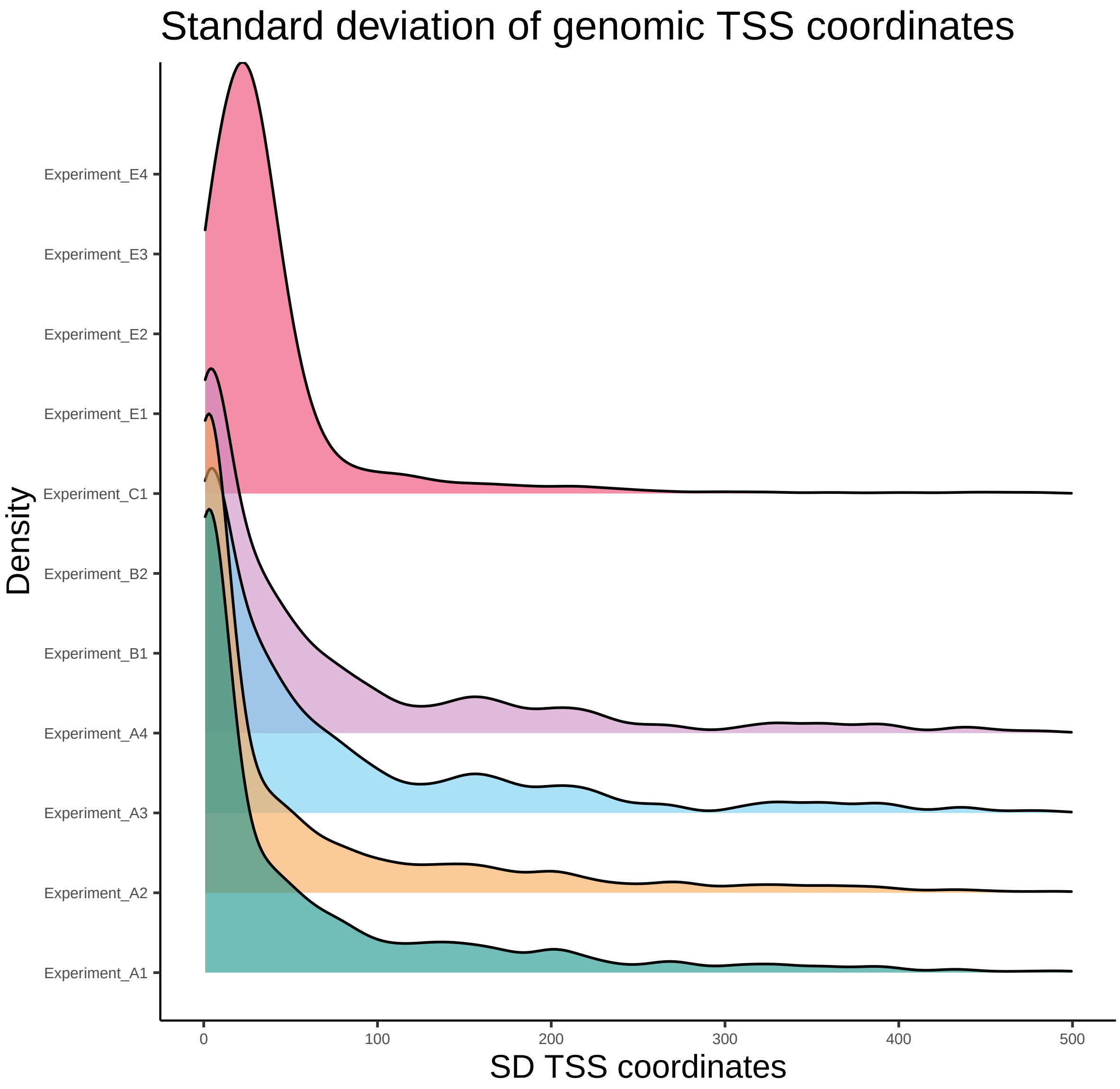

Standard deviation of genomic TSS coordinates
Density
Experiment_E4
Experiment_E3
Experiment_E2
Experiment_E1
Experiment_C1
Experiment_B2
Experiment_B1
Experiment_A4
Experiment_A3
Experiment_A2
Experiment_A1
0
100
200
300
400
500
SD TSS coordinates

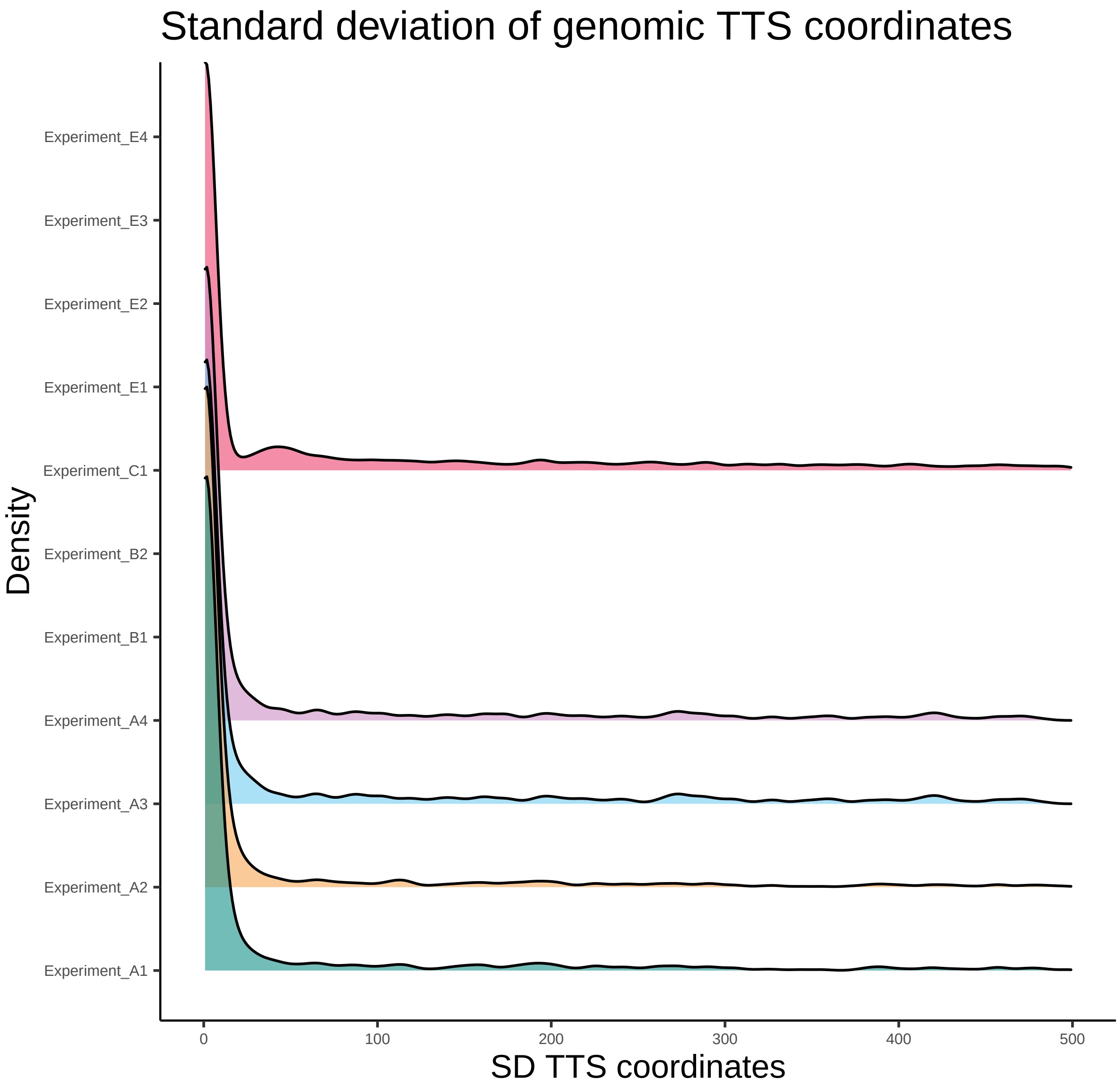
Standard deviation of genomic TTS coordinates
Density
Experiment_E4
Experiment_E3
Experiment_E2
Experiment_E1
Experiment_C1
Experiment_B2
Experiment_B1
Experiment_A4
Experiment_A3
Experiment_A2
Experiment_A1
0
100
200
300
400
500
SD TTS coordinates

*LRGASP metrics Comparison*

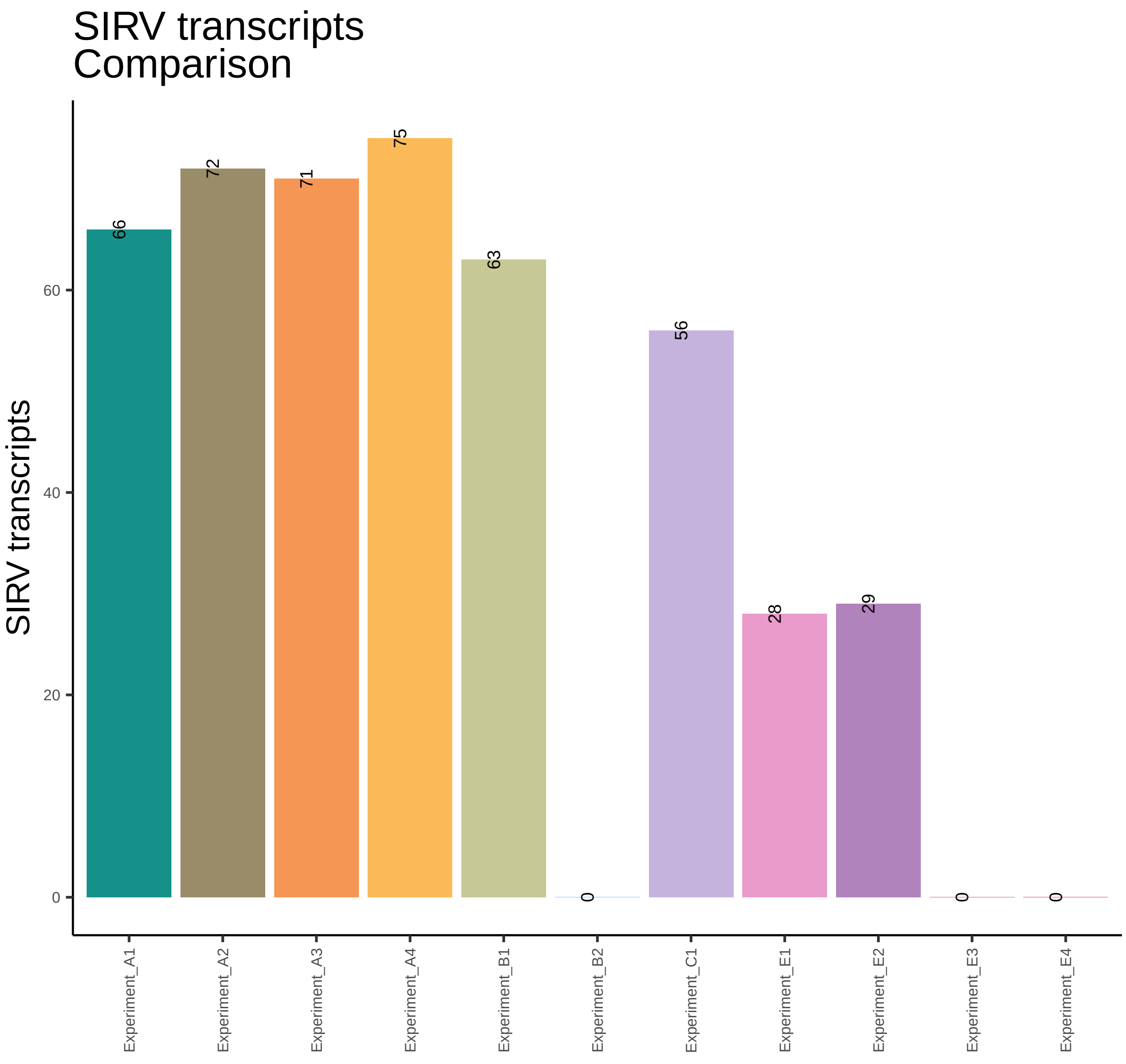

SIRV transcripts
Comparison
SIRV transcripts
0
20
40
60
66
72
71
75
63
0
56
28
29
0
0
Experiment_A1
Experiment_A2
Experiment_A3
Experiment_A4
Experiment_B1
Experiment_B2
Experiment_C1
Experiment_E1
Experiment_E2
Experiment_E3
Experiment_E4

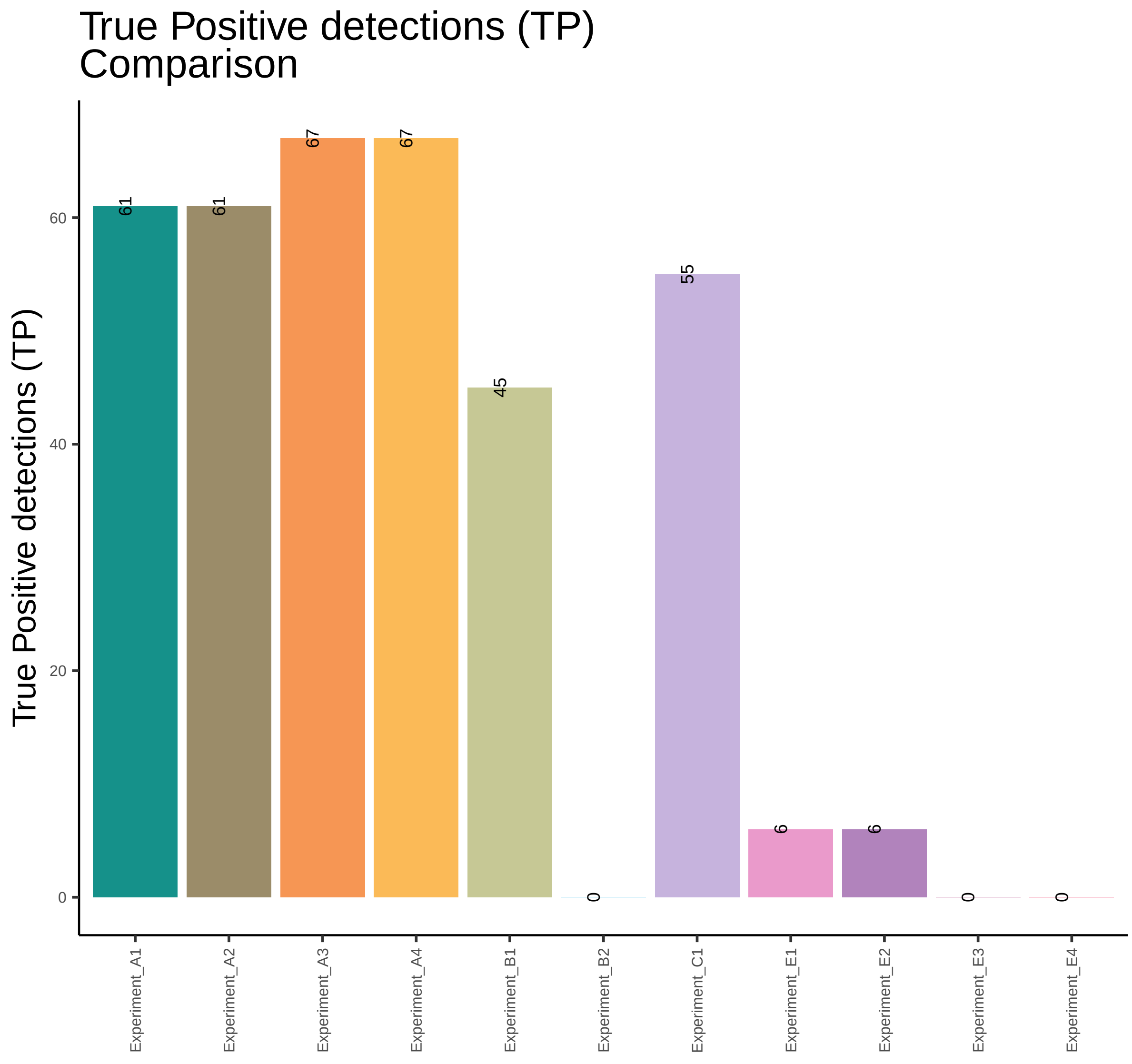

True Positive detections (TP)
Comparison
True Positive detections (TP)
0
20
40
60
61
61
67
67
45
0
55
6
6
0
0
Experiment_A1
Experiment_A2
Experiment_A3
Experiment_A4
Experiment_B1
Experiment_B2
Experiment_C1
Experiment_E1
Experiment_E2
Experiment_E3
Experiment_E4

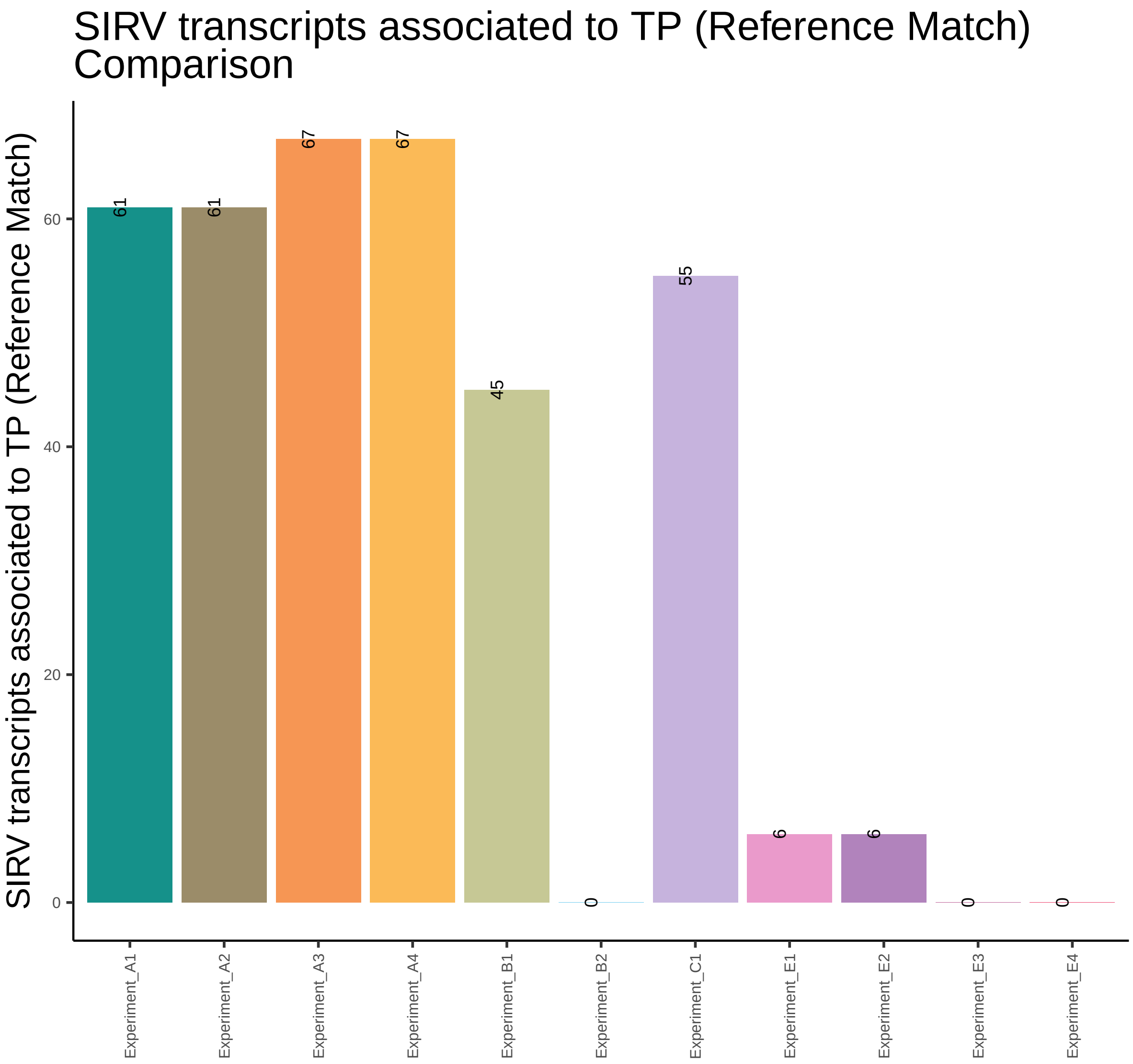

SIRV transcripts associated to TP (Reference Match)
Comparison
SIRV transcripts associated to TP (Reference Match)
0
20
40
60
61
61
67
67
45
0
55
6
6
0
0
Experiment_A1
Experiment_A2
Experiment_A3
Experiment_A4
Experiment_B1
Experiment_B2
Experiment_C1
Experiment_E1
Experiment_E2
Experiment_E3
Experiment_E4

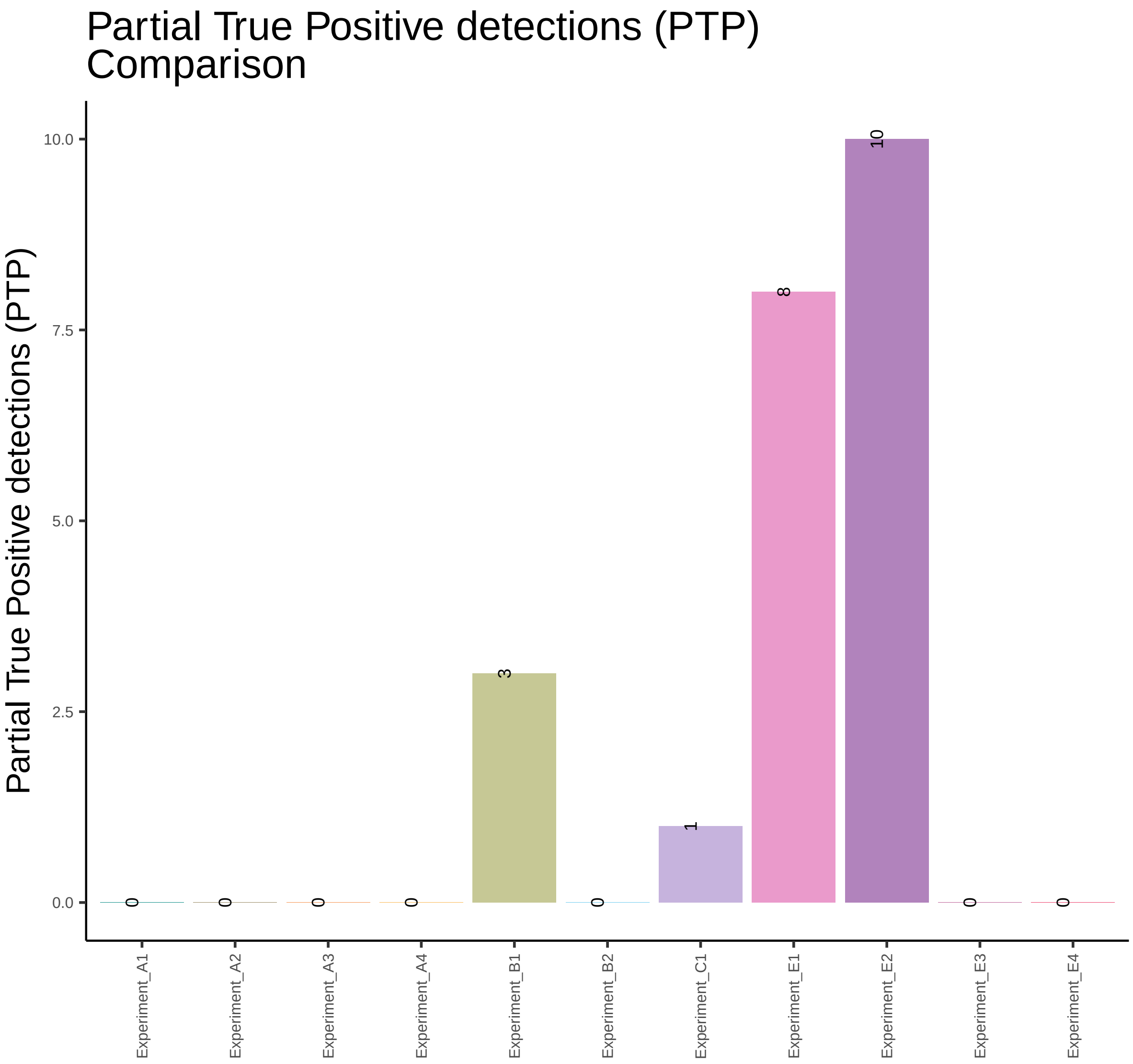

Partial True Positive detections (PTP)
Comparison
Partial True Positive detections (PTP)
10.0
7.5
5.0
2.5
0.0
0
0
0
0
3
0
1
8
10
0
0
Experiment_A1
Experiment_A2
Experiment_A3
Experiment_A4
Experiment_B1
Experiment_B2
Experiment_C1
Experiment_E1
Experiment_E2
Experiment_E3
Experiment_E4

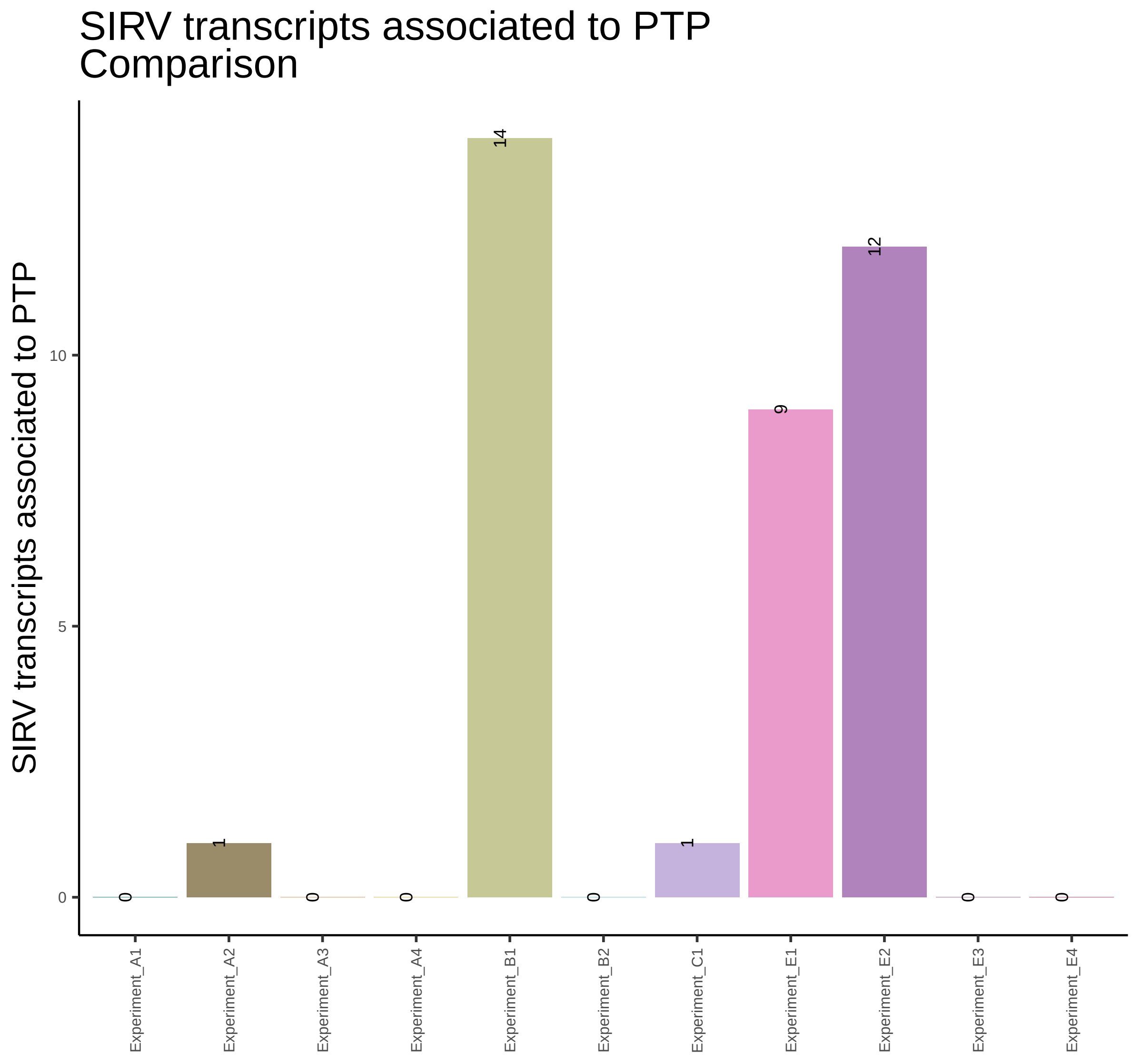
SIRV transcripts associated to PTP
Comparison
SIRV transcripts associated to PTP
0
5
10
0
1
0
0
14
0
1
9
12
0
0
Experiment_A1
Experiment_A2
Experiment_A3
Experiment_A4
Experiment_B1
Experiment_B2
Experiment_C1
Experiment_E1
Experiment_E2
Experiment_E3
Experiment_E4

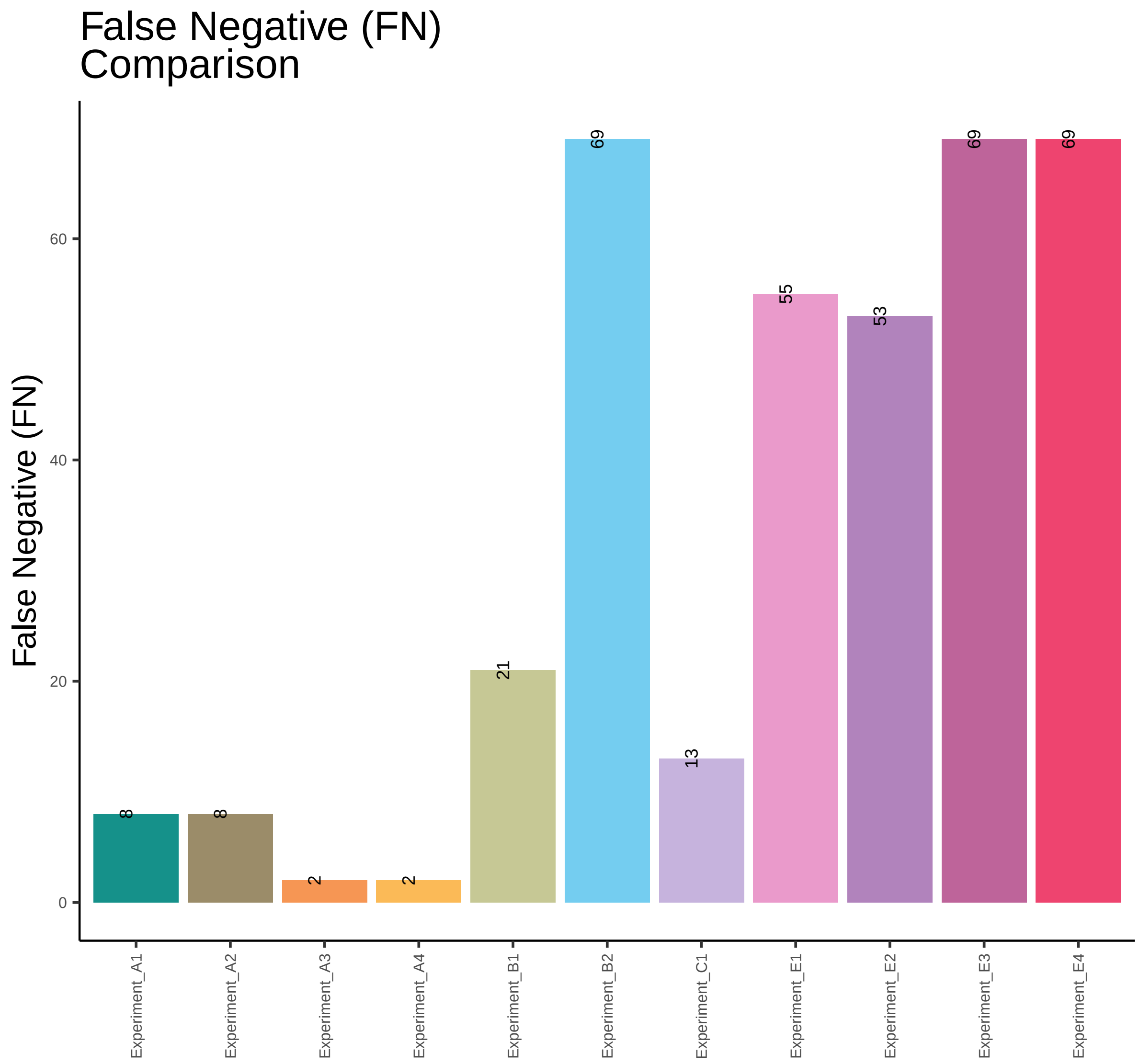

False Negative (FN) Comparison
False Negative (FN)
0
20
40
60
8
8
2
2
21
69
13
55
53
69
69
Experiment_A1
Experiment_A2
Experiment_A3
Experiment_A4
Experiment_B1
Experiment_B2
Experiment_C1
Experiment_E1
Experiment_E2
Experiment_E3
Experiment_E4

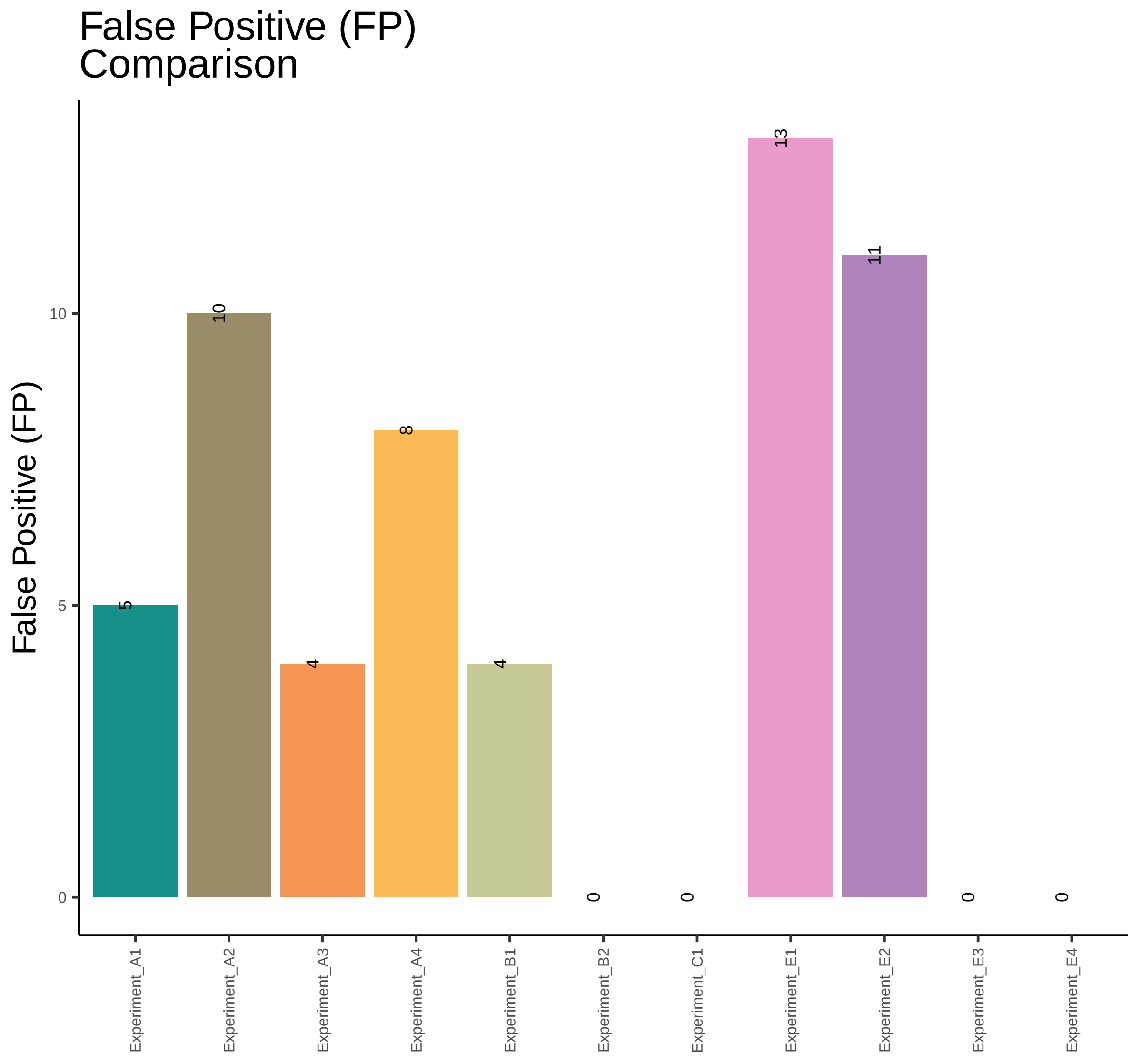

False Positive (FP)
Comparison
False Positive (FP)
0
5
10
5
10
4
8
4
0
0
13
11
0
0
Experiment_A1
Experiment_A2
Experiment_A3
Experiment_A4
Experiment_B1
Experiment_B2
Experiment_C1
Experiment_E1
Experiment_E2
Experiment_E3
Experiment_E4

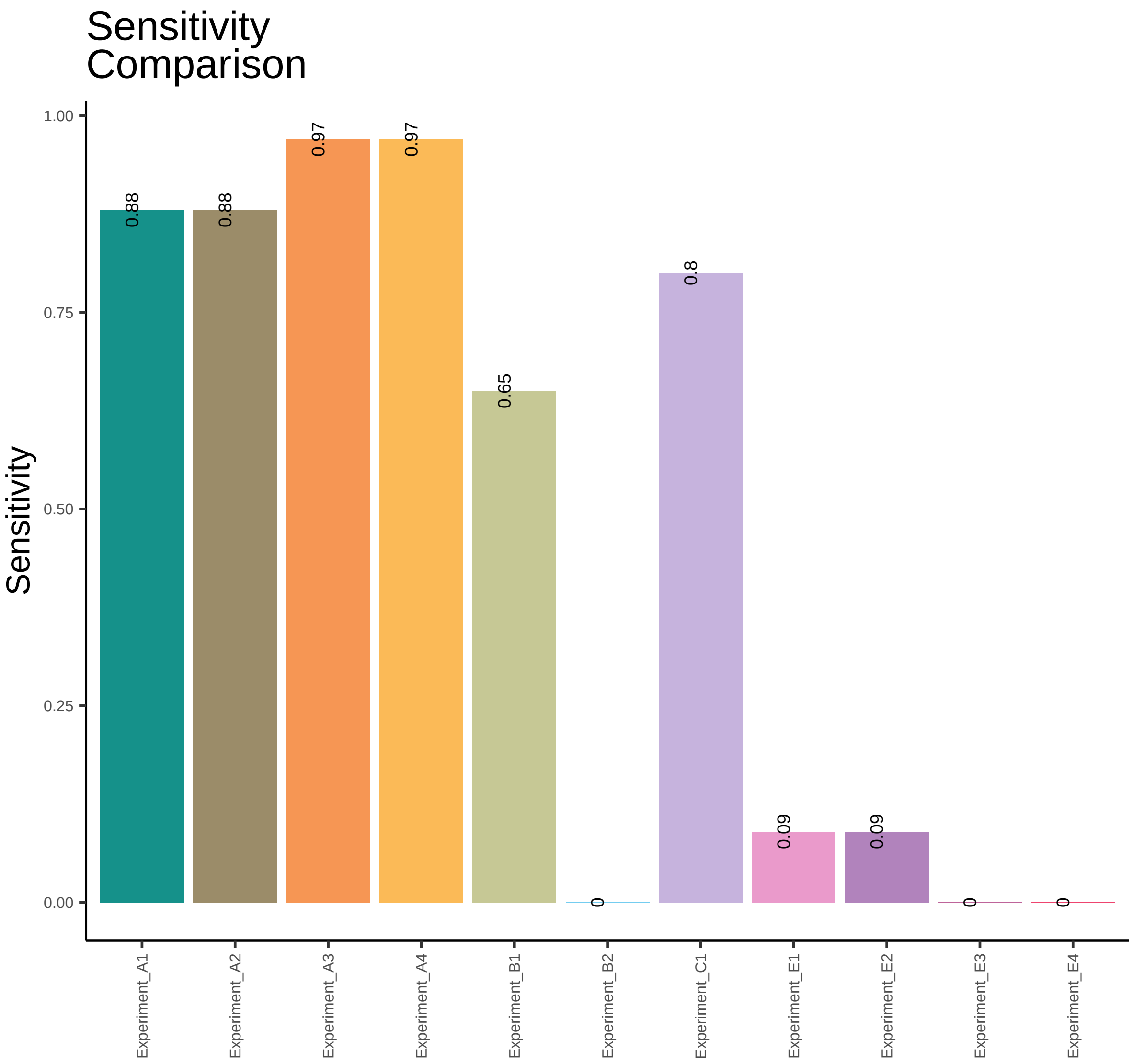
Sensitivity Comparison
Sensitivity
1.00
0.75
0.50
0.25
0.00
0.88
0.88
0.97
0.97
0.65
0
0.8
0.09
0.09
0
0
Experiment_A1
Experiment_A2
Experiment_A3
Experiment_A4
Experiment_B1
Experiment_B2
Experiment_C1
Experiment_E1
Experiment_E2
Experiment_E3
Experiment_E4

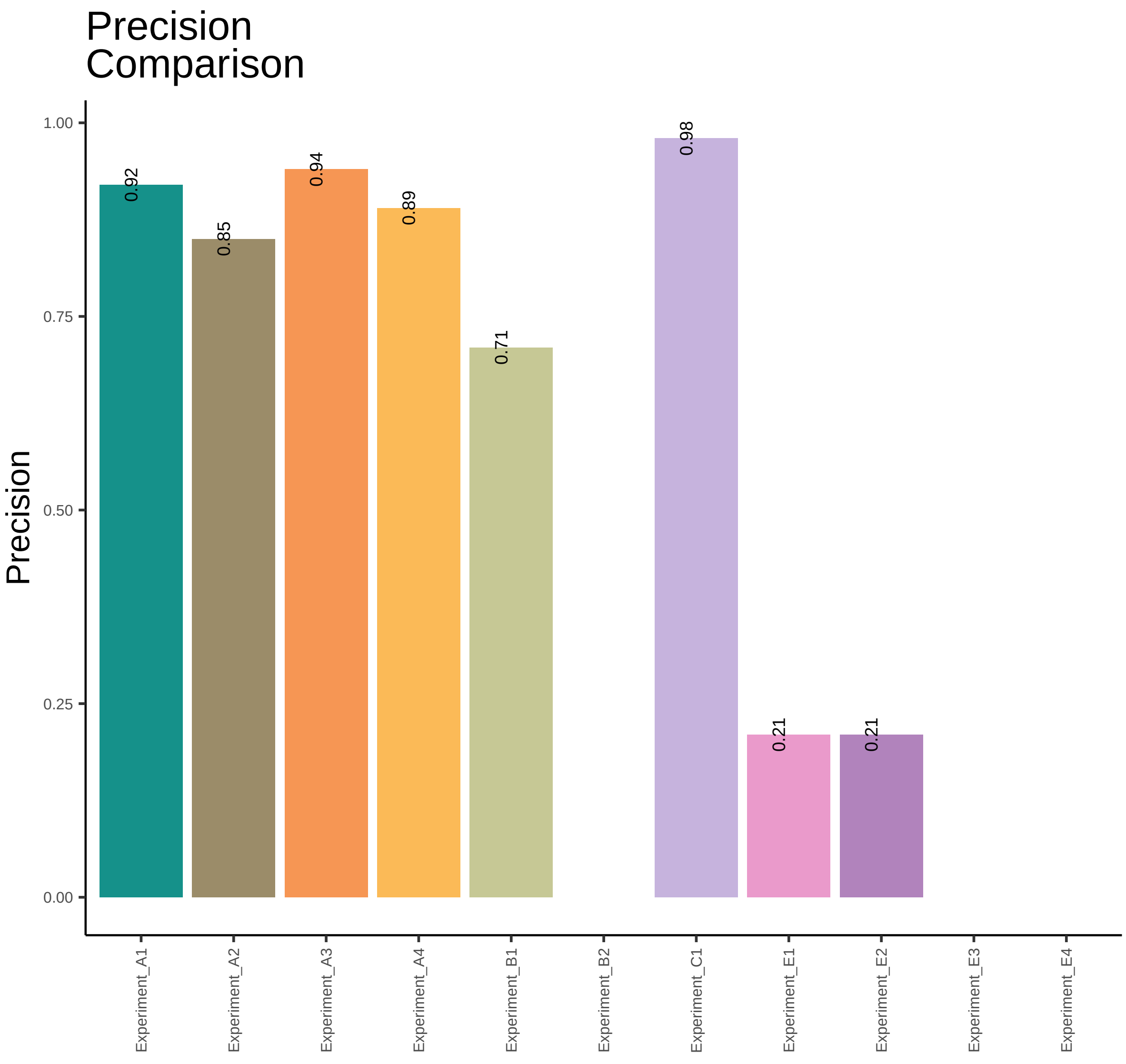
Precision
Comparison
Precision
1.00
0.75
0.50
0.25
0.00
0.92
0.85
0.94
0.89
0.71
0.98
0.21
0.21
Experiment_A1
Experiment_A2
Experiment_A3
Experiment_A4
Experiment_B1
Experiment_B2
Experiment_C1
Experiment_E1
Experiment_E2
Experiment_E3
Experiment_E4

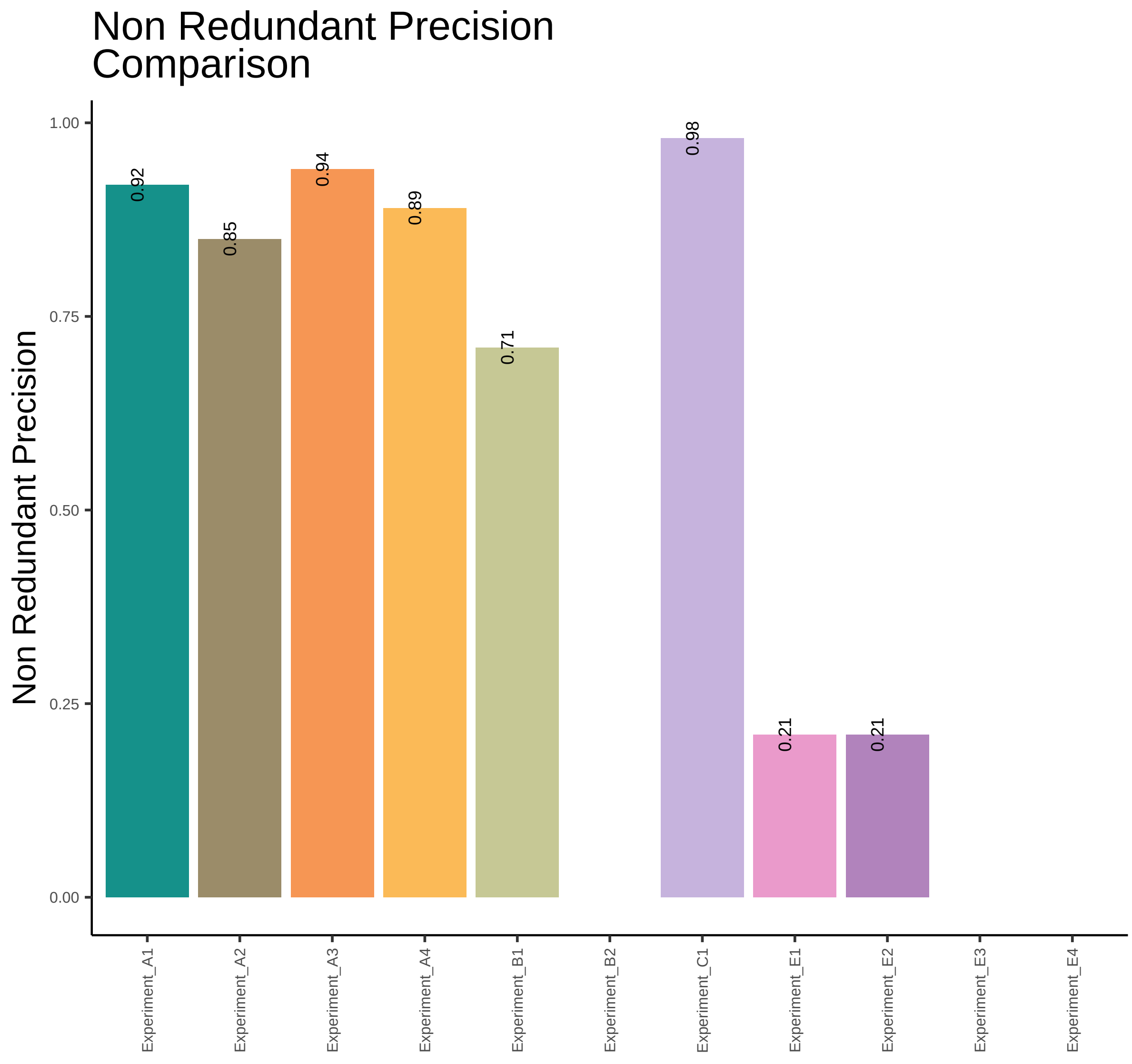

Non Redundant Precision
Comparison
Non Redundant Precision
1.00
0.75
0.50
0.25
0.00
0.92
0.85
0.94
0.89
0.71
0.98
0.21
0.21
Experiment_A1
Experiment_A2
Experiment_A3
Experiment_A4
Experiment_B1
Experiment_B2
Experiment_C1
Experiment_E1
Experiment_E2
Experiment_E3
Experiment_E4

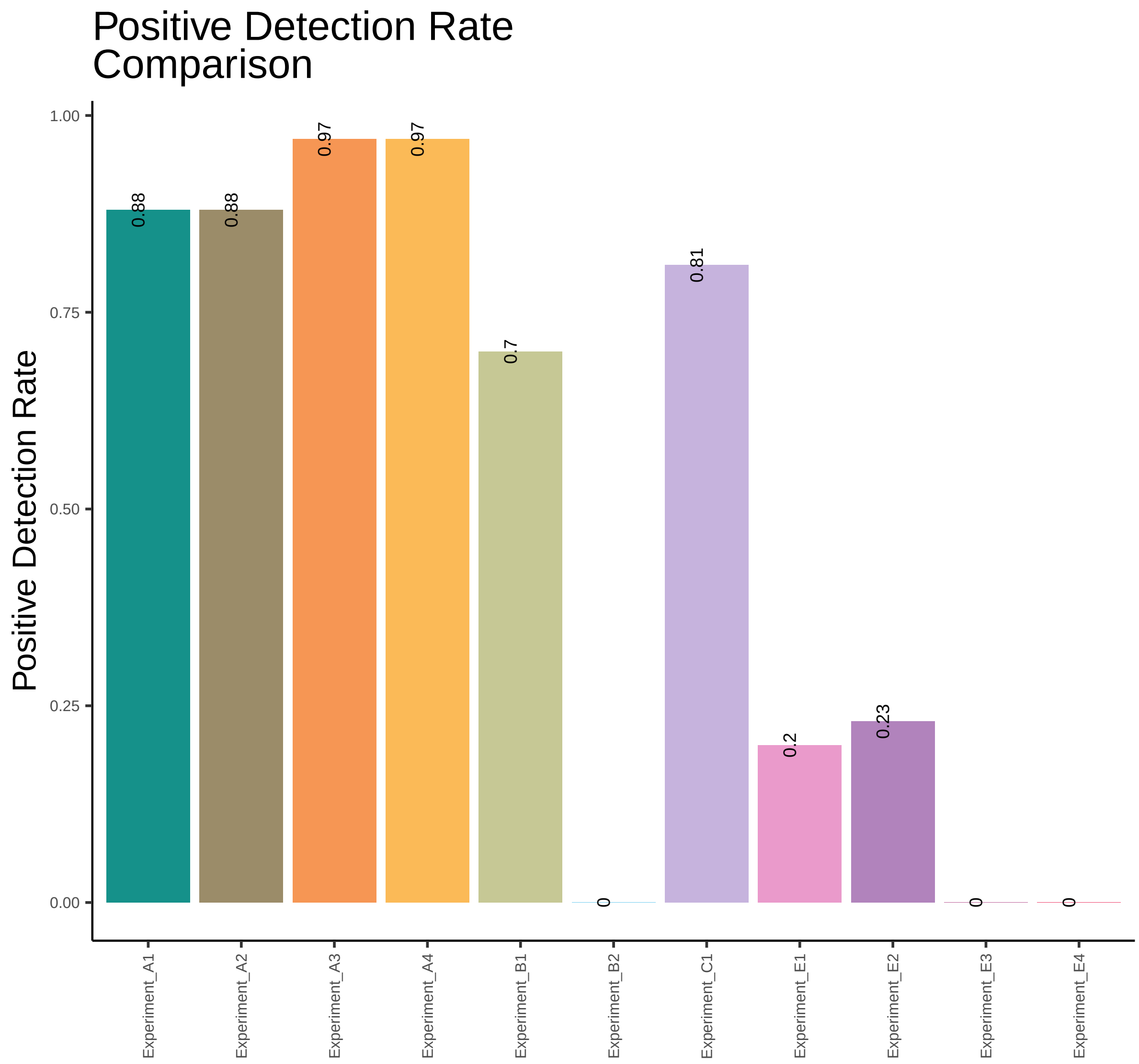

Positive Detection Rate
Comparison
Positive Detection Rate
1.00
0.75
0.50
0.25
0.00
0.88
0.88
0.97
0.97
0.7
0
0.81
0.2
0.23
0
0
Experiment_A1
Experiment_A2
Experiment_A3
Experiment_A4
Experiment_B1
Experiment_B2
Experiment_C1
Experiment_E1
Experiment_E2
Experiment_E3
Experiment_E4

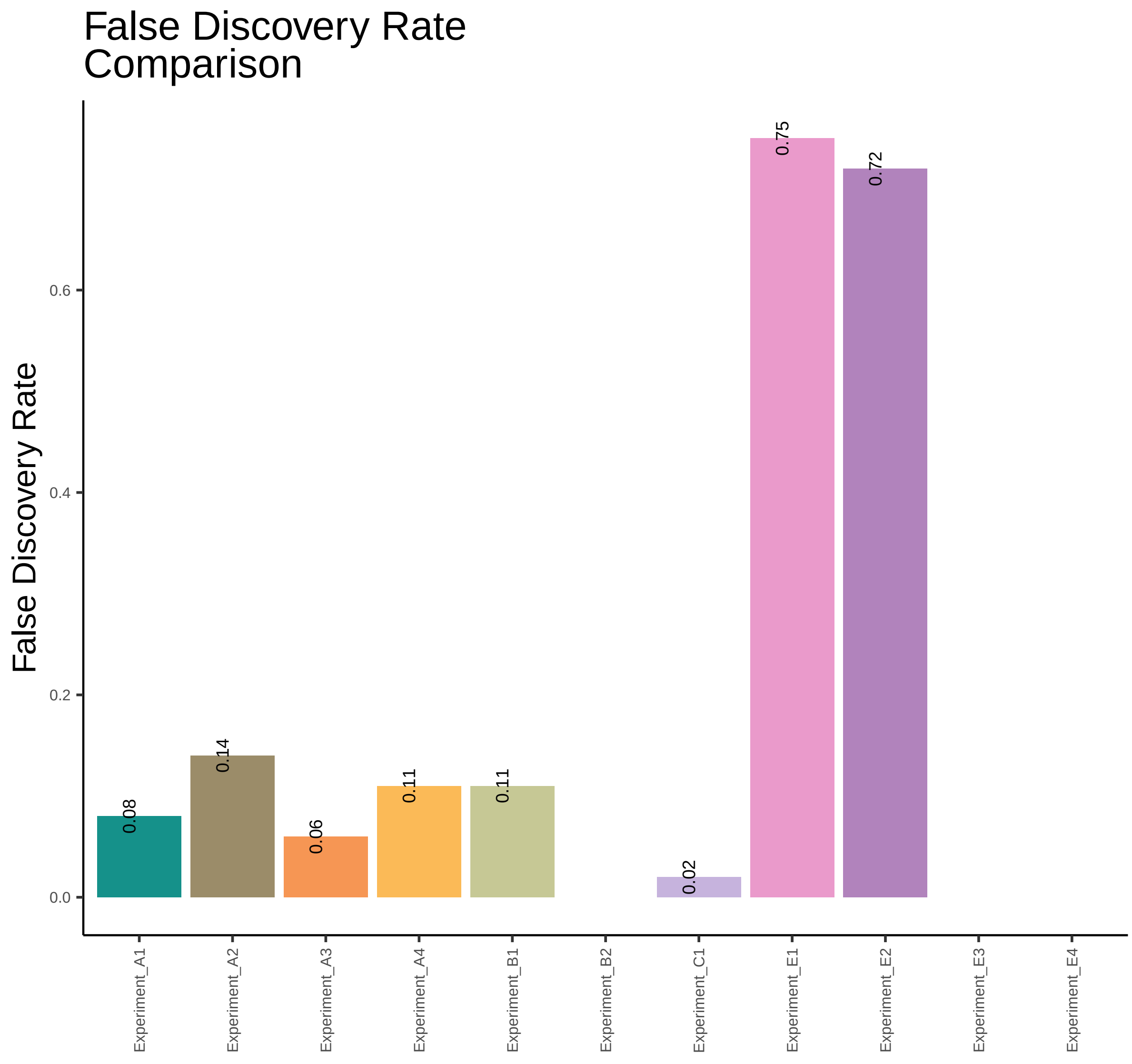
False Discovery Rate
Comparison
False Discovery Rate
0.0
0.2
0.4
0.6
0.08
0.14
0.06
0.11
0.11
0.02
0.75
0.72
Experiment_A1
Experiment_A2
Experiment_A3
Experiment_A4
Experiment_B1
Experiment_B2
Experiment_C1
Experiment_E1
Experiment_E2
Experiment_E3
Experiment_E4

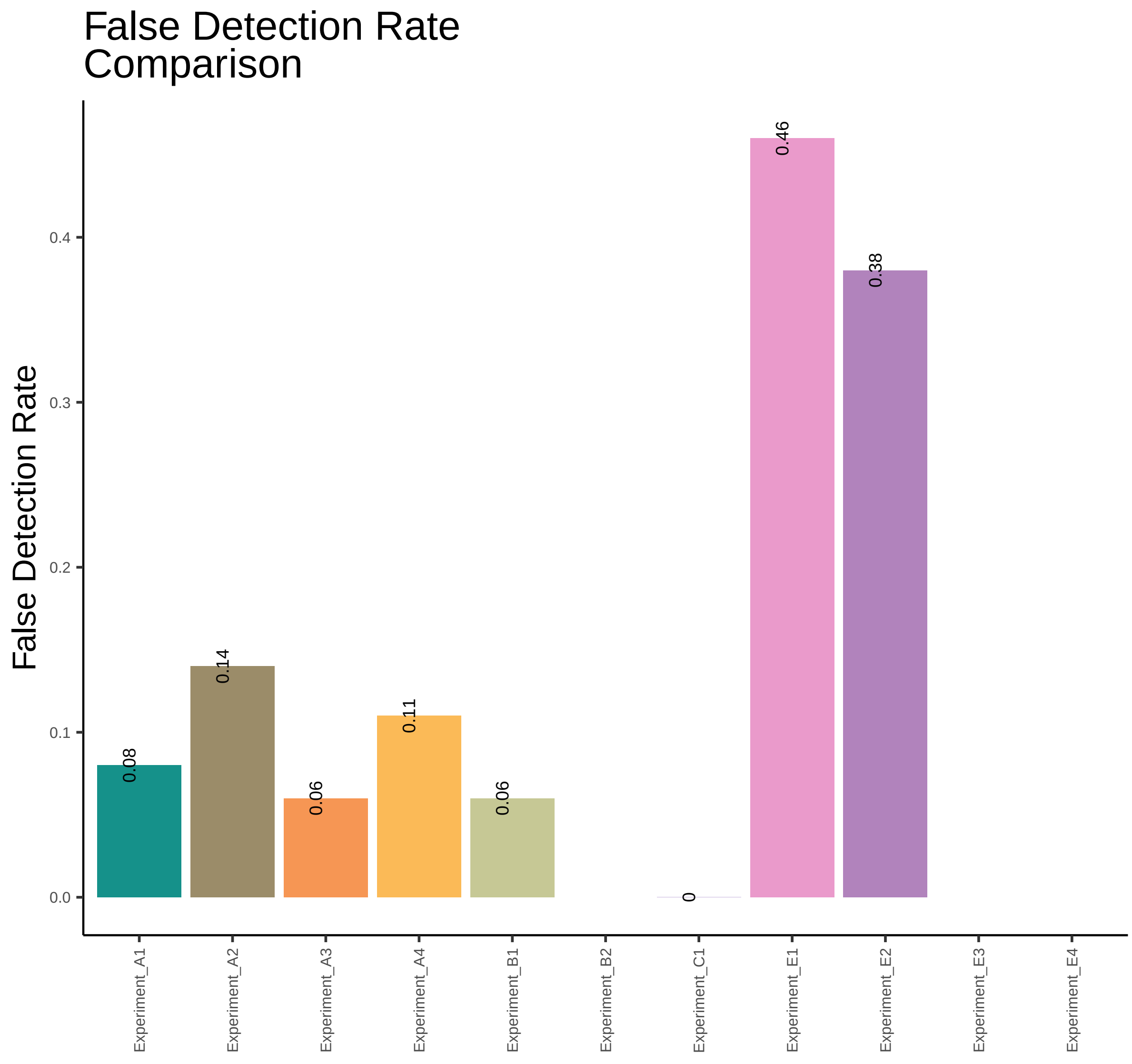

False Detection Rate
Comparison
False Detection Rate
0.0
0.1
0.2
0.3
0.4
0.08
0.14
0.06
0.11
0.06
0
0.46
0.38
Experiment_A1
Experiment_A2
Experiment_A3
Experiment_A4
Experiment_B1
Experiment_B2
Experiment_C1
Experiment_E1
Experiment_E2
Experiment_E3
Experiment_E4

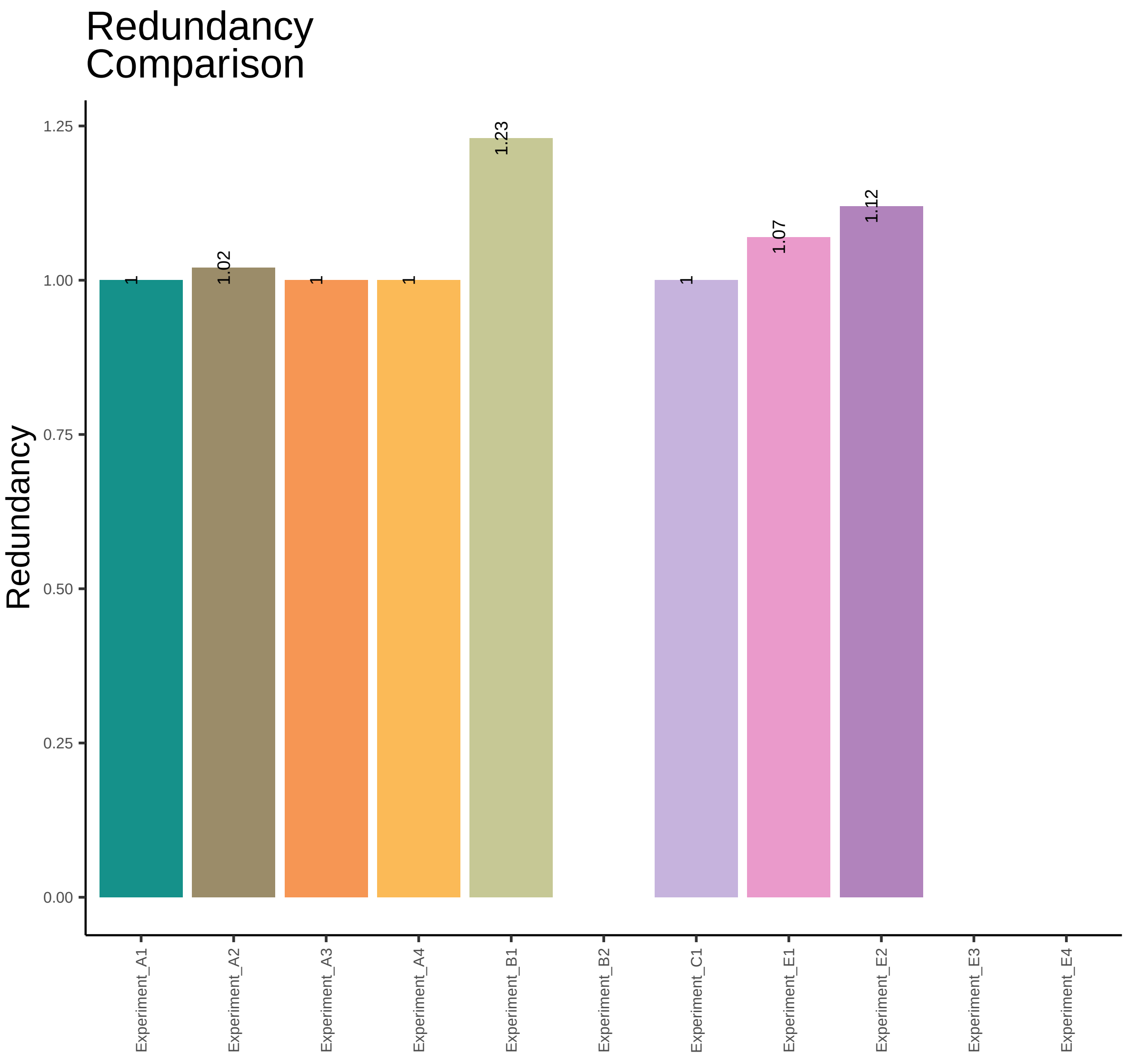
Redundancy
Comparison
Redundancy
1.25
1.00
0.75
0.50
0.25
0.00
1
1.02
1
1
1.23
1
1.07
1.12
Experiment_A1
Experiment_A2
Experiment_A3
Experiment_A4
Experiment_B1
Experiment_B2
Experiment_C1
Experiment_E1
Experiment_E2
Experiment_E3
Experiment_E4

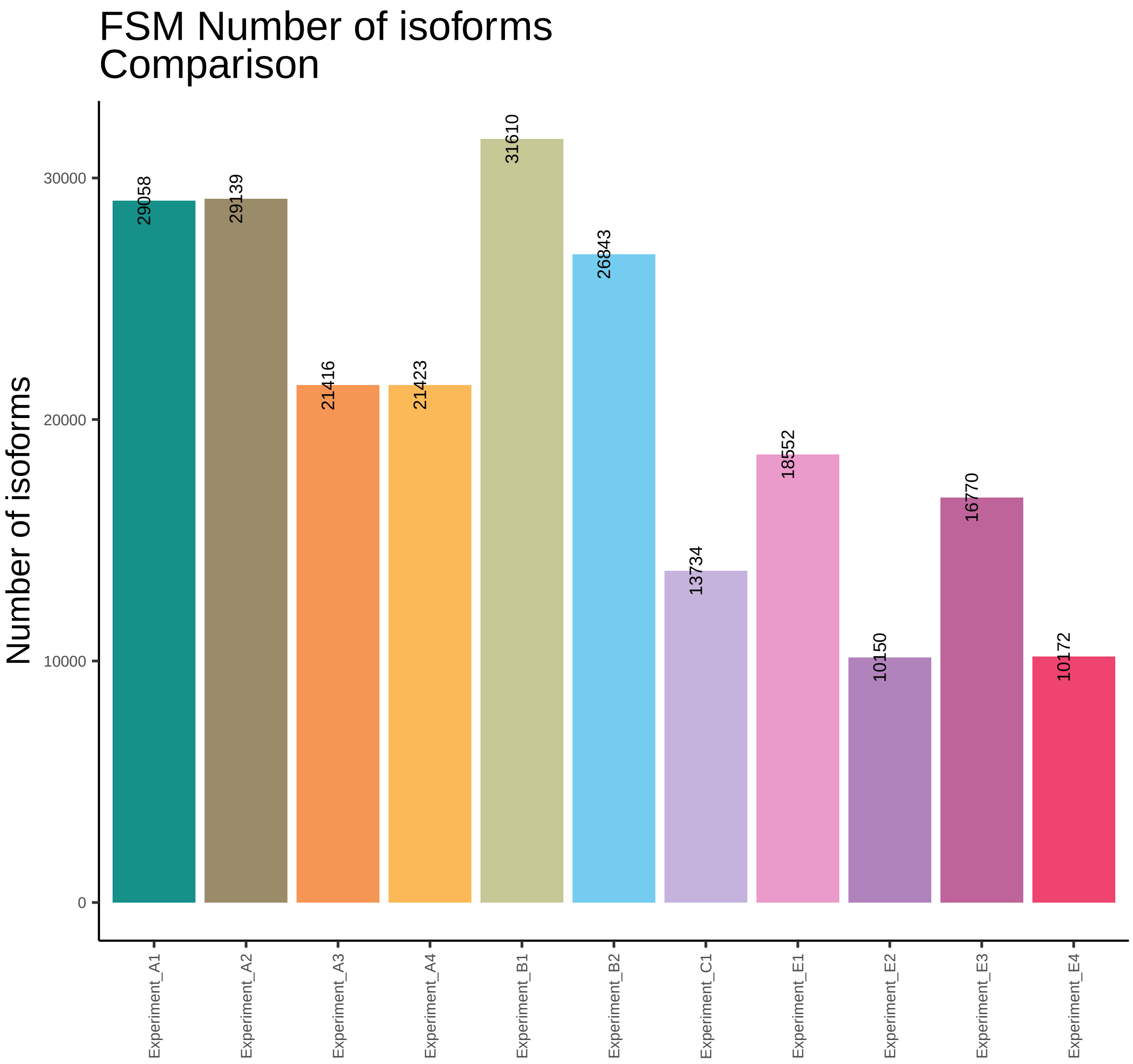
FSM Number of isoforms
Comparison
Number of isoforms
0
10000
20000
30000
29058
29139
21416
21423
31610
26843
13734
18552
10150
16770
10172
Experiment_A1
Experiment_A2
Experiment_A3
Experiment_A4
Experiment_B1
Experiment_B2
Experiment_C1
Experiment_E1
Experiment_E2
Experiment_E3
Experiment_E4

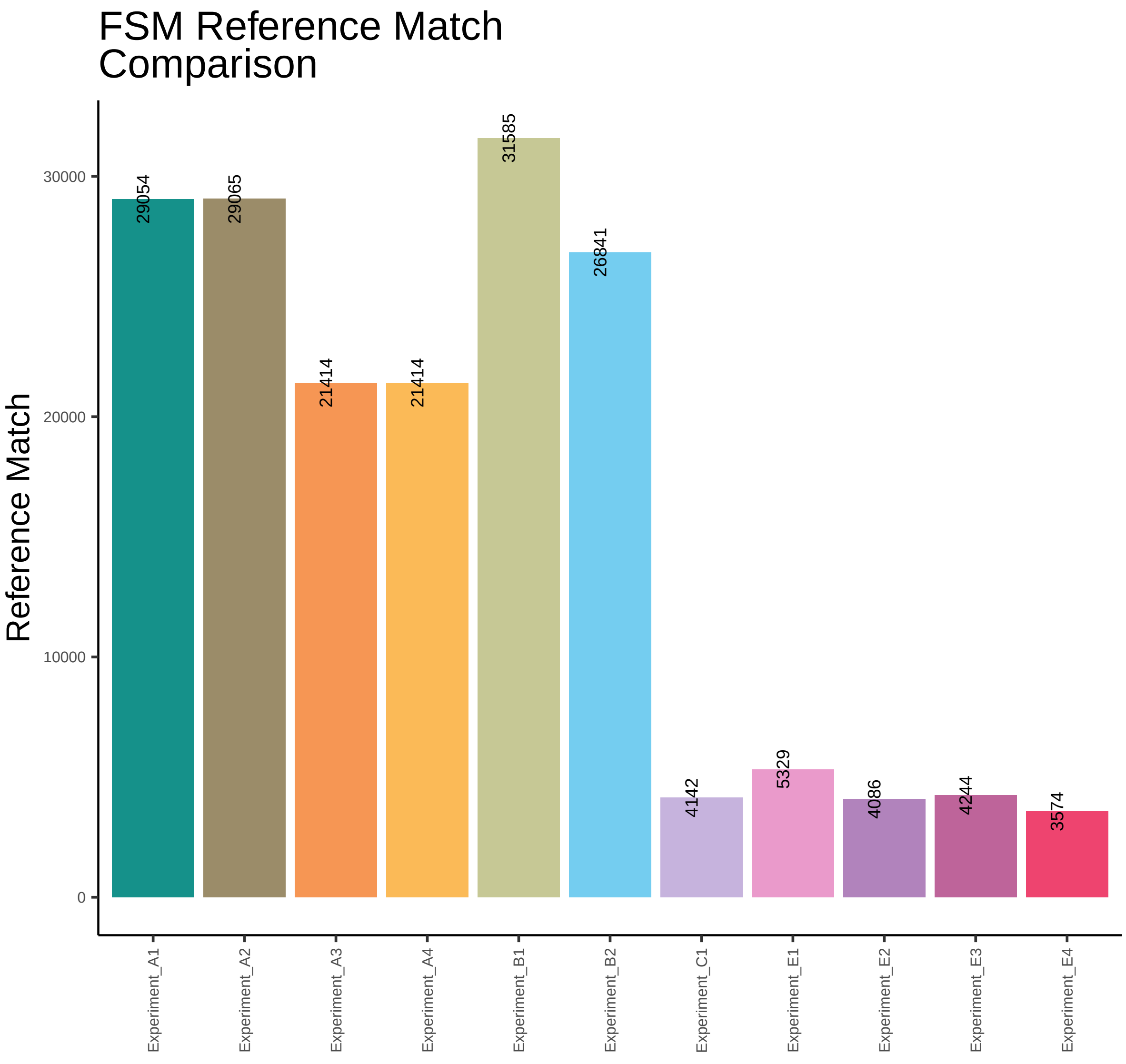
FSM Reference Match Comparison
Reference Match
30000
20000
10000
0
29054
29065
21414
21414
31585
26841
4142
5329
4086
4244
3574
Experiment_A1
Experiment_A2
Experiment_A3
Experiment_A4
Experiment_B1
Experiment_B2
Experiment_C1
Experiment_E1
Experiment_E2
Experiment_E3
Experiment_E4

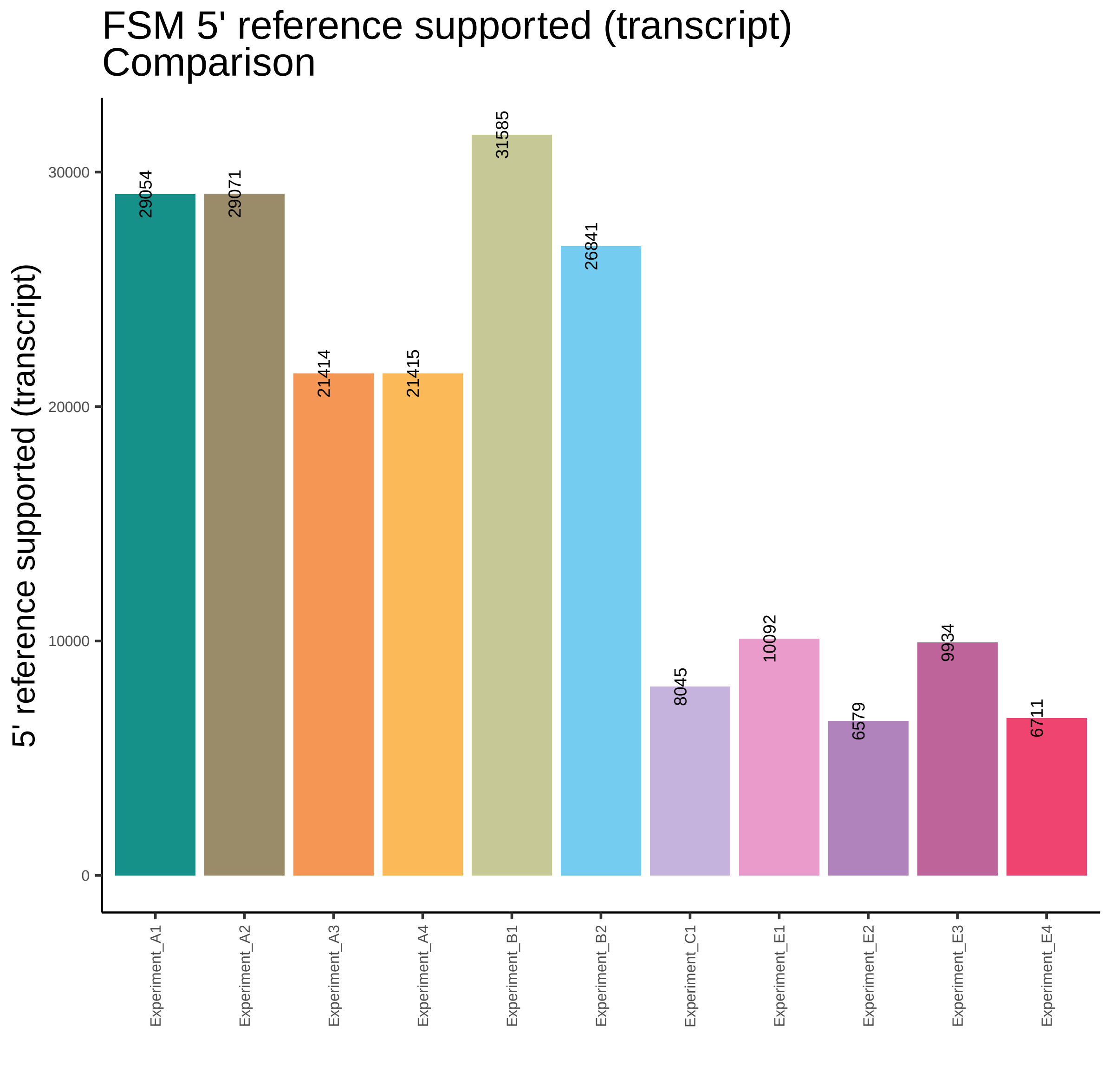
FSM 5' reference supported (transcript)
Comparison
5' reference supported (transcript)
0
10000
20000
30000
29054
29071
21414
21415
31585
26841
8045
10092
6579
9934
6711
Experiment_A1
Experiment_A2
Experiment_A3
Experiment_A4
Experiment_B1
Experiment_B2
Experiment_C1
Experiment_E1
Experiment_E2
Experiment_E3
Experiment_E4

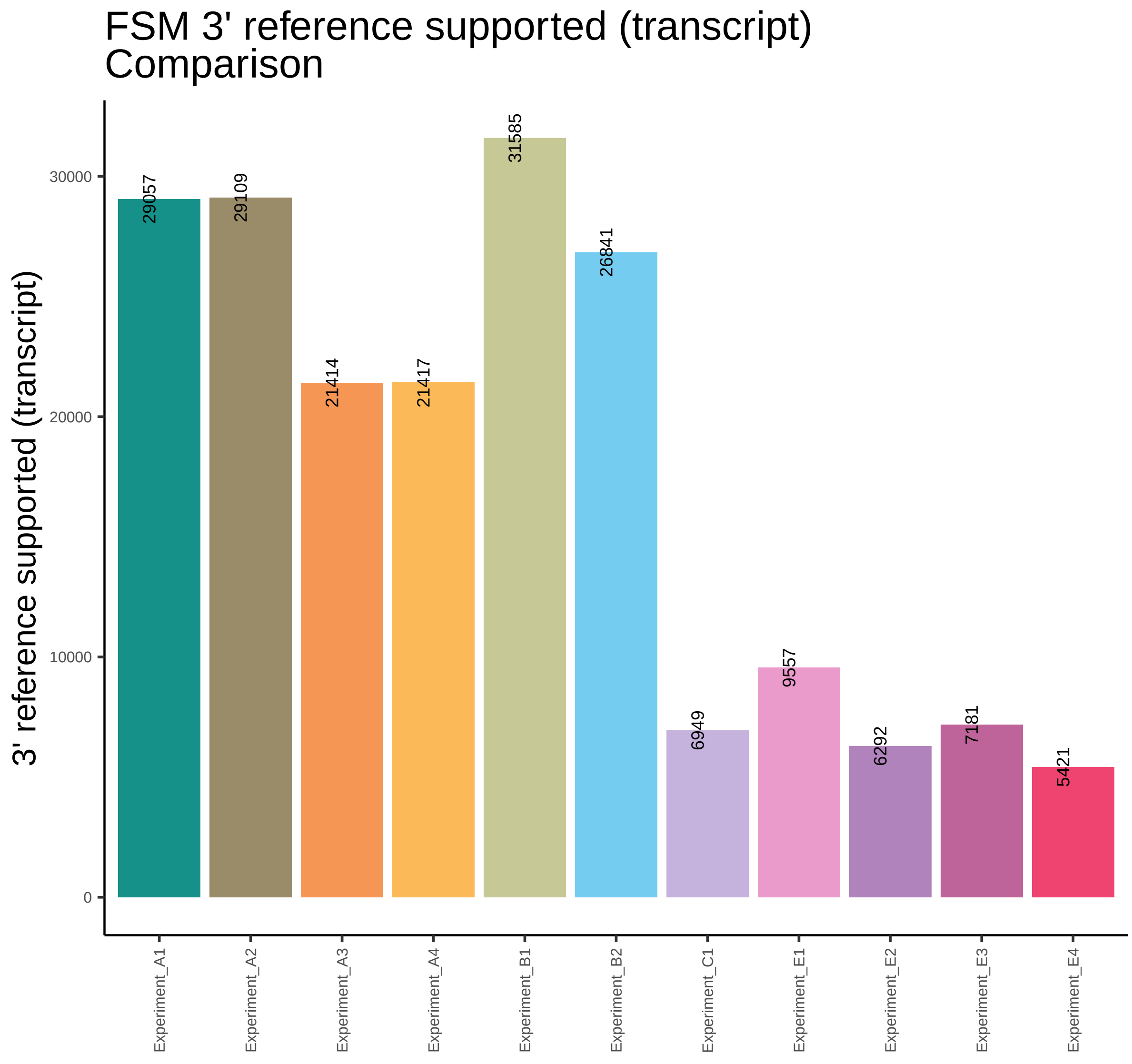
FSM 3' reference supported (transcript)
Comparison
3' reference supported (transcript)
0
10000
20000
30000
29057
29109
21414
21417
31585
26841
6949
9557
6292
7181
5421
Experiment_A1
Experiment_A2
Experiment_A3
Experiment_A4
Experiment_B1
Experiment_B2
Experiment_C1
Experiment_E1
Experiment_E2
Experiment_E3
Experiment_E4

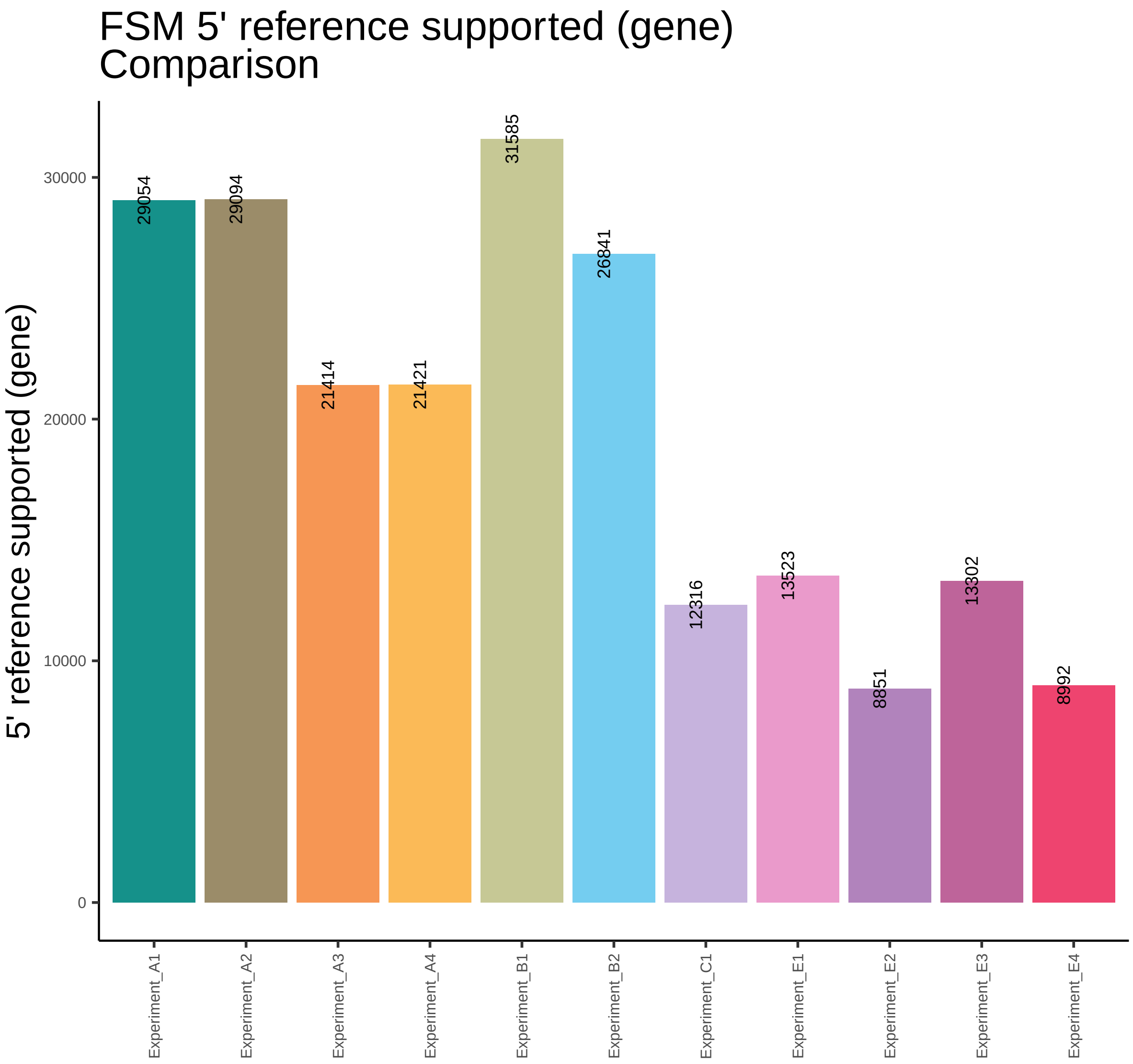
FSM 5' reference supported (gene)
Comparison
5' reference supported (gene)
0
10000
20000
30000
29054
29094
21414
21421
31585
26841
12316
13523
8851
13302
8992
Experiment_A1
Experiment_A2
Experiment_A3
Experiment_A4
Experiment_B1
Experiment_B2
Experiment_C1
Experiment_E1
Experiment_E2
Experiment_E3
Experiment_E4

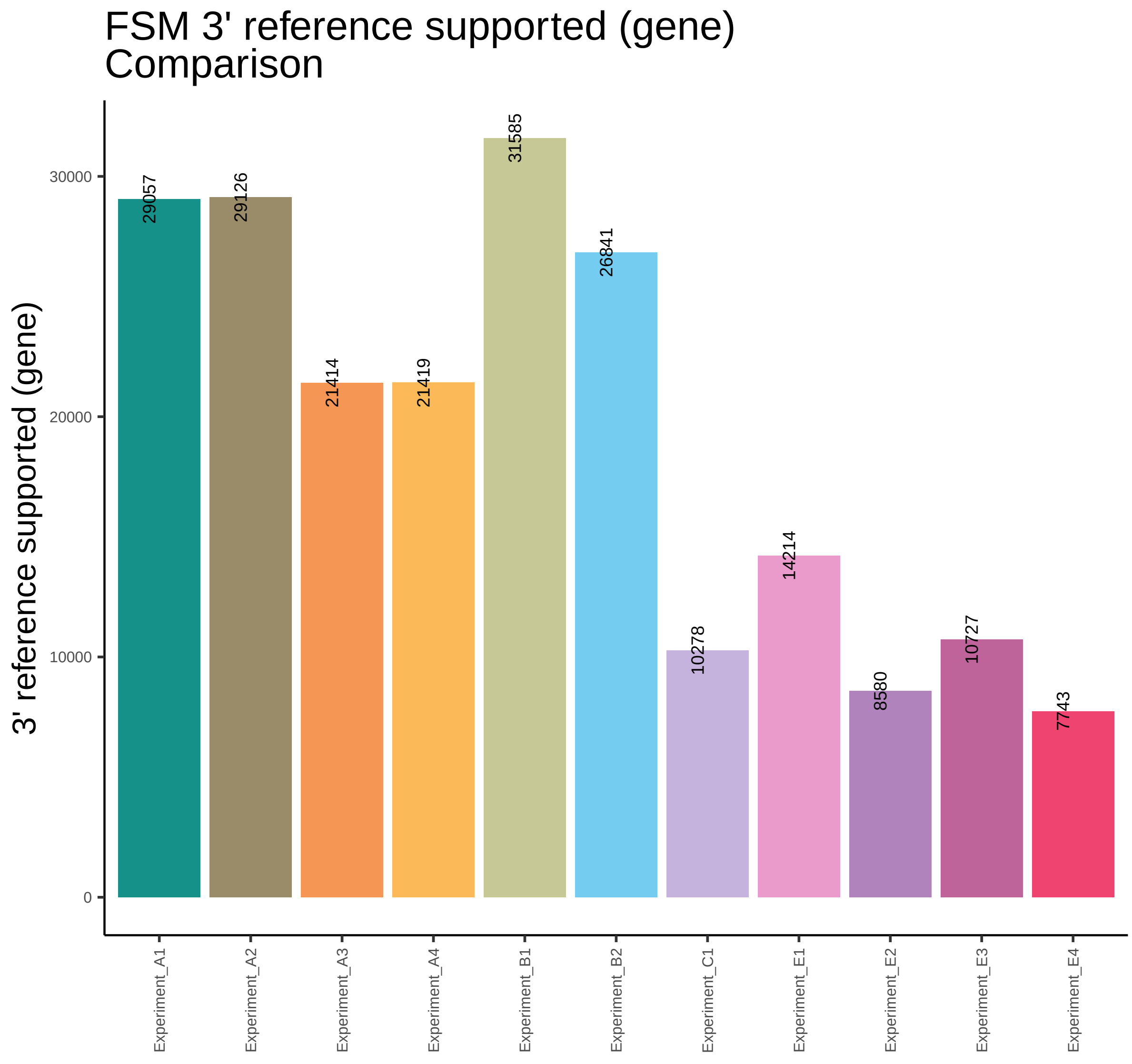
FSM 3' reference supported (gene)
Comparison
3' reference supported (gene)
0
10000
20000
30000
29057
29126
21414
21419
31585
26841
10278
14214
8580
10727
7743
Experiment_A1
Experiment_A2
Experiment_A3
Experiment_A4
Experiment_B1
Experiment_B2
Experiment_C1
Experiment_E1
Experiment_E2
Experiment_E3
Experiment_E4

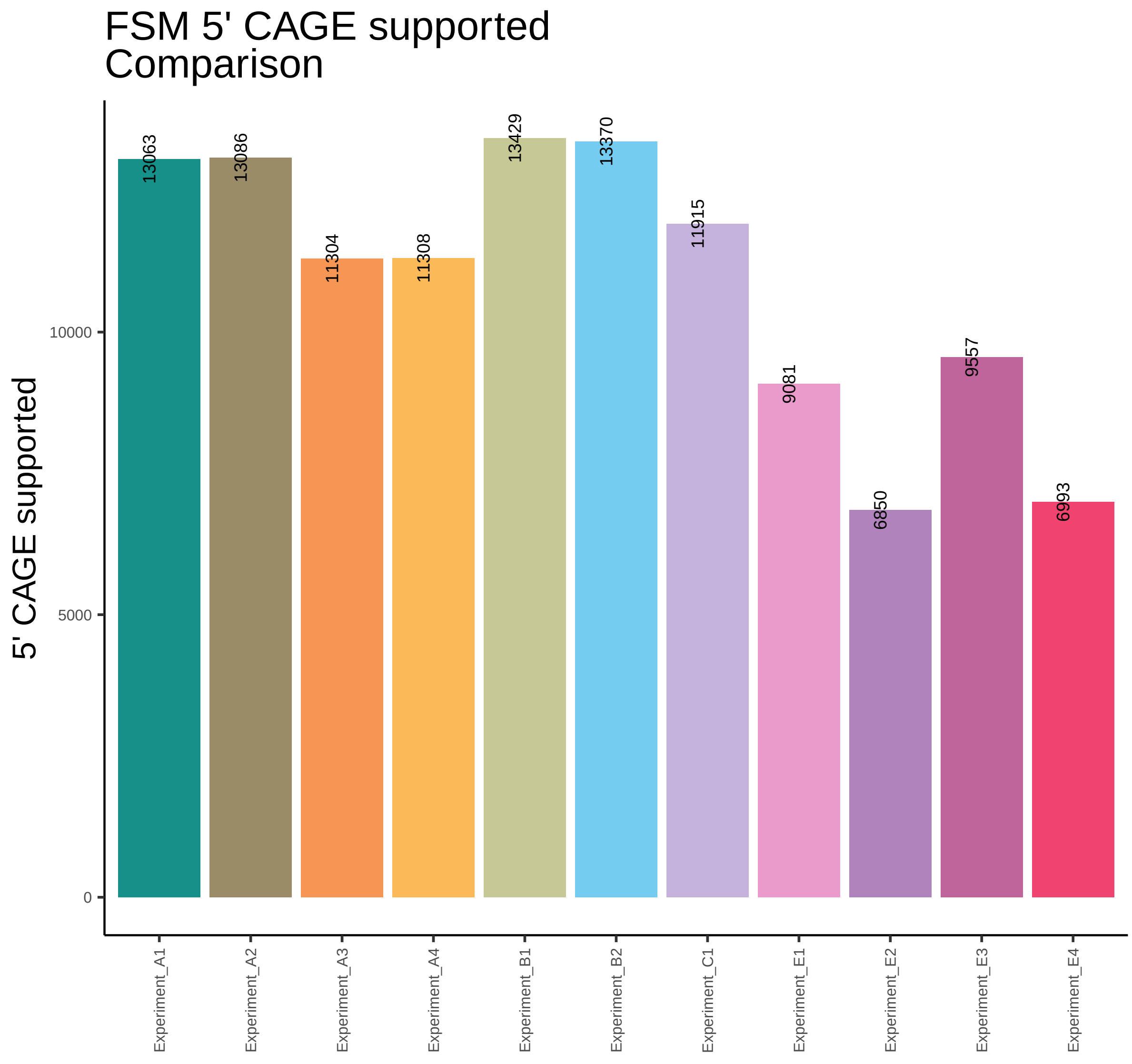

FSM 5' CAGE supported
Comparison
5' CAGE supported
0
5000
10000
13063
13086
11304
11308
13429
13370
11915
9081
6850
9557
6993
Experiment_A1
Experiment_A2
Experiment_A3
Experiment_A4
Experiment_B1
Experiment_B2
Experiment_C1
Experiment_E1
Experiment_E2
Experiment_E3
Experiment_E4

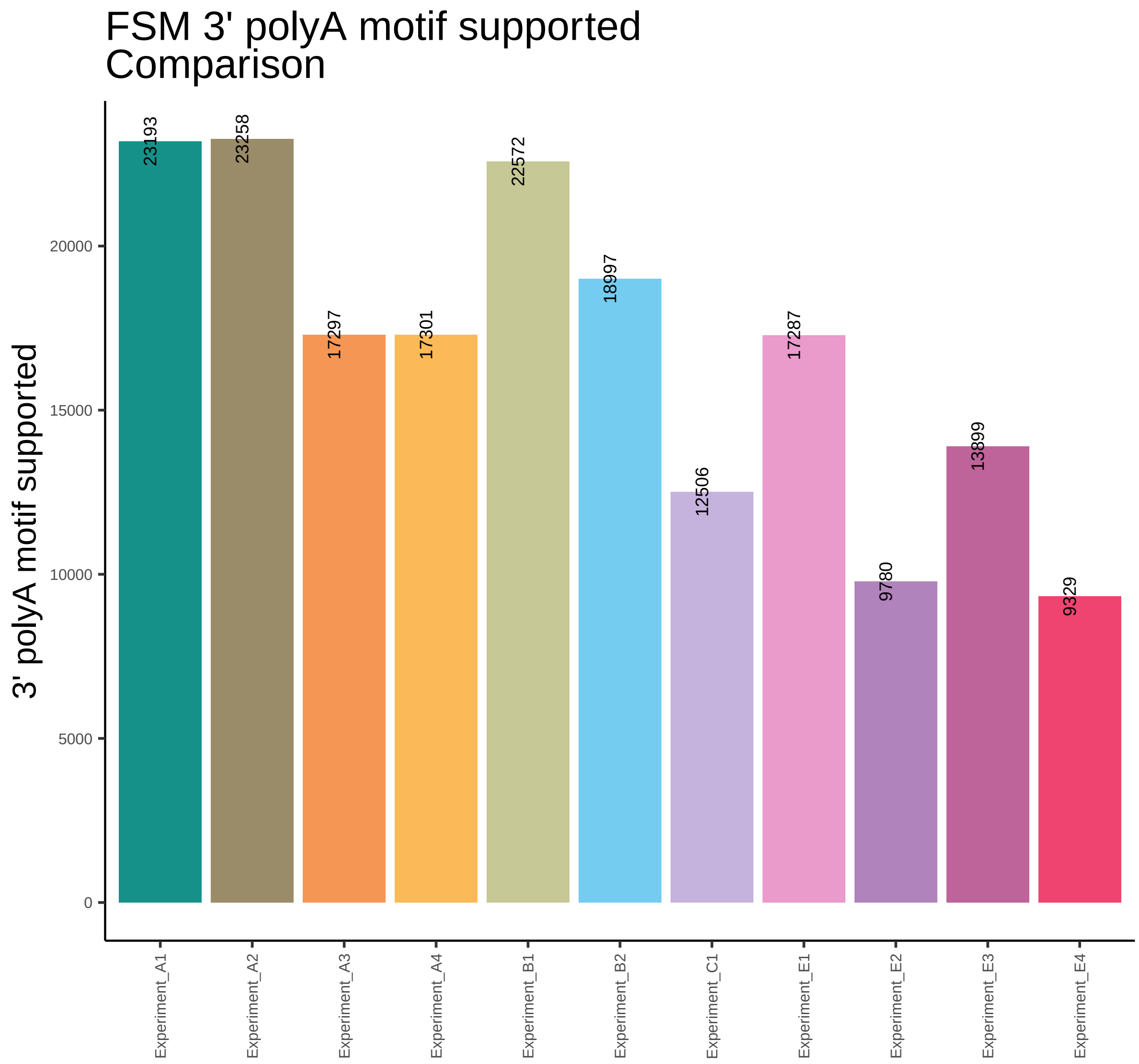
FSM 3' polyA motif supported
Comparison
3' polyA motif supported
0
5000
10000
15000
20000
23193
23258
17297
17301
22572
18997
12506
17287
9780
13899
9329
Experiment_A1
Experiment_A2
Experiment_A3
Experiment_A4
Experiment_B1
Experiment_B2
Experiment_C1
Experiment_E1
Experiment_E2
Experiment_E3
Experiment_E4

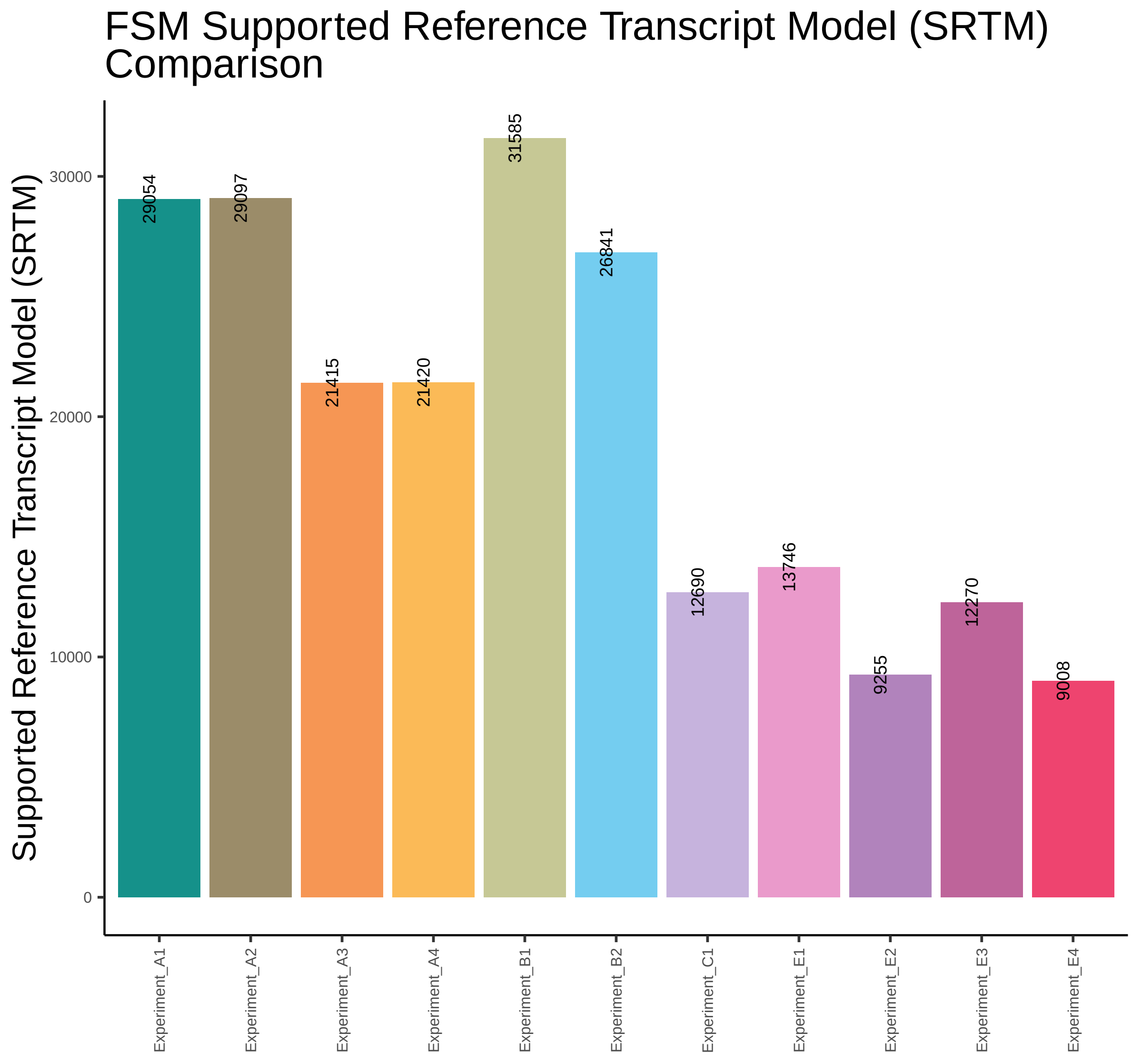
FSM Supported Reference Transcript Model (SRTM) Comparison
Supported Reference Transcript Model (SRTM)
30000
20000
10000
0
29054
29097
21415
21420
31585
26841
12690
13746
9255
12270
9008
Experiment_A1
Experiment_A2
Experiment_A3
Experiment_A4
Experiment_B1
Experiment_B2
Experiment_C1
Experiment_E1
Experiment_E2
Experiment_E3
Experiment_E4

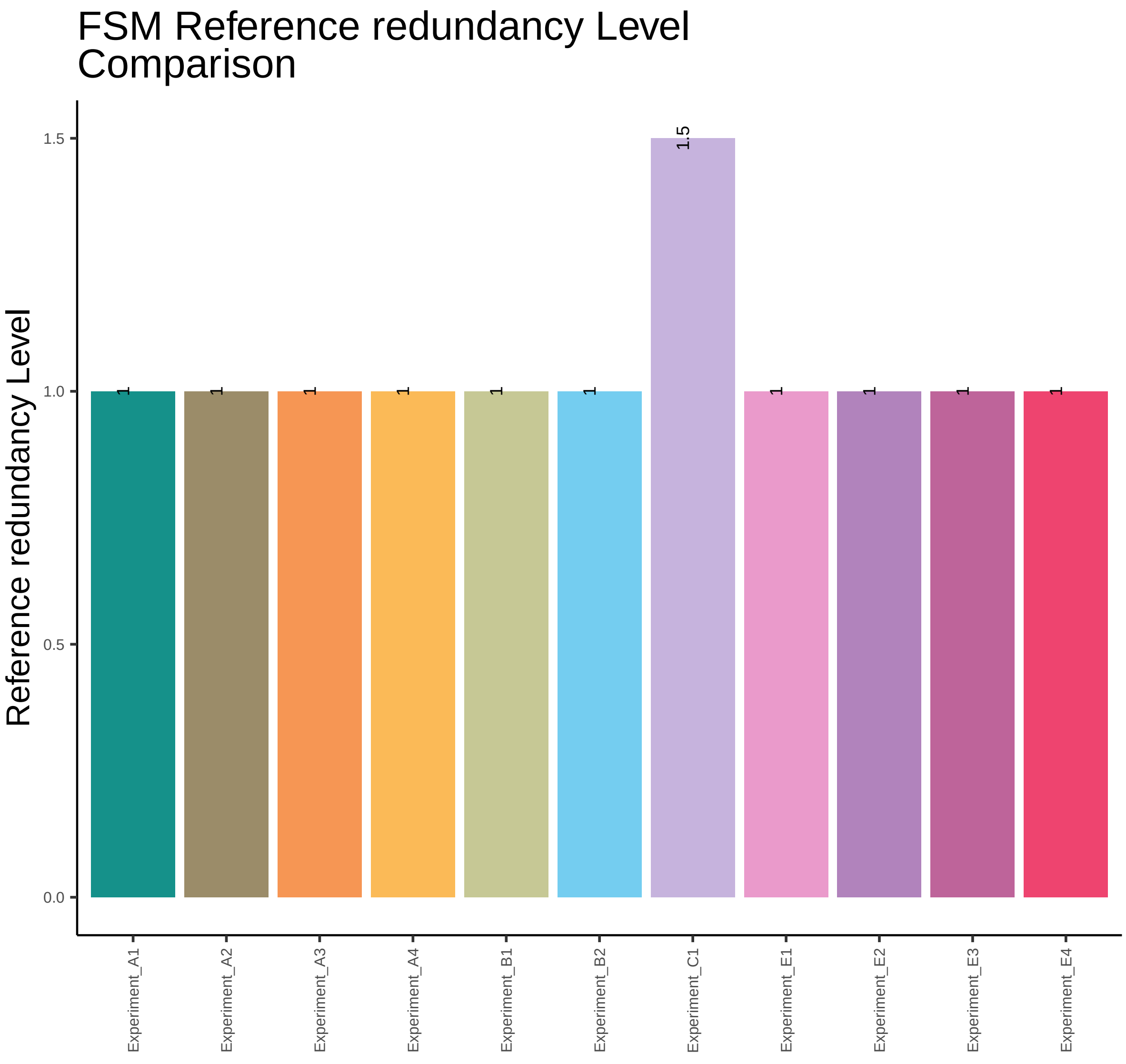
FSM Reference redundancy Level
Comparison
Reference redundancy Level
1.5
1
1
1
1
1
1
1
1
1
1
0.0
0.5
1.0
1.5
Experiment_A1
Experiment_A2
Experiment_A3
Experiment_A4
Experiment_B1
Experiment_B2
Experiment_C1
Experiment_E1
Experiment_E2
Experiment_E3
Experiment_E4

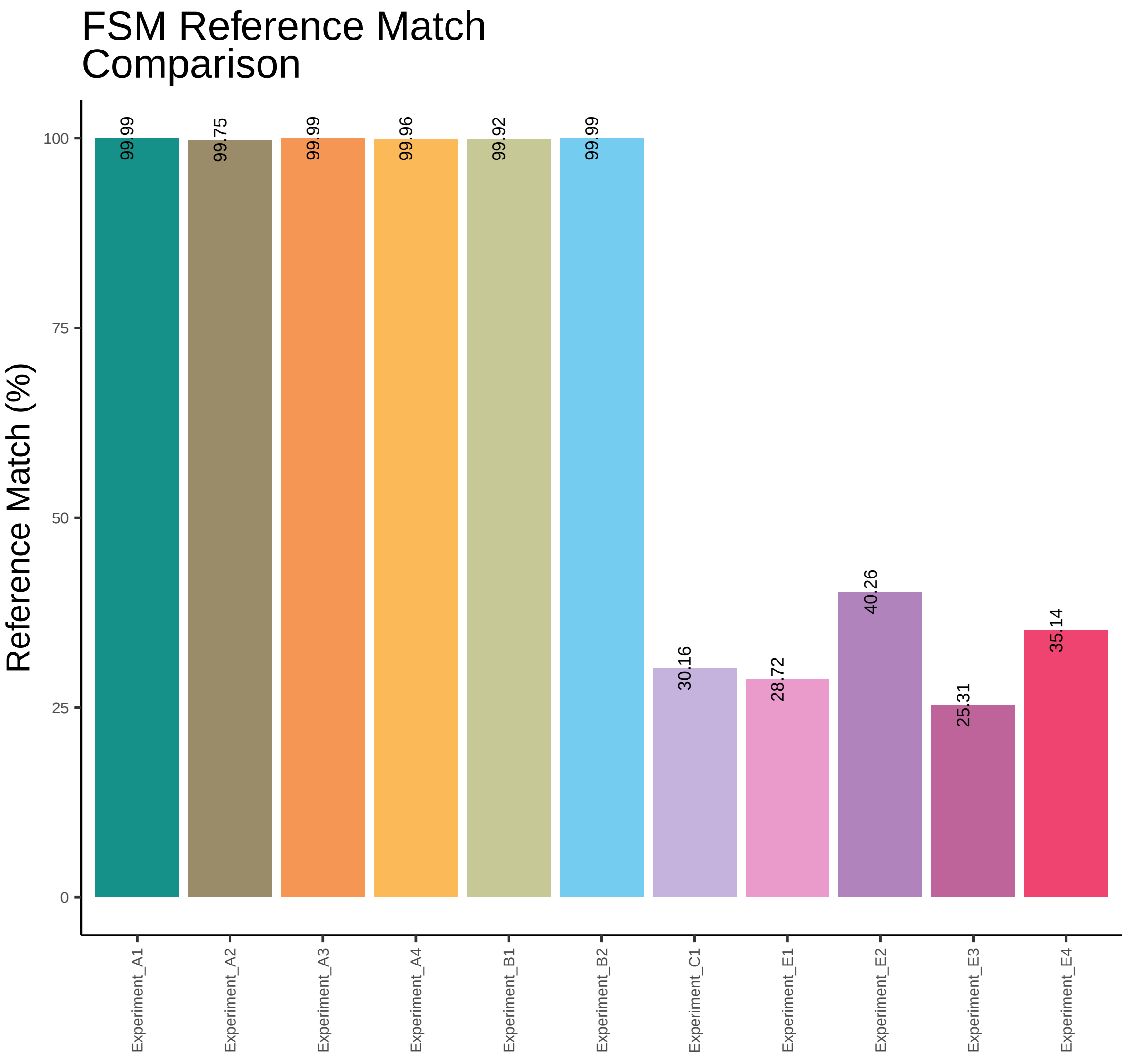
FSM Reference Match
Comparison
Reference Match (%)
100
75
50
25
0
99.99
99.75
99.99
99.96
99.92
99.99
30.16
28.72
40.26
25.31
35.14
Experiment_A1
Experiment_A2
Experiment_A3
Experiment_A4
Experiment_B1
Experiment_B2
Experiment_C1
Experiment_E1
Experiment_E2
Experiment_E3
Experiment_E4

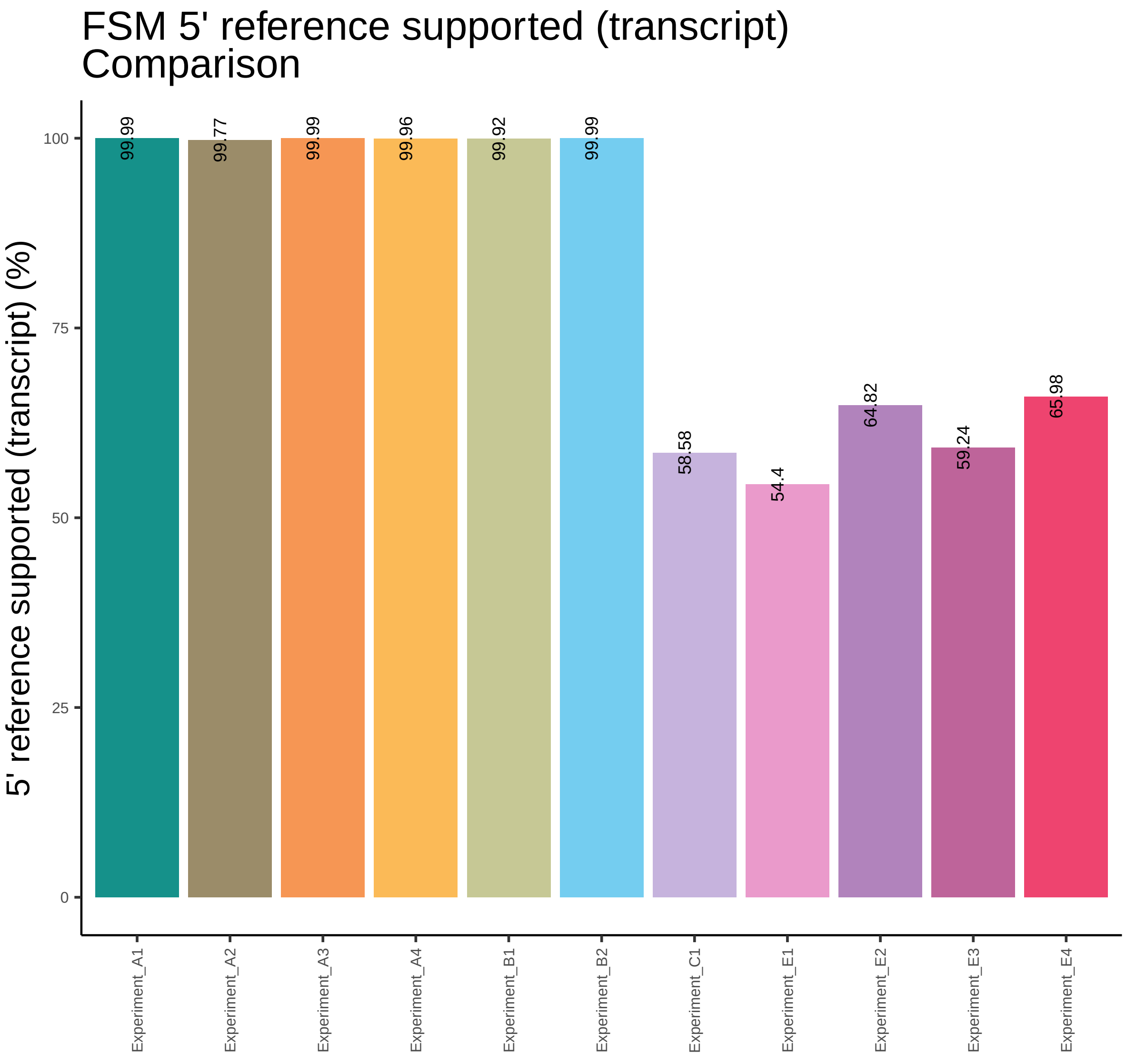

FSM 5' reference supported (transcript)
Comparison
5' reference supported (transcript) (%)
100
75
50
25
0
99.99
99.77
99.99
99.96
99.92
99.99
58.58
54.4
64.82
59.24
65.98
Experiment_A1
Experiment_A2
Experiment_A3
Experiment_A4
Experiment_B1
Experiment_B2
Experiment_C1
Experiment_E1
Experiment_E2
Experiment_E3
Experiment_E4

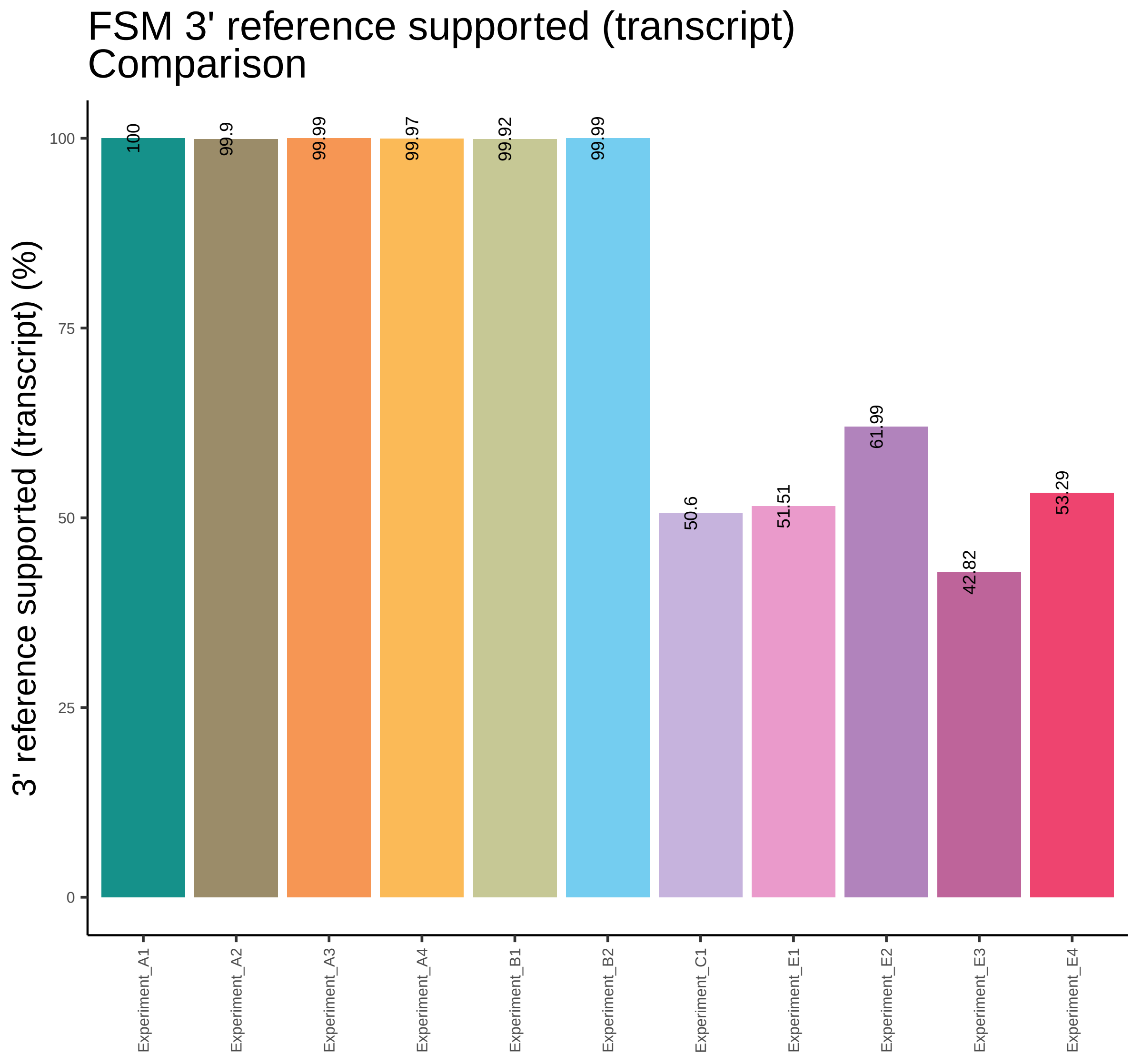

FSM 3' reference supported (transcript)
Comparison
3' reference supported (transcript) (%)
0
25
50
75
100
100
99.9
99.99
99.97
99.92
99.99
50.6
51.51
61.99
42.82
53.29
Experiment_A1
Experiment_A2
Experiment_A3
Experiment_A4
Experiment_B1
Experiment_B2
Experiment_C1
Experiment_E1
Experiment_E2
Experiment_E3
Experiment_E4

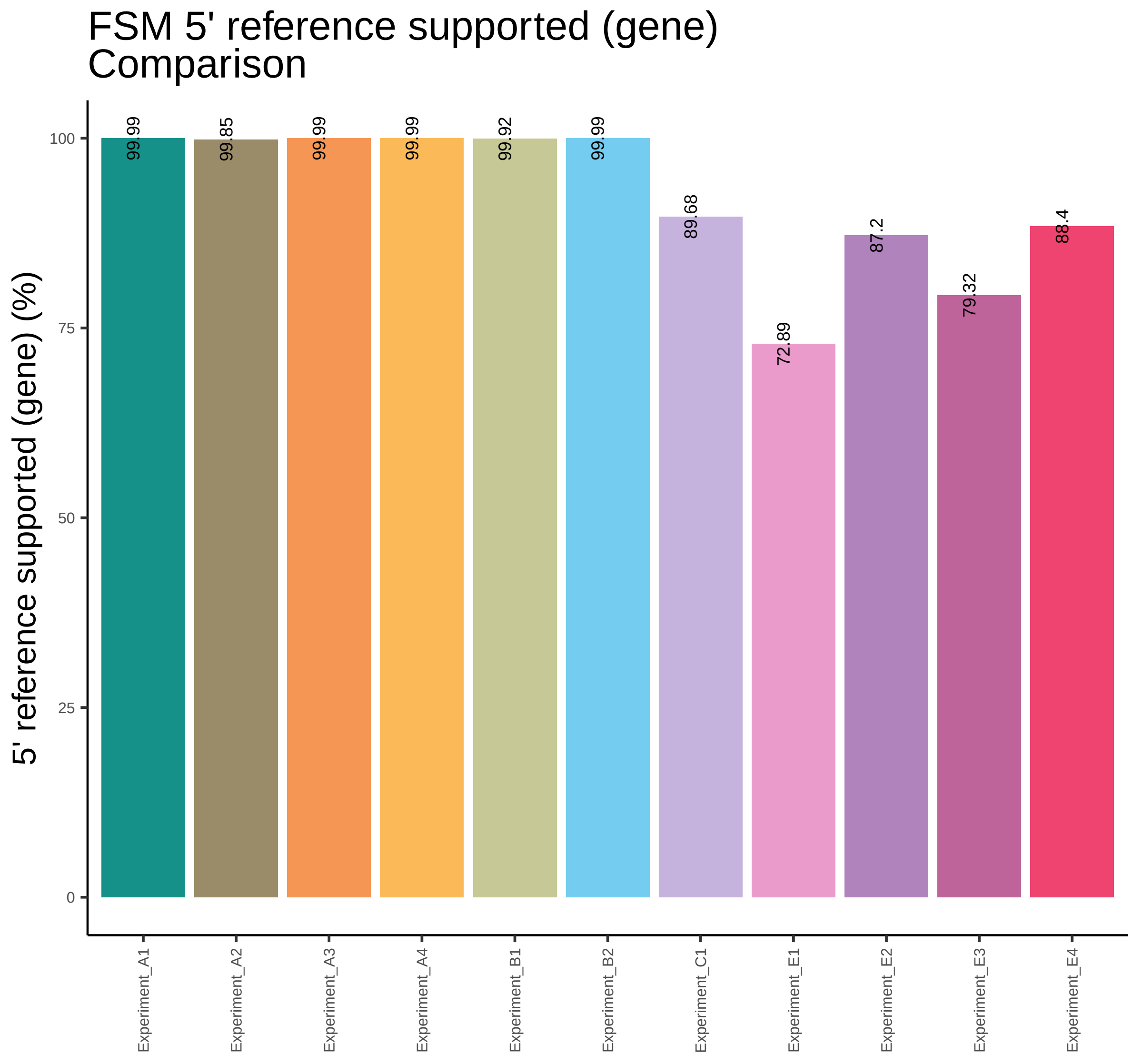

FSM 5' reference supported (gene)
Comparison
5' reference supported (gene) (%)
0
25
50
75
100
99.99
99.85
99.99
99.99
99.92
99.99
89.68
72.89
87.2
79.32
88.4
Experiment_A1
Experiment_A2
Experiment_A3
Experiment_A4
Experiment_B1
Experiment_B2
Experiment_C1
Experiment_E1
Experiment_E2
Experiment_E3
Experiment_E4

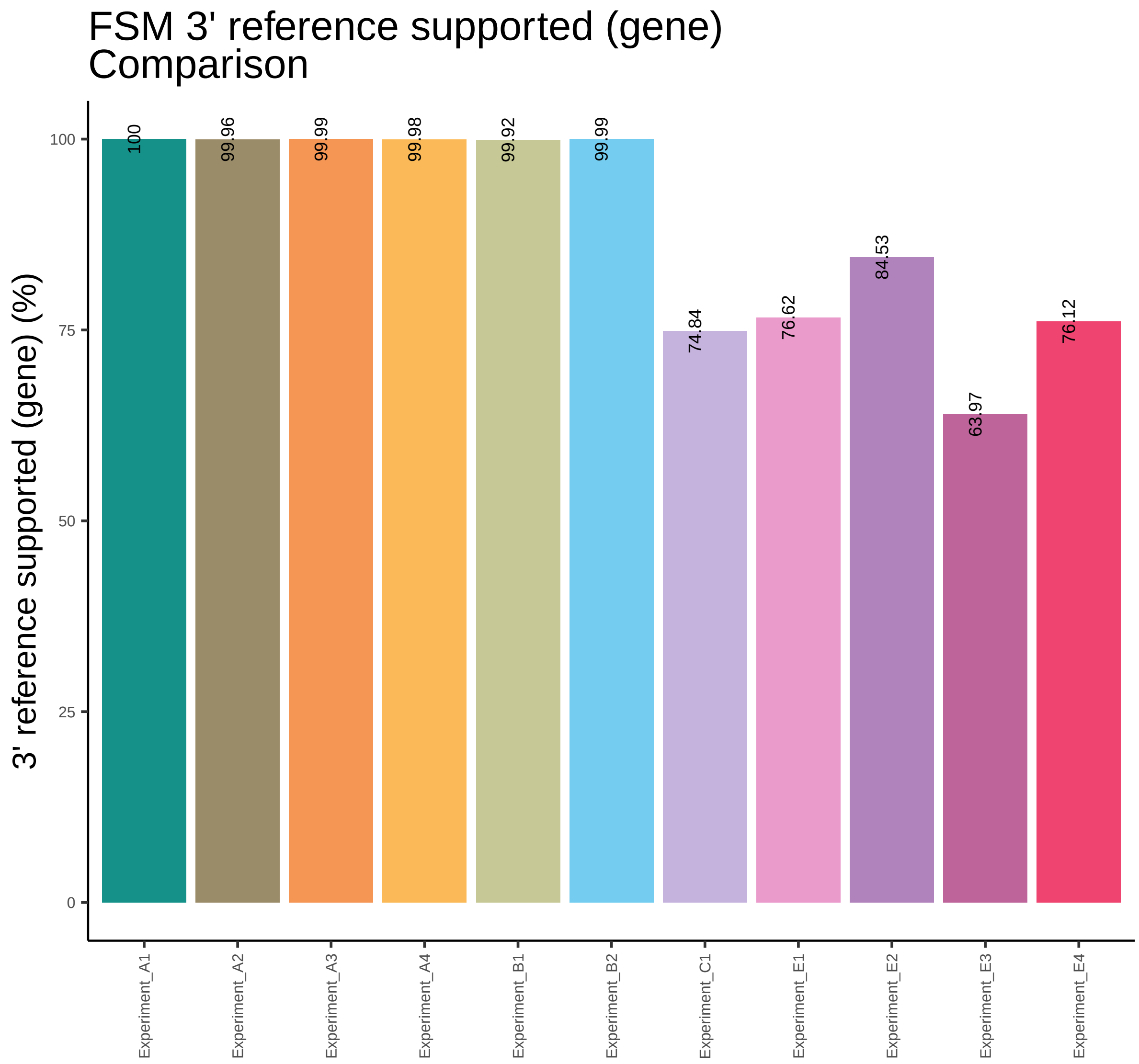

FSM 3' reference supported (gene)
Comparison
3' reference supported (gene) (%)
100
75
50
25
0
100
99.96
99.99
99.98
99.92
99.99
74.84
76.62
84.53
63.97
76.12
Experiment_A1
Experiment_A2
Experiment_A3
Experiment_A4
Experiment_B1
Experiment_B2
Experiment_C1
Experiment_E1
Experiment_E2
Experiment_E3
Experiment_E4

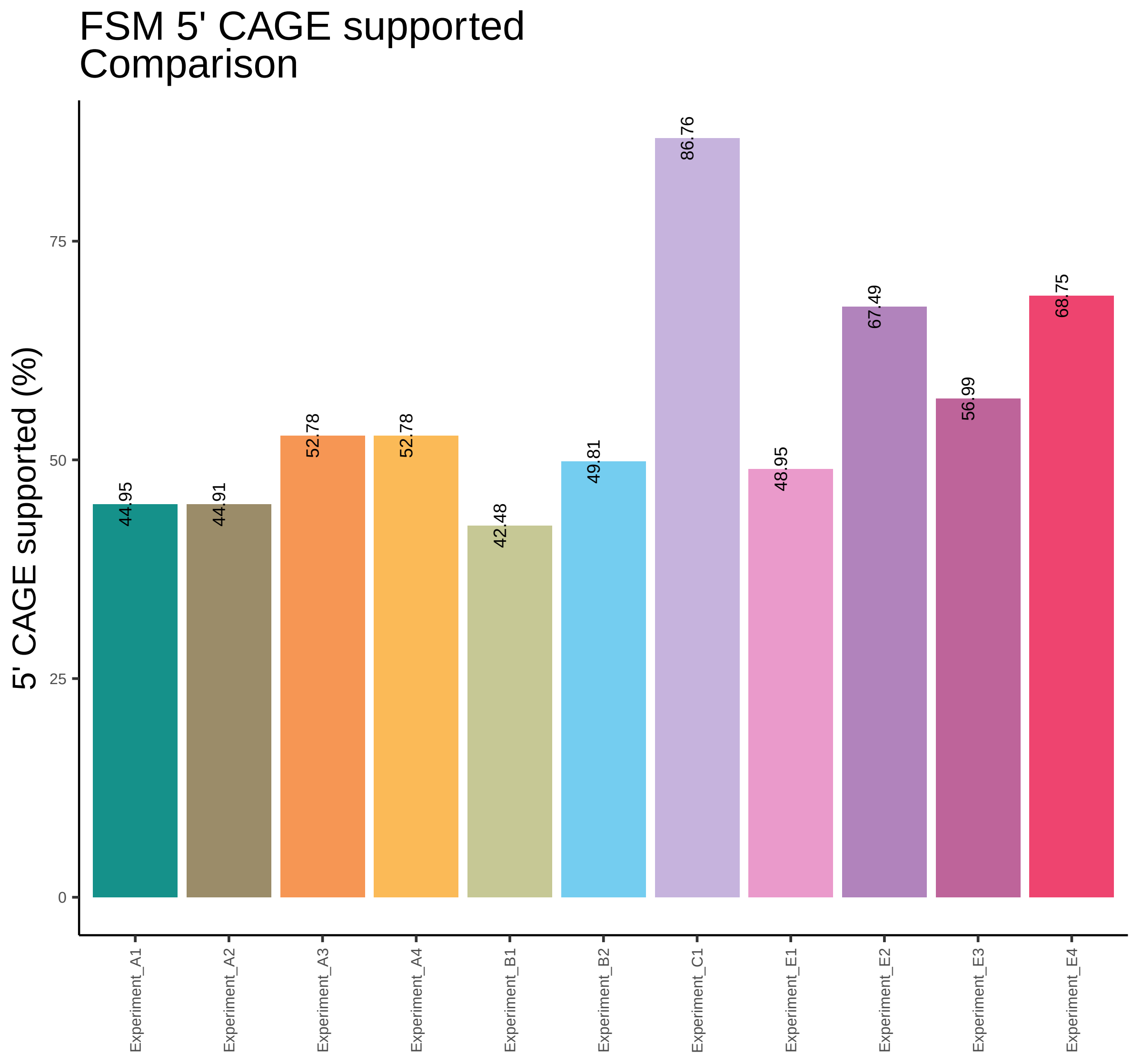
FSM 5' CAGE supported
Comparison
5' CAGE supported (%)
0
25
50
75
44.95
44.91
52.78
52.78
42.48
49.81
86.76
48.95
67.49
56.99
68.75
Experiment_A1
Experiment_A2
Experiment_A3
Experiment_A4
Experiment_B1
Experiment_B2
Experiment_C1
Experiment_E1
Experiment_E2
Experiment_E3
Experiment_E4

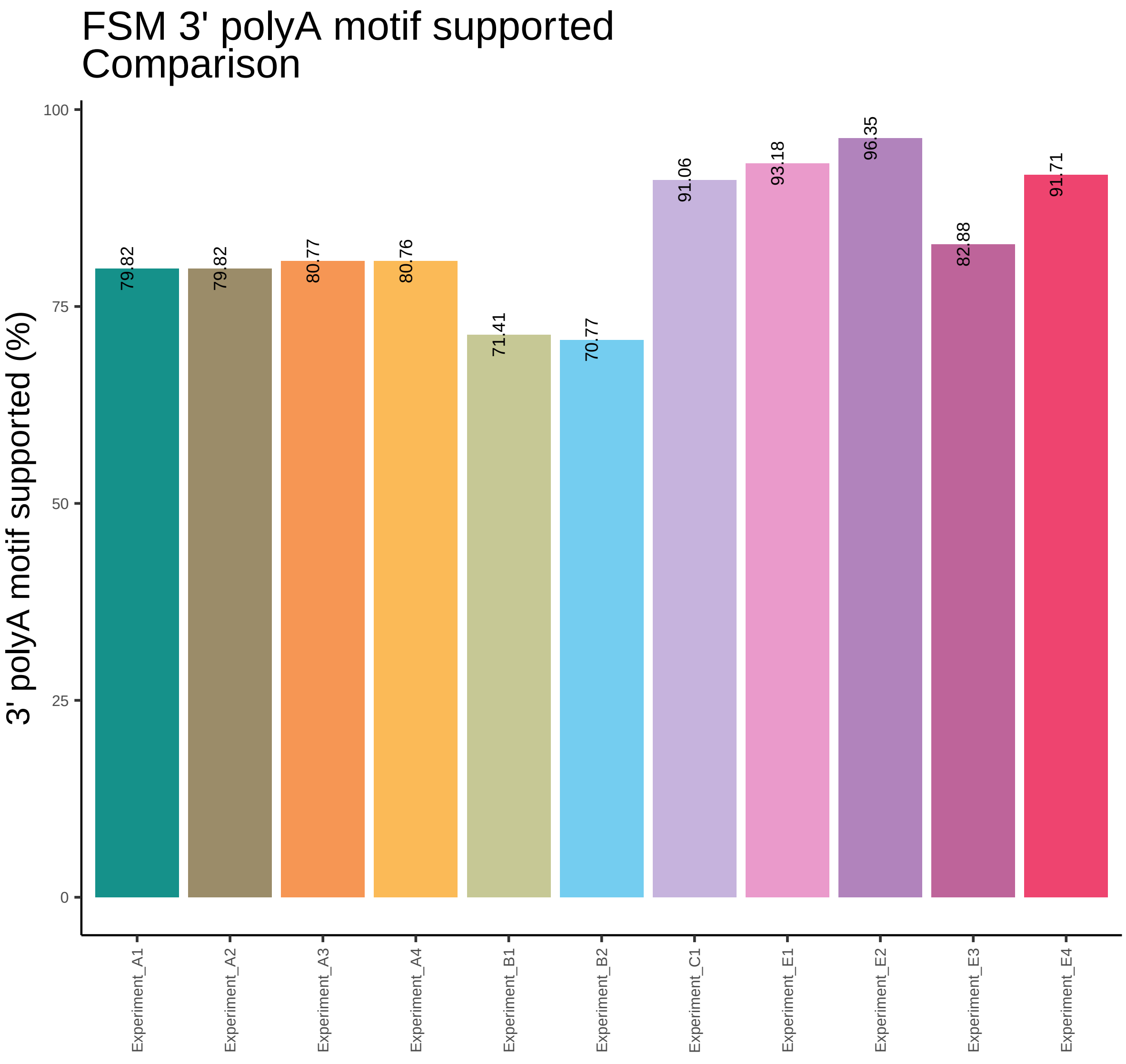
FSM 3' polyA motif supported
Comparison
3' polyA motif supported (%)
100
75
50
25
0
79.82
79.82
80.77
80.76
71.41
70.77
91.06
93.18
96.35
82.88
91.71
Experiment_A1
Experiment_A2
Experiment_A3
Experiment_A4
Experiment_B1
Experiment_B2
Experiment_C1
Experiment_E1
Experiment_E2
Experiment_E3
Experiment_E4

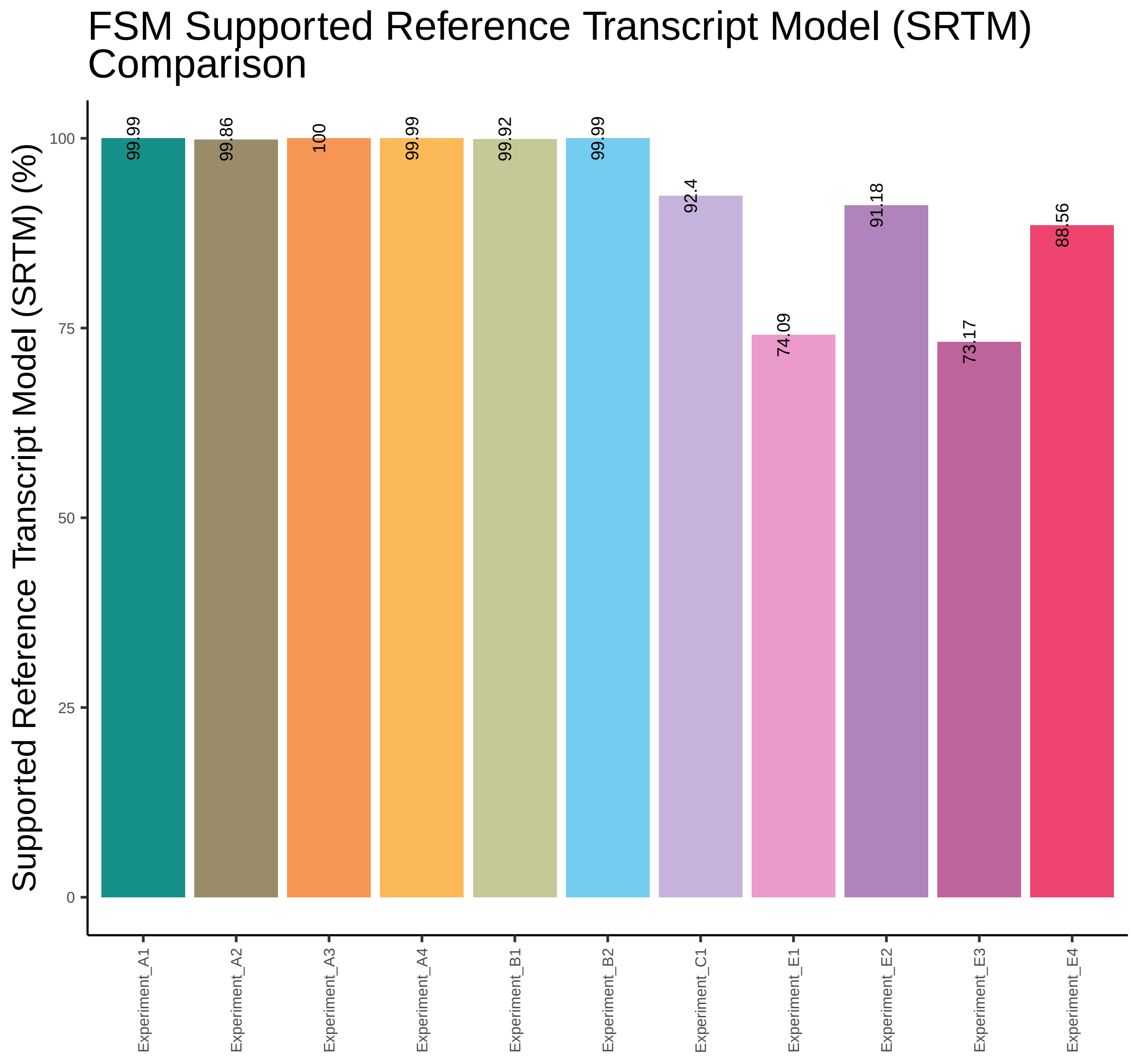

FSM Supported Reference Transcript Model (SRTM) Comparison
Supported Reference Transcript Model (SRTM) (%)
100
75
50
25
0
99.99
99.86
100
99.99
99.92
99.99
92.4
74.09
91.18
73.17
88.56
Experiment_A1
Experiment_A2
Experiment_A3
Experiment_A4
Experiment_B1
Experiment_B2
Experiment_C1
Experiment_E1
Experiment_E2
Experiment_E3
Experiment_E4

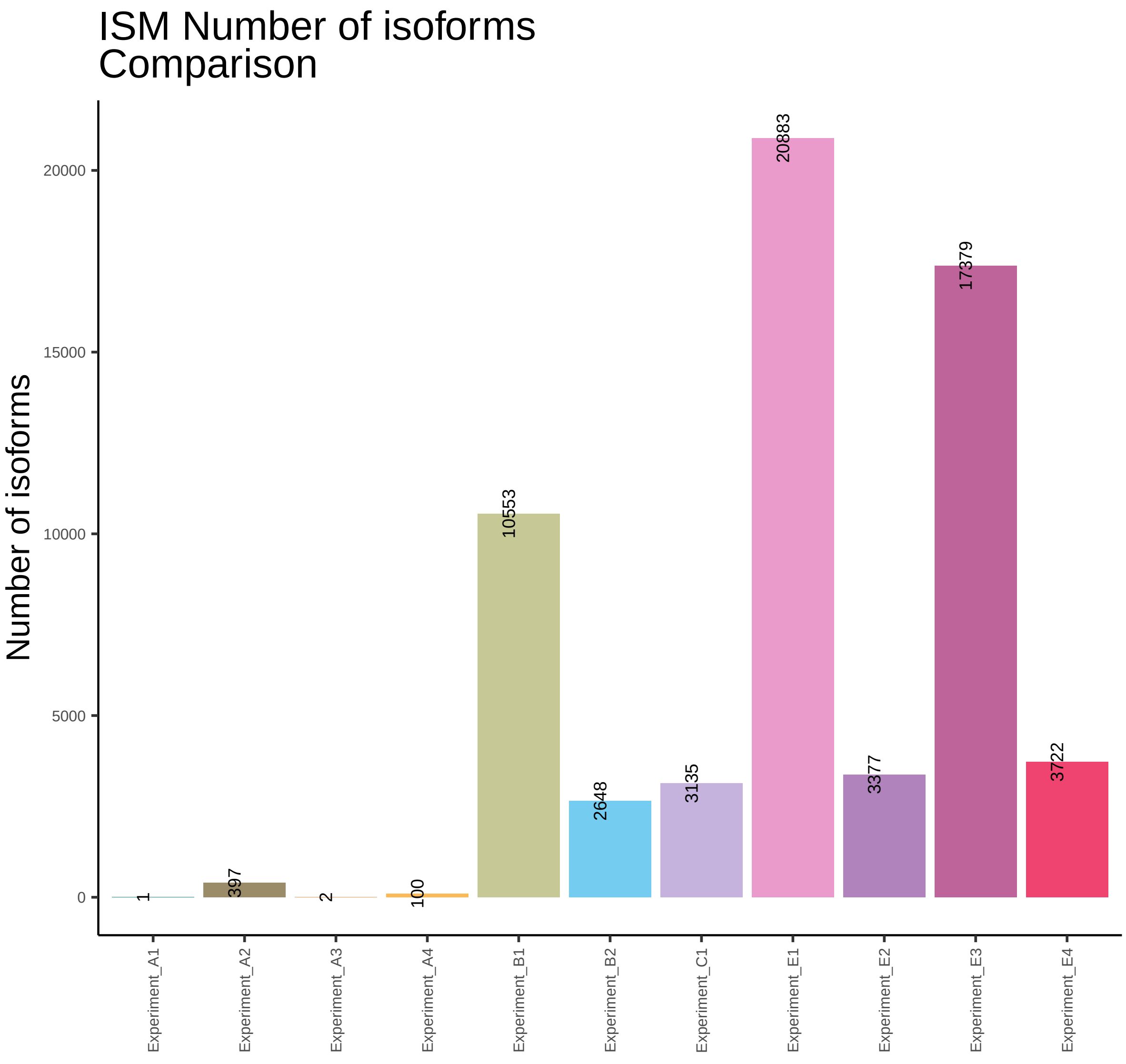
ISM Number of isoforms
Comparison
Number of isoforms
20000
15000
10000
5000
0
1
397
2
100
10553
2648
3135
20883
3377
17379
3722
Experiment_A1
Experiment_A2
Experiment_A3
Experiment_A4
Experiment_B1
Experiment_B2
Experiment_C1
Experiment_E1
Experiment_E2
Experiment_E3
Experiment_E4

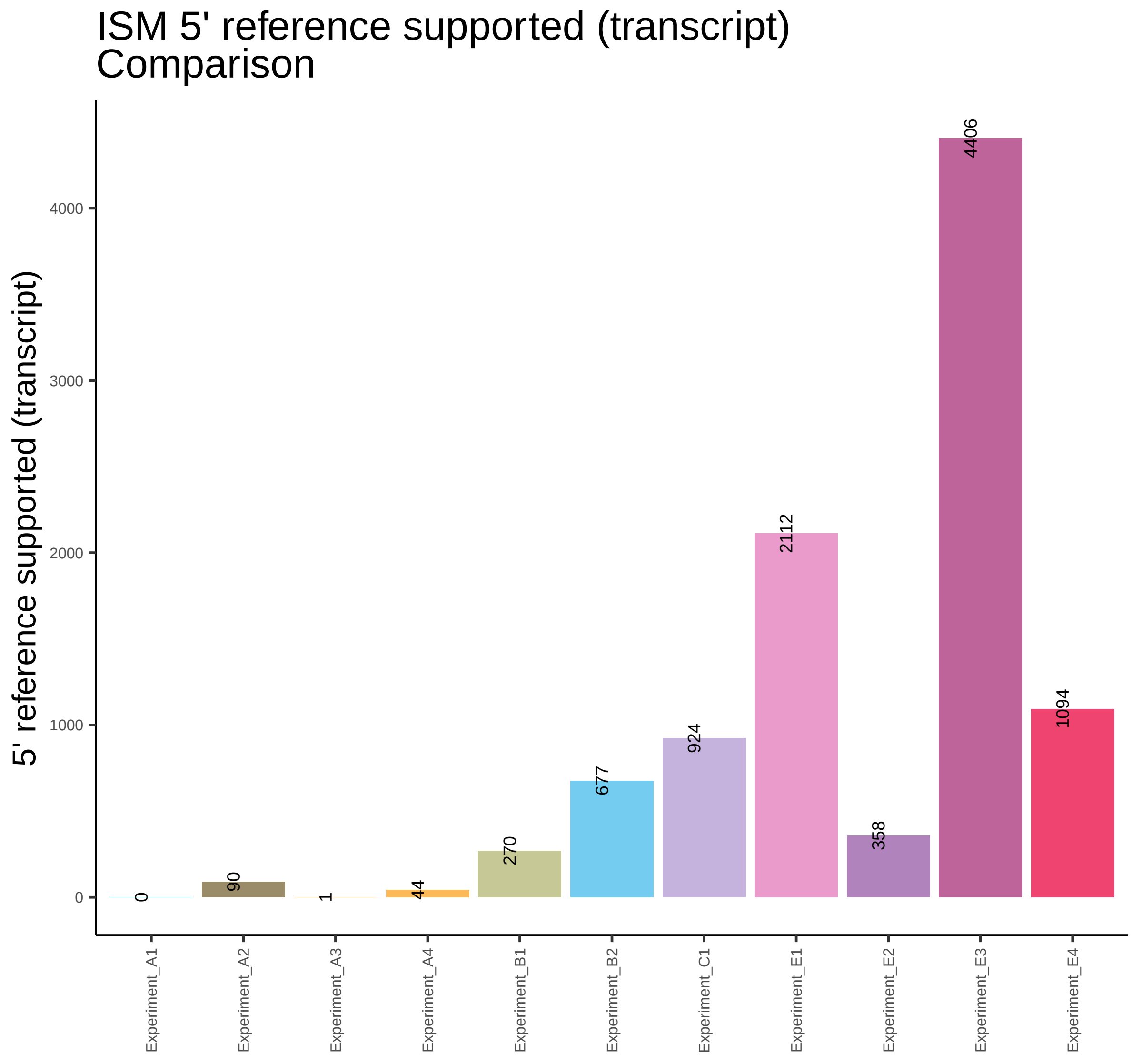

ISM 5' reference supported (transcript)
Comparison
5' reference supported (transcript)
0
1000
2000
3000
4000
0
90
1
44
270
677
924
2112
358
4406
1094
Experiment_A1
Experiment_A2
Experiment_A3
Experiment_A4
Experiment_B1
Experiment_B2
Experiment_C1
Experiment_E1
Experiment_E2
Experiment_E3
Experiment_E4

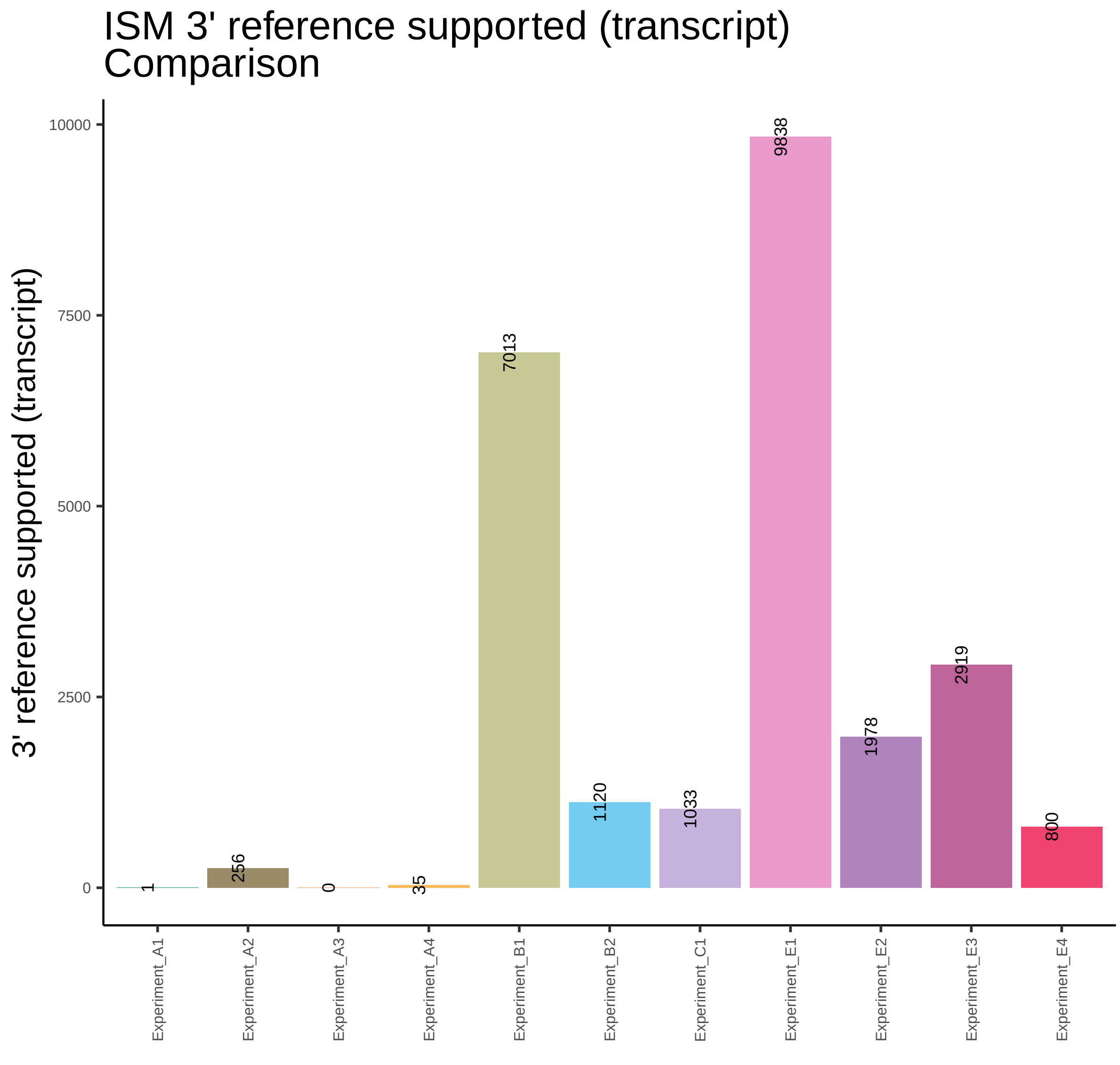
ISM 3' reference supported (transcript)
Comparison
3' reference supported (transcript)
10000
7500
5000
2500
0
1
256
0
35
7013
1120
1033
9838
1978
2919
800
Experiment_A1
Experiment_A2
Experiment_A3
Experiment_A4
Experiment_B1
Experiment_B2
Experiment_C1
Experiment_E1
Experiment_E2
Experiment_E3
Experiment_E4

### ISM 5' and 3' reference supported (gene) Comparison

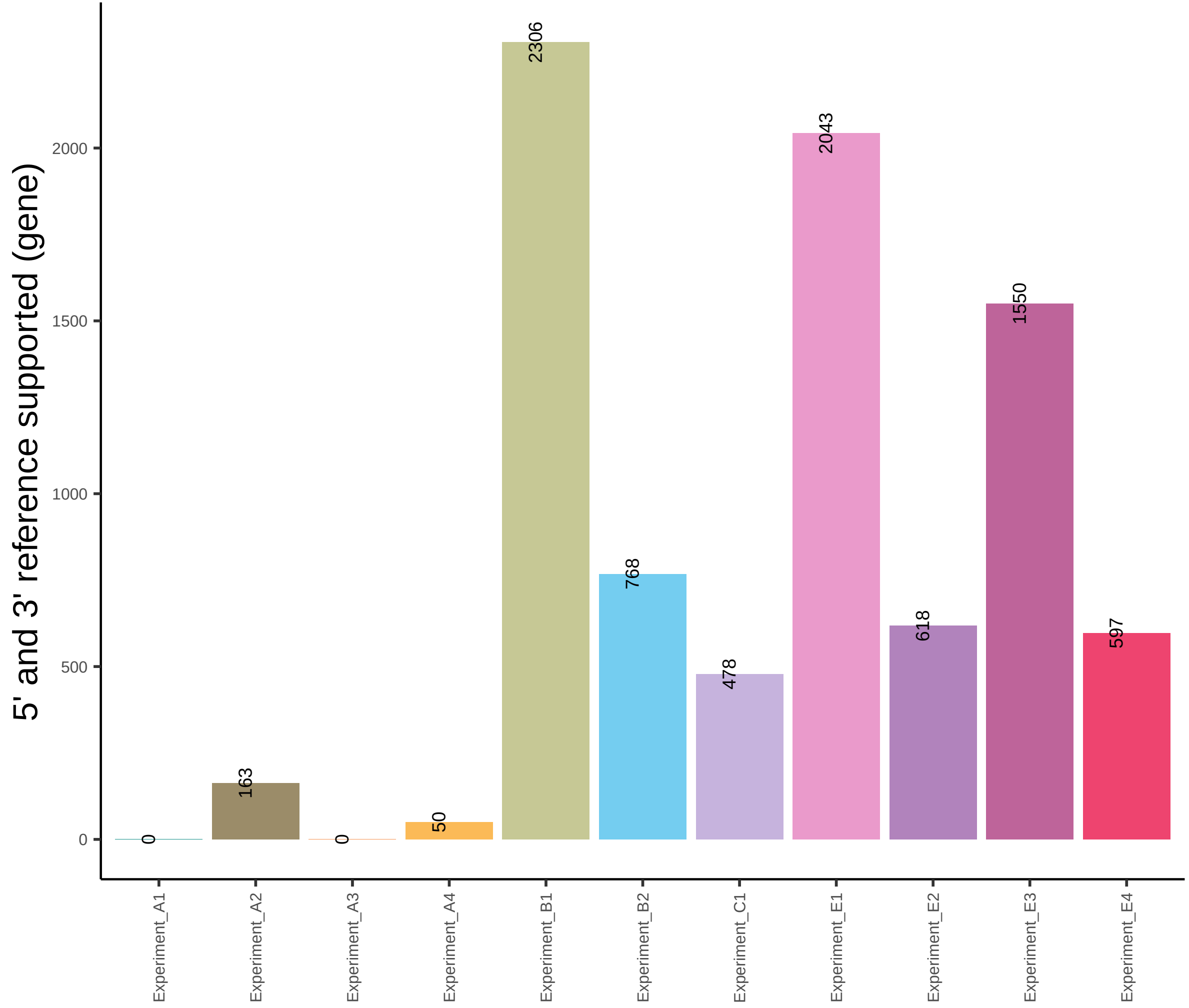

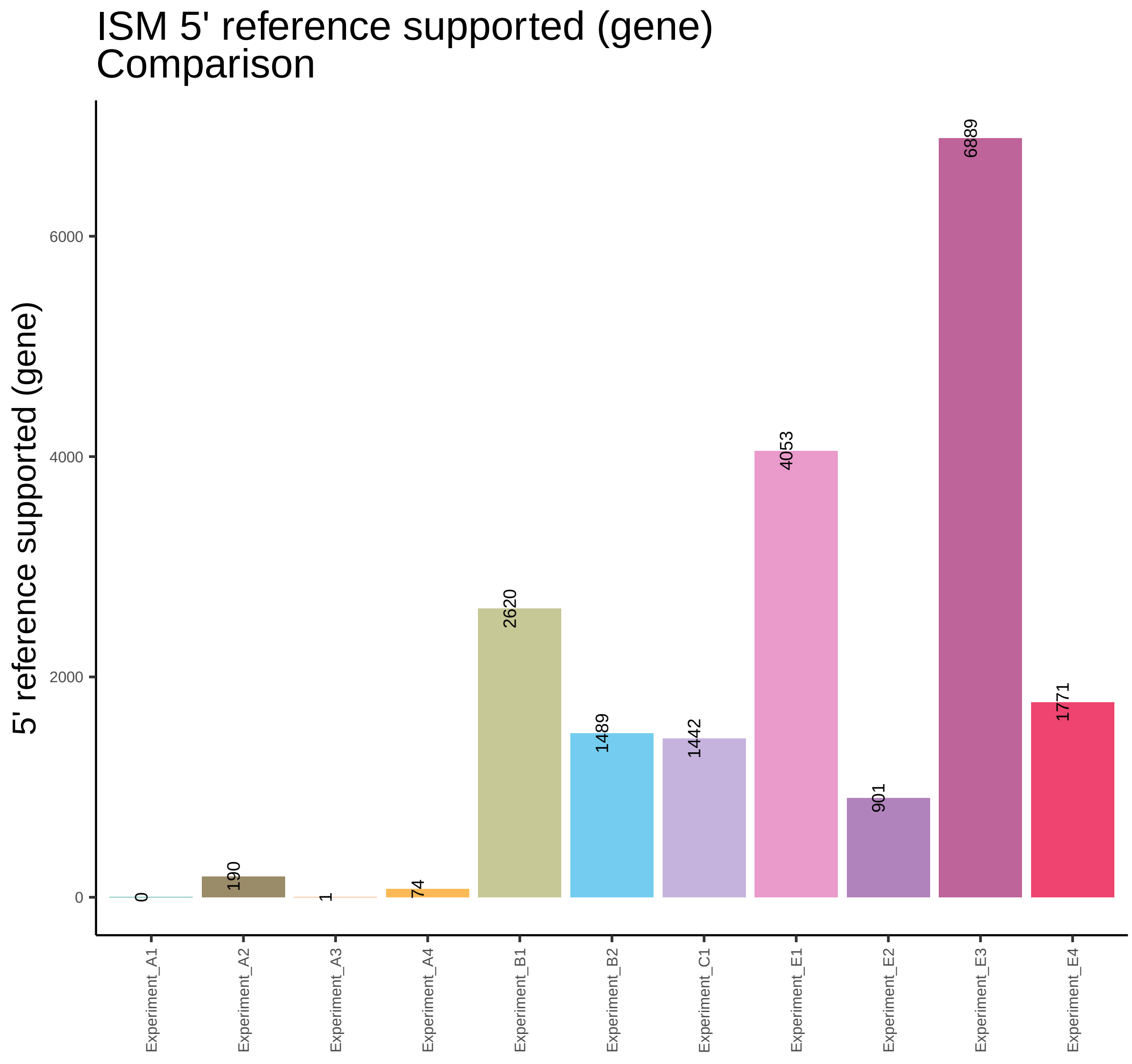

ISM 5' reference supported (gene)
Comparison
5' reference supported (gene)
0
2000
4000
6000
0
190
1
74
2620
1489
1442
4053
901
6889
1771
Experiment_A1
Experiment_A2
Experiment_A3
Experiment_A4
Experiment_B1
Experiment_B2
Experiment_C1
Experiment_E1
Experiment_E2
Experiment_E3
Experiment_E4

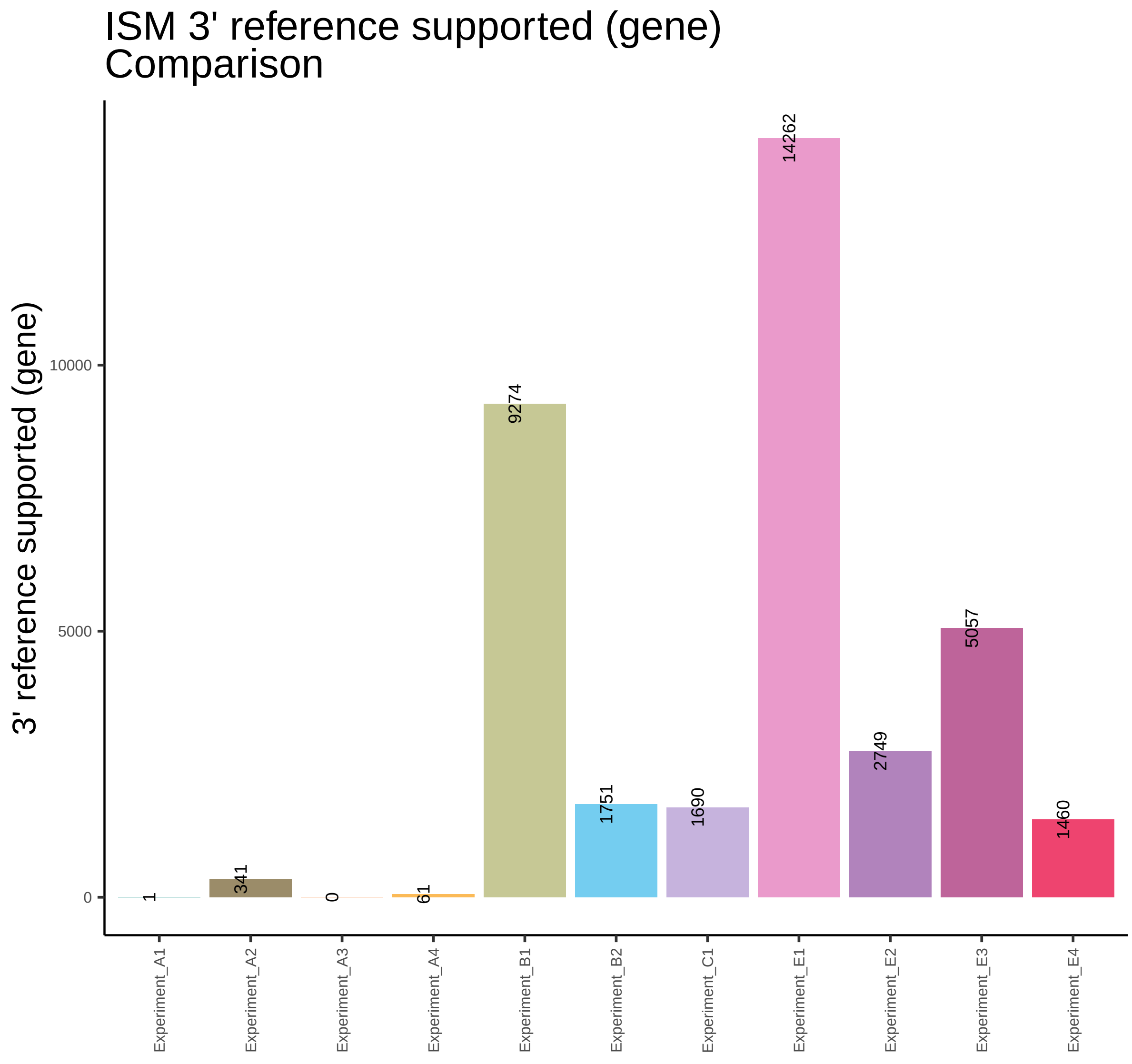

ISM 3' reference supported (gene)
Comparison
3' reference supported (gene)
10000
5000
0
1
341
0
61
9274
1751
1690
14262
2749
5057
1460
Experiment_A1
Experiment_A2
Experiment_A3
Experiment_A4
Experiment_B1
Experiment_B2
Experiment_C1
Experiment_E1
Experiment_E2
Experiment_E3
Experiment_E4

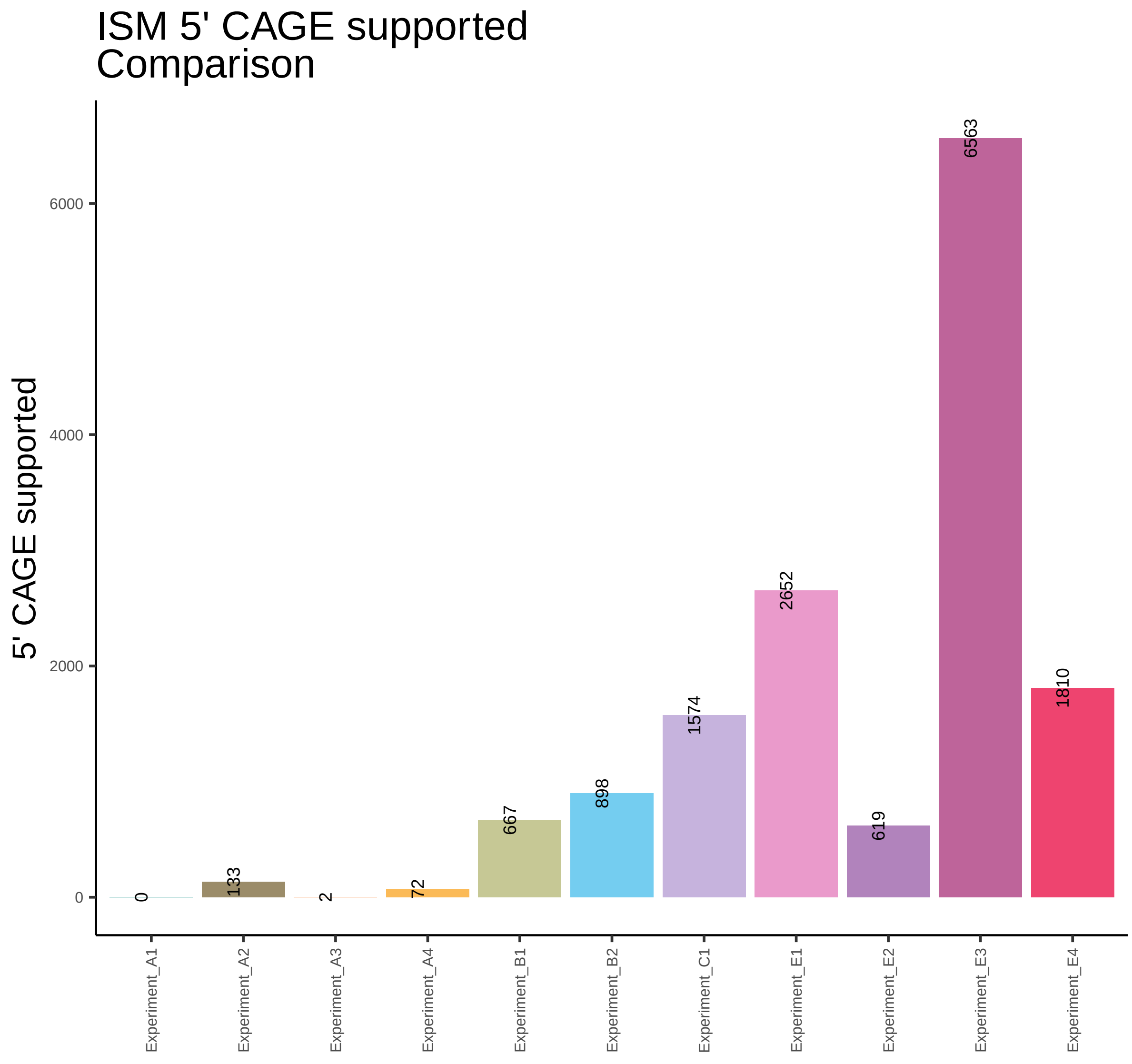

ISM 5' CAGE supported
Comparison
5' CAGE supported
6000
4000
2000
0
0
133
2
72
667
898
1574
2652
619
6563
1810
Experiment_A1
Experiment_A2
Experiment_A3
Experiment_A4
Experiment_B1
Experiment_B2
Experiment_C1
Experiment_E1
Experiment_E2
Experiment_E3
Experiment_E4

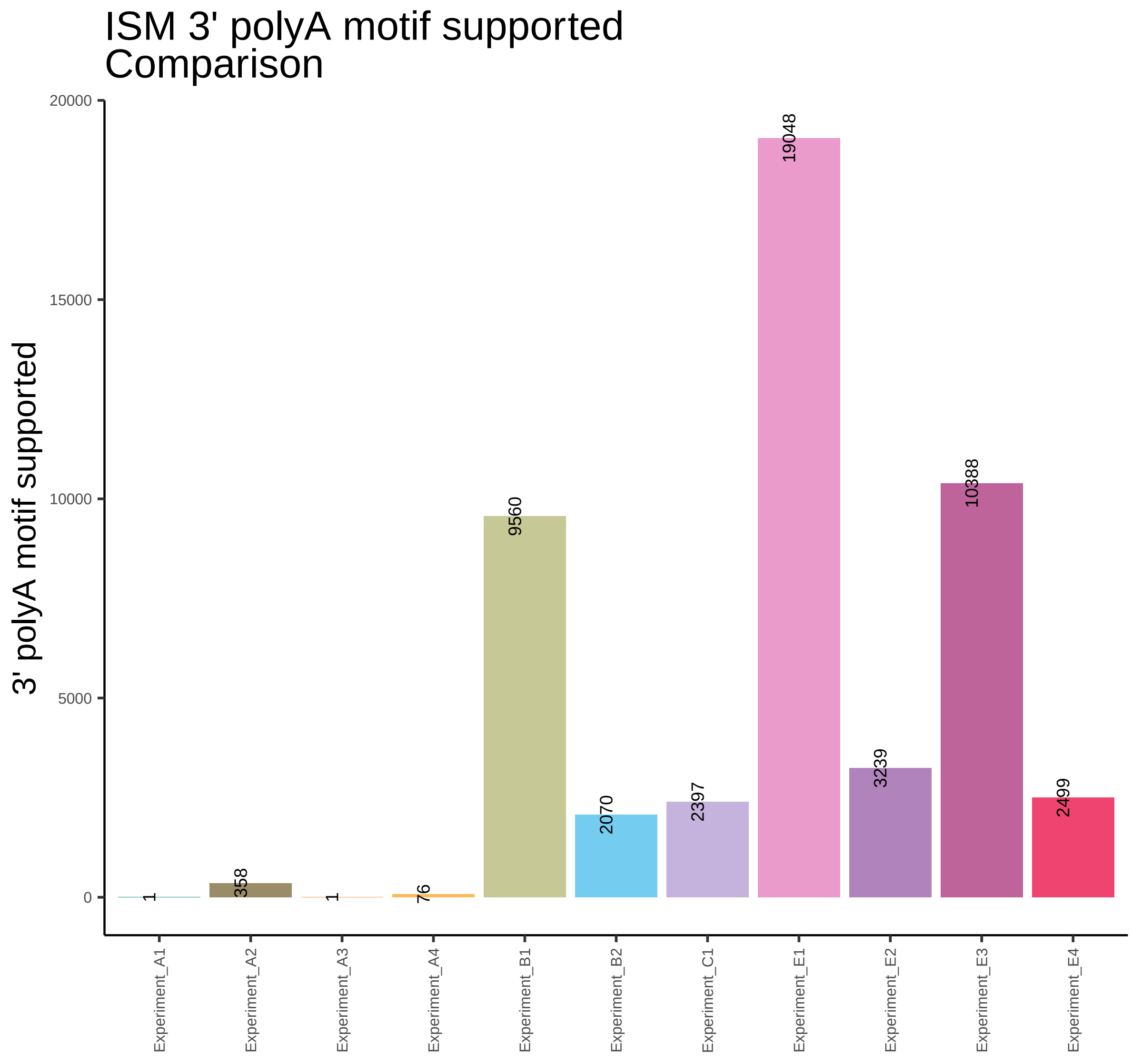

ISM 3' polyA motif supported
Comparison
3' polyA motif supported
20000
15000
10000
5000
0
1
358
1
76
9560
2070
2397
19048
3239
10388
2499
Experiment_A1
Experiment_A2
Experiment_A3
Experiment_A4
Experiment_B1
Experiment_B2
Experiment_C1
Experiment_E1
Experiment_E2
Experiment_E3
Experiment_E4

### ISM Supported Reference Transcript Model (SRTM) Comparison

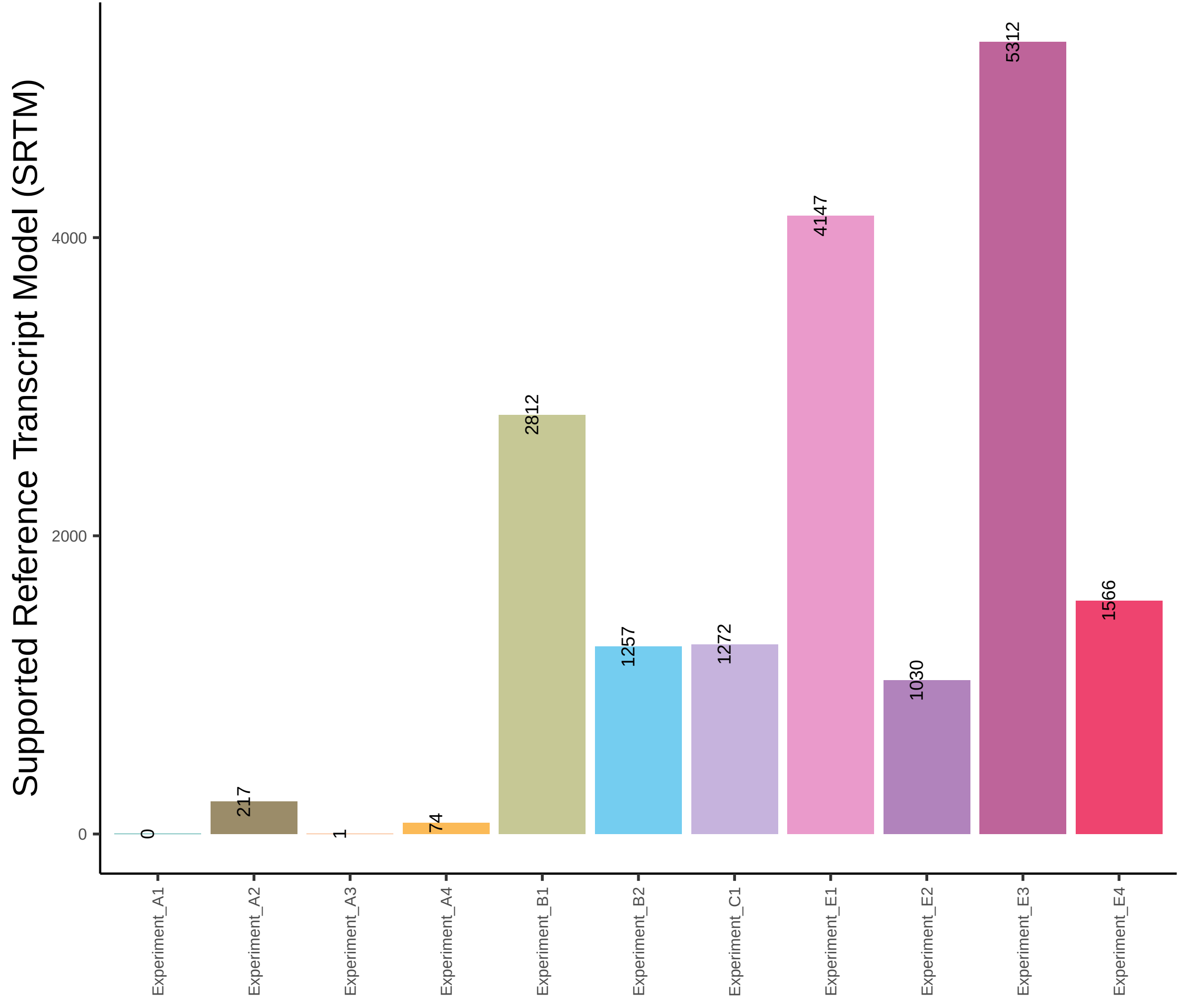

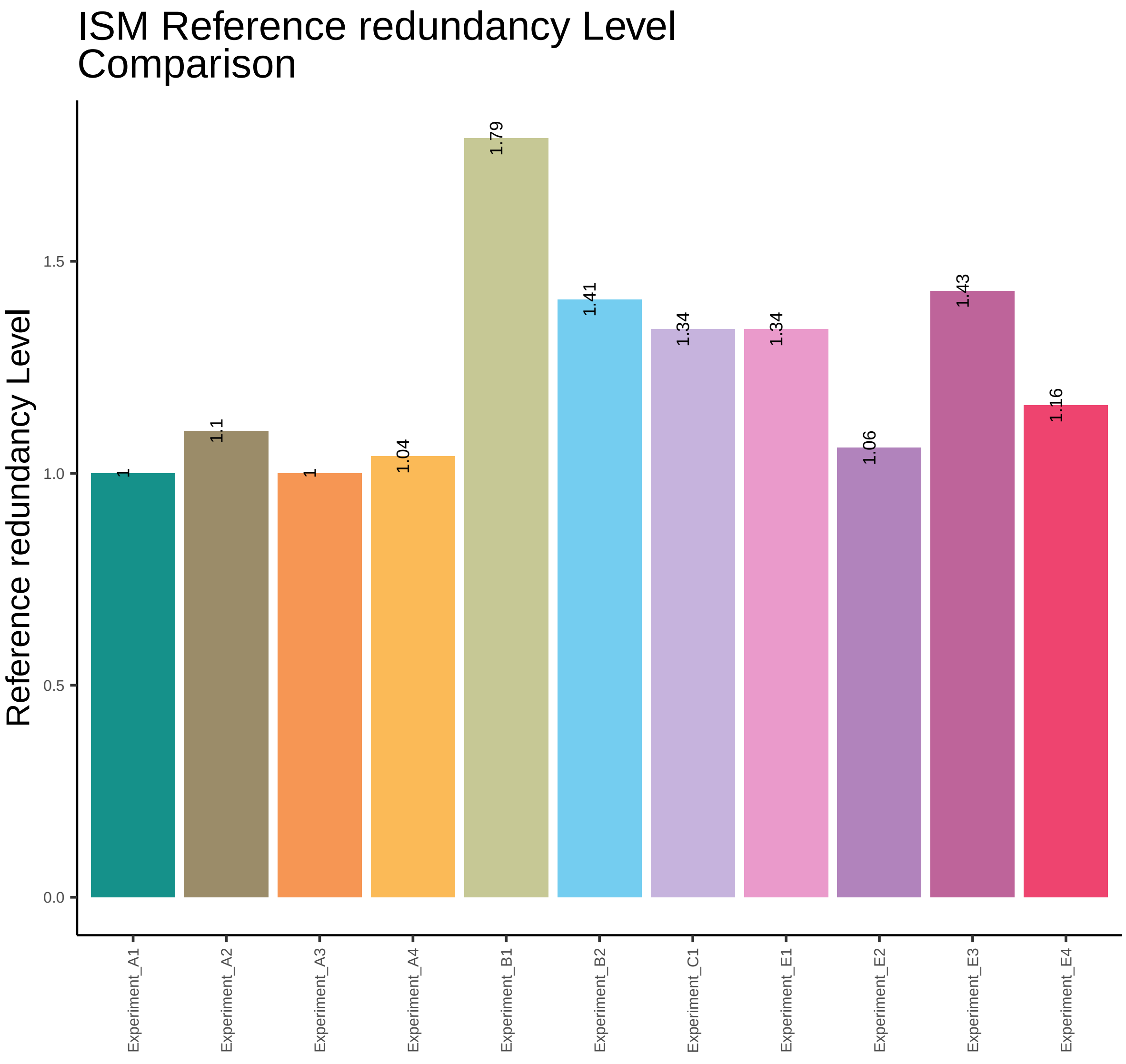
ISM Reference redundancy Level Comparison
Reference redundancy Level
0.0
0.5
1.0
1.5
1
1.1
1
1.04
1.79
1.41
1.34
1.34
1.06
1.43
1.16
Experiment_A1
Experiment_A2
Experiment_A3
Experiment_A4
Experiment_B1
Experiment_B2
Experiment_C1
Experiment_E1
Experiment_E2
Experiment_E3
Experiment_E4

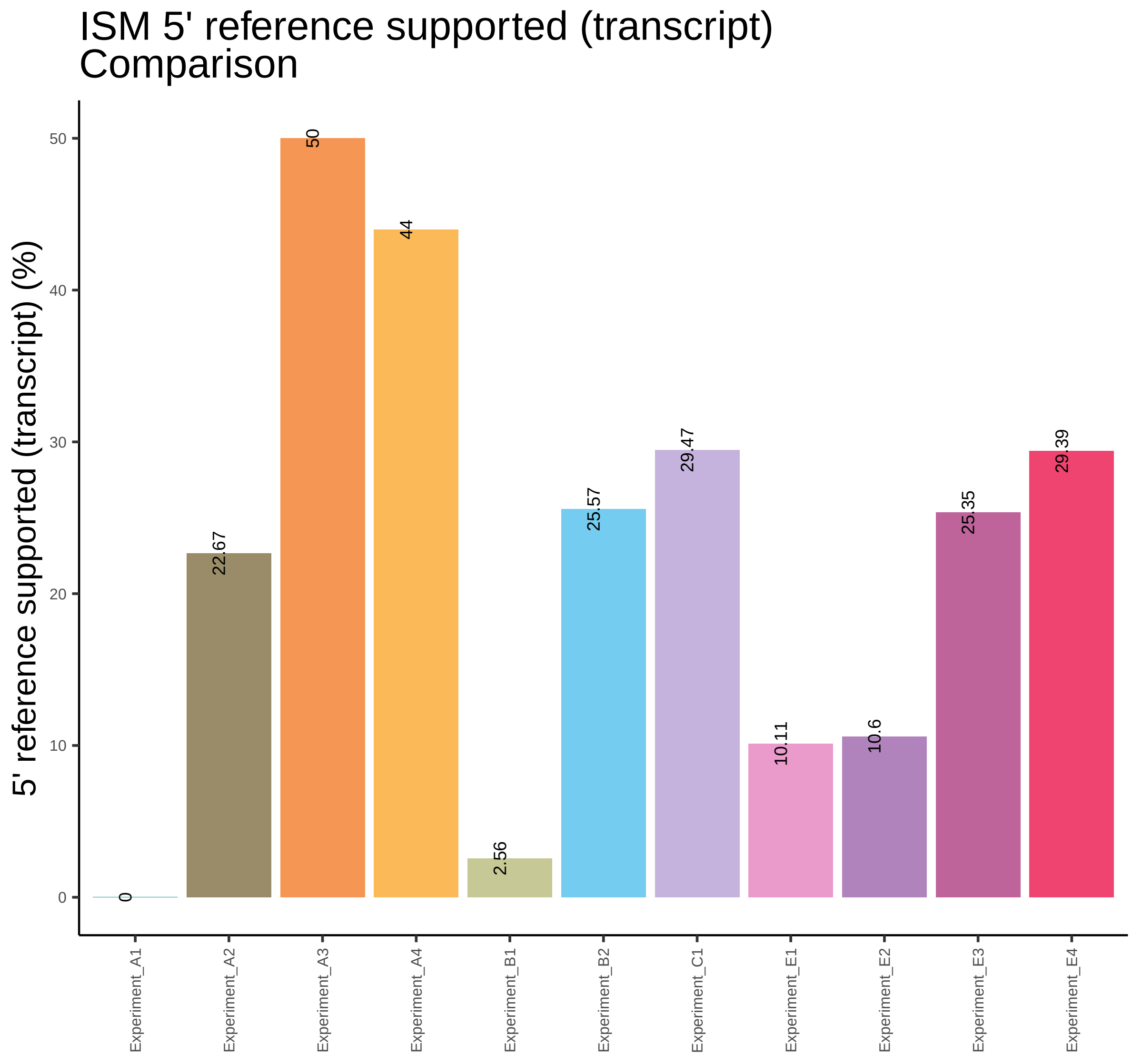
ISM 5' reference supported (transcript)
Comparison
5' reference supported (transcript) (%)
0
10
20
30
40
50
0
22.67
50
44
2.56
25.57
29.47
10.11
10.6
25.35
29.39
Experiment_A1
Experiment_A2
Experiment_A3
Experiment_A4
Experiment_B1
Experiment_B2
Experiment_C1
Experiment_E1
Experiment_E2
Experiment_E3
Experiment_E4

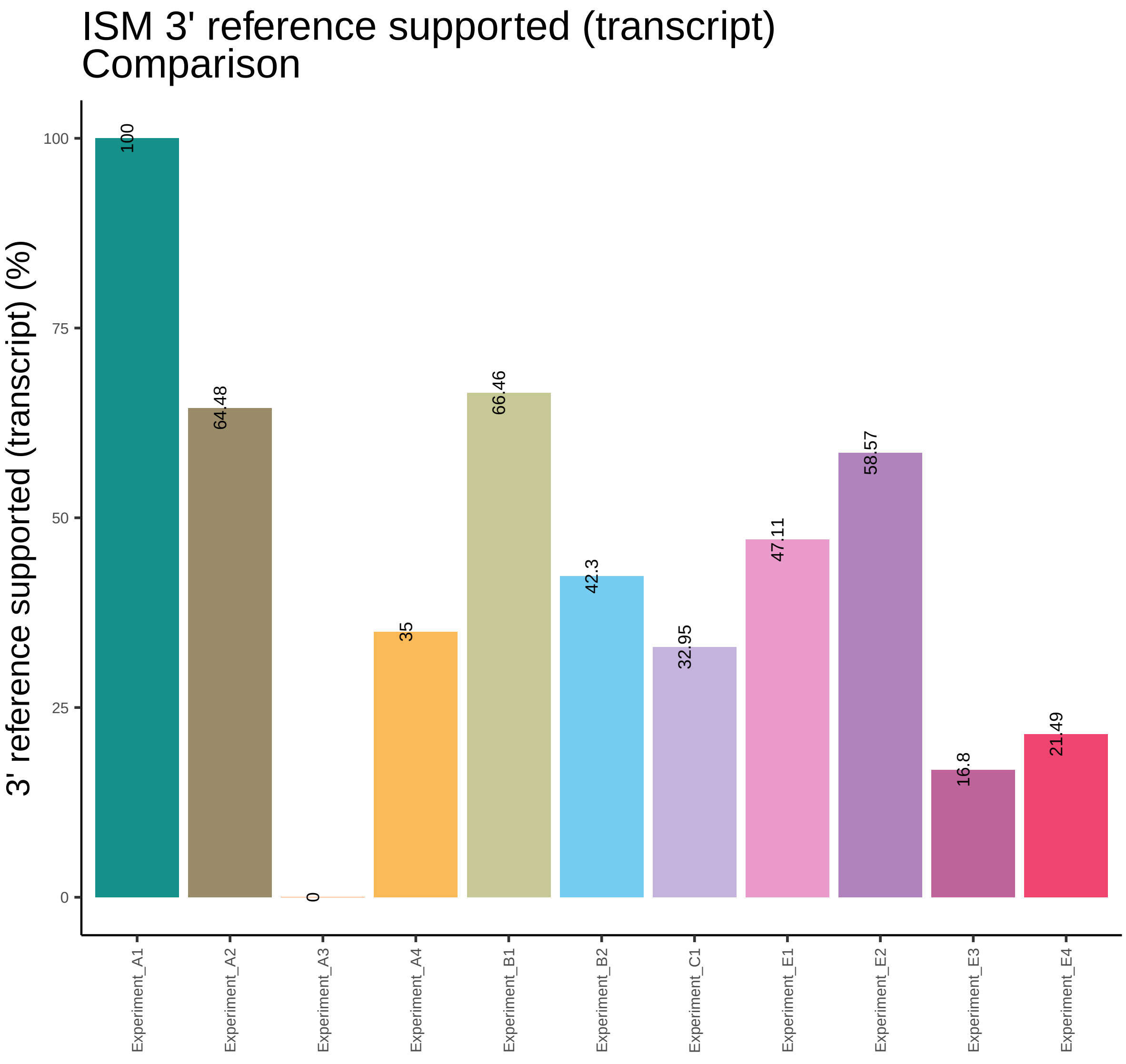
ISM 3' reference supported (transcript)
Comparison
3' reference supported (transcript) (%)
100
75
50
25
0
100
64.48
0
35
66.46
42.3
32.95
47.11
58.57
16.8
21.49
Experiment_A1
Experiment_A2
Experiment_A3
Experiment_A4
Experiment_B1
Experiment_B2
Experiment_C1
Experiment_E1
Experiment_E2
Experiment_E3
Experiment_E4

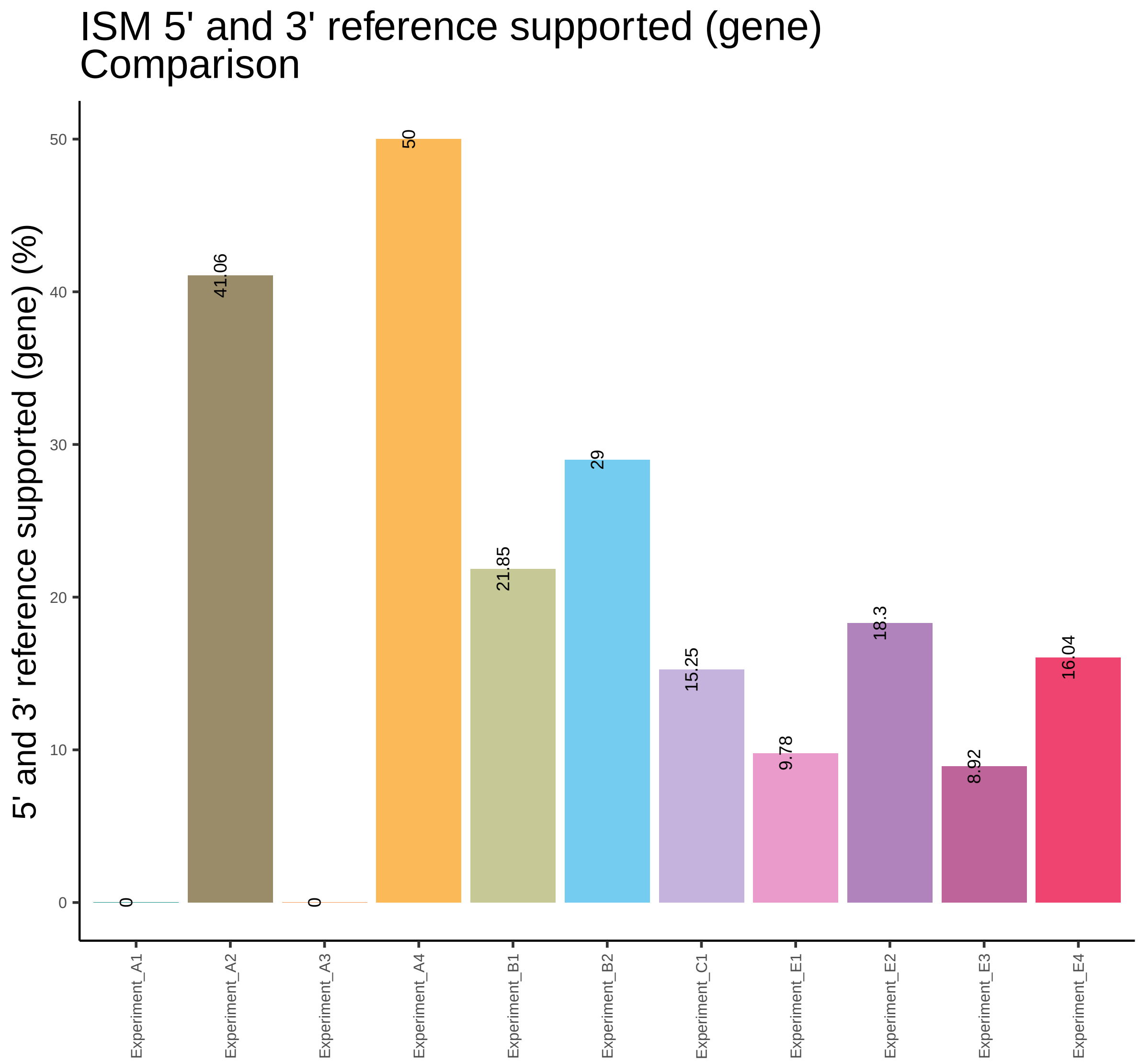
ISM 5' and 3' reference supported (gene)
Comparison
5' and 3' reference supported (gene) (%)
0
10
20
30
40
50
0
41.06
0
50
21.85
29
15.25
9.78
18.3
8.92
16.04
Experiment_A1
Experiment_A2
Experiment_A3
Experiment_A4
Experiment_B1
Experiment_B2
Experiment_C1
Experiment_E1
Experiment_E2
Experiment_E3
Experiment_E4

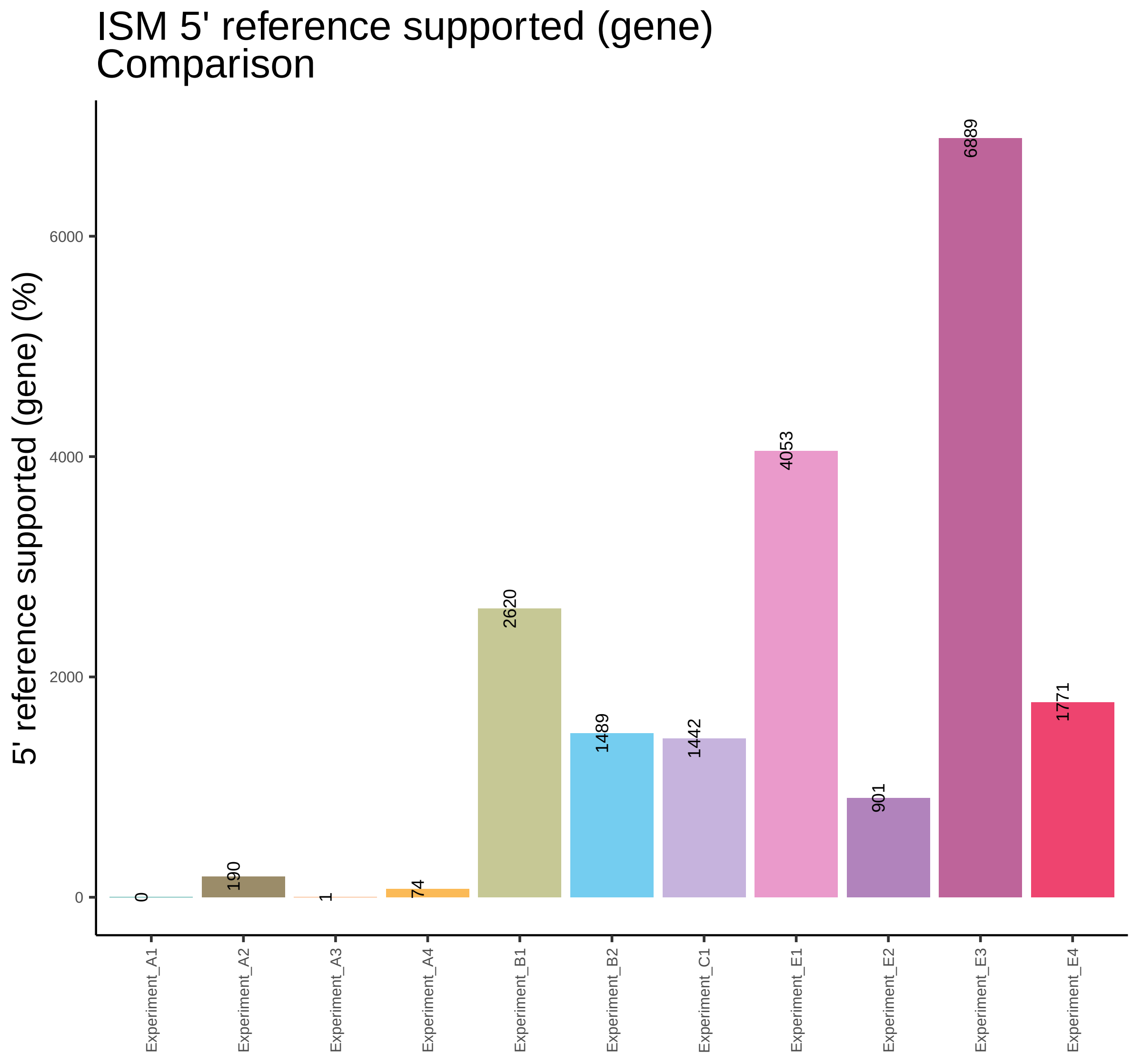

ISM 5' reference supported (gene)
Comparison
5' reference supported (gene) (%)
0
2000
4000
6000
0
190
1
74
2620
1489
1442
4053
901
6889
1771
Experiment_A1
Experiment_A2
Experiment_A3
Experiment_A4
Experiment_B1
Experiment_B2
Experiment_C1
Experiment_E1
Experiment_E2
Experiment_E3
Experiment_E4

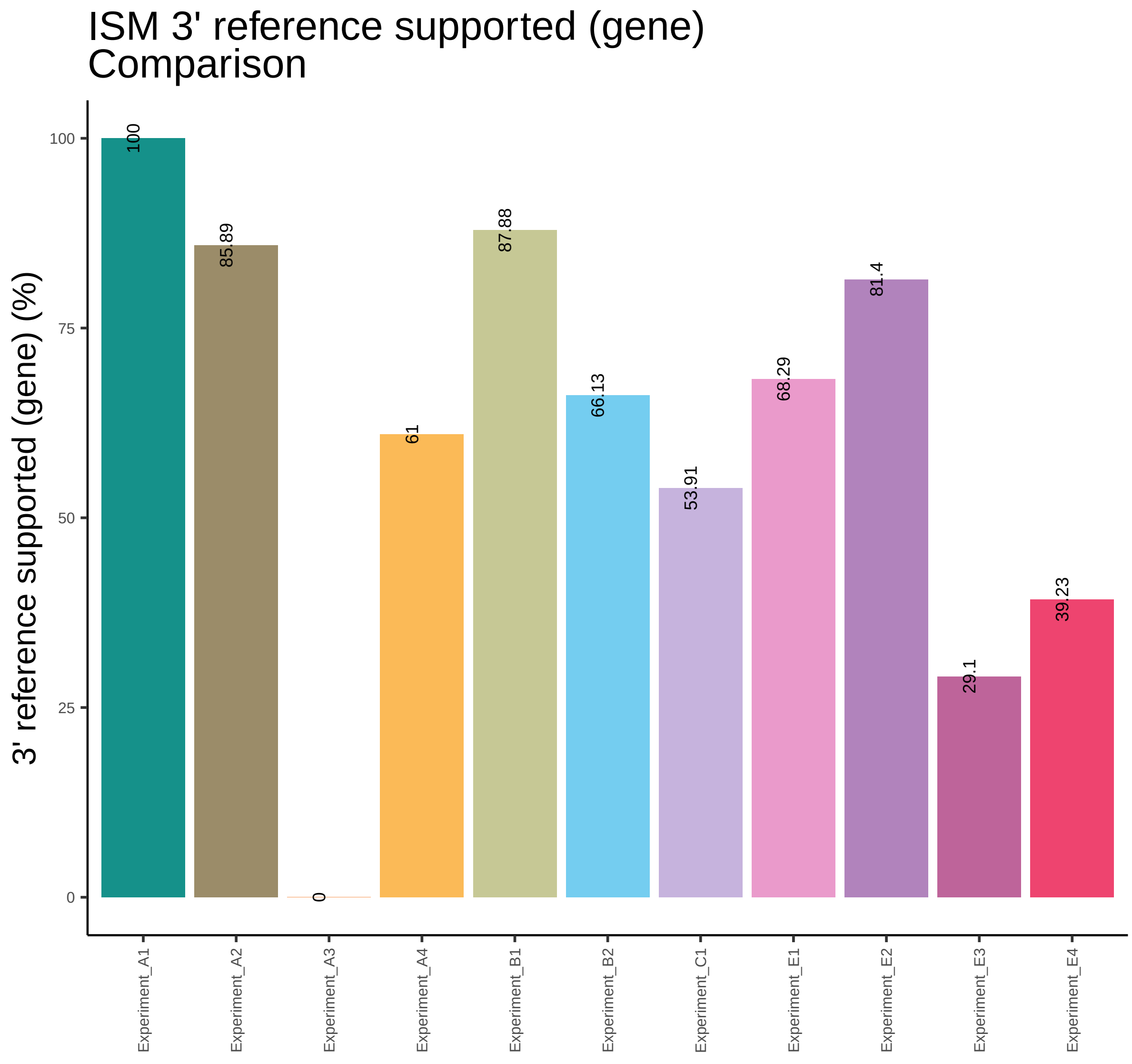

ISM 3' reference supported (gene)
Comparison
3' reference supported (gene) (%)
100
75
50
25
0
100
85.89
0
61
87.88
66.13
53.91
68.29
81.4
29.1
39.23
Experiment_A1
Experiment_A2
Experiment_A3
Experiment_A4
Experiment_B1
Experiment_B2
Experiment_C1
Experiment_E1
Experiment_E2
Experiment_E3
Experiment_E4

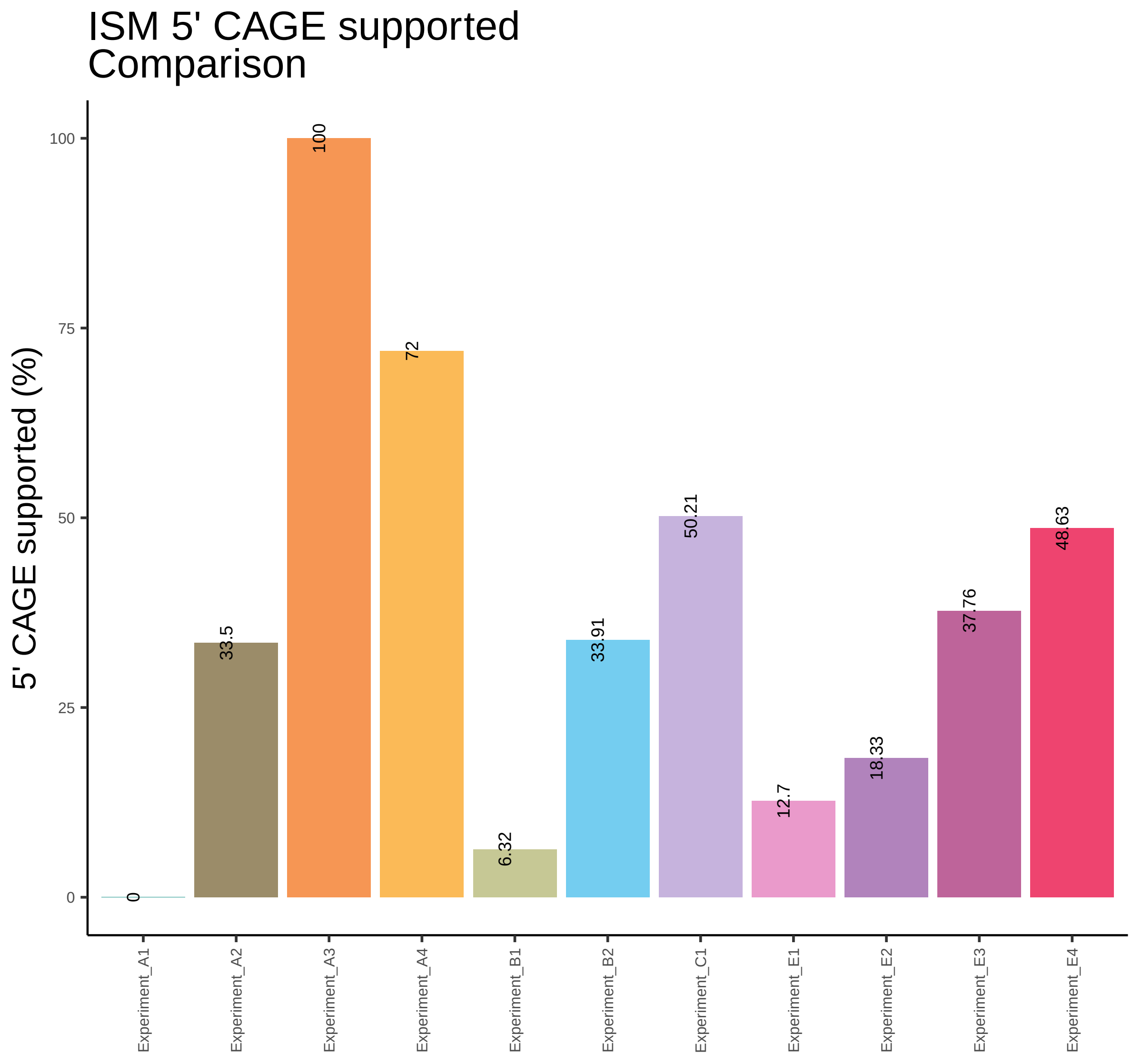
ISM 5' CAGE supported
Comparison
5' CAGE supported (%)
100
75
50
25
0
0
33.5
100
72
6.32
33.91
50.21
12.7
18.33
37.76
48.63
Experiment_A1
Experiment_A2
Experiment_A3
Experiment_A4
Experiment_B1
Experiment_B2
Experiment_C1
Experiment_E1
Experiment_E2
Experiment_E3
Experiment_E4

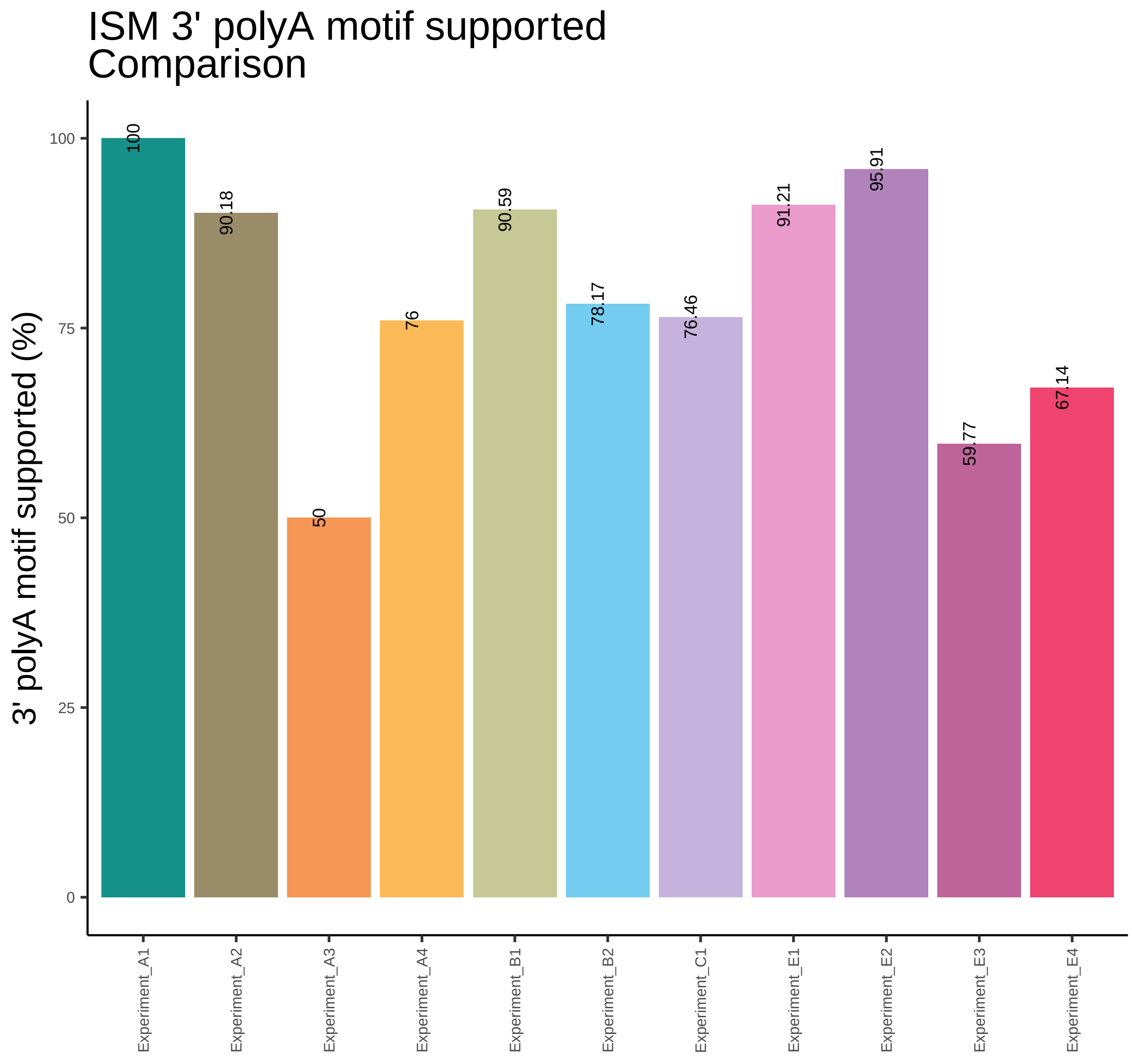
ISM 3' polyA motif supported
Comparison
3' polyA motif supported (%)
0
25
50
75
100
100
90.18
50
76
90.59
78.17
76.46
91.21
95.91
59.77
67.14
Experiment_A1
Experiment_A2
Experiment_A3
Experiment_A4
Experiment_B1
Experiment_B2
Experiment_C1
Experiment_E1
Experiment_E2
Experiment_E3
Experiment_E4

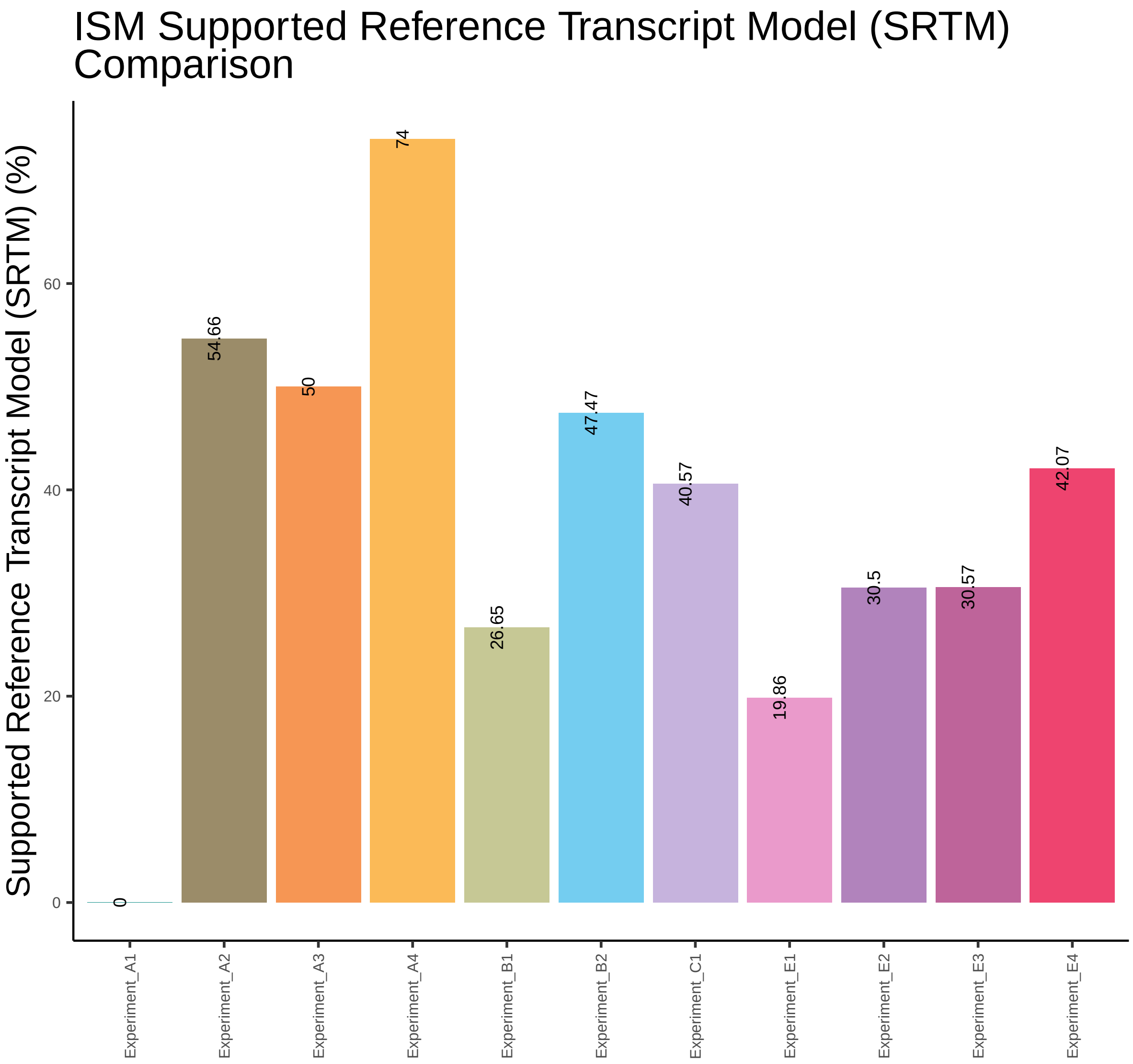
ISM Supported Reference Transcript Model (SRTM) Comparison
Supported Reference Transcript Model (SRTM) (%)
0
20
40
60
0
54.66
50
74
26.65
47.47
40.57
19.86
30.5
30.57
42.07
Experiment_A1
Experiment_A2
Experiment_A3
Experiment_A4
Experiment_B1
Experiment_B2
Experiment_C1
Experiment_E1
Experiment_E2
Experiment_E3
Experiment_E4

NIC Number of isoforms
Comparison

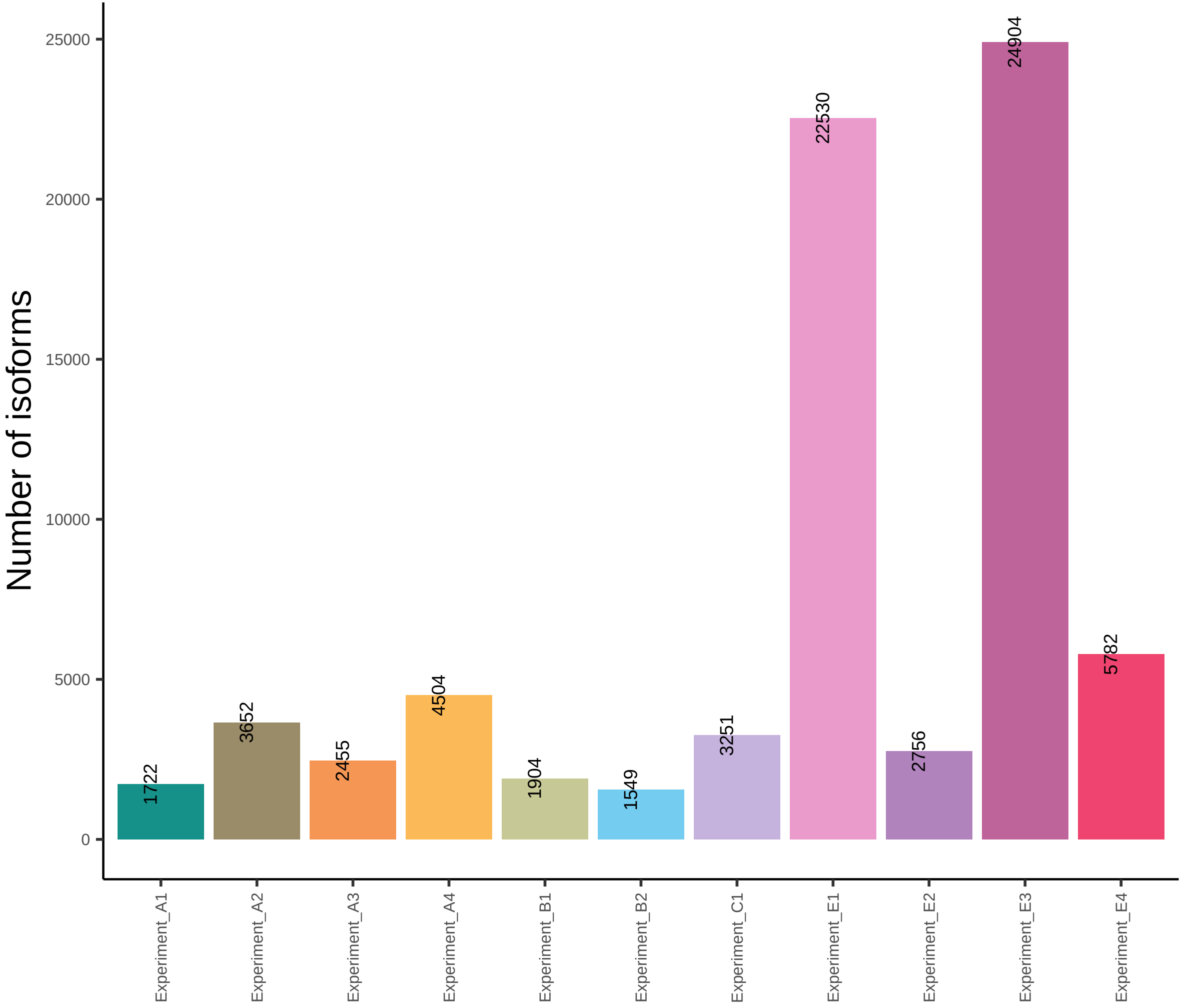

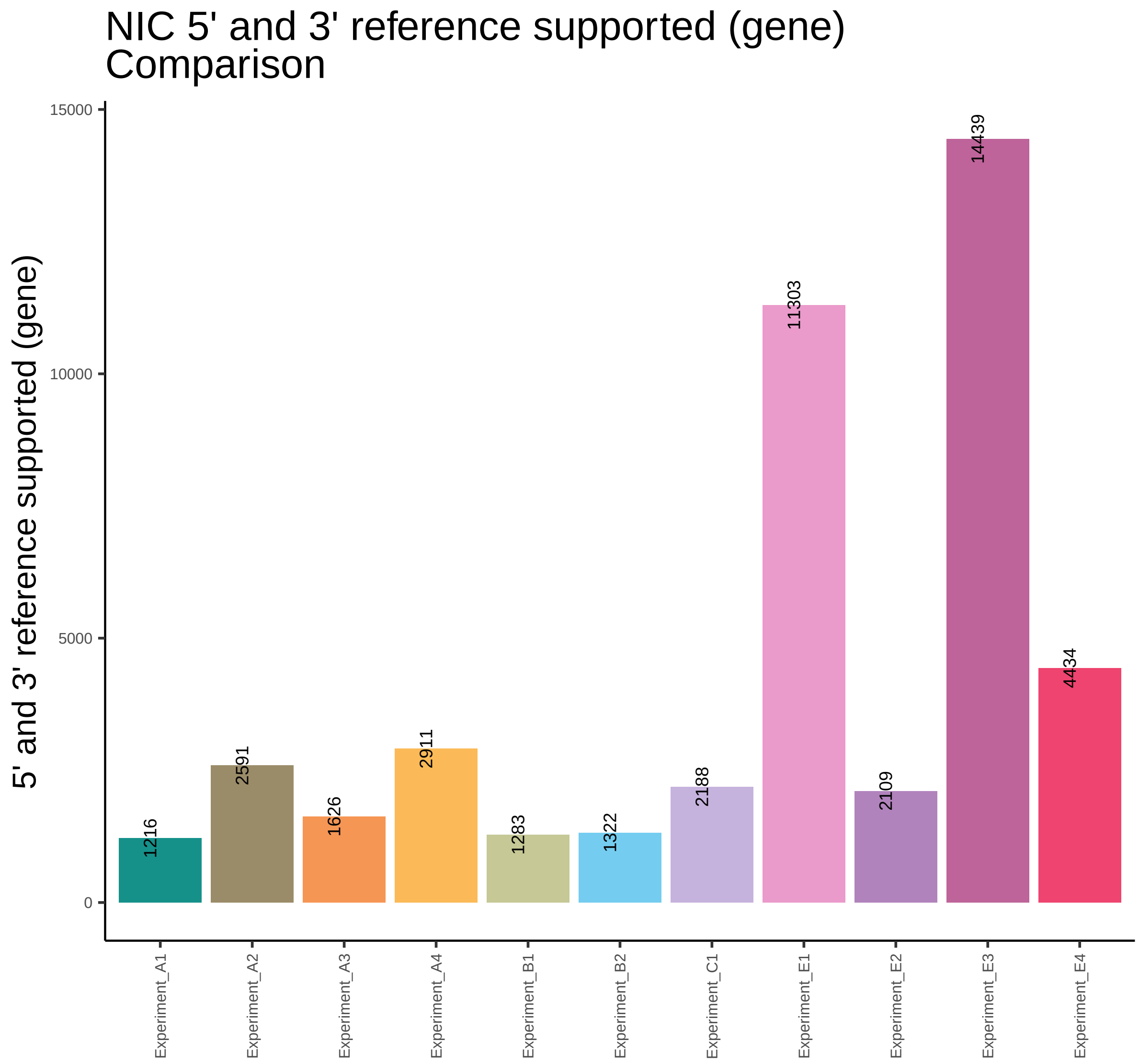

NIC 5' and 3' reference supported (gene)
Comparison
5' and 3' reference supported (gene)
15000
10000
5000
0
1216
2591
1626
2911
1283
1322
2188
11303
2109
14439
4434
Experiment_A1
Experiment_A2
Experiment_A3
Experiment_A4
Experiment_B1
Experiment_B2
Experiment_C1
Experiment_E1
Experiment_E2
Experiment_E3
Experiment_E4

### NIC 5' reference supported (gene) Comparison

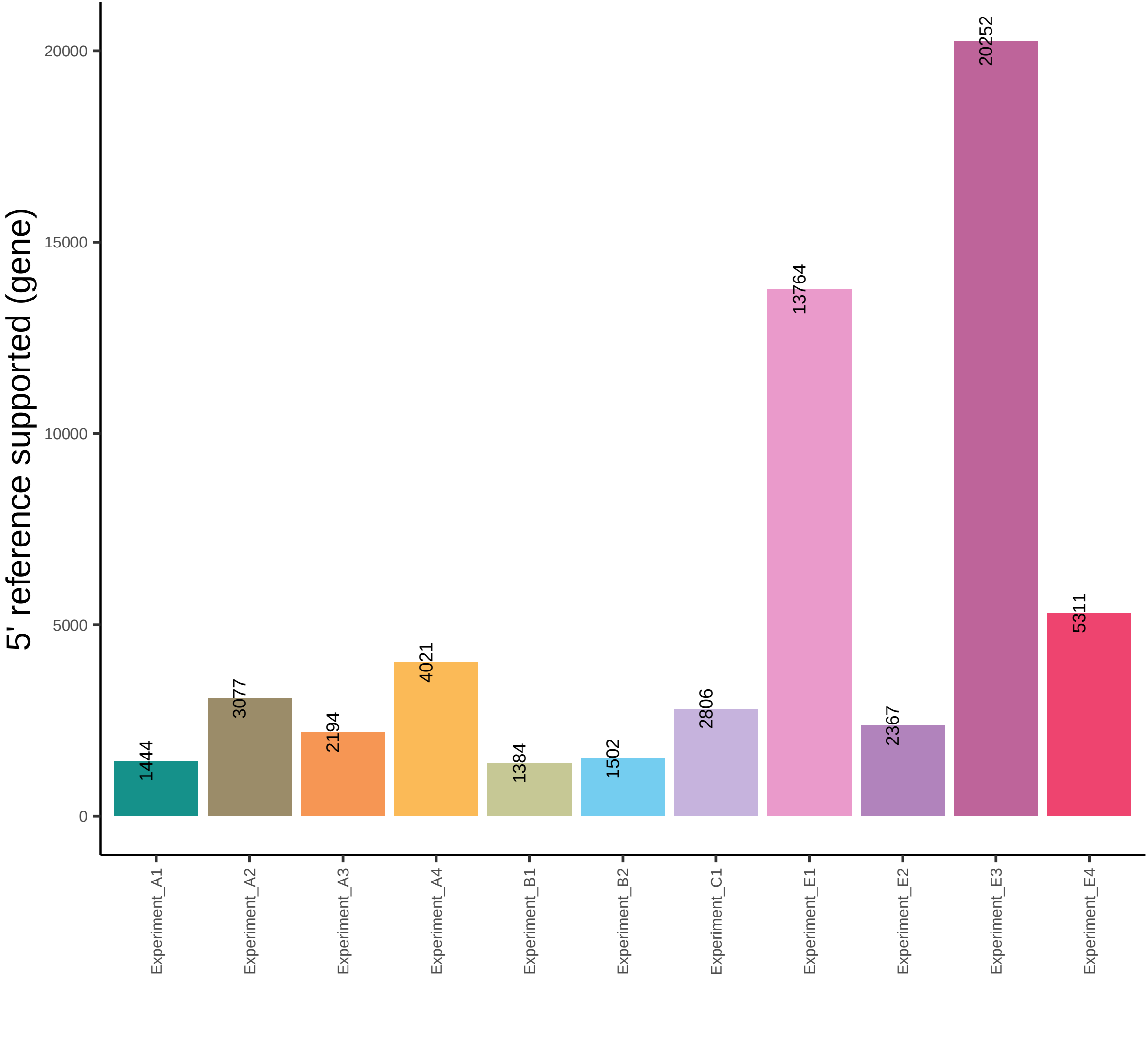

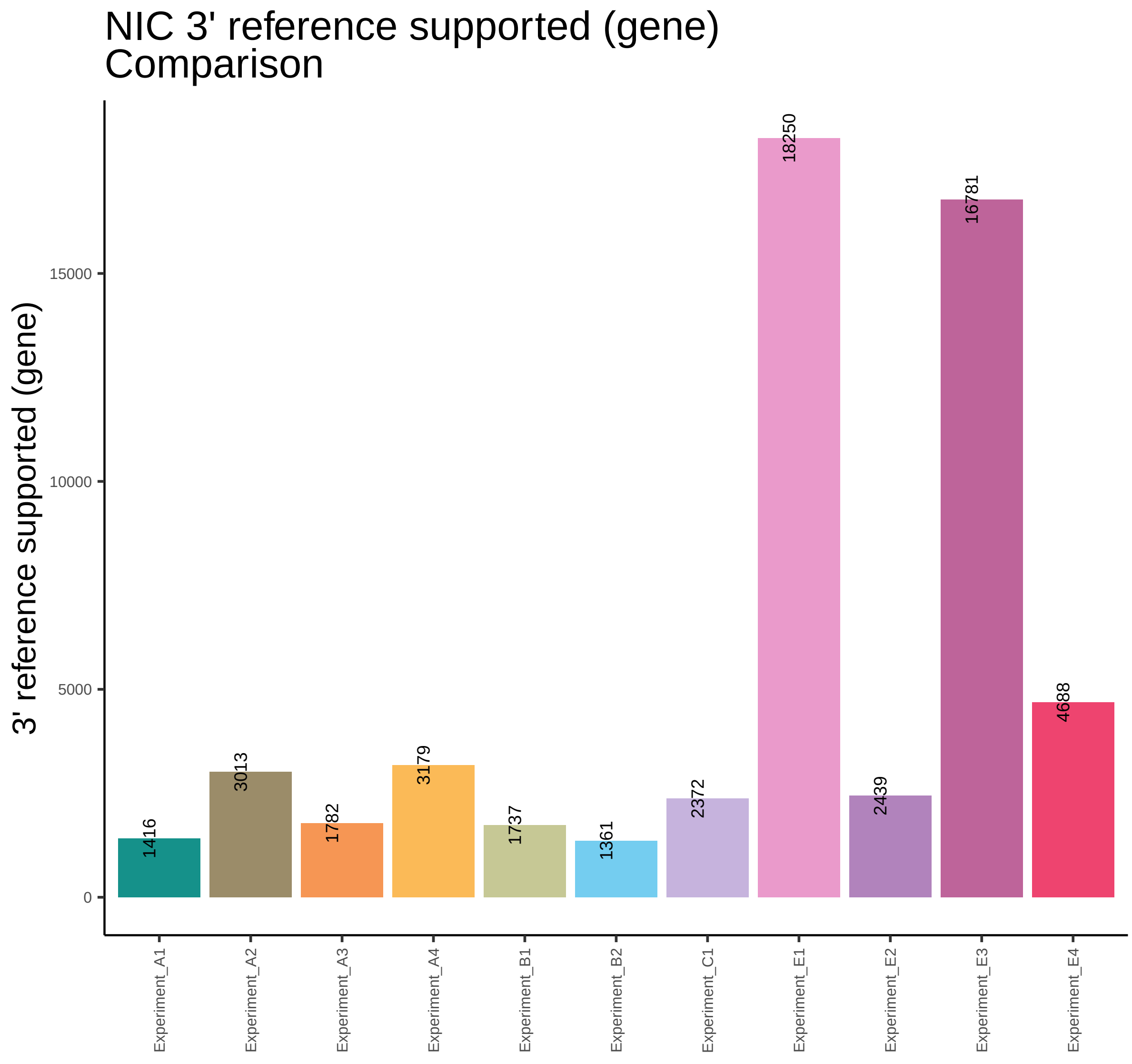
NIC 3' reference supported (gene)
Comparison
3' reference supported (gene)
0
5000
10000
15000
1416
3013
1782
3179
1737
1361
2372
18250
2439
16781
4688
Experiment_A1
Experiment_A2
Experiment_A3
Experiment_A4
Experiment_B1
Experiment_B2
Experiment_C1
Experiment_E1
Experiment_E2
Experiment_E3
Experiment_E4

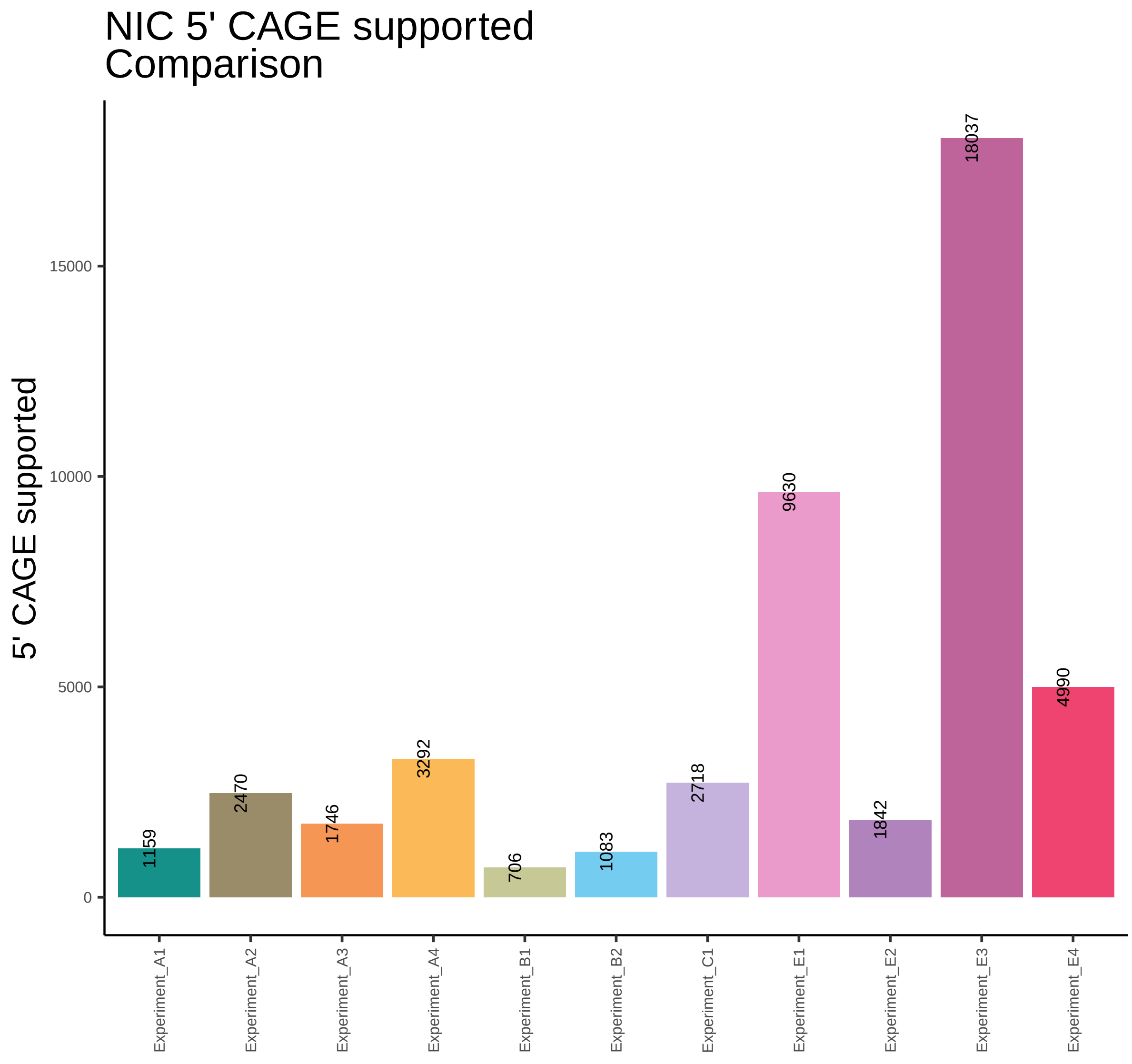
NIC 5' CAGE supported
Comparison
5' CAGE supported
15000
10000
5000
0
1159
2470
1746
3292
706
1083
2718
9630
1842
18037
4990
Experiment_A1
Experiment_A2
Experiment_A3
Experiment_A4
Experiment_B1
Experiment_B2
Experiment_C1
Experiment_E1
Experiment_E2
Experiment_E3
Experiment_E4

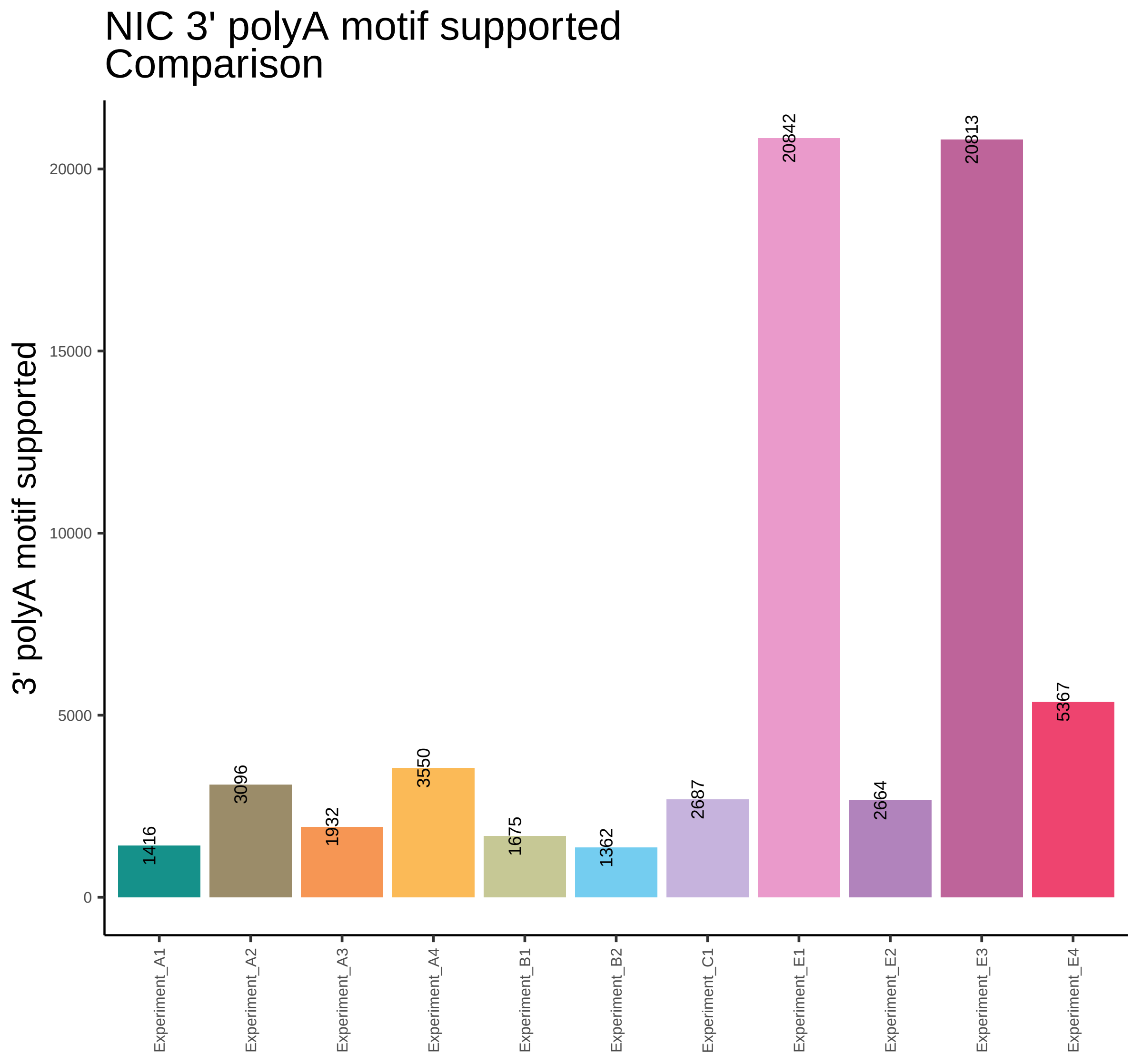

NIC 3' polyA motif supported
Comparison
3' polyA motif supported
20000
15000
10000
5000
0
1416
3096
1932
3550
1675
1362
2687
20842
2664
20813
5367
Experiment_A1
Experiment_A2
Experiment_A3
Experiment_A4
Experiment_B1
Experiment_B2
Experiment_C1
Experiment_E1
Experiment_E2
Experiment_E3
Experiment_E4

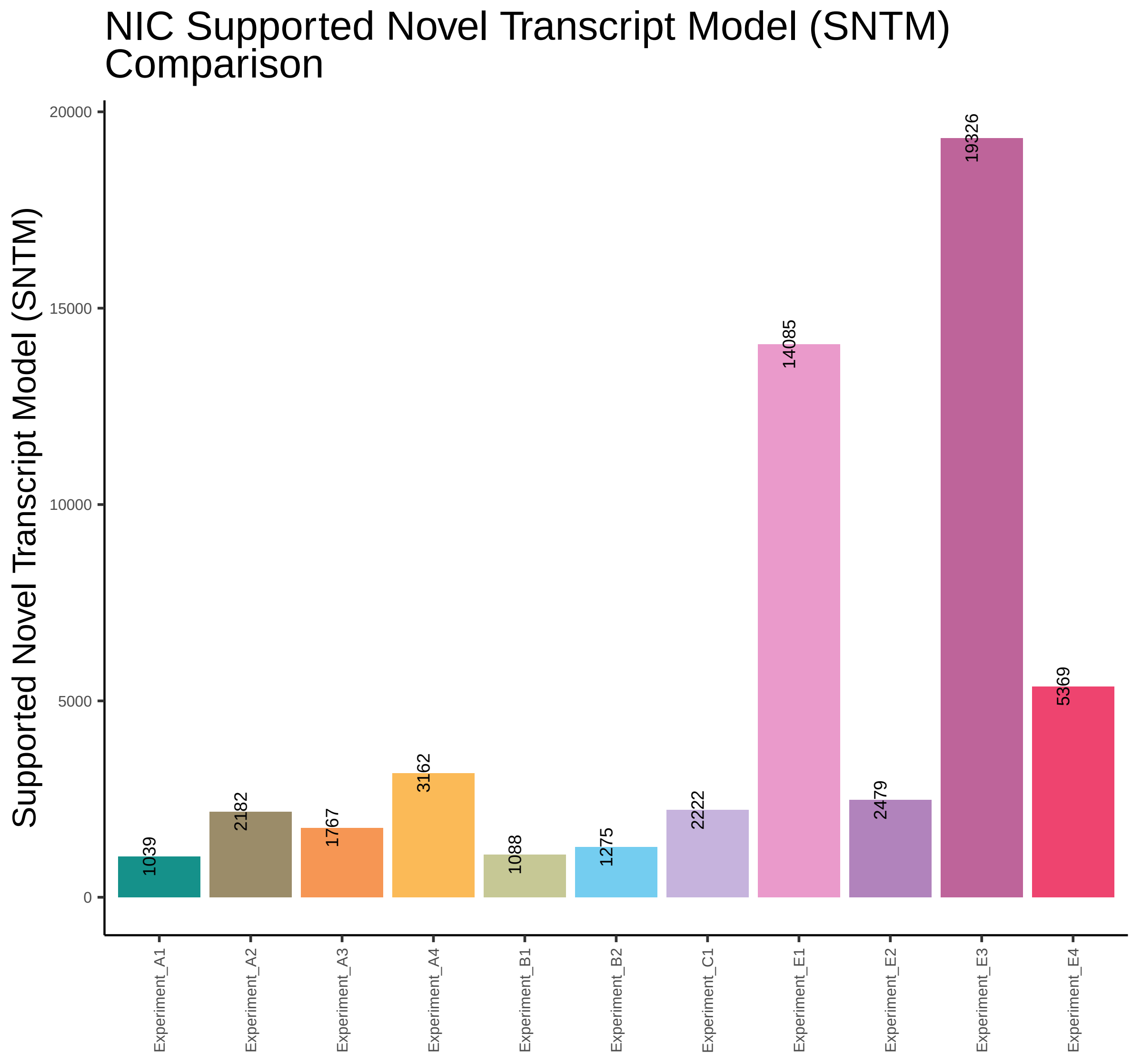

NIC Supported Novel Transcript Model (SNTM) Comparison
Supported Novel Transcript Model (SNTM)
20000
15000
10000
5000
0
1039
2182
1767
3162
1088
1275
2222
14085
2479
19326
5369
Experiment_A1
Experiment_A2
Experiment_A3
Experiment_A4
Experiment_B1
Experiment_B2
Experiment_C1
Experiment_E1
Experiment_E2
Experiment_E3
Experiment_E4

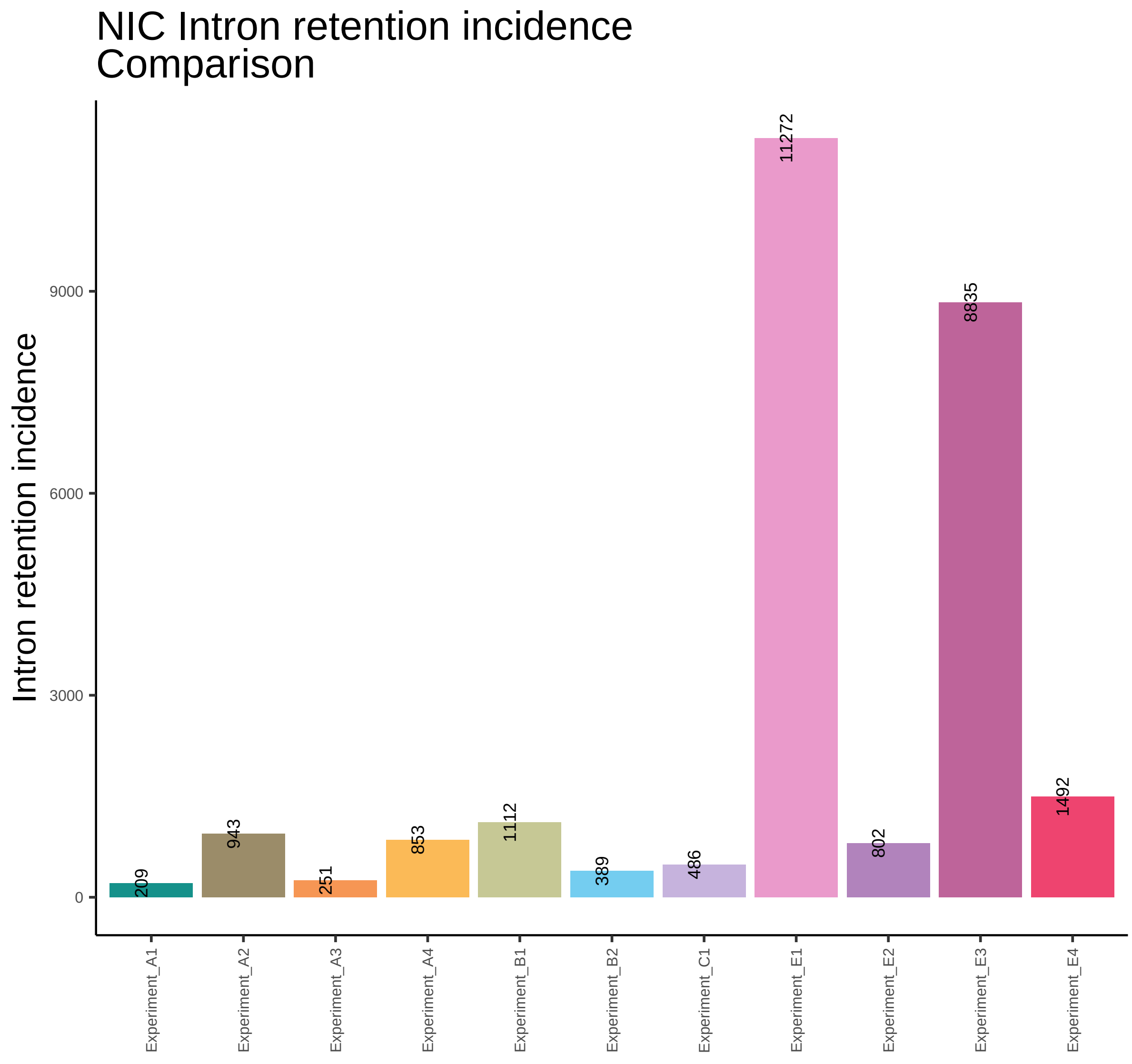

NIC Intron retention incidence
Comparison
Intron retention incidence
0
3000
6000
9000
209
943
251
853
1112
389
486
11272
802
8835
1492
Experiment_A1
Experiment_A2
Experiment_A3
Experiment_A4
Experiment_B1
Experiment_B2
Experiment_C1
Experiment_E1
Experiment_E2
Experiment_E3
Experiment_E4

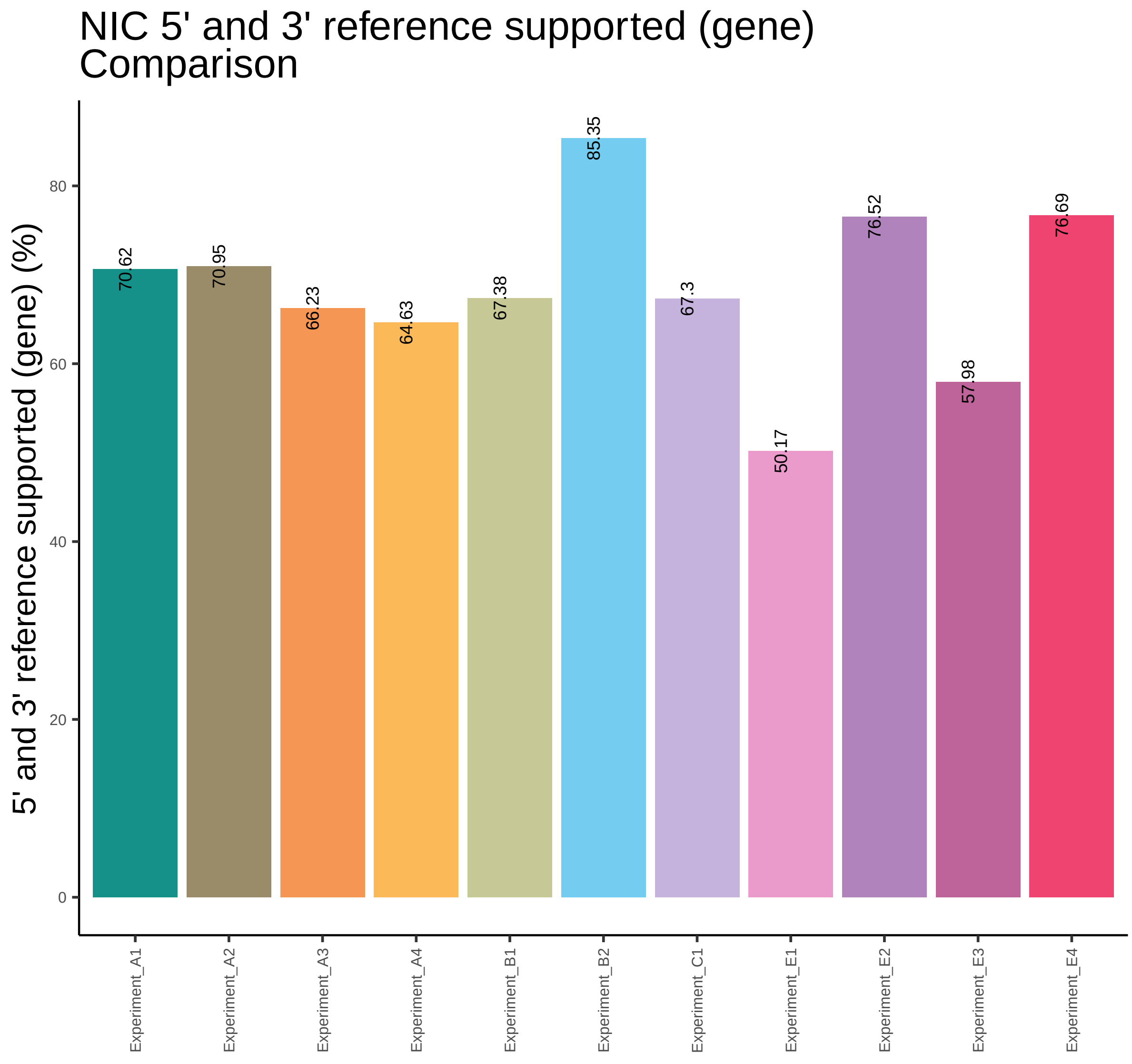
NIC 5' and 3' reference supported (gene)
Comparison
5' and 3' reference supported (gene) (%)
0
20
40
60
80
70.62
70.95
66.23
64.63
67.38
85.35
67.3
50.17
76.52
57.98
76.69
Experiment_A1
Experiment_A2
Experiment_A3
Experiment_A4
Experiment_B1
Experiment_B2
Experiment_C1
Experiment_E1
Experiment_E2
Experiment_E3
Experiment_E4

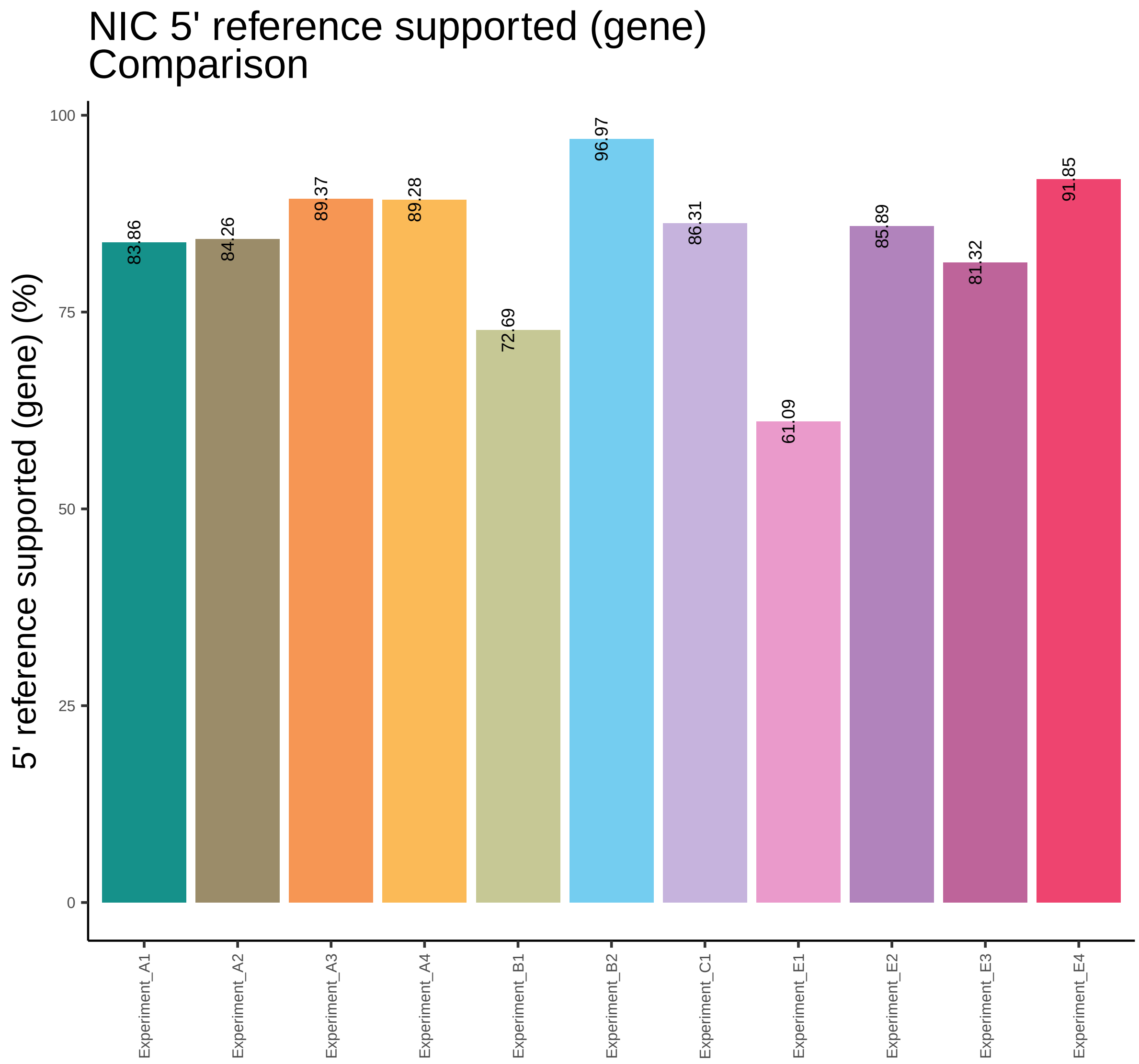
NIC 5' reference supported (gene)
Comparison
5' reference supported (gene) (%)
100
75
50
25
0
83.86
84.26
89.37
89.28
72.69
96.97
86.31
61.09
85.89
81.32
91.85
Experiment_A1
Experiment_A2
Experiment_A3
Experiment_A4
Experiment_B1
Experiment_B2
Experiment_C1
Experiment_E1
Experiment_E2
Experiment_E3
Experiment_E4

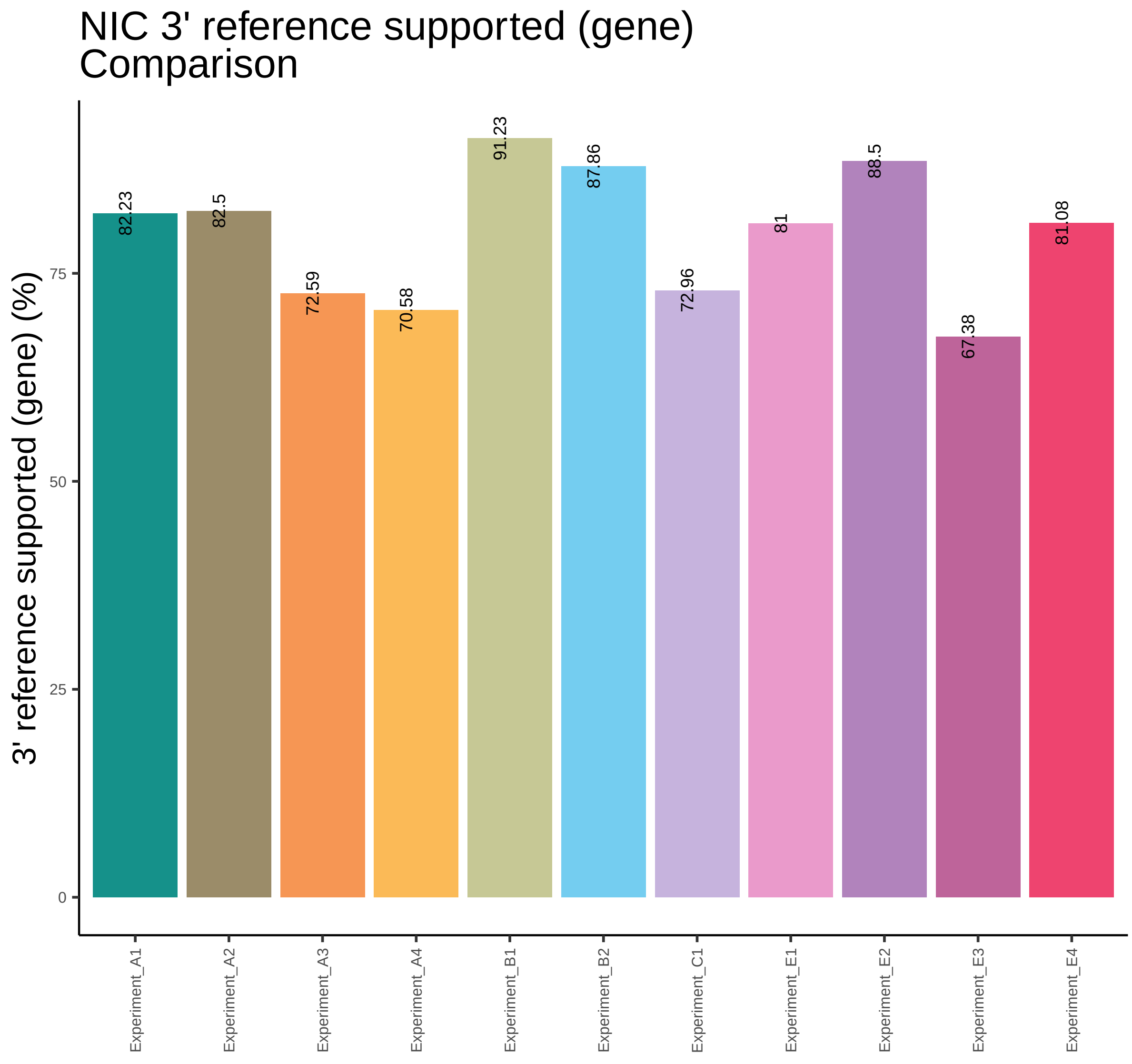

NIC 3' reference supported (gene)
Comparison
3' reference supported (gene) (%)
0
25
50
75
82.23
82.5
72.59
70.58
91.23
87.86
72.96
81
88.5
67.38
81.08
Experiment_A1
Experiment_A2
Experiment_A3
Experiment_A4
Experiment_B1
Experiment_B2
Experiment_C1
Experiment_E1
Experiment_E2
Experiment_E3
Experiment_E4

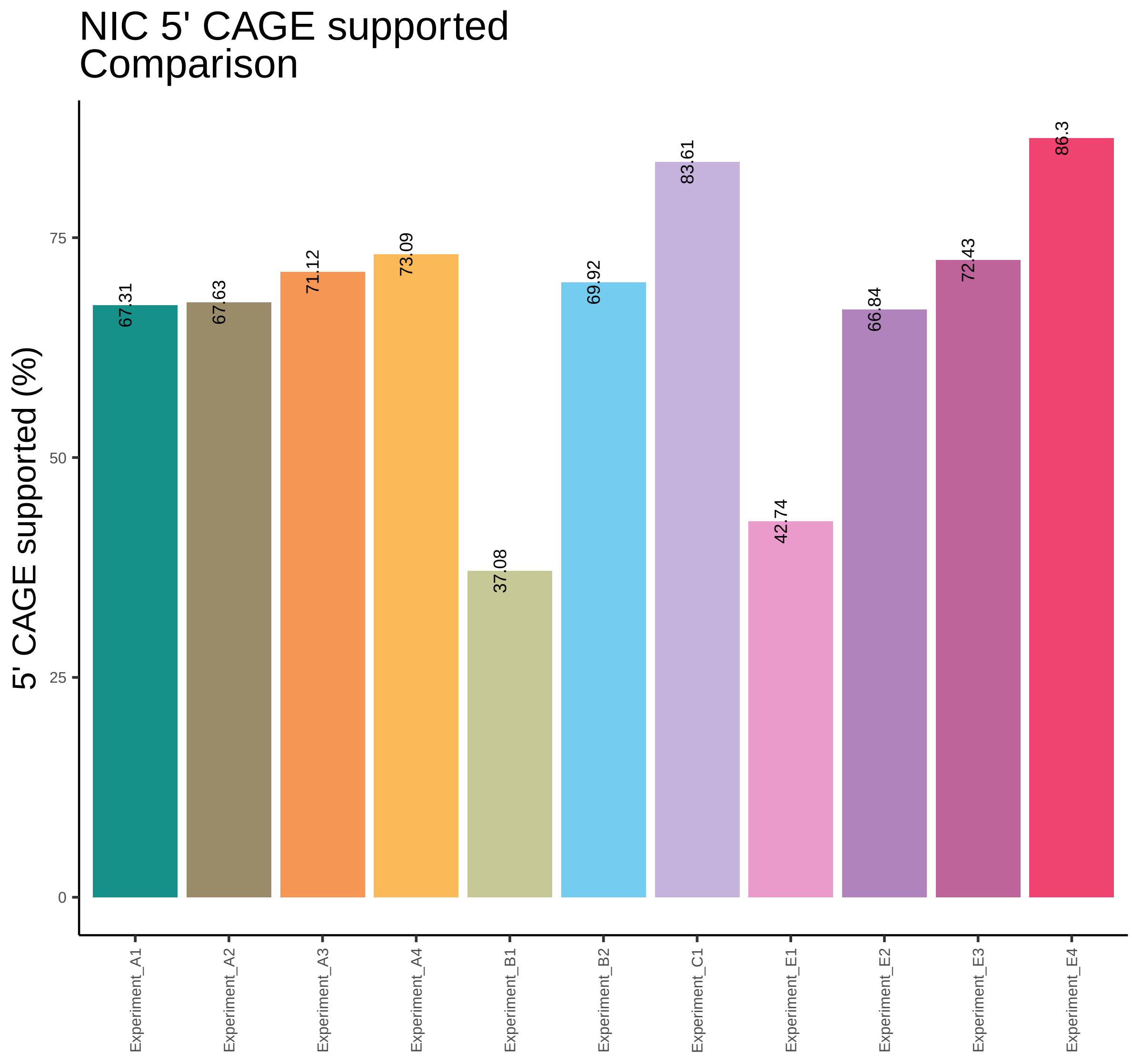

NIC 5' CAGE supported
Comparison
5' CAGE supported (%)
0
25
50
75
67.31
67.63
71.12
73.09
37.08
69.92
83.61
42.74
66.84
72.43
86.3
Experiment_A1
Experiment_A2
Experiment_A3
Experiment_A4
Experiment_B1
Experiment_B2
Experiment_C1
Experiment_E1
Experiment_E2
Experiment_E3
Experiment_E4

### NIC 3' polyA motif supported Comparison

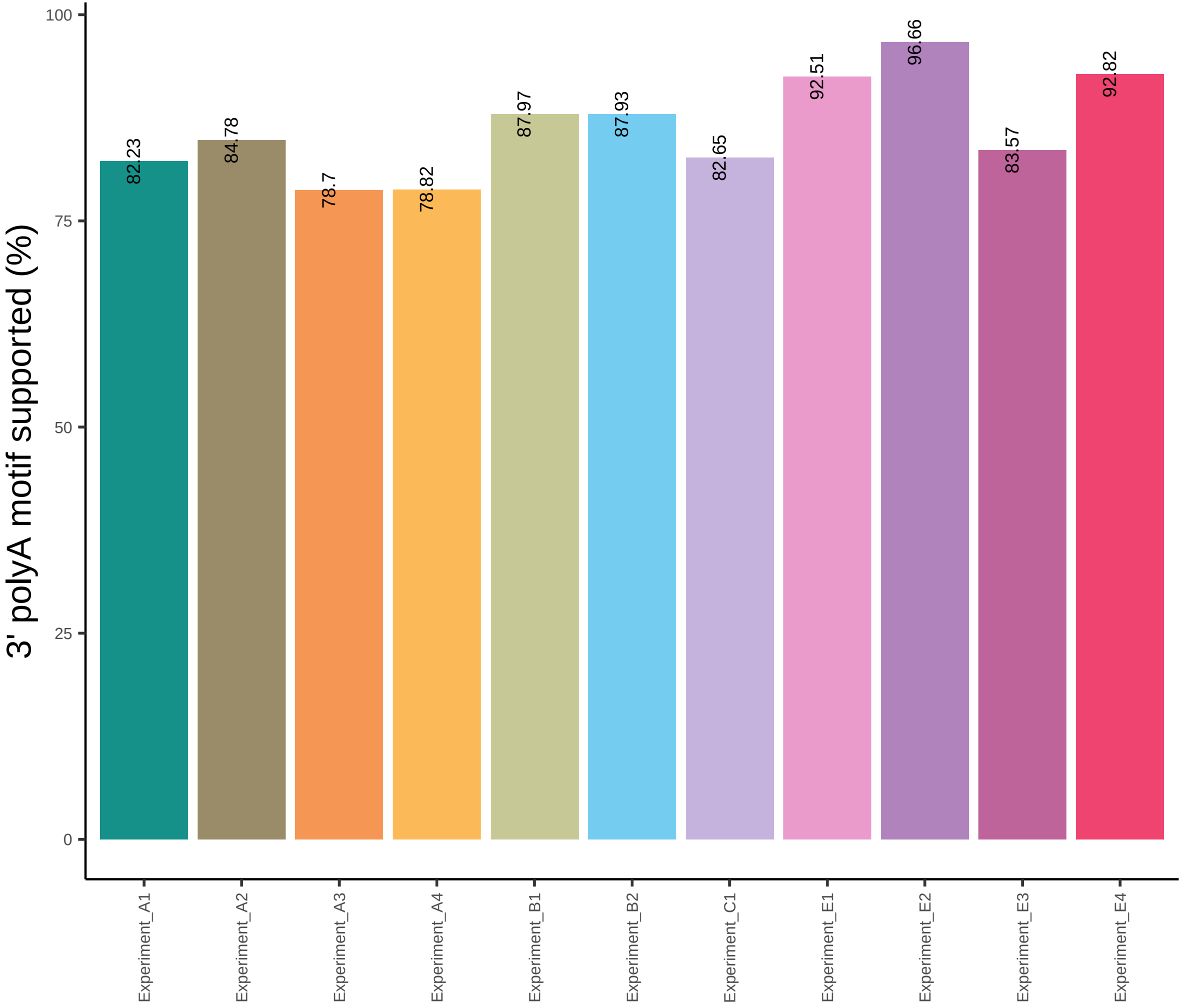

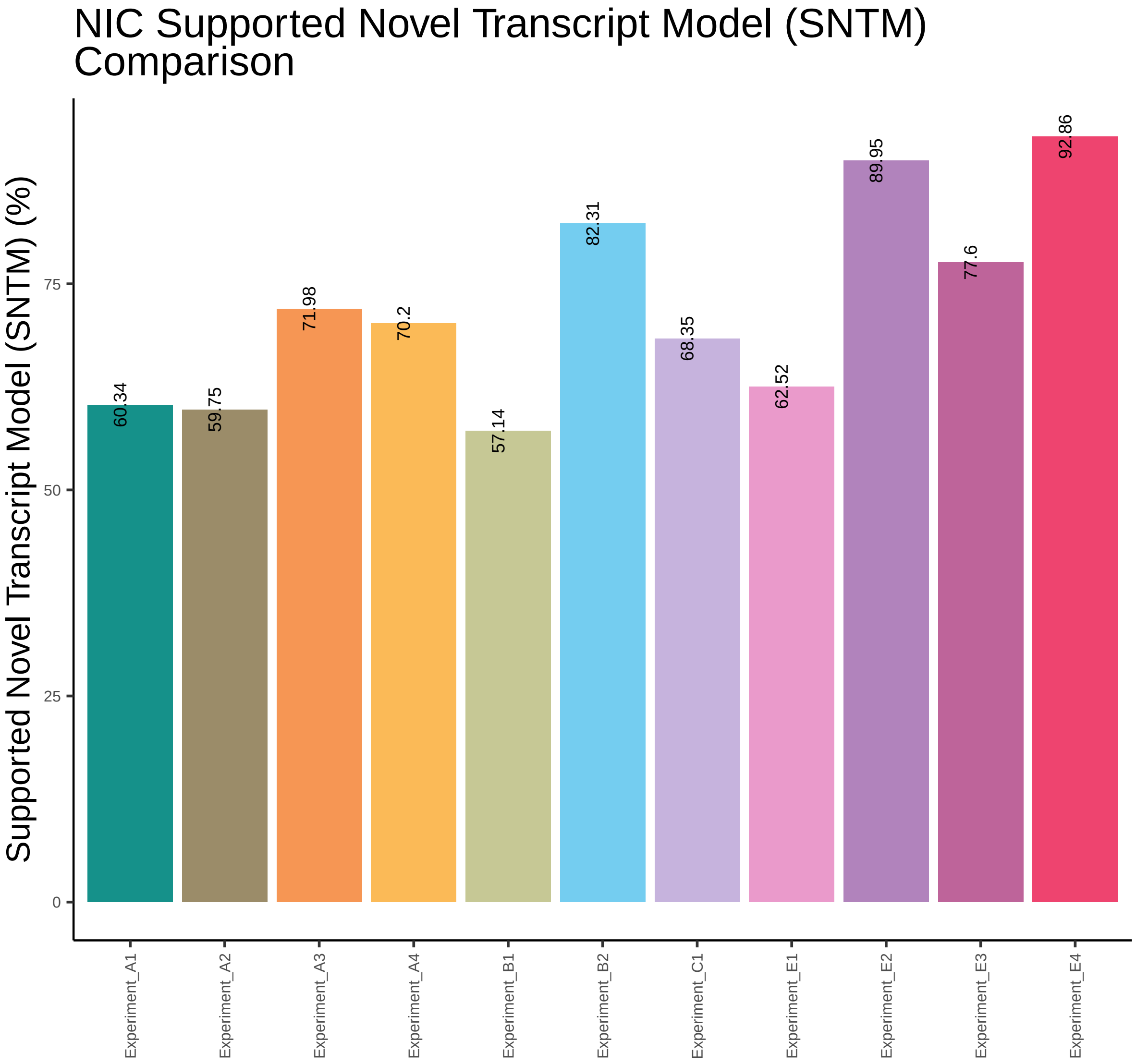

NIC Supported Novel Transcript Model (SNTM) Comparison
Supported Novel Transcript Model (SNTM) (%)
0
25
50
75
60.34
59.75
71.98
70.2
57.14
82.31
68.35
62.52
89.95
77.6
92.86
Experiment_A1
Experiment_A2
Experiment_A3
Experiment_A4
Experiment_B1
Experiment_B2
Experiment_C1
Experiment_E1
Experiment_E2
Experiment_E3
Experiment_E4

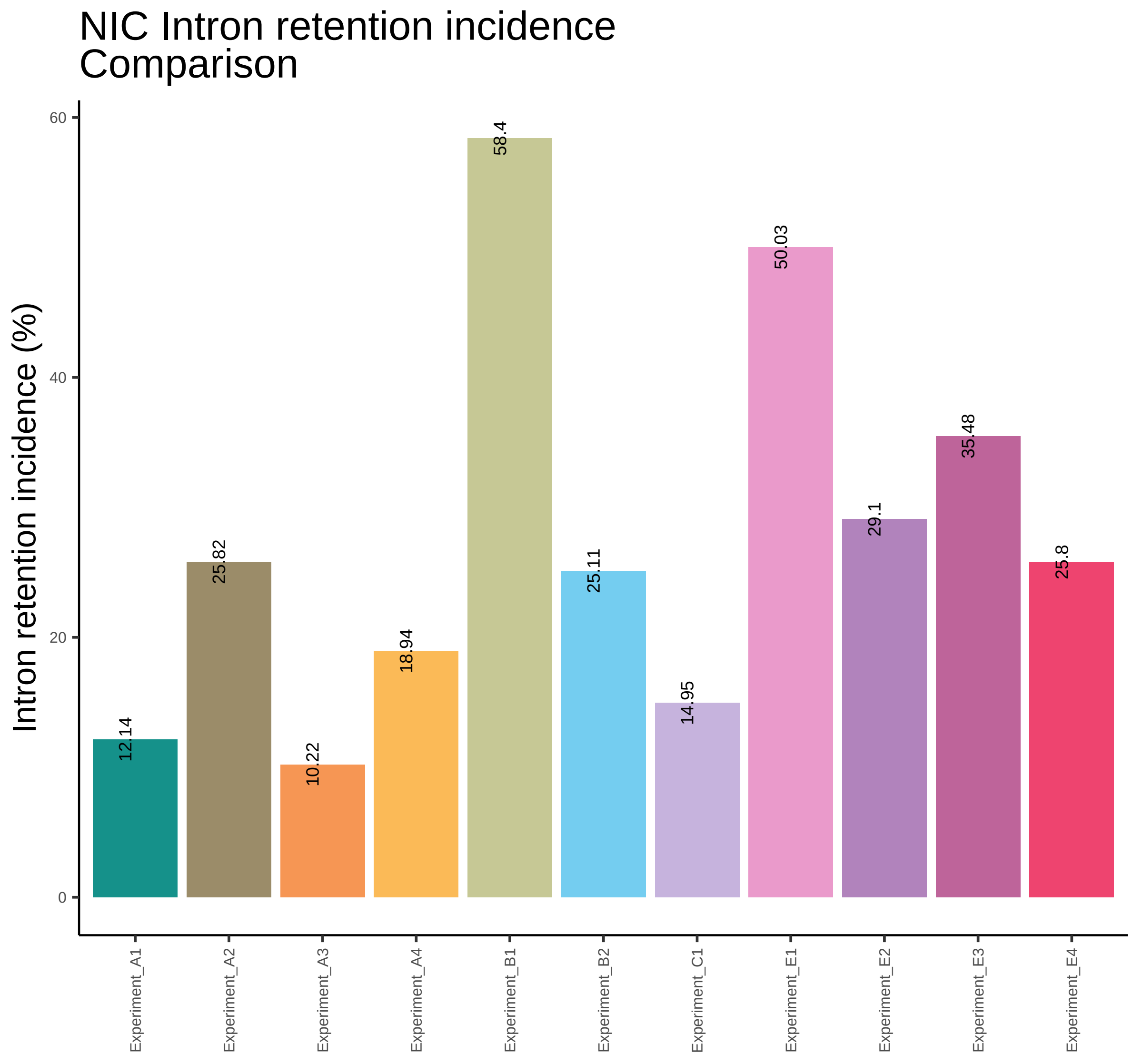

NIC Intron retention incidence
Comparison
Intron retention incidence (%)
0
20
40
60
12.14
25.82
10.22
18.94
58.4
25.11
14.95
50.03
29.1
35.48
25.8
Experiment_A1
Experiment_A2
Experiment_A3
Experiment_A4
Experiment_B1
Experiment_B2
Experiment_C1
Experiment_E1
Experiment_E2
Experiment_E3
Experiment_E4

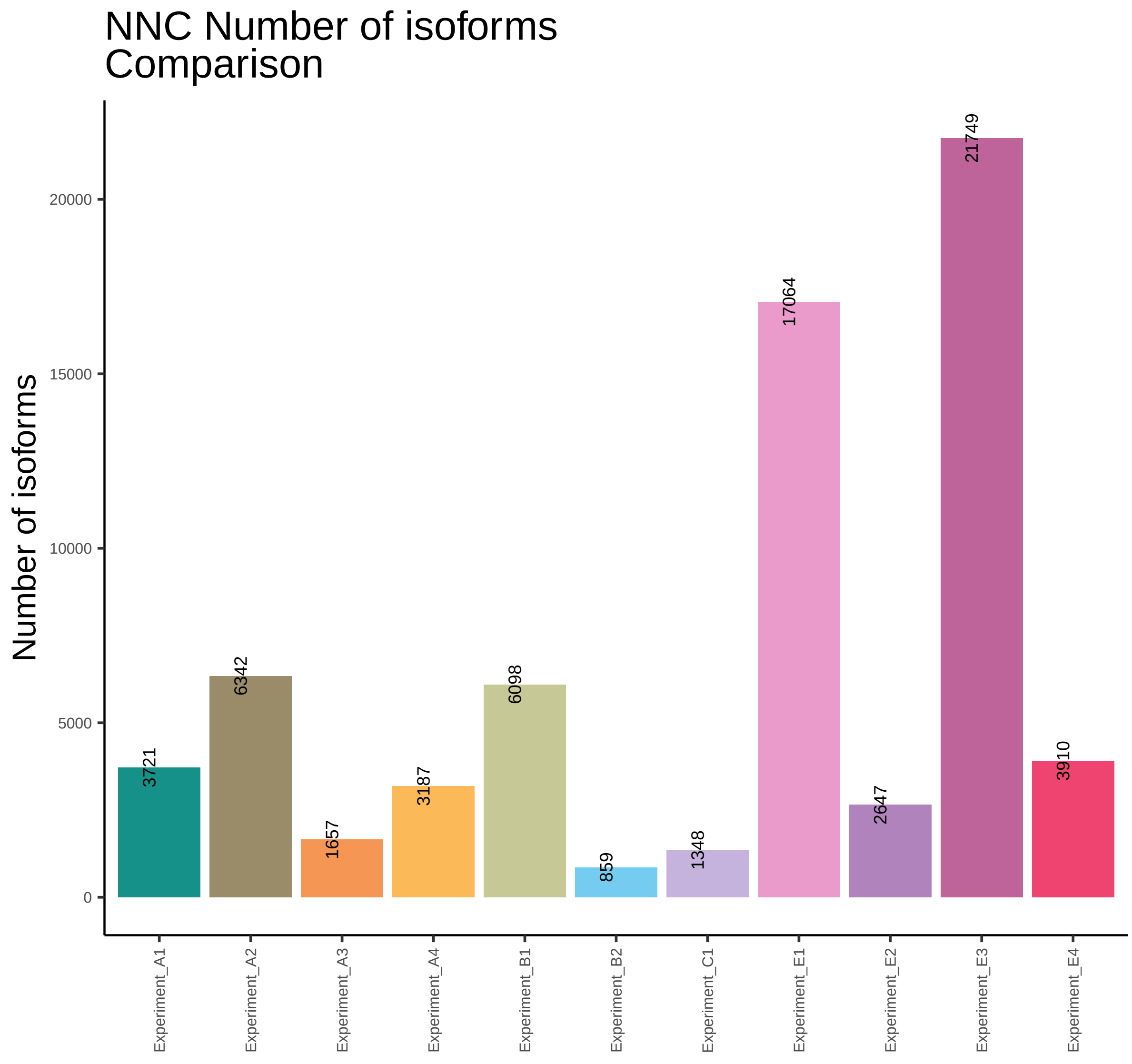
NNC Number of isoforms
Comparison
Number of isoforms
0
5000
10000
15000
20000
3721
6342
1657
3187
6098
859
1348
17064
2647
21749
3910
Experiment_A1
Experiment_A2
Experiment_A3
Experiment_A4
Experiment_B1
Experiment_B2
Experiment_C1
Experiment_E1
Experiment_E2
Experiment_E3
Experiment_E4

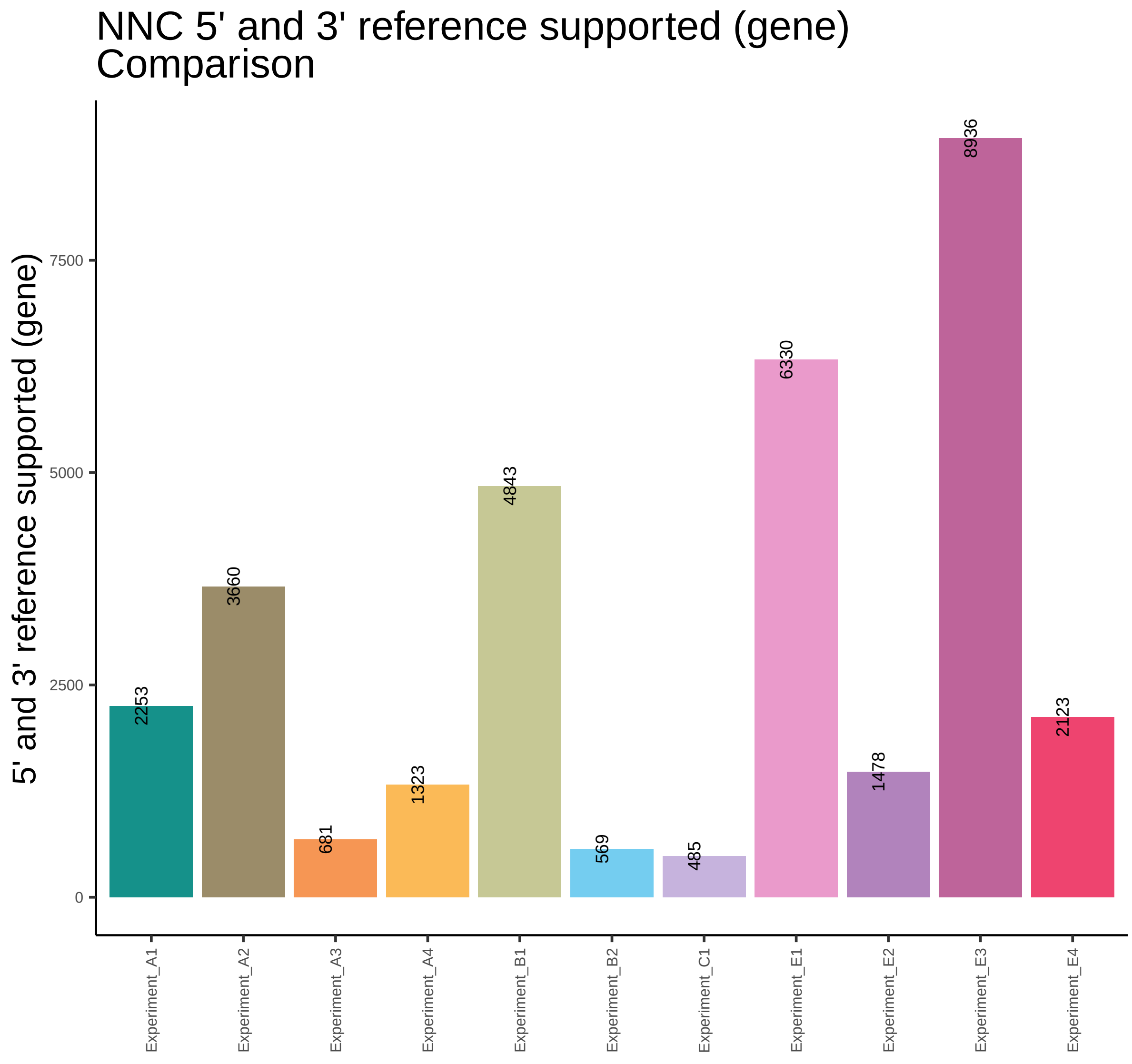
NNC 5' and 3' reference supported (gene)
Comparison
5' and 3' reference supported (gene)
0
2500
5000
7500
2253
3660
681
1323
4843
569
485
6330
1478
8936
2123
Experiment_A1
Experiment_A2
Experiment_A3
Experiment_A4
Experiment_B1
Experiment_B2
Experiment_C1
Experiment_E1
Experiment_E2
Experiment_E3
Experiment_E4

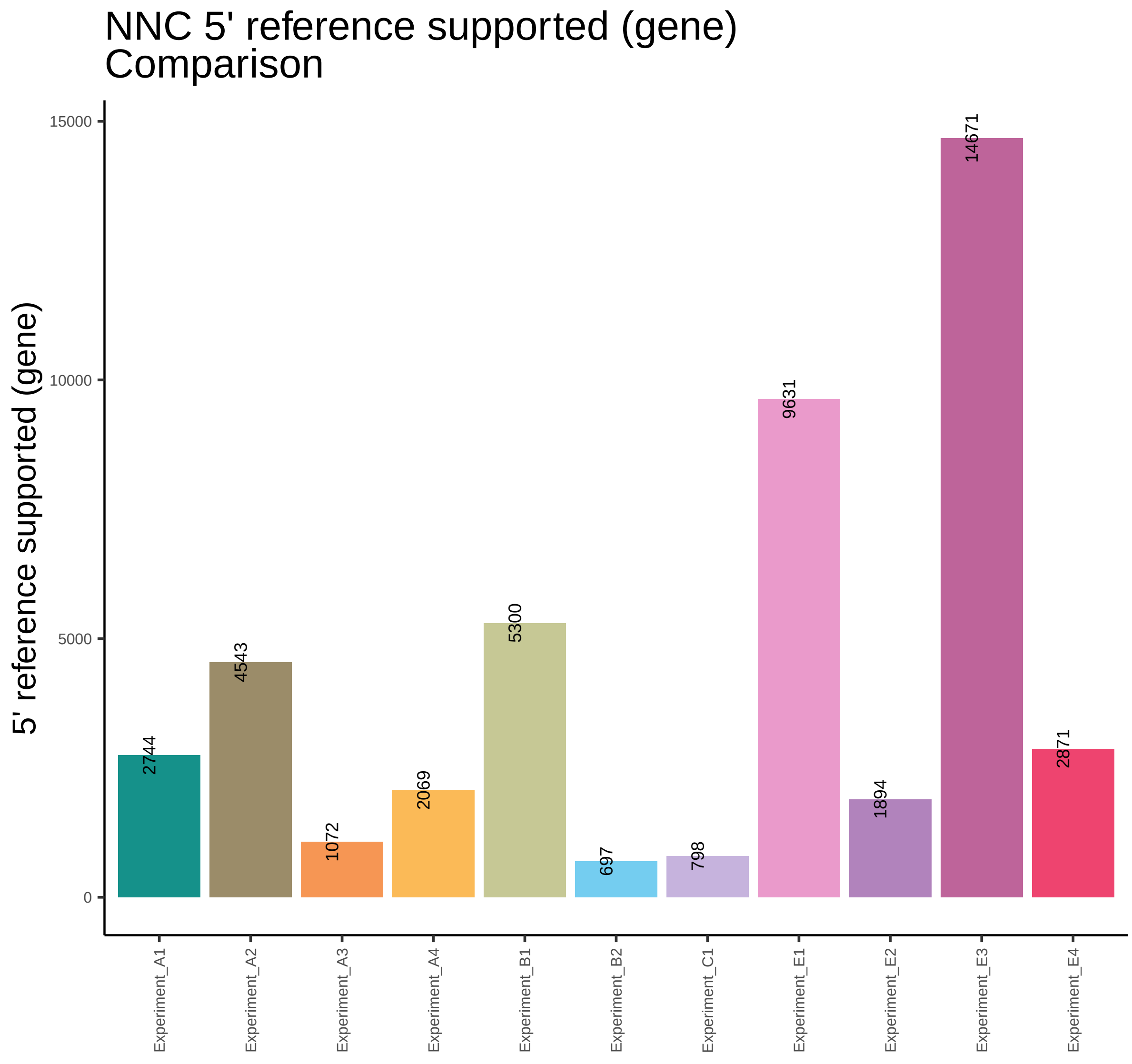
NNC 5' reference supported (gene)
Comparison
5' reference supported (gene)
15000
10000
5000
0
2744
4543
1072
2069
5300
697
798
9631
1894
14671
2871
Experiment_A1
Experiment_A2
Experiment_A3
Experiment_A4
Experiment_B1
Experiment_B2
Experiment_C1
Experiment_E1
Experiment_E2
Experiment_E3
Experiment_E4

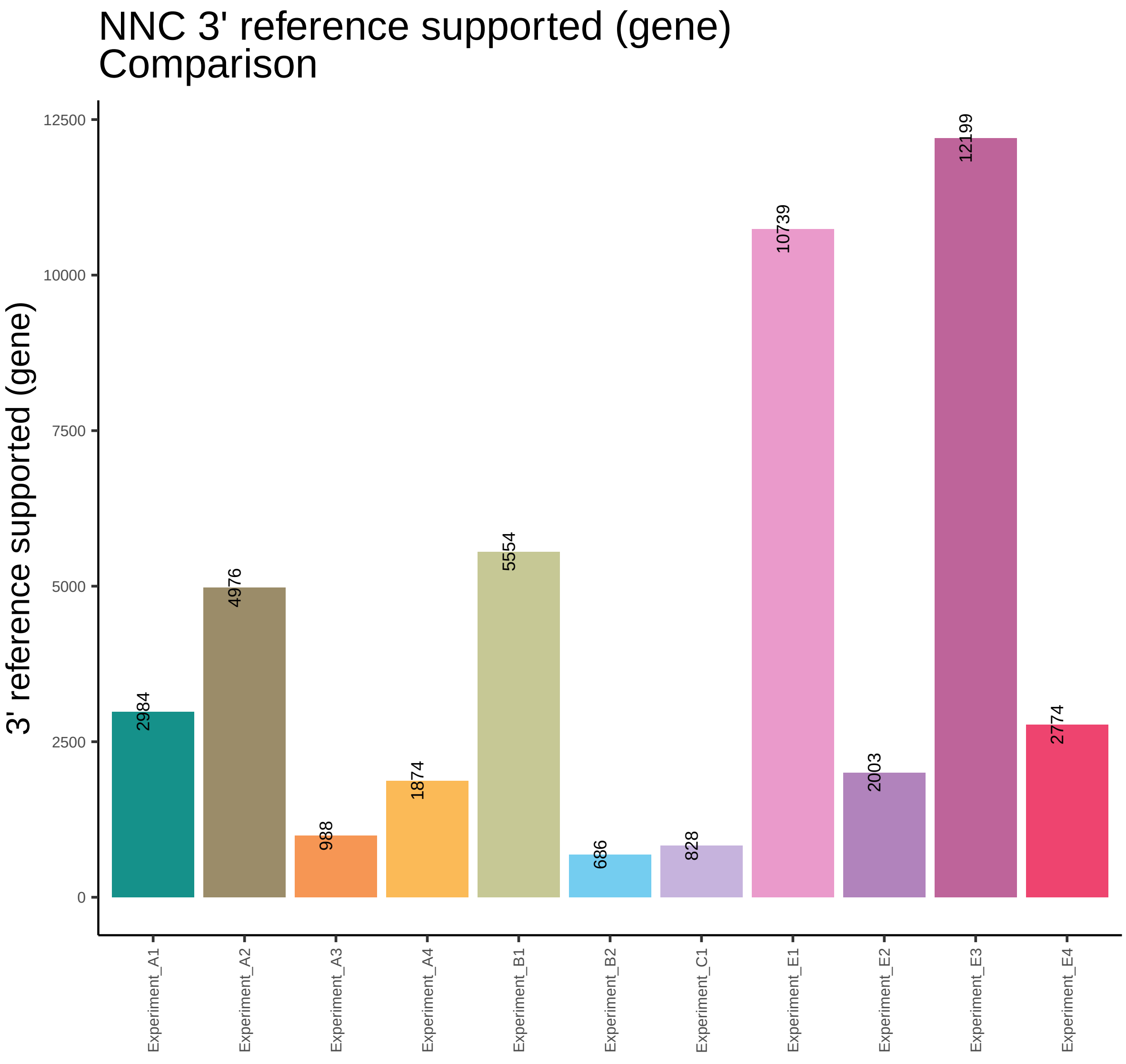
NNC 3' reference supported (gene)
Comparison
3' reference supported (gene)
12500
10000
7500
5000
2500
0
2984
4976
988
1874
5554
686
828
10739
2003
12199
2774
Experiment_A1
Experiment_A2
Experiment_A3
Experiment_A4
Experiment_B1
Experiment_B2
Experiment_C1
Experiment_E1
Experiment_E2
Experiment_E3
Experiment_E4

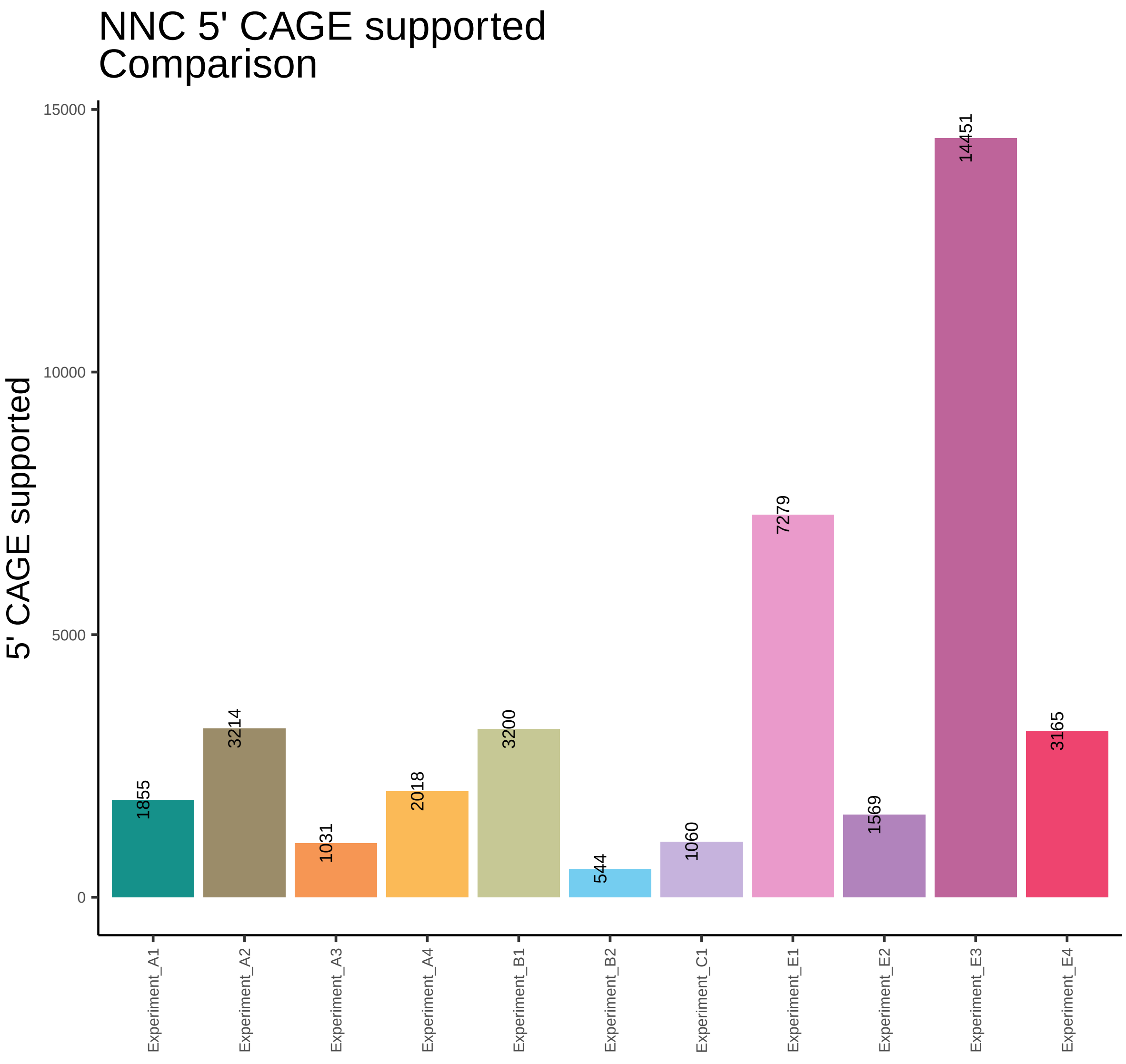
NNC 5' CAGE supported
Comparison
5' CAGE supported
15000
10000
5000
0
1855
3214
1031
2018
3200
544
1060
7279
1569
14451
3165
Experiment_A1
Experiment_A2
Experiment_A3
Experiment_A4
Experiment_B1
Experiment_B2
Experiment_C1
Experiment_E1
Experiment_E2
Experiment_E3
Experiment_E4

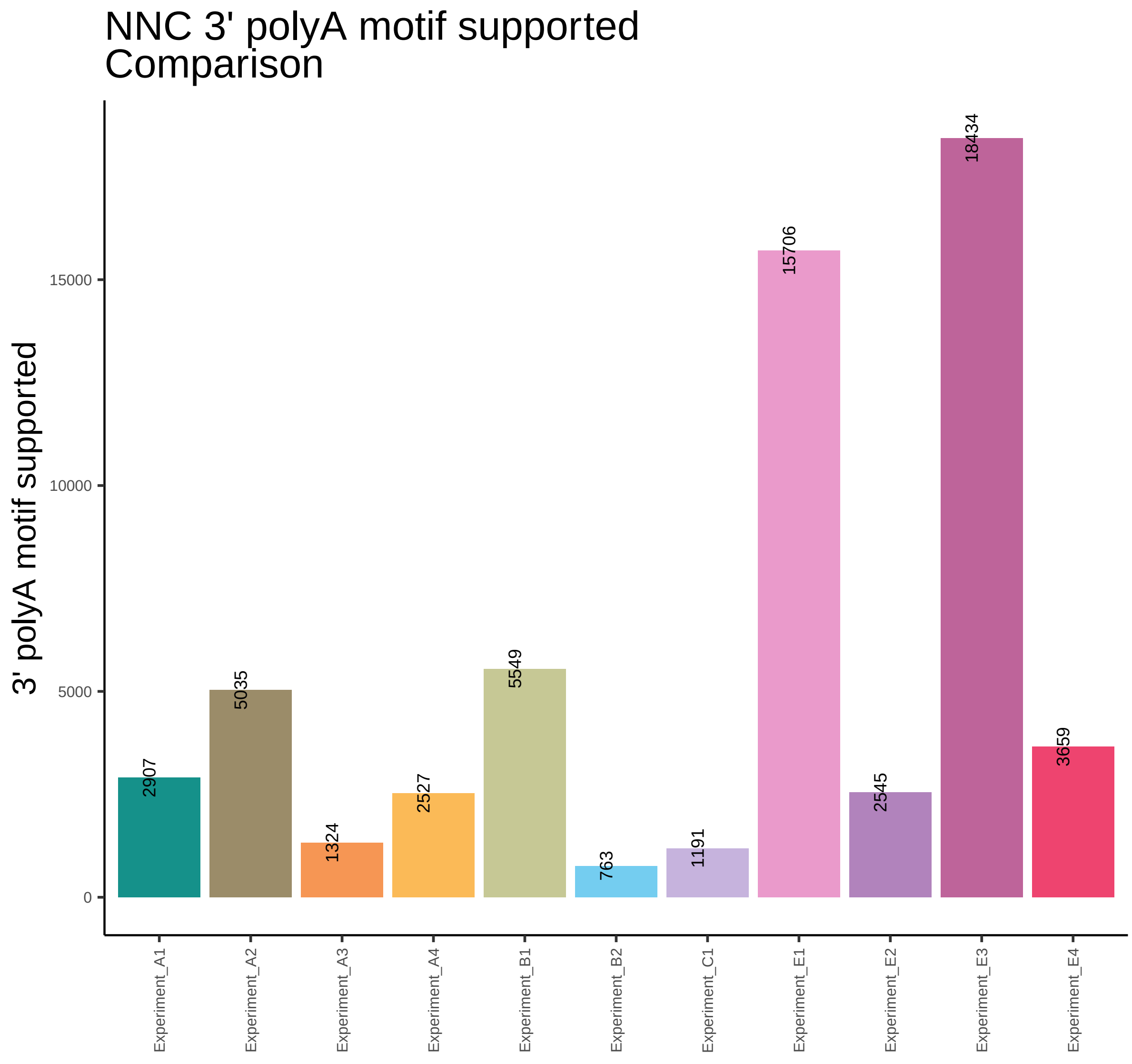
NNC 3' polyA motif supported
Comparison
3' polyA motif supported
0
5000
10000
15000
2907
5035
1324
2527
5549
763
1191
15706
2545
18434
3659
Experiment_A1
Experiment_A2
Experiment_A3
Experiment_A4
Experiment_B1
Experiment_B2
Experiment_C1
Experiment_E1
Experiment_E2
Experiment_E3
Experiment_E4

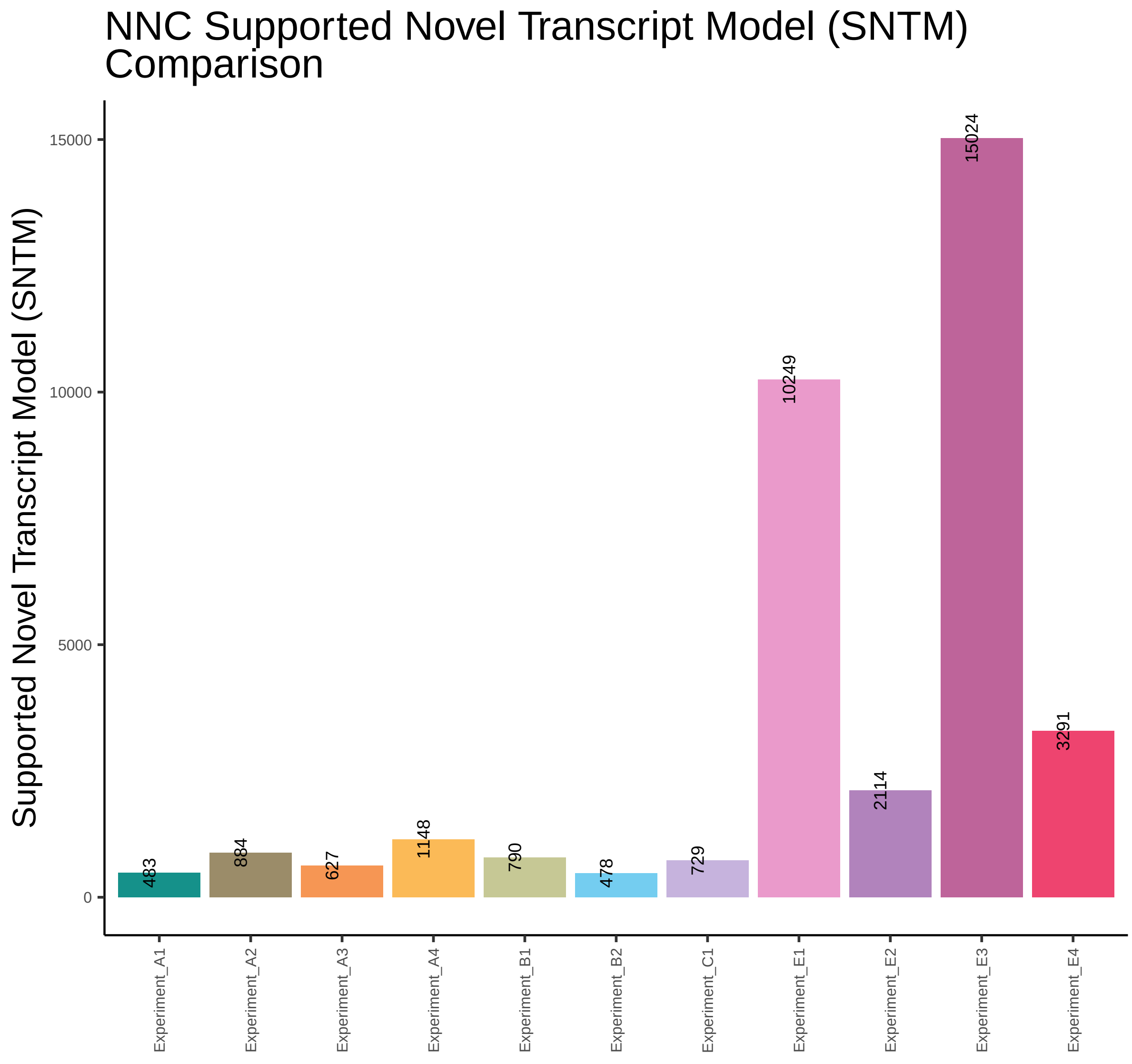
NNC Supported Novel Transcript Model (SNTM) Comparison
Supported Novel Transcript Model (SNTM)
15000
10000
5000
0
483
884
627
1148
790
478
729
10249
2114
15024
3291
Experiment_A1
Experiment_A2
Experiment_A3
Experiment_A4
Experiment_B1
Experiment_B2
Experiment_C1
Experiment_E1
Experiment_E2
Experiment_E3
Experiment_E4

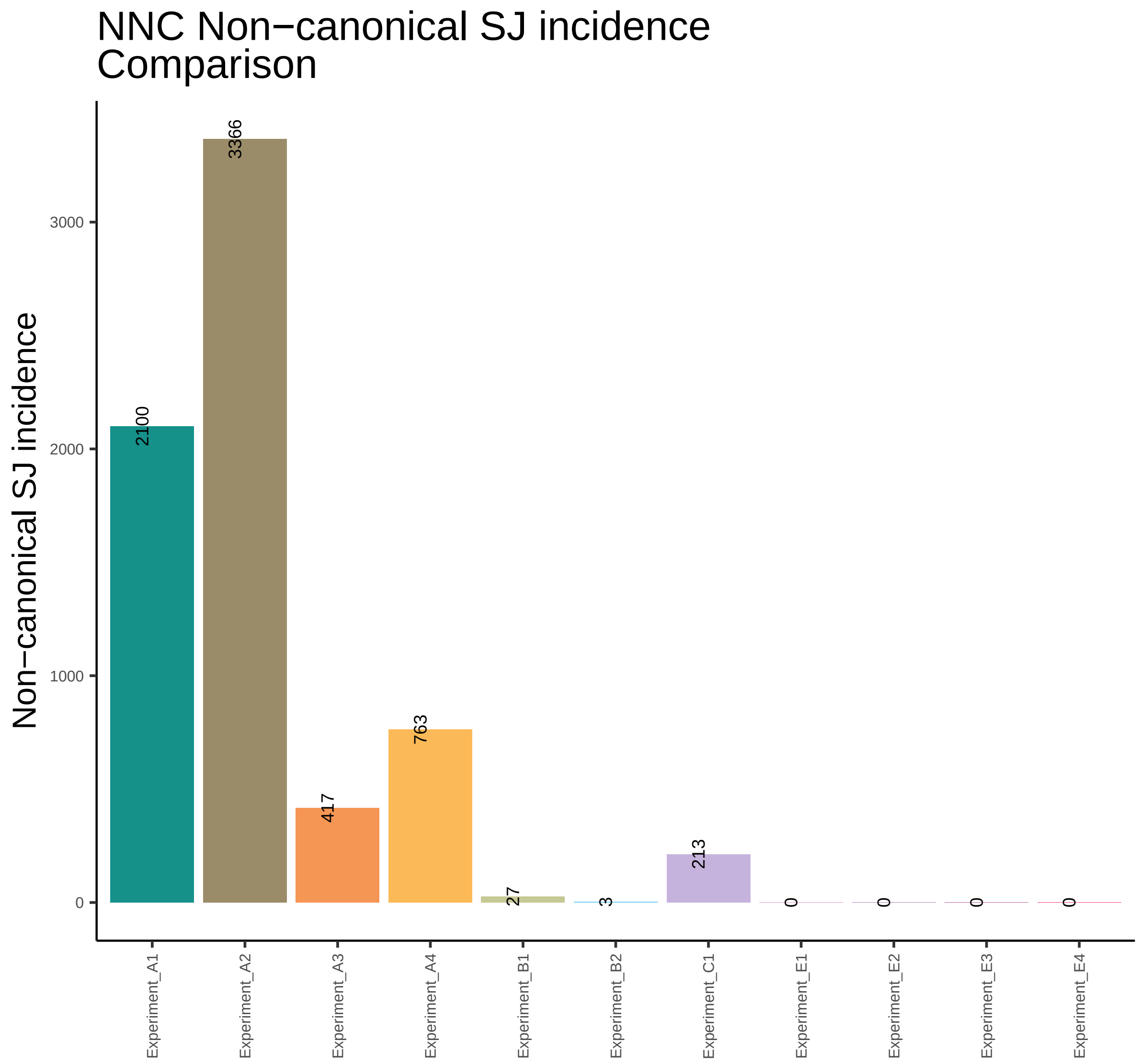

NNC Non−canonical SJ incidence
Comparison
Non−canonical SJ incidence
3000
2000
1000
0
3366
2100
417
763
27
3
213
0
0
0
0
Experiment_A1
Experiment_A2
Experiment_A3
Experiment_A4
Experiment_B1
Experiment_B2
Experiment_C1
Experiment_E1
Experiment_E2
Experiment_E3
Experiment_E4

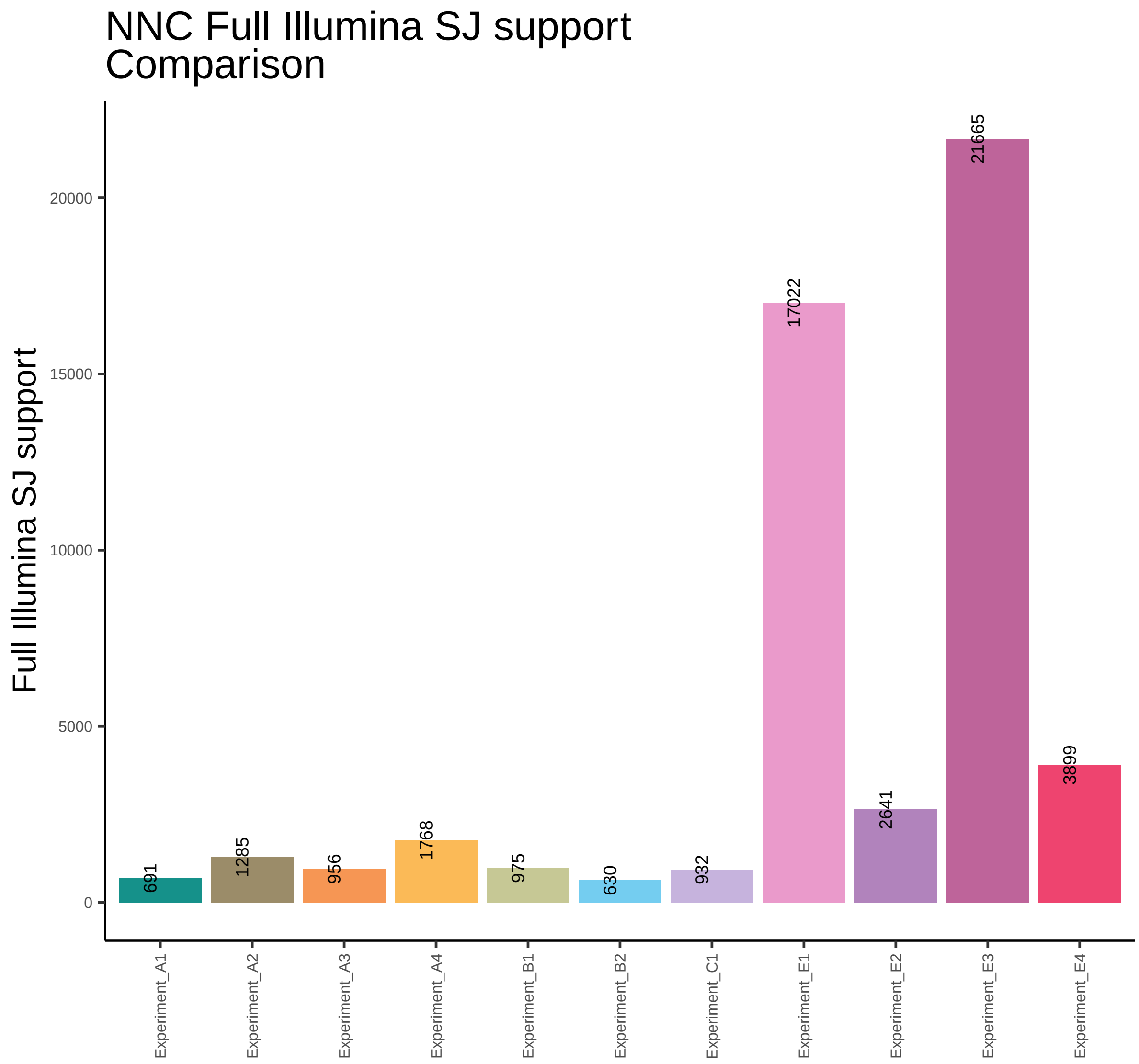
NNC Full Illumina SJ support
Comparison
Full Illumina SJ support
0
5000
10000
15000
20000
691
1285
956
1768
975
630
932
17022
2641
21665
3899
Experiment_A1
Experiment_A2
Experiment_A3
Experiment_A4
Experiment_B1
Experiment_B2
Experiment_C1
Experiment_E1
Experiment_E2
Experiment_E3
Experiment_E4

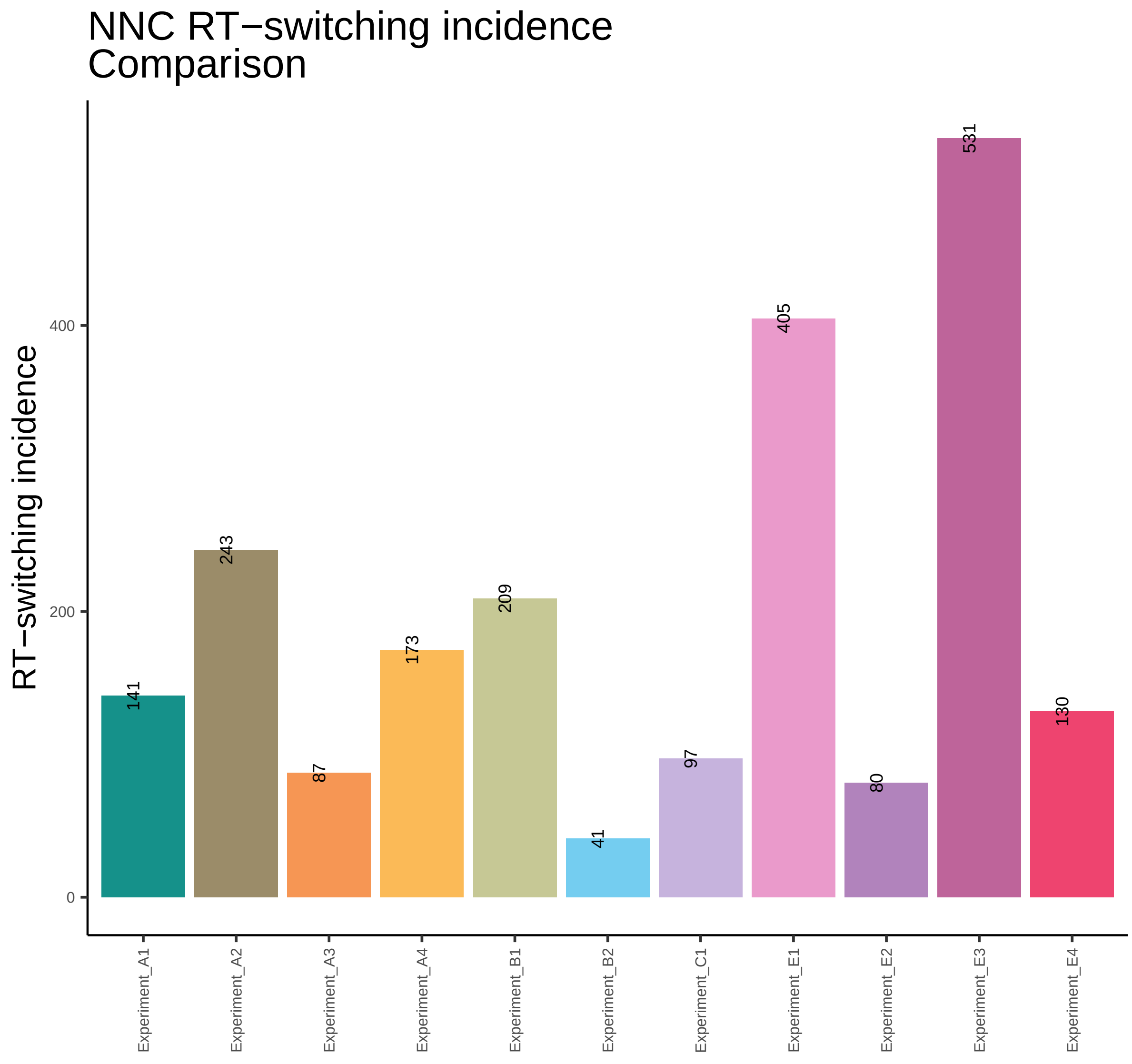

NNC RT−switching incidence
Comparison
RT−switching incidence
0
200
400
141
243
87
173
209
41
97
405
80
531
130
Experiment_A1
Experiment_A2
Experiment_A3
Experiment_A4
Experiment_B1
Experiment_B2
Experiment_C1
Experiment_E1
Experiment_E2
Experiment_E3
Experiment_E4

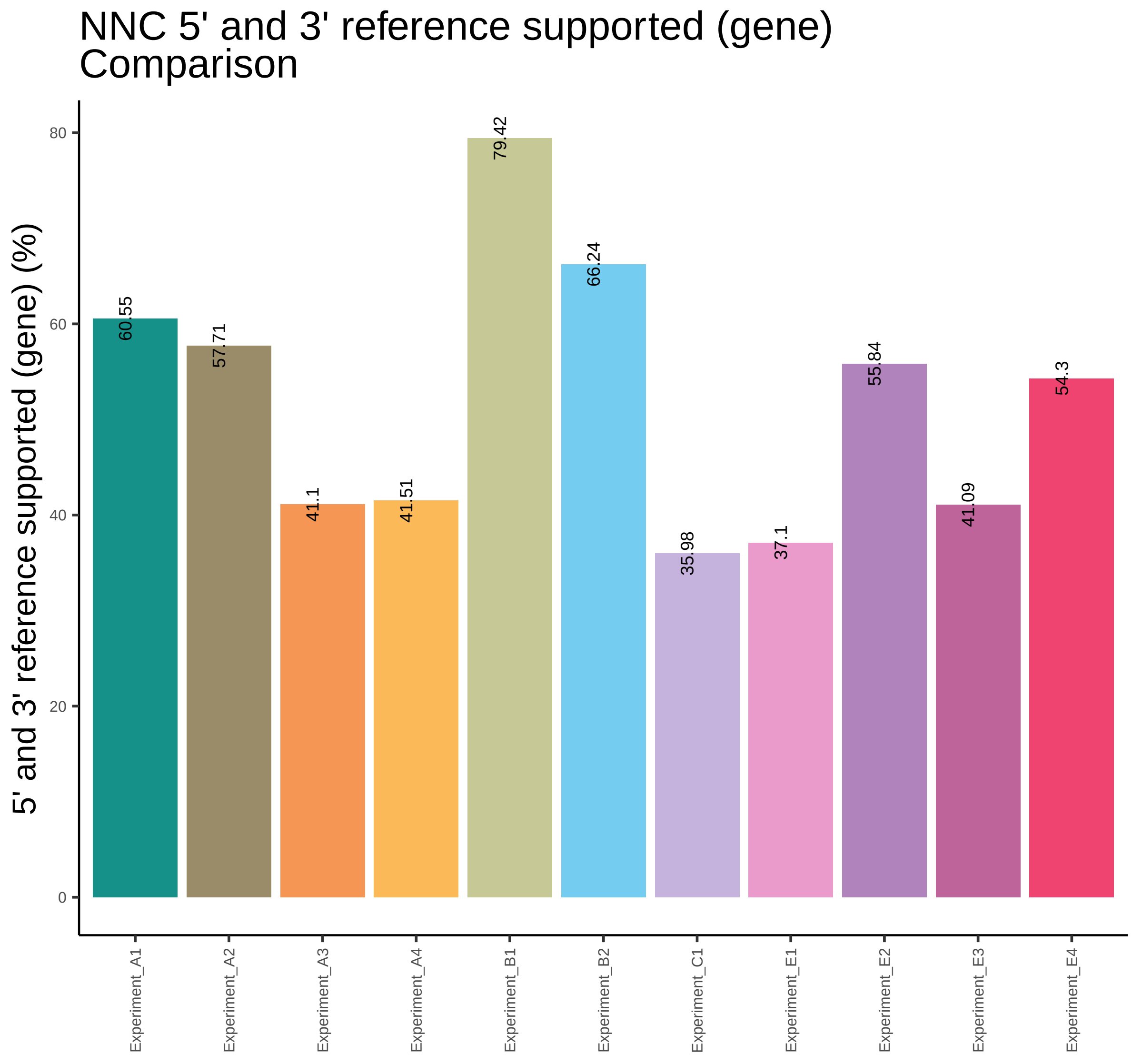

NNC 5' and 3' reference supported (gene)
Comparison
5' and 3' reference supported (gene) (%)
0
20
40
60
80
60.55
57.71
41.1
41.51
79.42
66.24
35.98
37.1
55.84
41.09
54.3
Experiment_A1
Experiment_A2
Experiment_A3
Experiment_A4
Experiment_B1
Experiment_B2
Experiment_C1
Experiment_E1
Experiment_E2
Experiment_E3
Experiment_E4

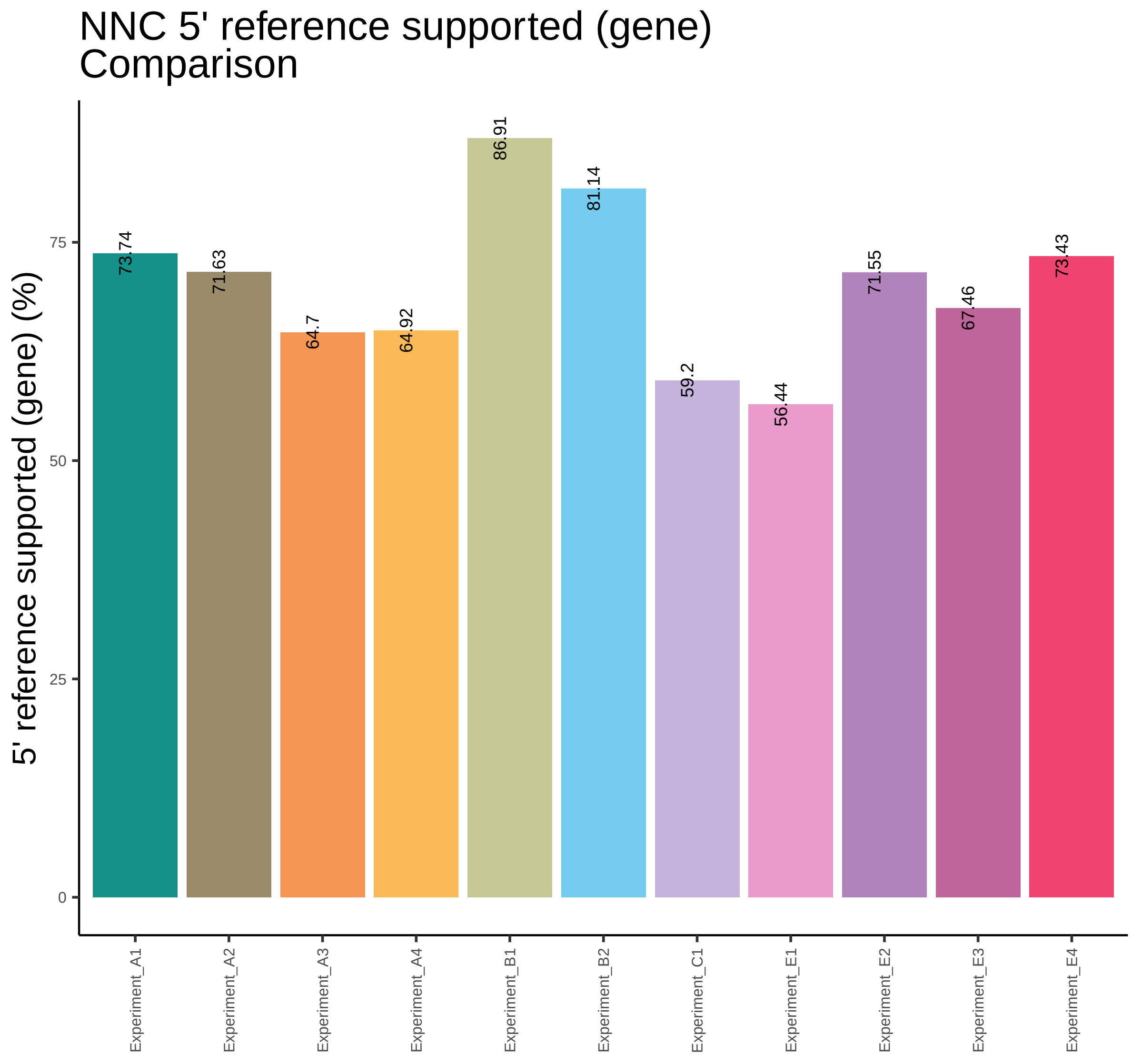
NNC 5' reference supported (gene)
Comparison
5' reference supported (gene) (%)
0
25
50
75
73.74
71.63
64.7
64.92
86.91
81.14
59.2
56.44
71.55
67.46
73.43
Experiment_A1
Experiment_A2
Experiment_A3
Experiment_A4
Experiment_B1
Experiment_B2
Experiment_C1
Experiment_E1
Experiment_E2
Experiment_E3
Experiment_E4

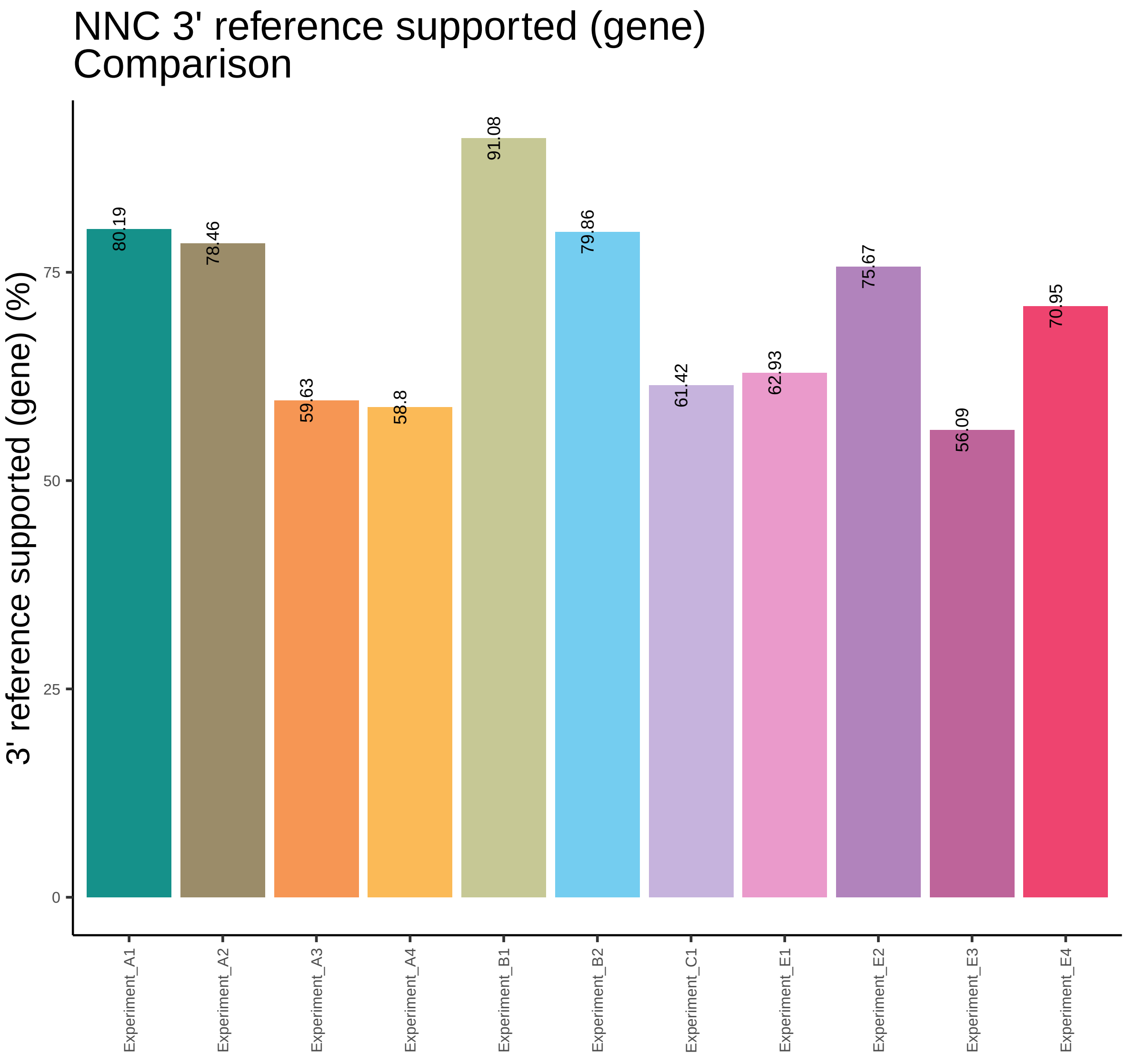
NNC 3' reference supported (gene)
Comparison
3' reference supported (gene) (%)
0
25
50
75
80.19
78.46
59.63
58.8
91.08
79.86
61.42
62.93
75.67
56.09
70.95
Experiment_A1
Experiment_A2
Experiment_A3
Experiment_A4
Experiment_B1
Experiment_B2
Experiment_C1
Experiment_E1
Experiment_E2
Experiment_E3
Experiment_E4

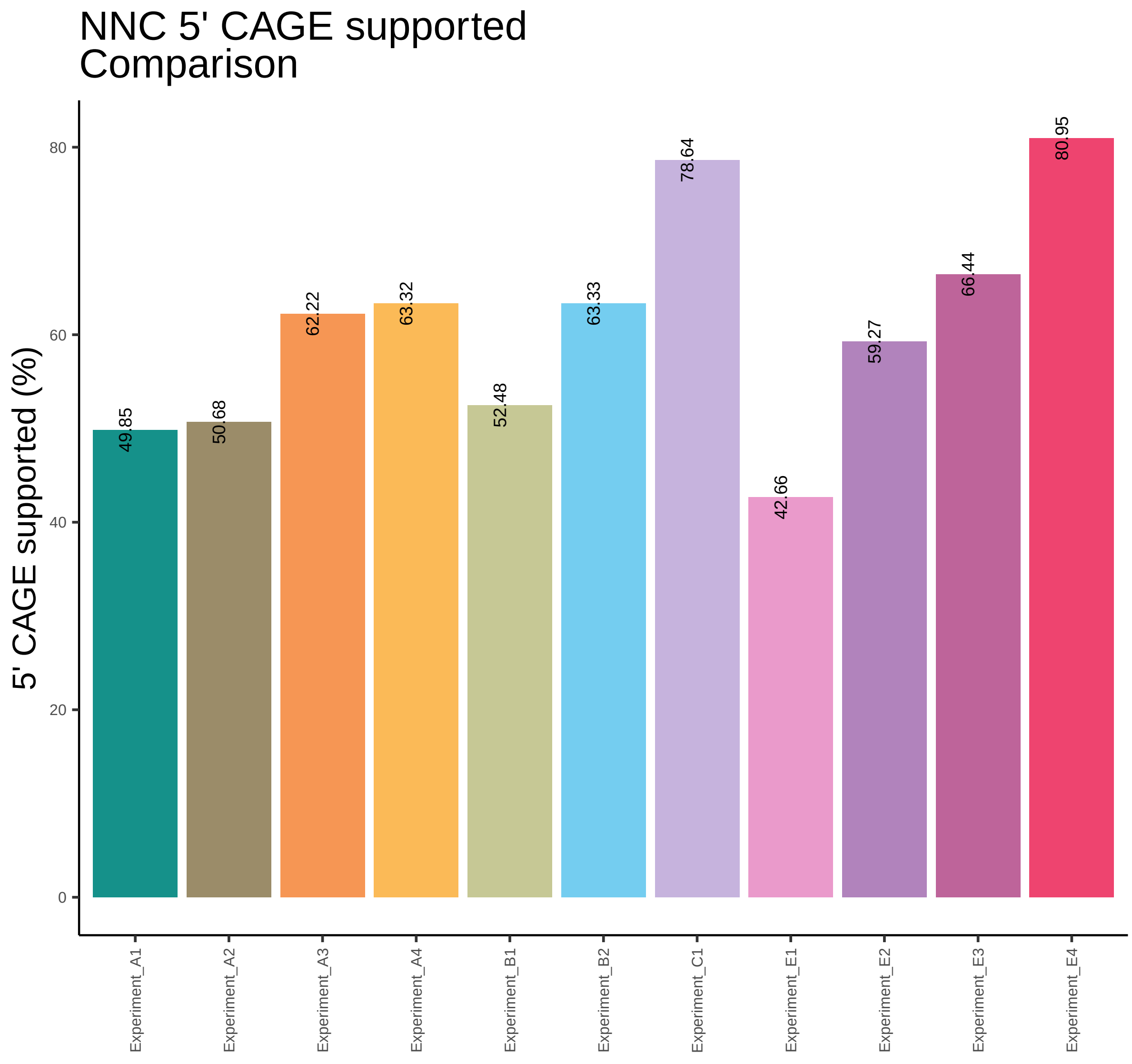
NNC 5' CAGE supported
Comparison
5' CAGE supported (%)
0
20
40
60
80
49.85
50.68
62.22
63.32
52.48
63.33
78.64
42.66
59.27
66.44
80.95
Experiment_A1
Experiment_A2
Experiment_A3
Experiment_A4
Experiment_B1
Experiment_B2
Experiment_C1
Experiment_E1
Experiment_E2
Experiment_E3
Experiment_E4

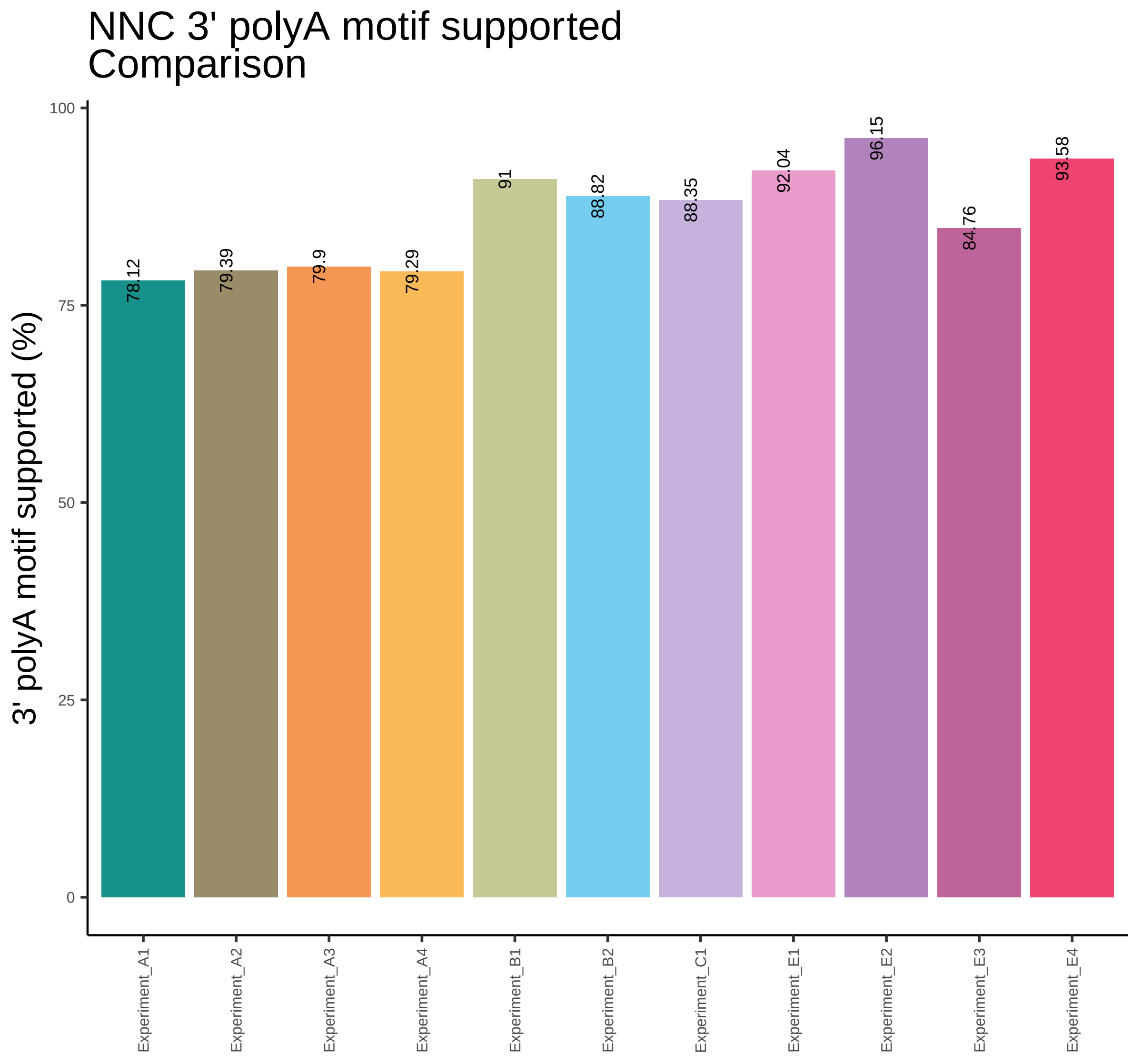
NNC 3' polyA motif supported
Comparison
3' polyA motif supported (%)
100
75
50
25
0
78.12
79.39
79.9
79.29
91
88.82
88.35
92.04
96.15
84.76
93.58
Experiment_A1
Experiment_A2
Experiment_A3
Experiment_A4
Experiment_B1
Experiment_B2
Experiment_C1
Experiment_E1
Experiment_E2
Experiment_E3
Experiment_E4

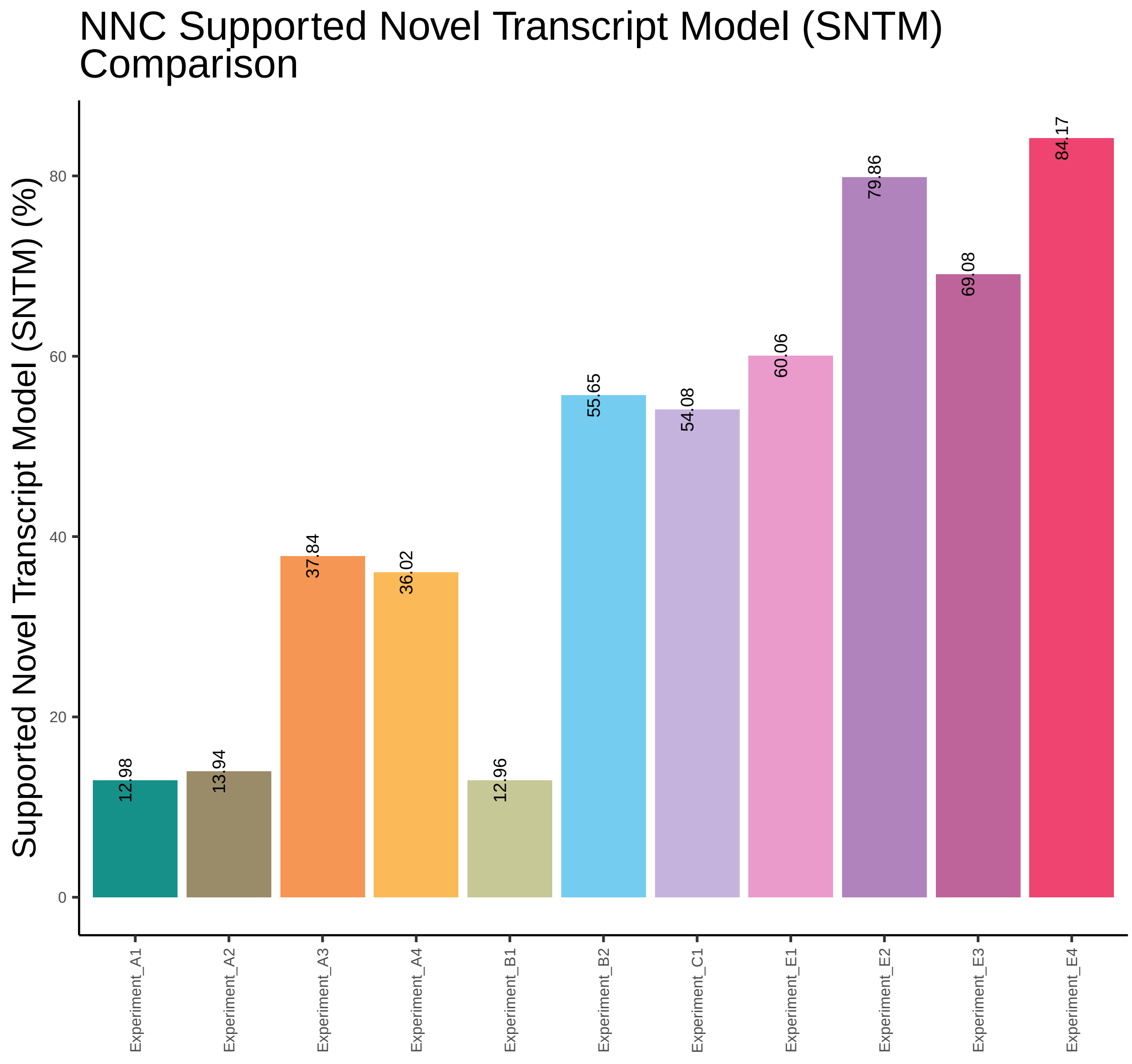
NNC Supported Novel Transcript Model (SNTM) Comparison
Supported Novel Transcript Model (SNTM) (%)
0
20
40
60
80
12.98
13.94
37.84
36.02
12.96
55.65
54.08
60.06
79.86
69.08
84.17
Experiment_A1
Experiment_A2
Experiment_A3
Experiment_A4
Experiment_B1
Experiment_B2
Experiment_C1
Experiment_E1
Experiment_E2
Experiment_E3
Experiment_E4

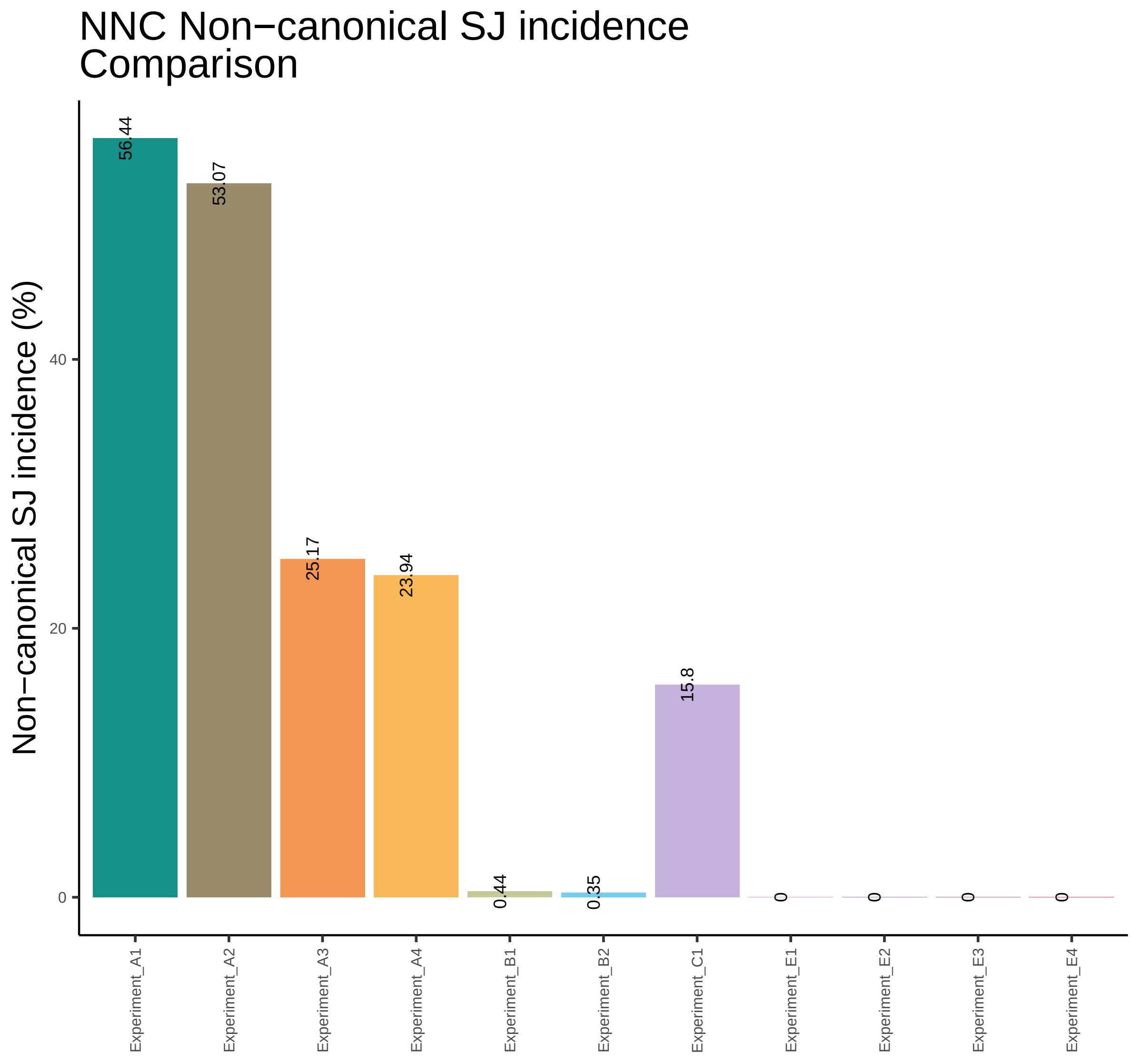
NNC Non−canonical SJ incidence
Comparison
Non−canonical SJ incidence (%)
0
20
40
56.44
53.07
25.17
23.94
0.44
0.35
15.8
0
0
0
0
Experiment_A1
Experiment_A2
Experiment_A3
Experiment_A4
Experiment_B1
Experiment_B2
Experiment_C1
Experiment_E1
Experiment_E2
Experiment_E3
Experiment_E4

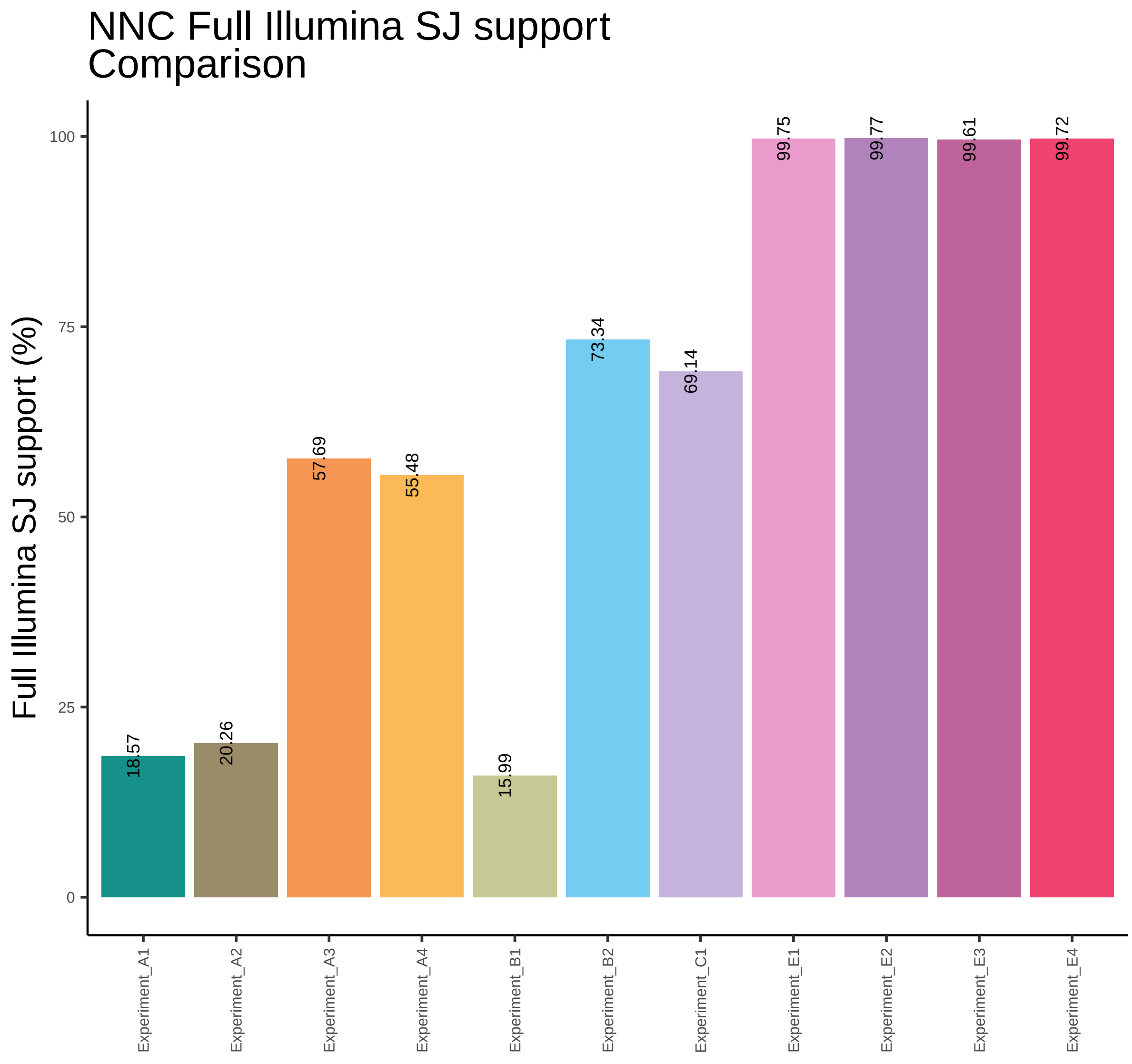
NNC Full Illumina SJ support Comparison
Full Illumina SJ support (%)
0
25
50
75
100
18.57
20.26
57.69
55.48
15.99
73.34
69.14
99.75
99.77
99.61
99.72
Experiment_A1
Experiment_A2
Experiment_A3
Experiment_A4
Experiment_B1
Experiment_B2
Experiment_C1
Experiment_E1
Experiment_E2
Experiment_E3
Experiment_E4

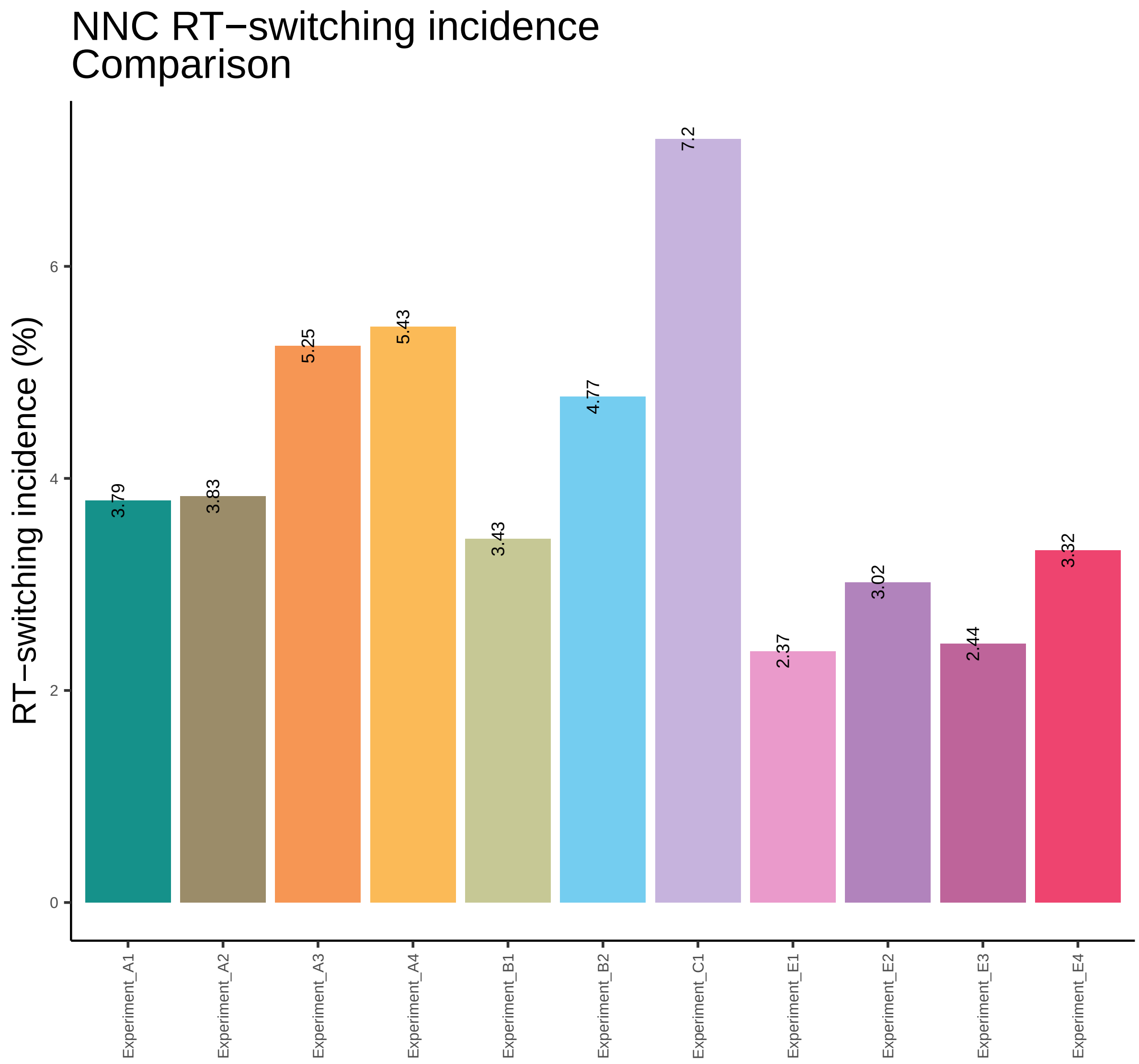
NNC RT−switching incidence
Comparison
RT−switching incidence (%)
0
2
4
6
3.79
3.83
5.25
5.43
3.43
4.77
7.2
2.37
3.02
2.44
3.32
Experiment_A1
Experiment_A2
Experiment_A3
Experiment_A4
Experiment_B1
Experiment_B2
Experiment_C1
Experiment_E1
Experiment_E2
Experiment_E3
Experiment_E4
